# Cistrome and transcriptome analysis identifies unique androgen receptor (AR) and AR- V7 splice variant chromatin binding and transcriptional activities

**DOI:** 10.1101/2021.12.06.471341

**Authors:** Paul Basil, Matthew J. Robertson, William E. Bingman, Amit K. Dash, William C. Krause, Ayesha A. Shafi, Badrajee Piyarathna, Cristian Coarfa, Nancy L. Weigel

## Abstract

The constitutively active androgen receptor (AR) splice variant, AR-V7, plays an important role in resistance to androgen deprivation therapy in castration resistant prostate cancer (CRPC). Studies seeking to determine whether AR-V7 is a partial mimic of the AR, or also has unique activities, and whether the AR-V7 cistrome contains unique binding sites have yielded conflicting results. One limitation in many studies has been the low level of AR variant compared to AR. Here, LNCaP and VCaP cell lines in which AR-V7 expression can be induced to match the level of AR, were used to compare the activities of AR and AR-V7. The two AR isoforms shared many targets, but overall had distinct transcriptomes. Optimal induction of novel targets sometimes required more receptor isoform than classical targets such as PSA. The isoforms displayed remarkably different cistromes with numerous differential binding sites. Some of the unique AR-V7 sites were located proximal to the transcription start sites (TSS). A *de novo* binding motif similar to a half ARE was identified in many AR-V7 preferential sites and, in contrast to conventional half ARE sites that bind AR-V7, FOXA1 was not enriched at these sites. This supports the concept that the AR isoforms have unique actions with the potential to serve as biomarkers or novel therapeutic targets.

## Introduction

Primary prostate cancer is androgen receptor (AR) signalling dependent; metastatic disease is treated with some form of androgen deprivation therapy (ADT). Initially responsive, some tumors develop resistance and are termed castration resistant prostate cancer (CRPC) ^1, 2^. Most CRPC remain AR-dependent. Alterations contributing to AR reactivation include elevated AR expression ^3^, local steroid synthesis ^4^, mutation of the ligand binding domain (LBD) ^5^, altered cell signalling and altered AR splicing. Alternative splicing causes expression of constitutively active AR splice variants that retain the amino-terminal transactivation domain and the DNA binding domain (DBD) but lack the LBD ^6–9^. The variants are particularly challenging to target therapeutically, since they lack the LBD, which is the primary target for most therapeutic efforts. The best characterized and most widely expressed variant is AR-V7. It contains exons 1, 2, 3 of AR and 16 unique amino acids ^7^ (Fig. 1a). Its expression has been correlated with resistance to second generation ADT in some, but not all studies ^10–13^. Studies seeking to evaluate the role of AR-V7 in prostate cancer, whether it can function independent of full length AR (AR), and whether it only regulates a sub-set of AR targets or also has unique targets have yielded conflicting results ^11, 14–16^. A major limitation in evaluating the role of AR-V7 is the lack of appropriate cell models. Many studies have relied on two cell lines, LNCaP95 (LN95) and 22RV1 ^14, 16–19^, which endogenously express AR-V7, but are both derivatives of tumors that did not originally express AR-V7. LN95 is a derivative of LNCaP cells ^20^ and 22RV1 cells were derived from an androgen dependent patient derived xenograft (PDX) that developed resistance subsequent to castration of the mouse ^21^. LN95 cells express quite low levels of AR-V7 ^22^ and 22RV1 cells express multiple variants ^23^. The AR-V7 expression levels in cell lines do not necessarily reflect AR-V7 levels in tumors. In the initial identification of AR-V7, at least some tumors were shown to express AR-V7 protein at levels comparable to that of AR in hormone dependent tumors ^7^. Consistent with this, a subsequent study of AR isoform mRNA levels in circulating tumor cells (CTCs) showed that AR-V7 mRNA levels were equivalent to or higher than the AR mRNA levels in many AR-V7 negative CTCs ^10^. Lines engineered to express variants have been generated by others. For example, Roggero et al. used engineered LNCaP cells expressing AR-V7 upon treatment with cumate, but the levels of AR-V7 were much lower than the levels of AR ^24^. Here, to address the capacity of AR-V7 to regulate transcription and chromatin binding relative to AR, LNCaP and VCaP cell derivatives, in which the levels of AR- V7 are controlled by treatment with doxycycline (Dox) ^25^, were used to equalize expression with levels equivalent to LNCaP AR.

**Figure 1.**
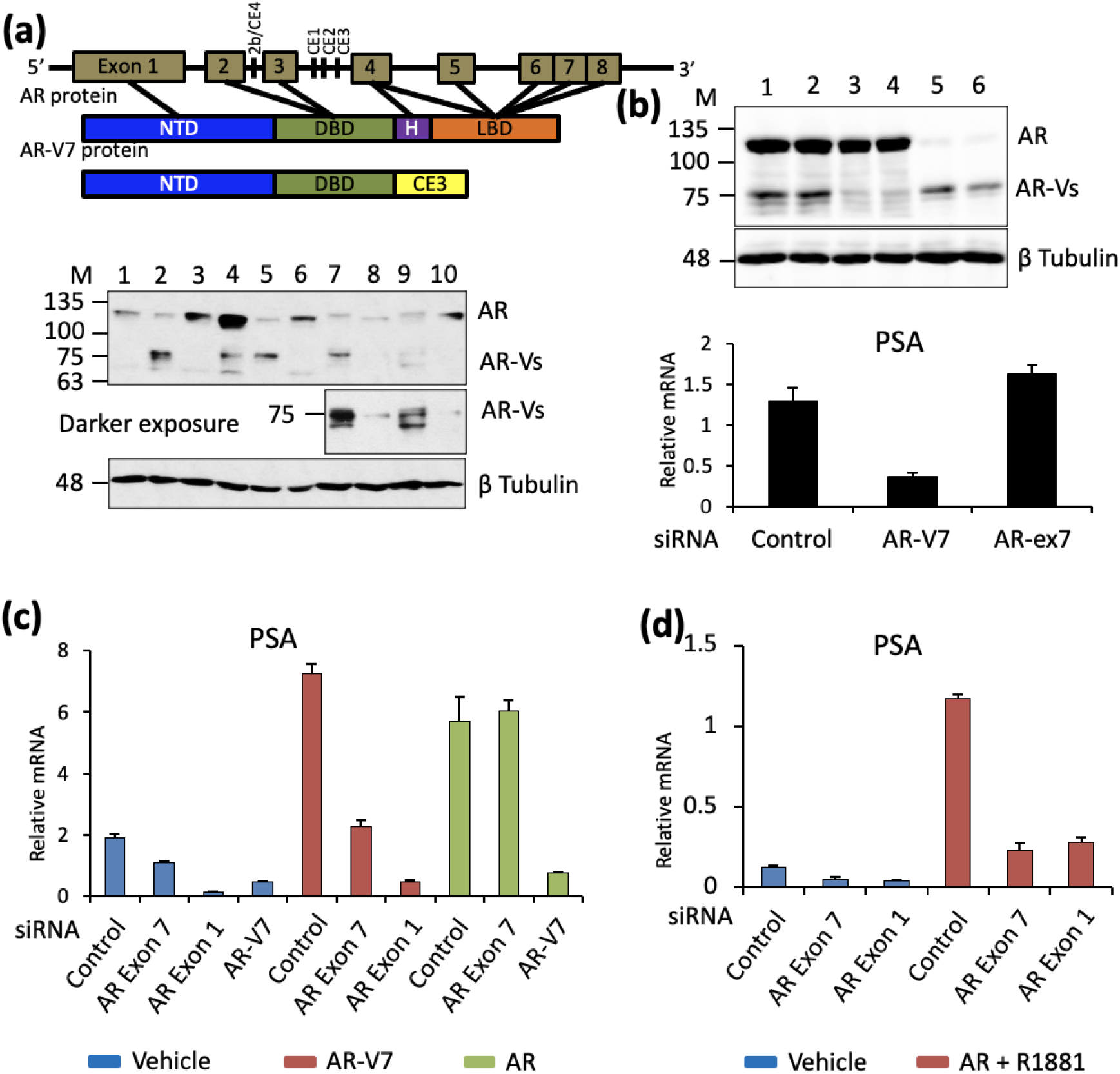
AR-V7 mediated induction of PSA is independent of AR. **a.** AR exon structure and the corresponding protein domain structures of AR and AR-V7 are shown. Western blot of 20 µg protein from cell lysates of 1. LNCaP cells treated with vehicle (0.1% ethanol), 2. LNCaP- AR-V7 treated with 20 ng/mL Dox, 3. LNCaP cells treated with 10 nM R1881 (a synthetic androgen), 4. VCaP cells treated with vehicle, 5. VCaP-V7 treated with 100 ng/mL Dox, 6. VCaP treated with 10 nM R1881, 7. 22RV1 cells treated with vehicle, 8. LN95 cells treated with vehicle, 9. 22RV1 cells treated with 10 nM R1881, 10. LN95 cells treated with 10 nM R1881. AR isoforms were detected with AR441 mAb, which recognizes AR isoforms that contain exon 1. **b**. Western blot confirmation of siRNA knock-down in LN95 cells treated with control siRNA (lanes 1,2), siRNA targeting AR-V7 (lanes 3,4) and exon-7 siRNA targeting AR (lanes 5,6) for 48 hours in charcoal stripped serum (CSS). RNA was isolated from parallel samples and PSA measured by qPCR and normalized to 18S RNA. **c.** LNCaP AR-V7 cells in CSS were treated with the indicated siRNAs for 48 hours followed by treatment with vehicle, R1881, or Dox overnight. RNA was isolated, PSA expression measured by qPCR and normalized to 18S RNA. **d**. LNCaP cells in CSS were treated with the indicated siRNAs for 48 hours followed by overnight treatment with vehicle or R1881. RNA was isolated and PSA measured as above. All qPCR data are plotted as a mean of three biological replicates from the same experiment and error bars are SEM. Each experiment was performed a minimum of three times and a representative experiment is shown.

## Results

### Expression of AR isoforms in cell models

The exon structure and protein domains of AR and AR-V7 are depicted in Fig. 1a. The western blot in Fig. 1a compares expression of AR and variant proteins in LNCaP, VCaP, 22RV1, LN95 cells and the LNCaP ^pHAGE AR-V7^ (LNCaP AR-V7) and VCaP ^pHAGE AR-V7^ (VCaP AR-V7) cells treated with Dox at levels used in subsequent studies. In the absence of hormone, the VCaP cells express some AR-V7 (Fig. 1a lane 4, but Dox treatment of VCaP AR-V7 cells increases levels in a dose dependent manner ^25^. In contrast, the parental LNCaP cells do not express AR- V7 (Fig. 1a lane 1). However, the LNCaP AR-V7 cells express low levels of AR-V7 in the absence of Dox (SFig. 1). Whereas the Dox treated cells express levels of AR-V7 comparable to R1881 (a synthetic androgen) bound AR in the parental lines, the level of AR variants in LN95 cells is much lower (compare Fig. 1a lanes 2 [LNCaP AR-V7], 5 [VCaP AR-V7] with lane 8 [LN95]). Although 22RV1 has substantial expression of variants, they are heterogeneous (Fig. 1a lanes 7, 9). LN95 cells primarily express the AR-V7 variant; treatment with an siRNA targeting CE3 greatly reduces variant expression (Fig. 1b). Although this siRNA would also deplete AR-V9, there is negligible AR-V9 expression in LN-95 cells (SFig. 2). In contrast, 22RV1 cells express substantial levels of AR-V9 (SFig. 2). Two separate siRNAs that target the 3’ untranslated region of AR-V7 as well as some additional variants, efficiently deplete AR- V7 in 22RV1 cells, but substantial amounts of other variants remain (SFig. 3). That these are other variants rather than AR proteolytic fragments is confirmed by the findings that an siRNA targeting exon 1, which is common to all AR isoforms and eliminates all isoforms (SFig. 3. lane 5), while an siRNA targeting exon 7 depletes AR, but not the variants (SFig. 3 lane 4).

### AR is not required for AR-V7 dependent induction of PSA

Since cell lines and tumours that express AR-V7 typically also express AR, the question of whether AR-V7 requires AR to function has been raised. In LN95 cells, depletion of AR-V7 reduces PSA expression, but depletion of AR does not (Fig. 1b). In the LNCaP AR-V7 model, which expresses low levels of AR-V7 in the absence of Dox treatment (SFig. 1), depletion of either AR (Exon 7) or AR-V7 in the absence of Dox induction reduces basal PSA expression (Fig. 1c). As expected, depleting AR using siRNA targeted to exon 7 reduces R1881 induced PSA expression; siRNA targeting exon 1, which depletes all isoforms, is even more effective. In contrast AR depletion has no effect on Dox induced PSA expression (Fig. 1c). Surprisingly, depletion of AR in parental LNCaP cells (Fig. 1d) using either an siRNA targeting exon 7 (unique to AR) or exon 1 (common to all isoforms) also reduced basal PSA expression suggesting that this parental line exhibits a low level of aberrant activation of AR independent of variants.

### AR isoform specific gene expression

RNA-seq data from LNCaP AR-V7 cells (LNCaP ^AR-V7/pLenti^) ^26^ and VCaP AR-V7 (VCaP ^pHAGE AR-V7^) cells in charcoal stripped serum (CSS) treated with vehicle, R1881 or Dox for 24 hours prior to harvesting were used for transcriptomic analyses. The reproducibility of the biological replicates prepared in separate experiments is shown in the principal component analysis (PCA) SFig. 4. The scatter plots of fold change in Fig. 2a (LNCaP model) and 2B (VCaP model) show that although there are many genes regulated by both isoforms (red), there also are unique targets (blue for AR and green for AR-V7) as well as some genes regulated in the opposite direction (black). The list of differentially expressed genes (DEGs) can be found in STable 3. Fig. 2c shows Venn diagrams depicting the numbers of common and uniquely up and down regulated genes in the two inducible models. In the LNCaP model, roughly 46% of the AR-regulated genes and 47% of the AR-V7 regulated genes are regulated by both isoforms, with 54% of AR-target genes and 53% of the AR-V7 target genes unique to each of the isoforms. Although the number of up and down-regulated genes are similar, there is a slight preference for upregulation by AR-V7. In the VCaP model, many more genes are regulated by AR than by AR-V7 (Fig. 2c), but about 34% of the AR-regulated genes and 70% of the AR-V7 regulated genes are regulated by both isoforms, and again, there are also unique AR-V7 regulated genes. In this line, too, AR-V7 shows a slight preference for induction of genes. The extent of overlap in DEGs between the two models is shown in SFig. 5. When total gene expression is compared, overlap ranges between 20-50% depending upon the specific comparison. When only isoform specific DEGs are compared, the total number of genes is reduced, and overlap diminishes to a range of 7-33%. Some of the variation is due to the differences in total number of genes regulated. For example, 7% (40) of AR-V7 specific down-regulated genes in LNCaP cells are also down-regulated in the VCaP model, but this set represents 21% of the genes specifically down-regulated by AR-V7 in the VCaP model.

**Figure 2.**
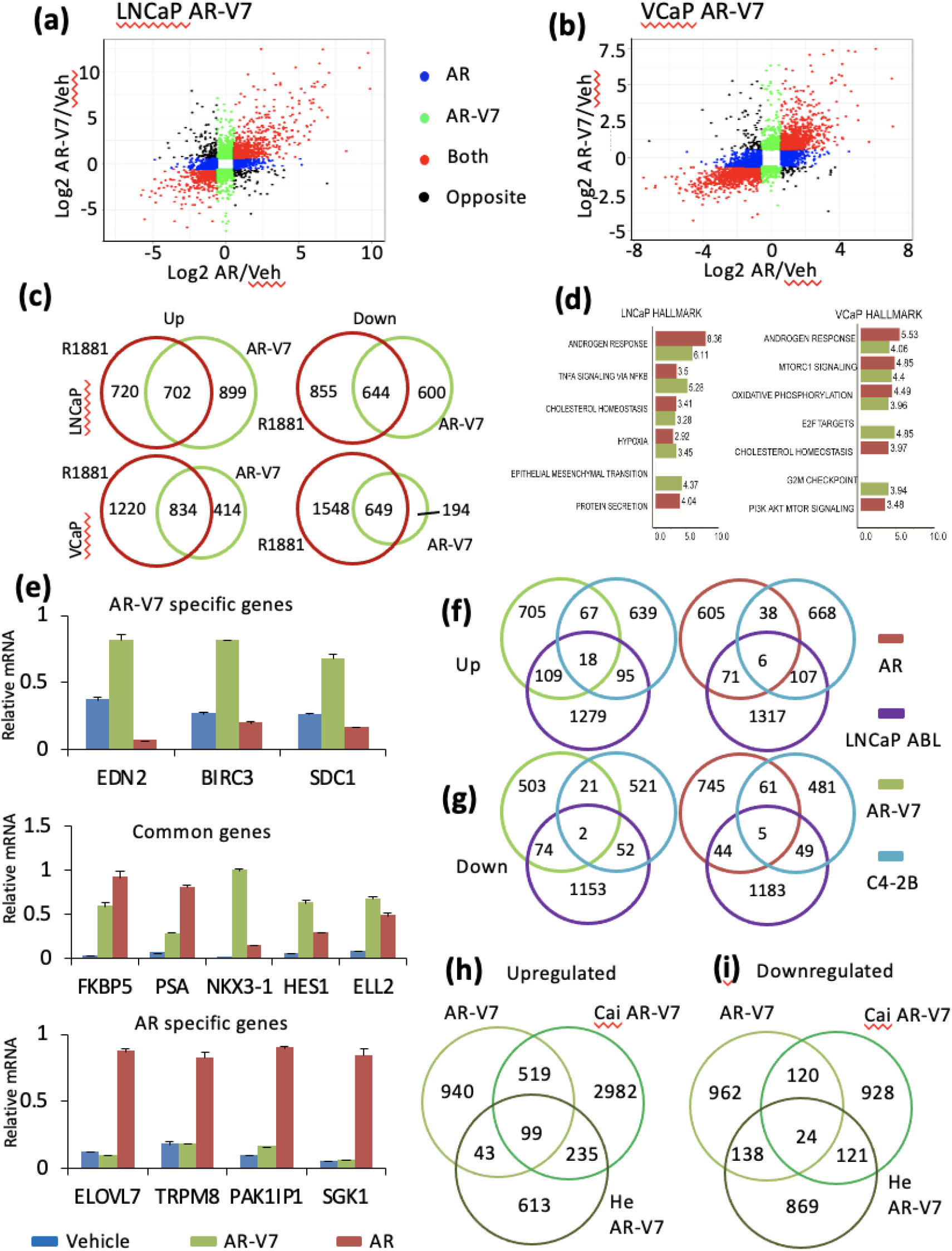
Differential Gene Regulation by AR isoforms: **a - b.** Scatter plot illustrating genes regulated uniquely by AR or AR-V7, regulated by both or regulated in the opposite direction (eg: AR induced and AR-V7 repressed); colored by expression pattern. Data are plotted as log2 fold change of gene expression in R1881 vs vehicle treated (x-axis) and Dox vs vehicle (0.1% ethanol) treated (y-axis) LNCaP-AR-V7 cells (**a**) and VCaP AR-V7 cells (**b)**. **c.** Venn diagram showing number and overlap of AR and AR-V7 regulated genes in LNCaP and VCaP models. FC>1.5, adjusted p<.05. Venn diagrams are plotted as circles proportional to the numbers. **d.** Comparison of top Hallmark Pathways upregulated by AR (red) and AR-V7 (green) in LNCaP (left) and VCaP (right) cell models. **e.** Gene expression profiles of genes specifically regulated by AR-V7, by both isoforms or by AR. **f.** Venn diagram comparing the number of unique AR- V7 induced genes (left) or unique AR induced genes (right) (FC > 1.5, adjusted *p* < .05) in LNCaP-V7 cells that overlap with previously reported AR upregulated genes in C4-2B (FC > 1.25, adjusted *p* < .05) ^27^ and LNCaP ABL models (FC > 1.5, adjusted *p* < .05) ^28^ identified by depleting AR in cells in CSS (plots not proportional). **g.** Venn diagram showing the number of unique AR-V7 (left) or AR (right) repressed genes (FC > 1.5, adjusted *p* < .05) in LNCaP-V7 cells overlapping AR regulated genes in C4-2B (FC > 1.25, adjusted *p* < .05) and LNCaP ABL models (FC > 1.5, adjusted *p* < .05) as above. **h.** Venn diagram showing number of unique AR- V7 up-regulated genes in LNCaP AR-V7 cells overlapping previously reported AR-V7 signatures in LN95 and 22RV1 cells identified either by depleting AR-V7 (Cai et al. ^19^) or all variants (He et al. ^17^). Unique AR-V7 genes from LNCaP AR-V7 (FC > 1.5, adjusted p < .05) were compared with AR-V7 signature from 22RV1 and LN95 (FC > 1.25, adjusted p < .05). **i.** Venn diagrams showing the down-regulated genes corresponding to the analysis in **h**. All qPCR plots are mean of three biological replicates from the same experiment and error bars are SEM. Venn diagrams **f – i** are plotting symmetrical circles and not proportionate to numbers.

Consistent with a significant overlap, GSEA analysis of Hallmark pathways shows that the androgen response is the most significant induced pathway for both isoforms in each line and cholesterol homeostasis is also included in the top five for both lines (Fig. 2d). However, there are substantial isoform and, particularly cell line specific differences in other pathways (Fig. 2d, SFig. 6, 7). Confirmation of select gene specificities is shown in Fig. 2e; qPCR analysis of the LNCaP AR-V7 model revealed genes preferentially regulated by AR-V7 (top), regulated by both although to different extents (middle) and regulated by AR (bottom).

Since AR dependent gene expression in CRPC cell lines differs substantially from hormone dependent activation in androgen sensitive lines, the AR-V7 (Fig. 2f) and AR unique genes (Fig. 2g) from the LNCaP model were compared with the sets of all AR dependent genes previously identified in C4-2B ^27^ and LNCaP ABL ^28^ CRPC cells that lack AR-V7 expression (STable 1). There was minimal overlap of any of the signatures (14% of AR-V7 upregulated genes overlapped with C4-2B and 9.5% with LNCaP ABL, SFig. 8) suggesting that the AR-V7 specific genes are not simply the subset of genes regulated when AR is activated in the absence of added hormone. The overlap between C4-2B and LNCaP ABL is of a similar magnitude. When all AR or AR-V7 genes in either the LNCaP or VCaP model are compared with regulated genes in C4-2B and LNCaP ABL, the total numbers of genes were increased, but the percentage overlap was similar (SFig. 8).

22RV1 cells express multiple variants and the total variant level exceeds that of AR (SFig. 3). When the LNCaP AR-V7 specific genes are compared with two 22RV1 data sets derived by depleting all variants ^17^ or AR-V7 ^19^ there is overlap between the LNCaP AR-V7 data and the 22RV1 data (Fig. 2h and i), but in each case, the majority of the genes are unique to the specific data set. Comparison with the VCaP AR-V7 genes using one of these data sets and a third 22RV1 data set ^29^ also yielded only modest overlaps (SFig. 9). This highlights the challenge of studying a mixed population of low levels of variants in the 22RV1 line. To directly compare with a model that endogenously expresses AR and AR-V7 only, we used the LN95 model. LN95 cells in CSS were depleted of AR (exon 7) or AR-V7 (CE3 siRNA) or treated with control siRNA as described for Fig. 1b; expression was analyzed by RNA-seq. LN95 cells also were treated with vehicle or 10 nM R1881, RNA isolated and expression analyzed by RNA-seq. Whereas about a thousand genes were regulated by AR-V7 depletion and about 1400 by R1881, only about 200 were regulated by depleting AR (SFig. 10). Of the 200, 100 were unique to AR depletion. There was very little overlap between AR depletion and either AR-V7 regulated genes or R1881 regulated genes. This suggests that the unliganded AR not only is relatively inactive in LN95 in CSS, but that it also is not required for regulation of most AR-V7 regulated genes. A comparison of gene expression in the LN95 model with the LNCaP AR-V7 model (SFig. 11) shows that 70% of the genes induced by R1881 in the LN95 model are also induced in the LNCaP model. AR-V7 is only weakly expressed in LN95 cells relative to AR (Fig. 1b).

Only 20% of the genes induced by AR-V7 in LN95 cells also are induced by AR-V7 in the LNCaP model. However, of the 282 genes upregulated by AR-V7 in LNCaP and by R1881 only in LN95, 236 are also regulated by R1881 in LNCaP and there is similar overlap for the downregulated genes (210 of 268). Failure to detect regulation of these genes by AR-V7 in LN95 may be due to the relatively low level of AR-V7.

### AR-V7 preferentially binds to the chromatin at proximal promoter sites

To determine whether AR and AR-V7 also differentially bind to chromatin, AR binding sites (ARBs) in LNCaP cells treated with R1881 and AR-V7 binding sites (V7Bs) in LNCaP AR-V7 cells treated with Dox were identified by ChIP-exo. The analyses yielded 87,095 AR binding peaks and 59,852 AR-V7 binding sites with 32,082 common sites detected in both analyses. (Fig. 3a). Specific sites were defined as those binding sites not overlapping (≥1 base pair) with the other isoform. The average relative intensities at the three types of sites (common, AR preferred, AR-V7 preferred) are also shown. Publicly available ChIP-seq data from vehicle treated LNCaP cells were used for comparison ^30^. An analysis of the regions 50 kb upstream and 5 kb downstream of the transcription start sites (TSS) revealed a remarkable preferential enrichment of AR-V7 near the TSS (Fig. 3b). A comparison of binding on genes separated as having more overall AR signal or AR-V7 signal revealed preferential binding of AR-V7 at the TSS in both cases (SFig. 12). Note that only binding 50 kb upstream from the TSS and 5 kb downstream is portrayed in the figure. ARBs were found associated with 2251 AR regulated genes and V7Bs associated with 2310 AR-V7 target genes in LNCaP AR-V7 cells (Fig. 3c). AR-V7 binding was detected at the TSS (within 100bp) of 267 AR-V7 induced genes and a smaller number of repressed genes (168) whereas AR bound at the TSS of relatively few regulated genes (29 up and 28 down). A motif analysis of ARBs and V7Bs within 100 bp of the TSS of up and down DEGs identified differential cofactor consensus motifs at these sites (SFig. 13). The consensus motifs for AR-V7 regulated genes differed markedly depending upon whether the genes were induced or repressed. Induced genes were enriched for ARE/GRE consensus motifs whereas the repressed genes showed predominantly Fox family motifs. In contrast genes up or down regulated by AR showed a similar abundance of both ARE and FOX family motifs. In sum, cistromic analyses indicate AR isoform specific binding in this model is induced by multiple mechanisms.

**Figure 3.**
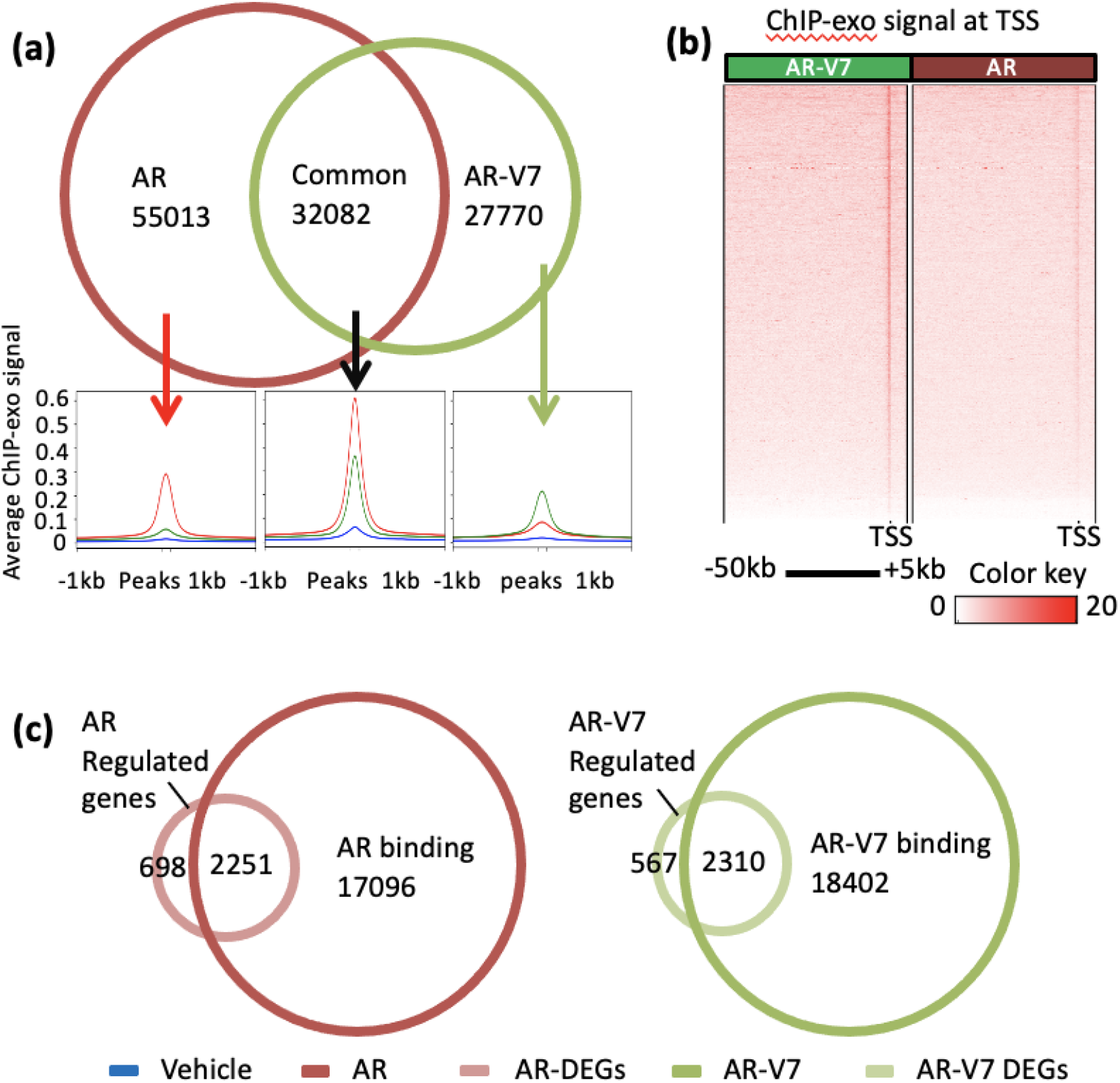
AR-V7 exhibits preferential binding to promoters in LNCaP AR-V7 cells. **a.** AR isoform specific and common binding sites identified in LNCaP cells treated with R1881 or LNCaP AR-V7 cells treated with Dox in 10% CSS medium. Average ChIP-exo signals at these corresponding subsets of binding sites (indicated by arrows) are plotted below. Average ChIP- seq intensities within 1 kb distance to the center of peaks plotted as a profile of each subset from previously reported LNCaP cells treated with vehicle (Blue) ^30^, LNCaP AR-V7 cells treated with 20 ng/mL Dox (green) and LNCaP cells treated with 10 nM R1881 (red) for 24 hours. **b.** AR and AR-V7 ChIP-exo signal within 50 kb upstream and 5 kb downstream of all the TSS in human genome build hg19 plotted as average signal for each 10 bp bin. TSS with only scores of zero were not included in the plot. **c.** Integrated RNA-seq and ChIP-exo analyses reveal the AR and AR-V7 induced and repressed direct targets. Venn diagram showing number of genes with AR or AR-V7 binding sites (annotated to the nearest TSS using HOMER/BEDTools, on the same DNA strand) overlapping differentially expressed genes (FC > 1.5, adjusted *p* < .05). All Venn diagrams are plotted as circles proportional to the numbers.

### AR-V7 recruits little to no AR to chromatin

The ChIP-exo analyses were performed with an antibody that recognizes both isoforms and thus the signal in response to Dox represents both AR-V7 and any AR that may be recruited, whereas the R1881 treatment was performed in parental LNCaP cells and the signal is a result only of AR binding. To more directly compare binding at select targets, ChIP qPCR was performed, and results compared to the ChIP-exo data. Fig. 4a, b confirm that AR binds more strongly to selected sites in TRPM8 and NKX3-1, and AR-V7 binds more strongly to a site in BIRC3 (Fig. 4c) when an antibody that recognizes both isoforms is used. To look at AR binding only, ChIP qPCR analysis was done with the C-19 antibody, which recognizes the C-terminus of AR and thus only full-length AR. R1881 dependent recruitment is detected at the same sites as when the Active Motif antibody is used (panels 4d-h). When the signal in response to AR-V7 induction (Dox) is compared, there is no enrichment of C-19 signal at the PSA enhancer (Fig. 4d), PSA promoter (Fig. 4f) or BIRC3 (Fig. 4h). However, for two genes where fold recruitment is very high, a small increase in C-19 signal is detected. For sites in FKBP5 (Fig. 4e) and EDN2 (Fig. 4g), fold recruitment relative to corresponding vehicle is 10% of the N-terminal Ab fold recruitment suggesting minimal recruitment of AR by AR-V7.

**Figure 4.**
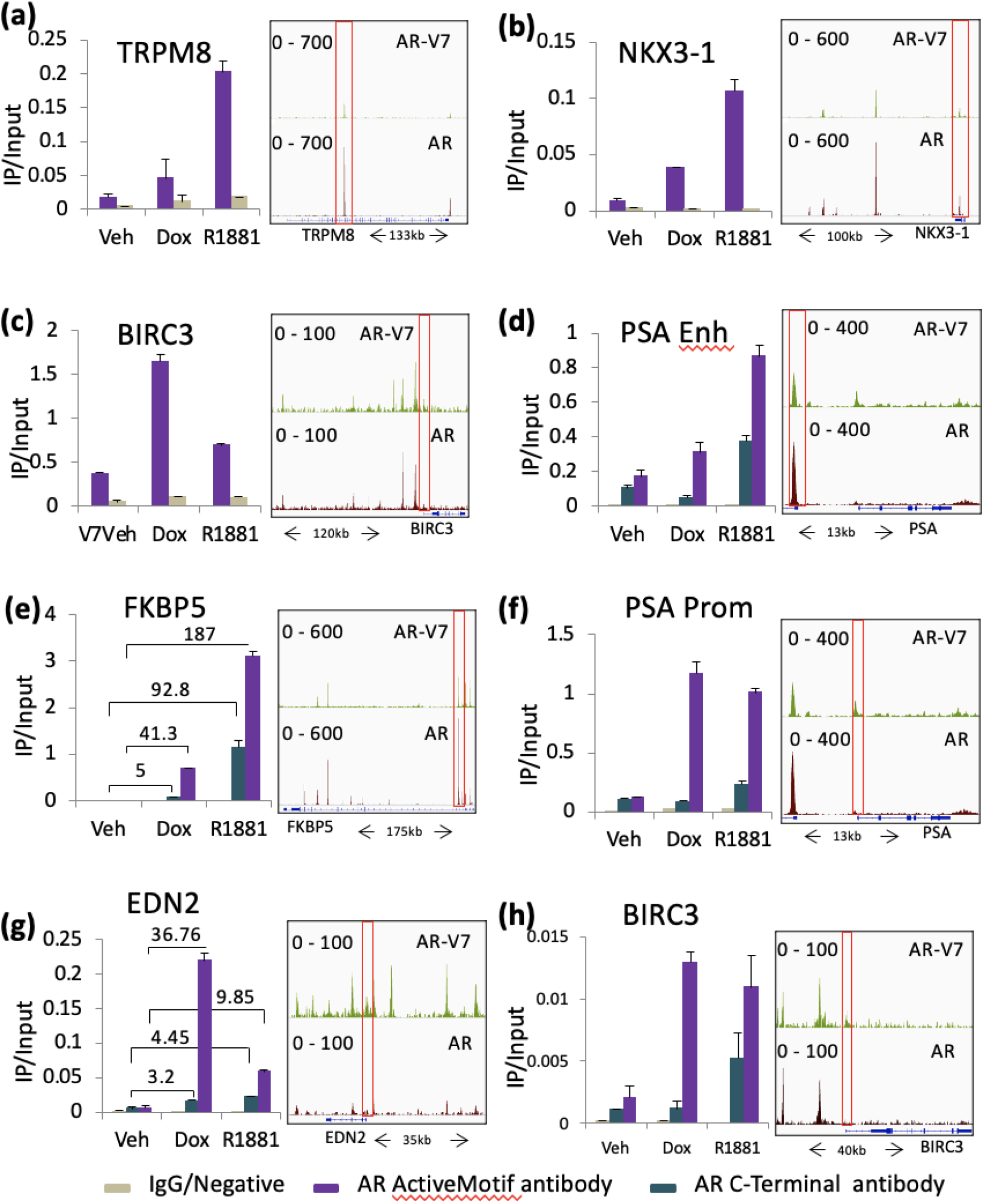
AR-V7 recruits only a small amount of AR to chromatin binding sites. **a-c.** ChIP- qPCR confirmation of AR and AR-V7 binding at **a**. gene induced by AR (TRPM8), **b**. gene induced by both isoforms (NKX3-1) and **c**. gene induced by AR-V7 (BIRC3). **d-h**. A comparison of recruitment measured by ChIP-qPCR following an immunoprecipitation by the Active Motif antibody that recognizes both AR isoforms or the C-19 antibody, which recognizes only AR. **d.** ChIP-qPCR quantification of AR and AR-V7 binding at the PSA enhancer site. **e**. ChIP-qPCR quantification of AR and AR-V7 binding at the FKBP5 promoter site. **f**. ChIP- qPCR quantification of AR and AR-V7 binding at the PSA promoter site. **g**. ChIP-qPCR quantification of AR and AR-V7 binding at the EDN2 promoter site. **h**. ChIP-qPCR quantification of AR and AR-V7 binding at the BIRC3 promoter site. ChIP-exo data using Active Motif antibody in 20 ng/mL Dox treated LNCaP AR-V7 cells and 10 nM R1881 treated LNCaP cells. Genome browser tracks visualizing ChIP-exo signal at the corresponding gene are plotted on the right with the site analyzed by ChIP qPCR enclosed in a red box. All qPCR plots are the ChIP signal normalized to input and data are plotted as a mean of two biological replicates from the same experiment; error bars are SEM. Each experiment was performed a minimum of three times and a representative experiment is shown.

### AR isoform preference is retained independent of isoform expression level for a sub-set of target genes

To assess the effect of AR-V7 levels on gene expression, the LNCaP AR-V7 cells were treated with vehicle, R1881, or various concentrations of Dox. SFig. 14 shows the relative levels of AR-V7 expression. In this model, 20 ng/ml yields optimal AR-V7 dependent induction of PSA gene expression but significantly less than that induced by R1881 (Fig. 5a). Some genes that are more highly induced by AR-V7 at 20 ng/ml than by 10 nM R1881 show an additional Dox dependent increase in expression ranging from 3-8 fold (Fig. 5a). In contrast, ELOVL7 and SGK1, AR specific genes, are not induced regardless of the level of AR-V7 induction (Fig. 5a). To assess whether genes preferentially induced by AR-V7 could be induced by AR at higher levels, an LNCaP line that expresses Flag tagged AR in response to Dox was developed (see Sfig. 15 and 16 for induction levels and a demonstration that the flag AR is active). PSA induction (Fig. 5b), TMPRSS2, SGK1, and NDRG1 induction (Sfig. 16) were not further increased by excess AR. In contrast for ELL2, which is somewhat induced by AR, increasing AR expression increased ELL2 expression 2X (Fig. 5b). EDN2 is induced by AR-V7 but repressed by AR. Increased expression of AR resulted in further repression of EDN2 at low levels of hormone, but comparable levels at 10 nM R881. There was no switch to induction. However, there is a dramatic increase in the induction of OLAH when AR is overexpressed (Fig. 5b). Thus, although there are examples where expression levels determine specificity, subsets of genes are isoform specific regardless of the level of isoform.

**Figure 5.**
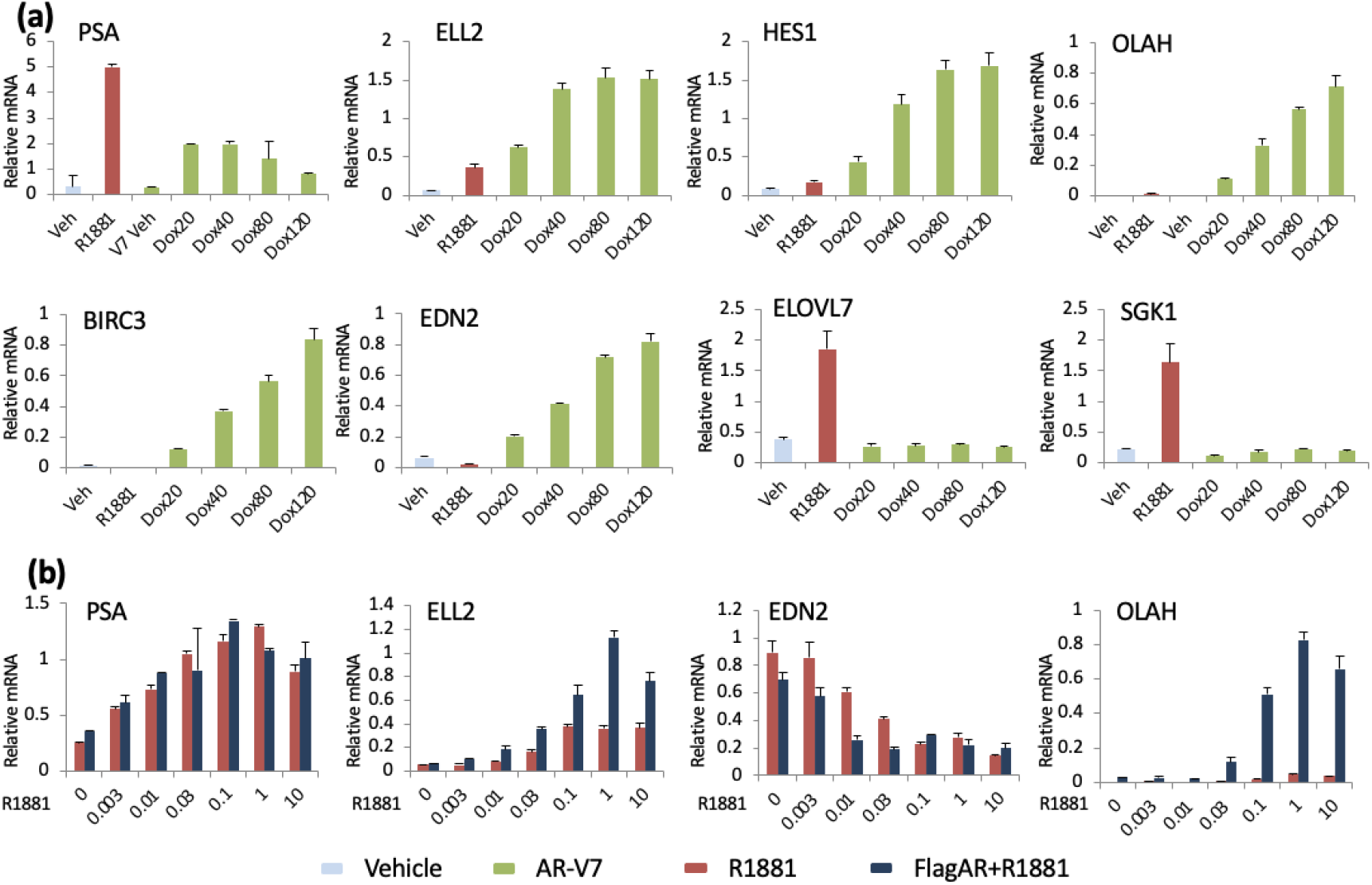
AR isoform concentration alters responsiveness for a sub-set of target genes. **a**. LNCaP AR-V7 cells in 10 % CSS were treated with vehicle (0.1% ethanol), 10 nM R1881 or varying levels of Dox (20-120 ng/mL) for 24 hours prior to isolation of RNA and measurement of target genes by qPCR. Gene expression of AR and AR-V7 induced genes (PSA, ELL2, HES1), AR specific genes (ELOVL7, SGK1) and AR-V7 specific genes (EDN2, OLAH, BIRC3) was quantified using qPCR assays. **b**. Cells were treated with vehicle or Dox (20 ng/mL) to induce FlagAR and the indicated concentration of R1881 (0-10 nM) for 24 hours, RNA isolated and gene expression of AR and AR-V7 induced genes (PSA, ELL2) and AR-V7 specific genes (EDN2, OLAH) was quantified using qPCR assays. All qPCR plots are expression values normalized to 18S on the *y*-axis and corresponding treatment on the *x*-axis. Data are plotted as the mean of three biological replicates from the same experiment; error bars are SEM. Each experiment was performed a minimum of three times and a representative experiment is shown.

### A role for FOXA1 in AR isoform specific gene expression

FOXA1 is a well-established pioneer factor for AR ^31^. An analysis of consensus motifs within 200 bp of the peak of AR isoform binding reveals that FOXA1 and related protein motifs are more abundant than consensus AREs, whereas the reverse is true for AR-V7 (Fig. 6a). Previous studies ^32^ have shown that AR dependent transcription can be roughly divided into three categories, genes that are regulated regardless of FOXA1 status, genes that require FOXA1 for AR dependent induction and those that are induced only when FOXA1 is depleted (Fig. 6b). FOXA1 can influence AR activity both through direct interaction at a composite element and indirectly by changing the overall chromatin structure. In previous studies, we had found that two genes that require FOXA1 for AR dependent induction and that contain composite elements were not induced by AR-V7, nor did AR-V7 bind to these sites in RASSF3 ^26^ or INPP4b ^33^. To test whether other genes show FOXA1 dependence, cells were depleted of FOXA1 (SFig. 17) and the capacity of AR or AR-V7 to regulate these genes was measured. Depletion of FOXA1 had no effect on AR-V7 mediated induction of FKBP5, although it slightly reduced R1881 dependent induction (Fig. 6c). There was no effect of depletion on AR-V7 dependent induction of BIRC3 nor did it cause BIRC3 to be regulated by R1881. In contrast, depletion of FOXA1 eliminated R1881 dependent induction of ELOVL7 (Fig. 6d). Surprisingly, SGK1, an AR dependent gene was further induced when FOXA1 was depleted (Fig. 6d). Although basal expression also was increased, AR-V7 expression did not further induce SGK1. That AR-V7 activity can be altered by FOXA1 status is shown in Fig. 6e. HES1 induction is reduced when FOXA1 is depleted, with no induction by R1881 in either condition. For EDN2, as reported previously, AR-V7 induces expression. However, this expression is increased when FOXA1 is depleted. The previously reported induction of EDN2 by AR when FOXA1 was depleted was confirmed, but levels of expression were very low compared to AR-V7 dependent induction (Fig. 6e).

**Figure 6.**
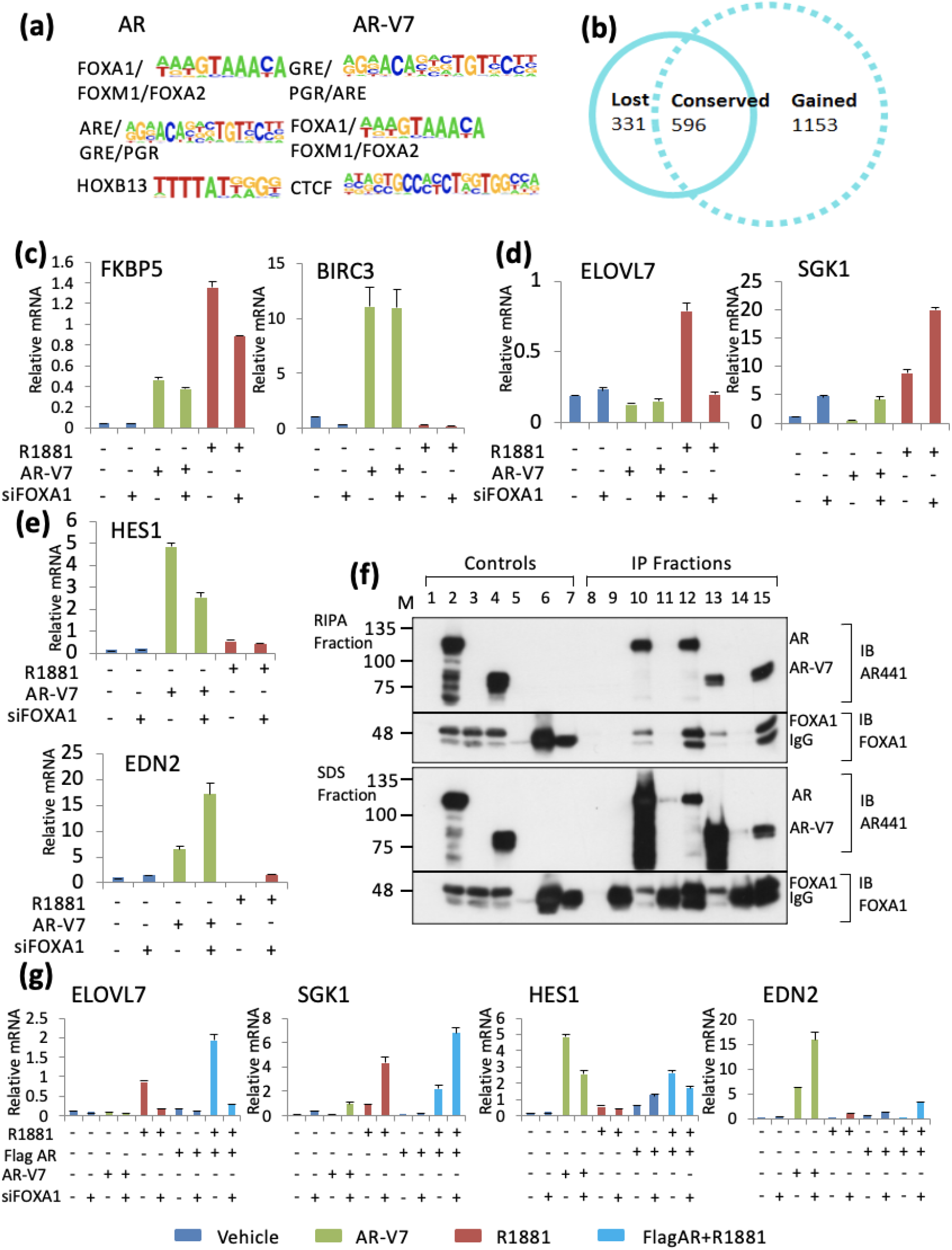
Role of FOXA1 in differential gene regulation. **a**. Top enriched consensus motifs (target vs background *p* < 0.01) identified in a motif analysis performed using HOMER, at AR and AR-V7 binding sites (Red and green circles in Fig. 3, panel a.). **b**. Venn diagram showing number of AR regulated genes lost, conserved or gained upon depleting FOXA1 (data obtained from GSE27824). **c-e**. LNCaP AR-V7 cells in CSS were treated with control or FOXA1 siRNA for 24 hours followed by 24 hour treatment with 10 nM R1881 or 20 ng/mL Dox prior to isolation of RNA and qPCR analysis. **C** FKBP5 and BIRC3 expression **d**. ELOVL7 expression (contains composite FOXA1:AR binding site) and SGK1 **e**. HES1 and EDN2. **f**. CoIP of AR isoforms with FOXA1 performed in HEK293 cells co-transfected with AR and FOXA1, AR- V7 and FOXA1 or FOXA1 alone as control. Lanes 1-7 are input controls. 1. HEK293 cells, 2. AR and FOXA1 transfected cells, 3. FOXA1 transfected cells, 4. AR-V7 and FOXA1 transfected cells, 5. 2 ul AR441 mAb antibody 6. 2 ul rabbit polyclonal IgG, 7. 2 ul FOXA1 antibody, 8-15. IP fractions. 8. FOXA1 transfected cells IP with AR441 monoclonal antibody, 9. FOXA1 transfected cells IP with IgG antibody, 10. AR and FOXA1 transfected cells IP with AR441 antibody, 11. AR and FOXA1 transfected cells IP with IgG antibody, 12. AR and FOXA1 transfected cells IP with FOXA1 antibody, 13. AR-V7 and FOXA1 transfected cells IP with AR441 antibody, 14. AR-V7 and FOXA1 transfected cells IP with IgG antibody, 15. AR-V7 and FOXA1 transfected cells IP with FOXA1 antibody. **g**. Depletion and treatments were performed as in Fig. 6, panel c; except that the Flag-AR line also was used. qPCR analysis measuring ELOVL7, SGK1, HES1 and EDN2 expression in FOXA1 depleted LNCaP AR-V7 and LNCaP AR cells treated with vehicle, 20 ng/mL Dox, 10 nM R1881 or Dox+R1881 in CSS. Green bars represent AR-V7 overexpressed condition, dark blue bars represent AR overexpressed and AR activated condition and red bars represent AR activated condition. All qPCR plots are mean expression values of three biological replicates normalized to 18S on the y-axis and error bars are SEM.

The sensitivity of some AR-V7 regulated genes to FOXA1 raised the question of whether any of the sensitivity could be due to direct interaction or whether the effects were indirect. The AR binding site for FOXA1 has been reported to be in the DBD and hinge region of AR ^34^ and early reports failed to detect interaction ^17, 32^. AR-V7 lacks the hinge region. To address this question, AR negative HEK293 cells were transfected with AR and FOXA1 expression vectors, with FOXA1 alone, or AR-V7 + FOXA1. Cell extracts were incubated with antibodies (AR, FOXA1, or control rabbit IgG), and immunoprecipitations performed as described in methods.

Complexes were sequentially extracted with RIPA (does not release IgG) to facilitate detection of FOXA1, followed by treatment with SDS sample buffer to release IgG and remaining proteins. Immunoprecipitation with AR441 showed that both AR isoforms can co-IP FOXA1 (Fig. 6f lanes 10, 13) and FOXA1 antibody also precipitated AR and AR-V7 (Fig. 6f lanes 12, 15)

To test whether AR overexpression overcomes the effects of FOXA1 depletion on AR mediated transcription, LNCaP AR cells were treated with control or FOXA1 siRNA followed by induction of AR isoforms and treatment with R1881. Although overexpression altered levels of gene expression, it did not alter the sensitivity of any of the genes to FOXA1 depletion suggesting that preference likely is not due to FOXA1 sequestering a limiting amount of AR (Fig. 6g).

### HOXB13 does not show preferential localization with AR-V7 specific sites and has mixed effects on AR-V7 activity

HOXB13 has been described as a factor required for AR-V7 binding and gene regulation in CRPC ^18^. A comparison of our ChIP-exo with published ChIP-seq data of transcription factor binding in LNCaP cells, reveals the expected enrichment of FOXA1 binding at AR specific and common binding sites (Fig. 7a); binding in the vicinity of AR-V7 specific sites is also enriched, but to a lesser extent than the other classes (Fig. 7a). HOXB13 is found associated with all three classes of sites, but, if anything, was more enriched at the AR specific sites (Fig. 7a). Using publicly available data sets, binding of three other factors, NFIB ^35^, CTCF ^36^, and cMyc ^37^ were examined, with the latter two showing modest enrichment at AR-V7 specific sites. To test the effect of HOXB13 depletion on AR isoform dependent gene expression, LNCaP AR-V7 cells were treated with control or HOXB13 siRNA followed by treatment with Dox or R1881 as described in methods. HOXB13 was substantially depleted (SFig. 18). Both hormone dependent ORM1 expression and AR-V7 dependent ORM1 expression were reduced by depletion (Fig. 7b) as was expected from previous studies ^18, 38^. However, if anything, depletion enhanced AR-V7 dependent induction of FKBP5 while having little effect on AR activity. SGK1 induction by R1881 also was dependent on HOXB13, but depletion did not induce responsiveness to AR-V7. A few AR-V7 preferential genes also were examined and all three showed enhanced induction when HOXB13 was depleted (Fig. 7c,d). At the genome level, HOXB13 and AR-V7 binding sites were not enriched relative to AR, HOXB13 binding (Fig. 7f). A comparison of motifs and transcription factor binding at transcription start sites is shown in SFig. 19a-d. Although the HOXB13 motif was found to be enriched to some extent at both AR and AR-V7 binding sites (SFig. 19d), an analysis of binding at the TSS shows modest enrichment of HOXB13 and no better correlation with AR-V7 than several other factors (SFig. 19b-c). When the analysis is further limited to TSS in genes that are regulated by AR or AR- V7 and bound by the corresponding AR isoforms, the differences are more profound. AR bound TSS sites are greatly enriched for FOXA1 binding whereas AR-V7 sites show minimal enrichment (SFig. 20a) with similar and weaker enrichment of HOXB13 at these sites (SFig. 18a,b). None of the other factors showed preferential enrichment with AR-V7 (SFig. 20b).

**Figure 7.**
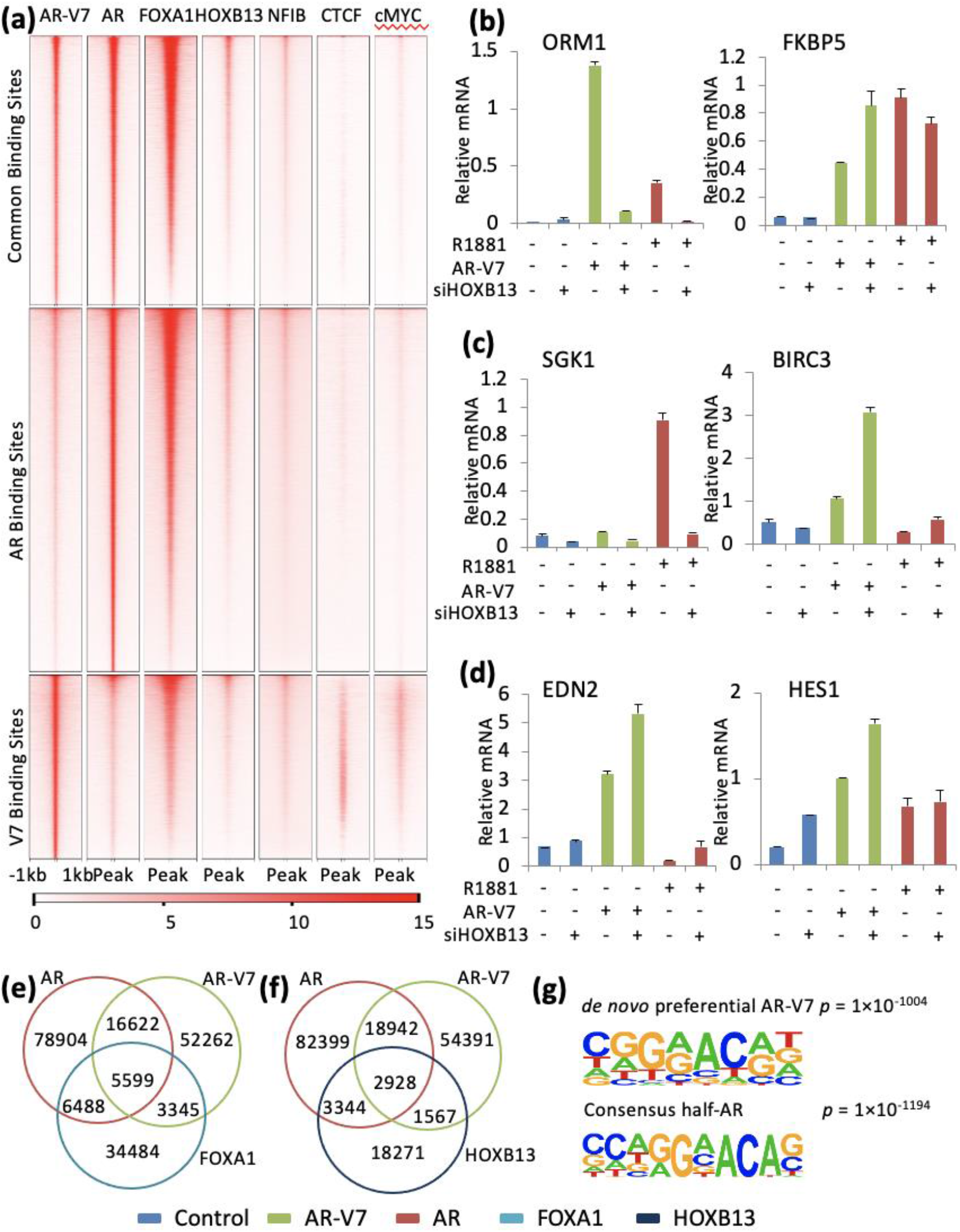
Role of other factors in differential gene regulation. **a.** AR, AR-V7, FOXA1, HOXB13, NFIB, CTCF, cMYC average ChIP-seq signal intensity per every 10bp bin at common, AR and AR-V7 binding sites identified in Fig. 3, panel a. Data were generated from LNCaP cells and ChIP signals plotted are within ±1 kb from the center of AR and AR-V7 binding sites using publicly available data listed in STable 1. **b-d**. qPCR analysis measuring gene expression upon HOXB13 depletion using the method previously described for FOXA1. All qPCR plots are expression values normalized to 18S on the *y*-axis and corresponding treatment on the *x*-axis. Data are plotted as mean of three biological replicates from the same experiment and error bars are SEM. Each experiment was performed a minimum of three times and a representative experiment is shown here. **e**. Venn diagram showing number of AR and AR-V7 binding sites in LNCaP AR-V7 cells overlapping FOXA1 binding sites in LNCaP cells (peaks with a local FC > 5 and at least a base pair overlap). **f**. Venn diagram showing number of AR and AR-V7 binding sites in LNCaP AR-V7 cells overlapping HOXB13 binding sites in LNCaP cells (peaks with a local FC > 5 and at least a base pair overlap). **g**. Consensus sequence reported by HOMER motif analysis of all AR-V7 peaks identified from LNCaP AR-V7. Motif enrichment *p value* for *de novo* motif and known half ARE motif were calculated using cumulative binomial distribution. Venn diagrams **e** and **f** are plotted as symmetrical circles and not proportional to numbers.

To look for other mechanisms for preferential binding, *de novo* motif analysis of all AR isoform binding sites was performed. A sequence (CGGAACAG) was detected as enriched in both the AR-V7 specific binding sites and in the sites common to AR and AR-V7, but not in the AR unique peaks or in the overall pool of AR binding sites (Fig. 7g). This site is similar to a consensus half-ARE and differs by one nucleotide from a consensus ZNF189 site (SFig. 21a). When AR, AR-V7, FOXA1 and ZNF189 binding are compared between *de novo* sites and consensus half AREs in AR-V7 bound sites, the *de novo* sites are preferential for AR-V7 with almost no FOXA1 or ZNF189 binding (SFig. 21b), whereas the consensus half AREs are strongly enriched for FOXA1 (SFig. 21c). Thus, the factors surrounding these classes of sites appear to be distinct and potentially impact the unique transcriptomic profiles of AR specific isoforms in prostate cancer models.

## Discussion

There is much interest in the role of splice variants, especially AR-V7, in CRPC; AR-V7- expressing tumors typically are refractory to second line ADT and there is a great need to develop therapies that target variants lacking the LBD and to identify variant actions that might be targeted therapeutically. The studies seeking to elucidate variant action have yielded conflicting results. There is good agreement that AR-V7 and other variants can regulate many of the same genes, but there is a substantial debate about whether AR-V7 and other variants also regulate a substantial set of unique targets or have minimal or no unique activities^16–20, 25, 26, 39^. Similarly, some investigators have identified unique AR-V7 chromatin binding sites, while others have described these as artifacts/unreliable ^16, 17, 19, 40^. Some investigators have concluded that AR-V7 acts with and/or requires AR for activity whereas others describe AR- V7 actions as independent of AR ^14, 16, 24, 29^. Many of the studies have been done in LN95 and 22RV1 cells ^14^, which endogenously express low level of variants, and/or also express other active variants ^41^, but neither cell line was derived from a tumor expressing variants ^20, 21^. These lines also have undergone changes during their derivation ^42^, so there is no AR only line suitable for a direct comparison. To more precisely compare the capacity of AR and AR-V7 to regulate gene expression and bind to chromatin, LNCaP and VCaP lines that express AR-V7 and an LNCaP line that expresses additional AR in response to Dox treatment were used. Isoform dependent gene expression was examined in these lines as well as in LN95 cells.

Previous studies have suggested that AR-V7 might be a “weak” AR. However, when expression levels of AR and AR-V7 are comparable in the LNCaP model, the number of genes regulated by the two isoforms are similar (Fig. 2c). While the isoforms regulate many of the same genes and androgen signalling is the primary Hallmark pathway for both isoforms, there also are many targets unique to each isoform (Fig. 2a,c). This is consistent with our finding that AR and AR- V7 differentially regulate metabolism in this model ^43^. Similar results were obtained in the VCaP model (Fig. 2b,c) although there appear to be more genes regulated by AR. Two potential contributors to this difference are the higher levels of AR in the VCaP model and the basal expression of AR-V7 diminishing sensitivity to the induced AR-V7.

Isoform specific gene regulation was also detected in the LN95 model (SFig. 10) when AR-V7 depleted cells were compared with R1881 treated cells. A previous report using LN95 derivative cell lines that expressed shRNA targeting AR, AR-V7, or GFP in response to Dox suggested that AR-V7 was predominantly a repressor ^16^. However, in all three of our models, AR-V7 induces a slightly higher proportion of total regulated genes than does AR. Even when the genes examined in the LNCaP model are restricted to the ones with AR-V7 binding (Fig. 3c) 1297 of 2310 genes (56.14%) bound by AR-V7 are upregulated whereas 1167 of 2251 genes (51.84%) bound by AR are upregulated. Part of the rationale for the repressor role for AR-V7 is that it fails to interact well with the classical LXXLL motifs known to recruit coactivators to the LBD of receptors such as estrogen receptor (ER). However, it is well established from molecular studies as well as Cryo-EM studies, that unlike ER, the primary interaction site for coactivators such as the SRC family is in the amino-terminus of AR ^44–46^. Thus, AR-V7 retains the primary AR transactivation domain although it does not recruit coregulators dependent on binding to the LBD.

Consistent with differential gene regulation, strong differential binding to chromatin was detected. In the LNCaP model, 87,095 chromatin binding sites were detected for AR versus 59,852 for AR-V7 with 32,082 common to both isoforms. The differential binding of AR isoforms to chromatin despite a common DBD, is a major factor in differential gene regulation. The finding that while some genes, including common targets such as PSA, are optimally expressed at levels of AR or AR-V7 comparable to AR in LNCaP cells while others are more highly regulated at higher levels of AR or AR-V7 (Fig. 5) likely explains some of the discrepancies in the literature. Many of the unique binding sites may be lower affinity and thus more receptor is required for detection of binding and gene regulation. This implies that not only are there quantitative differences in gene expression as a function of receptor level, but also changes in the specific genes regulated. The limited studies of AR-V7 expression in tumors and CTCs suggest that at least some human tumors express levels comparable to those in these studies ^7, 10^.

One question that has clinical relevance is what, if any, role AR plays in AR-V7 action. Depletion of AR in the LNCaP model with induced AR-V7 had no effect on PSA induction, nor did depletion in the LN95 line (Fig. 1). However, basal levels of PSA were reduced by AR depletion in LNCaP AR-V7 cells not treated with Dox; this raised the possibility that the very low levels of AR-V7, due to leakage of the system, does require AR for activity. However, somewhat to our surprise, depletion of AR in LNCaP cells also reduced basal PSA suggesting a low level of aberrant AR activation. A previous report that depletion of AR in LNCaP cells in CSS induced apoptosis ^47^ is consistent with AR retaining some activity in these cells despite hormone depletion. In the LN95 model, depletion of AR altered expression of about 200 genes whereas depletion of AR-V7 altered expression of nearly 1000 genes, of which only 61 overlapped with the AR genes. Thus, in this model, little or none of the AR-V7 dependent activity requires AR. Consistent with other studies suggesting that AR and AR-V7 can heterodimerize ^48^ and are present on the same chromatin site ^16^, we detected a small amount of recruitment of AR to some sites in chromatin in response to expression of AR-V7 in the LNCaP. However, the relative amounts indicate that the vast majority of the receptor is AR-V7 and thus, for the most part, not a heterodimer with AR. Although the two are colocalized, it also is possible that an AR-V7 homodimer is recruiting AR through the AR-V7 amino-terminal domain. Consistent with this idea is the report that the FXXLF motif in AR-V7 is required for heterodimerization ^24^. A recent study in LN95 cells ^14^ also confirms that there is little recruitment and no need for AR for AR-V7 chromatin binding.

All models exhibit common and isoform specific gene expression, but the overlap in either hormone dependent or AR-V7 dependent gene expression between models is variable and often rather modest. The strongest overlap detected was between R1881 treatment of LNCaP and LN95 cells (SFig. 11). The LN95 cells were derived from LNCaP and the expression levels of AR are similar. The LN95 cells show little AR dependent regulation in the absence of hormone (SFig. 10). This is in contrast to the CRPC derivatives of LNCaP, LNCaP ABL, and C4-2B (Fig. 2f,g) which express only AR. It was formally possible that the AR-V7 signature would be more similar to the aberrant AR activation in these lines, but the comparisons show little overlap (Fig. 2f,g). All other comparisons using our models were between receptors at different levels and/or in different backgrounds and both likely contribute to these differences. One challenge in studying variant actions is that they are constitutively active, so there is no good approach to study short term gene regulation, which presumably would show more overlap. In some cases, even studies using the same line (22RV1) using slightly different approaches also yield modest overlaps of genes detected (Fig. 2h,i) and another study found divergent AR-V7 cistromes in CRPC cells and tumors ^18^.

The two most likely mechanisms for differential regulation of target genes are differential recruitment of coregulators and differential binding to DNA, which could result from differences in direct binding or differential interactions with other DNA binding factors. As noted above, the ChIP-exo analysis reveals that not only are there common AR-V7 and AR sites, but also many isoform specific sites. The two isoforms contain the same DNA binding domain suggesting that much of the AR isoform specific binding relies on additional DNA binding proteins. Thus, the differences in activity may be predominantly a function of these differential actions rather than differential recruitment of coactivators and corepressors. However, the studies presented here support the conclusion that there are multiple mechanisms that contribute to differential binding and/or gene expression.

Although there are many preferential sites throughout the genome, there is marked preferential interaction of AR-V7 near the TSS (Fig. 3b and SFig. 12). This is consistent with recent studies that also detected enrichment of AR-V7 near TSS in 22RV1 cells ^17, 19^. However, less than 20% of genes differentially regulated by AR-V7 (435/2310) in the LNCaP model exhibit AR-V7 binding at the TSS. Moreover, an analysis of the consensus binding motifs surrounding these AR-V7 binding sites revealed that whereas upregulated genes contained AREs, binding regions in repressed genes were enriched for FOXA1 motifs suggesting additional specificity (SFig.13). In contrast there was much less AR binding at the TSS and no major differences in motifs for up and down-regulated genes.

Several factors (HOXB13, ZFX, and FOXA1) have been suggested to play a role in preferential actions of the AR isoforms ^18, 19, 26, 49^. ZFX has been suggested as a factor that contributes to AR-V7 dependent action at the TSS ^19^. Since most CpG island promoters bind ZFX ^50^, it is not surprising that transcription of these genes is affected by ZFX. However, the high frequency of ZFX binding suggests that it is not the primary factor determining whether AR-V7 will bind to the promoter and regulate transcription of the subset of AR-V7 genes with TSS binding. HOXB13 was reported to be required for AR-V7 dependent induction of unique target genes ^18^, but similar to a previous study of AR dependence ^38^, we see both loss and gain of function of AR and AR-V7 when HOXB13 is depleted (Fig. 7) and no compelling enrichment of HOXB13 colocalization with AR-V7 in our larger data set of AR-V7 regulated genes (Fig. 7, SFig. 20). Interestingly, induction of SGK1 by AR was strongly dependent upon HOXB13 expression whereas AR-V7 was inactive regardless of HOXB13 expression.

Although FOXA1 expression can modulate the activity of both AR and AR-V7, FOXA1 has a much more substantial impact on a subset of AR activities (Fig. 7). We previously reported differential responsiveness of AR and AR-V7 at genes with AR FOXA1 composite elements (RASSF3 ^26^ and INPP4B ^33^), whereas a gene (EDN2) ^26^ that is only induced by AR when FOXA1 is depleted is induced by AR-V7 regardless of FOXA1 status ^26^. One possible explanation for these differences was differential binding of FOXA1 to AR and AR-V7. Our studies show that AR-V7 can interact with FOXA1 as measured by a co-ip assay (Fig. 6e). However, the three genes (RASSF3, INPP4B, and ELOVL7) that we have examined that are induced only by AR and absolutely require FOXA1 for induction contain at least one composite element which binds AR, but not AR-V7 suggesting that there are differences in the capacity to bind to some of these elements. In our limited analyses of the role of FOXA1 expression in regulation of AR-V7 activity, we have seen no effect (BIRC3), a reduction in activity (HES1) and an increase in activity (EDN2) in response to FOXA1 depletion (Fig. 6c,e), but in no case do we see a complete loss of regulation as is seen for a subset of AR regulated genes. Interestingly, EDN2, which is induced by AR-V7, but repressed by AR in the presence of FOXA1 and is induced by both when FOXA1 is depleted, remains much more highly induced by AR-V7 even when AR is overexpressed (Fig. 6g) suggesting additional factors determining efficacy of induction.

In looking for other factors that might bind, we identified a *de novo* sequence similar to a half ARE (Fig. 7) that is much more highly enriched at AR-V7 binding sites than AR binding sites. These sites lack the abundant FOXA1 binding found associated with a classical half ARE (SFig. 21) raising the possibility that there may be an additional unknown factor contributing to AR- V7 binding.

One possible limitation of our transcription factor colocalization studies is that published data measuring localization of factors in parental LNCaP cells were used for all comparisons. Although these factors all bind to chromatin independent of receptors, it is formally possible that AR-V7 could induce relocalization or a change in expression of some factors and these changes would not have been detected in our studies.

Using inducible models as well as the LN95 model, both common and isoform specific transcriptional activities were detected in our studies. Consistent with this, when similar levels of AR and AR-V7 were expressed, the isoforms were detected both at common and unique sites using ChIP-exo. Depletion studies showed that AR-V7 functions independently of AR and at the selected sites examined, there was little recruitment of AR by AR-V7. The finding that expression levels of AR or AR-V7 can determine levels of gene induction, while not unexpected, provides a potential explanation for some of the studies that failed to detect variant specific activities.

Our studies in LN95 cells supported the finding of unique isoform dependent activities, but there are differences in the results among investigators who have studied either LN95 or 22RV1 cells. Different approaches have been taken and this may contribute to the different conclusions. Generally, investigators who have used stable expression or depletion approaches to produce separate cell lines that have elevated or reduced levels of AR isoforms and use these for comparisons detect fewer isoform specific differences in gene expression/binding than those who use transient methods (siRNA or PROTAC), so that all comparisons are within the same passages of a single cell line. Another potential confounding factor is the level of steroid depletion in charcoal stripped serum as has been suggested by Liang et al ^14^. Residual androgens can activate AR. For these studies, commercial batches of CSS were pre-screened for levels of aberrant AR activation in LNCaP cells (reduction in expression in response to enzalutamide) and batches with no background activity were used and/or serum was stripped in the lab and lack of activity confirmed in the same manner. The LNCaP (and LN95) AR have a mutant hormone binding domain and respond to a broader range of steroids. Thus, they may be more sensitive to residual steroids than lines with wild type receptor.

Our studies utilizing the inducible models are unique in that the levels of AR-V7 were comparable to LNCaP AR levels to determine the capacity of AR-V7 to bind to chromatin and regulate gene transcription. At this level of receptor, AR-V7 shows several unique activities, a capacity to bind to TSS, enrichment of binding to a novel ARE related sequence and differential functional interactions with FOXA1. Although coregulators presumably also play an important role, the vast majority of differentially regulated genes showed differential binding to chromatin implicating interactions with other DNA binding proteins and potentially different DNA sequences in the unique receptor activities. These studies were limited to defined cell lines, but the results suggest that the activities of AR-V7 in CRPC will be heavily dependent on the level of AR-V7 expressed in the tumor and serve as the preclinical foundation for discerning the isoform specific function in disease progression.

## Materials and Methods

### Cell lines

LNCaP, VCaP, 22RV1, and HEK 293T/17 cells were obtained from the American Type Culture Collection (ATCC) Manassas, VA and grown as recommended by the provider. The engineered derivatives of LNCaP and VCaP ^25^ were grown under the same conditions except that G418 was included to prevent loss of lentivirus. LN95 cells were kindly provided by Dr. Jun Luo, Johns Hopkins University, and are grown in phenol red free RPMI 1640 with 10% charcoal stripped serum (CSS). All cell lines have been authenticated by STR analysis and are maintained in culture for up to three months before a new batch is thawed. Mycoplasma testing is done routinely every few months.

### Chemicals

Cell culture media are purchased from Gibco, a division of ThermoFisher Scientifiic (Waltham MA). Fetal bovine serum (FBS) and CSS were purchased from Sigma Aldrich, St. Louis, MO. In some cases, FBS was stripped with charcoal dextran in the cell culture core to produce CSS. R1881 and doxycycline (Dox) are purchased from Sigma Aldrich and Enzalutamide from Selleck chemical LLC, Houston, TX. If not otherwise indicated, chemicals are reagent grade.

### Antibodies

An AR antibody (RRID: AB_2793341) that recognises the N-terminus of AR was used by Active Motif for ChIP-exo analysis and this was also used for ChIP-qPCR. Other antibodies used in the ChIP-qPCR assays include, AR-N20 (RRID:AB_1563391) Santa Cruz Biotechnology, Santa Cruz, CA, AR C-19 (RRID:AB_630864), rabbit anti mouse IgG (RRID:AB_228419). These antibodies and FOXA1 (RRID:AB_2104842), HOXB13 (GTX129245, GeneTex, Irvine, CA), Tubulin (RRID:AB_309885), Actin (RRID:AB_11004139), and AR-V7 (RRID:AB_2631057) antibodies were used for western blotting also. AR441 was used for western blotting and immunoprecipitation. It was grown in the BCM Protein and Monoclonal Antibody Production core, but also is sold as RRID AB_11000751 by ThermoFisher Scientific. FOXA1 ChIP validated antibody (RRID:AB_2104842) was used for CoIP experiments.

### Bacterial strains and plasmids

DH5a and TOP10 competent cells were purchased from Thermo Fisher Scientific, Waltham, MD and used for plasmid transformation and propagation as per manufacturer’s instructions. PCR amplified cDNA of AR (full length) was cloned from an available pCR3.1 expression vector ^51^ by fusing with 3X FLAG sequences at the beginning and HA-Tag at the end. This was cloned into the pHAGE lentiviral expression vector as previously described ^26^. pCR3.1 AR-V7 has been described previously ^52^ pLX302_FOXA1-V5 plasmid was purchased from Addgene, Watertown, MA.

### Transient RNA interference

Knock-down of FOXA1, HOXB13, AR, AR-V7 expression was performed using transient transfection of siRNA against the gene in comparison to non-targeted negative control siRNA^26^. Synthesised oligo duplexes were purchased from Sigma-Genosys, The Woodlands, TX (sequences and source listed in STable 2). Cells were transferred to medium containing CSS and transfected with 35-100 nM siRNA using the Lipofectamine RNAiMAX reagent (Invitrogen, Rockville IL) according to the manufacturer’s directions for a duration of 48-72 hours.

### Cell culture and treatment protocol

For experiments to test AR variant binding and activity, cells were ligand-starved for 24 hours using RPMI-1640 base medium supplemented with 10% CSS, before treating with vehicle (0.1% ethanol), 20 ng/mL (LNCaP lines) or 100 ng/mL Dox (VCaP) or 10 nM R1881. All cells were used within 25 passages of thawing and inducible cells were maintained under G418 selection and transferred to medium lacking G418 for individual experiments.

### Stable cell line generation using lentiviral production

Lentivirus was prepared using HEK293T/17 cells and a packaging system as reported earlier ^43^. The LNCaP cell line with inducible expression of flag tagged AR was generated by infection of lentivirus encoding Flag tagged AR, followed by G418 selection in RPMI-1640 medium for two weeks and termed LNCaP^pHAGE flAR^ (LNCaP AR).

### Reverse Transcriptase Quantitative PCR (RT-qPCR)

Cells were harvested in PureXtract RNAzol solution (GenDepot, Barker, TX), RNA isolated, cDNA synthesised from 500 µg total RNA using an AmphiRevert cDNA synthesis kit (GenDepot, Barker, TX). Primers (STable 2) and SYBR Green PCR Master Mix (Applied Biosystems by Life Technologies, TX) were used for q-PCR analysis using the StepOnePlus Real-Time PCR System (Applied Biosystems, Foster City, CA). All qPCR plots are expression values normalized to 18S on the y-axis and corresponding treatment on the x-axis. Data are plotted as the mean of three biological replicates from the same experiment; error bars are SEM.

### Co-immunoprecipitation (CoIP)

CoIP was performed from biological replicates of HEK293T/17 cells maintained in DMEM with 10% CSS. Briefly, HEK293T/17 cells were transfected with pLX302_FOXA1-V5, pLX302_FOXA1-V5 + pCR3.1 AR or pLX302_FOXA1-V5 + pCR3.1 AR-V7 expression vectors using 15 µl Lipofectamine 3000 Reagent and 10 µl P3000 reagent (Thermo Fisher Scientific, Waltham, MA) for 28 hours. After 24 hours of treatment pCR3.1 AR transfected cells were treated with 10 nM R1881 for 4 hours. Biological replicates of cell pellets were pooled together and lysed by incubating in 1X TESH lysis buffer (40X TESH made of 0.4 M Tris, 0.04 M EDTA and 0.48 M Monothioglycerol pH 7.6) ^53^ with 0.15 M NaCl on ice for 2 hours. Lysates were split into equal fractions of approximately 250 ug protein, and 2 ug of antibody or control IgG were added. Tubes were incubated for 1 hour at 4°C and then protein A/G agarose magnetic beads (Thermo Scientific) were added for 10 minutes at room temperature. Beads were washed three times in 1X TESH buffer with 0.15 M NaCl and resuspended in 40 µl of RIPA buffer (G-Biosciences, MO) on ice for 1 hour. The RIPA fraction was transferred to a new tube and a second fraction was extracted from the same beads using 40 µl 2X SDS sample loading buffer. SDS sample loading buffer was added to the RIPA fraction, samples were boiled at 100°C for 5 minutes. RIPA samples were run on a 7.5% SDS-PAGE gel with a set of controls and SDS samples on another gel with a second set of controls and proteins detected by western blotting, using Protran nitrocellulose membrane (PerkinElmer, Waltham, MA) and ECL2 western blot substrate (Pierce, ThermoScientific, IL) to visualise signals. Each nitrocellulose membrane was split to process the upper portion with AR441 mAb and the lower portions with the FOXA1 antibody.

### ChIP assays and ChIP-exo data analysis

ChIP assays were performed as previously described ^26^ with the modifications using AR antibody (ActiveMotif, Carlsbad, CA) or AR C-19 (Santa Cruz Biotechnology, Dallas, TX) as mentioned in the Supplementary methods (SMethods). Data generated using the ChIP-exo method ^54^ were used to map AR and AR-V7 interaction using AR antibody (ActiveMotif, Carlsbad, CA). ChIP-seq and ChIP-exo (STable 1) were analysed as described in SMethods, using the bowtie2 (version 2.3, default parameters) alignment software ^55^ and peaks were called using macs2 (version 2.1; -*q* <.05, otherwise mentioned) ^56^.

### RNA-seq data generation and analysis

In short, RNA-seq was performed using LN95 cells treated with either: 10 nM R1881 or an equivalent volume of the vehicle control. For knock-down experiments cells were treated with control siRNA, siRNA targeting exon7 or siRNA targeting cryptic exon of AR-V7 for 48 hours before harvesting and processed as described in SMethods. Total RNA was extracted, and column purified using an RNAeasy mini kit. (Qiagen). The clean RNA samples having RIN more than 8.0 were used for library preparation in the Genomic and RNA Profiling Core, Baylor College of Medicine. Libraries were constructed using a Truseq library preparation kit (Illumina, USA) following the manufacturer’s instructions. cDNA libraries were sequenced on the Illumina NextSeq-500 or NovaSeq-6000 platform in the Genomic and RNA Profiling Core facility, Baylor College of Medicine. This RNA-seq dataset with LNCaP AR-V7 and VCaP AR-V7 datasets listed in STable 1 was used to identify the AR and AR-V7 transcriptome. In short HISAT2 ^57^ and STAR ^58^ was used to align raw reads to human genome build hg19 (V24) gene expression was measured using featureCounts ^59^ and DEGs calculated using edgeR ^60^. Potential direct target DEGs were identified by integrating ARBs, V7Bs and DEGs as described in SMethods.

### Analysis of public datasets

We utilised public gene expression datasets and ChIP datasets listed in STable 1 for comparisons in this study. Raw sequence data was processed as per the previous description and SMethods.

### Statistical analysis

Three independent experiments were performed using separate batches of cells with biological triplicates within each experiment for qPCR assays. For western blot, qPCR and ChIP-qPCR assays, representative examples are presented although the experiments were performed a minimum of three times. Representative genes from subset of genes induced or repressed by FOXA1 or HOXB13 were tested using qPCR assays. Student’s *t*-test is used to calculate difference in mean gene expression between the treatment and control group.

## Data availability

The LN95 datasets generated during this study can be accessed at GSE184676.

## Acknowledgements

RNA-seq studies were performed in the Baylor College of Medicine (BCM) Genomic and RNA Profiling cor. ARR441 was prepared by the BCM Protein and Monoclonal Antibody Production core. The authors thank the Molecular and Cellular Biology BCM tissue culture core for assistance in growth and maintenance of cell cultures. The authors thank Dr. Hari Krishna Yalamanchili for his valuable suggestions on bioinformatic analyses and Dr. Harika Nagandla for assistance in preparing SFig. 1 and characterizing and maintaining LNCaP and VCaP AR- V7 cell lines.

## Author contributions

NLW, PB, and CC conceived the study. PB and WEB performed the experiments. AKD, WCK and AAS prepared samples for RNA-seq and ChIP-exo sequencing. PB, MJR, BP and CC performed data analyses. PB and NLW interpreted the results and wrote the manuscript with input from others. All authors reviewed the manuscript.

