## Supplementary material for "Cistrome and transcriptome analysis identifies unique androgen receptor (AR) and AR- V7 splice variant chromatin binding and transcriptional activities": Supplementary_bp copy.pdf

**Supplementary Methods**

**ChIP analyses**

ChIP assays were performed from nuclear extracts prepared in a water bath sonicator (Bioruptor from Diagenode, Denville, NJ). Samples were sonicated with 10 cycles of 30 seconds on and 30 seconds off each in a final volume of 1 mL. 5% of the lysate was reserved as input and the remainder was pre-cleared with 25 µL of Protein A/G Magnetic Agarose (Pierce, Thermo Scientific, Waltham, MA) conditioned with 20 µg/mL salmon sperm DNA. Lysates were split into equal portions and immunoprecipitations were performed with 50 µL of Magnetic Agarose using 2 µg of rabbit anti-mouse IgG (Invitrogen), 2 µg of AR antibody (ActiveMotif, Carlsbad, CA) or AR C-19 (Santa Cruz Biotechnology, Dallas, TX). Eluted ChIP fractions were quantified by qPCR and normalised using the primer sequences listed in STable 1.

ChIP-exo datasets in STable 1 were used to map AR and AR-V7 interaction with DNA at single base resolution. Cells in CSS were treated for 24 hours with 10 nM R1881 (LNCaP) or 20 ng/mL Dox (LNCaP AR-V7). Cells were fixed in 1% formaldehyde. Six independent experiments were performed, and cells shipped to ActiveMotif where like samples were combined and ChIP-exo performed using the ActiveMotif kit. ChIP-exo libraries were prepared using the TruSeq ChIP library preparation kit from Illumina, San Diego, CA and sequenced on an Illumina HiSeq 2000 platform.

**ChIP-seq data analysis**

ChIP-seq and ChIP-exo raw reads were aligned to the human reference genome from UCSC genome (hg19) by the bowtie2 (version 2.3, default parameters) alignment software [55]. Duplicate reads were ignored while calling peaks using macs2 (version 2.1; -*q* <.05, otherwise mentioned) with input as control when available [56]. Peaks overlapping (at least 1bp) with the “hg19 blacklist regions” reported by the ENCODE project [61] were also removed. Peaks were annotated as AR variant specific or common peaks based on their overlap in genomic coordinates (at least 1bp). Annotation function in HOMER (version 4.10) [62] was used to assign peaks to genes. In-house scripts and BEDTools (version 2.28.0) [63] were used to annotate proximal promoter (±100 bp from transcription start site, TSS) and distal enhancer peaks (upstream to transcription start site, without interfering genes and -100bp to TSS) using the hg19 RefSeq annotation as reference. Genes with either a promoter or enhancer peak were considered to be potential AR isoform targets and target genes with differential expression as possible direct targets. ChIP-seq intensities and read coverage were calculated (at each 10 bp bin, by extending 100 bp) from aligned reads at the reported peak regions. Average ChIP-seq intensities within ±1000 bp of peaks were plotted as profile and heatmaps using deepTools [64]. Overrepresentation of motif sequences in the peak locations were identified by the software HOMER, within 200 bp of reported peak summits.

**RNA-seq data generation**

RNA-seq datasets in STable 1 were used to probe the AR and AR-V7 transcriptome. In short, RNA-seq was performed using LN95 cells treated with either: 10 nM R1881 or an equivalent volume of the vehicle control. For knock-down experiments cells were treated with control siRNA, siRNA targeting exon7 or siRNA targeting cryptic exon of AR-V7 for 48 hours before harvesting. LNCaP $^{AR\text{-}V7/pLenti}$ cell samples [26] and VCaP AR-V7 were treated with either: 10 nM R1881, 0.25 ng/mL Dox, or an equivalent volume of the vehicle control. RNA was isolated After 24 hours. RNeasy column purification kit (Qiagen, MD) was used according to the manufacturer's protocol to extract RNA from all samples. The clean RNA samples having RIN more than 8.0 were used for library preparation. Library preparation was performed with the TruSeq™ RNA Sample Prep Kit v1 (Illumina, San Diego, CA, Cat#FC-122-100). LN95 cDNA libraries were sequenced on the Illumina NextSeq-500 or NovaSeq-6000 platform, LNCaP cDNA libraries were sequenced on the Hiseq 2500 platform, Illumina, San Diego, CA in the Genomic and RNA Profiling Core facility, Baylor College of Medicine and VCaP cDNA libraries were sequenced on the NextSeq 500 platform, (Illumina) by the University of Houston Sequencing core.

**RNA-seq data analysis**

RNA-seq data were used to identify the AR and AR-V7 transcriptome. Raw read sequences from fastq files for each sample were trimmed using Trim Galore (https://github.com/FelixKrueger/TrimGalore). After quality trimming, using a Phred score of 20, the sequences for each sample were aligned to the prebuilt hg19 genome from UCSC using HISAT2 [57] using default parameters. Aligned reads for each sample were then quantified using featureCounts [59] and a transcript annotation reference for the hg19 genome build (V24). Differential gene expression analysis of the protein coding genes was then performed using the Bioconductor package edgeR [60]. A change in expression for a protein coding gene was considered significant if it had a BH-adjusted p-value of less than or equal to 0.05. Gene signatures were generated by filtering the results for statistically significant genes that had at least the indicated log2 fold change in expression. Ranked gene lists of all the detected genes were created for each comparison using the log2 fold change results from the differential gene expression analysis. Gene set enrichment analysis (GSEA) of these ranked gene lists was then performed against the entire molecular signature database (MSigDB) using the java based GSEA software developed by UC San Diego and the Broad Institute [65]. Plots are not proportional unless specifically indicated.

**Analysis of public datasets**

We utilised public gene expression datasets and ChIP datasets listed in STable 1 for comparisons in this study. LNCaP ABL [28] and C4-2B [27] datasets were used for aberrant AR activation signature in prostate cancer cells expressing only AR. Datasets from 22RV1 were used to compare AR and AR-V7 transcriptomes in endogenous models [17,19]. Transcriptome dataset of FOXA1 KD in LNCaP cells was also used in this study. GEO2R function in the GEO database was used with BH multiple testing correction and other default parameters to test differential gene expression. Raw RNA-seq datasets from 22RV1 were also downloaded and analysed as previously mentioned. Other public ChIP datasets used in this study are also listed in STable 1. These include AR ChIP-exo data from LNCaP cells treated with vehicle control, FOXA1 ChIP-exo data from LNCaP cells treated with DHT and HOXB13 ChIP-seq data from LNCaP cells in 10% FBS. Raw sequence data was processed as per the previous description.

#### Supplementary Figure 1

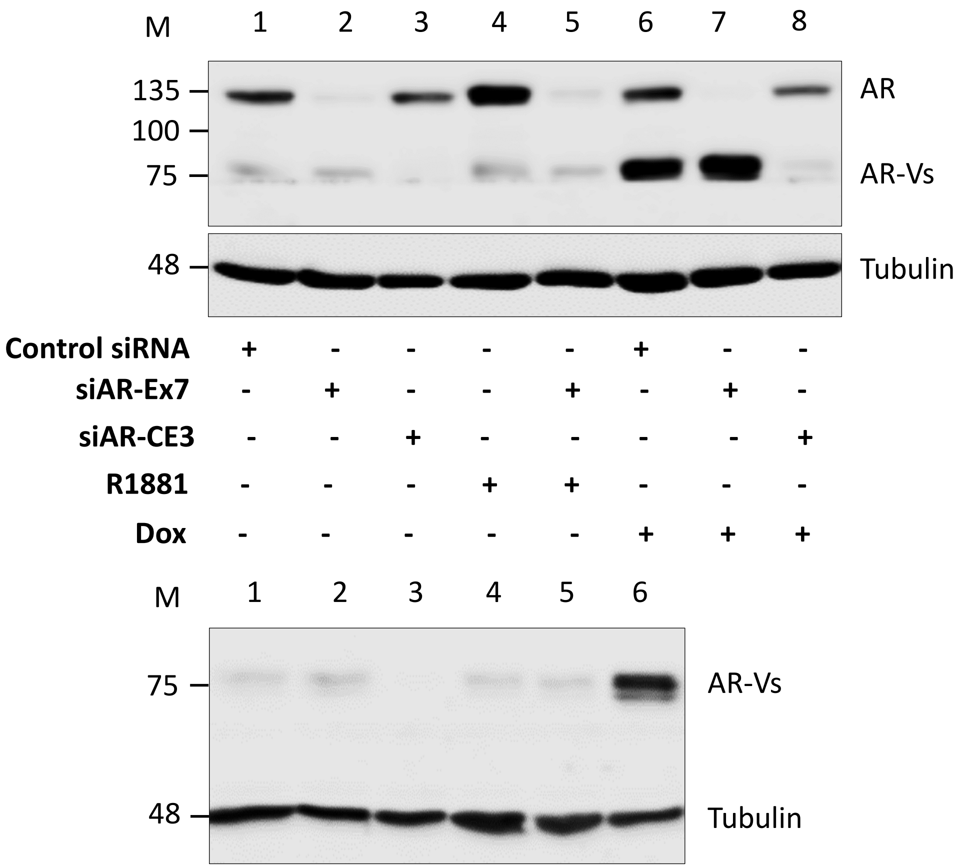


**Supplementary Figure 1. AR variant expression in LNCaP AR-V7 cells.** Cells in 10% CSS medium were treated with the indicated siRNAs for 72 hours, protein lysates prepared, separated by SDS gel electrophoresis and AR isoforms detected using AR441 (top) and AR-V7 antibodies (bottom). Lanes represent LNCaP AR-V7 cells treated with 1. Control siRNA, 2. siRNA against AR (Exon 7), 3. siRNA against AR-V7 (CE3), 4. 1 nM R1881, 5. siRNA against AR (exon 7) + 1 nM R1881, 6. Control siRNA + 20 ng/ml Dox, 7. siRNA against AR (Exon 7) + 20 ng/ml Dox. 8. siRNA against AR-V7 (CE3) + 20 ng/ml Dox.

**Supplementary Figure 2**

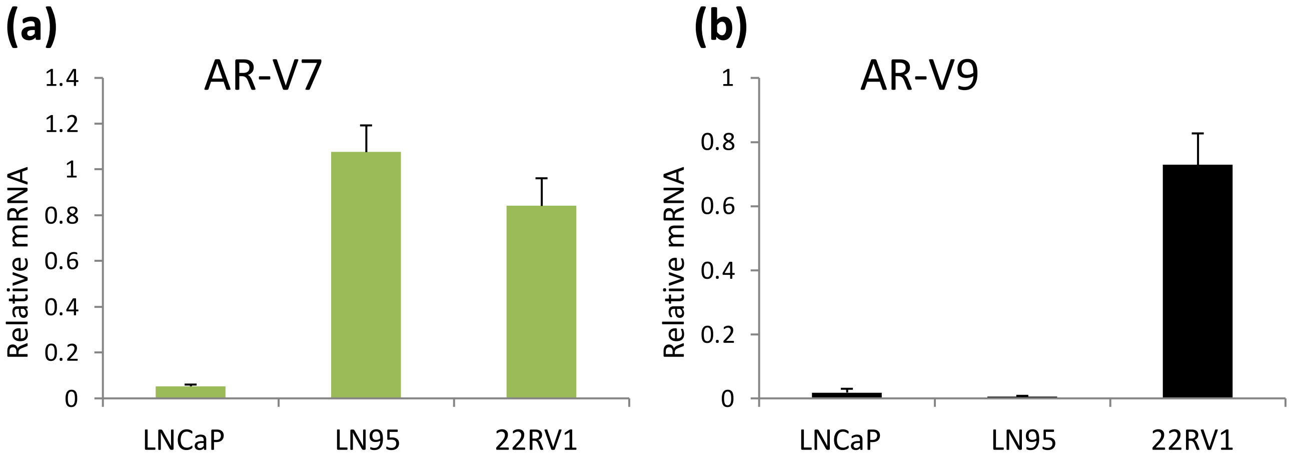


**Supplementary Figure 2. AR-V9 expression in different cell lines**. Cells were transferred to medium containing CSS, harvested after 24 hours and RNA isolated for qPCR analyses. **a**. AR-V7 expression measured using qPCR in LNCaP, LN95 and 22RV1 cells. **b**. AR-V9 expression measured using qPCR in LNCaP, LN95 and 22RV1 cells.

#### Supplementary Figure 3

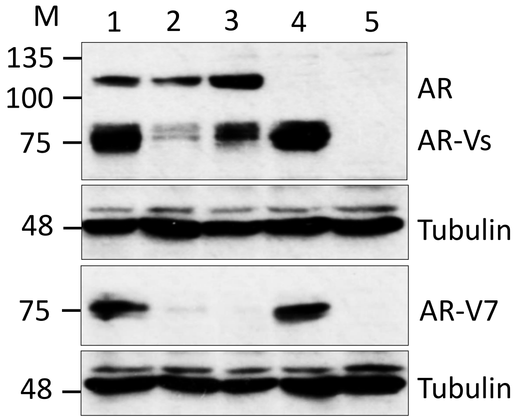


**Supplementary Figure 3. AR variant expression in 22RV1 cells.** Cells in CSS medium were treated with the indicated siRNAs for 72 hours, protein lysates prepared, separated by SDS gel electrophoresis and AR isoforms detected using AR441 (top) and AR-V7 antibodies (bottom). Lanes represent 22RV1 cells treated with 1. Control siRNA, 2. siRNA against AR-V7 II, 3. siRNA against AR-V7 III, 4. siRNA against AR (exon 7), 5. siRNA against all AR isoforms (exon 1).

#### Supplementary figure 4

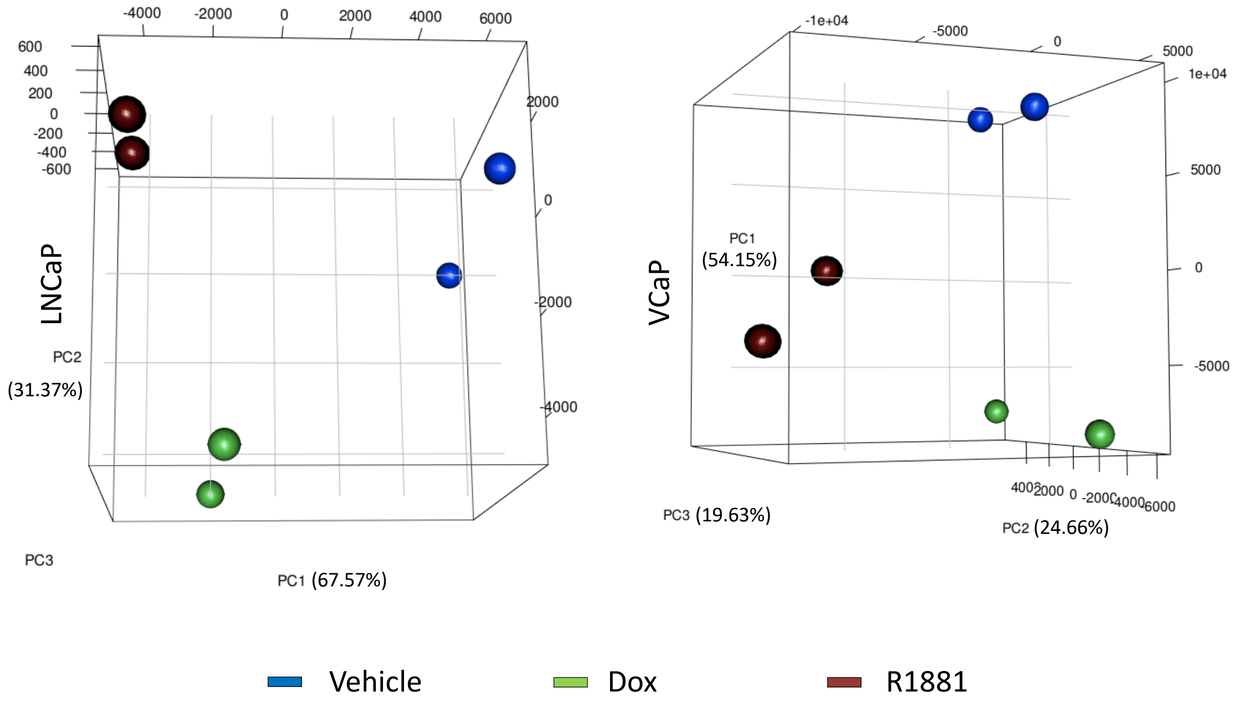


**Supplementary Figure 4. Principal component analysis for LNCaP and VCaP RNA-Seq datasets.** First three components are plotted, and they explained most of the variability. Vehicle treated, Dox treated and R1881 treated samples show a clear separation from each other. Replicates also behave similarly although there is some variation between samples.

**Supplementary figure 5**

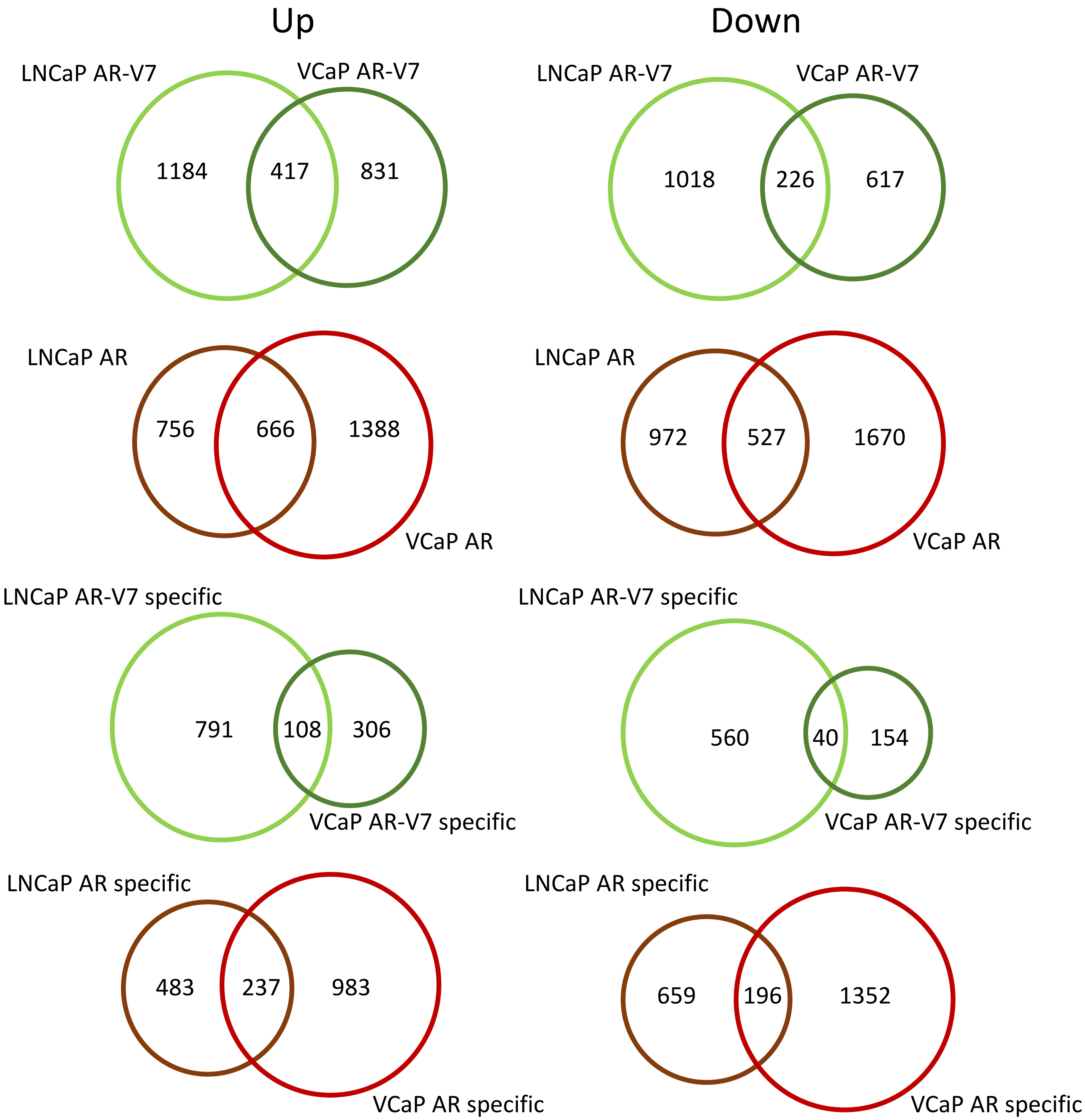


**Supplementary Figure 5. Comparison of AR isoform mediated gene expression in the LNCaP and VCaP models.** Venn diagrams were prepared to show the overlap between genes regulated by the AR isoforms in the two inducible models. Upper portion: Comparison of all genes up or down regulated by AR or AR-V7 at least 1.5 fold adjusted p <0.05. Lower portion: A comparison of genes specifically regulated by one isoform, but not the other. Venn diagrams are plotted as circles proportional to the numbers.

**Supplementary Figure 6**

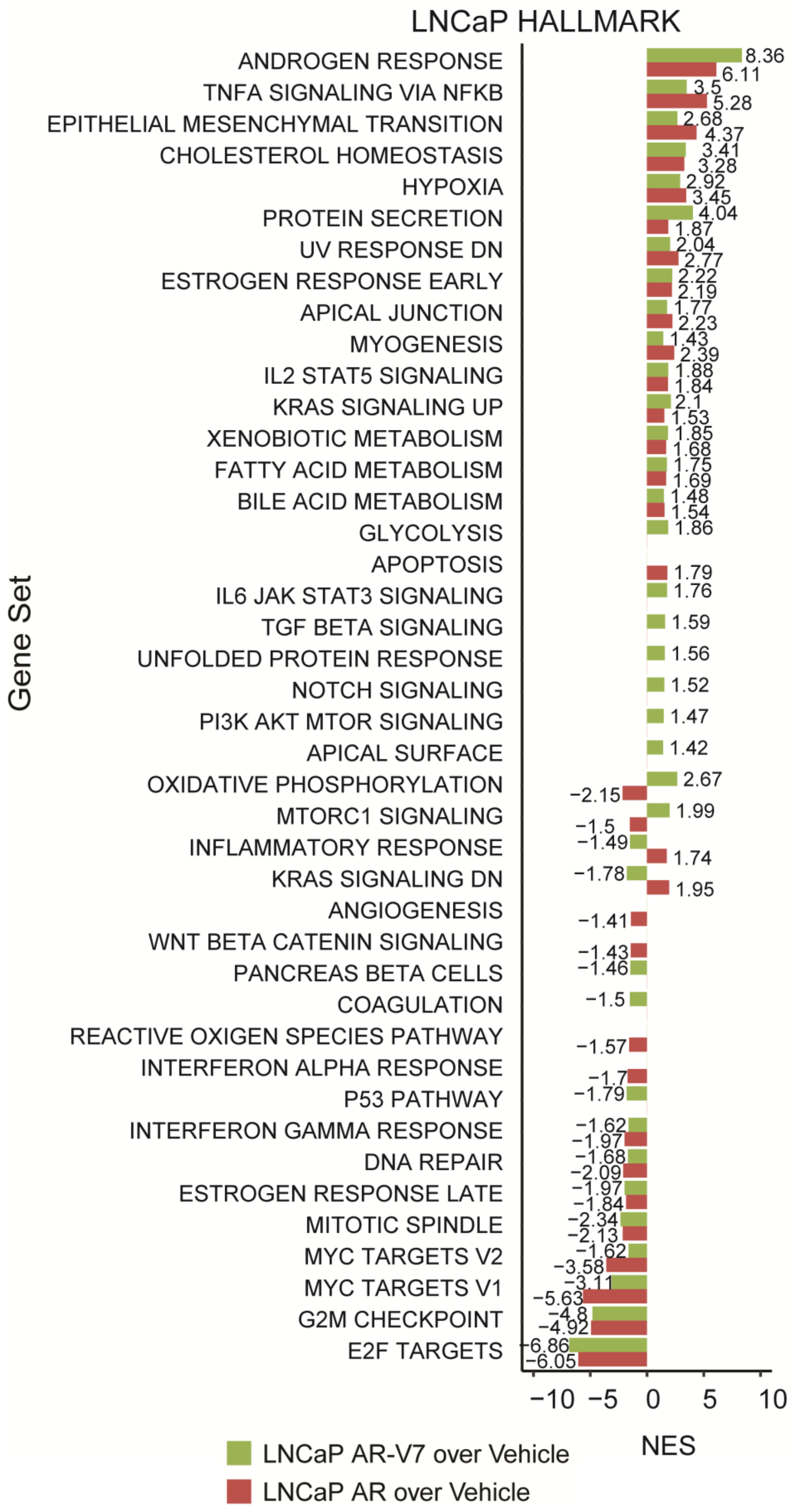


**Supplementary Figure 6. Hallmark pathways regulated in LNCaP AR-V7 cells.** All Hallmark pathways significantly regulated by AR isoforms in the LNCaP model. Rank is determined by AR-V7 value.

**Supplementary Figure 7**

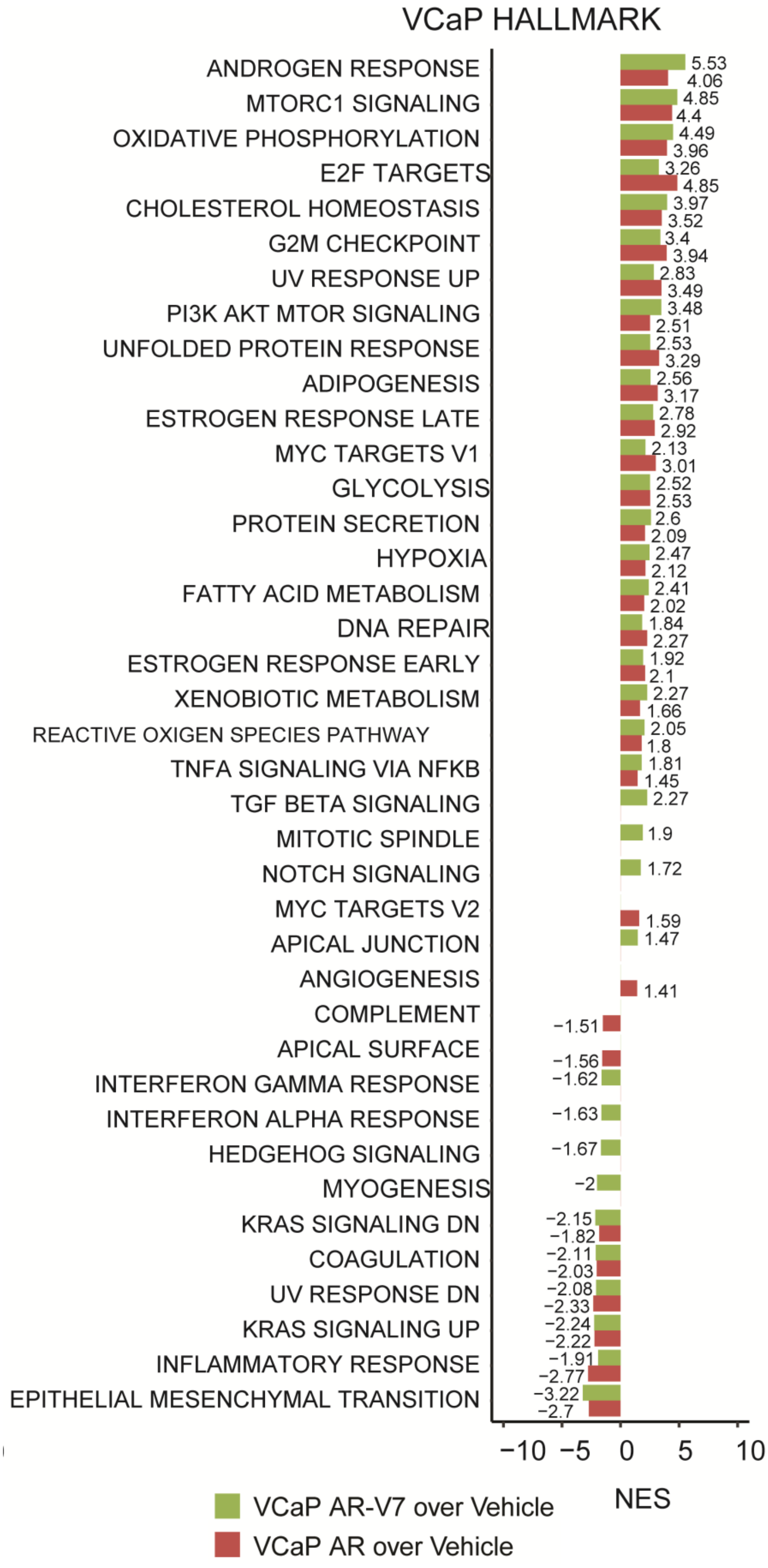


**Supplementary Figure 7. Hallmark pathways regulated in VCaP AR-V7 cells.** All Hallmark pathways significantly regulated by AR isoforms in the VCaP model. Rank is determined by AR-V7 value

**Supplementary figure 8**

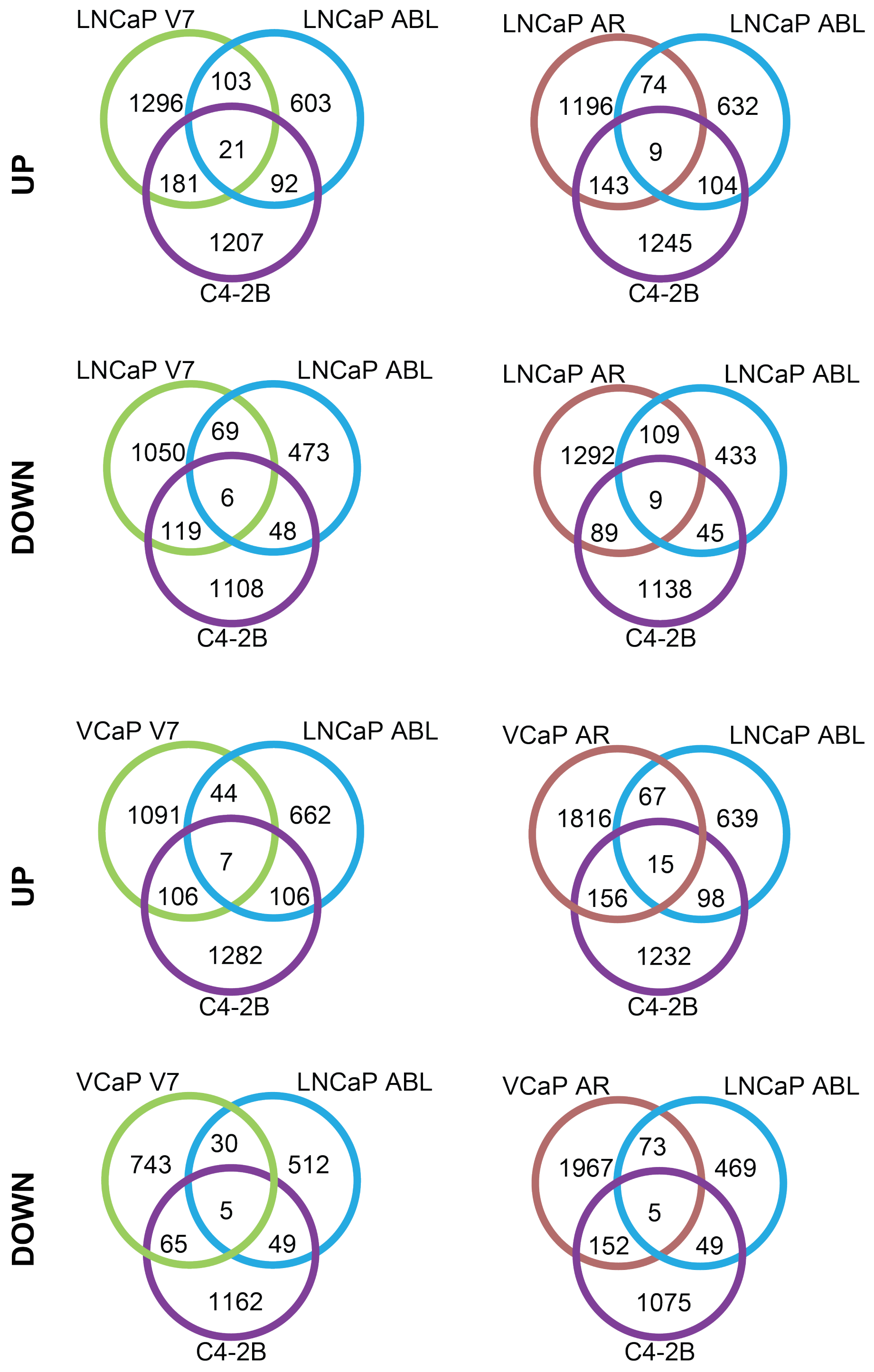


**Supplementary Figure 8. Comparison of Gene Expression in LNCaP AR-V7 and VCaP AR-V7 cells with LNCaP ABL and C4-2B cells.** All Genes up or down regulated by AR or AR-V7 in the two inducible models were compared with genes regulated by depletion of AR in the LNCaP ABL and C4-2B models. Only those genes with 1.5 fold regulation and $p<0.05$ were included. Left, AR-V7 comparisons; Right, AR comparisons. Venn diagrams are plotting symmetrical circles and not proportionate to numbers.

**Supplementary Figure 9**

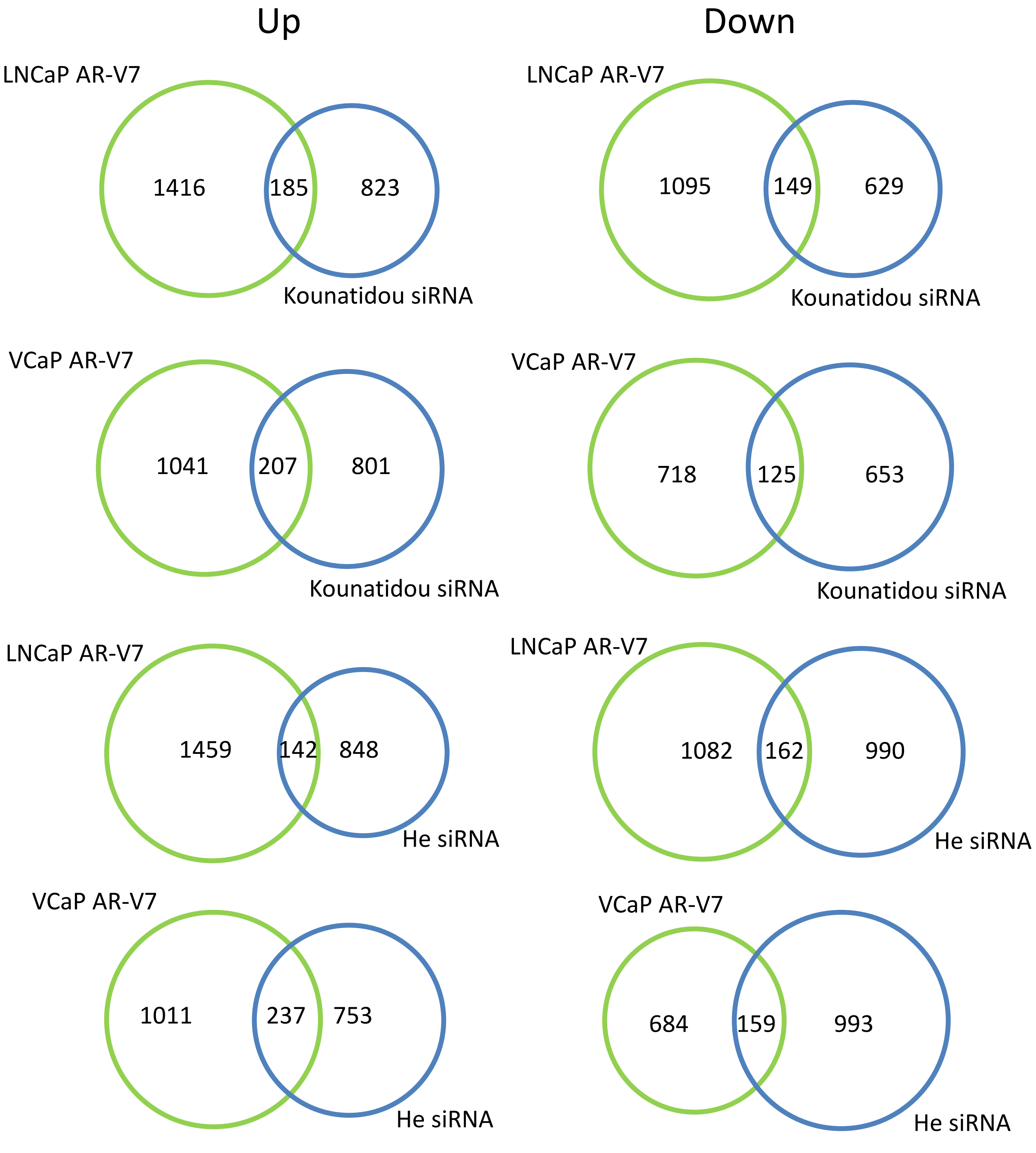


**Supplementary Figure 9. Comparison of genes regulated in the LNCaP AR-V7 and VCaP AR-V7 models with genes regulated in 22RV1 models.** Venn diagrams showing the overlaps between AR-V7 up or down-regulated genes from the engineered models with genes regulated by variants in two separate data sets in 22RV1 cells derived from depletion of variants using mixed siRNAs targeting variants(He) or by eliminating full length AR CRISPR followed by depletion of all AR variants using siRNA targeting exon 1 (Kounatidou). Venn diagrams are plotted as circles proportional to the numbers.

**Supplementary figure 10**

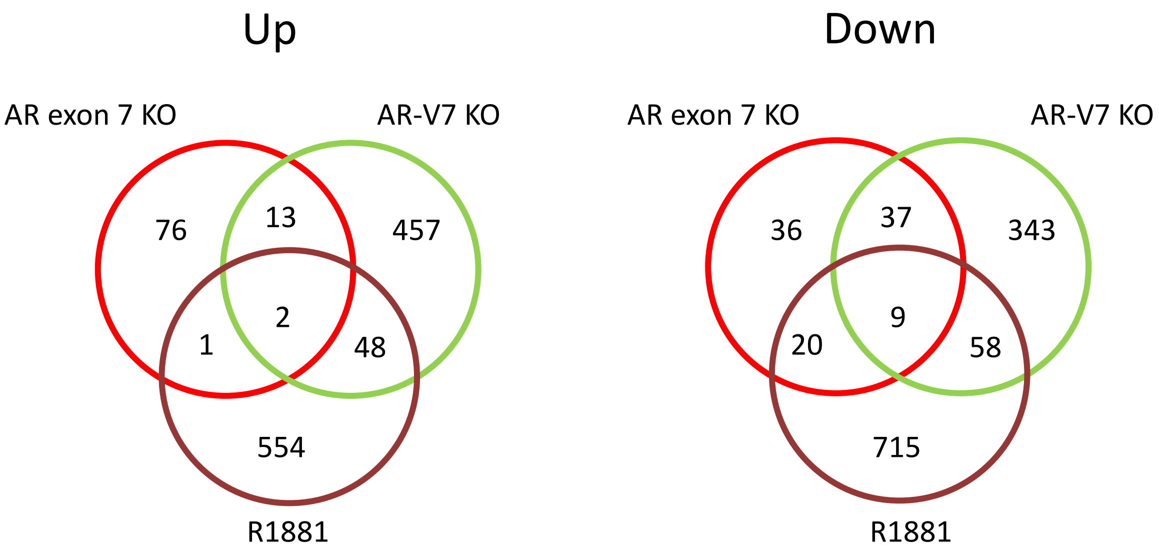


**Supplementary Figure 10. Comparison of genes regulated by AR-V7, AR, and R1881 in LN95 cells** RNA was isolated from LN-95 cells depleted of AR (si-exon 7), or AR-V7 (siCE3) or control siRNA. To identify R1881 dependent genes, cells were treated with vehicle or 10 nM R1881 and all samples were subjected to RNA-Seq. AR exon 7 KO (control si/exon 7si); AR-V7 KO (controlsi/CE3 si). R1881 (R1881/vehicle). Only those genes with 1.5 fold regulation and $p<0.05$ were included. Two biological replicates prepared in separate weeks were used for each treatment condition. Venn diagrams are plotting symmetrical circles and not proportionate to numbers.

**Supplementary figure 11**

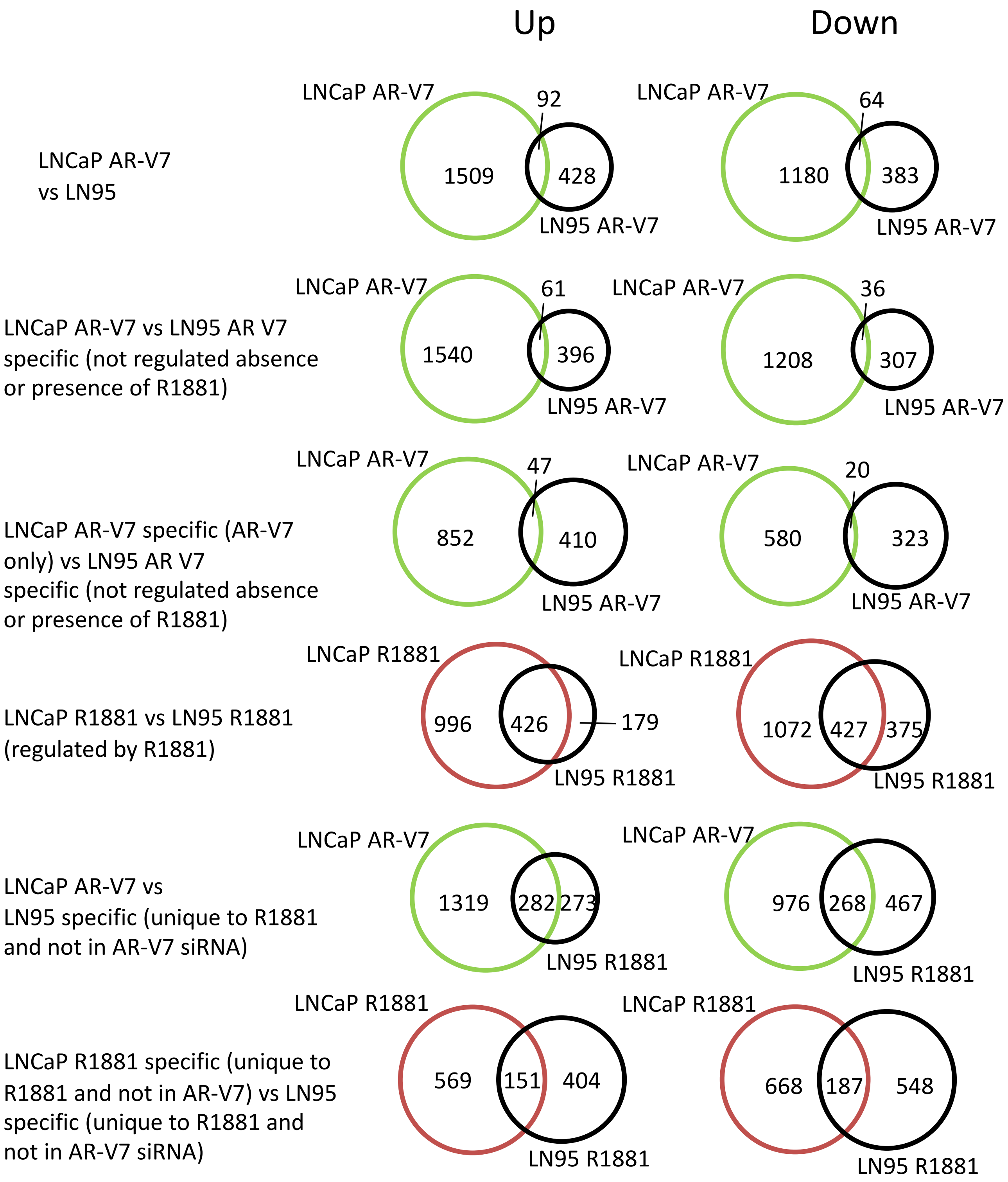


**Supplementary Figure 11. Comparison of genes regulated in the LN95 model and in the LNCaP AR-V7 model.** Genes from the sets described in SFig. 10 were compared to total AR-V7 regulated genes in both models or AR-V7 specific genes in both models. Total and specific R1881 regulated genes in both models also were compared. Finally, total AR-V7 regulated genes in the LNCaP model were compared with R1881 regulated genes in the LN95 model. Venn diagrams are plotted as circles proportional to the numbers.

**Supplementary Figure 12**

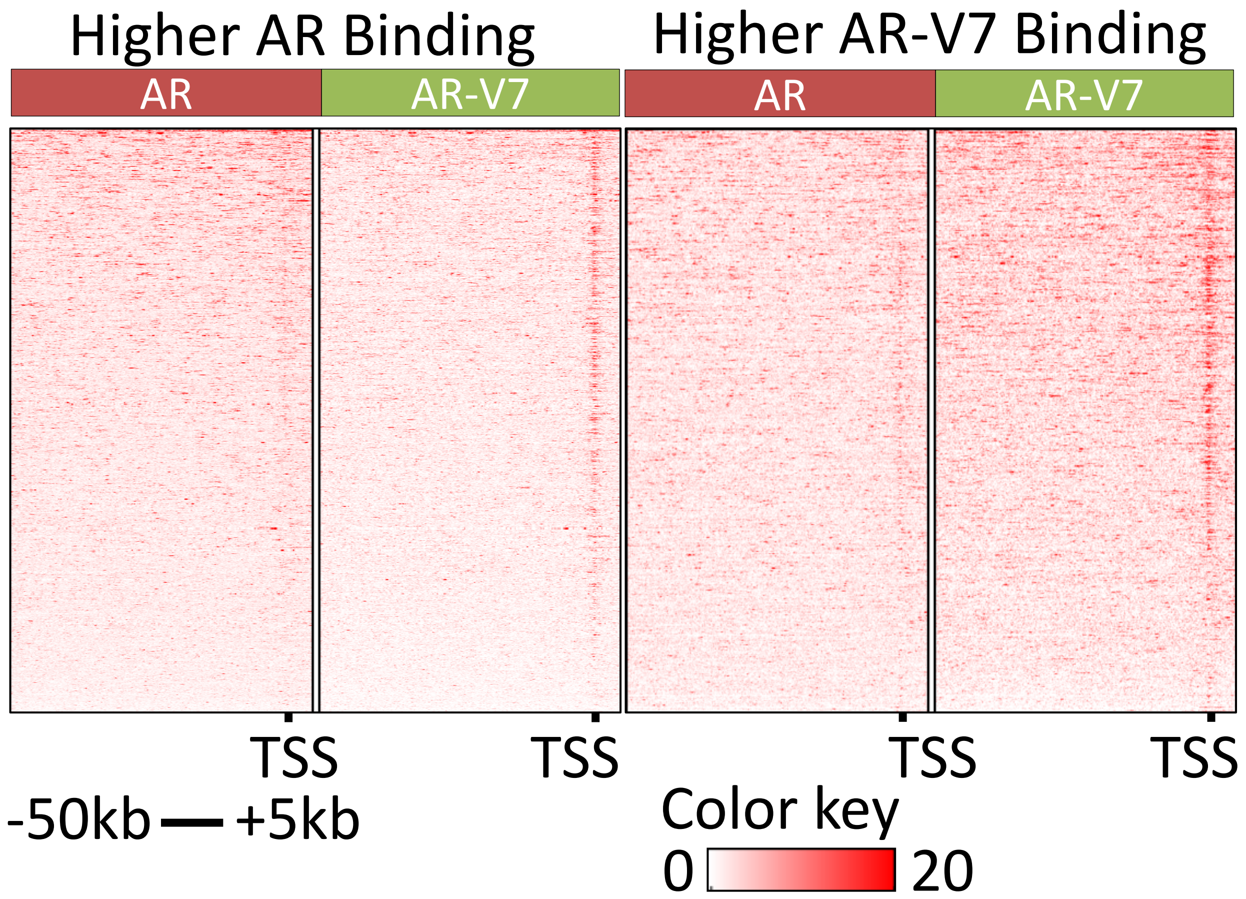


**Supplementary Figure 12. AR and AR-V7 Exhibit Differential Binding Patterns. Characterization of Transcription factor binding at Transcription Start Sites in the Human Genome. A**. Average ChIP-exo signal (normalized read counts) for AR and AR-V7 at TSS plotted as a profile within 50 kb upstream and 5 kb downstream. ChIP-exo signal > 4-fold is considered as higher binding.

### Supplementary Figure 13

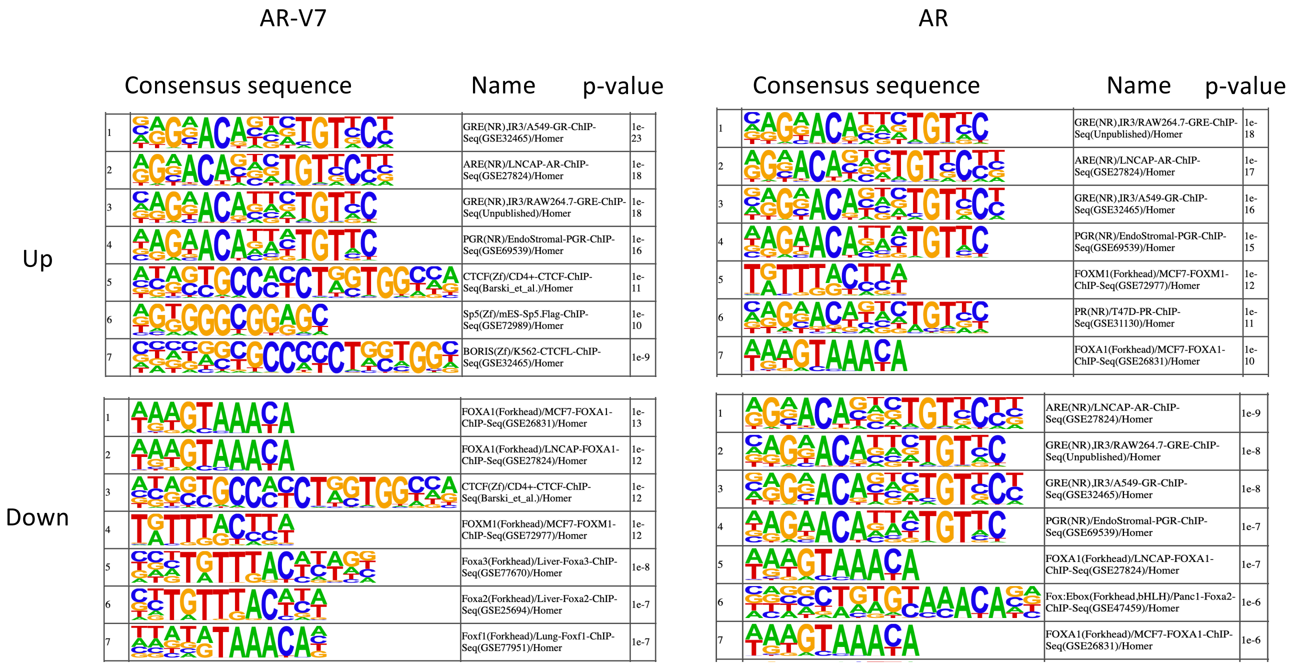


**Supplementary Figure 13. Consensus motifs enriched at AR and AR-V7 binding sites near TSS of DEGs.** Consensus motif sequences identified in a motif analysis using ARBs and ARVBs within 100bpof TSS of up and down DEGs. Homer findmotifsGenome.pl, -factor and other default parameters (target peaks vs background $p < 0.05$) were used.

**Supplementary Figure 14**

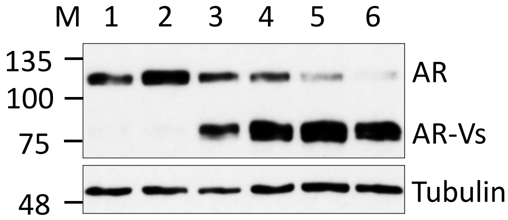


**Supplementary Figure 14. Induction of AR-V7 by Doxycycline.** LNCaP AR-V7 cells in CSS were treated with vehicle, 10 nM R1881, or the indicated doses of doxycycline for 24 hours, harvested, and AR isoforms detected by western blotting with AR 441 antibody and β Tubulin antibody. Note that the shorter exposure needed to portray the high levels of induction does not detect the low basal level of AR-V7 seen in the absence of Doxycyline. Lanes represent LNCaP AR-V7 cells treated with 1. Ethanol, 2. 10 nM R1881, 3-6. Doxycycline final concentration 3. Doxycycline 20 ng/mL, 4. Doxycycline 40 ng/mL, 5. Doxycycline 80 ng/mL, 6. Doxycycline 120 ng/mL.

**Supplementary Figure 15**

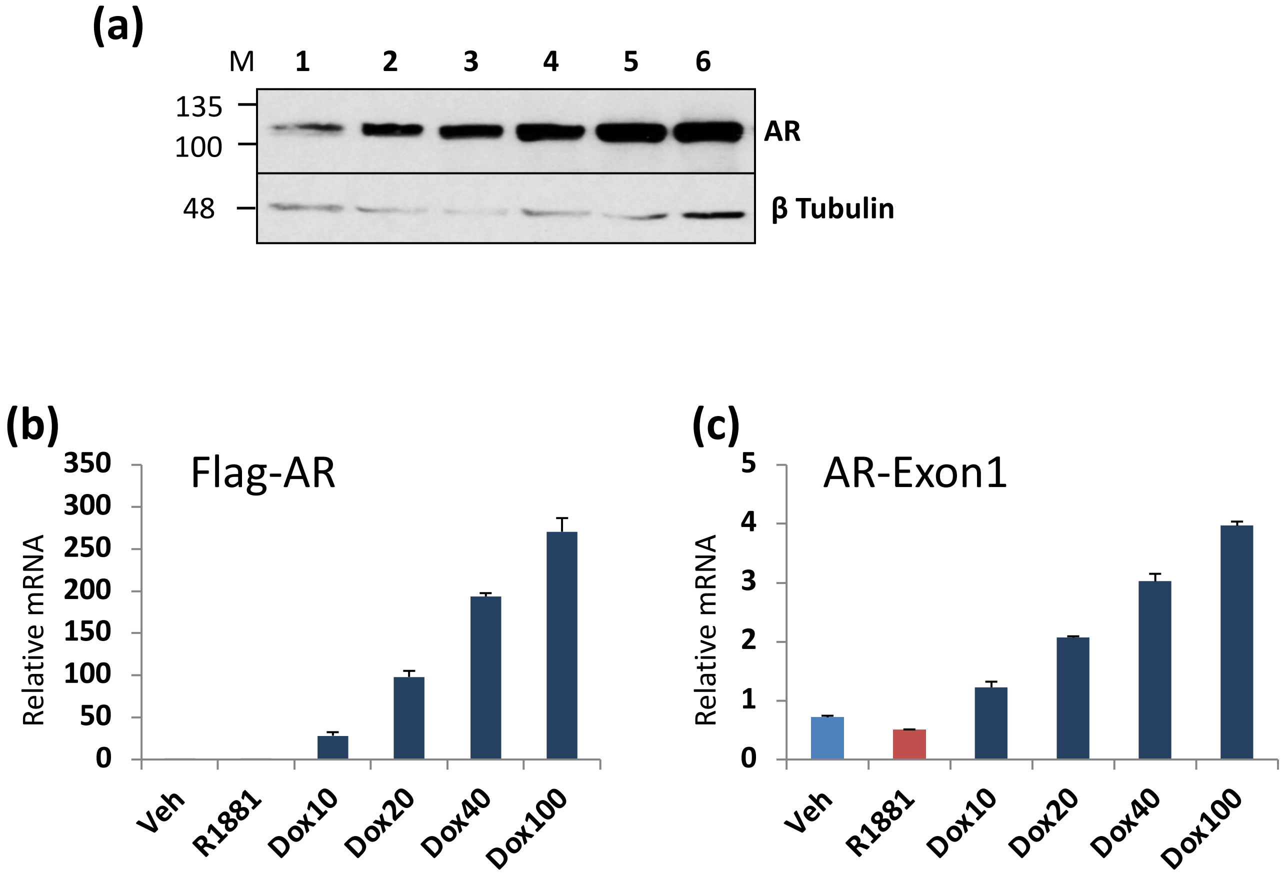


**Supplementary Figure 15. Doxycycline induced overexpression of AR.** The flag AR line in CSS was treated with vehicle (0.1% ethanol), 10 nM R1881 or the indicated doses of Doxycyclne for 24 hours, proteins isolated for western analyses, and RNA for mRNA expression. **a**. Western blot. Western blot using AR 441 antibody and β Tubulin antibody confirming increased expression of AR in response to different doses of Doxycycline. Lanes represent LNCaP AR cells treated with 1. Ethanol, 2. 10 nM R1881, 3-6. Doxycycline final concentration 3. Doxycycline 10 ng/mL, 4. Doxycycline 20 ng/mL, 5. Doxycycline 40 ng/mL, 6. Doxycycline 100 ng/mL. **b**. Samples prepared as described above were assayed using qPCR for the flag epitope. **c**. The same samples as in **b** were assayed by qPCR for AR-exon 1, a measure of total AR.

**Supplementary Figure 16**

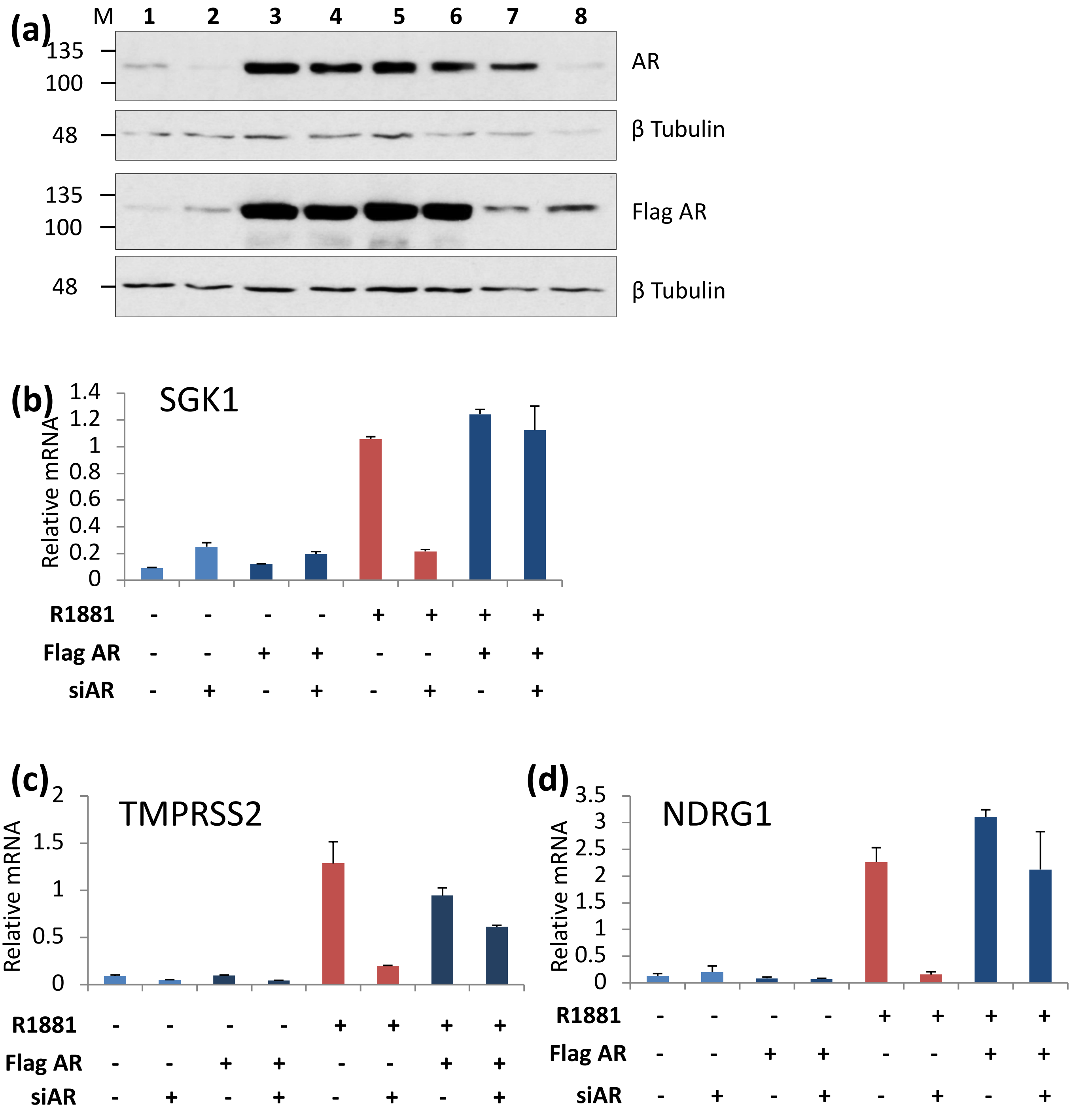


**Supplementary Figure 16. Confirmation of flag AR activity.** Flag AR cells in CSS were treated with si3'AR, or control siRNA for 48 hours and then treated with vehicle , R1881, or doxycycline (20 ng/ml) for 24 hours prior to harvest for analysis of protein and mRNA. **a**. Western blot using AR 441 antibody and β Tubulin (upper) and flag antibody and tubulin antibody (lower). Lanes represent LNCaP AR cells treated with 1. Vehicle, 2. Vehicle+si3'AR, 3. Dox, 4. Dox+si3'AR, 5. Dox+10 nM R1881, 6. Dox+10 nM R1881+si3'AR, 7. 10 nM R1881, 8. 10 nM R1881+si3'AR. **b-d**, qPCR analysis of **b**. SGK1, **c**, TMPRSS2 , and **d**, NDRG1 expression. Dox treatment was with 20ng/mL, si3'AR – represent cells treated with siControl.

#### Supplementary Figure 17

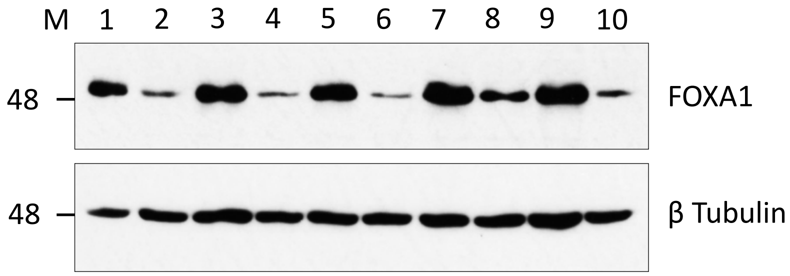


**Supplementary Figure 17. Depletion of FOXA1 in LNCaP AR-V7 and LNCaP AR cells.** Cells in CSS were treated with control siRNA or siFoXA1 for 24 hours, followed by treatment with vehicle, Doxycycline, or R1881 for another 24 hours.Western blot showing FOXA1 depletion using siRNA transfection in LNCaP AR-V7 (1 – 6) and LNCaP AR (7 – 10) cells 1. Vehicle, 2. Vehicle +siFOXA1, 3. 20 ng/mL final concentration Dox, 4. 20 ng/mL final concentration Doxy+siFOXA1, 5. 10 nM R1881, 6. 10 nM R1881+siFOXA1, 7. LNCaP AR Dox, 8. LNCaP AR Dox +siFOXA1, 9. LNCaP AR Dox+ 10 nM R1881, 10. LNCaP AR Dox +siFOXA1+ 10 nM R1881. Corresponding control cells that did not get siFOXA1 treatment were treated with siControl.

**Supplementary Figure 18**

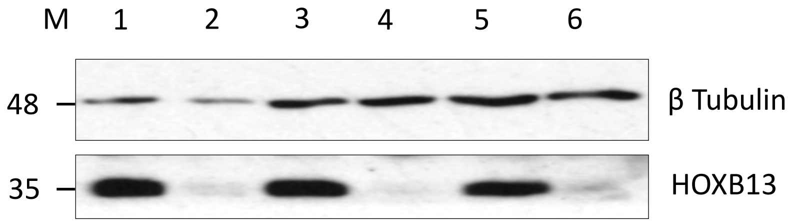


**Supplementary Figure 18. Depletion of HOXB13.** LNCaP cells in 10% CSS with treated with control or HOXB13 siRNA for 24 hours prior to addition of the indicated treatments for another 24 hours. HOXB13 was detected by western blotting.1. Ethanol, 2. Ethanol +siHOXB13, 3. 20ng/mL final concentration Dox, 4. 20ng/mL final concentration Dox+siHOXB13, 5. 10 nM R1881, 6. 10 nM R1881+siHOXB13. Corresponding control cells that didn't get siFOXA1 treatment were treated with siControl.

#### Supplementary Figure 19

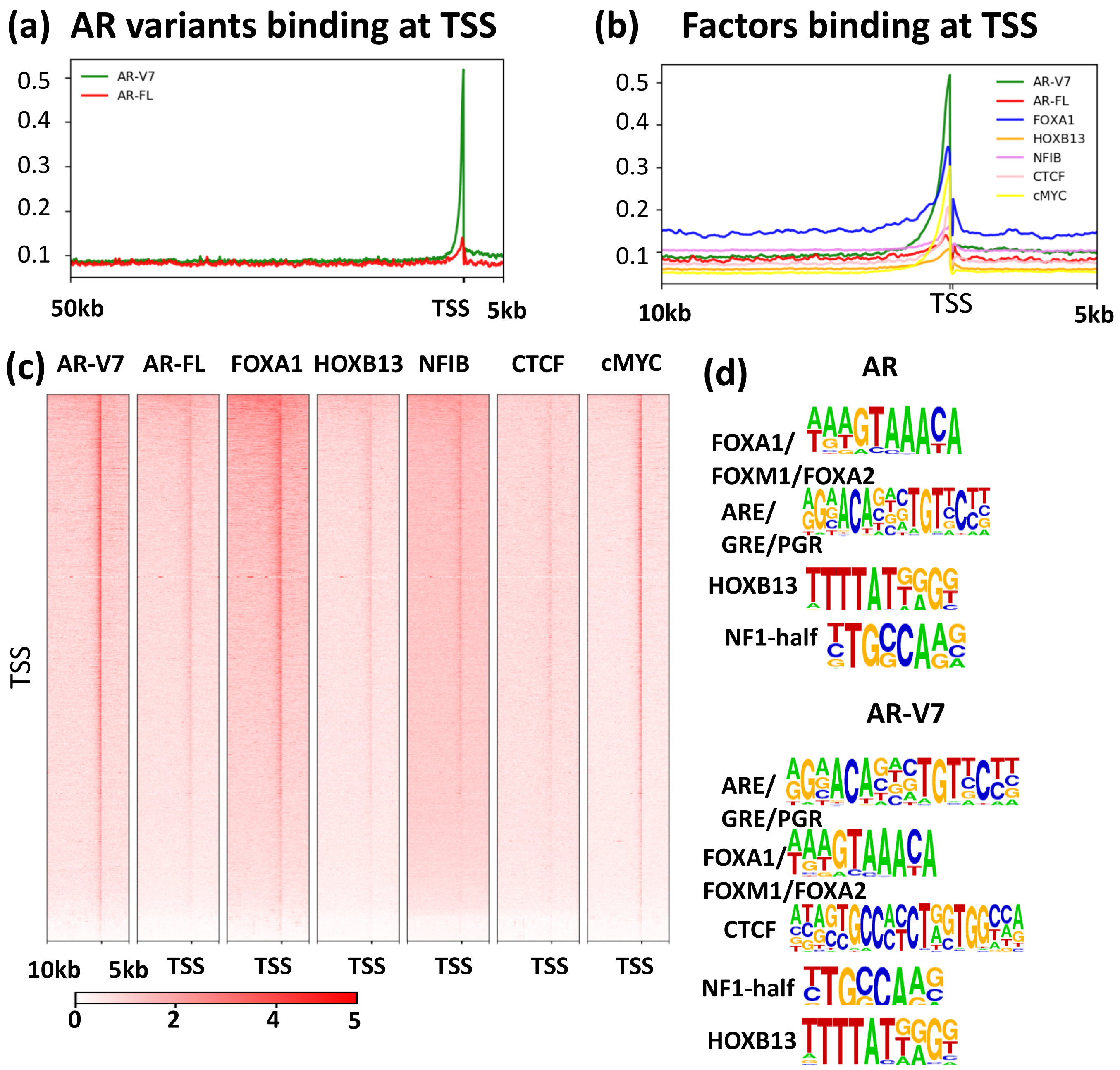


**Supplementary Figure 19. Transcription factor binding profile at Transcription Start Sites of DEGs. a**. Average ChIP-Exo signal for AR variants and **b.** ChIP signal for other factors at regulatory sites of DEGs, plotted as a profile within 1kb upstream and 1kb downstream. **c**. Heatmap visualizing average ChIP signal for AR variants and other factors at proximal promoter sites (ARBs and AR-V7 binding sites within 100bp to TSS) plotted within 1kb upstream to 1kb downstream of peak center. **d**. Consensus motif sequences identified in a motif analysis using Homer findmotifsGenome.pl, -factor and other default parameters (target peaks vs background $p < 0.05$) of AR and AR-V7 binding sites in Figure 3, panel A.

#### Supplementary Figure 20

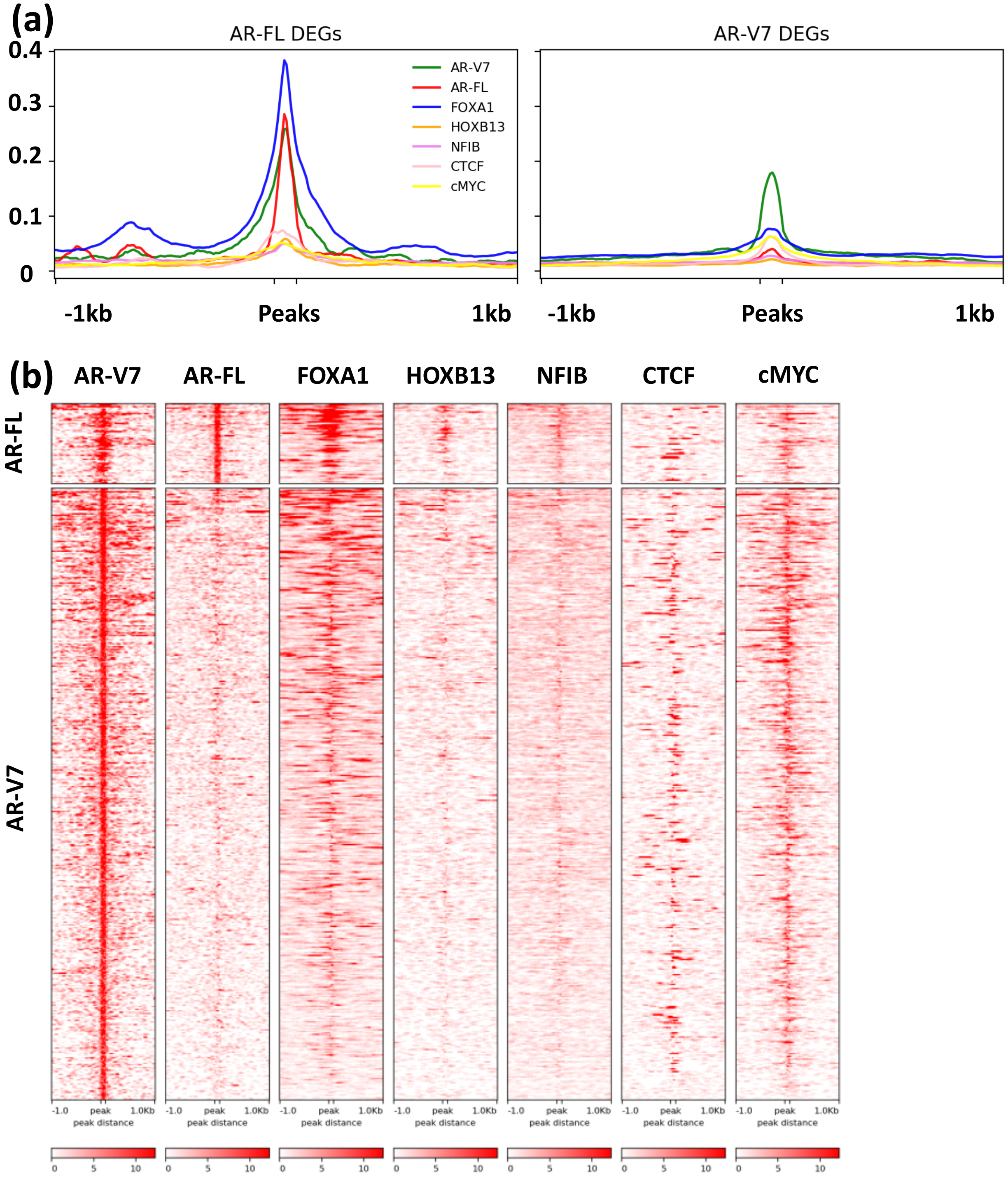


**Supplementary Figure 20.** Transcription factor binding profile at Transcription Start Sites of DEGs. **a**. Average ChIP signal for AR variants and other factors at regulatory sites of DEGs, plotted as a profile within 1 kb upstream and 1 kb downstream. **b**. Heatmap visualizing average ChIP signal for AR variants and other factors at proximal promoter sites (ARBs and AR-V7 binding sites within 100 bp to TSS) plotted within 1kb upstream to 1kb downstream of peak center.

**Supplementary Figure 21**

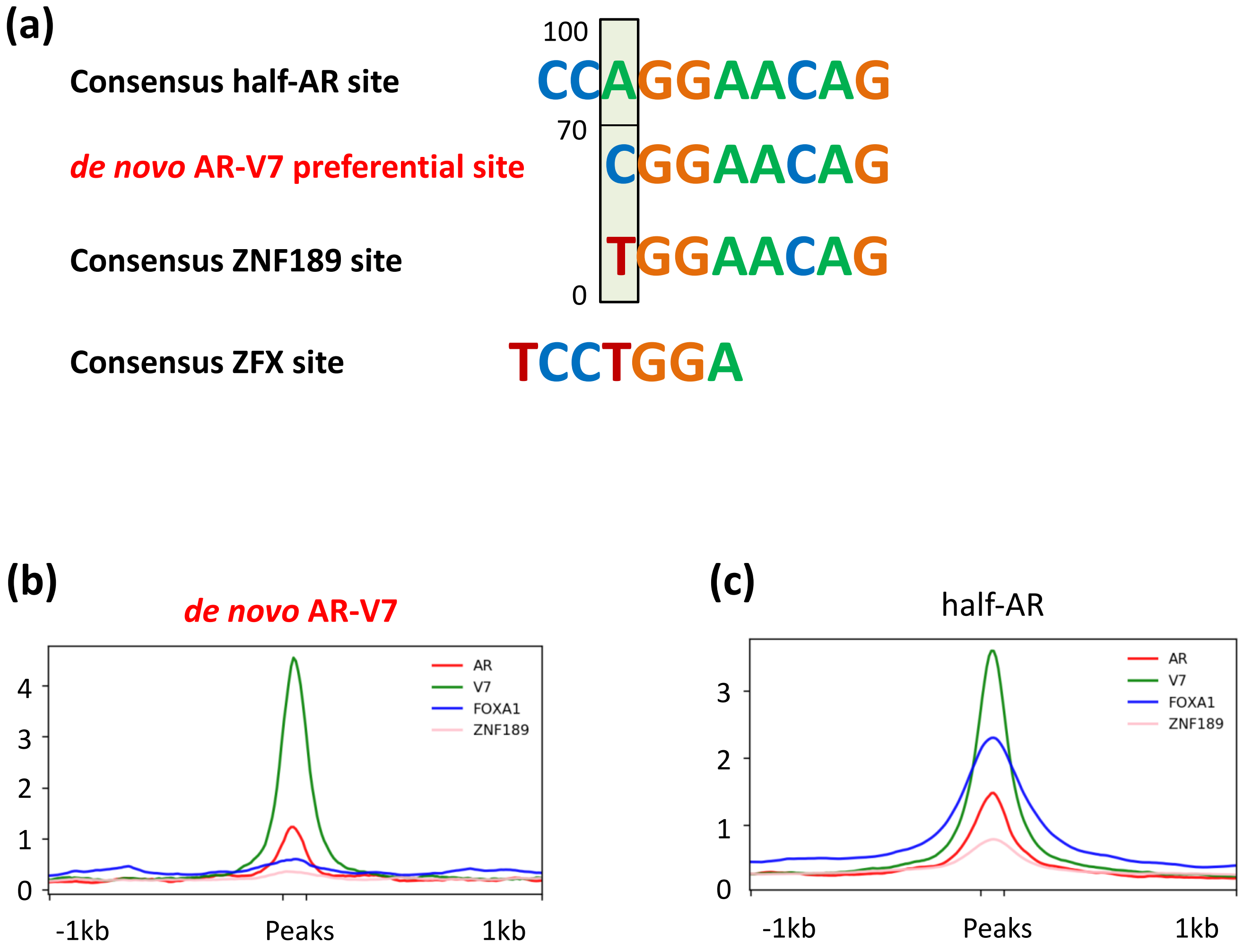


**Supplementary Figure 21.** AR-V7 *de novo* motifs enriched in a motif analysis within 20bp from peak center. **a**. Consensus half-AR motif, *de novo* AR-V7 motif identified, consensus ZNF189 motif and consensus ZFX motif showing sequence similarity (target peaks vs background $p < 0.05$). **b**. Average ChIP signal for AR variants and other factors at AR-V7 binding sites having *de novo* motif present (within 250bp from peak center). **c**. Average ChIP signal at AR-V7 binding sites with consensus half ARE motif (within 250bp from peak center) is plotted as a profile. b and c are plotted within 1 kb upstream to 1 kb downstream of peak center.

**STable1.** List of publicly available data sets used in this study.

| | ChIP-seq data | Data |
|---|---|---|
| 1 | **AR ChIP-exo (Data from this Lab)** | **GSE143906** |
| 2 | **AR-V7 ChIP-exo (Data from this Lab)** | **GSE143906** |
| 3 | AR ChIP-exo | GSE43785 |
| 4 | FOXA1 ChIP-exo | GSE43678 |
| 5 | HOXB13 ChIP-seq | GSE96652 |
| 6 | ZFX ChIP-seq | GSE94013 |
| 7 | NFIB ChIP-seq | PRJNA276666 |
| 8 | CTCF ChIP-seq | GSE38685 |
| 9 | cMYC ChIP-seq | GSE73995 |

| | RNA-seq data | |
|---|---|---|
| 1 | **LNCaP AR-V7 (Data from this Lab)** | **GSE143905** |
| 2 | **LN95 datasets (Data from this Lab)** | **GSE184676** |
| 3 | 22RV1 AR-V7 signature (He) | GSE80743 |
| 4 | 22RV1 AR-V7 signature (Cai) | GSE94013 |
| 5 | 22RV1 AR-V7 signature (Kounatidou) | GSE126306 |
| 6 | LNCaP ABL AR signature | GSE35126 |
| 7 | C4-2B AR signature | GSE40050 |
| 8 | FOXA1 KD | GSE27824 |

**STable 2.** Sequence information for Real-Time PCR Real-time PCR primer information for qPCR and ChIP-qPCR.

| Orientation | Gene | Sequence | Comments |
|---|---|---|---|
| Sense | 18S | ACCGCAGCTAGGAATAATGGA | qPCR |
| Antisense | 18S | GCCTCAGTTCCGAAAACCA | qPCR |
| Sense | PSA (KLK3) | ACCTGCACCCGGAGAGCT | qPCR |
| Antisense | PSA (KLK3) | TCACGGACAGGGTGAGGAAG | qPCR |
| Sense | ELL2 | GCTCAATTGCCTGGGATTTATAC | qPCR |
| Antisense | ELL2 | CTCTGCCTGGGTCATTCTTT | qPCR |
| Sense | SGK1 | CCTTGACGCTGGCTGTGA | qPCR |
| Antisense | SGK1 | TGACCCCGAGTTTACCGA | qPCR |
| Sense | TMPRSS2 | AGGATCGGTGTGTTCGCCTC | qPCR |
| Antisense | TMPRSS2 | CTCGTTCCAGTCGTCTTGGC | qPCR |
| Sense | NRDG1 | CATGAACCCCGGCAACC | qPCR |
| Antisense | NRDG1 | GCTCAATCTCCAGGTCGCG | qPCR |
| Sense | OLAH | GAGAGAGGAGACCAACCTAAGA | qPCR |
| Antisense | OLAH | CAGGGAAAGCAAATCAGCTTAAA | qPCR |
| Sense | ChIP -ve | AACCTCACTTTCATTGTTACTAGCCATA | ChIP qPCR |
| Antisense | ChIP -ve | CGCTCAAGGATGTCAGTAGCAT | ChIP qPCR |
| Sense | TRPM8 | CCCCTGGTGTGCATCTACATG | qPCR |
| Antisense | TRPM8 | CCCACCGTGTAGCCAAACAT | qPCR |
| Sense | PAK1IP1 | CCTCTTTGGGTTCGCTGTACAC | qPCR |
| Antisense | PAK1IP1 | TGAAGTCAGCCACAAGAGTCCAT | qPCR |
| Sense | ELOVL7 | TTTGCTGCAGGTGGTTTGGGAA | qPCR |
| Antisense | ELOVL7 | AAATACTTCTGGTAGGCTGGCCC | qPCR |
| Sense | TRPM8 Enh1 | AGCATCAGTGAAATTTGAGACA | ChIP qPCR |
| Antisense | TRPM8 Enh1 | GAGTTTGGGAAATGCTGAAGTG | ChIP qPCR |
| Sense | EDN2 | CTGCAGACGTGTTCCAGAC | qPCR |
| Antisense | EDN2 | AGGCTCTTGACTGTGGAAATG | qPCR |
| Sense | FKBP5 | GGATATACGCCAACATGTTCA | qPCR |
| Antisense | FKBP5 | CCATTGCTTTATTGGCCTCT | qPCR |
| Sense | NDRG1 | CATGAACCCCGGCAACC | qPCR |
| Antisense | NDRG1 | GCTCAATCTCCAGGTCGCG | qPCR |
| Sense | FOXA1 | GTGGCTCCAGGATGTTAGGA | qPCR |
| Antisense | FOXA1 | GAGTAGGCCTCCTGCGTGT | qPCR |
| Sense | NKX3-1 | GAGAGAGCCTTGGCCATATTC | ChIP qPCR |
| Antisense | NKX3-1 | GAGAAAGGACAACCAGGATGAG | ChIP qPCR |
| Sense | PSA Enh | GACCTACTCTGGAGGAACATATTG | ChIP qPCR |
| Antisense | PSA Enh | GGCTTGCTTACTGTCCTAGATAA | ChIP qPCR |
| Sense | BIRC3 | CGACAGGTGTCGCTTGAAA | ChIP qPCR |
| Antisense | BIRC3 | ACACACACACACACACACA | ChIP qPCR |
| Sense | FKBP5 | AGGTCAACTCTTGCTGGAAC | ChIP qPCR |
| Antisense | FKBP5 | ACTGGGAGTTGTTTGGTCTG | ChIP qPCR |

| | | | |
|---|---|---|---|
| Sense | PSA Prom | TGCATCCAGGGTGATCTAGTA | ChIP qPCR |
| Antisense | PSA Prom | GCTGGAGGCTGGACAAC | ChIP qPCR |
| Sense | ORM1 | GCATAATACAACCTGCTGTTCTG | qPCR |
| Antisense | ORM1 | TGTTCCCAACACAAAGAAATGG | qPCR |
| Sense | HOXB13 | TTGCCAGGGAGAACAGAAC | qPCR |
| Antisense | HOXB13 | CTGTACGGAATGCGTTTCTTG | qPCR |
| Sense | AR-V7 | GTCGTCTTCGGAAATGTTATGAA | qPCR |
| Antisense | AR-V7 | TGCCAACCCGGAATTTTTCT | qPCR |
| Sense | AR-V9 | AACAGAAGTACCTGTGCGCC | qPCR |
| Antisense | AR-V9 | TCAGGGTCTGGTCATTTTGA | qPCR |
| Antisense | AR-V9 | GCAAATGTCTCCAAAAAGCAGC | qPCR |
| Sense | AR Exon1 | GGATAGCTACTCCGGACATTACG | qPCR |
| Antisense | AR Exon1 | ATAGTCAATGGGGAAAAGATGGT | qPCR |
| Sense | AR Flag | ACAAAGACCATGACGGTGATTA | qPCR |
| Antisense | AR Flag | GAACAGATTCTGGAAAGCTCCT | qPCR |

**STable 3.** The list of differentially expressed genes (DEGs).

**LNCaP DEGs**

| Gene | log2FC R1881/Veh | log2FC Dox/Veh | Direction |
|---|---|---|---|
| NKX6-2 | -2.31 | 7.94 | Opposite |
| NR2F1 | -1.93 | 6.52 | Opposite |
| TNC | -0.62 | 6.38 | Opposite |
| ZNF727 | -1.60 | 6.18 | Opposite |
| EPHA3 | -0.84 | 5.62 | Opposite |
| KCNJ3 | -2.27 | 5.56 | Opposite |
| ZNF98 | -0.64 | 4.98 | Opposite |
| ALB | -1.18 | 4.23 | Opposite |
| SOHLH2 | -1.36 | 4.20 | Opposite |
| UBE2QL1 | -2.74 | 3.91 | Opposite |
| BIRC3 | -0.62 | 3.79 | Opposite |
| PKIA | -2.40 | 3.69 | Opposite |
| TFCP2L1 | -1.40 | 3.50 | Opposite |
| ZNF729 | -1.43 | 3.35 | Opposite |
| CYP7A1 | -3.17 | 3.34 | Opposite |
| CP | -2.14 | 3.32 | Opposite |
| ATOH8 | -1.06 | 3.31 | Opposite |
| ST6GALNAC2 | -0.61 | 3.26 | Opposite |
| ZNF99 | -1.66 | 3.19 | Opposite |
| KRT80 | -1.63 | 2.96 | Opposite |
| LUM | -1.73 | 2.93 | Opposite |
| KCNQ5 | -1.74 | 2.92 | Opposite |
| COL27A1 | -2.02 | 2.90 | Opposite |
| INSL5 | -1.39 | 2.86 | Opposite |
| UGT8 | -1.05 | 2.84 | Opposite |
| ADGRD1 | -0.77 | 2.72 | Opposite |
| OVOS2 | -1.38 | 2.70 | Opposite |
| EPAS1 | -0.76 | 2.64 | Opposite |
| EDN2 | -2.04 | 2.51 | Opposite |
| GCG | -2.67 | 2.49 | Opposite |
| ZNF676 | -0.63 | 2.46 | Opposite |
| KNG1 | -1.82 | 2.38 | Opposite |
| CCDC74A | -0.70 | 2.31 | Opposite |
| ARHGAP23 | -0.80 | 2.29 | Opposite |
| GSTA1 | -0.91 | 2.26 | Opposite |
| PLPPR1 | -3.95 | 2.25 | Opposite |
| NSG1 | -0.68 | 2.16 | Opposite |
| P3H2 | -1.05 | 2.12 | Opposite |
| ZFHX4 | -0.66 | 2.11 | Opposite |

**VCaP DEGs**

| Gene | log2FC R1881/Veh | log2FC Dox/Veh | Direction |
|---|---|---|---|
| HSD17B2 | 3.32 | -3.64 | Opposite |
| VWF | 0.72 | -2.15 | Opposite |
| SV2C | 1.43 | -1.92 | Opposite |
| PALMD | 0.94 | -1.60 | Opposite |
| TEAD2 | 2.73 | -1.37 | Opposite |
| LDLRAD4 | 0.68 | -1.35 | Opposite |
| PDE4B | 0.72 | -1.28 | Opposite |
| ALPK3 | 1.15 | -1.19 | Opposite |
| ZIC3 | 0.72 | -1.15 | Opposite |
| RPH3A | 3.44 | -1.14 | Opposite |
| SLC22A4 | 1.21 | -1.10 | Opposite |
| ALDH1A1 | 0.70 | -1.07 | Opposite |
| TMC7 | 0.61 | -1.06 | Opposite |
| UGT2B4 | 0.59 | -1.01 | Opposite |
| FPR3 | 0.93 | -1.00 | Opposite |
| GSDMC | 3.47 | -0.98 | Opposite |
| GNB4 | 1.27 | -0.96 | Opposite |
| ANXA8L1 | 0.98 | -0.94 | Opposite |
| CACNA2D1 | 1.00 | -0.94 | Opposite |
| TRPM8 | 1.99 | -0.93 | Opposite |
| GLB1L3 | 1.63 | -0.91 | Opposite |
| PIK3AP1 | 1.34 | -0.89 | Opposite |
| TFF1 | 0.69 | -0.87 | Opposite |
| KCNK6 | 1.00 | -0.87 | Opposite |
| RGS4 | 0.87 | -0.87 | Opposite |
| ROPN1B | 1.26 | -0.86 | Opposite |
| BOLA2B | 3.48 | -0.84 | Opposite |
| SLIT3 | 0.62 | -0.81 | Opposite |
| LGALS3 | 1.55 | -0.78 | Opposite |
| CHST8 | 0.74 | -0.78 | Opposite |
| GDF15 | 1.09 | -0.76 | Opposite |
| CNTD2 | 0.82 | -0.75 | Opposite |
| CDH13 | 1.21 | -0.74 | Opposite |
| TP53INP2 | 1.05 | -0.74 | Opposite |
| ADH1C | 1.80 | -0.71 | Opposite |
| PDE2A | 1.54 | -0.70 | Opposite |
| IGFL1 | 1.57 | -0.69 | Opposite |
| PDLIM3 | 0.66 | -0.69 | Opposite |
| GRHL1 | 0.58 | -0.66 | Opposite |

| | | | | | | | |
|---|---|---|---|---|---|---|---|
| SDC1 | -1.20 | 2.09 | Opposite | MYOF | 3.73 | -0.64 | Opposite |
| ASPH | -0.75 | 1.96 | Opposite | SASH1 | 1.17 | -0.64 | Opposite |
| GMPR | -1.23 | 1.95 | Opposite | CCDC53 | 0.74 | -0.63 | Opposite |
| XAGE1B | -1.71 | 1.95 | Opposite | PLCB1 | 0.69 | -0.63 | Opposite |
| PDE4D | -1.34 | 1.94 | Opposite | MAK | 1.99 | -0.63 | Opposite |
| TENM2 | -1.30 | 1.93 | Opposite | CYP1A1 | 0.71 | -0.63 | Opposite |
| ZDHHC15 | -1.02 | 1.91 | Opposite | VSIG10L | 1.89 | -0.62 | Opposite |
| GPR176 | -0.74 | 1.87 | Opposite | RNF152 | 1.25 | -0.62 | Opposite |
| C5orf46 | -1.46 | 1.82 | Opposite | SMPD3 | 1.43 | -0.62 | Opposite |
| ATP2B1 | -0.99 | 1.79 | Opposite | IQGAP2 | 0.72 | -0.61 | Opposite |
| BRDT | -1.75 | 1.77 | Opposite | TG | 2.30 | -0.61 | Opposite |
| KCNU1 | -3.87 | 1.72 | Opposite | CRABP2 | 2.29 | -0.61 | Opposite |
| USP2 | -0.96 | 1.69 | Opposite | ASB4 | 1.34 | -0.61 | Opposite |
| GRIK2 | -1.73 | 1.61 | Opposite | ZDHHC1 | 0.58 | -0.60 | Opposite |
| KIF17 | -1.40 | 1.61 | Opposite | SLC6A19 | 1.05 | -0.59 | Opposite |
| ADAMTS17 | -0.70 | 1.61 | Opposite | MESP2 | 1.14 | -0.58 | Opposite |
| GSTM4 | -0.71 | 1.59 | Opposite | ARID3A | 0.70 | -0.57 | AR-FL |
| NPTX2 | -0.98 | 1.54 | Opposite | SEC14L2 | 1.44 | -0.57 | AR-FL |
| MT2A | -0.66 | 1.54 | Opposite | UGT2B11 | 1.26 | -0.57 | AR-FL |
| RGMA | -0.78 | 1.49 | Opposite | ANKH | 0.73 | -0.57 | AR-FL |
| FSTL4 | -1.45 | 1.46 | Opposite | CYP4B1 | 1.38 | -0.57 | AR-FL |
| MICALCL | -0.84 | 1.44 | Opposite | FAM212B | 0.66 | -0.55 | AR-FL |
| ZNF257 | -1.05 | 1.40 | Opposite | GPR158 | 0.85 | -0.55 | AR-FL |
| MOV10L1 | -1.52 | 1.37 | Opposite | CA5B | 0.99 | -0.54 | AR-FL |
| REEP2 | -1.08 | 1.36 | Opposite | SCGB2A1 | 0.89 | -0.53 | AR-FL |
| PITPNM3 | -0.68 | 1.35 | Opposite | CREB3L1 | 2.38 | -0.53 | AR-FL |
| SDC2 | -1.11 | 1.34 | Opposite | SUSD2 | 0.73 | -0.51 | AR-FL |
| CAMK2D | -0.67 | 1.33 | Opposite | SHROOM1 | 0.68 | -0.50 | AR-FL |
| FCHO1 | -1.33 | 1.31 | Opposite | TRIM16 | 0.90 | -0.49 | AR-FL |
| OASL | -1.00 | 1.31 | Opposite | SPON2 | 0.87 | -0.49 | AR-FL |
| ST3GAL5 | -1.07 | 1.28 | Opposite | FAM69A | 1.09 | -0.49 | AR-FL |
| TNS1 | -1.56 | 1.24 | Opposite | DSC2 | 0.70 | -0.49 | AR-FL |
| VWDE | -0.86 | 1.22 | Opposite | LGR5 | 1.38 | -0.49 | AR-FL |
| COL9A3 | -1.02 | 1.21 | Opposite | LRCH1 | 1.12 | -0.49 | AR-FL |
| PARP8 | -1.09 | 1.18 | Opposite | LCK | 1.57 | -0.48 | AR-FL |
| CHAC2 | -0.66 | 1.17 | Opposite | EHBP1 | 0.70 | -0.48 | AR-FL |
| NOTCH2NL | -0.59 | 1.16 | Opposite | SAT1 | 1.24 | -0.45 | AR-FL |
| LAMC3 | -0.71 | 1.14 | Opposite | ELOVL7 | 1.12 | -0.45 | AR-FL |
| CES4A | -0.66 | 1.13 | Opposite | BEND4 | 0.60 | -0.45 | AR-FL |
| PRC1 | -0.62 | 1.13 | Opposite | VASH2 | 0.60 | -0.43 | AR-FL |
| IER5L | -1.17 | 1.02 | Opposite | CHRM1 | 1.16 | -0.43 | AR-FL |
| FAM49A | -0.93 | 1.01 | Opposite | RAB23 | 0.62 | -0.42 | AR-FL |
| BEX1 | -1.35 | 1.01 | Opposite | SOSTDC1 | 0.87 | -0.42 | AR-FL |
| DEGS2 | -1.44 | 1.00 | Opposite | CCDC71L | 0.96 | -0.42 | AR-FL |
| PRIMA1 | -1.96 | 1.00 | Opposite | ENDOD1 | 1.96 | -0.41 | AR-FL |
| PDGFRB | -0.78 | 1.00 | Opposite | ACER2 | 1.30 | -0.41 | AR-FL |

| | | | | | | | |
|---|---|---|---|---|---|---|---|
| MT1X | -0.74 | 0.98 | Opposite | ANXA2 | 1.48 | -0.41 | AR-FL |
| ZNF492 | -1.02 | 0.96 | Opposite | SERPINE2 | 0.62 | -0.41 | AR-FL |
| PGM2L1 | -1.54 | 0.95 | Opposite | SLC22A23 | 0.62 | -0.41 | AR-FL |
| C16orf89 | -2.30 | 0.95 | Opposite | SNX25 | 0.80 | -0.41 | AR-FL |
| GPR161 | -0.67 | 0.92 | Opposite | EMILIN1 | 0.60 | -0.41 | AR-FL |
| CCDC109B | -0.63 | 0.92 | Opposite | ANKRD37 | 2.57 | -0.40 | AR-FL |
| HAS3 | -1.25 | 0.91 | Opposite | ATP8A2 | 0.70 | -0.39 | AR-FL |
| SPATA6 | -0.95 | 0.91 | Opposite | MAP3K15 | 0.71 | -0.39 | AR-FL |
| CCDC74B | -0.81 | 0.90 | Opposite | RARB | 0.88 | -0.38 | AR-FL |
| EPB41L2 | -0.69 | 0.89 | Opposite | GPNMB | 0.66 | -0.37 | AR-FL |
| NEU4 | -1.01 | 0.89 | Opposite | RASSF3 | 0.61 | -0.37 | AR-FL |
| LPO | -1.82 | 0.89 | Opposite | POPDC3 | 1.10 | -0.37 | AR-FL |
| MSX2 | -0.88 | 0.88 | Opposite | SLC16A6 | 1.20 | -0.36 | AR-FL |
| TMEM59L | -1.48 | 0.88 | Opposite | ZNF792 | 0.60 | -0.35 | AR-FL |
| CCNO | -2.41 | 0.86 | Opposite | SH3RF3 | 0.82 | -0.35 | AR-FL |
| CEL | -1.09 | 0.83 | Opposite | HCAR1 | 0.66 | -0.35 | AR-FL |
| SDC4 | -1.67 | 0.83 | Opposite | TIGD2 | 0.67 | -0.34 | AR-FL |
| TMPRSS9 | -0.73 | 0.82 | Opposite | F3 | 1.22 | -0.34 | AR-FL |
| DDIT4 | -0.90 | 0.81 | Opposite | GREM2 | 1.58 | -0.34 | AR-FL |
| CATSPERB | -0.69 | 0.79 | Opposite | ZNF827 | 0.75 | -0.33 | AR-FL |
| TRPC1 | -1.56 | 0.78 | Opposite | GNA12 | 0.61 | -0.33 | AR-FL |
| C18orf54 | -0.86 | 0.78 | Opposite | DIRAS2 | 0.70 | -0.33 | AR-FL |
| LDOC1 | -0.72 | 0.77 | Opposite | SKIL | 0.81 | -0.32 | AR-FL |
| CACNA1G | -2.99 | 0.77 | Opposite | PHLDB3 | 1.11 | -0.32 | AR-FL |
| TIGD3 | -0.87 | 0.75 | Opposite | INSM1 | 0.99 | -0.32 | AR-FL |
| ACSF2 | -1.20 | 0.75 | Opposite | SHISA8 | 1.52 | -0.32 | AR-FL |
| SPAG8 | -0.78 | 0.75 | Opposite | ITPKB | 1.06 | -0.31 | AR-FL |
| FBXL2 | -0.73 | 0.72 | Opposite | NOD2 | 1.78 | -0.31 | AR-FL |
| CCDC155 | -0.97 | 0.71 | Opposite | SMN1 | 1.61 | -0.31 | AR-FL |
| FEZ1 | -0.69 | 0.70 | Opposite | AFAP1 | 0.79 | -0.31 | AR-FL |
| TRIM29 | -1.88 | 0.70 | Opposite | RAPGEF5 | 0.67 | -0.30 | AR-FL |
| KLRG2 | -1.33 | 0.70 | Opposite | ZNF703 | 0.71 | -0.30 | AR-FL |
| GPER1 | -0.69 | 0.69 | Opposite | FAM3D | 1.98 | -0.29 | AR-FL |
| TLR3 | -0.63 | 0.69 | Opposite | DDR2 | 0.86 | -0.29 | AR-FL |
| ZNF594 | -0.61 | 0.69 | Opposite | GMDS | 0.84 | -0.28 | AR-FL |
| CFAP69 | -0.63 | 0.69 | Opposite | DEGS1 | 0.60 | -0.28 | AR-FL |
| SLC12A4 | -0.71 | 0.69 | Opposite | SLC2A10 | 1.08 | -0.27 | AR-FL |
| MECOM | -3.55 | 0.68 | Opposite | SYTL1 | 0.75 | -0.27 | AR-FL |
| ASZ1 | -4.49 | 0.68 | Opposite | ADRB1 | 0.66 | -0.27 | AR-FL |
| RARRES3 | -1.14 | 0.67 | Opposite | OTUD7B | 1.17 | -0.27 | AR-FL |
| LEKR1 | -0.75 | 0.67 | Opposite | ELOVL2 | 1.16 | -0.26 | AR-FL |
| VEPH1 | -1.09 | 0.67 | Opposite | SLC10A4 | 0.92 | -0.26 | AR-FL |
| ADAM23 | -1.89 | 0.64 | Opposite | ETV6 | 0.74 | -0.25 | AR-FL |
| IFIT1 | -0.83 | 0.64 | Opposite | LAMC2 | 1.15 | -0.25 | AR-FL |
| IFI27L2 | -2.62 | 0.63 | Opposite | MME | 0.83 | -0.25 | AR-FL |
| TRIM54 | -1.36 | 0.62 | Opposite | TBC1D16 | 0.65 | -0.25 | AR-FL |

| | | | | | | | |
|---|---|---|---|---|---|---|---|
| FOSB | -0.60 | 0.60 | Opposite | SERPINA11 | 0.64 | -0.25 | AR-FL |
| SMC6 | -1.06 | 0.60 | Opposite | NBL1 | 1.27 | -0.25 | AR-FL |
| PPFIA4 | -0.80 | 0.59 | Opposite | TM4SF1 | 0.70 | -0.24 | AR-FL |
| NHLRC4 | -0.82 | 0.59 | Opposite | MTCL1 | 1.31 | -0.24 | AR-FL |
| SPOCK2 | -0.74 | 0.58 | Opposite | BAIAP2L1 | 0.60 | -0.24 | AR-FL |
| ZBBX | -0.79 | 0.58 | AR-FL | B3GNT5 | 0.66 | -0.23 | AR-FL |
| SMIM3 | -1.95 | 0.57 | AR-FL | CLDN8 | 1.81 | -0.23 | AR-FL |
| MAGED4B | -0.67 | 0.57 | AR-FL | SEL1L3 | 0.68 | -0.23 | AR-FL |
| MYO15A | -0.94 | 0.57 | AR-FL | FMOD | 1.31 | -0.23 | AR-FL |
| CNBD1 | -2.28 | 0.57 | AR-FL | CCDC160 | 1.56 | -0.22 | AR-FL |
| MET | -2.75 | 0.56 | AR-FL | LPAR2 | 0.83 | -0.22 | AR-FL |
| CA13 | -0.90 | 0.56 | AR-FL | FAM105A | 1.25 | -0.22 | AR-FL |
| ADRA1A | -1.91 | 0.55 | AR-FL | NKX3-1 | 0.88 | -0.21 | AR-FL |
| NOS3 | -0.60 | 0.55 | AR-FL | C3orf14 | 1.35 | -0.21 | AR-FL |
| AHNAK2 | -0.93 | 0.52 | AR-FL | TMEM210 | 1.23 | -0.21 | AR-FL |
| ZNF711 | -1.28 | 0.52 | AR-FL | EPS15 | 0.65 | -0.21 | AR-FL |
| TMEM45A | -0.77 | 0.52 | AR-FL | C20orf96 | 1.22 | -0.21 | AR-FL |
| ASS1 | -0.58 | 0.52 | AR-FL | GRK5 | 0.90 | -0.21 | AR-FL |
| SERPINB5 | -0.68 | 0.52 | AR-FL | SGK223 | 0.96 | -0.21 | AR-FL |
| CES3 | -0.96 | 0.51 | AR-FL | GUCY1A3 | 0.72 | -0.21 | AR-FL |
| PCP2 | -0.85 | 0.51 | AR-FL | UAP1 | 1.25 | -0.20 | AR-FL |
| GNG4 | -1.24 | 0.50 | AR-FL | ATP7B | 0.84 | -0.20 | AR-FL |
| QSOX1 | -0.79 | 0.50 | AR-FL | ALOX15 | 0.82 | -0.19 | AR-FL |
| RCN3 | -0.66 | 0.50 | AR-FL | PDE9A | 0.64 | -0.19 | AR-FL |
| HSPA4L | -1.51 | 0.49 | AR-FL | ARF6 | 0.71 | -0.19 | AR-FL |
| IRF1 | -0.70 | 0.48 | AR-FL | TJP2 | 1.03 | -0.19 | AR-FL |
| NFASC | -0.78 | 0.47 | AR-FL | ILDR1 | 0.62 | -0.18 | AR-FL |
| DYRK3 | -0.73 | 0.47 | AR-FL | TMEM135 | 0.68 | -0.18 | AR-FL |
| LEF1 | -0.79 | 0.47 | AR-FL | UCHL1 | 0.80 | -0.18 | AR-FL |
| UNC5CL | -0.75 | 0.47 | AR-FL | SPECC1L | 0.62 | -0.18 | AR-FL |
| EIF3C | -0.66 | 0.46 | AR-FL | POLB | 0.66 | -0.17 | AR-FL |
| ANKRD2 | -2.01 | 0.45 | AR-FL | SEC14L1 | 0.59 | -0.17 | AR-FL |
| PIFO | -0.82 | 0.45 | AR-FL | BCAR3 | 0.61 | -0.17 | AR-FL |
| KCNH3 | -1.22 | 0.45 | AR-FL | KLHL29 | 1.24 | -0.17 | AR-FL |
| SI | -4.83 | 0.45 | AR-FL | MMD | 0.80 | -0.16 | AR-FL |
| IRX3 | -1.09 | 0.45 | AR-FL | CEBPA | 0.75 | -0.16 | AR-FL |
| SPTB | -0.62 | 0.44 | AR-FL | PCCA | 0.66 | -0.16 | AR-FL |
| LRP1 | -0.76 | 0.44 | AR-FL | DNASE2B | 0.87 | -0.16 | AR-FL |
| RP5-1187M17.10 | -0.61 | 0.44 | AR-FL | SEMG2 | 2.47 | -0.16 | AR-FL |
| THRA | -0.74 | 0.43 | AR-FL | CARD6 | 0.69 | -0.15 | AR-FL |
| PLEKHA7 | -0.88 | 0.43 | AR-FL | PEX10 | 1.73 | -0.15 | AR-FL |
| ANKRD31 | -1.13 | 0.42 | AR-FL | ETS1 | 1.13 | -0.14 | AR-FL |
| NOVA1 | -1.87 | 0.41 | AR-FL | TTC39A | 0.60 | -0.13 | AR-FL |
| CAMK1D | -2.39 | 0.41 | AR-FL | SPATC1L | 0.85 | -0.13 | AR-FL |
| HNF4G | -0.82 | 0.40 | AR-FL | DLGAP1 | 0.82 | -0.12 | AR-FL |
| RAB40A | -0.65 | 0.39 | AR-FL | TP53I11 | 1.19 | -0.12 | AR-FL |

| | | | | | | | |
|---|---|---|---|---|---|---|---|
| IFIT3 | -0.78 | 0.39 | AR-FL | DMXL1 | 0.60 | -0.12 | AR-FL |
| TCF4 | -0.71 | 0.39 | AR-FL | HES2 | 2.49 | -0.12 | AR-FL |
| CDHR3 | -0.68 | 0.37 | AR-FL | LIMD2 | 1.22 | -0.11 | AR-FL |
| APBB1 | -0.81 | 0.37 | AR-FL | FMNL2 | 0.83 | -0.11 | AR-FL |
| FAM234B | -0.81 | 0.36 | AR-FL | TMEM55A | 0.69 | -0.11 | AR-FL |
| SLC1A1 | -0.65 | 0.35 | AR-FL | NKAIN1 | 0.58 | -0.11 | AR-FL |
| NAP1L5 | -1.63 | 0.35 | AR-FL | SOAT1 | 0.58 | -0.11 | AR-FL |
| MCTP1 | -1.92 | 0.35 | AR-FL | BHLHE40 | 1.31 | -0.10 | AR-FL |
| DIAPH2 | -0.82 | 0.35 | AR-FL | ABLIM2 | 0.80 | -0.10 | AR-FL |
| MICA | -1.34 | 0.35 | AR-FL | MALL | 1.27 | -0.10 | AR-FL |
| RIBC1 | -0.83 | 0.35 | AR-FL | BAZ1A | 0.64 | -0.10 | AR-FL |
| NUAK2 | -1.67 | 0.35 | AR-FL | ASPHD2 | 1.50 | -0.10 | AR-FL |
| CCDC146 | -0.94 | 0.34 | AR-FL | CSGALNACT1 | 2.52 | -0.09 | AR-FL |
| GRIN1 | -0.70 | 0.34 | AR-FL | ANXA5 | 0.78 | -0.09 | AR-FL |
| RAD9B | -0.64 | 0.33 | AR-FL | CORO2A | 1.40 | -0.09 | AR-FL |
| PBX4 | -0.75 | 0.32 | AR-FL | CRACR2A | 1.93 | -0.09 | AR-FL |
| F2R | -0.84 | 0.32 | AR-FL | OSMR | 0.76 | -0.09 | AR-FL |
| EPHX4 | -1.58 | 0.31 | AR-FL | MARVELD3 | 1.11 | -0.08 | AR-FL |
| SPATA18 | -1.01 | 0.31 | AR-FL | GJB2 | 0.93 | -0.08 | AR-FL |
| HLF | -0.80 | 0.30 | AR-FL | INTS10 | 1.13 | -0.08 | AR-FL |
| ENPP3 | -0.90 | 0.30 | AR-FL | PTPRN2 | 0.63 | -0.08 | AR-FL |
| ABCC5 | -0.79 | 0.29 | AR-FL | C9orf152 | 0.66 | -0.08 | AR-FL |
| GYLTL1B | -0.69 | 0.29 | AR-FL | ERBB2IP | 0.89 | -0.08 | AR-FL |
| KCNG1 | -1.06 | 0.28 | AR-FL | TCTN2 | 0.86 | -0.08 | AR-FL |
| ZNF385C | -0.89 | 0.28 | AR-FL | GLRX2 | 1.60 | -0.08 | AR-FL |
| ERBB2 | -0.59 | 0.28 | AR-FL | MEX3D | 0.67 | -0.08 | AR-FL |
| ATP1B1 | -1.07 | 0.28 | AR-FL | PDZRN3 | 0.78 | -0.07 | AR-FL |
| TRIB1 | -0.82 | 0.28 | AR-FL | RIPK4 | 0.64 | -0.07 | AR-FL |
| FBLN2 | -1.16 | 0.28 | AR-FL | SLC8B1 | 0.65 | -0.07 | AR-FL |
| PLXDC1 | -0.86 | 0.28 | AR-FL | KAZALD1 | 1.25 | -0.07 | AR-FL |
| EPHB2 | -0.89 | 0.27 | AR-FL | C1orf116 | 0.61 | -0.06 | AR-FL |
| STON2 | -1.17 | 0.27 | AR-FL | TSPAN33 | 0.65 | -0.06 | AR-FL |
| LCAT | -0.60 | 0.27 | AR-FL | AP1S2 | 0.58 | -0.06 | AR-FL |
| NLGN1 | -1.72 | 0.27 | AR-FL | PDGFA | 0.91 | -0.06 | AR-FL |
| PLCXD1 | -0.82 | 0.26 | AR-FL | FAM214A | 0.80 | -0.04 | AR-FL |
| MMP24 | -0.78 | 0.26 | AR-FL | PXDN | 0.66 | -0.04 | AR-FL |
| FAM167B | -0.76 | 0.25 | AR-FL | ZBTB24 | 0.69 | -0.04 | AR-FL |
| CIART | -0.82 | 0.24 | AR-FL | LMAN1L | 1.82 | -0.04 | AR-FL |
| NATD1 | -1.66 | 0.24 | AR-FL | STXBP1 | 0.64 | -0.04 | AR-FL |
| LYST | -0.64 | 0.24 | AR-FL | CEP55 | 0.68 | -0.04 | AR-FL |
| AP006285.2 | -0.70 | 0.24 | AR-FL | MMP15 | 1.05 | -0.04 | AR-FL |
| PKDREJ | -0.63 | 0.24 | AR-FL | GPCPD1 | 1.52 | -0.03 | AR-FL |
| ZIC2 | -0.73 | 0.24 | AR-FL | MALT1 | 1.06 | -0.03 | AR-FL |
| ANKS1B | -0.65 | 0.24 | AR-FL | TRAM2 | 0.92 | -0.03 | AR-FL |
| HLA-DQB1 | -0.79 | 0.24 | AR-FL | SEC62 | 0.58 | -0.03 | AR-FL |
| C1orf145 | -0.89 | 0.23 | AR-FL | RAB19 | 0.59 | -0.03 | AR-FL |

| | | | |
|---|---|---|---|
| LRRC73 | -1.13 | 0.22 | AR-FL |
| EBF4 | -0.71 | 0.22 | AR-FL |
| TBX6 | -0.87 | 0.22 | AR-FL |
| AR | -0.63 | 0.21 | AR-FL |
| KLHL13 | -0.62 | 0.21 | AR-FL |
| ALDH3B1 | -0.67 | 0.21 | AR-FL |
| MYLK | -0.94 | 0.20 | AR-FL |
| CBLN2 | -2.03 | 0.20 | AR-FL |
| MAST1 | -0.67 | 0.20 | AR-FL |
| LFNG | -1.08 | 0.20 | AR-FL |
| ANKRD16 | -0.85 | 0.19 | AR-FL |
| UGT2B17 | -2.31 | 0.19 | AR-FL |
| SCARF1 | -0.95 | 0.19 | AR-FL |
| KIAA1614 | -0.78 | 0.19 | AR-FL |
| LAMB1 | -1.10 | 0.19 | AR-FL |
| TPPP | -0.60 | 0.19 | AR-FL |
| IQCC | -0.62 | 0.19 | AR-FL |
| RSPH1 | -0.64 | 0.18 | AR-FL |
| SEMA6C | -0.64 | 0.18 | AR-FL |
| COL16A1 | -2.45 | 0.17 | AR-FL |
| MFSD4 | -1.01 | 0.17 | AR-FL |
| DENND5B | -0.88 | 0.17 | AR-FL |
| ST7 | -1.40 | 0.17 | AR-FL |
| NAB1 | -0.79 | 0.17 | AR-FL |
| PRKG2 | -0.64 | 0.16 | AR-FL |
| S1PR3 | -0.77 | 0.15 | AR-FL |
| CCDC69 | -1.59 | 0.15 | AR-FL |
| LASP1 | -1.25 | 0.15 | AR-FL |
| TCF7L2 | -0.88 | 0.14 | AR-FL |
| RMI1 | -0.67 | 0.14 | AR-FL |
| CABLES2 | -0.61 | 0.14 | AR-FL |
| HEG1 | -0.58 | 0.14 | AR-FL |
| ABCB9 | -0.59 | 0.13 | AR-FL |
| POLR3GL | -0.74 | 0.13 | AR-FL |
| ASMTL | -0.66 | 0.13 | AR-FL |
| STX16-NPEPL1 | -0.64 | 0.13 | AR-FL |
| TMEM38A | -0.58 | 0.13 | AR-FL |
| PCSK6 | -0.61 | 0.12 | AR-FL |
| DHDH | -1.12 | 0.12 | AR-FL |
| PNPLA3 | -1.11 | 0.11 | AR-FL |
| MAGEA2 | -0.79 | 0.11 | AR-FL |
| PDE6D | -0.60 | 0.11 | AR-FL |
| ACOXL | -0.59 | 0.11 | AR-FL |
| NUP93 | -1.35 | 0.11 | AR-FL |
| CLSTN3 | -0.74 | 0.10 | AR-FL |
| MACROD2 | -0.61 | 0.10 | AR-FL |
| ITSN2 | 0.82 | -0.03 | AR-FL |
| CRISP3 | 1.09 | -0.02 | AR-FL |
| TRAPPC5 | 0.72 | -0.02 | AR-FL |
| NDRG3 | 0.61 | -0.02 | AR-FL |
| OSTF1 | 0.83 | -0.01 | AR-FL |
| TJP1 | 0.68 | -0.01 | AR-FL |
| CEACAM1 | 1.43 | -0.01 | AR-FL |
| CD320 | 0.65 | -0.01 | AR-FL |
| ITGB5 | 0.67 | -0.01 | AR-FL |
| FBN2 | 1.29 | -0.01 | AR-FL |
| CD24 | 1.05 | 0.00 | AR-FL |
| SH3RF2 | 1.41 | 0.00 | AR-FL |
| CADM2 | 0.99 | 0.00 | AR-FL |
| INCA1 | 1.26 | 0.00 | AR-FL |
| MAP3K14 | -0.59 | 0.00 | AR-FL |
| CDK5R2 | -0.82 | 0.00 | AR-FL |
| C11orf70 | -2.33 | 0.00 | AR-FL |
| ADAM8 | -1.14 | 0.00 | AR-FL |
| TMEM67 | -0.58 | 0.00 | AR-FL |
| SNAPC1 | -0.60 | 0.00 | AR-FL |
| ERAP2 | -1.08 | 0.01 | AR-FL |
| ATP6V1C2 | -0.82 | 0.01 | AR-FL |
| PCSK6 | -0.81 | 0.01 | AR-FL |
| TOP2A | -0.60 | 0.01 | AR-FL |
| CTGF | -1.94 | 0.01 | AR-FL |
| EYS | -0.87 | 0.01 | AR-FL |
| FOXO3 | -1.05 | 0.02 | AR-FL |
| PRR22 | -1.07 | 0.02 | AR-FL |
| KCNQ2 | -0.59 | 0.02 | AR-FL |
| TAS2R31 | -1.01 | 0.03 | AR-FL |
| WDR97 | -0.69 | 0.03 | AR-FL |
| RGS11 | -0.91 | 0.03 | AR-FL |
| IGSF9B | -0.68 | 0.03 | AR-FL |
| C16orf45 | -1.71 | 0.03 | AR-FL |
| ENO2 | -1.06 | 0.03 | AR-FL |
| SMARCA2 | -1.07 | 0.03 | AR-FL |
| MTUS2 | -0.92 | 0.04 | AR-FL |
| PEG10 | -2.04 | 0.04 | AR-FL |
| CRIP1 | -1.50 | 0.04 | AR-FL |
| SLC4A8 | -0.91 | 0.04 | AR-FL |
| IRS1 | -1.43 | 0.04 | AR-FL |
| VWA8 | -0.75 | 0.04 | AR-FL |
| ARHGDIG | -0.80 | 0.04 | AR-FL |
| SORBS1 | -0.75 | 0.05 | AR-FL |
| LDB3 | -0.71 | 0.05 | AR-FL |
| SYT12 | -1.90 | 0.05 | AR-FL |

| | | | |
|---|---|---|---|
| DDIT4L | -1.48 | 0.10 | AR-FL |
| FANK1 | -0.68 | 0.10 | AR-FL |
| WEE1 | -0.67 | 0.10 | AR-FL |
| CYP2C18 | -0.58 | 0.10 | AR-FL |
| PAG1 | -0.69 | 0.09 | AR-FL |
| CCDC88B | -1.03 | 0.08 | AR-FL |
| SFR1 | -0.71 | 0.08 | AR-FL |
| EPHX1 | -0.65 | 0.08 | AR-FL |
| VTA1 | -0.59 | 0.08 | AR-FL |
| QRICH2 | -0.64 | 0.08 | AR-FL |
| CD82 | -0.76 | 0.07 | AR-FL |
| C2CD4D | -0.64 | 0.07 | AR-FL |
| MGAT5 | -0.60 | 0.07 | AR-FL |
| SUSD5 | -0.90 | 0.07 | AR-FL |
| PAQR8 | -0.97 | 0.07 | AR-FL |
| ARHGEF17 | -1.04 | 0.06 | AR-FL |
| KIAA1683 | -0.91 | 0.06 | AR-FL |
| CCDC148 | -1.15 | 0.06 | AR-FL |
| CHML | -0.62 | 0.06 | AR-FL |
| TMEM91 | -0.61 | 0.05 | AR-FL |
| F10 | -0.59 | 0.05 | AR-FL |
| RP11-1212A22.4 | -0.71 | 0.05 | AR-FL |
| C17orf53 | -0.94 | 0.05 | AR-FL |
| MLKL | -0.91 | 0.05 | AR-FL |
| IL17RD | -1.13 | 0.05 | AR-FL |
| HMGB2 | -0.77 | 0.05 | AR-FL |
| SH3BP5 | -0.68 | 0.05 | AR-FL |
| EPS8 | -1.06 | 0.04 | AR-FL |
| SMARCD3 | -1.24 | 0.04 | AR-FL |
| HPCAL4 | -1.21 | 0.04 | AR-FL |
| ZNF695 | -0.80 | 0.04 | AR-FL |
| GPC1 | -0.67 | 0.04 | AR-FL |
| WDR31 | -0.69 | 0.03 | AR-FL |
| PLAUR | -0.90 | 0.03 | AR-FL |
| SLC9A1 | -0.59 | 0.03 | AR-FL |
| BLM | -0.95 | 0.03 | AR-FL |
| ASB13 | -0.81 | 0.02 | AR-FL |
| CLEC2D | -0.64 | 0.02 | AR-FL |
| ZNF385A | -0.93 | 0.02 | AR-FL |
| FAM83D | -0.64 | 0.02 | AR-FL |
| ZFHX2 | -0.69 | 0.02 | AR-FL |
| KRTAP5-1 | -0.78 | 0.02 | AR-FL |
| PLK3 | -0.62 | 0.02 | AR-FL |
| ZNF519 | -0.73 | 0.01 | AR-FL |
| SEMA6A | -1.63 | 0.01 | AR-FL |
| EMID1 | -1.25 | 0.01 | AR-FL |
| PYGO1 | -1.42 | 0.05 | AR-FL |
| TTC23L | -0.95 | 0.06 | AR-FL |
| SYCE2 | -1.68 | 0.06 | AR-FL |
| SLC1A2 | -0.95 | 0.06 | AR-FL |
| ACRV1 | -0.70 | 0.06 | AR-FL |
| CARNS1 | -0.99 | 0.06 | AR-FL |
| CSAD | -0.58 | 0.06 | AR-FL |
| PRUNE2 | -1.35 | 0.06 | AR-FL |
| RAPGEFL1 | -1.09 | 0.06 | AR-FL |
| ZNF76 | -0.65 | 0.06 | AR-FL |
| MAP3K12 | -0.68 | 0.07 | AR-FL |
| A2ML1 | -1.28 | 0.07 | AR-FL |
| DEPDC4 | -0.77 | 0.07 | AR-FL |
| AMH | -0.59 | 0.07 | AR-FL |
| EMID1 | -0.71 | 0.07 | AR-FL |
| ZNF671 | -0.58 | 0.07 | AR-FL |
| CLDN1 | -0.99 | 0.07 | AR-FL |
| SRD5A1 | -0.93 | 0.07 | AR-FL |
| IFT172 | -0.61 | 0.08 | AR-FL |
| TOR4A | -1.17 | 0.08 | AR-FL |
| UBE2L6 | -0.81 | 0.08 | AR-FL |
| LCA5L | -0.64 | 0.08 | AR-FL |
| PHKG1 | -0.77 | 0.08 | AR-FL |
| NAV1 | -0.88 | 0.08 | AR-FL |
| IFITM10 | -0.73 | 0.08 | AR-FL |
| ACTA2 | -0.68 | 0.08 | AR-FL |
| CPNE7 | -0.98 | 0.08 | AR-FL |
| NFIL3 | -1.29 | 0.09 | AR-FL |
| EPB41L2 | -0.59 | 0.09 | AR-FL |
| ALDH3B1 | -0.94 | 0.09 | AR-FL |
| SLC9A5 | -0.86 | 0.09 | AR-FL |
| PLA2G7 | -1.39 | 0.09 | AR-FL |
| AKNA | -0.67 | 0.09 | AR-FL |
| SNPH | -0.79 | 0.09 | AR-FL |
| WBP2NL | -0.92 | 0.10 | AR-FL |
| RAB6B | -0.95 | 0.10 | AR-FL |
| PPFIBP2 | -0.71 | 0.11 | AR-FL |
| TMEM65 | -0.86 | 0.11 | AR-FL |
| ARSK | -0.84 | 0.11 | AR-FL |
| CLIP3 | -1.51 | 0.11 | AR-FL |
| ZNF19 | -1.31 | 0.11 | AR-FL |
| ACBD7 | -0.88 | 0.11 | AR-FL |
| PDLIM1 | -1.46 | 0.12 | AR-FL |
| TFAP2E | -0.60 | 0.12 | AR-FL |
| NME5 | -0.76 | 0.12 | AR-FL |
| ZNF132 | -0.63 | 0.12 | AR-FL |

| | | | |
|---|---|---|---|
| TCTEX1D4 | -1.12 | 0.01 | AR-FL |
| ARHGEF2 | -0.74 | 0.01 | AR-FL |
| CEP152 | -0.76 | 0.00 | AR-FL |
| SOWAHD | -1.12 | 0.00 | AR-FL |
| TMEM132A | -0.78 | 0.00 | AR-FL |
| MAN1A1 | -3.29 | 0.00 | AR-FL |
| SYT4 | 0.02 | -7.26 | AR-V7 |
| KCNH8 | 0.08 | -6.48 | AR-V7 |
| C5orf42 | 0.18 | -6.22 | AR-V7 |
| OBSL1 | 0.01 | -5.66 | AR-V7 |
| DSC3 | 0.02 | -5.38 | AR-V7 |
| PDE10A | 0.18 | -3.81 | AR-V7 |
| EBF2 | 0.13 | -3.70 | AR-V7 |
| MSI1 | 0.27 | -3.41 | AR-V7 |
| PSMD5 | 0.37 | -3.35 | AR-V7 |
| FAM213A | 0.16 | -3.30 | AR-V7 |
| SLC4A4 | 0.53 | -3.05 | AR-V7 |
| GUCY1B3 | 0.06 | -2.98 | AR-V7 |
| PHF11 | 0.21 | -2.96 | AR-V7 |
| KCND3 | 0.14 | -2.92 | AR-V7 |
| RLN1 | 0.11 | -2.79 | AR-V7 |
| NDUFA4L2 | 0.06 | -2.59 | AR-V7 |
| FOXD1 | 0.46 | -2.58 | AR-V7 |
| SP140L | 0.33 | -2.49 | AR-V7 |
| DNALI1 | 0.27 | -2.46 | AR-V7 |
| MDGA1 | 0.28 | -2.37 | AR-V7 |
| PXDN | 0.58 | -2.35 | AR-V7 |
| PHTF1 | 0.41 | -2.34 | AR-V7 |
| TMEM45B | 0.51 | -2.29 | AR-V7 |
| STOX2 | 0.02 | -2.12 | AR-V7 |
| ECHDC1 | 0.02 | -1.87 | AR-V7 |
| UTRN | 0.53 | -1.87 | AR-V7 |
| TMEM178B | 0.39 | -1.83 | AR-V7 |
| ADRA2A | 0.09 | -1.77 | AR-V7 |
| PIK3AP1 | 0.02 | -1.76 | AR-V7 |
| PCBP3 | 0.04 | -1.73 | AR-V7 |
| RNF152 | 0.40 | -1.72 | AR-V7 |
| FAM110C | 0.02 | -1.71 | AR-V7 |
| RNF150 | 0.52 | -1.71 | AR-V7 |
| PRPH | 0.10 | -1.68 | AR-V7 |
| DNAH8 | 0.05 | -1.65 | AR-V7 |
| PLXDC2 | 0.55 | -1.56 | AR-V7 |
| FAM129A | 0.15 | -1.53 | AR-V7 |
| SP100 | 0.17 | -1.52 | AR-V7 |
| IFITM2 | 0.32 | -1.52 | AR-V7 |
| MCTP2 | 0.58 | -1.50 | AR-V7 |
| COL27A1 | -2.08 | 0.12 | AR-FL |
| PLPP2 | -1.09 | 0.12 | AR-FL |
| LETM2 | -0.64 | 0.13 | AR-FL |
| ENO3 | -0.92 | 0.13 | AR-FL |
| CYP3A5 | -1.13 | 0.13 | AR-FL |
| TMEM179 | -1.36 | 0.13 | AR-FL |
| CEP126 | -2.73 | 0.13 | AR-FL |
| TGFBR2 | -0.86 | 0.14 | AR-FL |
| ETS2 | -1.41 | 0.14 | AR-FL |
| DNAH1 | -0.78 | 0.14 | AR-FL |
| FBXL2 | -0.77 | 0.14 | AR-FL |
| ERC2 | -1.14 | 0.14 | AR-FL |
| SAXO2 | -0.92 | 0.14 | AR-FL |
| TANC2 | -0.73 | 0.15 | AR-FL |
| HYPK | -0.67 | 0.15 | AR-FL |
| RNF207 | -0.75 | 0.16 | AR-FL |
| TMEM91 | -1.02 | 0.16 | AR-FL |
| FAM19A5 | -0.89 | 0.16 | AR-FL |
| IFT80 | -0.68 | 0.16 | AR-FL |
| IGFBP2 | -0.68 | 0.16 | AR-FL |
| CCDC93 | -0.61 | 0.17 | AR-FL |
| RERG | -2.06 | 0.18 | AR-FL |
| SEMA7A | -1.19 | 0.18 | AR-FL |
| PARP14 | -1.11 | 0.18 | AR-FL |
| PLEKHH2 | -0.88 | 0.18 | AR-FL |
| ZNF337 | -0.66 | 0.19 | AR-FL |
| ASPHD1 | -0.76 | 0.20 | AR-FL |
| STOX2 | -0.76 | 0.20 | AR-FL |
| FAM198A | -1.53 | 0.20 | AR-FL |
| ADM5 | -1.06 | 0.20 | AR-FL |
| TREX2 | -0.60 | 0.20 | AR-FL |
| SDC2 | -2.29 | 0.21 | AR-FL |
| EIF4A1 | -0.68 | 0.21 | AR-FL |
| FERMT2 | -1.14 | 0.21 | AR-FL |
| PNMA3 | -1.60 | 0.21 | AR-FL |
| TMC3 | -0.84 | 0.22 | AR-FL |
| RAP1GDS1 | -0.62 | 0.22 | AR-FL |
| USP46 | -0.63 | 0.23 | AR-FL |
| SGSM2 | -0.75 | 0.23 | AR-FL |
| CDKN2A | -1.41 | 0.23 | AR-FL |
| LRRC4 | -2.09 | 0.24 | AR-FL |
| FMNL3 | -0.81 | 0.24 | AR-FL |
| NAT8L | -0.93 | 0.24 | AR-FL |
| RP11-849H4.2 | -0.69 | 0.24 | AR-FL |
| NGFRAP1 | -0.76 | 0.24 | AR-FL |
| MROH8 | -1.03 | 0.25 | AR-FL |

| | | | | | | | |
|---|---|---|---|---|---|---|---|
| SCARF2 | 0.08 | -1.46 | AR-V7 | RIPK2 | -1.01 | 0.25 | AR-FL |
| CLEC7A | 0.42 | -1.46 | AR-V7 | RCCD1 | -1.10 | 0.25 | AR-FL |
| UST | 0.52 | -1.45 | AR-V7 | NTSR1 | -1.74 | 0.25 | AR-FL |
| CACNA1D | 0.11 | -1.39 | AR-V7 | MAMDC4 | -0.74 | 0.25 | AR-FL |
| NAALADL2 | 0.07 | -1.37 | AR-V7 | TUBE1 | -0.67 | 0.26 | AR-FL |
| OSBPL10 | 0.25 | -1.35 | AR-V7 | ATOH1 | -2.28 | 0.26 | AR-FL |
| GYG2 | 0.02 | -1.34 | AR-V7 | AGTR1 | -2.14 | 0.26 | AR-FL |
| SLC16A3 | 0.54 | -1.34 | AR-V7 | AF011889.5 | -1.10 | 0.27 | AR-FL |
| SLC1A5 | 0.31 | -1.33 | AR-V7 | FLT4 | -1.00 | 0.27 | AR-FL |
| SMAGP | 0.40 | -1.33 | AR-V7 | FAM169A | -0.89 | 0.28 | AR-FL |
| RINL | 0.02 | -1.31 | AR-V7 | SACS | -0.81 | 0.28 | AR-FL |
| TP53I11 | 0.46 | -1.30 | AR-V7 | PTGER4 | -1.66 | 0.28 | AR-FL |
| HEY1 | 0.22 | -1.29 | AR-V7 | ITGB8 | -1.48 | 0.28 | AR-FL |
| GPC6 | 0.56 | -1.26 | AR-V7 | SLC5A3 | -0.72 | 0.29 | AR-FL |
| ABHD17C | 0.30 | -1.24 | AR-V7 | C2orf15 | -1.04 | 0.30 | AR-FL |
| OSBPL6 | 0.37 | -1.24 | AR-V7 | SYNGR1 | -1.09 | 0.31 | AR-FL |
| SPIRE1 | 0.13 | -1.23 | AR-V7 | IQGAP3 | -0.71 | 0.31 | AR-FL |
| MYO6 | 0.38 | -1.22 | AR-V7 | FAM102B | -1.47 | 0.31 | AR-FL |
| CAB39L | 0.19 | -1.22 | AR-V7 | WIF1 | -0.70 | 0.31 | AR-FL |
| TANC1 | 0.05 | -1.17 | AR-V7 | NKD1 | -0.73 | 0.31 | AR-FL |
| FAM135A | 0.23 | -1.16 | AR-V7 | PDE4A | -1.64 | 0.32 | AR-FL |
| CRIP2 | 0.53 | -1.16 | AR-V7 | SUSD4 | -1.11 | 0.32 | AR-FL |
| RNF24 | 0.18 | -1.13 | AR-V7 | MXRA7 | -0.81 | 0.32 | AR-FL |
| SPRY1 | 0.40 | -1.11 | AR-V7 | THRA | -1.22 | 0.32 | AR-FL |
| CYP2E1 | 0.37 | -1.08 | AR-V7 | ADD2 | -0.84 | 0.33 | AR-FL |
| DTNA | 0.12 | -1.08 | AR-V7 | DGAT2 | -0.69 | 0.33 | AR-FL |
| CPT1A | 0.11 | -1.06 | AR-V7 | HS3ST1 | -0.77 | 0.33 | AR-FL |
| COLGALT2 | 0.38 | -1.05 | AR-V7 | TLR5 | -0.71 | 0.34 | AR-FL |
| MED27 | 0.07 | -1.05 | AR-V7 | TMEM59L | -1.28 | 0.35 | AR-FL |
| SNAP91 | 0.41 | -1.04 | AR-V7 | TPH1 | -0.64 | 0.35 | AR-FL |
| NEDD4 | 0.18 | -1.03 | AR-V7 | GJC1 | -0.67 | 0.35 | AR-FL |
| RCBTB2 | 0.08 | -1.02 | AR-V7 | KCND1 | -1.36 | 0.35 | AR-FL |
| SAPCD2 | 0.03 | -1.02 | AR-V7 | ITGA7 | -1.02 | 0.35 | AR-FL |
| NRIP1 | 0.22 | -1.00 | AR-V7 | MNS1 | -0.65 | 0.35 | AR-FL |
| TTC7B | 0.02 | -0.98 | AR-V7 | TSSK4 | -0.84 | 0.35 | AR-FL |
| DGKB | 0.46 | -0.97 | AR-V7 | MYLK | -1.12 | 0.36 | AR-FL |
| PACSIN1 | 0.23 | -0.94 | AR-V7 | SNX22 | -2.01 | 0.36 | AR-FL |
| HCN2 | 0.58 | -0.94 | AR-V7 | PCYOX1L | -0.60 | 0.36 | AR-FL |
| CYP2J2 | 0.08 | -0.94 | AR-V7 | FLRT1 | -0.74 | 0.37 | AR-FL |
| ERGIC1 | 0.50 | -0.94 | AR-V7 | SEMA6D | -1.49 | 0.38 | AR-FL |
| TMC1 | 0.22 | -0.90 | AR-V7 | PIWIL4 | -0.74 | 0.38 | AR-FL |
| SMAD9 | 0.11 | -0.89 | AR-V7 | BIRC3 | -2.16 | 0.39 | AR-FL |
| SPON2 | 0.32 | -0.89 | AR-V7 | ZNF23 | -0.81 | 0.39 | AR-FL |
| EPB41L4B | 0.23 | -0.87 | AR-V7 | BCL6 | -0.82 | 0.39 | AR-FL |
| RALGAPA2 | 0.32 | -0.87 | AR-V7 | YAP1 | -0.67 | 0.40 | AR-FL |
| EFNB2 | 0.27 | -0.85 | AR-V7 | ALCAM | -1.50 | 0.41 | AR-FL |

| | | | | | | | |
|---|---|---|---|---|---|---|---|
| ARID3A | 0.01 | -0.85 | AR-V7 | CHAC1 | -0.66 | 0.41 | AR-FL |
| LAD1 | 0.11 | -0.85 | AR-V7 | TNC | -4.93 | 0.41 | AR-FL |
| DNPEP | 0.18 | -0.84 | AR-V7 | CCDC30 | -0.71 | 0.41 | AR-FL |
| SLC19A1 | 0.03 | -0.82 | AR-V7 | UACA | -0.66 | 0.42 | AR-FL |
| NVL | 0.04 | -0.82 | AR-V7 | NAV2 | -1.36 | 0.43 | AR-FL |
| SLC25A48 | 0.05 | -0.81 | AR-V7 | PCBP3 | -0.80 | 0.44 | AR-FL |
| PVRL1 | 0.13 | -0.81 | AR-V7 | SNX33 | -1.08 | 0.45 | AR-FL |
| RIMS1 | 0.30 | -0.81 | AR-V7 | PLXNC1 | -0.59 | 0.46 | AR-FL |
| ADAMTS1 | 0.54 | -0.81 | AR-V7 | SCN3B | -0.62 | 0.46 | AR-FL |
| R3HDM2 | 0.52 | -0.81 | AR-V7 | TMEM63C | -1.42 | 0.47 | AR-FL |
| DDAH1 | 0.20 | -0.80 | AR-V7 | FOXE1 | -1.37 | 0.47 | AR-FL |
| MARVELD1 | 0.03 | -0.79 | AR-V7 | RADIL | -0.75 | 0.48 | AR-FL |
| IMPG1 | 0.42 | -0.79 | AR-V7 | MECOM | -2.28 | 0.48 | AR-FL |
| ADARB1 | 0.27 | -0.78 | AR-V7 | GFOD1 | -0.79 | 0.50 | AR-FL |
| ME1 | 0.30 | -0.78 | AR-V7 | JAZF1 | -0.66 | 0.50 | AR-FL |
| ABCE1 | 0.01 | -0.77 | AR-V7 | DHRSX | -1.10 | 0.51 | AR-FL |
| FAM222A | 0.11 | -0.76 | AR-V7 | TM7SF3 | -0.67 | 0.51 | AR-FL |
| DOCK9 | 0.05 | -0.76 | AR-V7 | RGS2 | -1.09 | 0.54 | AR-FL |
| ZNF75D | 0.02 | -0.75 | AR-V7 | ADAM12 | -0.94 | 0.54 | AR-FL |
| MID2 | 0.02 | -0.75 | AR-V7 | IGFBP1 | -0.91 | 0.54 | AR-FL |
| CDK5R1 | 0.40 | -0.75 | AR-V7 | TBC1D32 | -0.60 | 0.54 | AR-FL |
| GTF3A | 0.07 | -0.75 | AR-V7 | RNF122 | -0.96 | 0.55 | AR-FL |
| TOP2B | 0.06 | -0.74 | AR-V7 | QSOX1 | -1.46 | 0.57 | AR-FL |
| AGAP1 | 0.24 | -0.73 | AR-V7 | PTGER3 | -1.39 | 0.57 | AR-FL |
| RPS6KA6 | 0.02 | -0.72 | AR-V7 | ARHGAP23 | -1.32 | 0.57 | AR-FL |
| KCTD1 | 0.38 | -0.72 | AR-V7 | FAM229B | -1.27 | 0.57 | AR-FL |
| RELL2 | 0.46 | -0.72 | AR-V7 | ST3GAL5 | -0.60 | 0.59 | Opposite |
| ERBB3 | 0.07 | -0.71 | AR-V7 | ADGRF1 | -2.89 | 0.60 | Opposite |
| UCK2 | 0.19 | -0.71 | AR-V7 | LAMA4 | -0.77 | 0.62 | Opposite |
| TNFSF4 | 0.48 | -0.71 | AR-V7 | PKIB | -1.19 | 0.62 | Opposite |
| GPRIN1 | 0.37 | -0.71 | AR-V7 | VEGFC | -1.80 | 0.67 | Opposite |
| ACTL10 | 0.16 | -0.70 | AR-V7 | WNT11 | -2.31 | 0.67 | Opposite |
| DDO | 0.20 | -0.69 | AR-V7 | SOX6 | -1.41 | 0.67 | Opposite |
| ZSCAN31 | 0.11 | -0.69 | AR-V7 | NEURL1 | -1.11 | 0.67 | Opposite |
| ZC3HAV1L | 0.29 | -0.69 | AR-V7 | SCARF1 | -0.66 | 0.71 | Opposite |
| LRP3 | 0.34 | -0.69 | AR-V7 | SCRN1 | -0.70 | 0.73 | Opposite |
| LITAF | 0.12 | -0.68 | AR-V7 | NPTX1 | -0.74 | 0.74 | Opposite |
| SPTBN2 | 0.38 | -0.68 | AR-V7 | FAM46A | -1.18 | 0.78 | Opposite |
| TUBB6 | 0.24 | -0.68 | AR-V7 | CRISPLD1 | -1.66 | 0.78 | Opposite |
| TNKS2 | 0.04 | -0.68 | AR-V7 | TEX9 | -1.24 | 0.78 | Opposite |
| OGDH | 0.27 | -0.67 | AR-V7 | SYN1 | -0.65 | 0.81 | Opposite |
| UBE2E2 | 0.15 | -0.67 | AR-V7 | RSPH1 | -0.76 | 0.81 | Opposite |
| GLO1 | 0.03 | -0.66 | AR-V7 | THBS1 | -2.14 | 0.82 | Opposite |
| MTCL1 | 0.36 | -0.66 | AR-V7 | MYH15 | -1.08 | 0.83 | Opposite |
| COTL1 | 0.33 | -0.66 | AR-V7 | MDH1B | -1.49 | 0.85 | Opposite |
| ZNF827 | 0.44 | -0.66 | AR-V7 | CHAC2 | -0.71 | 0.87 | Opposite |

| | | | | | | | |
|---|---|---|---|---|---|---|---|
| TUBG1 | 0.23 | -0.65 | AR-V7 | TMEM74B | -0.59 | 0.87 | Opposite |
| FAM162A | 0.00 | -0.65 | AR-V7 | KCNQ3 | -0.61 | 0.88 | Opposite |
| GMDS | 0.25 | -0.65 | AR-V7 | THRB | -0.97 | 0.89 | Opposite |
| RYR3 | 0.06 | -0.65 | AR-V7 | DISC1 | -1.58 | 0.93 | Opposite |
| PPFIA2 | 0.01 | -0.65 | AR-V7 | SLC1A1 | -0.84 | 0.95 | Opposite |
| FBXL17 | 0.27 | -0.65 | AR-V7 | LSAMP | -1.60 | 0.95 | Opposite |
| BVES | 0.54 | -0.65 | AR-V7 | SEMA4G | -0.99 | 0.95 | Opposite |
| FAM84B | 0.01 | -0.64 | AR-V7 | MT1G | -0.63 | 0.96 | Opposite |
| SERPINB6 | 0.02 | -0.64 | AR-V7 | NR3C2 | -1.25 | 1.01 | Opposite |
| IFNAR2 | 0.21 | -0.62 | AR-V7 | AK8 | -1.01 | 1.01 | Opposite |
| ZNRF2 | 0.09 | -0.61 | AR-V7 | PRIMA1 | -0.94 | 1.03 | Opposite |
| TBL1XR1 | 0.07 | -0.61 | AR-V7 | ZNF185 | -1.33 | 1.04 | Opposite |
| SLC25A5 | 0.25 | -0.61 | AR-V7 | CXorf57 | -0.62 | 1.05 | Opposite |
| TXNRD1 | 0.11 | -0.61 | AR-V7 | IGFBP5 | -1.12 | 1.06 | Opposite |
| MPHOSPH6 | 0.28 | -0.60 | AR-V7 | BTG1 | -0.83 | 1.10 | Opposite |
| SERINC5 | 0.43 | -0.60 | AR-V7 | ETNK2 | -1.05 | 1.10 | Opposite |
| DUSP2 | 0.46 | -0.60 | AR-V7 | ABLIM3 | -0.71 | 1.14 | Opposite |
| SETDB2 | 0.16 | -0.60 | AR-V7 | PDK4 | -7.12 | 1.16 | Opposite |
| NUDT3 | 0.02 | -0.60 | AR-V7 | ARX | -1.01 | 1.23 | Opposite |
| EHBP1 | 0.38 | -0.60 | AR-V7 | ANK2 | -0.96 | 1.36 | Opposite |
| GOLM1 | 0.43 | -0.60 | AR-V7 | ADRA2B | -1.35 | 1.40 | Opposite |
| TRIM14 | 0.09 | -0.59 | AR-V7 | TMEFF2 | -0.91 | 1.44 | Opposite |
| WDR4 | 0.05 | -0.59 | AR-V7 | TRPA1 | -3.21 | 1.51 | Opposite |
| WDR72 | 0.07 | -0.59 | AR-V7 | OCA2 | -1.50 | 1.53 | Opposite |
| ACTR3C | 0.02 | -0.59 | AR-V7 | PDZD2 | -1.17 | 1.59 | Opposite |
| DAAM1 | 0.31 | -0.58 | AR-V7 | GATSL2 | -1.06 | 1.60 | Opposite |
| VAV2 | 0.02 | -0.58 | AR-V7 | EPHA4 | -1.58 | 1.64 | Opposite |
| KIAA1107 | -0.15 | 0.58 | AR-V7 | ETV5 | -1.37 | 1.67 | Opposite |
| TCAP | -0.15 | 0.58 | AR-V7 | USP2 | -1.57 | 1.88 | Opposite |
| HEMK1 | -0.05 | 0.58 | AR-V7 | DUOX1 | -1.46 | 1.90 | Opposite |
| ODF2L | -0.10 | 0.58 | AR-V7 | GRIN3A | -0.86 | 1.91 | Opposite |
| DUSP19 | -0.24 | 0.58 | AR-V7 | POU3F1 | -1.95 | 1.91 | Opposite |
| KCNJ11 | -0.27 | 0.59 | AR-V7 | CCDC109B | -0.59 | 1.96 | Opposite |
| KLHDC9 | -0.11 | 0.59 | AR-V7 | MSX2 | -0.88 | 2.07 | Opposite |
| LOXL2 | -0.39 | 0.59 | AR-V7 | NR5A2 | -0.95 | 2.24 | Opposite |
| KIF6 | -0.27 | 0.59 | AR-V7 | ADORA1 | -0.81 | 2.38 | Opposite |
| EIF4A2 | -0.14 | 0.59 | AR-V7 | HOPX | -3.53 | 2.40 | Opposite |
| INTU | -0.14 | 0.59 | AR-V7 | NPY2R | -0.71 | 2.54 | Opposite |
| CFAP61 | -0.16 | 0.59 | AR-V7 | MAATS1 | -1.90 | 2.66 | Opposite |
| TMEM62 | -0.40 | 0.59 | AR-V7 | ATP2B2 | -2.61 | 2.69 | Opposite |
| MAP1B | -0.30 | 0.59 | AR-V7 | NGFR | -0.62 | 2.79 | Opposite |
| ZC2HC1C | -0.29 | 0.59 | AR-V7 | RASL11B | -1.55 | 2.81 | Opposite |
| SEC61A2 | -0.10 | 0.60 | AR-V7 | DGKG | -2.38 | 2.82 | Opposite |
| ARHGAP4 | -0.19 | 0.60 | AR-V7 | CHN2 | -1.38 | 2.95 | Opposite |
| ENKD1 | -0.08 | 0.60 | AR-V7 | KLHL31 | -0.84 | 3.02 | Opposite |
| AAMDC | -0.13 | 0.60 | AR-V7 | WISP2 | -1.68 | 3.12 | Opposite |

| | | | | | | | |
|---|---|---|---|---|---|---|---|
| SSPO | -0.16 | 0.60 | AR-V7 | NCMAP | -1.50 | 3.18 | Opposite |
| ULBP3 | -0.27 | 0.60 | AR-V7 | TAGAP | -1.53 | 3.23 | Opposite |
| GNPTAB | -0.31 | 0.60 | AR-V7 | UPP1 | -0.72 | 3.24 | Opposite |
| STAU2 | -0.07 | 0.61 | AR-V7 | MYH7 | -4.03 | 3.29 | Opposite |
| CCDC78 | -0.24 | 0.61 | AR-V7 | PSTPIP2 | -0.65 | 3.59 | Opposite |
| C14orf79 | -0.05 | 0.61 | AR-V7 | TEK | -1.22 | 4.09 | Opposite |
| DNAJA4 | -0.01 | 0.61 | AR-V7 | UBE2QL1 | -2.45 | 4.31 | Opposite |
| CASC1 | -0.46 | 0.61 | AR-V7 | ACVR1C | -1.78 | 4.44 | Opposite |
| CITED4 | -0.30 | 0.62 | AR-V7 | SULT1C2 | -0.58 | 4.77 | Opposite |
| DYX1C1 | -0.31 | 0.62 | AR-V7 | PLXNA4 | -1.13 | 5.04 | Opposite |
| TECTA | -0.43 | 0.63 | AR-V7 | PROC | -0.84 | 5.25 | Opposite |
| COL1A1 | -0.22 | 0.63 | AR-V7 | CABS1 | -2.28 | 5.27 | Opposite |
| RECK | -0.02 | 0.63 | AR-V7 | UGT3A2 | 0.38 | -3.24 | AR-V7 |
| IQSEC2 | -0.03 | 0.64 | AR-V7 | SP8 | 0.09 | -1.70 | AR-V7 |
| VAMP5 | -0.40 | 0.65 | AR-V7 | DNAJC22 | 0.29 | -1.33 | AR-V7 |
| SDC3 | -0.31 | 0.65 | AR-V7 | CSTA | 0.25 | -1.32 | AR-V7 |
| TRIM17 | -0.35 | 0.65 | AR-V7 | ABCG1 | 0.14 | -1.32 | AR-V7 |
| ELK1 | -0.03 | 0.65 | AR-V7 | MAGEB17 | 0.28 | -1.31 | AR-V7 |
| TLE1 | -0.20 | 0.66 | AR-V7 | DNAH8 | 0.46 | -1.27 | AR-V7 |
| ZC2HC1A | -0.13 | 0.66 | AR-V7 | HAO1 | 0.27 | -1.26 | AR-V7 |
| RABL2B | -0.18 | 0.66 | AR-V7 | INPP4B | 0.06 | -1.25 | AR-V7 |
| NBPF14 | -0.30 | 0.67 | AR-V7 | FGFR2 | 0.22 | -1.24 | AR-V7 |
| TCIRG1 | -0.09 | 0.67 | AR-V7 | HGD | 0.32 | -1.23 | AR-V7 |
| REC8 | -0.01 | 0.67 | AR-V7 | FFAR2 | 0.03 | -1.17 | AR-V7 |
| CEBPB | -0.06 | 0.67 | AR-V7 | KCNH8 | 0.26 | -1.14 | AR-V7 |
| ZNF66 | -0.20 | 0.67 | AR-V7 | SPTSSB | 0.25 | -1.14 | AR-V7 |
| CCNB1IP1 | -0.08 | 0.68 | AR-V7 | TMC1 | 0.04 | -1.08 | AR-V7 |
| BBS12 | 0.00 | 0.68 | AR-V7 | PLA2R1 | 0.23 | -1.07 | AR-V7 |
| C9orf173 | -0.44 | 0.68 | AR-V7 | FRMD5 | 0.50 | -1.07 | AR-V7 |
| FSD1L | -0.25 | 0.68 | AR-V7 | SERPINA1 | 0.33 | -1.02 | AR-V7 |
| ANKRD26 | -0.41 | 0.68 | AR-V7 | LCP1 | 0.23 | -1.00 | AR-V7 |
| CNPY4 | -0.08 | 0.69 | AR-V7 | KCNK3 | 0.12 | -0.97 | AR-V7 |
| RASSF4 | -0.30 | 0.69 | AR-V7 | MGAT4A | 0.22 | -0.96 | AR-V7 |
| PLXNA1 | -0.06 | 0.69 | AR-V7 | LNX1 | 0.46 | -0.94 | AR-V7 |
| VWA5A | -0.03 | 0.69 | AR-V7 | LHFP | 0.25 | -0.93 | AR-V7 |
| CPT1B | -0.02 | 0.70 | AR-V7 | RUNX2 | 0.05 | -0.93 | AR-V7 |
| MKL1 | -0.06 | 0.70 | AR-V7 | ITGA6 | 0.11 | -0.92 | AR-V7 |
| CYSTM1 | -0.25 | 0.70 | AR-V7 | LRP2BP | 0.19 | -0.89 | AR-V7 |
| CD37 | -0.18 | 0.70 | AR-V7 | REPS2 | 0.26 | -0.89 | AR-V7 |
| TNFAIP8L1 | -0.09 | 0.70 | AR-V7 | NCALD | 0.35 | -0.88 | AR-V7 |
| LMNTD2 | -0.18 | 0.71 | AR-V7 | SPRED1 | 0.22 | -0.88 | AR-V7 |
| TMEM64 | -0.30 | 0.71 | AR-V7 | TMEM133 | 0.07 | -0.87 | AR-V7 |
| RBPMS | -0.15 | 0.71 | AR-V7 | STAP2 | 0.23 | -0.87 | AR-V7 |
| GPT | -0.40 | 0.71 | AR-V7 | CGNL1 | 0.17 | -0.83 | AR-V7 |
| RABL2A | -0.29 | 0.71 | AR-V7 | CMTM8 | 0.14 | -0.82 | AR-V7 |
| SLC12A2 | -0.20 | 0.72 | AR-V7 | IGSF21 | 0.50 | -0.80 | AR-V7 |

| | | | |
|---|---|---|---|
| RANBP17 | -0.17 | 0.72 | AR-V7 |
| HSPA12B | -0.09 | 0.72 | AR-V7 |
| CDH8 | -0.40 | 0.73 | AR-V7 |
| GGT7 | -0.21 | 0.73 | AR-V7 |
| ACYP1 | -0.23 | 0.73 | AR-V7 |
| WDR19 | -0.16 | 0.73 | AR-V7 |
| FRMD4B | -0.42 | 0.73 | AR-V7 |
| TMC2 | -0.27 | 0.74 | AR-V7 |
| UACA | -0.43 | 0.74 | AR-V7 |
| ZNF467 | -0.21 | 0.74 | AR-V7 |
| CCDC189 | -0.22 | 0.74 | AR-V7 |
| C8G | -0.14 | 0.75 | AR-V7 |
| LIPH | -0.54 | 0.75 | AR-V7 |
| MOK | -0.07 | 0.75 | AR-V7 |
| PIANP | -0.26 | 0.76 | AR-V7 |
| ARNTL2 | -0.38 | 0.77 | AR-V7 |
| FAM217B | -0.55 | 0.78 | AR-V7 |
| C1orf228 | -0.53 | 0.78 | AR-V7 |
| TM7SF3 | -0.51 | 0.78 | AR-V7 |
| PCSK4 | -0.10 | 0.78 | AR-V7 |
| LCN12 | -0.54 | 0.78 | AR-V7 |
| SBSPON | -0.28 | 0.78 | AR-V7 |
| CCDC62 | -0.57 | 0.79 | AR-V7 |
| GOLT1A | -0.23 | 0.80 | AR-V7 |
| CDC14A | -0.53 | 0.80 | AR-V7 |
| NBN | -0.07 | 0.81 | AR-V7 |
| MIDN | -0.15 | 0.81 | AR-V7 |
| SLC16A9 | -0.01 | 0.81 | AR-V7 |
| ALKBH6 | -0.02 | 0.82 | AR-V7 |
| TJP3 | -0.20 | 0.83 | AR-V7 |
| PDE4A | -0.51 | 0.83 | AR-V7 |
| AQP1 | -0.21 | 0.83 | AR-V7 |
| NOXRED1 | -0.26 | 0.83 | AR-V7 |
| ZNF502 | -0.04 | 0.83 | AR-V7 |
| ZNF185 | -0.38 | 0.83 | AR-V7 |
| PRICKLE4 | -0.37 | 0.83 | AR-V7 |
| TSC22D3 | -0.06 | 0.83 | AR-V7 |
| AIFM2 | -0.34 | 0.83 | AR-V7 |
| ARX | -0.31 | 0.83 | AR-V7 |
| AKAP7 | -0.11 | 0.84 | AR-V7 |
| RNF32 | -0.21 | 0.84 | AR-V7 |
| GRAMD3 | -0.10 | 0.84 | AR-V7 |
| CCDC30 | -0.39 | 0.84 | AR-V7 |
| POGLUT1 | -0.29 | 0.85 | AR-V7 |
| TMEM56-RWDD3 | -0.11 | 0.85 | AR-V7 |
| TTC21A | -0.12 | 0.85 | AR-V7 |
| GPX2 | 0.15 | -0.79 | AR-V7 |
| ACOX2 | 0.40 | -0.78 | AR-V7 |
| SLA2 | 0.58 | -0.77 | AR-V7 |
| THSD4 | 0.31 | -0.77 | AR-V7 |
| OVGP1 | 0.22 | -0.76 | AR-V7 |
| IGSF5 | 0.16 | -0.76 | AR-V7 |
| TBX2 | 0.36 | -0.76 | AR-V7 |
| B4GALNT3 | 0.39 | -0.76 | AR-V7 |
| SRPX | 0.06 | -0.75 | AR-V7 |
| TACSTD2 | 0.43 | -0.73 | AR-V7 |
| BBS4 | 0.52 | -0.70 | AR-V7 |
| PLD2 | 0.03 | -0.70 | AR-V7 |
| SH3KBP1 | 0.08 | -0.69 | AR-V7 |
| GPR160 | 0.12 | -0.68 | AR-V7 |
| CDK17 | 0.28 | -0.66 | AR-V7 |
| SDK2 | 0.23 | -0.66 | AR-V7 |
| PKP1 | 0.36 | -0.65 | AR-V7 |
| LENG9 | 0.16 | -0.65 | AR-V7 |
| FAM162A | 0.07 | -0.64 | AR-V7 |
| ENTPD5 | 0.58 | -0.63 | AR-V7 |
| SMPDL3B | 0.02 | -0.63 | AR-V7 |
| RUNX1 | 0.05 | -0.62 | AR-V7 |
| TPM4 | 0.26 | -0.62 | AR-V7 |
| ONECUT2 | 0.07 | -0.60 | AR-V7 |
| RTN2 | 0.17 | -0.60 | AR-V7 |
| USP53 | 0.00 | -0.59 | AR-V7 |
| EML6 | 0.22 | -0.59 | AR-V7 |
| CARD9 | 0.28 | -0.59 | AR-V7 |
| PREX1 | 0.24 | -0.59 | AR-V7 |
| DNAH2 | 0.05 | -0.58 | AR-V7 |
| NRTN | -0.28 | 0.58 | AR-V7 |
| SLC6A8 | -0.09 | 0.58 | AR-V7 |
| DUSP9 | -0.15 | 0.60 | AR-V7 |
| PER2 | -0.25 | 0.60 | AR-V7 |
| MASP2 | -0.08 | 0.60 | AR-V7 |
| QSER1 | -0.41 | 0.60 | AR-V7 |
| C11orf54 | -0.06 | 0.60 | AR-V7 |
| DGCR6 | -0.07 | 0.61 | AR-V7 |
| PGAP2 | -0.25 | 0.61 | AR-V7 |
| APOH | -0.31 | 0.62 | AR-V7 |
| FBXO21 | -0.03 | 0.63 | AR-V7 |
| SRRM5 | -0.05 | 0.63 | AR-V7 |
| EME2 | -0.03 | 0.63 | AR-V7 |
| MXD3 | -0.53 | 0.64 | AR-V7 |
| ELK1 | -0.32 | 0.64 | AR-V7 |
| ITPR2 | -0.30 | 0.64 | AR-V7 |

| | | | | | | | |
|---|---|---|---|---|---|---|---|
| RIPK2 | -0.21 | 0.86 | AR-V7 | C18orf54 | -0.44 | 0.66 | AR-V7 |
| WDR66 | -0.03 | 0.86 | AR-V7 | ANOS1 | -0.43 | 0.66 | AR-V7 |
| CATIP | -0.02 | 0.86 | AR-V7 | C5orf34 | -0.51 | 0.67 | AR-V7 |
| MAP2K6 | -0.49 | 0.87 | AR-V7 | TUBGCP6 | -0.12 | 0.67 | AR-V7 |
| SLC6A16 | -0.30 | 0.87 | AR-V7 | GMPR | -0.24 | 0.68 | AR-V7 |
| UNC45A | -0.09 | 0.88 | AR-V7 | PNRC2 | -0.48 | 0.69 | AR-V7 |
| HPCAL1 | -0.02 | 0.89 | AR-V7 | NPIPB11 | -0.05 | 0.69 | AR-V7 |
| SYNGR3 | -0.03 | 0.89 | AR-V7 | PHTF2 | -0.10 | 0.70 | AR-V7 |
| PRUNE2 | -0.19 | 0.89 | AR-V7 | VPS33B | -0.16 | 0.70 | AR-V7 |
| CFAP43 | -0.10 | 0.89 | AR-V7 | AHNAK | -0.27 | 0.71 | AR-V7 |
| CFAP53 | -0.10 | 0.89 | AR-V7 | ENC1 | -0.45 | 0.71 | AR-V7 |
| LIX1L | -0.02 | 0.89 | AR-V7 | DGKH | -0.02 | 0.71 | AR-V7 |
| RASL11A | -0.08 | 0.89 | AR-V7 | WDR62 | -0.02 | 0.71 | AR-V7 |
| KIF21B | -0.34 | 0.90 | AR-V7 | SMC6 | -0.28 | 0.71 | AR-V7 |
| TSNAXIP1 | -0.09 | 0.90 | AR-V7 | LIG1 | -0.20 | 0.72 | AR-V7 |
| TAS1R3 | -0.21 | 0.90 | AR-V7 | NBN | -0.27 | 0.72 | AR-V7 |
| METRNL | -0.20 | 0.91 | AR-V7 | ITM2C | -0.16 | 0.73 | AR-V7 |
| TNFSF15 | -0.21 | 0.92 | AR-V7 | DPP6 | -0.35 | 0.74 | AR-V7 |
| TRIP6 | -0.12 | 0.93 | AR-V7 | PLEKHG4B | -0.14 | 0.74 | AR-V7 |
| RNF144B | -0.27 | 0.94 | AR-V7 | C17orf97 | -0.29 | 0.75 | AR-V7 |
| MAATS1 | -0.39 | 0.94 | AR-V7 | GLDC | -0.04 | 0.75 | AR-V7 |
| AMH | -0.41 | 0.94 | AR-V7 | IL10RA | -0.14 | 0.75 | AR-V7 |
| C17orf97 | -0.14 | 0.94 | AR-V7 | CEP152 | -0.13 | 0.76 | AR-V7 |
| DAB2IP | -0.56 | 0.94 | AR-V7 | POGLUT1 | -0.42 | 0.76 | AR-V7 |
| SGSM1 | -0.27 | 0.95 | AR-V7 | NUAK2 | -0.53 | 0.77 | AR-V7 |
| DKKL1 | -0.07 | 0.95 | AR-V7 | UNC5B | -0.33 | 0.77 | AR-V7 |
| EDA | -0.55 | 0.95 | AR-V7 | RMI2 | -0.05 | 0.78 | AR-V7 |
| SLC7A7 | -0.06 | 0.96 | AR-V7 | HSPA4L | -0.34 | 0.80 | AR-V7 |
| ARSK | -0.27 | 0.96 | AR-V7 | CYP46A1 | -0.07 | 0.80 | AR-V7 |
| RAB31 | -0.38 | 0.96 | AR-V7 | KANK1 | -0.35 | 0.80 | AR-V7 |
| MSH4 | -0.13 | 0.97 | AR-V7 | APITD1 | -0.11 | 0.81 | AR-V7 |
| FUOM | -0.54 | 0.98 | AR-V7 | CFAP69 | -0.35 | 0.83 | AR-V7 |
| MORN3 | -0.26 | 0.98 | AR-V7 | MRPS25 | -0.41 | 0.83 | AR-V7 |
| ITPR3 | -0.01 | 0.98 | AR-V7 | SDC3 | -0.33 | 0.84 | AR-V7 |
| ARHGAP28 | -0.14 | 0.99 | AR-V7 | ZNF287 | -0.09 | 0.85 | AR-V7 |
| ID1 | -0.20 | 0.99 | AR-V7 | TECTA | -0.17 | 0.86 | AR-V7 |
| APOF | -0.38 | 1.01 | AR-V7 | NETO1 | -0.20 | 0.86 | AR-V7 |
| SLC41A2 | -0.31 | 1.01 | AR-V7 | CASK | -0.25 | 0.88 | AR-V7 |
| CHST3 | -0.07 | 1.02 | AR-V7 | HELZ2 | -0.07 | 0.88 | AR-V7 |
| CDKN1A | -0.36 | 1.03 | AR-V7 | NRCAM | -0.55 | 0.89 | AR-V7 |
| FUT3 | -0.13 | 1.04 | AR-V7 | RP5-1187M17.10 | -0.47 | 0.90 | AR-V7 |
| MFGE8 | -0.18 | 1.04 | AR-V7 | CES2 | -0.48 | 0.91 | AR-V7 |
| ZSCAN18 | -0.26 | 1.04 | AR-V7 | RAP2A | -0.44 | 0.93 | AR-V7 |
| NR3C2 | -0.09 | 1.04 | AR-V7 | PARP12 | -0.40 | 0.94 | AR-V7 |
| ADAMTSL5 | -0.27 | 1.04 | AR-V7 | BRCA2 | -0.04 | 0.95 | AR-V7 |

| | | | |
|---|---|---|---|
| SPAG4 | -0.14 | 1.04 | AR-V7 |
| METTL7A | -0.22 | 1.04 | AR-V7 |
| CNTNAP1 | -0.20 | 1.04 | AR-V7 |
| GADD45A | -0.30 | 1.04 | AR-V7 |
| RCCD1 | -0.51 | 1.05 | AR-V7 |
| C12orf75 | -0.20 | 1.05 | AR-V7 |
| MLLT3 | -0.21 | 1.05 | AR-V7 |
| NHLRC1 | -0.33 | 1.07 | AR-V7 |
| GUCY1A2 | -0.39 | 1.07 | AR-V7 |
| NDUFAF5 | -0.01 | 1.08 | AR-V7 |
| IQCD | -0.32 | 1.09 | AR-V7 |
| IFIT2 | -0.37 | 1.09 | AR-V7 |
| PTPRG | -0.34 | 1.10 | AR-V7 |
| DLL1 | -0.21 | 1.10 | AR-V7 |
| JADE1 | -0.09 | 1.11 | AR-V7 |
| UVRAG | -0.19 | 1.11 | AR-V7 |
| SELO | -0.12 | 1.11 | AR-V7 |
| COL4A2 | -0.29 | 1.12 | AR-V7 |
| PAQR5 | -0.07 | 1.12 | AR-V7 |
| PARP14 | -0.26 | 1.12 | AR-V7 |
| FSIP1 | -0.34 | 1.14 | AR-V7 |
| TAF1L | -0.21 | 1.14 | AR-V7 |
| CCDC191 | -0.01 | 1.15 | AR-V7 |
| KANK3 | -0.40 | 1.15 | AR-V7 |
| NUDT8 | -0.13 | 1.16 | AR-V7 |
| TUBE1 | -0.46 | 1.16 | AR-V7 |
| MYO1F | -0.25 | 1.19 | AR-V7 |
| TNFRSF12A | -0.48 | 1.21 | AR-V7 |
| CCNJ | -0.03 | 1.22 | AR-V7 |
| LINGO4 | -0.26 | 1.23 | AR-V7 |
| PRKAR2B | -0.48 | 1.24 | AR-V7 |
| HIST1H4E | -0.02 | 1.24 | AR-V7 |
| PHLDA1 | -0.24 | 1.24 | AR-V7 |
| IGF2 | -0.27 | 1.27 | AR-V7 |
| SCNN1A | -0.41 | 1.27 | AR-V7 |
| LRRIQ1 | -0.03 | 1.31 | AR-V7 |
| FAM161B | -0.03 | 1.31 | AR-V7 |
| ATL1 | -0.27 | 1.34 | AR-V7 |
| S100A10 | -0.37 | 1.36 | AR-V7 |
| PTGFR | -0.23 | 1.37 | AR-V7 |
| FABP6 | -0.35 | 1.37 | AR-V7 |
| LGI4 | -0.29 | 1.38 | AR-V7 |
| TTC16 | -0.35 | 1.40 | AR-V7 |
| HIST2H2AA4 | -0.27 | 1.40 | AR-V7 |
| MICAL2 | -0.06 | 1.40 | AR-V7 |
| PDLIM1 | -0.13 | 1.43 | AR-V7 |
| UNC5A | -0.38 | 0.95 | AR-V7 |
| PRDM1 | -0.25 | 0.98 | AR-V7 |
| MPV17L | -0.28 | 0.99 | AR-V7 |
| ARNTL | -0.27 | 1.00 | AR-V7 |
| FAM101B | -0.34 | 1.01 | AR-V7 |
| IFT57 | -0.58 | 1.02 | AR-V7 |
| TGS1 | -0.04 | 1.03 | AR-V7 |
| MYBL1 | -0.22 | 1.03 | AR-V7 |
| ZNF84 | -0.35 | 1.05 | AR-V7 |
| PRKAR2B | -0.11 | 1.05 | AR-V7 |
| IQUB | -0.02 | 1.05 | AR-V7 |
| B4GALT6 | -0.35 | 1.06 | AR-V7 |
| ASIC3 | -0.09 | 1.06 | AR-V7 |
| CADPS2 | -0.13 | 1.06 | AR-V7 |
| ARMCX4 | -0.20 | 1.10 | AR-V7 |
| CSPG5 | -0.14 | 1.11 | AR-V7 |
| TSNAXIP1 | -0.36 | 1.12 | AR-V7 |
| MAMLD1 | -0.46 | 1.15 | AR-V7 |
| C3orf58 | -0.22 | 1.18 | AR-V7 |
| SPATA13 | -0.16 | 1.20 | AR-V7 |
| RND3 | -0.33 | 1.20 | AR-V7 |
| IER5L | -0.08 | 1.23 | AR-V7 |
| ADRA2A | -0.23 | 1.25 | AR-V7 |
| HYDIN | -0.12 | 1.26 | AR-V7 |
| PTPRG | -0.07 | 1.29 | AR-V7 |
| TACC2 | -0.08 | 1.34 | AR-V7 |
| SELO | -0.21 | 1.35 | AR-V7 |
| ZNF189 | -0.05 | 1.47 | AR-V7 |
| HAUS7 | -0.01 | 1.49 | AR-V7 |
| GCLC | -0.49 | 1.54 | AR-V7 |
| PDGFC | -0.10 | 1.56 | AR-V7 |
| TNS1 | -0.38 | 1.58 | AR-V7 |
| ABCC3 | -0.54 | 1.72 | AR-V7 |
| PPP1R1B | -0.42 | 1.75 | AR-V7 |
| CYP4F3 | -0.54 | 1.79 | AR-V7 |
| FRMD4B | -0.03 | 1.89 | AR-V7 |
| CACNA1G | -0.37 | 1.90 | AR-V7 |
| CADPS | -0.35 | 1.90 | AR-V7 |
| KCNG1 | -0.22 | 1.91 | AR-V7 |
| PHLPP2 | -0.12 | 1.93 | AR-V7 |
| AUTS2 | -0.40 | 1.94 | AR-V7 |
| DOK4 | 0.00 | 2.01 | AR-V7 |
| SGPP2 | -0.32 | 2.02 | AR-V7 |
| PMAIP1 | -0.18 | 2.16 | AR-V7 |
| SPSB1 | -0.17 | 2.19 | AR-V7 |
| PAPPA2 | -0.29 | 2.54 | AR-V7 |

| | | | |
|---|---|---|---|
| PHTF2 | -0.36 | 1.43 | AR-V7 |
| HADH | -0.16 | 1.47 | AR-V7 |
| ALDH3B2 | -0.37 | 1.49 | AR-V7 |
| PLIN2 | -0.14 | 1.51 | AR-V7 |
| RNF112 | -0.05 | 1.53 | AR-V7 |
| PCDHB8 | -0.21 | 1.55 | AR-V7 |
| MSRB3 | -0.39 | 1.55 | AR-V7 |
| MMP25 | -0.42 | 1.57 | AR-V7 |
| TSPAN12 | -0.07 | 1.61 | AR-V7 |
| P2RX2 | -0.06 | 1.62 | AR-V7 |
| DNAAF1 | -0.17 | 1.67 | AR-V7 |
| ABCC8 | -0.21 | 1.69 | AR-V7 |
| PXYLP1 | -0.30 | 1.74 | AR-V7 |
| AKNA | -0.09 | 1.75 | AR-V7 |
| TMEM74B | -0.31 | 1.80 | AR-V7 |
| IL32 | -0.48 | 1.81 | AR-V7 |
| ZNF208 | -0.06 | 1.86 | AR-V7 |
| ARL4A | -0.29 | 1.87 | AR-V7 |
| SAMD4A | -0.12 | 1.88 | AR-V7 |
| DYNC1I1 | -0.27 | 1.88 | AR-V7 |
| OPRL1 | -0.23 | 1.93 | AR-V7 |
| HBEGF | -0.01 | 1.98 | AR-V7 |
| CACNA1H | -0.31 | 1.98 | AR-V7 |
| SYCP3 | -0.44 | 2.04 | AR-V7 |
| ALCAM | -0.18 | 2.05 | AR-V7 |
| SSFA2 | -0.32 | 2.07 | AR-V7 |
| ABCA4 | -0.09 | 2.11 | AR-V7 |
| PPP1R1B | -0.35 | 2.15 | AR-V7 |
| STAB1 | -0.31 | 2.16 | AR-V7 |
| CSPG4 | -0.28 | 2.22 | AR-V7 |
| FAM229B | -0.42 | 2.30 | AR-V7 |
| HIST2H2AA3 | -0.20 | 2.31 | AR-V7 |
| HSPB8 | -0.26 | 2.33 | AR-V7 |
| PDZK1IP1 | -0.55 | 2.40 | AR-V7 |
| EFHD1 | -0.30 | 2.44 | AR-V7 |
| CYP3A5 | -0.02 | 2.56 | AR-V7 |
| ADD3 | -0.31 | 2.60 | AR-V7 |
| CNTN5 | -0.37 | 2.61 | AR-V7 |
| RDH10 | -0.45 | 2.63 | AR-V7 |
| CACNA2D4 | -0.03 | 2.79 | AR-V7 |
| CDK18 | -0.42 | 2.82 | AR-V7 |
| IRGM | -0.35 | 3.11 | AR-V7 |
| NLGN4X | -0.28 | 3.11 | AR-V7 |
| NPY1R | -0.28 | 3.15 | AR-V7 |
| PDZD2 | -0.58 | 3.43 | AR-V7 |
| PGF | -0.52 | 3.78 | AR-V7 |
| ANO1 | -0.19 | 2.57 | AR-V7 |
| ADRA2C | -0.15 | 2.63 | AR-V7 |
| GRIK4 | -0.01 | 3.62 | AR-V7 |
| AFF2 | -0.20 | 3.96 | AR-V7 |
| SERPINA5 | -0.05 | 4.52 | AR-V7 |
| SCNN1A | -0.26 | 4.71 | AR-V7 |
| MS4A2 | -0.34 | 4.73 | AR-V7 |
| SCG2 | -0.17 | 4.92 | AR-V7 |
| NRG2 | -0.53 | 4.98 | AR-V7 |
| CFAP46 | -0.06 | 5.12 | AR-V7 |
| LAPTM5 | -0.05 | 5.21 | AR-V7 |
| CD99 | -0.28 | 5.97 | AR-V7 |
| TMC2 | -0.06 | 6.32 | AR-V7 |
| NTS | -4.56 | -4.28 | Both Isoforms |
| ATF6B | -1.09 | -4.08 | Both Isoforms |
| WDR46 | -0.80 | -3.87 | Both Isoforms |
| TDGF1 | -4.03 | -3.20 | Both Isoforms |
| ABCB5 | -3.13 | -3.02 | Both Isoforms |
| TMPRSS11F | -1.53 | -2.95 | Both Isoforms |
| LRRC2 | -5.14 | -2.93 | Both Isoforms |
| SMCO3 | -3.96 | -2.80 | Both Isoforms |
| SCUBE2 | -3.31 | -2.76 | Both Isoforms |
| LMX1A | -2.62 | -2.76 | Both Isoforms |
| ADGRL3 | -9.02 | -2.75 | Both Isoforms |
| C2CD4A | -4.24 | -2.71 | Both Isoforms |
| ALS2CL | -2.50 | -2.68 | Both Isoforms |
| SALL4 | -3.05 | -2.66 | Both Isoforms |
| COL14A1 | -4.36 | -2.65 | Both Isoforms |
| ASZ1 | -7.70 | -2.64 | Both Isoforms |
| HABP2 | -3.68 | -2.59 | Both Isoforms |
| CPED1 | -4.00 | -2.58 | Both Isoforms |
| GCG | -3.81 | -2.49 | Both Isoforms |
| B3GNT6 | -4.31 | -2.47 | Both Isoforms |
| PPFIA2 | -4.43 | -2.41 | Both Isoforms |
| DCHS2 | -5.38 | -2.41 | Both Isoforms |
| INHBA | -2.40 | -2.39 | Both Isoforms |
| CTTNBP2 | -4.11 | -2.38 | Both Isoforms |
| PTPRR | -2.77 | -2.37 | Both Isoforms |
| IFIT3 | -1.85 | -2.36 | Both Isoforms |
| CTNNA3 | -4.59 | -2.34 | Both Isoforms |
| REG4 | -3.53 | -2.33 | Both Isoforms |
| FOXQ1 | -3.26 | -2.33 | Both Isoforms |
| SOX30 | -4.04 | -2.32 | Both Isoforms |
| CA8 | -3.55 | -2.32 | Both Isoforms |
| BEND5 | -4.15 | -2.31 | Both Isoforms |
| GPRIN3 | -1.91 | -2.30 | Both Isoforms |

| | | | | | | | |
|---|---|---|---|---|---|---|---|
| CRISPLD1 | -0.25 | 3.96 | AR-V7 | KRT2 | -1.03 | -2.30 | Both Isoforms |
| CLDN1 | -0.35 | 4.55 | AR-V7 | LUM | -4.94 | -2.26 | Both Isoforms |
| TRIM49 | -0.56 | 4.77 | AR-V7 | NPIPA2 | -0.85 | -2.26 | Both Isoforms |
| PAGE1 | -0.30 | 4.90 | AR-V7 | SATB1 | -4.46 | -2.25 | Both Isoforms |
| FIBCD1 | -0.06 | 5.75 | AR-V7 | CAV1 | -2.74 | -2.22 | Both Isoforms |
| TRIM49C | -0.51 | 5.87 | AR-V7 | SLC4A4 | -1.29 | -2.21 | Both Isoforms |
| SIGLEC1 | -0.11 | 6.45 | AR-V7 | ABCA1 | -3.22 | -2.20 | Both Isoforms |
| PROC | -0.54 | 7.11 | AR-V7 | CACNA2D3 | -5.03 | -2.17 | Both Isoforms |
| NTNG1 | 1.72 | -5.43 | Opposite | NAALAD2 | -4.28 | -2.16 | Both Isoforms |
| RGS16 | 0.67 | -4.49 | Opposite | TNFSF10 | -1.24 | -2.15 | Both Isoforms |
| KCNN2 | 1.63 | -3.39 | Opposite | LTF | -2.96 | -2.14 | Both Isoforms |
| COL12A1 | 2.40 | -3.27 | Opposite | NAV3 | -1.49 | -2.12 | Both Isoforms |
| TWIST1 | 2.83 | -3.26 | Opposite | CFTR | -5.31 | -2.11 | Both Isoforms |
| PARM1 | 2.02 | -2.97 | Opposite | PGM5 | -4.11 | -2.11 | Both Isoforms |
| FEV | 1.28 | -2.67 | Opposite | PTPRB | -3.50 | -2.10 | Both Isoforms |
| MYOF | 1.33 | -2.64 | Opposite | C12orf60 | -2.13 | -2.10 | Both Isoforms |
| IL1RAPL1 | 0.59 | -2.62 | Opposite | C8orf4 | -0.86 | -2.10 | Both Isoforms |
| PALMD | 0.76 | -2.53 | Opposite | IL33 | -7.33 | -2.09 | Both Isoforms |
| AIM1 | 1.06 | -2.42 | Opposite | TSPAN12 | -2.48 | -2.09 | Both Isoforms |
| SDR9C7 | 3.18 | -2.25 | Opposite | SLC7A14 | -4.29 | -2.09 | Both Isoforms |
| DUSP27 | 0.62 | -2.17 | Opposite | DLGAP2 | -3.04 | -2.08 | Both Isoforms |
| LXN | 2.13 | -2.14 | Opposite | LHX9 | -4.23 | -2.06 | Both Isoforms |
| PDE3A | 6.91 | -2.13 | Opposite | AOX1 | -2.25 | -2.05 | Both Isoforms |
| TMEM150C | 2.17 | -1.97 | Opposite | WNT2 | -6.81 | -2.04 | Both Isoforms |
| FAM65B | 1.35 | -1.96 | Opposite | APELA | -2.32 | -2.04 | Both Isoforms |
| ETV1 | 0.60 | -1.83 | Opposite | UGT2B15 | -5.32 | -2.03 | Both Isoforms |
| APOL3 | 0.69 | -1.79 | Opposite | IL1RAPL1 | -2.55 | -2.03 | Both Isoforms |
| CNTNAP2 | 1.66 | -1.79 | Opposite | FLT3 | -3.85 | -2.02 | Both Isoforms |
| BEND4 | 1.53 | -1.76 | Opposite | GLIPR1 | -3.00 | -2.02 | Both Isoforms |
| SLC22A10 | 1.25 | -1.48 | Opposite | RAPGEF4 | -1.88 | -1.99 | Both Isoforms |
| TRPM8 | 3.96 | -1.47 | Opposite | WNT5B | -2.26 | -1.99 | Both Isoforms |
| CNGB3 | 1.64 | -1.47 | Opposite | CXXC4 | -3.75 | -1.99 | Both Isoforms |
| KCNJ2 | 1.09 | -1.43 | Opposite | SNTB1 | -4.62 | -1.99 | Both Isoforms |
| IGSF5 | 1.36 | -1.34 | Opposite | PI15 | -2.81 | -1.99 | Both Isoforms |
| KREMEN2 | 0.73 | -1.31 | Opposite | IL2RG | -2.04 | -1.99 | Both Isoforms |
| MAGEA1 | 0.63 | -1.28 | Opposite | RNF125 | -2.34 | -1.98 | Both Isoforms |
| FGFR3 | 1.05 | -1.26 | Opposite | CDCA7L | -2.69 | -1.98 | Both Isoforms |
| HHIPL2 | 3.54 | -1.24 | Opposite | TNFSF15 | -4.08 | -1.96 | Both Isoforms |
| OSBPL7 | 1.06 | -1.20 | Opposite | TPK1 | -1.01 | -1.96 | Both Isoforms |
| CYP1A1 | 0.67 | -1.19 | Opposite | NIPSNAP3B | -4.41 | -1.96 | Both Isoforms |
| SLC43A1 | 1.10 | -1.19 | Opposite | PCED1B | -2.74 | -1.95 | Both Isoforms |
| RAI2 | 0.85 | -1.19 | Opposite | CLEC3A | -7.84 | -1.95 | Both Isoforms |
| DEGS1 | 0.88 | -1.18 | Opposite | THEG | -2.24 | -1.92 | Both Isoforms |
| FAT1 | 0.89 | -1.18 | Opposite | FST | -6.77 | -1.92 | Both Isoforms |
| KLHL29 | 1.33 | -1.10 | Opposite | ERP27 | -3.84 | -1.92 | Both Isoforms |
| DSC1 | 3.38 | -1.10 | Opposite | TPO | -0.87 | -1.91 | Both Isoforms |

| | | | |
|---|---|---|---|
| RANBP3L | 1.14 | -1.08 | Opposite |
| IL20RA | 1.16 | -1.06 | Opposite |
| INHBE | 0.63 | -1.06 | Opposite |
| CLDN8 | 1.27 | -1.04 | Opposite |
| SERPINE2 | 0.64 | -1.04 | Opposite |
| GCNT2 | 0.94 | -1.03 | Opposite |
| WNT5A | 1.40 | -1.03 | Opposite |
| MAGEC2 | 0.77 | -0.99 | Opposite |
| ARFGEF3 | 0.92 | -0.99 | Opposite |
| RFPL2 | 1.70 | -0.98 | Opposite |
| LCP1 | 1.70 | -0.97 | Opposite |
| FMO5 | 0.67 | -0.95 | Opposite |
| PPM1E | 0.66 | -0.95 | Opposite |
| MESP1 | 0.99 | -0.94 | Opposite |
| C9orf152 | 1.46 | -0.91 | Opposite |
| TACSTD2 | 0.84 | -0.90 | Opposite |
| GRPR | 2.94 | -0.90 | Opposite |
| WNT7B | 1.67 | -0.87 | Opposite |
| ADORA2A | 2.21 | -0.86 | Opposite |
| GNB4 | 2.08 | -0.86 | Opposite |
| GADD45G | 0.88 | -0.86 | Opposite |
| SMAP1 | 1.03 | -0.84 | Opposite |
| RBM47 | 0.62 | -0.81 | Opposite |
| RAP1GAP | 1.09 | -0.80 | Opposite |
| AQP4 | 3.34 | -0.79 | Opposite |
| SRSF12 | 1.54 | -0.77 | Opposite |
| SASH1 | 1.94 | -0.76 | Opposite |
| OAS1 | 0.82 | -0.76 | Opposite |
| RGMB | 0.82 | -0.74 | Opposite |
| CAMK2N2 | 0.96 | -0.74 | Opposite |
| MED12L | 0.61 | -0.73 | Opposite |
| SEMA3C | 1.75 | -0.73 | Opposite |
| PKP1 | 3.97 | -0.71 | Opposite |
| LRRC36 | 2.09 | -0.70 | Opposite |
| MZB1 | 1.31 | -0.70 | Opposite |
| HLA-C | 0.76 | -0.69 | Opposite |
| TJP1 | 0.83 | -0.69 | Opposite |
| PROB1 | 0.99 | -0.65 | Opposite |
| TNS3 | 2.13 | -0.65 | Opposite |
| SP9 | 3.14 | -0.63 | Opposite |
| GALNT18 | 0.60 | -0.62 | Opposite |
| CDC42BPA | 0.62 | -0.60 | Opposite |
| OPHN1 | 1.03 | -0.60 | Opposite |
| DDR2 | 0.97 | -0.58 | Opposite |
| RAPGEF5 | 0.70 | -0.58 | AR-FL |
| DEFB132 | 1.71 | -0.55 | AR-FL |
| MCC | -1.48 | -1.90 | Both Isoforms |
| LIX1 | -3.32 | -1.88 | Both Isoforms |
| WIPF1 | -1.18 | -1.88 | Both Isoforms |
| ALPL | -2.75 | -1.87 | Both Isoforms |
| MAN1A1 | -2.55 | -1.87 | Both Isoforms |
| PDE7B | -2.75 | -1.87 | Both Isoforms |
| ESRRG | -1.08 | -1.87 | Both Isoforms |
| NRIP3 | -2.46 | -1.87 | Both Isoforms |
| GIMAP2 | -2.55 | -1.86 | Both Isoforms |
| IGFBP3 | -4.48 | -1.86 | Both Isoforms |
| ARHGAP20 | -4.24 | -1.85 | Both Isoforms |
| PRKG2 | -2.00 | -1.85 | Both Isoforms |
| FREM2 | -1.93 | -1.85 | Both Isoforms |
| IPCEF1 | -2.03 | -1.84 | Both Isoforms |
| SULF1 | -4.78 | -1.84 | Both Isoforms |
| UPB1 | -2.85 | -1.83 | Both Isoforms |
| SYTL5 | -2.09 | -1.83 | Both Isoforms |
| C1QTNF1 | -2.29 | -1.82 | Both Isoforms |
| FOLH1 | -3.22 | -1.82 | Both Isoforms |
| SYTL2 | -3.72 | -1.82 | Both Isoforms |
| HS3ST3B1 | -4.61 | -1.82 | Both Isoforms |
| C8orf74 | -1.10 | -1.82 | Both Isoforms |
| DIXDC1 | -2.46 | -1.81 | Both Isoforms |
| DENND2A | -2.81 | -1.81 | Both Isoforms |
| ROS1 | -3.87 | -1.81 | Both Isoforms |
| SLC24A3 | -2.19 | -1.80 | Both Isoforms |
| EBF3 | -2.26 | -1.80 | Both Isoforms |
| CCDC83 | -3.52 | -1.80 | Both Isoforms |
| AC021106.1 | -2.74 | -1.80 | Both Isoforms |
| MAT1A | -3.27 | -1.80 | Both Isoforms |
| PEAK1 | -2.23 | -1.79 | Both Isoforms |
| BNC2 | -3.04 | -1.79 | Both Isoforms |
| UGT2B10 | -7.56 | -1.78 | Both Isoforms |
| GPR85 | -5.55 | -1.78 | Both Isoforms |
| GRM4 | -1.92 | -1.78 | Both Isoforms |
| CYBRD1 | -2.61 | -1.78 | Both Isoforms |
| SOX11 | -2.33 | -1.77 | Both Isoforms |
| LRRC31 | -1.70 | -1.77 | Both Isoforms |
| KLF5 | -1.26 | -1.77 | Both Isoforms |
| DACH1 | -3.10 | -1.76 | Both Isoforms |
| VAV3 | -3.15 | -1.76 | Both Isoforms |
| RUNDC3B | -3.31 | -1.76 | Both Isoforms |
| ADAM19 | -3.58 | -1.76 | Both Isoforms |
| LURAP1L | -2.17 | -1.75 | Both Isoforms |
| LRRIQ4 | -1.32 | -1.75 | Both Isoforms |
| EDARADD | -0.82 | -1.75 | Both Isoforms |

| | | | | | | | |
|---|---|---|---|---|---|---|---|
| LRCH1 | 2.49 | -0.55 | AR-FL | COL5A2 | -3.65 | -1.75 | Both Isoforms |
| TTC39A | 0.88 | -0.55 | AR-FL | OSBPL6 | -0.81 | -1.74 | Both Isoforms |
| CPEB2 | 0.70 | -0.53 | AR-FL | MUCL1 | -2.32 | -1.74 | Both Isoforms |
| MMEL1 | 0.93 | -0.52 | AR-FL | BMP2 | -3.84 | -1.74 | Both Isoforms |
| MIA3 | 0.82 | -0.50 | AR-FL | C11orf45 | -3.96 | -1.74 | Both Isoforms |
| ZNF385B | 2.24 | -0.50 | AR-FL | SEMA3E | -1.77 | -1.73 | Both Isoforms |
| SLC5A4 | 1.73 | -0.49 | AR-FL | SLC13A2 | -4.16 | -1.73 | Both Isoforms |
| TNK2 | 0.92 | -0.49 | AR-FL | HIST4H4 | -3.68 | -1.73 | Both Isoforms |
| NEO1 | 0.63 | -0.48 | AR-FL | TFPI | -4.47 | -1.72 | Both Isoforms |
| LSM6 | 0.59 | -0.47 | AR-FL | PDGFRB | -3.21 | -1.72 | Both Isoforms |
| JAM3 | 1.56 | -0.47 | AR-FL | NUAK1 | -1.10 | -1.71 | Both Isoforms |
| KAT2B | 0.96 | -0.47 | AR-FL | IL36RN | -2.85 | -1.70 | Both Isoforms |
| OLA1 | 0.76 | -0.47 | AR-FL | BCL11A | -2.97 | -1.70 | Both Isoforms |
| TTC39C | 0.81 | -0.46 | AR-FL | ZNF521 | -3.51 | -1.70 | Both Isoforms |
| CDC42EP2 | 0.87 | -0.45 | AR-FL | REEP1 | -2.36 | -1.69 | Both Isoforms |
| DPP4 | 1.72 | -0.44 | AR-FL | GPR149 | -3.83 | -1.69 | Both Isoforms |
| PLA2G4F | 0.68 | -0.44 | AR-FL | STXBP5L | -2.46 | -1.69 | Both Isoforms |
| HERC6 | 0.60 | -0.44 | AR-FL | PACRG | -2.73 | -1.69 | Both Isoforms |
| USP10 | 0.87 | -0.44 | AR-FL | RBP4 | -3.43 | -1.69 | Both Isoforms |
| DNASE2B | 0.78 | -0.43 | AR-FL | UGT2B17 | -9.65 | -1.68 | Both Isoforms |
| PRRT2 | 0.87 | -0.42 | AR-FL | KIAA1210 | -3.02 | -1.68 | Both Isoforms |
| LRP4 | 2.04 | -0.42 | AR-FL | SV2B | -0.65 | -1.68 | Both Isoforms |
| RNF223 | 0.85 | -0.42 | AR-FL | IHH | -1.69 | -1.68 | Both Isoforms |
| SMARCA1 | 1.23 | -0.40 | AR-FL | NAP1L3 | -3.33 | -1.67 | Both Isoforms |
| ST6GAL1 | 0.91 | -0.40 | AR-FL | CCDC80 | -3.31 | -1.67 | Both Isoforms |
| GRHL2 | 0.97 | -0.39 | AR-FL | SEMA5A | -0.75 | -1.66 | Both Isoforms |
| VSIG10L | 0.75 | -0.39 | AR-FL | NTRK2 | -2.74 | -1.65 | Both Isoforms |
| NCAPD3 | 2.52 | -0.37 | AR-FL | P2RY1 | -2.07 | -1.65 | Both Isoforms |
| GPR158 | 1.70 | -0.37 | AR-FL | STON1 | -1.17 | -1.64 | Both Isoforms |
| KCTD3 | 1.15 | -0.37 | AR-FL | BACE2 | -1.85 | -1.64 | Both Isoforms |
| CHD7 | 0.85 | -0.36 | AR-FL | C11orf85 | -2.66 | -1.64 | Both Isoforms |
| EXOC6 | 0.60 | -0.35 | AR-FL | TM4SF20 | -3.86 | -1.63 | Both Isoforms |
| SNX25 | 1.65 | -0.35 | AR-FL | POF1B | -2.13 | -1.63 | Both Isoforms |
| ALDH1A3 | 2.89 | -0.35 | AR-FL | CNGA1 | -0.92 | -1.63 | Both Isoforms |
| HID1 | 0.91 | -0.34 | AR-FL | NT5E | -2.50 | -1.62 | Both Isoforms |
| ADRBK1 | 0.67 | -0.33 | AR-FL | ANXA3 | -1.11 | -1.62 | Both Isoforms |
| MYCBP2 | 0.77 | -0.33 | AR-FL | EPHA2 | -1.25 | -1.62 | Both Isoforms |
| GNA12 | 1.04 | -0.33 | AR-FL | LEPR | -1.98 | -1.62 | Both Isoforms |
| GALK2 | 0.68 | -0.32 | AR-FL | POU2F3 | -2.39 | -1.62 | Both Isoforms |
| PAPSS1 | 0.86 | -0.32 | AR-FL | KLRG1 | -0.96 | -1.61 | Both Isoforms |
| TEX2 | 1.21 | -0.32 | AR-FL | TLR1 | -4.15 | -1.61 | Both Isoforms |
| NPDC1 | 0.78 | -0.32 | AR-FL | NEGR1 | -3.55 | -1.60 | Both Isoforms |
| NSMAF | 1.16 | -0.32 | AR-FL | FZD10 | -2.14 | -1.60 | Both Isoforms |
| TRIM33 | 1.00 | -0.31 | AR-FL | MET | -4.25 | -1.60 | Both Isoforms |
| SPECC1L | 1.40 | -0.30 | AR-FL | AKR1C3 | -2.96 | -1.60 | Both Isoforms |
| HMG20B | 0.66 | -0.30 | AR-FL | FAM110C | -1.36 | -1.60 | Both Isoforms |

| | | | |
|---|---|---|---|
| ZNF608 | 1.16 | -0.30 | AR-FL |
| DOCK5 | 0.80 | -0.30 | AR-FL |
| CNDP2 | 0.83 | -0.29 | AR-FL |
| KIF22 | 0.75 | -0.29 | AR-FL |
| FNIP2 | 0.76 | -0.29 | AR-FL |
| RAB3D | 0.79 | -0.29 | AR-FL |
| GCFC2 | 0.88 | -0.29 | AR-FL |
| SLC2A4 | 1.21 | -0.27 | AR-FL |
| SYTL1 | 0.67 | -0.26 | AR-FL |
| MAP7 | 0.77 | -0.26 | AR-FL |
| IQGAP2 | 2.39 | -0.26 | AR-FL |
| VPS26B | 1.47 | -0.25 | AR-FL |
| SYNGR2 | 0.83 | -0.24 | AR-FL |
| FHIT | 1.37 | -0.24 | AR-FL |
| EMC2 | 0.75 | -0.24 | AR-FL |
| EPS15 | 1.01 | -0.24 | AR-FL |
| RUNX1 | 3.32 | -0.23 | AR-FL |
| COL4A3BP | 1.24 | -0.23 | AR-FL |
| ELOVL2 | 3.87 | -0.23 | AR-FL |
| ARFIP1 | 0.65 | -0.23 | AR-FL |
| RCBTB1 | 0.75 | -0.23 | AR-FL |
| ATP10A | 1.10 | -0.22 | AR-FL |
| TDRD1 | 1.10 | -0.22 | AR-FL |
| PSD3 | 1.17 | -0.21 | AR-FL |
| INTS10 | 1.21 | -0.21 | AR-FL |
| ABHD17B | 0.73 | -0.21 | AR-FL |
| ATP8B1 | 0.66 | -0.21 | AR-FL |
| NHSL2 | 0.68 | -0.21 | AR-FL |
| LRRC16A | 1.88 | -0.20 | AR-FL |
| TRPM2 | 0.73 | -0.20 | AR-FL |
| ALDH4A1 | 0.90 | -0.19 | AR-FL |
| GRHL1 | 0.64 | -0.19 | AR-FL |
| PLXNB3 | 0.63 | -0.19 | AR-FL |
| CHST11 | 0.66 | -0.18 | AR-FL |
| GPRC5B | 1.00 | -0.17 | AR-FL |
| SOAT1 | 1.44 | -0.17 | AR-FL |
| SGK1 | 3.25 | -0.17 | AR-FL |
| CARTPT | 0.82 | -0.16 | AR-FL |
| NUP58 | 0.76 | -0.16 | AR-FL |
| CITED1 | 1.19 | -0.16 | AR-FL |
| ENDOD1 | 1.85 | -0.16 | AR-FL |
| PCDH1 | 1.62 | -0.16 | AR-FL |
| OAZ3 | 0.58 | -0.16 | AR-FL |
| COPRS | 0.61 | -0.16 | AR-FL |
| GNAQ | 1.03 | -0.15 | AR-FL |
| ALPK3 | 1.33 | -0.15 | AR-FL |
| LYPD6B | -2.05 | -1.59 | Both Isoforms |
| CSRP2 | -2.15 | -1.59 | Both Isoforms |
| FBN1 | -2.22 | -1.58 | Both Isoforms |
| MGP | -2.64 | -1.58 | Both Isoforms |
| CACNB2 | -2.42 | -1.58 | Both Isoforms |
| B3GALT5 | -2.76 | -1.58 | Both Isoforms |
| SHH | -4.69 | -1.58 | Both Isoforms |
| AMIGO2 | -3.70 | -1.58 | Both Isoforms |
| SERPINI1 | -2.37 | -1.57 | Both Isoforms |
| SAMD13 | -1.08 | -1.57 | Both Isoforms |
| FAM19A2 | -1.87 | -1.56 | Both Isoforms |
| TXNRD2 | -1.74 | -1.56 | Both Isoforms |
| SLC30A10 | -3.02 | -1.56 | Both Isoforms |
| EDNRA | -2.56 | -1.56 | Both Isoforms |
| ZNF711 | -3.30 | -1.55 | Both Isoforms |
| APOL4 | -0.81 | -1.55 | Both Isoforms |
| NEB | -2.90 | -1.55 | Both Isoforms |
| UGT2B7 | -3.56 | -1.54 | Both Isoforms |
| FAM171B | -2.37 | -1.53 | Both Isoforms |
| TTLL2 | -2.76 | -1.53 | Both Isoforms |
| ZNF624 | -1.07 | -1.53 | Both Isoforms |
| SYNPO | -2.99 | -1.53 | Both Isoforms |
| SSTR1 | -1.85 | -1.53 | Both Isoforms |
| SYT4 | -0.60 | -1.52 | Both Isoforms |
| GTF2IRD2B | -1.42 | -1.51 | Both Isoforms |
| EPB41L4A | -2.96 | -1.51 | Both Isoforms |
| COL8A2 | -2.54 | -1.51 | Both Isoforms |
| PTGER2 | -3.02 | -1.51 | Both Isoforms |
| MDGA1 | -2.15 | -1.51 | Both Isoforms |
| SORL1 | -1.61 | -1.51 | Both Isoforms |
| TMEM176B | -2.86 | -1.50 | Both Isoforms |
| CDH26 | -1.81 | -1.50 | Both Isoforms |
| HMGCS2 | -0.77 | -1.50 | Both Isoforms |
| ZNF536 | -3.28 | -1.49 | Both Isoforms |
| ATP8B4 | -0.80 | -1.48 | Both Isoforms |
| ADCY2 | -0.96 | -1.48 | Both Isoforms |
| PLAGL1 | -0.66 | -1.48 | Both Isoforms |
| TRIB1 | -3.04 | -1.47 | Both Isoforms |
| SNX10 | -2.12 | -1.47 | Both Isoforms |
| CD200 | -5.55 | -1.47 | Both Isoforms |
| UMODL1 | -0.81 | -1.47 | Both Isoforms |
| MAPK4 | -0.92 | -1.46 | Both Isoforms |
| FTCD | -2.02 | -1.46 | Both Isoforms |
| PAX1 | -2.32 | -1.46 | Both Isoforms |
| NOV | -4.29 | -1.46 | Both Isoforms |
| TAL2 | -2.16 | -1.46 | Both Isoforms |

| | | | |
|---|---|---|---|
| L3MBTL3 | 0.85 | -0.15 | AR-FL |
| RBM45 | 0.58 | -0.14 | AR-FL |
| PASK | 0.78 | -0.14 | AR-FL |
| ACTR3 | 0.84 | -0.14 | AR-FL |
| ANTXR2 | 0.58 | -0.14 | AR-FL |
| SOD3 | 0.71 | -0.13 | AR-FL |
| BHLHA15 | 2.05 | -0.12 | AR-FL |
| PITHD1 | 0.62 | -0.12 | AR-FL |
| AP2S1 | 0.68 | -0.12 | AR-FL |
| CARD10 | 0.63 | -0.11 | AR-FL |
| AF131216.1 | 1.31 | -0.11 | AR-FL |
| OR5B2 | 5.20 | -0.11 | AR-FL |
| RHPN2 | 0.62 | -0.10 | AR-FL |
| HIST2H4B | 0.60 | -0.10 | AR-FL |
| SEC63 | 0.63 | -0.09 | AR-FL |
| OTUD7B | 1.43 | -0.09 | AR-FL |
| C2orf72 | 0.83 | -0.09 | AR-FL |
| C6orf132 | 0.72 | -0.09 | AR-FL |
| MTFP1 | 1.22 | -0.08 | AR-FL |
| ZDHHC21 | 0.59 | -0.08 | AR-FL |
| SPATC1L | 0.76 | -0.07 | AR-FL |
| BBS4 | 0.95 | -0.07 | AR-FL |
| UHRF2 | 0.63 | -0.07 | AR-FL |
| UAP1 | 2.12 | -0.07 | AR-FL |
| PRKAR2A | 0.58 | -0.06 | AR-FL |
| GSR | 1.04 | -0.06 | AR-FL |
| ATP6V1C1 | 0.61 | -0.06 | AR-FL |
| NUP98 | 0.67 | -0.06 | AR-FL |
| WWC2 | 1.09 | -0.06 | AR-FL |
| TEP1 | 0.67 | -0.06 | AR-FL |
| CD9 | 0.77 | -0.05 | AR-FL |
| SEMA4A | 0.60 | -0.05 | AR-FL |
| SNX16 | 0.99 | -0.05 | AR-FL |
| SPATA24 | 0.65 | -0.05 | AR-FL |
| WWP1 | 0.71 | -0.05 | AR-FL |
| RUSC1 | 0.67 | -0.05 | AR-FL |
| TECPR1 | 1.14 | -0.05 | AR-FL |
| PCMT1 | 0.76 | -0.04 | AR-FL |
| IL1R1 | 0.85 | -0.04 | AR-FL |
| CSK | 0.61 | -0.04 | AR-FL |
| EPS8L1 | 1.78 | -0.04 | AR-FL |
| SERINC2 | 1.07 | -0.04 | AR-FL |
| UBE2D1 | 0.66 | -0.04 | AR-FL |
| UNC5C | 0.59 | -0.03 | AR-FL |
| GSPT1 | 0.63 | -0.03 | AR-FL |
| ZNF615 | 0.76 | -0.03 | AR-FL |

| | | | |
|---|---|---|---|
| CDH17 | -2.72 | -1.45 | Both Isoforms |
| SYT2 | -1.62 | -1.45 | Both Isoforms |
| LOXL4 | -3.20 | -1.45 | Both Isoforms |
| ADAMTSL2 | -1.89 | -1.45 | Both Isoforms |
| PITPNC1 | -1.84 | -1.45 | Both Isoforms |
| ST7 | -2.52 | -1.44 | Both Isoforms |
| BMX | -4.20 | -1.44 | Both Isoforms |
| A1CF | -2.53 | -1.44 | Both Isoforms |
| P2RX7 | -3.70 | -1.44 | Both Isoforms |
| HGF | -3.77 | -1.44 | Both Isoforms |
| RGS17 | -1.28 | -1.43 | Both Isoforms |
| PGR | -2.65 | -1.43 | Both Isoforms |
| FGL2 | -3.67 | -1.43 | Both Isoforms |
| PLEKHG7 | -2.95 | -1.42 | Both Isoforms |
| PHEX | -1.54 | -1.42 | Both Isoforms |
| CDH10 | -3.80 | -1.42 | Both Isoforms |
| KCNB1 | -2.39 | -1.42 | Both Isoforms |
| ZYG11A | -1.33 | -1.42 | Both Isoforms |
| TCP11L2 | -1.64 | -1.42 | Both Isoforms |
| LRRC9 | -1.30 | -1.41 | Both Isoforms |
| PIK3IP1 | -2.33 | -1.41 | Both Isoforms |
| SPTLC3 | -1.92 | -1.41 | Both Isoforms |
| CLDN11 | -3.29 | -1.40 | Both Isoforms |
| BMF | -0.61 | -1.40 | Both Isoforms |
| IGF2BP2 | -0.77 | -1.40 | Both Isoforms |
| LPAR1 | -0.98 | -1.40 | Both Isoforms |
| CD8A | -2.68 | -1.39 | Both Isoforms |
| PKNOX2 | -2.37 | -1.39 | Both Isoforms |
| GLI3 | -2.34 | -1.39 | Both Isoforms |
| SLC13A3 | -1.77 | -1.39 | Both Isoforms |
| BCAT1 | -2.91 | -1.39 | Both Isoforms |
| ZIC2 | -3.78 | -1.39 | Both Isoforms |
| CAPRIN2 | -1.86 | -1.39 | Both Isoforms |
| NR1D2 | -1.79 | -1.39 | Both Isoforms |
| C5 | -2.42 | -1.38 | Both Isoforms |
| PDE1B | -2.44 | -1.38 | Both Isoforms |
| ANKRD66 | -2.49 | -1.38 | Both Isoforms |
| KLF7 | -2.09 | -1.37 | Both Isoforms |
| FAM89A | -1.63 | -1.37 | Both Isoforms |
| PLPPR4 | -2.25 | -1.37 | Both Isoforms |
| RHCE | -1.29 | -1.37 | Both Isoforms |
| FZD7 | -1.86 | -1.36 | Both Isoforms |
| CLIC5 | -3.07 | -1.36 | Both Isoforms |
| KLHL24 | -1.35 | -1.36 | Both Isoforms |
| CDON | -1.26 | -1.36 | Both Isoforms |
| TIAM2 | -1.35 | -1.36 | Both Isoforms |

| | | | | | | | |
|---|---|---|---|---|---|---|---|
| NFKBIE | 0.58 | -0.02 | AR-FL | SKIDA1 | -0.91 | -1.36 | Both Isoforms |
| SLAIN2 | 1.00 | -0.02 | AR-FL | TMEM266 | -2.54 | -1.36 | Both Isoforms |
| TADA3 | 0.61 | -0.02 | AR-FL | KIAA1211 | -0.86 | -1.36 | Both Isoforms |
| IPPK | 0.64 | -0.02 | AR-FL | LAMB4 | -2.28 | -1.35 | Both Isoforms |
| RAB6C | 0.68 | -0.01 | AR-FL | C20orf195 | -2.15 | -1.35 | Both Isoforms |
| MAZ | 0.63 | -0.01 | AR-FL | NR3C1 | -2.04 | -1.35 | Both Isoforms |
| HEBP2 | 1.24 | -0.01 | AR-FL | OPTN | -1.99 | -1.35 | Both Isoforms |
| SPATS2L | 0.66 | -0.01 | AR-FL | SYN2 | -2.46 | -1.34 | Both Isoforms |
| TIMM23 | 0.64 | -0.01 | AR-FL | EVX1 | -2.90 | -1.34 | Both Isoforms |
| KIAA0355 | 0.86 | -0.01 | AR-FL | NATD1 | -2.14 | -1.34 | Both Isoforms |
| EML1 | 1.25 | 0.00 | AR-FL | ADH1B | -4.18 | -1.34 | Both Isoforms |
| RYBP | 0.65 | 0.00 | AR-FL | MEP1A | -7.48 | -1.34 | Both Isoforms |
| KCNG3 | 1.74 | 0.00 | AR-FL | KCTD16 | -2.84 | -1.34 | Both Isoforms |
| POLR3E | 0.91 | 0.00 | AR-FL | CDC14A | -2.81 | -1.33 | Both Isoforms |
| NRG3 | -2.61 | -6.90 | Both Isoforms | SLC22A11 | -0.98 | -1.33 | Both Isoforms |
| MBNL2 | -0.91 | -6.23 | Both Isoforms | AKR1C1 | -1.85 | -1.33 | Both Isoforms |
| LRRN1 | -3.48 | -6.20 | Both Isoforms | SH3GL3 | -2.90 | -1.33 | Both Isoforms |
| DAPK1 | -5.53 | -6.00 | Both Isoforms | MMP10 | -1.82 | -1.32 | Both Isoforms |
| CSRNP3 | -1.81 | -5.40 | Both Isoforms | GPC5 | -1.47 | -1.32 | Both Isoforms |
| VARS2 | -3.00 | -4.88 | Both Isoforms | SH3BP5 | -1.40 | -1.32 | Both Isoforms |
| EYA2 | -1.05 | -4.71 | Both Isoforms | MR1 | -1.68 | -1.32 | Both Isoforms |
| PRSS1 | -1.14 | -4.70 | Both Isoforms | CCDC158 | -2.68 | -1.32 | Both Isoforms |
| FRAS1 | -2.26 | -4.47 | Both Isoforms | POU5F1B | -1.33 | -1.31 | Both Isoforms |
| GRIK1 | -2.10 | -4.46 | Both Isoforms | LRRTM3 | -6.20 | -1.31 | Both Isoforms |
| OR51E1 | -3.00 | -4.40 | Both Isoforms | CDK18 | -2.82 | -1.31 | Both Isoforms |
| ST6GALNAC5 | -2.74 | -4.39 | Both Isoforms | CD44 | -0.66 | -1.31 | Both Isoforms |
| SALL4 | -4.77 | -4.21 | Both Isoforms | LRRC7 | -2.17 | -1.31 | Both Isoforms |
| ADAM2 | -1.60 | -4.19 | Both Isoforms | TSPAN8 | -0.97 | -1.31 | Both Isoforms |
| IHH | -3.51 | -4.13 | Both Isoforms | CCDC67 | -1.78 | -1.31 | Both Isoforms |
| FAM189A1 | -1.73 | -3.92 | Both Isoforms | CDH3 | -2.23 | -1.31 | Both Isoforms |
| ABCA12 | -2.88 | -3.78 | Both Isoforms | TMEM159 | -1.64 | -1.31 | Both Isoforms |

| | | | | | | | |
|---|---|---|---|---|---|---|---|
| ARHGDIG | -0.59 | -3.76 | Both Isoforms | AMN | -3.16 | -1.31 | Both Isoforms |
| RLN2 | -1.12 | -3.67 | Both Isoforms | ADGRD1 | -2.63 | -1.30 | Both Isoforms |
| NAV3 | -1.94 | -3.65 | Both Isoforms | SYT13 | -1.20 | -1.30 | Both Isoforms |
| PLA2G2F | -5.19 | -3.61 | Both Isoforms | LANCL3 | -1.85 | -1.30 | Both Isoforms |
| COL3A1 | -3.39 | -3.61 | Both Isoforms | PPP1R14C | -3.42 | -1.30 | Both Isoforms |
| COL5A2 | -2.88 | -3.60 | Both Isoforms | PPP2R5A | -1.75 | -1.30 | Both Isoforms |
| LTF | -2.49 | -3.50 | Both Isoforms | ADGRG6 | -3.90 | -1.30 | Both Isoforms |
| OR51E2 | -0.82 | -3.49 | Both Isoforms | FAM184A | -2.87 | -1.30 | Both Isoforms |
| OPRK1 | -4.43 | -3.45 | Both Isoforms | GATA5 | -1.46 | -1.30 | Both Isoforms |
| SLC6A20 | -5.80 | -3.41 | Both Isoforms | ACKR3 | -2.01 | -1.30 | Both Isoforms |
| ACPP | -0.61 | -3.37 | Both Isoforms | TMEM200A | -5.49 | -1.30 | Both Isoforms |
| CCDC83 | -4.81 | -3.17 | Both Isoforms | LTBP1 | -2.95 | -1.30 | Both Isoforms |
| SLC5A1 | -1.88 | -3.15 | Both Isoforms | CDH12 | -1.93 | -1.30 | Both Isoforms |
| CERK | -2.81 | -3.09 | Both Isoforms | KIF26B | -1.15 | -1.30 | Both Isoforms |
| DLX3 | -3.16 | -3.02 | Both Isoforms | AFP | -0.82 | -1.30 | Both Isoforms |
| AMIGO2 | -5.06 | -2.97 | Both Isoforms | SLC4A10 | -3.15 | -1.29 | Both Isoforms |
| SCN8A | -1.15 | -2.96 | Both Isoforms | P4HA3 | -1.87 | -1.29 | Both Isoforms |
| SIRPA | -0.92 | -2.95 | Both Isoforms | PTGFR | -0.92 | -1.29 | Both Isoforms |
| GALNT14 | -2.50 | -2.93 | Both Isoforms | CNIH3 | -0.71 | -1.29 | Both Isoforms |
| ODAM | -1.56 | -2.86 | Both Isoforms | RSAD2 | -1.59 | -1.29 | Both Isoforms |
| TLL1 | -2.79 | -2.86 | Both Isoforms | NCAM2 | -3.22 | -1.29 | Both Isoforms |
| IL1RN | -3.54 | -2.85 | Both Isoforms | SLC3A1 | -2.85 | -1.29 | Both Isoforms |
| CXorf36 | -5.63 | -2.84 | Both Isoforms | BNC1 | -2.49 | -1.29 | Both Isoforms |
| CD274 | -1.50 | -2.83 | Both Isoforms | PLCXD3 | -2.58 | -1.29 | Both Isoforms |
| BCL11B | -1.20 | -2.80 | Both Isoforms | CSMD2 | -1.08 | -1.28 | Both Isoforms |

| | | | | | | | |
|---|---|---|---|---|---|---|---|
| KIFC1 | -2.52 | -2.77 | Both Isoforms | CHST11 | -1.31 | -1.28 | Both Isoforms |
| VAV3 | -3.01 | -2.76 | Both Isoforms | KLHDC8A | -5.81 | -1.28 | Both Isoforms |
| ACSS1 | -1.46 | -2.74 | Both Isoforms | FRAS1 | -2.51 | -1.28 | Both Isoforms |
| ELF5 | -0.60 | -2.70 | Both Isoforms | GATA6 | -0.89 | -1.28 | Both Isoforms |
| PHF1 | -2.00 | -2.69 | Both Isoforms | SLC10A5 | -1.72 | -1.28 | Both Isoforms |
| MYRIP | -1.82 | -2.67 | Both Isoforms | MUC13 | -0.95 | -1.28 | Both Isoforms |
| PLD1 | -4.56 | -2.66 | Both Isoforms | BLNK | -1.02 | -1.27 | Both Isoforms |
| TMC7 | -0.85 | -2.64 | Both Isoforms | TMEM45B | -1.19 | -1.27 | Both Isoforms |
| RIMS4 | -2.13 | -2.64 | Both Isoforms | IL16 | -0.67 | -1.27 | Both Isoforms |
| TSHZ3 | -1.90 | -2.63 | Both Isoforms | ACSL5 | -0.80 | -1.27 | Both Isoforms |
| GBP3 | -0.65 | -2.63 | Both Isoforms | KCNK13 | -1.81 | -1.27 | Both Isoforms |
| LRRC31 | -4.53 | -2.63 | Both Isoforms | PRTG | -1.86 | -1.27 | Both Isoforms |
| LYPD6B | -2.40 | -2.63 | Both Isoforms | CSRNP3 | -2.11 | -1.27 | Both Isoforms |
| DPYSL3 | -0.82 | -2.61 | Both Isoforms | ANO2 | -1.93 | -1.27 | Both Isoforms |
| SUSD4 | -2.01 | -2.60 | Both Isoforms | SLC9A9 | -1.15 | -1.27 | Both Isoforms |
| CAMK2N1 | -4.21 | -2.60 | Both Isoforms | CDH7 | -1.06 | -1.27 | Both Isoforms |
| TMEM179 | -1.15 | -2.58 | Both Isoforms | CECR2 | -1.69 | -1.27 | Both Isoforms |
| EIF3CL | -1.69 | -2.57 | Both Isoforms | FAM175A | -2.74 | -1.26 | Both Isoforms |
| CACHD1 | -2.36 | -2.54 | Both Isoforms | SOX5 | -2.17 | -1.26 | Both Isoforms |
| LRRC4C | -2.41 | -2.51 | Both Isoforms | MYB | -2.32 | -1.26 | Both Isoforms |
| VWF | -1.03 | -2.51 | Both Isoforms | PLA2G2A | -3.26 | -1.26 | Both Isoforms |
| PCDH11Y | -3.43 | -2.50 | Both Isoforms | TRERF1 | -1.81 | -1.26 | Both Isoforms |
| AIDA | -1.59 | -2.49 | Both Isoforms | ZNF385B | -2.18 | -1.26 | Both Isoforms |
| ABCG1 | -0.98 | -2.48 | Both Isoforms | CEMIP | -1.57 | -1.26 | Both Isoforms |
| PLA2G2A | -3.71 | -2.48 | Both Isoforms | BRINP1 | -1.38 | -1.26 | Both Isoforms |

| | | | | | | | |
|---|---|---|---|---|---|---|---|
| EFHC2 | -1.95 | -2.48 | Both Isoforms | AKAP3 | -2.66 | -1.25 | Both Isoforms |
| RELN | -1.13 | -2.43 | Both Isoforms | TBX6 | -1.90 | -1.25 | Both Isoforms |
| FN1 | -0.92 | -2.42 | Both Isoforms | ADAMTSL3 | -2.98 | -1.25 | Both Isoforms |
| IL36RN | -1.17 | -2.40 | Both Isoforms | MID1 | -1.73 | -1.25 | Both Isoforms |
| MYB | -2.65 | -2.39 | Both Isoforms | MEF2C | -2.60 | -1.25 | Both Isoforms |
| DDC | -2.20 | -2.39 | Both Isoforms | HEPACAM2 | -1.25 | -1.25 | Both Isoforms |
| PCED1B | -3.99 | -2.37 | Both Isoforms | CMBL | -2.45 | -1.25 | Both Isoforms |
| LYPD6 | -1.06 | -2.36 | Both Isoforms | CDK5R1 | -1.17 | -1.25 | Both Isoforms |
| TTC9 | -0.84 | -2.35 | Both Isoforms | GPR146 | -0.81 | -1.25 | Both Isoforms |
| MMP16 | -1.83 | -2.33 | Both Isoforms | SCGB1D2 | -1.67 | -1.25 | Both Isoforms |
| SLC7A5 | -2.61 | -2.31 | Both Isoforms | PPARG | -1.18 | -1.25 | Both Isoforms |
| DCBLD2 | -1.20 | -2.30 | Both Isoforms | ITGA9 | -3.76 | -1.25 | Both Isoforms |
| LRRC8C | -1.08 | -2.30 | Both Isoforms | ALX4 | -2.96 | -1.24 | Both Isoforms |
| REEP1 | -3.16 | -2.29 | Both Isoforms | DAPK1 | -1.61 | -1.24 | Both Isoforms |
| GLP1R | -1.14 | -2.27 | Both Isoforms | CD83 | -2.19 | -1.24 | Both Isoforms |
| CDH3 | -2.67 | -2.23 | Both Isoforms | FCGBP | -0.61 | -1.23 | Both Isoforms |
| RTN4RL1 | -1.83 | -2.23 | Both Isoforms | SLC6A4 | -1.58 | -1.23 | Both Isoforms |
| FFAR2 | -3.17 | -2.19 | Both Isoforms | SLC30A4 | -1.12 | -1.23 | Both Isoforms |
| CAMK2B | -1.43 | -2.19 | Both Isoforms | KIAA0922 | -1.98 | -1.23 | Both Isoforms |
| MAPK4 | -1.34 | -2.19 | Both Isoforms | SUPT3H | -1.86 | -1.23 | Both Isoforms |
| KCNK5 | -1.37 | -2.14 | Both Isoforms | ALB | -3.52 | -1.23 | Both Isoforms |
| KIAA1211L | -1.43 | -2.12 | Both Isoforms | CD109 | -2.35 | -1.23 | Both Isoforms |
| SMCO3 | -1.14 | -2.11 | Both Isoforms | DUSP5 | -1.01 | -1.23 | Both Isoforms |
| HOXB9 | -0.67 | -2.10 | Both Isoforms | NOVA1 | -2.86 | -1.22 | Both Isoforms |
| ETNPPL | -1.97 | -2.09 | Both Isoforms | HRH1 | -1.05 | -1.22 | Both Isoforms |

| | | | | | | | |
|---|---|---|---|---|---|---|---|
| PLCH1 | -2.11 | -2.09 | Both Isoforms | SRGAP3 | -1.15 | -1.22 | Both Isoforms |
| P2RX7 | -1.06 | -2.09 | Both Isoforms | FAM101A | -1.57 | -1.22 | Both Isoforms |
| PTPRK | -1.14 | -2.08 | Both Isoforms | WNK4 | -2.89 | -1.22 | Both Isoforms |
| MATN2 | -4.39 | -2.06 | Both Isoforms | CCT6B | -1.31 | -1.22 | Both Isoforms |
| TULP4 | -1.58 | -2.04 | Both Isoforms | GBGT1 | -1.85 | -1.22 | Both Isoforms |
| IGFBP3 | -0.61 | -2.03 | Both Isoforms | SCN4B | -2.34 | -1.21 | Both Isoforms |
| FLT4 | -2.10 | -2.03 | Both Isoforms | NDUFA4L2 | -1.69 | -1.21 | Both Isoforms |
| MANEA | -2.05 | -2.01 | Both Isoforms | POU6F1 | -2.14 | -1.21 | Both Isoforms |
| TOX3 | -0.98 | -2.01 | Both Isoforms | FAM47E | -1.76 | -1.21 | Both Isoforms |
| C1orf95 | -2.12 | -1.99 | Both Isoforms | NPNT | -2.10 | -1.21 | Both Isoforms |
| OLFM2 | -1.01 | -1.98 | Both Isoforms | C4orf32 | -0.93 | -1.21 | Both Isoforms |
| CDK8 | -1.78 | -1.97 | Both Isoforms | CAPN13 | -2.41 | -1.21 | Both Isoforms |
| CUTA | -2.15 | -1.97 | Both Isoforms | NFATC4 | -1.45 | -1.21 | Both Isoforms |
| TMOD1 | -1.12 | -1.96 | Both Isoforms | IER5 | -2.50 | -1.21 | Both Isoforms |
| NEB | -2.03 | -1.96 | Both Isoforms | SALL2 | -2.04 | -1.21 | Both Isoforms |
| LRRC7 | -1.94 | -1.96 | Both Isoforms | LRRN1 | -3.26 | -1.21 | Both Isoforms |
| SCD5 | -0.73 | -1.96 | Both Isoforms | ACRBP | -2.96 | -1.21 | Both Isoforms |
| TGM6 | -1.41 | -1.94 | Both Isoforms | APLF | -2.38 | -1.20 | Both Isoforms |
| LGI2 | -2.20 | -1.93 | Both Isoforms | RP11-166B2.1 | -2.01 | -1.20 | Both Isoforms |
| DNAJC6 | -0.71 | -1.87 | Both Isoforms | CHRNA7 | -1.72 | -1.20 | Both Isoforms |
| FAM198A | -2.21 | -1.87 | Both Isoforms | PCOLCE2 | -2.21 | -1.20 | Both Isoforms |
| BARD1 | -2.09 | -1.86 | Both Isoforms | GNG2 | -2.59 | -1.20 | Both Isoforms |
| ABHD1 | -0.72 | -1.86 | Both Isoforms | MFSD4 | -2.20 | -1.20 | Both Isoforms |
| TARSL2 | -1.12 | -1.85 | Both Isoforms | FYN | -3.05 | -1.19 | Both Isoforms |
| ZSWIM5 | -1.60 | -1.85 | Both Isoforms | POLR2M | -2.20 | -1.19 | Both Isoforms |

| | | | | | | | |
|---|---|---|---|---|---|---|---|
| TMEM163 | -1.76 | -1.84 | Both Isoforms | EIF3C | -1.25 | -1.19 | Both Isoforms |
| POLR3G | -1.51 | -1.84 | Both Isoforms | KCNJ3 | -3.59 | -1.19 | Both Isoforms |
| TGFA | -1.42 | -1.84 | Both Isoforms | GAB1 | -0.98 | -1.19 | Both Isoforms |
| ANTXR1 | -3.16 | -1.83 | Both Isoforms | VILL | -1.18 | -1.19 | Both Isoforms |
| LUZP2 | -1.43 | -1.83 | Both Isoforms | SLC7A5 | -1.76 | -1.19 | Both Isoforms |
| AKR1C2 | -0.86 | -1.83 | Both Isoforms | ANXA10 | -1.21 | -1.19 | Both Isoforms |
| CMPK2 | -1.24 | -1.82 | Both Isoforms | SLC24A2 | -1.88 | -1.19 | Both Isoforms |
| SESN1 | -0.92 | -1.81 | Both Isoforms | STK32A | -2.76 | -1.18 | Both Isoforms |
| PCDH11X | -4.12 | -1.80 | Both Isoforms | SPRY1 | -1.15 | -1.18 | Both Isoforms |
| BCL2 | -1.14 | -1.80 | Both Isoforms | TMEM220 | -1.19 | -1.18 | Both Isoforms |
| ARHGAP22 | -1.75 | -1.79 | Both Isoforms | GLS2 | -1.51 | -1.18 | Both Isoforms |
| CA12 | -0.88 | -1.79 | Both Isoforms | ZSCAN31 | -0.78 | -1.18 | Both Isoforms |
| SLIT1 | -2.08 | -1.79 | Both Isoforms | CMPK2 | -1.12 | -1.18 | Both Isoforms |
| DEPTOR | -1.38 | -1.78 | Both Isoforms | TRPC1 | -1.66 | -1.18 | Both Isoforms |
| HTRA1 | -1.85 | -1.77 | Both Isoforms | RBFOX3 | -3.25 | -1.18 | Both Isoforms |
| IL27RA | -3.53 | -1.77 | Both Isoforms | FOXN3 | -0.82 | -1.18 | Both Isoforms |
| ADCY1 | -1.74 | -1.76 | Both Isoforms | PRKD1 | -1.39 | -1.18 | Both Isoforms |
| CHAC1 | -1.40 | -1.76 | Both Isoforms | YPEL1 | -1.61 | -1.17 | Both Isoforms |
| FAM184B | -1.27 | -1.76 | Both Isoforms | TSHZ1 | -1.75 | -1.17 | Both Isoforms |
| GPR63 | -1.72 | -1.75 | Both Isoforms | DPY19L2 | -2.70 | -1.17 | Both Isoforms |
| SLITRK3 | -3.64 | -1.74 | Both Isoforms | FGL1 | -0.74 | -1.17 | Both Isoforms |
| PAK3 | -0.67 | -1.73 | Both Isoforms | COL16A1 | -2.67 | -1.17 | Both Isoforms |
| BEST1 | -2.20 | -1.73 | Both Isoforms | TNNT1 | -1.25 | -1.17 | Both Isoforms |
| ARHGAP9 | -1.10 | -1.73 | Both Isoforms | FRRS1L | -2.78 | -1.17 | Both Isoforms |
| C8orf34 | -0.59 | -1.72 | Both Isoforms | NELL2 | -1.27 | -1.17 | Both Isoforms |

| | | | | | | | |
|---|---|---|---|---|---|---|---|
| JAKMIP1 | -1.24 | -1.72 | Both Isoforms | TMPRSS4 | -0.59 | -1.17 | Both Isoforms |
| CHRD | -1.46 | -1.71 | Both Isoforms | GDA | -0.58 | -1.17 | Both Isoforms |
| NOV | -3.78 | -1.71 | Both Isoforms | NIPAL2 | -0.67 | -1.16 | Both Isoforms |
| FOXO6 | -1.89 | -1.71 | Both Isoforms | FAM189A1 | -0.72 | -1.16 | Both Isoforms |
| SLC6A9 | -0.87 | -1.71 | Both Isoforms | CHST15 | -1.46 | -1.16 | Both Isoforms |
| TMEM158 | -1.29 | -1.69 | Both Isoforms | BEND7 | -1.95 | -1.16 | Both Isoforms |
| ABCB11 | -9.44 | -1.69 | Both Isoforms | IZUMO1R | -2.72 | -1.16 | Both Isoforms |
| ENO2 | -1.65 | -1.68 | Both Isoforms | ZMYND15 | -2.11 | -1.16 | Both Isoforms |
| CERS1 | -1.41 | -1.68 | Both Isoforms | LTA4H | -1.86 | -1.16 | Both Isoforms |
| AIM1L | -1.76 | -1.68 | Both Isoforms | HHAT | -1.02 | -1.16 | Both Isoforms |
| CD200 | -4.86 | -1.68 | Both Isoforms | INSM2 | -2.47 | -1.16 | Both Isoforms |
| UBE2L6 | -1.29 | -1.68 | Both Isoforms | N4BP2L1 | -2.19 | -1.15 | Both Isoforms |
| CCDC80 | -3.79 | -1.67 | Both Isoforms | NSG1 | -2.75 | -1.15 | Both Isoforms |
| ARL15 | -0.80 | -1.66 | Both Isoforms | HNF4G | -2.28 | -1.15 | Both Isoforms |
| FRMD3 | -1.29 | -1.66 | Both Isoforms | RASSF10 | -2.47 | -1.15 | Both Isoforms |
| GPR27 | -0.74 | -1.66 | Both Isoforms | NEURL3 | -2.68 | -1.15 | Both Isoforms |
| SCARA3 | -0.93 | -1.65 | Both Isoforms | PPM1N | -1.96 | -1.15 | Both Isoforms |
| TMEM173 | -0.66 | -1.65 | Both Isoforms | RARRES2 | -2.42 | -1.15 | Both Isoforms |
| SQRDL | -0.65 | -1.64 | Both Isoforms | FZD3 | -1.15 | -1.15 | Both Isoforms |
| CSTA | -1.08 | -1.64 | Both Isoforms | HLX | -2.08 | -1.15 | Both Isoforms |
| NINL | -2.33 | -1.63 | Both Isoforms | CX3CL1 | -3.34 | -1.15 | Both Isoforms |
| FRMPD1 | -0.92 | -1.62 | Both Isoforms | SEMA6C | -2.20 | -1.14 | Both Isoforms |
| HOXC4 | -1.24 | -1.61 | Both Isoforms | PLSCR4 | -2.10 | -1.14 | Both Isoforms |
| UNC5B | -0.76 | -1.59 | Both Isoforms | IYD | -1.36 | -1.14 | Both Isoforms |
| RGPD5 | -1.58 | -1.59 | Both Isoforms | IL17RD | -2.55 | -1.14 | Both Isoforms |

| | | | | | | | |
|---|---|---|---|---|---|---|---|
| PDZD4 | -0.76 | -1.59 | Both Isoforms | NOL4 | -2.11 | -1.14 | Both Isoforms |
| MAGED4 | -1.66 | -1.58 | Both Isoforms | GPD1L | -0.64 | -1.14 | Both Isoforms |
| SH3PXD2A | -0.92 | -1.58 | Both Isoforms | GDF9 | -1.30 | -1.14 | Both Isoforms |
| SIDT1 | -1.61 | -1.56 | Both Isoforms | ASAH2 | -1.19 | -1.14 | Both Isoforms |
| NAALAD2 | -1.44 | -1.55 | Both Isoforms | LRRC4C | -2.53 | -1.14 | Both Isoforms |
| PKNOX2 | -1.29 | -1.54 | Both Isoforms | LPCAT4 | -2.42 | -1.13 | Both Isoforms |
| PARK2 | -1.77 | -1.54 | Both Isoforms | TMEM26 | -0.82 | -1.13 | Both Isoforms |
| CSF1 | -0.61 | -1.54 | Both Isoforms | PCGF5 | -1.77 | -1.13 | Both Isoforms |
| TEX19 | -0.96 | -1.53 | Both Isoforms | ITGB6 | -0.85 | -1.13 | Both Isoforms |
| AUTS2 | -1.09 | -1.53 | Both Isoforms | ITM2A | -1.91 | -1.13 | Both Isoforms |
| SOX8 | -1.37 | -1.52 | Both Isoforms | RTN4RL1 | -1.41 | -1.13 | Both Isoforms |
| PHGDH | -0.86 | -1.52 | Both Isoforms | FGGY | -1.45 | -1.13 | Both Isoforms |
| PCK2 | -0.69 | -1.52 | Both Isoforms | CDX2 | -1.03 | -1.13 | Both Isoforms |
| STON1 | -0.94 | -1.51 | Both Isoforms | NET1 | -0.79 | -1.12 | Both Isoforms |
| ACKR3 | -2.80 | -1.51 | Both Isoforms | PBX1 | -1.54 | -1.12 | Both Isoforms |
| PDX1 | -0.83 | -1.50 | Both Isoforms | STBD1 | -0.99 | -1.12 | Both Isoforms |
| ZNF84 | -1.79 | -1.50 | Both Isoforms | IL6ST | -1.17 | -1.12 | Both Isoforms |
| REG4 | -2.27 | -1.49 | Both Isoforms | ELFN1 | -1.40 | -1.12 | Both Isoforms |
| TSPAN5 | -1.42 | -1.49 | Both Isoforms | SHISA6 | -0.75 | -1.12 | Both Isoforms |
| SVIP | -0.90 | -1.49 | Both Isoforms | PPM1L | -1.06 | -1.12 | Both Isoforms |
| FGF13 | -1.62 | -1.48 | Both Isoforms | CCDC169 | -1.82 | -1.12 | Both Isoforms |
| DIO1 | -1.86 | -1.48 | Both Isoforms | CLEC2D | -1.19 | -1.11 | Both Isoforms |
| ZMAT1 | -1.38 | -1.48 | Both Isoforms | KIF5A | -2.92 | -1.11 | Both Isoforms |
| FRMD6 | -0.73 | -1.47 | Both Isoforms | MYZAP | -2.67 | -1.11 | Both Isoforms |
| OSR2 | -1.92 | -1.46 | Both Isoforms | IGSF3 | -0.65 | -1.11 | Both Isoforms |

| | | | | | | | |
|---|---|---|---|---|---|---|---|
| FAM171A2 | -1.04 | -1.46 | Both Isoforms | CCNG2 | -1.47 | -1.11 | Both Isoforms |
| ST3GAL1 | -0.77 | -1.46 | Both Isoforms | FBLN2 | -1.88 | -1.11 | Both Isoforms |
| KAZN | -1.12 | -1.45 | Both Isoforms | ENOX1 | -1.31 | -1.11 | Both Isoforms |
| RNF165 | -1.49 | -1.44 | Both Isoforms | PPM1J | -0.80 | -1.11 | Both Isoforms |
| PTPN13 | -0.66 | -1.44 | Both Isoforms | PPP1R3C | -0.68 | -1.11 | Both Isoforms |
| KLHL1 | -0.69 | -1.44 | Both Isoforms | HSD17B6 | -1.95 | -1.11 | Both Isoforms |
| BAHCC1 | -1.00 | -1.43 | Both Isoforms | ADAMTS3 | -1.30 | -1.11 | Both Isoforms |
| PKIB | -2.81 | -1.43 | Both Isoforms | EFNB2 | -0.82 | -1.11 | Both Isoforms |
| CPT1C | -0.87 | -1.42 | Both Isoforms | RASGEF1B | -1.21 | -1.11 | Both Isoforms |
| EMCN | -2.61 | -1.42 | Both Isoforms | SLC17A4 | -2.52 | -1.11 | Both Isoforms |
| NRTN | -0.75 | -1.42 | Both Isoforms | ROR2 | -2.51 | -1.11 | Both Isoforms |
| NANOS1 | -1.38 | -1.42 | Both Isoforms | C1QTNF6 | -1.21 | -1.10 | Both Isoforms |
| NLRC5 | -1.96 | -1.41 | Both Isoforms | HOXD8 | -0.98 | -1.10 | Both Isoforms |
| FAM20C | -1.80 | -1.41 | Both Isoforms | ENPP5 | -2.13 | -1.10 | Both Isoforms |
| PRKAR1B | -1.44 | -1.41 | Both Isoforms | NOX1 | -2.15 | -1.10 | Both Isoforms |
| ACTRT3 | -2.63 | -1.41 | Both Isoforms | NRXN1 | -3.47 | -1.10 | Both Isoforms |
| INHBB | -1.66 | -1.41 | Both Isoforms | HOXB6 | -1.23 | -1.10 | Both Isoforms |
| SOWAHB | -1.20 | -1.41 | Both Isoforms | MTMR11 | -0.79 | -1.10 | Both Isoforms |
| COLEC12 | -2.61 | -1.40 | Both Isoforms | PPP2R2B | -2.72 | -1.10 | Both Isoforms |
| UBA7 | -0.60 | -1.39 | Both Isoforms | ZNF334 | -1.90 | -1.09 | Both Isoforms |
| SOBP | -1.55 | -1.39 | Both Isoforms | ARHGAP9 | -2.20 | -1.09 | Both Isoforms |
| KSR2 | -1.41 | -1.39 | Both Isoforms | EPHB6 | -0.79 | -1.09 | Both Isoforms |
| IKZF3 | -1.08 | -1.38 | Both Isoforms | GPR37 | -0.85 | -1.09 | Both Isoforms |
| ATP2C2 | -0.64 | -1.37 | Both Isoforms | VIL1 | -2.52 | -1.09 | Both Isoforms |
| EML6 | -2.60 | -1.37 | Both Isoforms | AKR7A3 | -1.55 | -1.09 | Both Isoforms |

| | | | | | | | |
|---|---|---|---|---|---|---|---|
| HAP1 | -1.72 | -1.37 | Both Isoforms | PADI2 | -1.73 | -1.09 | Both Isoforms |
| MT1G | -1.20 | -1.36 | Both Isoforms | HNMT | -1.25 | -1.09 | Both Isoforms |
| SRCIN1 | -3.02 | -1.36 | Both Isoforms | FHOD3 | -1.02 | -1.09 | Both Isoforms |
| CAPN5 | -0.88 | -1.36 | Both Isoforms | ZNF491 | -0.79 | -1.09 | Both Isoforms |
| SVIL | -1.47 | -1.36 | Both Isoforms | KCNIP2 | -2.73 | -1.09 | Both Isoforms |
| C1QTNF3 | -1.27 | -1.35 | Both Isoforms | ZNF699 | -0.81 | -1.08 | Both Isoforms |
| AIF1L | -0.85 | -1.35 | Both Isoforms | ZNF385C | -2.78 | -1.08 | Both Isoforms |
| TMEM51 | -1.00 | -1.34 | Both Isoforms | SELENBP1 | -1.89 | -1.08 | Both Isoforms |
| RAI14 | -1.28 | -1.34 | Both Isoforms | AKAP2 | -2.30 | -1.08 | Both Isoforms |
| PSPH | -0.62 | -1.32 | Both Isoforms | GFI1 | -1.75 | -1.08 | Both Isoforms |
| TMEM181 | -0.59 | -1.32 | Both Isoforms | HTRA1 | -2.74 | -1.08 | Both Isoforms |
| KNDC1 | -0.66 | -1.32 | Both Isoforms | COL17A1 | -1.59 | -1.08 | Both Isoforms |
| PARD3B | -0.92 | -1.31 | Both Isoforms | FOXF2 | -1.77 | -1.08 | Both Isoforms |
| CNTN2 | -3.06 | -1.31 | Both Isoforms | FRMD6 | -1.36 | -1.07 | Both Isoforms |
| PACRG | -1.46 | -1.31 | Both Isoforms | NID1 | -1.56 | -1.07 | Both Isoforms |
| OAS3 | -0.77 | -1.31 | Both Isoforms | SCIN | -0.82 | -1.07 | Both Isoforms |
| SH3KBP1 | -0.80 | -1.30 | Both Isoforms | PTH1R | -5.08 | -1.07 | Both Isoforms |
| SLC16A7 | -1.01 | -1.30 | Both Isoforms | KIRREL | -3.78 | -1.07 | Both Isoforms |
| SLCO5A1 | -1.13 | -1.30 | Both Isoforms | TNFRSF11A | -0.80 | -1.07 | Both Isoforms |
| ASNS | -0.73 | -1.29 | Both Isoforms | AIM1L | -1.62 | -1.07 | Both Isoforms |
| LHFPL2 | -0.74 | -1.28 | Both Isoforms | ROBO3 | -0.68 | -1.07 | Both Isoforms |
| ARNT2 | -0.74 | -1.27 | Both Isoforms | NEU4 | -3.34 | -1.07 | Both Isoforms |
| ALOXE3 | -0.75 | -1.27 | Both Isoforms | SEMA6A | -0.99 | -1.06 | Both Isoforms |
| SAMD11 | -1.05 | -1.27 | Both Isoforms | NEFL | -2.00 | -1.06 | Both Isoforms |
| SLC30A10 | -2.40 | -1.27 | Both Isoforms | GCOM1 | -4.05 | -1.06 | Both Isoforms |

| | | | | | | | |
|---|---|---|---|---|---|---|---|
| TUBA4A | -0.89 | -1.27 | Both Isoforms | ADGRL2 | -1.07 | -1.06 | Both Isoforms |
| PSAT1 | -1.04 | -1.26 | Both Isoforms | MXI1 | -3.07 | -1.06 | Both Isoforms |
| PLGRKT | -0.64 | -1.25 | Both Isoforms | ITGA2 | -1.33 | -1.06 | Both Isoforms |
| TINAGL1 | -1.90 | -1.25 | Both Isoforms | TMEM158 | -0.95 | -1.06 | Both Isoforms |
| MARK1 | -1.48 | -1.24 | Both Isoforms | 2-Mar | -0.94 | -1.06 | Both Isoforms |
| SRGAP3 | -1.25 | -1.24 | Both Isoforms | LASP1 | -1.72 | -1.06 | Both Isoforms |
| ZYG11A | -1.63 | -1.24 | Both Isoforms | ERBB3 | -0.59 | -1.06 | Both Isoforms |
| STK3 | -0.83 | -1.24 | Both Isoforms | SSPN | -2.05 | -1.06 | Both Isoforms |
| MAPKAPK3 | -1.11 | -1.22 | Both Isoforms | NLGN1 | -2.70 | -1.05 | Both Isoforms |
| LRRC34 | -2.37 | -1.22 | Both Isoforms | OXTR | -2.02 | -1.05 | Both Isoforms |
| ANO9 | -0.87 | -1.22 | Both Isoforms | CEP68 | -2.85 | -1.05 | Both Isoforms |
| PTRF | -1.09 | -1.22 | Both Isoforms | RAP2B | -2.58 | -1.05 | Both Isoforms |
| CWH43 | -0.76 | -1.21 | Both Isoforms | XYLT1 | -0.76 | -1.05 | Both Isoforms |
| CLDND2 | -1.07 | -1.21 | Both Isoforms | DSC3 | -0.64 | -1.05 | Both Isoforms |
| APBA1 | -0.70 | -1.21 | Both Isoforms | CABYR | -1.54 | -1.05 | Both Isoforms |
| SLC29A1 | -1.22 | -1.21 | Both Isoforms | HOXC13 | -0.86 | -1.05 | Both Isoforms |
| FRMPD3 | -0.94 | -1.20 | Both Isoforms | ADM | -1.44 | -1.05 | Both Isoforms |
| SLC9A3 | -2.15 | -1.20 | Both Isoforms | BCO1 | -1.43 | -1.05 | Both Isoforms |
| POU5F1B | -0.76 | -1.20 | Both Isoforms | SERPINB9 | -1.97 | -1.05 | Both Isoforms |
| DMGDH | -0.97 | -1.19 | Both Isoforms | PARP8 | -2.29 | -1.05 | Both Isoforms |
| SLC16A14 | -1.25 | -1.19 | Both Isoforms | PCMTD1 | -0.67 | -1.04 | Both Isoforms |
| NCOA7 | -1.32 | -1.18 | Both Isoforms | SGTB | -1.38 | -1.04 | Both Isoforms |
| MYT1 | -1.29 | -1.18 | Both Isoforms | MCTP1 | -2.82 | -1.04 | Both Isoforms |
| ADAM11 | -0.60 | -1.18 | Both Isoforms | FAM131B | -0.94 | -1.04 | Both Isoforms |
| EFNB3 | -0.99 | -1.16 | Both Isoforms | KLHDC1 | -1.22 | -1.04 | Both Isoforms |

| | | | | |
|---|---|---|---|---|
| ZDHHC14 | | -0.68 | -1.16 | Both Isoforms |
| ARHGAP27 | | -1.48 | -1.16 | Both Isoforms |
| NEDD9 | | -1.02 | -1.16 | Both Isoforms |
| PKP4 | | -1.24 | -1.15 | Both Isoforms |
| AGBL3 | | -0.96 | -1.15 | Both Isoforms |
| MYNN | | -1.58 | -1.15 | Both Isoforms |
| EXPH5 | | -0.86 | -1.14 | Both Isoforms |
| SLC16A10 | | -0.78 | -1.14 | Both Isoforms |
| CPNE3 | | -0.81 | -1.14 | Both Isoforms |
| ASIC1 | | -1.09 | -1.14 | Both Isoforms |
| | 3-Mar | -0.75 | -1.14 | Both Isoforms |
| C16orf45 | | -2.28 | -1.14 | Both Isoforms |
| MAPT | | -1.34 | -1.12 | Both Isoforms |
| GPR160 | | -0.90 | -1.12 | Both Isoforms |
| GSAP | | -0.99 | -1.12 | Both Isoforms |
| CABLES1 | | -0.81 | -1.11 | Both Isoforms |
| PCDHGA12 | | -0.67 | -1.11 | Both Isoforms |
| TBL1Y | | -0.93 | -1.11 | Both Isoforms |
| SLC44A1 | | -1.26 | -1.11 | Both Isoforms |
| C5orf30 | | -1.45 | -1.11 | Both Isoforms |
| LRRC16B | | -0.98 | -1.11 | Both Isoforms |
| SRMS | | -2.05 | -1.11 | Both Isoforms |
| C3orf70 | | -1.13 | -1.10 | Both Isoforms |
| NWD1 | | -1.74 | -1.10 | Both Isoforms |
| | 3-Sep | -0.85 | -1.10 | Both Isoforms |

| | | | |
|---|---|---|---|
| GJA3 | -1.83 | -1.04 | Both Isoforms |
| TNFSF4 | -0.94 | -1.04 | Both Isoforms |
| SLC12A6 | -0.67 | -1.04 | Both Isoforms |
| TSHZ3 | -1.99 | -1.04 | Both Isoforms |
| PAPSS1 | -0.81 | -1.04 | Both Isoforms |
| ATP8A1 | -1.46 | -1.04 | Both Isoforms |
| DLX3 | -1.01 | -1.04 | Both Isoforms |
| SI | -4.20 | -1.03 | Both Isoforms |
| MTURN | -2.33 | -1.03 | Both Isoforms |
| CCDC122 | -1.38 | -1.03 | Both Isoforms |
| ACOXL | -1.33 | -1.03 | Both Isoforms |
| HS3ST3A1 | -3.07 | -1.03 | Both Isoforms |
| TMEM27 | -1.16 | -1.03 | Both Isoforms |
| HSPA12B | -1.26 | -1.03 | Both Isoforms |
| SGF29 | -1.37 | -1.03 | Both Isoforms |
| SPATA17 | -1.39 | -1.03 | Both Isoforms |
| PLEKHA2 | -1.19 | -1.02 | Both Isoforms |
| BACH2 | -2.17 | -1.02 | Both Isoforms |
| HOXC12 | -1.09 | -1.02 | Both Isoforms |
| RORB | -2.08 | -1.02 | Both Isoforms |
| C11orf95 | -1.36 | -1.02 | Both Isoforms |
| CALHM2 | -1.74 | -1.02 | Both Isoforms |
| COL28A1 | -1.09 | -1.02 | Both Isoforms |
| LIAS | -1.54 | -1.02 | Both Isoforms |
| ESYT3 | -1.40 | -1.02 | Both Isoforms |

| | | | | | | | |
|---|---|---|---|---|---|---|---|
| SYNGAP1 | -0.66 | -1.09 | Both Isoforms | MIA3 | -1.12 | -1.01 | Both Isoforms |
| DOCK3 | -0.82 | -1.09 | Both Isoforms | LGR4 | -1.06 | -1.01 | Both Isoforms |
| ZNF629 | -0.97 | -1.09 | Both Isoforms | DDX25 | -1.76 | -1.01 | Both Isoforms |
| PABPC5 | -1.08 | -1.09 | Both Isoforms | C7orf31 | -1.26 | -1.01 | Both Isoforms |
| SERPINI1 | -2.09 | -1.09 | Both Isoforms | DBP | -2.17 | -1.01 | Both Isoforms |
| KCNAB2 | -1.13 | -1.09 | Both Isoforms | FRY | -1.23 | -1.01 | Both Isoforms |
| B4GALNT1 | -1.05 | -1.08 | Both Isoforms | YPEL2 | -0.70 | -1.01 | Both Isoforms |
| NDUFAF4 | -0.82 | -1.07 | Both Isoforms | C15orf48 | -2.40 | -1.01 | Both Isoforms |
| ZNF367 | -0.95 | -1.07 | Both Isoforms | NKX2-1 | -0.83 | -1.01 | Both Isoforms |
| SLC2A9 | -1.54 | -1.06 | Both Isoforms | CDH4 | -1.80 | -1.01 | Both Isoforms |
| STMN3 | -0.62 | -1.06 | Both Isoforms | AGO4 | -1.11 | -1.01 | Both Isoforms |
| IL17D | -0.89 | -1.06 | Both Isoforms | RINL | -1.37 | -1.01 | Both Isoforms |
| SELENBP1 | -1.74 | -1.05 | Both Isoforms | HFM1 | -2.65 | -1.01 | Both Isoforms |
| TNNT2 | -0.97 | -1.05 | Both Isoforms | DNAH5 | -1.64 | -1.01 | Both Isoforms |
| ASAP2 | -1.43 | -1.05 | Both Isoforms | FOXD3 | -0.96 | -1.00 | Both Isoforms |
| MX1 | -0.60 | -1.04 | Both Isoforms | TBX19 | -0.58 | -1.00 | Both Isoforms |
| PLCD3 | -0.84 | -1.04 | Both Isoforms | CA13 | -0.95 | -1.00 | Both Isoforms |
| GPR143 | -0.98 | -1.03 | Both Isoforms | BTN3A2 | -2.19 | -1.00 | Both Isoforms |
| GABRA3 | -0.88 | -1.03 | Both Isoforms | GSTO2 | -2.46 | -1.00 | Both Isoforms |
| RALB | -0.83 | -1.03 | Both Isoforms | PAK1 | -1.32 | -1.00 | Both Isoforms |
| CRYL1 | -0.61 | -1.03 | Both Isoforms | SYT15 | -1.50 | -1.00 | Both Isoforms |
| GLI3 | -1.50 | -1.03 | Both Isoforms | SCN2A | -1.63 | -1.00 | Both Isoforms |
| EBF3 | -1.07 | -1.03 | Both Isoforms | ELFN2 | -1.32 | -1.00 | Both Isoforms |
| KIAA1211 | -1.12 | -1.02 | Both Isoforms | SLC8A2 | -1.61 | -1.00 | Both Isoforms |
| SYTL2 | -2.27 | -1.02 | Both Isoforms | MX1 | -2.18 | -1.00 | Both Isoforms |

| | | | | | | | | |
|---|---|---|---|---|---|---|---|---|
| FBLN7 | | -1.46 | -1.02 | Both Isoforms | EIF4EBP3 | -1.88 | -1.00 | Both Isoforms |
| WNT2B | | -1.27 | -1.02 | Both Isoforms | LRIG3 | -1.47 | -1.00 | Both Isoforms |
| VANGL2 | | -0.74 | -1.02 | Both Isoforms | GARNL3 | -1.18 | -1.00 | Both Isoforms |
| TRERF1 | | -0.90 | -1.02 | Both Isoforms | MLLT3 | -1.17 | -1.00 | Both Isoforms |
| EFNA3 | | -0.68 | -1.01 | Both Isoforms | STOM | -1.67 | -1.00 | Both Isoforms |
| FRMD5 | | -1.83 | -1.01 | Both Isoforms | GLCCI1 | -0.65 | -1.00 | Both Isoforms |
| GLCCI1 | | -1.03 | -1.01 | Both Isoforms | PYGL | -2.35 | -0.99 | Both Isoforms |
| CCDC14 | | -0.95 | -1.01 | Both Isoforms | ARHGAP22 | -2.77 | -0.99 | Both Isoforms |
| RNF43 | | -1.41 | -1.01 | Both Isoforms | PARK2 | -2.55 | -0.99 | Both Isoforms |
| NR1D1 | | -1.15 | -1.01 | Both Isoforms | LCA5 | -1.76 | -0.99 | Both Isoforms |
| ALX4 | | -1.68 | -1.01 | Both Isoforms | TMPRSS13 | -1.47 | -0.99 | Both Isoforms |
| KIF7 | | -0.95 | -1.01 | Both Isoforms | BEX2 | -1.55 | -0.99 | Both Isoforms |
| LRRN2 | | -1.84 | -1.01 | Both Isoforms | ZCCHC7 | -0.87 | -0.99 | Both Isoforms |
| ZNF365 | | -1.17 | -1.01 | Both Isoforms | EXPH5 | -1.57 | -0.99 | Both Isoforms |
| SKA3 | | -0.73 | -1.01 | Both Isoforms | ICAM5 | -2.46 | -0.99 | Both Isoforms |
| RGS10 | | -0.84 | -1.00 | Both Isoforms | C1QTNF3 | -1.40 | -0.99 | Both Isoforms |
| FAM81A | | -0.81 | -1.00 | Both Isoforms | REXO2 | -1.33 | -0.99 | Both Isoforms |
| CCNJL | | -1.61 | -1.00 | Both Isoforms | SMAD3 | -2.40 | -0.99 | Both Isoforms |
| ADAP1 | | -1.83 | -1.00 | Both Isoforms | PRR26 | -1.97 | -0.99 | Both Isoforms |
| | 11-Sep | -0.74 | -1.00 | Both Isoforms | DUSP16 | -0.68 | -0.99 | Both Isoforms |
| ABCG2 | | -2.65 | -1.00 | Both Isoforms | PLK2 | -2.80 | -0.98 | Both Isoforms |
| PCDHGB4 | | -0.79 | -1.00 | Both Isoforms | SERPINA10 | -3.36 | -0.98 | Both Isoforms |
| CSMD1 | | -1.38 | -1.00 | Both Isoforms | BDH2 | -1.30 | -0.98 | Both Isoforms |
| PNMA3 | | -0.67 | -1.00 | Both Isoforms | RNF19A | -1.18 | -0.98 | Both Isoforms |
| FZD3 | | -1.76 | -1.00 | Both Isoforms | AKAP6 | -1.31 | -0.98 | Both Isoforms |

| | | | | | | | |
|---|---|---|---|---|---|---|---|
| DTX3 | -1.01 | -0.99 | Both Isoforms | ATXN7L1 | -2.01 | -0.98 | Both Isoforms |
| CA11 | -0.86 | -0.99 | Both Isoforms | LRRC16B | -1.60 | -0.98 | Both Isoforms |
| HOXC13 | -1.37 | -0.99 | Both Isoforms | FRMPD2 | -0.68 | -0.98 | Both Isoforms |
| RPP25 | -1.23 | -0.99 | Both Isoforms | MANBA | -1.45 | -0.98 | Both Isoforms |
| SCRN1 | -0.93 | -0.99 | Both Isoforms | ADGRA2 | -2.83 | -0.98 | Both Isoforms |
| GOLGA6L10 | -2.45 | -0.99 | Both Isoforms | PARD3B | -0.83 | -0.98 | Both Isoforms |
| COL5A1 | -2.01 | -0.99 | Both Isoforms | MRAS | -0.64 | -0.97 | Both Isoforms |
| ETS1 | -0.74 | -0.98 | Both Isoforms | GJC3 | -1.43 | -0.97 | Both Isoforms |
| SBK1 | -0.61 | -0.98 | Both Isoforms | TNIK | -0.70 | -0.97 | Both Isoforms |
| ZNF117 | -1.05 | -0.98 | Both Isoforms | RASAL1 | -1.16 | -0.97 | Both Isoforms |
| SMAD1 | -0.78 | -0.98 | Both Isoforms | CAMK1D | -1.46 | -0.97 | Both Isoforms |
| CNIH3 | -1.06 | -0.97 | Both Isoforms | PCDHB12 | -1.41 | -0.97 | Both Isoforms |
| MCU | -0.81 | -0.97 | Both Isoforms | KIAA1462 | -0.97 | -0.97 | Both Isoforms |
| SH3BGRL2 | -0.88 | -0.97 | Both Isoforms | STAT6 | -1.22 | -0.97 | Both Isoforms |
| SHC4 | -1.13 | -0.97 | Both Isoforms | ZNF608 | -0.60 | -0.97 | Both Isoforms |
| ISM2 | -1.37 | -0.97 | Both Isoforms | PRSS27 | -1.11 | -0.97 | Both Isoforms |
| CNGA1 | -1.53 | -0.97 | Both Isoforms | CABLES1 | -1.41 | -0.97 | Both Isoforms |
| PLEKHA2 | -0.77 | -0.97 | Both Isoforms | RASSF5 | -1.88 | -0.97 | Both Isoforms |
| BMX | -1.52 | -0.97 | Both Isoforms | ALDH1L2 | -2.75 | -0.96 | Both Isoforms |
| CTB-55O6.8 | -1.17 | -0.96 | Both Isoforms | BTC | -1.50 | -0.96 | Both Isoforms |
| CDCA7L | -1.54 | -0.96 | Both Isoforms | DENND2C | -1.83 | -0.96 | Both Isoforms |
| WASF1 | -0.70 | -0.96 | Both Isoforms | FAM227B | -0.97 | -0.96 | Both Isoforms |
| ACRV1 | -0.88 | -0.96 | Both Isoforms | FAM155A | -1.19 | -0.96 | Both Isoforms |
| GRB10 | -1.85 | -0.96 | Both Isoforms | SMTNL2 | -2.20 | -0.96 | Both Isoforms |
| MID1 | -0.88 | -0.95 | Both Isoforms | NEO1 | -0.84 | -0.96 | Both Isoforms |

| | | | | | | | |
|---|---|---|---|---|---|---|---|
| CFAP45 | -1.03 | -0.95 | Both Isoforms | KCTD12 | -2.49 | -0.96 | Both Isoforms |
| AC138969.4 | -0.95 | -0.95 | Both Isoforms | GAL3ST1 | -3.22 | -0.96 | Both Isoforms |
| PLEKHG5 | -0.73 | -0.95 | Both Isoforms | ZNF345 | -1.32 | -0.96 | Both Isoforms |
| EGLN3 | -1.88 | -0.95 | Both Isoforms | CPA6 | -2.11 | -0.96 | Both Isoforms |
| ADRBK2 | -0.59 | -0.95 | Both Isoforms | BCL2L10 | -1.71 | -0.96 | Both Isoforms |
| GSDMD | -1.34 | -0.94 | Both Isoforms | NADK2 | -1.30 | -0.95 | Both Isoforms |
| NUP54 | -0.81 | -0.94 | Both Isoforms | RAI2 | -1.01 | -0.95 | Both Isoforms |
| ADAMTSL4 | -1.06 | -0.94 | Both Isoforms | NR4A2 | -2.91 | -0.95 | Both Isoforms |
| PRKD1 | -1.36 | -0.94 | Both Isoforms | CDKL5 | -1.13 | -0.95 | Both Isoforms |
| CORO1A | -1.57 | -0.94 | Both Isoforms | TMEM173 | -1.60 | -0.95 | Both Isoforms |
| DTX4 | -1.26 | -0.94 | Both Isoforms | RPS18 | -1.10 | -0.95 | Both Isoforms |
| TRPS1 | -1.22 | -0.94 | Both Isoforms | CNKSR3 | -1.95 | -0.95 | Both Isoforms |
| STOX1 | -2.34 | -0.94 | Both Isoforms | C4orf36 | -1.40 | -0.95 | Both Isoforms |
| IFIT5 | -0.79 | -0.94 | Both Isoforms | PRICKLE1 | -1.42 | -0.95 | Both Isoforms |
| E2F2 | -1.30 | -0.93 | Both Isoforms | DAPK2 | -1.71 | -0.95 | Both Isoforms |
| IKZF2 | -0.93 | -0.93 | Both Isoforms | ZNF594 | -1.42 | -0.95 | Both Isoforms |
| FZD7 | -0.78 | -0.92 | Both Isoforms | WNT8B | -2.17 | -0.95 | Both Isoforms |
| IGIP | -1.10 | -0.92 | Both Isoforms | PCDHGB7 | -2.20 | -0.95 | Both Isoforms |
| PCNA | -0.81 | -0.92 | Both Isoforms | SAMD4A | -1.93 | -0.94 | Both Isoforms |
| FOLH1 | -0.79 | -0.92 | Both Isoforms | SYNGAP1 | -1.49 | -0.94 | Both Isoforms |
| MCM5 | -0.97 | -0.92 | Both Isoforms | ZFP92 | -1.73 | -0.94 | Both Isoforms |
| HELLS | -1.32 | -0.92 | Both Isoforms | MKX | -2.51 | -0.94 | Both Isoforms |
| C1QTNF6 | -1.85 | -0.91 | Both Isoforms | SGMS1 | -0.95 | -0.94 | Both Isoforms |
| APBA2 | -0.80 | -0.91 | Both Isoforms | NUDT4 | -1.06 | -0.94 | Both Isoforms |
| CCDC85B | -0.73 | -0.91 | Both Isoforms | ZNF33B | -1.24 | -0.94 | Both Isoforms |

| | | | | | | | |
|---|---|---|---|---|---|---|---|
| HOXA10 | -1.35 | -0.91 | Both Isoforms | NOB1 | -1.27 | -0.94 | Both Isoforms |
| ANKRD46 | -0.62 | -0.91 | Both Isoforms | QKI | -1.70 | -0.94 | Both Isoforms |
| ENKUR | -0.75 | -0.91 | Both Isoforms | PAX2 | -1.59 | -0.94 | Both Isoforms |
| IL36G | -3.21 | -0.91 | Both Isoforms | ABTB2 | -1.14 | -0.94 | Both Isoforms |
| BLNK | -1.07 | -0.91 | Both Isoforms | ZNF660 | -2.88 | -0.94 | Both Isoforms |
| VASH1 | -1.09 | -0.91 | Both Isoforms | MAP3K7CL | -1.14 | -0.94 | Both Isoforms |
| DPYSL5 | -2.22 | -0.91 | Both Isoforms | ASPH | -2.07 | -0.93 | Both Isoforms |
| RHBG | -1.75 | -0.90 | Both Isoforms | DSG2 | -0.71 | -0.93 | Both Isoforms |
| WDR62 | -1.01 | -0.90 | Both Isoforms | LIPM | -1.27 | -0.93 | Both Isoforms |
| LMX1B | -1.08 | -0.90 | Both Isoforms | CCSER1 | -1.28 | -0.93 | Both Isoforms |
| E2F1 | -0.94 | -0.90 | Both Isoforms | TRIM7 | -1.24 | -0.93 | Both Isoforms |
| SAMHD1 | -0.71 | -0.90 | Both Isoforms | SPINK1 | -1.51 | -0.93 | Both Isoforms |
| TEAD4 | -0.71 | -0.90 | Both Isoforms | CLRN3 | -2.13 | -0.93 | Both Isoforms |
| ABHD18 | -1.11 | -0.90 | Both Isoforms | MTSS1 | -1.07 | -0.93 | Both Isoforms |
| ATP11C | -1.33 | -0.89 | Both Isoforms | BASP1 | -1.39 | -0.93 | Both Isoforms |
| FBXL7 | -1.69 | -0.89 | Both Isoforms | CYP27B1 | -1.65 | -0.93 | Both Isoforms |
| BROX | -0.60 | -0.89 | Both Isoforms | CD55 | -1.74 | -0.93 | Both Isoforms |
| KIAA1324 | -2.13 | -0.89 | Both Isoforms | NCAM1 | -1.62 | -0.93 | Both Isoforms |
| PBX1 | -1.65 | -0.89 | Both Isoforms | ZIC5 | -1.87 | -0.93 | Both Isoforms |
| MREG | -1.20 | -0.89 | Both Isoforms | TGM3 | -2.42 | -0.93 | Both Isoforms |
| ANXA9 | -1.02 | -0.89 | Both Isoforms | ULBP1 | -2.23 | -0.93 | Both Isoforms |
| SLC6A11 | -0.73 | -0.88 | Both Isoforms | CACNB4 | -1.75 | -0.93 | Both Isoforms |
| SLFN5 | -1.28 | -0.88 | Both Isoforms | CASP4 | -1.60 | -0.93 | Both Isoforms |
| PIK3IP1 | -1.43 | -0.88 | Both Isoforms | MS4A8 | -0.66 | -0.93 | Both Isoforms |
| CUEDC1 | -0.61 | -0.88 | Both Isoforms | LGALS4 | -0.72 | -0.93 | Both Isoforms |

| | | | | | | | |
|---|---|---|---|---|---|---|---|
| VOPP1 | -0.93 | -0.88 | Both Isoforms | BFSP1 | -1.15 | -0.93 | Both Isoforms |
| RGS17 | -0.88 | -0.88 | Both Isoforms | PRSS3 | -1.32 | -0.93 | Both Isoforms |
| MND1 | -1.09 | -0.88 | Both Isoforms | CELF5 | -1.84 | -0.93 | Both Isoforms |
| BCHE | -2.70 | -0.88 | Both Isoforms | INHBB | -1.20 | -0.92 | Both Isoforms |
| RFX2 | -1.03 | -0.88 | Both Isoforms | ITGA2B | -1.88 | -0.92 | Both Isoforms |
| RASGRP1 | -0.99 | -0.87 | Both Isoforms | ACCS | -1.59 | -0.92 | Both Isoforms |
| IGSF9 | -0.83 | -0.87 | Both Isoforms | SCN8A | -1.75 | -0.92 | Both Isoforms |
| STC2 | -1.13 | -0.87 | Both Isoforms | CPNE4 | -0.70 | -0.92 | Both Isoforms |
| B4GALNT4 | -0.91 | -0.87 | Both Isoforms | CADM1 | -0.62 | -0.92 | Both Isoforms |
| FYN | -2.15 | -0.87 | Both Isoforms | ADD3 | -2.13 | -0.92 | Both Isoforms |
| ARHGAP42 | -0.83 | -0.87 | Both Isoforms | DCLK2 | -1.23 | -0.92 | Both Isoforms |
| TNFRSF25 | -0.92 | -0.86 | Both Isoforms | SNX24 | -0.66 | -0.92 | Both Isoforms |
| GRK5 | -1.19 | -0.86 | Both Isoforms | ISX | -0.81 | -0.91 | Both Isoforms |
| CBR3 | -0.96 | -0.86 | Both Isoforms | PPP1R12B | -1.26 | -0.91 | Both Isoforms |
| COL9A2 | -1.04 | -0.86 | Both Isoforms | TMPRSS9 | -1.96 | -0.91 | Both Isoforms |
| AVPI1 | -0.78 | -0.86 | Both Isoforms | TAF4B | -1.62 | -0.91 | Both Isoforms |
| SUSD6 | -1.23 | -0.85 | Both Isoforms | MB21D2 | -3.08 | -0.91 | Both Isoforms |
| HIBADH | -0.97 | -0.85 | Both Isoforms | PRX | -1.49 | -0.91 | Both Isoforms |
| TMEM133 | -0.83 | -0.85 | Both Isoforms | FAM234B | -0.92 | -0.91 | Both Isoforms |
| KHDRBS3 | -0.95 | -0.85 | Both Isoforms | KHDRBS3 | -1.85 | -0.91 | Both Isoforms |
| JDP2 | -1.29 | -0.85 | Both Isoforms | BBC3 | -2.38 | -0.91 | Both Isoforms |
| SYTL3 | -0.65 | -0.85 | Both Isoforms | KLC3 | -1.38 | -0.91 | Both Isoforms |
| GXYLT2 | -1.01 | -0.85 | Both Isoforms | CREB5 | -1.94 | -0.91 | Both Isoforms |
| TIMP2 | -1.12 | -0.85 | Both Isoforms | NABP1 | -1.50 | -0.91 | Both Isoforms |
| NKD1 | -0.91 | -0.85 | Both Isoforms | IMPA1 | -0.84 | -0.91 | Both Isoforms |

| | | | | | | | |
|---|---|---|---|---|---|---|---|
| RAD54L | -1.15 | -0.84 | Both Isoforms | LRRIQ3 | -1.36 | -0.91 | Both Isoforms |
| ZBTB43 | -1.25 | -0.84 | Both Isoforms | HOXD13 | -1.51 | -0.91 | Both Isoforms |
| TMEM121 | -0.93 | -0.84 | Both Isoforms | SLC18B1 | -2.67 | -0.91 | Both Isoforms |
| CDCA7 | -0.97 | -0.84 | Both Isoforms | RAVER2 | -1.52 | -0.91 | Both Isoforms |
| FA2H | -0.99 | -0.84 | Both Isoforms | DEPTOR | -1.68 | -0.90 | Both Isoforms |
| CECR1 | -1.24 | -0.84 | Both Isoforms | ODF3B | -0.97 | -0.90 | Both Isoforms |
| CDON | -1.03 | -0.84 | Both Isoforms | CHMP4C | -1.16 | -0.90 | Both Isoforms |
| CDC45 | -1.11 | -0.83 | Both Isoforms | ZRANB3 | -1.77 | -0.90 | Both Isoforms |
| GINS2 | -0.71 | -0.83 | Both Isoforms | TTC21B | -0.80 | -0.90 | Both Isoforms |
| SLC27A2 | -1.10 | -0.83 | Both Isoforms | GAD1 | -1.86 | -0.90 | Both Isoforms |
| XXYLT1 | -0.58 | -0.83 | Both Isoforms | NMNAT3 | -1.97 | -0.90 | Both Isoforms |
| EFR3B | -1.40 | -0.83 | Both Isoforms | ANKRD36 | -1.78 | -0.90 | Both Isoforms |
| NEIL3 | -0.59 | -0.83 | Both Isoforms | TSPAN5 | -1.27 | -0.90 | Both Isoforms |
| CENPU | -0.88 | -0.83 | Both Isoforms | HS2ST1 | -0.71 | -0.90 | Both Isoforms |
| C4orf32 | -1.21 | -0.83 | Both Isoforms | JMY | -1.02 | -0.90 | Both Isoforms |
| KCNQ1 | -1.05 | -0.83 | Both Isoforms | MCAM | -3.89 | -0.90 | Both Isoforms |
| ZNF624 | -0.76 | -0.82 | Both Isoforms | RARG | -2.10 | -0.90 | Both Isoforms |
| ABCA1 | -4.16 | -0.82 | Both Isoforms | SP100 | -1.40 | -0.90 | Both Isoforms |
| FMNL3 | -1.26 | -0.82 | Both Isoforms | CCDC110 | -1.06 | -0.90 | Both Isoforms |
| PRELID3A | -0.76 | -0.82 | Both Isoforms | CCDC28A | -1.37 | -0.90 | Both Isoforms |
| RABL3 | -0.61 | -0.81 | Both Isoforms | ARL4C | -0.67 | -0.90 | Both Isoforms |
| GNAZ | -1.19 | -0.81 | Both Isoforms | RYR2 | -1.74 | -0.90 | Both Isoforms |
| SLC22A11 | -0.71 | -0.81 | Both Isoforms | LAMB1 | -1.35 | -0.90 | Both Isoforms |
| MCM10 | -1.30 | -0.81 | Both Isoforms | MBTD1 | -1.35 | -0.89 | Both Isoforms |
| CCNE2 | -1.29 | -0.81 | Both Isoforms | KRCC1 | -1.19 | -0.89 | Both Isoforms |

| | | | | | | | |
|---|---|---|---|---|---|---|---|
| IGFBP2 | -0.82 | -0.80 | Both Isoforms | VPS37D | -0.82 | -0.89 | Both Isoforms |
| CHEK2 | -0.71 | -0.80 | Both Isoforms | SYNPO2 | -3.48 | -0.89 | Both Isoforms |
| TGIF2 | -0.62 | -0.80 | Both Isoforms | GCNT1 | -1.00 | -0.89 | Both Isoforms |
| RAVER2 | -0.87 | -0.80 | Both Isoforms | DTX4 | -2.32 | -0.89 | Both Isoforms |
| TYMS | -0.76 | -0.80 | Both Isoforms | BAG2 | -1.09 | -0.89 | Both Isoforms |
| HOXC12 | -0.90 | -0.80 | Both Isoforms | ANKRD30B | -2.29 | -0.89 | Both Isoforms |
| KHK | -1.20 | -0.80 | Both Isoforms | FNDC4 | -1.42 | -0.89 | Both Isoforms |
| MCM3 | -0.94 | -0.80 | Both Isoforms | POU2F2 | -0.83 | -0.89 | Both Isoforms |
| GRAMD1C | -1.06 | -0.79 | Both Isoforms | MATN2 | -1.01 | -0.89 | Both Isoforms |
| TONSL | -0.95 | -0.79 | Both Isoforms | PIGR | -2.44 | -0.89 | Both Isoforms |
| PIPOX | -0.70 | -0.78 | Both Isoforms | RASSF2 | -0.59 | -0.89 | Both Isoforms |
| C4orf46 | -1.17 | -0.78 | Both Isoforms | JAK3 | -1.80 | -0.89 | Both Isoforms |
| DNAH2 | -0.88 | -0.78 | Both Isoforms | HHIPL1 | -1.16 | -0.88 | Both Isoforms |
| EEF1A2 | -0.72 | -0.78 | Both Isoforms | RLTPR | -1.66 | -0.88 | Both Isoforms |
| LPCAT4 | -1.58 | -0.78 | Both Isoforms | C20orf194 | -1.54 | -0.88 | Both Isoforms |
| WNT10A | -0.67 | -0.78 | Both Isoforms | SLIT1 | -1.85 | -0.88 | Both Isoforms |
| SLC48A1 | -0.97 | -0.78 | Both Isoforms | TMEM38B | -0.96 | -0.88 | Both Isoforms |
| CTSH | -1.03 | -0.77 | Both Isoforms | MAML2 | -1.08 | -0.88 | Both Isoforms |
| ADGRV1 | -1.23 | -0.77 | Both Isoforms | ZNF425 | -1.04 | -0.88 | Both Isoforms |
| RCC1 | -0.76 | -0.77 | Both Isoforms | ABHD14B | -1.58 | -0.88 | Both Isoforms |
| POLA2 | -0.88 | -0.77 | Both Isoforms | HECTD2 | -0.64 | -0.88 | Both Isoforms |
| C20orf27 | -0.74 | -0.77 | Both Isoforms | ARL10 | -2.31 | -0.88 | Both Isoforms |
| TMEM117 | -1.08 | -0.77 | Both Isoforms | SH3BGR | -1.21 | -0.88 | Both Isoforms |
| TERT | -1.43 | -0.77 | Both Isoforms | OTUD1 | -1.90 | -0.88 | Both Isoforms |
| TRIM16L | -0.59 | -0.77 | Both Isoforms | GPRASP2 | -1.09 | -0.88 | Both Isoforms |

| | | | | | | | |
|---|---|---|---|---|---|---|---|
| FAM134B | -1.78 | -0.77 | Both Isoforms | POTEI | -0.71 | -0.87 | Both Isoforms |
| KLHL23 | -0.60 | -0.76 | Both Isoforms | FOXD2 | -1.70 | -0.87 | Both Isoforms |
| XRCC2 | -1.20 | -0.76 | Both Isoforms | DYRK3 | -1.99 | -0.87 | Both Isoforms |
| CPAMD8 | -2.37 | -0.76 | Both Isoforms | NCOA7 | -2.32 | -0.87 | Both Isoforms |
| MAP3K8 | -1.44 | -0.75 | Both Isoforms | FERMT1 | -1.34 | -0.87 | Both Isoforms |
| DPYSL2 | -0.98 | -0.75 | Both Isoforms | ZSCAN12 | -0.59 | -0.87 | Both Isoforms |
| GATA2 | -0.68 | -0.75 | Both Isoforms | ITGB2 | -1.64 | -0.87 | Both Isoforms |
| PRAC1 | -1.08 | -0.75 | Both Isoforms | RELB | -1.54 | -0.87 | Both Isoforms |
| C11orf95 | -1.10 | -0.74 | Both Isoforms | CBR4 | -1.50 | -0.87 | Both Isoforms |
| MCM6 | -0.76 | -0.74 | Both Isoforms | R3HDM2 | -0.87 | -0.87 | Both Isoforms |
| MAGI1 | -0.63 | -0.74 | Both Isoforms | DNAJC18 | -1.89 | -0.87 | Both Isoforms |
| SLC10A5 | -0.69 | -0.74 | Both Isoforms | C8orf58 | -0.97 | -0.87 | Both Isoforms |
| PURA | -0.85 | -0.74 | Both Isoforms | ANKRD26 | -1.49 | -0.87 | Both Isoforms |
| MYBL2 | -0.92 | -0.74 | Both Isoforms | WDR88 | -1.62 | -0.87 | Both Isoforms |
| HACL1 | -0.88 | -0.74 | Both Isoforms | LRRC37B | -1.08 | -0.87 | Both Isoforms |
| WDR91 | -0.92 | -0.73 | Both Isoforms | DSE | -0.70 | -0.87 | Both Isoforms |
| GVQW2 | -1.32 | -0.73 | Both Isoforms | MPP1 | -1.52 | -0.87 | Both Isoforms |
| EXO1 | -1.38 | -0.73 | Both Isoforms | ADAM22 | -1.61 | -0.87 | Both Isoforms |
| TBC1D31 | -0.74 | -0.73 | Both Isoforms | NR2F1 | -3.05 | -0.87 | Both Isoforms |
| SLC1A3 | -1.10 | -0.73 | Both Isoforms | BTN3A3 | -1.19 | -0.87 | Both Isoforms |
| SMKR1 | -0.93 | -0.73 | Both Isoforms | TGFBR3L | -1.49 | -0.87 | Both Isoforms |
| HOXC9 | -1.89 | -0.73 | Both Isoforms | SH3BGRL2 | -1.12 | -0.87 | Both Isoforms |
| IL4R | -0.60 | -0.73 | Both Isoforms | PROCR | -1.29 | -0.86 | Both Isoforms |
| HOXB5 | -0.73 | -0.73 | Both Isoforms | TFCP2L1 | -1.62 | -0.86 | Both Isoforms |
| CLSPN | -1.06 | -0.73 | Both Isoforms | ZNF853 | -2.00 | -0.86 | Both Isoforms |

| | | | | | | | |
|---|---|---|---|---|---|---|---|
| KATNAL2 | -1.00 | -0.72 | Both Isoforms | C4orf50 | -1.86 | -0.86 | Both Isoforms |
| DNAH10 | -1.43 | -0.72 | Both Isoforms | LAMA3 | -0.97 | -0.86 | Both Isoforms |
| RECQL4 | -0.86 | -0.72 | Both Isoforms | PAX6 | -1.15 | -0.86 | Both Isoforms |
| SLC34A3 | -0.61 | -0.72 | Both Isoforms | BHLHE41 | -1.05 | -0.86 | Both Isoforms |
| OIP5 | -0.85 | -0.72 | Both Isoforms | NLGN3 | -0.72 | -0.86 | Both Isoforms |
| CYFIP2 | -0.91 | -0.72 | Both Isoforms | PELI2 | -1.87 | -0.86 | Both Isoforms |
| BEND3 | -0.60 | -0.72 | Both Isoforms | N4BP2 | -0.71 | -0.86 | Both Isoforms |
| CRMP1 | -0.97 | -0.71 | Both Isoforms | NFU1 | -1.20 | -0.86 | Both Isoforms |
| RCAN3 | -0.90 | -0.71 | Both Isoforms | GRB10 | -1.69 | -0.86 | Both Isoforms |
| ORC1 | -1.12 | -0.71 | Both Isoforms | PDE10A | -1.32 | -0.86 | Both Isoforms |
| RIMKLA | -0.65 | -0.71 | Both Isoforms | TAS1R3 | -1.64 | -0.86 | Both Isoforms |
| GALNT10 | -0.62 | -0.71 | Both Isoforms | EFCC1 | -1.54 | -0.86 | Both Isoforms |
| FERMT1 | -0.67 | -0.71 | Both Isoforms | FBXO2 | -1.71 | -0.86 | Both Isoforms |
| TP53 | -1.12 | -0.71 | Both Isoforms | FAM49A | -1.45 | -0.86 | Both Isoforms |
| ZC3H6 | -0.78 | -0.71 | Both Isoforms | GSC | -0.86 | -0.85 | Both Isoforms |
| TMEM38B | -1.23 | -0.71 | Both Isoforms | DPY19L1 | -0.81 | -0.85 | Both Isoforms |
| ANKRD6 | -1.12 | -0.70 | Both Isoforms | ZNF503 | -1.33 | -0.85 | Both Isoforms |
| DENND1B | -1.16 | -0.70 | Both Isoforms | RDH16 | -1.03 | -0.85 | Both Isoforms |
| EPM2A | -1.13 | -0.70 | Both Isoforms | DNAJC12 | -0.96 | -0.85 | Both Isoforms |
| BID | -1.06 | -0.70 | Both Isoforms | THEM5 | -1.90 | -0.85 | Both Isoforms |
| DSCC1 | -0.88 | -0.70 | Both Isoforms | ZCWPW1 | -1.15 | -0.85 | Both Isoforms |
| ITGB4 | -1.46 | -0.70 | Both Isoforms | TMEM45A | -2.15 | -0.85 | Both Isoforms |
| POLD1 | -0.77 | -0.70 | Both Isoforms | MYRF | -1.66 | -0.85 | Both Isoforms |
| NIPSNAP3A | -1.19 | -0.70 | Both Isoforms | ZMYND12 | -1.52 | -0.85 | Both Isoforms |
| SSTR5 | -2.85 | -0.70 | Both Isoforms | TPSG1 | -0.76 | -0.85 | Both Isoforms |

| | | | |
|---|---|---|---|
| DAPK2 | -1.44 | -0.69 | Both Isoforms |
| MVB12B | -0.88 | -0.69 | Both Isoforms |
| RCC2 | -0.63 | -0.69 | Both Isoforms |
| BMF | -1.96 | -0.69 | Both Isoforms |
| VASN | -0.63 | -0.69 | Both Isoforms |
| MCM2 | -1.23 | -0.69 | Both Isoforms |
| MGAT4A | -0.66 | -0.69 | Both Isoforms |
| TP53INP1 | -1.47 | -0.69 | Both Isoforms |
| MANBA | -1.08 | -0.69 | Both Isoforms |
| MB | -1.22 | -0.69 | Both Isoforms |
| NFIX | -0.72 | -0.69 | Both Isoforms |
| TMEM229B | -1.31 | -0.69 | Both Isoforms |
| CLCN5 | -0.63 | -0.69 | Both Isoforms |
| MCM4 | -1.02 | -0.69 | Both Isoforms |
| SLC25A19 | -1.04 | -0.68 | Both Isoforms |
| EVC | -0.81 | -0.68 | Both Isoforms |
| PPP1R14C | -2.44 | -0.68 | Both Isoforms |
| BBC3 | -1.17 | -0.68 | Both Isoforms |
| GPC4 | -0.69 | -0.68 | Both Isoforms |
| OAF | -0.58 | -0.68 | Both Isoforms |
| ITPR1 | -0.64 | -0.68 | Both Isoforms |
| APOBEC3B | -0.69 | -0.68 | Both Isoforms |
| GAS6 | -0.87 | -0.68 | Both Isoforms |
| GAREML | -0.73 | -0.68 | Both Isoforms |
| CRAT | -0.83 | -0.68 | Both Isoforms |

| | | | |
|---|---|---|---|
| LIX1L | -1.68 | -0.85 | Both Isoforms |
| DRC7 | -0.85 | -0.85 | Both Isoforms |
| SYNE1 | -0.83 | -0.85 | Both Isoforms |
| CPT1B | -0.87 | -0.84 | Both Isoforms |
| H2AFJ | -0.82 | -0.84 | Both Isoforms |
| GSTA1 | -1.08 | -0.84 | Both Isoforms |
| PLIN1 | -1.47 | -0.84 | Both Isoforms |
| RCAN3 | -0.87 | -0.84 | Both Isoforms |
| UBA7 | -0.84 | -0.84 | Both Isoforms |
| CDK14 | -0.85 | -0.84 | Both Isoforms |
| GID4 | -0.98 | -0.84 | Both Isoforms |
| ZNF572 | -1.38 | -0.84 | Both Isoforms |
| SETSIP | -0.66 | -0.84 | Both Isoforms |
| FABP5 | -2.23 | -0.84 | Both Isoforms |
| RNF212 | -2.33 | -0.84 | Both Isoforms |
| DZIP3 | -1.43 | -0.84 | Both Isoforms |
| TARSL2 | -1.15 | -0.84 | Both Isoforms |
| HECA | -1.28 | -0.84 | Both Isoforms |
| PPM1M | -1.17 | -0.84 | Both Isoforms |
| BEX4 | -1.40 | -0.84 | Both Isoforms |
| TEX15 | -1.71 | -0.84 | Both Isoforms |
| LTBP2 | -1.96 | -0.83 | Both Isoforms |
| ZNF391 | -1.69 | -0.83 | Both Isoforms |
| STARD9 | -1.66 | -0.83 | Both Isoforms |
| C12orf56 | -0.58 | -0.83 | Both Isoforms |

| | | | | | | | | |
|---|---|---|---|---|---|---|---|---|
| | 6-Sep | -1.13 | -0.68 | Both Isoforms | GSN | -0.67 | -0.83 | Both Isoforms |
| MTURN | | -0.97 | -0.68 | Both Isoforms | RGL3 | -2.15 | -0.83 | Both Isoforms |
| SLC9A5 | | -0.66 | -0.68 | Both Isoforms | KIAA1257 | -1.28 | -0.83 | Both Isoforms |
| TIPIN | | -0.77 | -0.68 | Both Isoforms | TRIM29 | -1.57 | -0.83 | Both Isoforms |
| ARHGAP20 | | -2.35 | -0.68 | Both Isoforms | SESN3 | -2.38 | -0.83 | Both Isoforms |
| TUBB4A | | -1.48 | -0.68 | Both Isoforms | DOCK2 | -1.34 | -0.83 | Both Isoforms |
| CDT1 | | -1.11 | -0.67 | Both Isoforms | PTPN14 | -0.80 | -0.83 | Both Isoforms |
| FANCD2 | | -0.84 | -0.67 | Both Isoforms | RNF180 | -1.01 | -0.83 | Both Isoforms |
| FIGNL1 | | -1.14 | -0.67 | Both Isoforms | TTLL6 | -0.83 | -0.83 | Both Isoforms |
| GATS | | -0.63 | -0.67 | Both Isoforms | PCDHB11 | -0.90 | -0.83 | Both Isoforms |
| GJC1 | | -0.80 | -0.67 | Both Isoforms | SLC9B2 | -2.03 | -0.83 | Both Isoforms |
| BTG2 | | -1.08 | -0.67 | Both Isoforms | RGS10 | -0.78 | -0.83 | Both Isoforms |
| LRIG3 | | -1.30 | -0.66 | Both Isoforms | PLCE1 | -1.17 | -0.82 | Both Isoforms |
| PIGZ | | -1.08 | -0.66 | Both Isoforms | FOXB2 | -2.21 | -0.82 | Both Isoforms |
| POLQ | | -1.07 | -0.66 | Both Isoforms | THG1L | -1.39 | -0.82 | Both Isoforms |
| F12 | | -0.61 | -0.66 | Both Isoforms | AUH | -0.86 | -0.82 | Both Isoforms |
| BUB1B | | -0.94 | -0.66 | Both Isoforms | AIDA | -1.68 | -0.82 | Both Isoforms |
| PKN3 | | -0.73 | -0.66 | Both Isoforms | NLRP1 | -2.43 | -0.82 | Both Isoforms |
| APCDD1 | | -1.18 | -0.66 | Both Isoforms | SMIM10L2A | -2.03 | -0.82 | Both Isoforms |
| GALNT3 | | -1.27 | -0.66 | Both Isoforms | MBOAT1 | -1.85 | -0.82 | Both Isoforms |
| ESCO2 | | -0.96 | -0.66 | Both Isoforms | SVIL | -1.51 | -0.82 | Both Isoforms |
| PAK1 | | -1.29 | -0.65 | Both Isoforms | PCDHA13 | -1.36 | -0.82 | Both Isoforms |
| OCLN | | -0.96 | -0.65 | Both Isoforms | MYC | -2.21 | -0.82 | Both Isoforms |
| NKD2 | | -1.37 | -0.65 | Both Isoforms | RBM7 | -0.59 | -0.82 | Both Isoforms |
| NDUFAF6 | | -0.83 | -0.65 | Both Isoforms | ZNF230 | -0.89 | -0.82 | Both Isoforms |

| | | | | | | | |
|---|---|---|---|---|---|---|---|
| SNRK | -0.64 | -0.65 | Both Isoforms | RYR1 | -2.70 | -0.82 | Both Isoforms |
| FBXO2 | -2.16 | -0.65 | Both Isoforms | CAMLG | -1.01 | -0.82 | Both Isoforms |
| FAM184A | -1.10 | -0.65 | Both Isoforms | GPR27 | -2.28 | -0.82 | Both Isoforms |
| E2F8 | -0.99 | -0.65 | Both Isoforms | SLC6A11 | -2.17 | -0.81 | Both Isoforms |
| PARD6G | -0.85 | -0.65 | Both Isoforms | JDP2 | -1.58 | -0.81 | Both Isoforms |
| MCM7 | -0.85 | -0.65 | Both Isoforms | EIF3CL | -1.30 | -0.81 | Both Isoforms |
| PHC1 | -0.88 | -0.65 | Both Isoforms | NMNAT2 | -1.23 | -0.81 | Both Isoforms |
| C14orf80 | -0.89 | -0.64 | Both Isoforms | PRR18 | -2.26 | -0.81 | Both Isoforms |
| PTPRR | -1.45 | -0.64 | Both Isoforms | ITGA1 | -1.04 | -0.81 | Both Isoforms |
| FBXO24 | -1.01 | -0.64 | Both Isoforms | NMT2 | -1.62 | -0.81 | Both Isoforms |
| ASF1B | -1.20 | -0.64 | Both Isoforms | AR | -2.83 | -0.81 | Both Isoforms |
| CPNE1 | -0.60 | -0.64 | Both Isoforms | SLC25A18 | -0.86 | -0.81 | Both Isoforms |
| COL18A1 | -0.87 | -0.64 | Both Isoforms | MAP3K8 | -0.64 | -0.81 | Both Isoforms |
| MYC | -0.66 | -0.64 | Both Isoforms | CNNM2 | -0.91 | -0.81 | Both Isoforms |
| DTL | -1.16 | -0.64 | Both Isoforms | ATP11C | -0.82 | -0.81 | Both Isoforms |
| LMNB1 | -0.88 | -0.64 | Both Isoforms | NLN | -1.16 | -0.81 | Both Isoforms |
| FHOD3 | -1.80 | -0.63 | Both Isoforms | SIX1 | -1.09 | -0.81 | Both Isoforms |
| GRB14 | -0.71 | -0.63 | Both Isoforms | RAI14 | -0.60 | -0.80 | Both Isoforms |
| RBL1 | -0.99 | -0.63 | Both Isoforms | MYO7A | -2.69 | -0.80 | Both Isoforms |
| ADORA1 | -2.40 | -0.63 | Both Isoforms | SCNN1G | -1.31 | -0.80 | Both Isoforms |
| SLC6A19 | -1.55 | -0.63 | Both Isoforms | VWDE | -1.79 | -0.80 | Both Isoforms |
| TMPRSS13 | -1.43 | -0.63 | Both Isoforms | RILPL2 | -0.97 | -0.80 | Both Isoforms |
| CELSR2 | -1.08 | -0.63 | Both Isoforms | PCP4 | -0.59 | -0.80 | Both Isoforms |
| CENPM | -0.69 | -0.63 | Both Isoforms | PTCH1 | -1.33 | -0.80 | Both Isoforms |
| DNMT1 | -0.64 | -0.63 | Both Isoforms | ASB14 | -0.62 | -0.80 | Both Isoforms |

| | | | | | | | |
|---|---|---|---|---|---|---|---|
| LRRC37A3 | -1.00 | -0.63 | Both Isoforms | CYBB | -1.05 | -0.80 | Both Isoforms |
| PRR36 | -0.66 | -0.63 | Both Isoforms | MREG | -1.09 | -0.80 | Both Isoforms |
| GLIS2 | -1.00 | -0.63 | Both Isoforms | SNX30 | -1.56 | -0.80 | Both Isoforms |
| PER3 | -0.70 | -0.62 | Both Isoforms | ZNF296 | -1.58 | -0.80 | Both Isoforms |
| RAB30 | -0.64 | -0.62 | Both Isoforms | PLCG2 | -1.49 | -0.80 | Both Isoforms |
| CHST15 | -0.65 | -0.62 | Both Isoforms | ABTB1 | -1.45 | -0.80 | Both Isoforms |
| WDR76 | -1.02 | -0.62 | Both Isoforms | PROZ | -1.42 | -0.80 | Both Isoforms |
| SKA1 | -0.92 | -0.62 | Both Isoforms | PDP1 | -0.84 | -0.79 | Both Isoforms |
| TRNP1 | -1.28 | -0.61 | Both Isoforms | DOCK10 | -2.74 | -0.79 | Both Isoforms |
| TUB | -0.66 | -0.61 | Both Isoforms | ZNF284 | -1.72 | -0.79 | Both Isoforms |
| ST6GALNAC4 | -1.05 | -0.61 | Both Isoforms | DOCK3 | -1.09 | -0.79 | Both Isoforms |
| DDB2 | -1.28 | -0.61 | Both Isoforms | MAPRE2 | -1.11 | -0.79 | Both Isoforms |
| E2F7 | -1.05 | -0.61 | Both Isoforms | PIFO | -0.68 | -0.79 | Both Isoforms |
| ADAM22 | -0.77 | -0.61 | Both Isoforms | BAHCC1 | -2.31 | -0.79 | Both Isoforms |
| ZNF680 | -0.74 | -0.61 | Both Isoforms | GSTA4 | -1.39 | -0.79 | Both Isoforms |
| ARHGAP29 | -0.97 | -0.61 | Both Isoforms | SAMD11 | -0.71 | -0.79 | Both Isoforms |
| N4BP3 | -1.01 | -0.61 | Both Isoforms | PCDHB10 | -0.85 | -0.79 | Both Isoforms |
| UHRF1 | -1.13 | -0.60 | Both Isoforms | KLK14 | -1.17 | -0.79 | Both Isoforms |
| TMCC2 | -0.72 | -0.60 | Both Isoforms | NOA1 | -0.69 | -0.79 | Both Isoforms |
| CHAF1A | -0.88 | -0.60 | Both Isoforms | TMOD2 | -1.72 | -0.79 | Both Isoforms |
| CHAF1B | -0.95 | -0.60 | Both Isoforms | GABRD | -0.74 | -0.79 | Both Isoforms |
| EPHA10 | -0.58 | -0.60 | Both Isoforms | GRAMD1C | -0.86 | -0.79 | Both Isoforms |
| GNG7 | -0.81 | -0.60 | Both Isoforms | IL22RA1 | -1.50 | -0.79 | Both Isoforms |
| RAD51 | -0.96 | -0.60 | Both Isoforms | CREBRF | -1.51 | -0.79 | Both Isoforms |
| FAM182B | -0.83 | -0.60 | Both Isoforms | RAC2 | -1.35 | -0.78 | Both Isoforms |

| | | | | | | | |
|---|---|---|---|---|---|---|---|
| TBX19 | -1.08 | -0.60 | Both Isoforms | CDK7 | -0.90 | -0.78 | Both Isoforms |
| ASCL5 | -0.76 | -0.60 | Both Isoforms | CCL2 | -2.13 | -0.78 | Both Isoforms |
| FBXO5 | -0.63 | -0.60 | Both Isoforms | SIDT1 | -1.36 | -0.78 | Both Isoforms |
| FANCA | -0.77 | -0.60 | Both Isoforms | SMAD1 | -1.24 | -0.78 | Both Isoforms |
| CENPH | -0.68 | -0.60 | Both Isoforms | SCAI | -0.73 | -0.78 | Both Isoforms |
| WNT4 | -1.36 | -0.59 | Both Isoforms | ADORA2B | -1.30 | -0.78 | Both Isoforms |
| BEND7 | -1.10 | -0.59 | Both Isoforms | PROSER2 | -0.68 | -0.78 | Both Isoforms |
| ABTB2 | -1.22 | -0.59 | Both Isoforms | SPATS2L | -0.87 | -0.78 | Both Isoforms |
| GLCE | -0.61 | -0.59 | Both Isoforms | RPS6KA6 | -1.20 | -0.78 | Both Isoforms |
| SOX11 | -1.06 | -0.59 | Both Isoforms | ACVR2B | -0.66 | -0.78 | Both Isoforms |
| PCDHB11 | -0.60 | -0.59 | Both Isoforms | TGIF2 | -1.19 | -0.78 | Both Isoforms |
| FOXM1 | -0.66 | -0.59 | Both Isoforms | POU3F3 | -1.82 | -0.78 | Both Isoforms |
| ADGRL1 | -0.62 | -0.59 | Both Isoforms | NT5C | -1.11 | -0.78 | Both Isoforms |
| WDR54 | -0.65 | -0.58 | Both Isoforms | GLI2 | -1.97 | -0.78 | Both Isoforms |
| VWA2 | -3.56 | -0.58 | Both Isoforms | PTK7 | -1.32 | -0.78 | Both Isoforms |
| SORL1 | -1.71 | -0.58 | Both Isoforms | ADPRHL1 | -1.12 | -0.78 | Both Isoforms |
| LNX2 | -0.73 | -0.58 | Both Isoforms | DMGDH | -1.44 | -0.78 | Both Isoforms |
| CBX6 | -0.80 | -0.58 | AR-FL | AGA | -0.83 | -0.78 | Both Isoforms |
| LAPTM4B | -0.98 | -0.58 | AR-FL | EIF4B | -1.03 | -0.78 | Both Isoforms |
| AC013461.1 | -0.61 | -0.58 | AR-FL | ARRDC3 | -1.44 | -0.78 | Both Isoforms |
| ADD2 | -2.39 | -0.57 | AR-FL | PGM2 | -1.15 | -0.78 | Both Isoforms |
| HMGA1 | -0.81 | -0.57 | AR-FL | MAP2K5 | -1.00 | -0.78 | Both Isoforms |
| ARMCX4 | -0.74 | -0.57 | AR-FL | OSR2 | -1.29 | -0.78 | Both Isoforms |
| NCAPH | -0.99 | -0.57 | AR-FL | LNPEP | -0.86 | -0.78 | Both Isoforms |
| DPY19L1 | -0.98 | -0.57 | AR-FL | CERK | -2.01 | -0.77 | Both Isoforms |
| DACT3 | -1.08 | -0.57 | AR-FL | ZNF658 | -1.13 | -0.77 | Both Isoforms |
| BMP8B | -0.70 | -0.57 | AR-FL | FAAH2 | -0.97 | -0.77 | Both Isoforms |
| ADCY3 | -0.65 | -0.57 | AR-FL | TMCO6 | -1.58 | -0.77 | Both Isoforms |
| GEMIN4 | -0.64 | -0.57 | AR-FL | PARD6G | -1.17 | -0.77 | Both Isoforms |
| HOXB3 | -0.67 | -0.57 | AR-FL | TMEM136 | -1.44 | -0.77 | Both Isoforms |
| PODXL | -0.77 | -0.57 | AR-FL | PCNXL2 | -0.62 | -0.77 | Both Isoforms |
| ME2 | -0.86 | -0.57 | AR-FL | PRAC1 | -1.09 | -0.77 | Both Isoforms |

| | | | |
|---|---|---|---|
| CENPV | -0.65 | -0.57 | AR-FL |
| XRCC6BP1 | -0.59 | -0.56 | AR-FL |
| ENHO | -0.97 | -0.56 | AR-FL |
| GLDC | -1.14 | -0.56 | AR-FL |
| TLL2 | -1.08 | -0.56 | AR-FL |
| SLC9A7 | -0.84 | -0.56 | AR-FL |
| CDCA5 | -1.08 | -0.56 | AR-FL |
| PKMYT1 | -1.04 | -0.56 | AR-FL |
| MKI67 | -0.63 | -0.56 | AR-FL |
| BRCA1 | -1.01 | -0.56 | AR-FL |
| FUT8 | -0.84 | -0.56 | AR-FL |
| WNT10B | -0.79 | -0.55 | AR-FL |
| SESN2 | -1.16 | -0.55 | AR-FL |
| GTF2IRD2B | -1.16 | -0.55 | AR-FL |
| EML4 | -0.58 | -0.55 | AR-FL |
| CCNA2 | -0.78 | -0.55 | AR-FL |
| PIK3C2B | -0.73 | -0.55 | AR-FL |
| CCDC102B | -1.29 | -0.55 | AR-FL |
| NOTCH1 | -0.80 | -0.55 | AR-FL |
| ADGRB2 | -1.79 | -0.55 | AR-FL |
| ST5 | -1.36 | -0.55 | AR-FL |
| ITGA2 | -0.87 | -0.55 | AR-FL |
| ESPN | -1.97 | -0.55 | AR-FL |
| RALGPS1 | -0.75 | -0.55 | AR-FL |
| ZNRF3 | -0.59 | -0.55 | AR-FL |
| RNF208 | -0.90 | -0.55 | AR-FL |
| ATXN7L2 | -0.64 | -0.55 | AR-FL |
| CACNA2D2 | -0.82 | -0.54 | AR-FL |
| CD83 | -1.52 | -0.54 | AR-FL |
| FANCB | -1.00 | -0.54 | AR-FL |
| ZMYND15 | -1.06 | -0.54 | AR-FL |
| OTUD3 | -0.62 | -0.54 | AR-FL |
| LTA4H | -0.69 | -0.54 | AR-FL |
| RNASEH2A | -0.65 | -0.54 | AR-FL |
| TMEM144 | -1.14 | -0.54 | AR-FL |
| EFCAB11 | -1.02 | -0.54 | AR-FL |
| MAP7D3 | -1.13 | -0.54 | AR-FL |
| HOXB6 | -0.77 | -0.54 | AR-FL |
| CCDC151 | -1.03 | -0.54 | AR-FL |
| AGO4 | -0.78 | -0.54 | AR-FL |
| SPTLC3 | -1.55 | -0.53 | AR-FL |
| CCDC92 | -0.79 | -0.53 | AR-FL |
| STXBP6 | -1.29 | -0.53 | AR-FL |
| NIPAL1 | -1.00 | -0.53 | AR-FL |
| TNFRSF21 | -0.93 | -0.53 | AR-FL |
| FDX1 | -0.64 | -0.53 | AR-FL |
| THNSL1 | -2.07 | -0.77 | Both Isoforms |
| HTRA3 | -1.34 | -0.77 | Both Isoforms |
| CRTAM | -1.33 | -0.77 | Both Isoforms |
| CTSK | -0.75 | -0.77 | Both Isoforms |
| ZNF704 | -1.26 | -0.77 | Both Isoforms |
| PKHD1 | -0.86 | -0.77 | Both Isoforms |
| IFRD1 | -0.70 | -0.77 | Both Isoforms |
| KCNJ2 | -1.23 | -0.77 | Both Isoforms |
| RAB6C | -0.77 | -0.77 | Both Isoforms |
| LSM11 | -1.60 | -0.76 | Both Isoforms |
| RBMS2 | -0.97 | -0.76 | Both Isoforms |
| NCOA2 | -1.03 | -0.76 | Both Isoforms |
| ELOVL4 | -0.78 | -0.76 | Both Isoforms |
| PCID2 | -0.85 | -0.76 | Both Isoforms |
| C1orf145 | -2.26 | -0.76 | Both Isoforms |
| LACE1 | -0.59 | -0.76 | Both Isoforms |
| ACY3 | -0.80 | -0.76 | Both Isoforms |
| FN1 | -0.62 | -0.76 | Both Isoforms |
| TTC23 | -1.37 | -0.76 | Both Isoforms |
| POLR3C | -1.19 | -0.76 | Both Isoforms |
| DSEL | -2.24 | -0.76 | Both Isoforms |
| FAM83G | -0.66 | -0.76 | Both Isoforms |
| RGS22 | -1.35 | -0.76 | Both Isoforms |
| ERICH2 | -1.65 | -0.76 | Both Isoforms |
| CTNND2 | -0.87 | -0.76 | Both Isoforms |
| SDCCAG8 | -0.96 | -0.76 | Both Isoforms |
| EFCAB2 | -0.88 | -0.76 | Both Isoforms |
| C14orf37 | -2.24 | -0.76 | Both Isoforms |
| NBPF1 | -1.74 | -0.76 | Both Isoforms |
| ZNF667 | -1.76 | -0.76 | Both Isoforms |
| ARNT2 | -0.93 | -0.75 | Both Isoforms |
| IMPDH2 | -1.43 | -0.75 | Both Isoforms |
| CCDC191 | -1.98 | -0.75 | Both Isoforms |
| WSCD1 | -1.94 | -0.75 | Both Isoforms |
| RBBP8NL | -1.08 | -0.75 | Both Isoforms |
| MBLAC2 | -1.15 | -0.75 | Both Isoforms |
| FAM134B | -1.57 | -0.75 | Both Isoforms |
| ACER3 | -1.00 | -0.75 | Both Isoforms |
| BANK1 | -0.66 | -0.75 | Both Isoforms |
| CDH2 | -2.95 | -0.75 | Both Isoforms |
| TMEM123 | -0.67 | -0.75 | Both Isoforms |
| KCNK15 | -3.03 | -0.75 | Both Isoforms |
| CLIP4 | -1.55 | -0.75 | Both Isoforms |
| FEZ1 | -1.69 | -0.75 | Both Isoforms |
| IMMP2L | -0.67 | -0.75 | Both Isoforms |
| PHC1 | -1.68 | -0.75 | Both Isoforms |

| | | | | | | | |
|---|---|---|---|---|---|---|---|
| TIAM2 | -0.94 | -0.53 | AR-FL | ZNF182 | -0.95 | -0.75 | Both Isoforms |
| MELK | -0.75 | -0.53 | AR-FL | LAMP3 | -0.75 | -0.75 | Both Isoforms |
| LOXL3 | -1.79 | -0.53 | AR-FL | KIZ | -1.18 | -0.75 | Both Isoforms |
| C11orf70 | -1.89 | -0.53 | AR-FL | WNT2B | -2.00 | -0.74 | Both Isoforms |
| PLEKHS1 | -1.51 | -0.53 | AR-FL | EYA3 | -1.10 | -0.74 | Both Isoforms |
| KIAA0101 | -0.63 | -0.53 | AR-FL | ZNF559-ZNF177 | -1.08 | -0.74 | Both Isoforms |
| THG1L | -0.73 | -0.53 | AR-FL | PDLIM2 | -0.64 | -0.74 | Both Isoforms |
| CDC6 | -0.62 | -0.53 | AR-FL | CALCOCO1 | -1.58 | -0.74 | Both Isoforms |
| BCAM | -0.79 | -0.52 | AR-FL | HIST1H2BF | -0.73 | -0.74 | Both Isoforms |
| CACNB2 | -1.20 | -0.52 | AR-FL | PARP3 | -1.30 | -0.74 | Both Isoforms |
| TTC23L | -1.22 | -0.52 | AR-FL | MPP7 | -0.67 | -0.74 | Both Isoforms |
| INPP5J | -0.74 | -0.52 | AR-FL | ODAM | -0.85 | -0.74 | Both Isoforms |
| LIG1 | -0.83 | -0.52 | AR-FL | HUNK | -0.95 | -0.74 | Both Isoforms |
| EPHB6 | -0.80 | -0.52 | AR-FL | POLG2 | -1.43 | -0.74 | Both Isoforms |
| IMPDH2 | -0.60 | -0.52 | AR-FL | PEX11G | -0.73 | -0.74 | Both Isoforms |
| ZNF462 | -0.71 | -0.52 | AR-FL | ADCY7 | -2.54 | -0.74 | Both Isoforms |
| RAB17 | -0.67 | -0.52 | AR-FL | CNOT6L | -0.99 | -0.74 | Both Isoforms |
| NT5M | -0.62 | -0.52 | AR-FL | PLCH1 | -0.67 | -0.73 | Both Isoforms |
| GJB1 | -0.59 | -0.52 | AR-FL | DPH5 | -1.30 | -0.73 | Both Isoforms |
| TSPAN15 | -1.05 | -0.52 | AR-FL | XRN2 | -0.71 | -0.73 | Both Isoforms |
| SYNE2 | -0.65 | -0.52 | AR-FL | DHRS7 | -0.62 | -0.73 | Both Isoforms |
| LHX4 | -0.67 | -0.52 | AR-FL | RFX3 | -1.28 | -0.73 | Both Isoforms |
| RNF130 | -0.59 | -0.52 | AR-FL | TRIQK | -0.91 | -0.73 | Both Isoforms |
| NPIPA7 | -0.81 | -0.52 | AR-FL | CPNE1 | -0.79 | -0.73 | Both Isoforms |
| ATXN7L1 | -0.85 | -0.52 | AR-FL | PCMTD2 | -1.28 | -0.73 | Both Isoforms |
| C1orf233 | -0.58 | -0.51 | AR-FL | CFAP161 | -0.96 | -0.73 | Both Isoforms |
| SDHAF3 | -0.60 | -0.51 | AR-FL | IGFBP4 | -1.19 | -0.73 | Both Isoforms |
| DGKH | -1.70 | -0.51 | AR-FL | NFKB1 | -1.35 | -0.73 | Both Isoforms |
| GTF2IRD2 | -1.45 | -0.51 | AR-FL | RPL22L1 | -0.67 | -0.73 | Both Isoforms |
| ADORA2B | -0.85 | -0.51 | AR-FL | DPYSL3 | -0.89 | -0.73 | Both Isoforms |
| C5orf34 | -1.03 | -0.51 | AR-FL | KIF12 | -1.24 | -0.73 | Both Isoforms |
| GSC | -0.67 | -0.51 | AR-FL | CDK19 | -0.76 | -0.73 | Both Isoforms |
| C1orf53 | -0.89 | -0.51 | AR-FL | DZANK1 | -0.75 | -0.73 | Both Isoforms |
| FAM212B | -0.98 | -0.51 | AR-FL | STOX1 | -3.08 | -0.73 | Both Isoforms |
| ASTN2 | -0.94 | -0.51 | AR-FL | DOK7 | -0.84 | -0.73 | Both Isoforms |
| GARNL3 | -1.05 | -0.51 | AR-FL | NHSL2 | -1.40 | -0.72 | Both Isoforms |
| DOK1 | -1.57 | -0.51 | AR-FL | PRELID3A | -1.00 | -0.72 | Both Isoforms |
| FANCI | -0.60 | -0.50 | AR-FL | IPMK | -0.96 | -0.72 | Both Isoforms |
| MLXIPL | -1.62 | -0.50 | AR-FL | SMAGP | -0.65 | -0.72 | Both Isoforms |
| STRBP | -0.98 | -0.50 | AR-FL | ANKRD46 | -0.63 | -0.72 | Both Isoforms |
| TCOF1 | -0.78 | -0.50 | AR-FL | ANKRD16 | -1.37 | -0.72 | Both Isoforms |
| GAS2L3 | -0.81 | -0.50 | AR-FL | CAMK2N1 | -1.60 | -0.72 | Both Isoforms |
| PMS2 | -0.86 | -0.50 | AR-FL | UPK3A | -1.96 | -0.72 | Both Isoforms |
| RFX3 | -1.61 | -0.50 | AR-FL | MYT1 | -0.83 | -0.72 | Both Isoforms |
| ISYNA1 | -0.79 | -0.50 | AR-FL | IFIT2 | -0.91 | -0.72 | Both Isoforms |
| HSD17B6 | -0.69 | -0.50 | AR-FL | MRM1 | -1.15 | -0.72 | Both Isoforms |

| | | | | | | | |
|---|---|---|---|---|---|---|---|
| SH3RF1 | -1.07 | -0.50 | AR-FL | TMEM243 | -1.01 | -0.72 | Both Isoforms |
| DCPS | -0.61 | -0.49 | AR-FL | KLHL35 | -1.49 | -0.72 | Both Isoforms |
| HRSP12 | -0.89 | -0.49 | AR-FL | UGT8 | -1.47 | -0.72 | Both Isoforms |
| ERV3-1 | -0.63 | -0.49 | AR-FL | NR1D1 | -1.37 | -0.72 | Both Isoforms |
| RELL1 | -1.06 | -0.49 | AR-FL | ATG2B | -1.15 | -0.72 | Both Isoforms |
| KIF18B | -1.03 | -0.49 | AR-FL | CCDC14 | -1.16 | -0.72 | Both Isoforms |
| LIN9 | -1.13 | -0.49 | AR-FL | PAQR8 | -1.25 | -0.72 | Both Isoforms |
| NFAT5 | -0.90 | -0.49 | AR-FL | BAZ2B | -0.97 | -0.71 | Both Isoforms |
| NAT8L | -1.69 | -0.48 | AR-FL | CYTH3 | -1.05 | -0.71 | Both Isoforms |
| EZH2 | -0.73 | -0.48 | AR-FL | RNLS | -0.70 | -0.71 | Both Isoforms |
| KIAA1958 | -0.79 | -0.48 | AR-FL | RNF217 | -2.15 | -0.71 | Both Isoforms |
| AJUBA | -1.22 | -0.48 | AR-FL | GRB14 | -1.77 | -0.71 | Both Isoforms |
| HECTD2 | -0.77 | -0.48 | AR-FL | STON2 | -1.09 | -0.71 | Both Isoforms |
| RFC2 | -0.68 | -0.48 | AR-FL | IRX2 | -0.61 | -0.71 | Both Isoforms |
| CYB5RL | -0.69 | -0.48 | AR-FL | ADA | -1.18 | -0.71 | Both Isoforms |
| FAM149A | -0.71 | -0.48 | AR-FL | GPR156 | -1.34 | -0.71 | Both Isoforms |
| MACROD1 | -0.58 | -0.48 | AR-FL | RBMX | -0.74 | -0.71 | Both Isoforms |
| RNF144A | -0.69 | -0.48 | AR-FL | PIWIL2 | -5.30 | -0.71 | Both Isoforms |
| RTKN2 | -0.65 | -0.48 | AR-FL | PTPRU | -1.74 | -0.71 | Both Isoforms |
| ARHGAP11A | -0.61 | -0.47 | AR-FL | NT5C3A | -0.66 | -0.71 | Both Isoforms |
| SALL2 | -1.31 | -0.47 | AR-FL | ZC3H8 | -0.85 | -0.71 | Both Isoforms |
| PGAP1 | -1.20 | -0.47 | AR-FL | ARMC4 | -2.99 | -0.71 | Both Isoforms |
| RGL1 | -0.74 | -0.47 | AR-FL | SETDB2 | -0.70 | -0.71 | Both Isoforms |
| TPX2 | -0.71 | -0.47 | AR-FL | GLTSCR1L | -0.82 | -0.71 | Both Isoforms |
| KANK2 | -0.76 | -0.47 | AR-FL | CCDC171 | -1.44 | -0.71 | Both Isoforms |
| NFKB1 | -0.90 | -0.47 | AR-FL | EIF4E3 | -2.13 | -0.70 | Both Isoforms |
| H2AFY2 | -0.71 | -0.47 | AR-FL | PLCB4 | -1.04 | -0.70 | Both Isoforms |
| CENPA | -0.66 | -0.47 | AR-FL | FKBP7 | -0.70 | -0.70 | Both Isoforms |
| INSIG2 | -0.70 | -0.47 | AR-FL | BID | -0.64 | -0.70 | Both Isoforms |
| PIK3R3 | -1.06 | -0.47 | AR-FL | IRF2BP2 | -1.07 | -0.70 | Both Isoforms |
| RTN4R | -0.94 | -0.47 | AR-FL | ELN | -0.78 | -0.70 | Both Isoforms |
| LRP5L | -0.67 | -0.47 | AR-FL | PROS1 | -0.77 | -0.70 | Both Isoforms |
| PLK1 | -0.64 | -0.46 | AR-FL | IGBP1 | -1.02 | -0.70 | Both Isoforms |
| ATP11A | -0.93 | -0.46 | AR-FL | SAMD5 | -1.49 | -0.70 | Both Isoforms |
| RBBP8 | -0.73 | -0.46 | AR-FL | TRAM1L1 | -0.83 | -0.70 | Both Isoforms |
| SLC22A2 | -1.41 | -0.46 | AR-FL | KCNJ14 | -1.32 | -0.70 | Both Isoforms |
| RNF19A | -0.59 | -0.46 | AR-FL | CCDC177 | -0.91 | -0.70 | Both Isoforms |
| MAD2L1 | -0.69 | -0.46 | AR-FL | METTL20 | -0.72 | -0.70 | Both Isoforms |
| PROZ | -0.71 | -0.46 | AR-FL | SH2D2A | -1.95 | -0.69 | Both Isoforms |
| KIF15 | -0.88 | -0.46 | AR-FL | FOXN2 | -0.61 | -0.69 | Both Isoforms |
| IRX2 | -0.66 | -0.46 | AR-FL | CAMK2D | -1.46 | -0.69 | Both Isoforms |
| KIF11 | -0.78 | -0.46 | AR-FL | NTM | -1.31 | -0.69 | Both Isoforms |
| BZRAP1 | -0.70 | -0.46 | AR-FL | PRR5L | -1.53 | -0.69 | Both Isoforms |
| CALD1 | -0.76 | -0.45 | AR-FL | HAUS4 | -1.27 | -0.69 | Both Isoforms |
| IRF2BPL | -0.80 | -0.45 | AR-FL | HOXB9 | -1.01 | -0.69 | Both Isoforms |
| CKAP2L | -0.83 | -0.45 | AR-FL | POLR3G | -1.05 | -0.69 | Both Isoforms |

| | | | |
|---|---|---|---|
| KCTD17 | -0.73 | -0.45 | AR-FL |
| ZBTB34 | -0.74 | -0.45 | AR-FL |
| TDP1 | -0.59 | -0.45 | AR-FL |
| IL15RA | -0.64 | -0.45 | AR-FL |
| NAIP | -0.58 | -0.45 | AR-FL |
| ZBTB47 | -0.65 | -0.45 | AR-FL |
| CEP131 | -0.65 | -0.45 | AR-FL |
| DBP | -1.09 | -0.44 | AR-FL |
| AURKB | -1.07 | -0.44 | AR-FL |
| GPD1 | -0.83 | -0.44 | AR-FL |
| HFE | -0.95 | -0.44 | AR-FL |
| KALRN | -1.58 | -0.44 | AR-FL |
| BMP1 | -1.17 | -0.44 | AR-FL |
| WDR90 | -0.62 | -0.44 | AR-FL |
| MAP1A | -1.39 | -0.44 | AR-FL |
| NDC80 | -0.76 | -0.44 | AR-FL |
| THAP10 | -0.88 | -0.44 | AR-FL |
| ARHGEF40 | -1.00 | -0.44 | AR-FL |
| SMIM10L2A | -0.95 | -0.44 | AR-FL |
| LRRC20 | -0.75 | -0.44 | AR-FL |
| WDR88 | -0.72 | -0.44 | AR-FL |
| TM7SF2 | -0.74 | -0.44 | AR-FL |
| GLS2 | -1.07 | -0.43 | AR-FL |
| H2AFX | -0.79 | -0.43 | AR-FL |
| TGM3 | -3.69 | -0.43 | AR-FL |
| ELOVL6 | -1.02 | -0.43 | AR-FL |
| HSPA12A | -0.85 | -0.43 | AR-FL |
| 9-Sep | -0.62 | -0.43 | AR-FL |
| MCC | -2.27 | -0.43 | AR-FL |
| ZNF736 | -0.86 | -0.42 | AR-FL |
| SESN3 | -1.21 | -0.42 | AR-FL |
| C16orf59 | -0.77 | -0.42 | AR-FL |
| SLC13A4 | -1.55 | -0.42 | AR-FL |
| TLR5 | -0.98 | -0.42 | AR-FL |
| C21orf58 | -0.72 | -0.42 | AR-FL |
| LGALS3BP | -0.59 | -0.42 | AR-FL |
| NUDT11 | -0.70 | -0.42 | AR-FL |
| NET1 | -0.74 | -0.42 | AR-FL |
| CLYBL | -1.25 | -0.42 | AR-FL |
| RDM1 | -0.94 | -0.42 | AR-FL |
| DMBX1 | -1.62 | -0.42 | AR-FL |
| KITLG | -1.05 | -0.42 | AR-FL |
| SSBP2 | -2.36 | -0.41 | AR-FL |
| TICRR | -0.92 | -0.41 | AR-FL |
| FAM177B | -1.79 | -0.41 | AR-FL |
| ADCY7 | -2.53 | -0.41 | AR-FL |

| | | | |
|---|---|---|---|
| SDC4 | -1.30 | -0.69 | Both Isoforms |
| SP110 | -1.30 | -0.69 | Both Isoforms |
| SOX1 | -1.51 | -0.69 | Both Isoforms |
| TCAIM | -0.66 | -0.69 | Both Isoforms |
| ZNF485 | -0.75 | -0.69 | Both Isoforms |
| ANKRD22 | -1.26 | -0.69 | Both Isoforms |
| ATG9B | -1.38 | -0.69 | Both Isoforms |
| SIK2 | -0.80 | -0.69 | Both Isoforms |
| IKZF2 | -0.95 | -0.69 | Both Isoforms |
| ANKRD50 | -2.19 | -0.69 | Both Isoforms |
| RASAL2 | -1.85 | -0.69 | Both Isoforms |
| TEC | -0.88 | -0.69 | Both Isoforms |
| LMCD1 | -0.61 | -0.69 | Both Isoforms |
| SPINK5 | -1.31 | -0.69 | Both Isoforms |
| DCLK1 | -1.03 | -0.69 | Both Isoforms |
| IL7 | -1.91 | -0.69 | Both Isoforms |
| DHX40 | -1.04 | -0.69 | Both Isoforms |
| ALKBH2 | -1.23 | -0.69 | Both Isoforms |
| ARHGEF10 | -0.72 | -0.68 | Both Isoforms |
| RASD2 | -0.71 | -0.68 | Both Isoforms |
| PPP1R3B | -0.68 | -0.68 | Both Isoforms |
| STX7 | -1.16 | -0.68 | Both Isoforms |
| FXYD5 | -1.47 | -0.68 | Both Isoforms |
| RAB39B | -1.01 | -0.68 | Both Isoforms |
| GNG4 | -1.92 | -0.68 | Both Isoforms |
| METAP2 | -0.69 | -0.68 | Both Isoforms |
| OBSCN | -2.69 | -0.68 | Both Isoforms |
| SPTBN5 | -1.14 | -0.68 | Both Isoforms |
| ERICH5 | -1.30 | -0.68 | Both Isoforms |
| CYP4V2 | -0.99 | -0.68 | Both Isoforms |
| SPRY2 | -1.31 | -0.68 | Both Isoforms |
| EFHC2 | -1.34 | -0.68 | Both Isoforms |
| ITGB1BP2 | -1.22 | -0.68 | Both Isoforms |
| AFF1 | -0.80 | -0.68 | Both Isoforms |
| ANTXR1 | -1.76 | -0.68 | Both Isoforms |
| SLC25A26 | -1.09 | -0.68 | Both Isoforms |
| HBP1 | -0.99 | -0.68 | Both Isoforms |
| GOLIM4 | -0.94 | -0.68 | Both Isoforms |
| PCDHGA1 | -0.78 | -0.68 | Both Isoforms |
| NHLRC4 | -1.89 | -0.68 | Both Isoforms |
| UBASH3B | -1.74 | -0.68 | Both Isoforms |
| DDX60 | -1.06 | -0.68 | Both Isoforms |
| NPM1 | -0.99 | -0.68 | Both Isoforms |
| FMO5 | -1.39 | -0.67 | Both Isoforms |
| EFCAB11 | -1.17 | -0.67 | Both Isoforms |
| TMC8 | -1.02 | -0.67 | Both Isoforms |

| | | | | | | | |
|---|---|---|---|---|---|---|---|
| RP11-723O4.6 | -0.87 | -0.41 | AR-FL | APP | -0.90 | -0.67 | Both Isoforms |
| FRMPD2 | -1.62 | -0.41 | AR-FL | FMN2 | -1.75 | -0.67 | Both Isoforms |
| KIF20B | -0.82 | -0.41 | AR-FL | GPR150 | -1.55 | -0.67 | Both Isoforms |
| HSD3B7 | -0.66 | -0.41 | AR-FL | FAM71F2 | -1.40 | -0.67 | Both Isoforms |
| MFI2 | -0.76 | -0.41 | AR-FL | ZADH2 | -1.07 | -0.67 | Both Isoforms |
| HAUS8 | -0.90 | -0.41 | AR-FL | TATDN3 | -0.87 | -0.67 | Both Isoforms |
| DEPDC1B | -0.59 | -0.41 | AR-FL | CBLN3 | -0.94 | -0.67 | Both Isoforms |
| PSRC1 | -0.73 | -0.41 | AR-FL | RAMP2 | -3.08 | -0.67 | Both Isoforms |
| PVRL4 | -0.59 | -0.41 | AR-FL | FAM216A | -1.40 | -0.67 | Both Isoforms |
| PATZ1 | -0.77 | -0.41 | AR-FL | ZNF581 | -1.48 | -0.67 | Both Isoforms |
| CCDC28B | -0.98 | -0.41 | AR-FL | ZNF277 | -0.91 | -0.67 | Both Isoforms |
| GINS4 | -0.84 | -0.41 | AR-FL | GTPBP8 | -0.68 | -0.67 | Both Isoforms |
| RASSF5 | -1.54 | -0.41 | AR-FL | ZNF354C | -0.84 | -0.67 | Both Isoforms |
| C16orf74 | -0.93 | -0.40 | AR-FL | ZDBF2 | -1.06 | -0.67 | Both Isoforms |
| PTPRB | -1.94 | -0.40 | AR-FL | NEDD9 | -0.63 | -0.67 | Both Isoforms |
| SLC16A2 | -0.73 | -0.40 | AR-FL | BMP2K | -1.13 | -0.67 | Both Isoforms |
| ITGB1BP1 | -0.62 | -0.40 | AR-FL | ARHGEF25 | -1.49 | -0.67 | Both Isoforms |
| TNFAIP2 | -1.43 | -0.40 | AR-FL | CC2D2A | -1.00 | -0.67 | Both Isoforms |
| SLC18B1 | -1.25 | -0.40 | AR-FL | DSP | -0.59 | -0.66 | Both Isoforms |
| DOC2A | -0.94 | -0.39 | AR-FL | BTN2A2 | -1.29 | -0.66 | Both Isoforms |
| DSEL | -1.72 | -0.39 | AR-FL | RORC | -1.78 | -0.66 | Both Isoforms |
| CEP78 | -0.72 | -0.39 | AR-FL | RCBTB2 | -1.05 | -0.66 | Both Isoforms |
| PRIM1 | -1.01 | -0.39 | AR-FL | OGFRL1 | -0.98 | -0.66 | Both Isoforms |
| IRF5 | -0.69 | -0.39 | AR-FL | VWA5B1 | -1.74 | -0.66 | Both Isoforms |
| FHAD1 | -1.05 | -0.39 | AR-FL | SLC25A43 | -0.66 | -0.66 | Both Isoforms |
| VPS37D | -0.87 | -0.39 | AR-FL | COLCA2 | -2.28 | -0.66 | Both Isoforms |
| IL1RAP | -1.07 | -0.39 | AR-FL | MAP7D2 | -1.35 | -0.66 | Both Isoforms |
| WLS | -1.46 | -0.39 | AR-FL | AGPAT4 | -1.80 | -0.66 | Both Isoforms |
| FAM102A | -0.76 | -0.39 | AR-FL | MSRB3 | -1.37 | -0.66 | Both Isoforms |
| SLC20A2 | -0.62 | -0.39 | AR-FL | SHE | -0.98 | -0.66 | Both Isoforms |
| RFXAP | -0.65 | -0.39 | AR-FL | SAMD9 | -1.54 | -0.66 | Both Isoforms |
| TK1 | -0.96 | -0.39 | AR-FL | SFTPA2 | -1.13 | -0.66 | Both Isoforms |
| CMTM1 | -0.66 | -0.39 | AR-FL | GPR176 | -1.19 | -0.66 | Both Isoforms |
| SPC24 | -0.98 | -0.39 | AR-FL | FAM110B | -1.56 | -0.66 | Both Isoforms |
| FANCG | -0.66 | -0.39 | AR-FL | RAB40B | -1.06 | -0.66 | Both Isoforms |
| SYNM | -0.74 | -0.39 | AR-FL | CDNF | -1.76 | -0.66 | Both Isoforms |
| KNTC1 | -0.83 | -0.39 | AR-FL | GAS2 | -1.46 | -0.65 | Both Isoforms |
| SYBU | -1.01 | -0.39 | AR-FL | POLR1D | -1.08 | -0.65 | Both Isoforms |
| RBP5 | -1.27 | -0.39 | AR-FL | ADGRE5 | -0.64 | -0.65 | Both Isoforms |
| GINS1 | -0.89 | -0.38 | AR-FL | STAG2 | -0.87 | -0.65 | Both Isoforms |
| GPR19 | -0.71 | -0.38 | AR-FL | ZC3H6 | -0.97 | -0.65 | Both Isoforms |
| CENPP | -0.65 | -0.38 | AR-FL | RP11-111M22.2 | -1.27 | -0.65 | Both Isoforms |
| SCLT1 | -0.84 | -0.38 | AR-FL | RAB17 | -0.83 | -0.65 | Both Isoforms |
| RAPGEF6 | -0.71 | -0.38 | AR-FL | CMYA5 | -1.67 | -0.65 | Both Isoforms |
| GLTPD2 | -0.67 | -0.38 | AR-FL | ANXA9 | -0.68 | -0.65 | Both Isoforms |
| FAM53B | -0.60 | -0.38 | AR-FL | PLEKHG4 | -0.82 | -0.65 | Both Isoforms |

| | | | | | | | |
|---|---|---|---|---|---|---|---|
| ORC6 | -1.02 | -0.38 | AR-FL | KANK3 | -1.29 | -0.65 | Both Isoforms |
| UPK3A | -1.23 | -0.37 | AR-FL | LETMD1 | -1.31 | -0.65 | Both Isoforms |
| COLQ | -1.33 | -0.37 | AR-FL | SESN1 | -1.30 | -0.65 | Both Isoforms |
| RFC3 | -0.81 | -0.37 | AR-FL | PTPRT | -4.55 | -0.65 | Both Isoforms |
| PLS1 | -0.92 | -0.37 | AR-FL | NKX2-3 | -1.19 | -0.65 | Both Isoforms |
| HSPG2 | -1.22 | -0.37 | AR-FL | OLFM2 | -1.03 | -0.65 | Both Isoforms |
| PBK | -0.66 | -0.37 | AR-FL | FBXL4 | -0.79 | -0.65 | Both Isoforms |
| ENOSF1 | -0.65 | -0.36 | AR-FL | PCNXL4 | -1.11 | -0.65 | Both Isoforms |
| LRRCC1 | -0.59 | -0.36 | AR-FL | SIX2 | -1.39 | -0.65 | Both Isoforms |
| ARRB2 | -0.74 | -0.36 | AR-FL | BEND6 | -1.76 | -0.65 | Both Isoforms |
| ESPL1 | -1.19 | -0.36 | AR-FL | SIAE | -1.09 | -0.65 | Both Isoforms |
| UBE2T | -0.61 | -0.36 | AR-FL | OARD1 | -0.89 | -0.64 | Both Isoforms |
| NPIPA8 | -0.70 | -0.36 | AR-FL | CDKL2 | -2.42 | -0.64 | Both Isoforms |
| RAD51AP1 | -0.74 | -0.36 | AR-FL | PRSS23 | -1.40 | -0.64 | Both Isoforms |
| RAB37 | -0.95 | -0.36 | AR-FL | WASF1 | -0.81 | -0.64 | Both Isoforms |
| TEF | -0.60 | -0.36 | AR-FL | LRRK1 | -0.96 | -0.64 | Both Isoforms |
| BRCA2 | -1.05 | -0.36 | AR-FL | E2F5 | -0.90 | -0.64 | Both Isoforms |
| YPEL1 | -1.03 | -0.35 | AR-FL | C5orf56 | -0.75 | -0.64 | Both Isoforms |
| STARD9 | -0.67 | -0.35 | AR-FL | TPM2 | -3.19 | -0.64 | Both Isoforms |
| ARHGAP11B | -0.65 | -0.35 | AR-FL | LTBP4 | -1.29 | -0.64 | Both Isoforms |
| TAPT1 | -0.77 | -0.35 | AR-FL | MYO15B | -1.12 | -0.64 | Both Isoforms |
| HAUS5 | -0.71 | -0.35 | AR-FL | TGIF1 | -0.68 | -0.64 | Both Isoforms |
| PKIG | -0.85 | -0.35 | AR-FL | SIRPA | -1.52 | -0.64 | Both Isoforms |
| TMEM116 | -0.61 | -0.35 | AR-FL | MRPL10 | -0.82 | -0.64 | Both Isoforms |
| DGAT2 | -1.18 | -0.35 | AR-FL | NPR2 | -0.89 | -0.64 | Both Isoforms |
| TMEM106C | -0.65 | -0.34 | AR-FL | ATXN1 | -0.83 | -0.64 | Both Isoforms |
| CARNS1 | -0.71 | -0.34 | AR-FL | CHGB | -1.60 | -0.64 | Both Isoforms |
| LIF | -1.54 | -0.34 | AR-FL | ANKAR | -0.65 | -0.64 | Both Isoforms |
| GSG2 | -1.01 | -0.34 | AR-FL | ZNF224 | -1.43 | -0.64 | Both Isoforms |
| NUSAP1 | -0.67 | -0.34 | AR-FL | APEX1 | -0.81 | -0.64 | Both Isoforms |
| ATAD5 | -0.88 | -0.34 | AR-FL | ARHGAP24 | -1.18 | -0.63 | Both Isoforms |
| SULT2B1 | -0.97 | -0.34 | AR-FL | AMZ1 | -0.87 | -0.63 | Both Isoforms |
| FEN1 | -0.59 | -0.34 | AR-FL | NPTN | -0.67 | -0.63 | Both Isoforms |
| OXTR | -0.85 | -0.34 | AR-FL | EXOSC5 | -1.03 | -0.63 | Both Isoforms |
| TSPAN11 | -1.42 | -0.34 | AR-FL | FAM92A1 | -1.19 | -0.63 | Both Isoforms |
| CASC5 | -0.83 | -0.34 | AR-FL | OPRK1 | -4.69 | -0.63 | Both Isoforms |
| KCNS3 | -0.89 | -0.33 | AR-FL | RAD9B | -1.26 | -0.63 | Both Isoforms |
| VDR | -0.70 | -0.33 | AR-FL | PGM2L1 | -1.29 | -0.63 | Both Isoforms |
| MAN1C1 | -1.13 | -0.33 | AR-FL | CAV2 | -1.84 | -0.63 | Both Isoforms |
| DNMT3B | -0.59 | -0.33 | AR-FL | RBM43 | -0.95 | -0.63 | Both Isoforms |
| MNS1 | -0.86 | -0.33 | AR-FL | CALN1 | -3.20 | -0.63 | Both Isoforms |
| NRGN | -0.88 | -0.33 | AR-FL | CA1 | -1.48 | -0.63 | Both Isoforms |
| AUNIP | -1.38 | -0.33 | AR-FL | SCG5 | -2.95 | -0.63 | Both Isoforms |
| POLN | -0.71 | -0.33 | AR-FL | ATAD3C | -1.63 | -0.63 | Both Isoforms |
| DNA2 | -0.87 | -0.33 | AR-FL | OSBPL8 | -1.07 | -0.63 | Both Isoforms |
| FAM19A2 | -1.46 | -0.33 | AR-FL | TFAP4 | -1.28 | -0.63 | Both Isoforms |

| | | | |
|---|---|---|---|
| SCAPER | -0.65 | -0.32 | AR-FL |
| UGT2B15 | -3.34 | -0.32 | AR-FL |
| CCNF | -0.70 | -0.32 | AR-FL |
| MAPRE2 | -1.08 | -0.32 | AR-FL |
| SFI1 | -0.69 | -0.32 | AR-FL |
| C1R | -1.23 | -0.32 | AR-FL |
| GAREM | -0.85 | -0.32 | AR-FL |
| NBPF26 | -1.81 | -0.32 | AR-FL |
| CLMN | -0.64 | -0.32 | AR-FL |
| QDPR | -0.60 | -0.32 | AR-FL |
| GFOD1 | -0.80 | -0.32 | AR-FL |
| PIF1 | -0.80 | -0.32 | AR-FL |
| CDH26 | -2.25 | -0.32 | AR-FL |
| RNF217 | -1.28 | -0.32 | AR-FL |
| STXBP5L | -2.52 | -0.32 | AR-FL |
| GRAMD4 | -0.92 | -0.32 | AR-FL |
| RFC5 | -0.78 | -0.32 | AR-FL |
| PRMT9 | -0.61 | -0.32 | AR-FL |
| PROM2 | -0.78 | -0.32 | AR-FL |
| RARG | -0.63 | -0.31 | AR-FL |
| SH3BP1 | -0.82 | -0.31 | AR-FL |
| DIS3L | -0.82 | -0.31 | AR-FL |
| PSMC3IP | -0.69 | -0.31 | AR-FL |
| SGOL1 | -0.62 | -0.31 | AR-FL |
| CEP68 | -0.89 | -0.31 | AR-FL |
| ANP32E | -0.90 | -0.31 | AR-FL |
| RMI2 | -1.02 | -0.31 | AR-FL |
| ELFN2 | -1.59 | -0.31 | AR-FL |
| GFPT2 | -1.06 | -0.30 | AR-FL |
| NPAS1 | -0.75 | -0.30 | AR-FL |
| TOX2 | -1.08 | -0.30 | AR-FL |
| CDCA4 | -0.84 | -0.30 | AR-FL |
| RNF128 | -0.99 | -0.30 | AR-FL |
| NCAPG | -0.67 | -0.30 | AR-FL |
| RELT | -1.32 | -0.30 | AR-FL |
| CENPF | -0.63 | -0.30 | AR-FL |
| FAM198B | -2.85 | -0.30 | AR-FL |
| CEP128 | -0.69 | -0.29 | AR-FL |
| EGFL7 | -0.70 | -0.29 | AR-FL |
| THOC3 | -0.64 | -0.29 | AR-FL |
| DNM3 | -1.48 | -0.29 | AR-FL |
| CENPK | -0.71 | -0.29 | AR-FL |
| RAB36 | -0.74 | -0.29 | AR-FL |
| RRM2 | -0.70 | -0.29 | AR-FL |
| PHLDA3 | -0.76 | -0.28 | AR-FL |
| LRRC37B | -0.65 | -0.28 | AR-FL |
| EFNB3 | -1.12 | -0.63 | Both Isoforms |
| PLEKHH1 | -0.63 | -0.62 | Both Isoforms |
| FDXR | -0.79 | -0.62 | Both Isoforms |
| CACNA1B | -1.42 | -0.62 | Both Isoforms |
| ZNF286A | -0.90 | -0.62 | Both Isoforms |
| ABHD1 | -1.35 | -0.62 | Both Isoforms |
| DIEXF | -1.08 | -0.62 | Both Isoforms |
| BZW2 | -0.91 | -0.62 | Both Isoforms |
| CAMKMT | -0.94 | -0.62 | Both Isoforms |
| ATG14 | -1.01 | -0.62 | Both Isoforms |
| POLR1E | -1.40 | -0.62 | Both Isoforms |
| SLC6A9 | -1.24 | -0.62 | Both Isoforms |
| ING3 | -0.95 | -0.62 | Both Isoforms |
| PSD | -1.97 | -0.62 | Both Isoforms |
| KCNH1 | -1.38 | -0.62 | Both Isoforms |
| ANKRD63 | -2.36 | -0.62 | Both Isoforms |
| WFIKKN1 | -1.26 | -0.62 | Both Isoforms |
| CTSF | -1.69 | -0.62 | Both Isoforms |
| TAF1D | -1.25 | -0.62 | Both Isoforms |
| SLC16A10 | -0.60 | -0.62 | Both Isoforms |
| RARRES3 | -0.77 | -0.62 | Both Isoforms |
| FAM117B | -0.89 | -0.62 | Both Isoforms |
| CCDC112 | -0.73 | -0.62 | Both Isoforms |
| XK | -0.89 | -0.62 | Both Isoforms |
| POLL | -1.14 | -0.62 | Both Isoforms |
| WDR35 | -1.16 | -0.62 | Both Isoforms |
| FSD1 | -2.50 | -0.62 | Both Isoforms |
| CROCC | -1.31 | -0.62 | Both Isoforms |
| HOXA13 | -1.64 | -0.61 | Both Isoforms |
| ANKRD65 | -1.02 | -0.61 | Both Isoforms |
| DDX60L | -0.65 | -0.61 | Both Isoforms |
| SEPSECS | -1.12 | -0.61 | Both Isoforms |
| MAPK15 | -2.12 | -0.61 | Both Isoforms |
| ATG16L2 | -1.81 | -0.61 | Both Isoforms |
| NMB | -1.00 | -0.61 | Both Isoforms |
| HDAC5 | -0.90 | -0.61 | Both Isoforms |
| ADCK3 | -0.71 | -0.61 | Both Isoforms |
| HIST1H2AC | -0.91 | -0.61 | Both Isoforms |
| KIAA0040 | -0.85 | -0.61 | Both Isoforms |
| FOXO4 | -0.95 | -0.61 | Both Isoforms |
| ZNF763 | -1.02 | -0.61 | Both Isoforms |
| C1orf106 | -0.70 | -0.61 | Both Isoforms |
| PIM1 | -0.71 | -0.61 | Both Isoforms |
| ZNF239 | -1.18 | -0.61 | Both Isoforms |
| CDKL1 | -0.97 | -0.61 | Both Isoforms |
| ADAM11 | -1.78 | -0.61 | Both Isoforms |

| | | | |
|---|---|---|---|
| CDR2L | -0.84 | -0.28 | AR-FL |
| STXBP1 | -0.75 | -0.28 | AR-FL |
| GLMN | -0.77 | -0.28 | AR-FL |
| PTPRU | -0.59 | -0.28 | AR-FL |
| RAB19 | -0.67 | -0.27 | AR-FL |
| ATP4B | -0.99 | -0.27 | AR-FL |
| BNIPL | -0.77 | -0.27 | AR-FL |
| CHEK1 | -0.77 | -0.27 | AR-FL |
| RPAP2 | -0.62 | -0.27 | AR-FL |
| TMEM266 | -0.62 | -0.27 | AR-FL |
| WDHD1 | -0.77 | -0.27 | AR-FL |
| STAG3 | -0.69 | -0.27 | AR-FL |
| TMC8 | -0.64 | -0.27 | AR-FL |
| ZNF596 | -0.69 | -0.27 | AR-FL |
| HS6ST2 | -2.51 | -0.27 | AR-FL |
| BOC | -1.00 | -0.26 | AR-FL |
| ULBP1 | -0.98 | -0.26 | AR-FL |
| TRIM45 | -1.15 | -0.26 | AR-FL |
| NDUFB4 | -0.96 | -0.26 | AR-FL |
| PLIN4 | -1.56 | -0.26 | AR-FL |
| TLN2 | -0.91 | -0.26 | AR-FL |
| TRIP13 | -0.60 | -0.26 | AR-FL |
| CIT | -0.74 | -0.26 | AR-FL |
| CISH | -1.17 | -0.26 | AR-FL |
| CCNG2 | -0.90 | -0.26 | AR-FL |
| IL17RB | -0.73 | -0.25 | AR-FL |
| STK32A | -0.76 | -0.25 | AR-FL |
| EME1 | -0.99 | -0.25 | AR-FL |
| NEK1 | -0.69 | -0.25 | AR-FL |
| CLIP3 | -0.90 | -0.25 | AR-FL |
| ARHGEF39 | -0.79 | -0.25 | AR-FL |
| SCNN1D | -0.74 | -0.25 | AR-FL |
| NFATC2 | -1.37 | -0.25 | AR-FL |
| DZIP1L | -1.86 | -0.24 | AR-FL |
| CAMKMT | -0.65 | -0.24 | AR-FL |
| FDXR | -0.98 | -0.24 | AR-FL |
| ANKRD36C | -0.65 | -0.24 | AR-FL |
| PALM | -0.99 | -0.24 | AR-FL |
| TOP2A | -1.01 | -0.24 | AR-FL |
| RFC4 | -0.70 | -0.24 | AR-FL |
| NUF2 | -0.75 | -0.23 | AR-FL |
| DONSON | -0.59 | -0.23 | AR-FL |
| SNAI1 | -1.36 | -0.23 | AR-FL |
| ELF3 | -0.65 | -0.23 | AR-FL |
| PTP4A3 | -0.59 | -0.23 | AR-FL |
| EIF4E3 | -1.22 | -0.23 | AR-FL |
| CARF | -1.36 | -0.61 | Both Isoforms |
| AVPR1B | -0.99 | -0.61 | Both Isoforms |
| GALNT14 | -1.82 | -0.61 | Both Isoforms |
| N4BP3 | -1.85 | -0.60 | Both Isoforms |
| BCL2 | -0.63 | -0.60 | Both Isoforms |
| COL4A4 | -2.57 | -0.60 | Both Isoforms |
| DIAPH2 | -0.89 | -0.60 | Both Isoforms |
| CCNG1 | -0.64 | -0.60 | Both Isoforms |
| KNG1 | -2.16 | -0.60 | Both Isoforms |
| BMP7 | -1.54 | -0.60 | Both Isoforms |
| NFATC1 | -0.79 | -0.60 | Both Isoforms |
| SRI | -0.68 | -0.60 | Both Isoforms |
| RFESD | -0.82 | -0.60 | Both Isoforms |
| RPUSD4 | -0.96 | -0.60 | Both Isoforms |
| AMER3 | -0.71 | -0.60 | Both Isoforms |
| STK26 | -0.70 | -0.60 | Both Isoforms |
| DMBX1 | -1.37 | -0.60 | Both Isoforms |
| LMBR1L | -1.28 | -0.60 | Both Isoforms |
| C8orf34 | -1.41 | -0.60 | Both Isoforms |
| HOXC9 | -1.23 | -0.60 | Both Isoforms |
| GCSAM | -1.47 | -0.60 | Both Isoforms |
| MAP3K5 | -1.08 | -0.60 | Both Isoforms |
| DLX6 | -1.01 | -0.60 | Both Isoforms |
| RAB11FIP4 | -0.60 | -0.60 | Both Isoforms |
| MTIF2 | -0.63 | -0.60 | Both Isoforms |
| AGTPBP1 | -0.62 | -0.60 | Both Isoforms |
| CD302 | -1.59 | -0.60 | Both Isoforms |
| CLHC1 | -0.77 | -0.60 | Both Isoforms |
| TNFRSF18 | -1.68 | -0.60 | Both Isoforms |
| RNF144B | -0.82 | -0.60 | Both Isoforms |
| ACAA2 | -1.22 | -0.60 | Both Isoforms |
| MAP7D3 | -1.51 | -0.59 | Both Isoforms |
| TSEN2 | -0.84 | -0.59 | Both Isoforms |
| TRAK1 | -0.79 | -0.59 | Both Isoforms |
| SIRT3 | -0.75 | -0.59 | Both Isoforms |
| SLC7A11 | -1.22 | -0.59 | Both Isoforms |
| COL4A3 | -1.88 | -0.59 | Both Isoforms |
| TRIP6 | -1.00 | -0.59 | Both Isoforms |
| LHX6 | -1.51 | -0.59 | Both Isoforms |
| SLC25A24 | -0.60 | -0.59 | Both Isoforms |
| FHL3 | -1.02 | -0.59 | Both Isoforms |
| LARP6 | -0.62 | -0.59 | Both Isoforms |
| CRBN | -0.86 | -0.59 | Both Isoforms |
| CCNDBP1 | -0.76 | -0.59 | Both Isoforms |
| NPM2 | -1.41 | -0.59 | Both Isoforms |
| TRIM59 | -0.77 | -0.59 | Both Isoforms |

| | | | |
|---|---|---|---|
| FUS | -0.60 | -0.23 | AR-FL |
| ZGRF1 | -0.92 | -0.23 | AR-FL |
| AGPAT2 | -0.60 | -0.23 | AR-FL |
| PLK4 | -0.99 | -0.23 | AR-FL |
| FGFR1 | -0.62 | -0.23 | AR-FL |
| KIF23 | -0.82 | -0.23 | AR-FL |
| MNX1 | -0.93 | -0.23 | AR-FL |
| ZC3H12B | -0.86 | -0.23 | AR-FL |
| ARL6IP6 | -0.67 | -0.22 | AR-FL |
| CCDC150 | -0.65 | -0.22 | AR-FL |
| LRRC75A | -1.44 | -0.22 | AR-FL |
| ELN | -0.94 | -0.22 | AR-FL |
| KIF14 | -0.69 | -0.22 | AR-FL |
| TCTEX1D2 | -0.82 | -0.22 | AR-FL |
| SPATA33 | -0.63 | -0.22 | AR-FL |
| CEP72 | -0.79 | -0.21 | AR-FL |
| TMPO | -0.58 | -0.21 | AR-FL |
| ZNF724P | -0.80 | -0.21 | AR-FL |
| PEG3 | -2.46 | -0.21 | AR-FL |
| TMOD2 | -0.99 | -0.21 | AR-FL |
| WNK2 | -1.21 | -0.21 | AR-FL |
| TRAIP | -0.81 | -0.21 | AR-FL |
| NT5DC2 | -0.65 | -0.21 | AR-FL |
| NOTCH3 | -1.08 | -0.20 | AR-FL |
| VMO1 | -0.92 | -0.20 | AR-FL |
| NBEA | -0.66 | -0.20 | AR-FL |
| C9orf43 | -0.59 | -0.20 | AR-FL |
| UBE2C | -0.67 | -0.20 | AR-FL |
| KIF2C | -0.89 | -0.20 | AR-FL |
| FAM86B2 | -1.47 | -0.20 | AR-FL |
| ADRA2C | -1.42 | -0.20 | AR-FL |
| PAX1 | -2.13 | -0.20 | AR-FL |
| FES | -1.45 | -0.19 | AR-FL |
| DPF3 | -0.91 | -0.19 | AR-FL |
| ATG16L2 | -1.34 | -0.19 | AR-FL |
| GNG13 | -0.68 | -0.19 | AR-FL |
| BAMBI | -1.77 | -0.19 | AR-FL |
| MXRA8 | -0.62 | -0.19 | AR-FL |
| SMO | -0.67 | -0.19 | AR-FL |
| CDH24 | -1.29 | -0.19 | AR-FL |
| ZPLD1 | -0.65 | -0.19 | AR-FL |
| IER5 | -1.12 | -0.19 | AR-FL |
| OR2B6 | -0.84 | -0.18 | AR-FL |
| MXD3 | -0.68 | -0.18 | AR-FL |
| TARBP1 | -0.76 | -0.18 | AR-FL |
| PITX1 | -0.92 | -0.18 | AR-FL |

| | | | |
|---|---|---|---|
| F8 | -1.48 | -0.59 | Both Isoforms |
| DCBLD2 | -1.52 | -0.59 | Both Isoforms |
| OGDHL | -1.21 | -0.59 | Both Isoforms |
| NINL | -0.87 | -0.59 | Both Isoforms |
| GPC6 | -1.92 | -0.59 | Both Isoforms |
| ABHD18 | -0.74 | -0.59 | Both Isoforms |
| NPAS1 | -2.26 | -0.59 | Both Isoforms |
| HS6ST2 | -1.49 | -0.59 | Both Isoforms |
| TSNARE1 | -1.43 | -0.58 | Both Isoforms |
| MYO5B | -0.61 | -0.58 | Both Isoforms |
| OTX1 | -0.74 | -0.58 | Both Isoforms |
| ZKSCAN7 | -1.29 | -0.58 | Both Isoforms |
| KIAA1328 | -0.99 | -0.58 | Both Isoforms |
| SUMF2 | -0.94 | -0.58 | Both Isoforms |
| MOB3C | -0.80 | -0.58 | Both Isoforms |
| ZC2HC1A | -0.83 | -0.58 | Both Isoforms |
| RBBP9 | -0.99 | -0.58 | AR-FL |
| FAM60A | -0.69 | -0.58 | AR-FL |
| CCDC102A | -1.19 | -0.58 | AR-FL |
| CBLB | -0.60 | -0.58 | AR-FL |
| SMAD7 | -0.83 | -0.58 | AR-FL |
| ATM | -1.10 | -0.58 | AR-FL |
| IRS2 | -1.53 | -0.58 | AR-FL |
| ARG2 | -1.13 | -0.58 | AR-FL |
| TSPAN7 | -1.09 | -0.58 | AR-FL |
| HDAC9 | -1.27 | -0.58 | AR-FL |
| CCDC77 | -0.80 | -0.58 | AR-FL |
| ANKRD6 | -0.79 | -0.58 | AR-FL |
| PDPR | -0.91 | -0.58 | AR-FL |
| FGD3 | -0.98 | -0.58 | AR-FL |
| CHRNB1 | -0.87 | -0.58 | AR-FL |
| TPPP | -1.35 | -0.58 | AR-FL |
| PRR16 | -2.68 | -0.58 | AR-FL |
| RSL24D1 | -0.85 | -0.57 | AR-FL |
| VSTM5 | -0.87 | -0.57 | AR-FL |
| PLTP | -1.43 | -0.57 | AR-FL |
| RPAP2 | -0.68 | -0.57 | AR-FL |
| ADAT2 | -1.40 | -0.57 | AR-FL |
| GKAP1 | -0.64 | -0.57 | AR-FL |
| TSPAN2 | -2.09 | -0.57 | AR-FL |
| ZNF251 | -1.05 | -0.57 | AR-FL |
| FAS | -1.23 | -0.57 | AR-FL |
| NOS3 | -1.00 | -0.57 | AR-FL |
| RNF43 | -0.83 | -0.57 | AR-FL |
| ZFP14 | -0.94 | -0.57 | AR-FL |
| HIST3H2A | -0.93 | -0.57 | AR-FL |

| | | | | | | | |
|---|---|---|---|---|---|---|---|
| AK7 | -0.66 | -0.18 | AR-FL | SERP2 | -1.68 | -0.57 | AR-FL |
| DPF1 | -0.85 | -0.18 | AR-FL | ZFP2 | -1.04 | -0.57 | AR-FL |
| DNM1 | -0.63 | -0.18 | AR-FL | EPS8 | -1.01 | -0.57 | AR-FL |
| EXTL1 | -2.21 | -0.18 | AR-FL | PPP1R9A | -0.99 | -0.57 | AR-FL |
| RPS17 | -1.35 | -0.18 | AR-FL | C2orf70 | -1.72 | -0.57 | AR-FL |
| POU2F3 | -0.78 | -0.18 | AR-FL | TMTC1 | -1.84 | -0.57 | AR-FL |
| MGME1 | -0.62 | -0.17 | AR-FL | EDA | -2.28 | -0.57 | AR-FL |
| SPG20 | -1.67 | -0.17 | AR-FL | SEMA6B | -1.75 | -0.56 | AR-FL |
| KIAA0922 | -0.59 | -0.17 | AR-FL | BTN3A1 | -1.28 | -0.56 | AR-FL |
| HMGCS2 | -0.62 | -0.17 | AR-FL | BCAS4 | -1.59 | -0.56 | AR-FL |
| FSCN1 | -1.13 | -0.17 | AR-FL | SOX13 | -1.90 | -0.56 | AR-FL |
| KLLN | -0.89 | -0.17 | AR-FL | DDC | -2.01 | -0.56 | AR-FL |
| NABP1 | -0.61 | -0.17 | AR-FL | C14orf159 | -1.10 | -0.56 | AR-FL |
| FBXO44 | -0.83 | -0.17 | AR-FL | MRPL45 | -0.60 | -0.56 | AR-FL |
| NAP1L1 | -0.80 | -0.17 | AR-FL | ZNF343 | -1.05 | -0.56 | AR-FL |
| FCGBP | -0.76 | -0.17 | AR-FL | KIAA1147 | -0.86 | -0.56 | AR-FL |
| MAN2A2 | -0.73 | -0.17 | AR-FL | IMPACT | -0.76 | -0.56 | AR-FL |
| ATP8A1 | -0.93 | -0.17 | AR-FL | ZDHHC17 | -0.59 | -0.56 | AR-FL |
| ATP6AP1L | -0.88 | -0.16 | AR-FL | SPATA7 | -0.95 | -0.56 | AR-FL |
| TNIK | -0.80 | -0.16 | AR-FL | TIMP2 | -1.13 | -0.56 | AR-FL |
| MMP11 | -0.66 | -0.16 | AR-FL | LANCL1 | -0.82 | -0.56 | AR-FL |
| INPP5B | -0.62 | -0.16 | AR-FL | KCNK5 | -0.94 | -0.56 | AR-FL |
| ARHGAP19 | -0.60 | -0.16 | AR-FL | TTC36 | -0.98 | -0.56 | AR-FL |
| KIF3C | -0.70 | -0.16 | AR-FL | BACH1 | -0.73 | -0.56 | AR-FL |
| CDCA3 | -0.73 | -0.16 | AR-FL | PSD4 | -1.03 | -0.56 | AR-FL |
| CENPE | -0.68 | -0.15 | AR-FL | RIMBP2 | -1.24 | -0.56 | AR-FL |
| CDCA8 | -0.64 | -0.15 | AR-FL | TRIM23 | -0.85 | -0.56 | AR-FL |
| NES | -1.03 | -0.15 | AR-FL | HTRA2 | -0.62 | -0.56 | AR-FL |
| CCNB2 | -0.70 | -0.15 | AR-FL | ZNF516 | -0.90 | -0.56 | AR-FL |
| ROBO1 | -0.95 | -0.15 | AR-FL | TMEM116 | -0.89 | -0.56 | AR-FL |
| ZNF703 | -0.82 | -0.15 | AR-FL | PRCP | -0.98 | -0.56 | AR-FL |
| FAM57B | -0.65 | -0.15 | AR-FL | RGAG4 | -0.78 | -0.56 | AR-FL |
| C2orf81 | -0.86 | -0.15 | AR-FL | AVPI1 | -0.83 | -0.56 | AR-FL |
| DIAPH3 | -0.79 | -0.14 | AR-FL | MATR3 | -0.90 | -0.56 | AR-FL |
| SCAI | -0.59 | -0.14 | AR-FL | MATR3 | -0.67 | -0.56 | AR-FL |
| MAGI3 | -0.83 | -0.14 | AR-FL | GSAP | -0.97 | -0.56 | AR-FL |
| ACACB | -0.82 | -0.14 | AR-FL | ZMAT1 | -2.47 | -0.56 | AR-FL |
| PSIP1 | -0.58 | -0.14 | AR-FL | BCL2L13 | -0.81 | -0.56 | AR-FL |
| RASL10B | -2.33 | -0.14 | AR-FL | EPHA8 | -1.83 | -0.56 | AR-FL |
| CLMP | -0.90 | -0.14 | AR-FL | CTTNBP2NL | -1.01 | -0.56 | AR-FL |
| ALDH1L2 | -0.96 | -0.13 | AR-FL | SNX5 | -0.95 | -0.56 | AR-FL |
| FAM102B | -1.16 | -0.13 | AR-FL | ARHGAP29 | -0.67 | -0.56 | AR-FL |
| ALDH5A1 | -0.74 | -0.13 | AR-FL | CD101 | -0.93 | -0.55 | AR-FL |
| SIPA1L1 | -0.69 | -0.13 | AR-FL | PCYOX1 | -0.74 | -0.55 | AR-FL |
| SPTBN1 | -0.76 | -0.13 | AR-FL | APPL1 | -0.79 | -0.55 | AR-FL |
| C19orf57 | -0.80 | -0.13 | AR-FL | ELF5 | -0.92 | -0.55 | AR-FL |

| | | | | | | | |
|---|---|---|---|---|---|---|---|
| CSAD | -0.65 | -0.13 | AR-FL | ANO9 | -0.68 | -0.55 | AR-FL |
| PDIK1L | -0.61 | -0.13 | AR-FL | RP11-1212A22.4 | -0.66 | -0.55 | AR-FL |
| BMP6 | -0.97 | -0.12 | AR-FL | OPRL1 | -0.92 | -0.55 | AR-FL |
| AMN1 | -0.63 | -0.12 | AR-FL | CATSPERG | -0.82 | -0.55 | AR-FL |
| PRSS23 | -0.69 | -0.12 | AR-FL | ZNF546 | -1.18 | -0.55 | AR-FL |
| CENPI | -0.61 | -0.12 | AR-FL | JRKL | -1.11 | -0.55 | AR-FL |
| RCOR2 | -1.30 | -0.12 | AR-FL | SLC25A36 | -0.81 | -0.55 | AR-FL |
| FAM101B | -1.12 | -0.11 | AR-FL | GPR135 | -0.99 | -0.55 | AR-FL |
| CCDC180 | -1.35 | -0.11 | AR-FL | GHR | -1.58 | -0.55 | AR-FL |
| NINJ1 | -0.97 | -0.11 | AR-FL | SNED1 | -0.64 | -0.55 | AR-FL |
| ENPP1 | -0.66 | -0.11 | AR-FL | ATP2C1 | -0.69 | -0.55 | AR-FL |
| PLCE1 | -0.68 | -0.11 | AR-FL | OAF | -1.13 | -0.55 | AR-FL |
| C3orf52 | -0.66 | -0.10 | AR-FL | ZNF25 | -1.18 | -0.55 | AR-FL |
| MTFR2 | -0.67 | -0.10 | AR-FL | TBC1D1 | -0.89 | -0.55 | AR-FL |
| ZNF738 | -0.92 | -0.10 | AR-FL | C3orf17 | -1.08 | -0.55 | AR-FL |
| NEIL1 | -0.96 | -0.10 | AR-FL | CD6 | -2.37 | -0.55 | AR-FL |
| NAV1 | -0.71 | -0.10 | AR-FL | ADAM7 | -2.52 | -0.55 | AR-FL |
| RASAL2 | -0.60 | -0.10 | AR-FL | PLEKHG1 | -1.74 | -0.55 | AR-FL |
| MXI1 | -0.66 | -0.10 | AR-FL | PTCHD2 | -0.67 | -0.55 | AR-FL |
| GDPD5 | -1.07 | -0.10 | AR-FL | ZNF596 | -0.68 | -0.55 | AR-FL |
| DBF4B | -0.68 | -0.09 | AR-FL | ZBTB41 | -0.87 | -0.55 | AR-FL |
| KLHL3 | -0.75 | -0.09 | AR-FL | RSBN1 | -0.65 | -0.54 | AR-FL |
| EFCC1 | -1.14 | -0.09 | AR-FL | METAP1D | -0.62 | -0.54 | AR-FL |
| POLD3 | -0.77 | -0.09 | AR-FL | ZFYVE9 | -0.71 | -0.54 | AR-FL |
| TNFRSF10D | -0.77 | -0.08 | AR-FL | HEG1 | -2.17 | -0.54 | AR-FL |
| PELI2 | -0.83 | -0.08 | AR-FL | C15orf41 | -0.64 | -0.54 | AR-FL |
| EML5 | -1.10 | -0.08 | AR-FL | CBFA2T2 | -0.58 | -0.54 | AR-FL |
| PCOLCE | -1.03 | -0.08 | AR-FL | KCNIP4 | -1.32 | -0.54 | AR-FL |
| TCEA3 | -0.64 | -0.08 | AR-FL | MED14 | -0.61 | -0.54 | AR-FL |
| MYOM2 | -0.66 | -0.08 | AR-FL | PLEKHN1 | -0.88 | -0.54 | AR-FL |
| TTYH2 | -0.88 | -0.08 | AR-FL | IGDCC4 | -1.53 | -0.54 | AR-FL |
| LRRC49 | -0.78 | -0.07 | AR-FL | ITGB4 | -0.83 | -0.54 | AR-FL |
| RTTN | -0.93 | -0.07 | AR-FL | C14orf93 | -1.61 | -0.54 | AR-FL |
| PARD6A | -0.70 | -0.07 | AR-FL | AKR1E2 | -0.93 | -0.54 | AR-FL |
| OSBPL3 | -0.62 | -0.07 | AR-FL | TMEM62 | -1.02 | -0.54 | AR-FL |
| CLUL1 | -1.12 | -0.06 | AR-FL | ING4 | -0.71 | -0.54 | AR-FL |
| C14orf93 | -0.63 | -0.06 | AR-FL | LMO4 | -1.34 | -0.54 | AR-FL |
| CBX2 | -0.63 | -0.06 | AR-FL | TOP1MT | -1.18 | -0.54 | AR-FL |
| KIAA0895 | -0.66 | -0.06 | AR-FL | MYH10 | -0.85 | -0.54 | AR-FL |
| SORBS3 | -0.59 | -0.06 | AR-FL | ZNF283 | -0.76 | -0.54 | AR-FL |
| SLC2A11 | -0.58 | -0.06 | AR-FL | ATP6AP1L | -1.45 | -0.54 | AR-FL |
| GOLGA8A | -0.59 | -0.06 | AR-FL | SSBP2 | -1.44 | -0.53 | AR-FL |
| PROCR | -0.93 | -0.06 | AR-FL | METTL8 | -0.81 | -0.53 | AR-FL |
| SYT15 | -0.77 | -0.05 | AR-FL | EIF3E | -0.98 | -0.53 | AR-FL |
| THUMPD2 | -0.59 | -0.05 | AR-FL | NLE1 | -1.02 | -0.53 | AR-FL |

| | | | | | | | |
|---|---|---|---|---|---|---|---|
| IRX5 | -1.53 | -0.05 | AR-FL | WDR59 | -1.04 | -0.53 | AR-FL |
| RAB6B | -0.91 | -0.05 | AR-FL | LRRN2 | -3.20 | -0.53 | AR-FL |
| C2orf15 | -0.64 | -0.05 | AR-FL | UBE3D | -0.96 | -0.53 | AR-FL |
| SUGCT | -0.76 | -0.05 | AR-FL | L1CAM | -1.38 | -0.53 | AR-FL |
| C16orf96 | -0.90 | -0.05 | AR-FL | LRPPRC | -0.88 | -0.53 | AR-FL |
| TSPAN3 | -0.89 | -0.04 | AR-FL | THADA | -0.78 | -0.53 | AR-FL |
| DET1 | -0.59 | -0.04 | AR-FL | KCTD18 | -0.62 | -0.53 | AR-FL |
| SORBS1 | -0.82 | -0.04 | AR-FL | PALM2 | -1.75 | -0.53 | AR-FL |
| TMEM65 | -0.59 | -0.04 | AR-FL | TEKT4 | -1.84 | -0.53 | AR-FL |
| FBXO15 | -0.85 | -0.04 | AR-FL | ASS1 | -1.09 | -0.53 | AR-FL |
| PLEKHD1 | -0.88 | -0.04 | AR-FL | PLEKHB1 | -1.29 | -0.53 | AR-FL |
| ZCWPW1 | -0.62 | -0.04 | AR-FL | TCEAL1 | -0.68 | -0.53 | AR-FL |
| CEP126 | -0.62 | -0.04 | AR-FL | C12orf66 | -1.05 | -0.53 | AR-FL |
| KDM4D | -0.62 | -0.03 | AR-FL | SCARA3 | -0.86 | -0.53 | AR-FL |
| FHL3 | -0.61 | -0.03 | AR-FL | GCA | -1.01 | -0.53 | AR-FL |
| BRIP1 | -0.99 | -0.03 | AR-FL | NR6A1 | -1.15 | -0.53 | AR-FL |
| HJURP | -0.74 | -0.02 | AR-FL | PYGM | -1.51 | -0.53 | AR-FL |
| KCNIP3 | -1.04 | -0.02 | AR-FL | USP54 | -1.13 | -0.53 | AR-FL |
| HIST1H2BO | -0.58 | -0.02 | AR-FL | NPHP3 | -0.68 | -0.53 | AR-FL |
| TCFL5 | -0.63 | -0.02 | AR-FL | MLANA | -0.59 | -0.53 | AR-FL |
| KIF24 | -0.76 | -0.02 | AR-FL | EIF3D | -0.73 | -0.53 | AR-FL |
| SSTR2 | -0.63 | -0.02 | AR-FL | SOX18 | -1.51 | -0.52 | AR-FL |
| FAM78A | -0.84 | -0.02 | AR-FL | UTP14A | -0.87 | -0.52 | AR-FL |
| LTBP1 | -1.39 | -0.02 | AR-FL | ZFP36L2 | -1.35 | -0.52 | AR-FL |
| ASPM | -0.62 | -0.02 | AR-FL | MCMDC2 | -1.47 | -0.52 | AR-FL |
| CDK1 | -0.83 | -0.02 | AR-FL | SOCS7 | -0.59 | -0.52 | AR-FL |
| QSER1 | -1.01 | -0.02 | AR-FL | FKTN | -0.83 | -0.52 | AR-FL |
| RASL11B | -2.21 | -0.01 | AR-FL | AMHR2 | -1.66 | -0.52 | AR-FL |
| CYP2W1 | -1.14 | -0.01 | AR-FL | SLC25A34 | -1.16 | -0.52 | AR-FL |
| SUV420H2 | -0.63 | -0.01 | AR-FL | EIF4A2 | -0.69 | -0.52 | AR-FL |
| GALK1 | -0.94 | -0.01 | AR-FL | UBXN7 | -0.88 | -0.52 | AR-FL |
| C5orf45 | -0.60 | -0.01 | AR-FL | BBS7 | -0.93 | -0.52 | AR-FL |
| TEX22 | -0.59 | -0.01 | AR-FL | PLCH2 | -1.54 | -0.52 | AR-FL |
| VASH2 | -0.65 | -0.01 | AR-FL | NRXN2 | -1.37 | -0.52 | AR-FL |
| SMAD3 | -1.28 | 0.00 | AR-FL | ANXA2R | -1.10 | -0.52 | AR-FL |
| TXNIP | -0.93 | 0.00 | AR-FL | BLOC1S6 | -0.62 | -0.52 | AR-FL |
| SSBP3 | -0.86 | 0.00 | AR-FL | SNX15 | -1.00 | -0.52 | AR-FL |
| OPN3 | -0.58 | 0.00 | AR-FL | TMEM117 | -1.13 | -0.52 | AR-FL |
| SPAG16 | -0.33 | -11.04 | AR-V7 | PPOX | -1.06 | -0.52 | AR-FL |
| HENMT1 | -0.22 | -6.26 | AR-V7 | C5orf47 | -0.76 | -0.52 | AR-FL |
| TMSB4X | -0.53 | -4.04 | AR-V7 | ZP3 | -0.94 | -0.52 | AR-FL |
| VIPR2 | -0.22 | -3.72 | AR-V7 | RHOBTB1 | -1.99 | -0.52 | AR-FL |
| SAMD5 | -0.38 | -3.18 | AR-V7 | CCNB1IP1 | -1.36 | -0.52 | AR-FL |
| SHC3 | -0.55 | -3.10 | AR-V7 | KCND3 | -0.61 | -0.52 | AR-FL |
| ANKRD65 | -0.43 | -3.04 | AR-V7 | SMKR1 | -1.80 | -0.51 | AR-FL |
| TNFRSF14 | -0.01 | -3.00 | AR-V7 | EIF2S3L | -0.72 | -0.51 | AR-FL |

| | | | | | | | |
|---|---|---|---|---|---|---|---|
| SLC40A1 | -0.31 | -2.98 | AR-V7 | MARK1 | -0.88 | -0.51 | AR-FL |
| SYNE1 | -0.54 | -2.97 | AR-V7 | PABPC4 | -0.79 | -0.51 | AR-FL |
| CNNM1 | -0.19 | -2.90 | AR-V7 | MACROD2 | -1.47 | -0.51 | AR-FL |
| CADM1 | -0.47 | -2.56 | AR-V7 | TATDN1 | -0.79 | -0.51 | AR-FL |
| CNTN4 | -0.35 | -2.46 | AR-V7 | ZFP90 | -1.18 | -0.51 | AR-FL |
| GDPD1 | -0.58 | -2.40 | AR-V7 | SLC7A2 | -0.65 | -0.51 | AR-FL |
| APOL6 | -0.28 | -2.20 | AR-V7 | ARID5A | -1.21 | -0.51 | AR-FL |
| SLC7A11 | -0.22 | -2.18 | AR-V7 | TRMT5 | -1.17 | -0.51 | AR-FL |
| ATXN1 | -0.23 | -2.09 | AR-V7 | ZNF285 | -0.78 | -0.51 | AR-FL |
| PGBD5 | -0.24 | -2.02 | AR-V7 | SPNS2 | -1.19 | -0.51 | AR-FL |
| GPRIN2 | -0.57 | -2.00 | AR-V7 | ZNF383 | -1.19 | -0.51 | AR-FL |
| CDH13 | -0.20 | -2.00 | AR-V7 | ABLIM1 | -0.66 | -0.51 | AR-FL |
| GPX2 | -0.43 | -1.99 | AR-V7 | IRAK1BP1 | -1.38 | -0.51 | AR-FL |
| PDGFA | -0.43 | -1.97 | AR-V7 | HTR6 | -1.31 | -0.51 | AR-FL |
| LONRF2 | -0.31 | -1.91 | AR-V7 | RFX7 | -0.67 | -0.51 | AR-FL |
| SORBS2 | -0.16 | -1.89 | AR-V7 | KIF3C | -0.72 | -0.51 | AR-FL |
| GIMAP2 | -0.42 | -1.84 | AR-V7 | SLC25A6 | -0.95 | -0.51 | AR-FL |
| B4GALNT3 | -0.08 | -1.82 | AR-V7 | KDSR | -0.93 | -0.51 | AR-FL |
| BANK1 | -0.54 | -1.81 | AR-V7 | GOLGA8A | -1.52 | -0.51 | AR-FL |
| MME | -0.17 | -1.77 | AR-V7 | HSD17B11 | -1.04 | -0.51 | AR-FL |
| SLC30A4 | -0.26 | -1.77 | AR-V7 | DDN | -1.53 | -0.51 | AR-FL |
| QKI | -0.19 | -1.76 | AR-V7 | TBC1D5 | -0.77 | -0.51 | AR-FL |
| LRFN2 | -0.43 | -1.73 | AR-V7 | FBXO4 | -0.99 | -0.51 | AR-FL |
| PLPP2 | -0.58 | -1.71 | AR-V7 | STK33 | -1.22 | -0.51 | AR-FL |
| PARP10 | -0.43 | -1.70 | AR-V7 | CRMP1 | -1.37 | -0.51 | AR-FL |
| SYT7 | -0.08 | -1.60 | AR-V7 | SLC12A8 | -0.59 | -0.51 | AR-FL |
| CCDC122 | 0.00 | -1.60 | AR-V7 | ASB11 | -1.01 | -0.50 | AR-FL |
| KIAA2022 | -0.07 | -1.59 | AR-V7 | TGFBR3 | -1.47 | -0.50 | AR-FL |
| RGS11 | -0.45 | -1.58 | AR-V7 | CPNE3 | -0.70 | -0.50 | AR-FL |
| BCAS1 | -0.34 | -1.54 | AR-V7 | SLC26A10 | -2.38 | -0.50 | AR-FL |
| SMPDL3B | -0.46 | -1.52 | AR-V7 | LRRC8D | -0.60 | -0.50 | AR-FL |
| CDC42EP1 | -0.30 | -1.52 | AR-V7 | UNC93B1 | -1.18 | -0.50 | AR-FL |
| DSC2 | -0.40 | -1.52 | AR-V7 | FLNB | -0.61 | -0.50 | AR-FL |
| TMEM40 | -0.33 | -1.51 | AR-V7 | NRIP2 | -0.63 | -0.50 | AR-FL |
| PLCG2 | -0.51 | -1.49 | AR-V7 | PDE5A | -0.83 | -0.50 | AR-FL |
| ZC3H12C | -0.56 | -1.49 | AR-V7 | SOBP | -2.54 | -0.50 | AR-FL |
| SV2C | -0.37 | -1.48 | AR-V7 | GABRA5 | -1.90 | -0.50 | AR-FL |
| FOXN3 | -0.11 | -1.47 | AR-V7 | DENND5A | -0.73 | -0.50 | AR-FL |
| PDIA2 | -0.57 | -1.47 | AR-V7 | HRSP12 | -1.26 | -0.50 | AR-FL |
| RTN1 | -0.21 | -1.45 | AR-V7 | NUP133 | -0.71 | -0.50 | AR-FL |
| LRBA | -0.52 | -1.43 | AR-V7 | LNP1 | -0.77 | -0.50 | AR-FL |
| LARGE | -0.18 | -1.41 | AR-V7 | JAKMIP3 | -0.67 | -0.50 | AR-FL |
| RLTPR | -0.36 | -1.37 | AR-V7 | CCDC27 | -2.17 | -0.50 | AR-FL |
| PAM | -0.15 | -1.37 | AR-V7 | NFASC | -1.64 | -0.50 | AR-FL |
| CGNL1 | -0.21 | -1.37 | AR-V7 | NTPCR | -0.93 | -0.50 | AR-FL |
| SLC39A8 | -0.55 | -1.34 | AR-V7 | SNAI1 | -0.86 | -0.50 | AR-FL |

| | | | |
|---|---|---|---|
| MFSD6L | -0.30 | -1.32 | AR-V7 |
| NXPH4 | -0.16 | -1.32 | AR-V7 |
| CDK14 | -0.12 | -1.32 | AR-V7 |
| DSE | -0.48 | -1.31 | AR-V7 |
| SLC9A9 | -0.14 | -1.31 | AR-V7 |
| CBR1 | -0.13 | -1.31 | AR-V7 |
| SATB2 | -0.57 | -1.29 | AR-V7 |
| NQO1 | -0.09 | -1.28 | AR-V7 |
| ENOX1 | -0.27 | -1.27 | AR-V7 |
| BICD1 | -0.12 | -1.27 | AR-V7 |
| SLC35G1 | -0.49 | -1.27 | AR-V7 |
| PLXNA2 | -0.08 | -1.25 | AR-V7 |
| ASPHD1 | -0.11 | -1.25 | AR-V7 |
| PELI1 | -0.36 | -1.24 | AR-V7 |
| KIAA1024 | -0.52 | -1.23 | AR-V7 |
| KCNQ2 | -0.47 | -1.21 | AR-V7 |
| PPARG | -0.38 | -1.21 | AR-V7 |
| CTSC | -0.29 | -1.19 | AR-V7 |
| DCHS1 | -0.28 | -1.18 | AR-V7 |
| SLC13A3 | -0.27 | -1.18 | AR-V7 |
| RASEF | -0.42 | -1.17 | AR-V7 |
| PITPNC1 | -0.41 | -1.17 | AR-V7 |
| CXXC5 | -0.53 | -1.14 | AR-V7 |
| KLF8 | -0.35 | -1.14 | AR-V7 |
| FAXC | -0.57 | -1.13 | AR-V7 |
| SLC23A1 | -0.10 | -1.12 | AR-V7 |
| KIF1A | -0.54 | -1.12 | AR-V7 |
| PLCB1 | -0.43 | -1.11 | AR-V7 |
| CNTNAP3 | -0.23 | -1.11 | AR-V7 |
| PPP2R2C | -0.47 | -1.11 | AR-V7 |
| AEBP1 | -0.25 | -1.10 | AR-V7 |
| BLOC1S6 | -0.31 | -1.09 | AR-V7 |
| PLK2 | -0.20 | -1.09 | AR-V7 |
| NPTXR | -0.01 | -1.08 | AR-V7 |
| SLC6A6 | -0.52 | -1.07 | AR-V7 |
| FTCDNL1 | -0.34 | -1.07 | AR-V7 |
| HIVEP3 | -0.42 | -1.06 | AR-V7 |
| SLC16A1 | -0.09 | -1.05 | AR-V7 |
| STAMBPL1 | -0.18 | -1.05 | AR-V7 |
| NETO1 | -0.48 | -1.04 | AR-V7 |
| COL26A1 | -0.17 | -1.03 | AR-V7 |
| PIM1 | -0.33 | -1.02 | AR-V7 |
| APBB2 | -0.17 | -1.02 | AR-V7 |
| SERAC1 | -0.43 | -1.02 | AR-V7 |
| VEGFB | -0.04 | -1.02 | AR-V7 |
| AP3B2 | -0.43 | -1.01 | AR-V7 |

| | | | |
|---|---|---|---|
| TRIM21 | -0.76 | -0.50 | AR-FL |
| ERBB2 | -1.11 | -0.50 | AR-FL |
| VAT1 | -0.68 | -0.50 | AR-FL |
| RBMX2 | -0.72 | -0.50 | AR-FL |
| FAM120C | -1.18 | -0.50 | AR-FL |
| PTAR1 | -0.73 | -0.50 | AR-FL |
| FAXDC2 | -1.31 | -0.50 | AR-FL |
| PATZ1 | -0.88 | -0.50 | AR-FL |
| MLXIPL | -1.41 | -0.49 | AR-FL |
| TMEM245 | -0.83 | -0.49 | AR-FL |
| ZNF395 | -1.21 | -0.49 | AR-FL |
| AIFM3 | -0.75 | -0.49 | AR-FL |
| FAM198B | -1.07 | -0.49 | AR-FL |
| FSCN1 | -1.50 | -0.49 | AR-FL |
| ZSCAN30 | -0.80 | -0.49 | AR-FL |
| LRRC34 | -0.62 | -0.49 | AR-FL |
| LPCAT2 | -0.91 | -0.49 | AR-FL |
| C5orf45 | -1.43 | -0.49 | AR-FL |
| DNAH10 | -0.61 | -0.49 | AR-FL |
| DPY19L4 | -0.78 | -0.49 | AR-FL |
| ZNF514 | -1.03 | -0.49 | AR-FL |
| TMEM128 | -0.77 | -0.49 | AR-FL |
| C1R | -0.98 | -0.49 | AR-FL |
| MMP11 | -0.69 | -0.49 | AR-FL |
| RASGEF1C | -2.07 | -0.49 | AR-FL |
| CCDC88A | -1.74 | -0.49 | AR-FL |
| ANKRD13B | -1.06 | -0.49 | AR-FL |
| ATXN3 | -0.64 | -0.49 | AR-FL |
| MEX3A | -0.69 | -0.49 | AR-FL |
| CA11 | -2.53 | -0.49 | AR-FL |
| C16orf62 | -0.94 | -0.49 | AR-FL |
| TP53 | -1.03 | -0.49 | AR-FL |
| ZNRF1 | -0.61 | -0.49 | AR-FL |
| BLOC1S5 | -0.67 | -0.49 | AR-FL |
| LGI2 | -1.92 | -0.49 | AR-FL |
| USP27X | -1.05 | -0.49 | AR-FL |
| ARRDC2 | -0.87 | -0.49 | AR-FL |
| PCBP2 | -0.69 | -0.49 | AR-FL |
| GAB2 | -1.04 | -0.49 | AR-FL |
| AEBP2 | -0.64 | -0.48 | AR-FL |
| ZNF362 | -0.80 | -0.48 | AR-FL |
| LOXL3 | -1.60 | -0.48 | AR-FL |
| WDR3 | -0.69 | -0.48 | AR-FL |
| NUDT6 | -1.24 | -0.48 | AR-FL |
| EIF2A | -0.68 | -0.48 | AR-FL |
| NFAT5 | -1.04 | -0.48 | AR-FL |

| | | | | | | | |
|---|---|---|---|---|---|---|---|
| STEAP3 | -0.27 | -1.01 | AR-V7 | TTK | -0.97 | -0.48 | AR-FL |
| PPP2R2A | -0.57 | -1.00 | AR-V7 | DYRK2 | -0.72 | -0.48 | AR-FL |
| SLC9A3R1 | -0.21 | -1.00 | AR-V7 | DIS3L | -1.08 | -0.48 | AR-FL |
| ISX | -0.57 | -1.00 | AR-V7 | HMG20A | -0.74 | -0.48 | AR-FL |
| UQCC1 | -0.35 | -1.00 | AR-V7 | MEIS1 | -2.37 | -0.48 | AR-FL |
| STAC3 | -0.29 | -1.00 | AR-V7 | ZKSCAN8 | -1.13 | -0.48 | AR-FL |
| RAB23 | -0.25 | -0.98 | AR-V7 | DNM1 | -1.17 | -0.48 | AR-FL |
| C17orf51 | -0.29 | -0.98 | AR-V7 | HSPG2 | -1.27 | -0.48 | AR-FL |
| MBP | -0.02 | -0.97 | AR-V7 | PRIM1 | -1.22 | -0.48 | AR-FL |
| CEBPA | -0.54 | -0.97 | AR-V7 | BRWD3 | -0.85 | -0.48 | AR-FL |
| TRIM7 | -0.38 | -0.97 | AR-V7 | ADAMTS13 | -1.60 | -0.48 | AR-FL |
| CCDC67 | -0.29 | -0.96 | AR-V7 | KLC4 | -0.69 | -0.48 | AR-FL |
| FLRT1 | -0.48 | -0.96 | AR-V7 | GABPB2 | -1.15 | -0.48 | AR-FL |
| BOK | -0.26 | -0.96 | AR-V7 | RMND1 | -1.05 | -0.48 | AR-FL |
| XPO5 | -0.20 | -0.96 | AR-V7 | UNC13D | -1.96 | -0.48 | AR-FL |
| DHRS13 | -0.23 | -0.96 | AR-V7 | HEBP1 | -0.83 | -0.48 | AR-FL |
| MTHFD2 | -0.54 | -0.95 | AR-V7 | EFCAB13 | -1.63 | -0.48 | AR-FL |
| CDKN2C | -0.57 | -0.95 | AR-V7 | CTH | -0.81 | -0.48 | AR-FL |
| EIF4EBP1 | -0.32 | -0.95 | AR-V7 | ZNF510 | -0.68 | -0.48 | AR-FL |
| EPHB3 | -0.17 | -0.95 | AR-V7 | ADGRV1 | -1.13 | -0.48 | AR-FL |
| NCAM2 | -0.10 | -0.94 | AR-V7 | NT5M | -1.97 | -0.47 | AR-FL |
| STBD1 | -0.57 | -0.94 | AR-V7 | VAMP7 | -1.01 | -0.47 | AR-FL |
| ANKRD44 | -0.45 | -0.94 | AR-V7 | ZDHHC14 | -0.60 | -0.47 | AR-FL |
| TPD52L1 | -0.01 | -0.94 | AR-V7 | DET1 | -0.80 | -0.47 | AR-FL |
| TUSC1 | -0.52 | -0.93 | AR-V7 | LONRF2 | -1.00 | -0.47 | AR-FL |
| SLC25A21 | -0.39 | -0.93 | AR-V7 | CLNS1A | -0.85 | -0.47 | AR-FL |
| TMTC4 | -0.13 | -0.93 | AR-V7 | DSTYK | -0.67 | -0.47 | AR-FL |
| ABLIM1 | -0.54 | -0.92 | AR-V7 | LYRM9 | -1.37 | -0.47 | AR-FL |
| ITGA6 | -0.06 | -0.92 | AR-V7 | SHF | -1.26 | -0.47 | AR-FL |
| CBFA2T2 | -0.08 | -0.91 | AR-V7 | RSL1D1 | -0.84 | -0.47 | AR-FL |
| TTC39B | -0.57 | -0.91 | AR-V7 | CTSC | -0.92 | -0.47 | AR-FL |
| SNTA1 | -0.35 | -0.90 | AR-V7 | SETD9 | -0.66 | -0.47 | AR-FL |
| SETBP1 | -0.09 | -0.90 | AR-V7 | GSPT2 | -0.60 | -0.47 | AR-FL |
| MAPK13 | -0.33 | -0.90 | AR-V7 | LEF1 | -0.68 | -0.47 | AR-FL |
| PDE3B | -0.12 | -0.90 | AR-V7 | KIF17 | -0.89 | -0.47 | AR-FL |
| AVEN | -0.17 | -0.89 | AR-V7 | TDRD5 | -0.66 | -0.47 | AR-FL |
| TTC34 | -0.57 | -0.89 | AR-V7 | EVC | -1.16 | -0.47 | AR-FL |
| PTK6 | -0.47 | -0.89 | AR-V7 | KLF11 | -0.87 | -0.47 | AR-FL |
| ACOX2 | -0.12 | -0.88 | AR-V7 | PXK | -0.80 | -0.47 | AR-FL |
| ARHGAP18 | -0.49 | -0.88 | AR-V7 | POT1 | -0.71 | -0.47 | AR-FL |
| GARS | -0.35 | -0.88 | AR-V7 | MTR | -1.12 | -0.47 | AR-FL |
| AHI1 | -0.15 | -0.88 | AR-V7 | CLCN5 | -0.66 | -0.47 | AR-FL |
| SLCO4A1 | -0.28 | -0.88 | AR-V7 | FAIM | -0.60 | -0.47 | AR-FL |
| GPHN | -0.23 | -0.88 | AR-V7 | ZNF566 | -0.75 | -0.47 | AR-FL |
| NIPAL2 | -0.23 | -0.88 | AR-V7 | GMCL1 | -0.72 | -0.47 | AR-FL |
| IFI35 | -0.57 | -0.87 | AR-V7 | ALOX12B | -0.60 | -0.47 | AR-FL |

| | | | | | | | |
|---|---|---|---|---|---|---|---|
| SHMT2 | -0.02 | -0.87 | AR-V7 | C12orf74 | -2.58 | -0.47 | AR-FL |
| FAM3B | -0.46 | -0.86 | AR-V7 | ZNF862 | -1.13 | -0.47 | AR-FL |
| CGN | -0.27 | -0.86 | AR-V7 | WASL | -0.64 | -0.47 | AR-FL |
| UBE3D | -0.19 | -0.86 | AR-V7 | C2CD5 | -0.87 | -0.47 | AR-FL |
| REXO2 | -0.39 | -0.86 | AR-V7 | CFD | -0.69 | -0.47 | AR-FL |
| ITGA3 | -0.57 | -0.86 | AR-V7 | MGAT3 | -1.88 | -0.47 | AR-FL |
| SHB | -0.02 | -0.86 | AR-V7 | RASIP1 | -1.26 | -0.47 | AR-FL |
| KDM6A | -0.47 | -0.86 | AR-V7 | PRDM11 | -0.68 | -0.46 | AR-FL |
| MAST4 | -0.20 | -0.85 | AR-V7 | RGL1 | -1.14 | -0.46 | AR-FL |
| PCDHGA9 | -0.46 | -0.84 | AR-V7 | MKS1 | -0.83 | -0.46 | AR-FL |
| YPEL2 | -0.46 | -0.84 | AR-V7 | ZSWIM5 | -0.82 | -0.46 | AR-FL |
| DTNB | -0.38 | -0.84 | AR-V7 | ZSCAN23 | -1.56 | -0.46 | AR-FL |
| NCOR2 | -0.41 | -0.84 | AR-V7 | RCN3 | -1.04 | -0.46 | AR-FL |
| MDN1 | -0.13 | -0.84 | AR-V7 | PFKM | -0.92 | -0.46 | AR-FL |
| RTEL1-TNFRSF6B | -0.35 | -0.83 | AR-V7 | PTPN4 | -0.75 | -0.46 | AR-FL |
| SH3RF3 | -0.22 | -0.83 | AR-V7 | MRPS31 | -0.73 | -0.46 | AR-FL |
| IFIH1 | -0.20 | -0.82 | AR-V7 | ATF4 | -0.83 | -0.46 | AR-FL |
| IMMP2L | -0.39 | -0.81 | AR-V7 | LOXL1 | -0.98 | -0.46 | AR-FL |
| ASAP3 | -0.42 | -0.81 | AR-V7 | BZRAP1 | -2.00 | -0.46 | AR-FL |
| TMEM238 | -0.48 | -0.80 | AR-V7 | TXNRD3 | -1.00 | -0.46 | AR-FL |
| HGH1 | -0.07 | -0.80 | AR-V7 | IRF8 | -2.65 | -0.46 | AR-FL |
| SAMD13 | -0.52 | -0.80 | AR-V7 | WDR12 | -0.86 | -0.46 | AR-FL |
| PANK1 | -0.46 | -0.80 | AR-V7 | FREM1 | -1.90 | -0.46 | AR-FL |
| TUBA1B | -0.22 | -0.80 | AR-V7 | MRE11A | -0.94 | -0.46 | AR-FL |
| TRIM25 | -0.56 | -0.80 | AR-V7 | TRPC7 | -0.83 | -0.46 | AR-FL |
| DST | -0.53 | -0.79 | AR-V7 | MYO1B | -0.82 | -0.46 | AR-FL |
| TRAPPC9 | -0.35 | -0.79 | AR-V7 | ANKRD36B | -1.31 | -0.46 | AR-FL |
| MTHFD1L | -0.29 | -0.79 | AR-V7 | C3orf18 | -0.86 | -0.46 | AR-FL |
| RAB9B | -0.25 | -0.79 | AR-V7 | PDE6B | -1.76 | -0.46 | AR-FL |
| GSTA4 | -0.26 | -0.78 | AR-V7 | COCH | -0.80 | -0.46 | AR-FL |
| SLC4A3 | -0.45 | -0.78 | AR-V7 | GAL3ST2 | -1.38 | -0.46 | AR-FL |
| ADK | -0.25 | -0.78 | AR-V7 | MFSD7 | -1.25 | -0.46 | AR-FL |
| NETO2 | -0.52 | -0.78 | AR-V7 | ALDH5A1 | -1.19 | -0.45 | AR-FL |
| ATIC | -0.42 | -0.78 | AR-V7 | KIAA1551 | -0.59 | -0.45 | AR-FL |
| PDSS2 | -0.40 | -0.77 | AR-V7 | NEIL1 | -1.52 | -0.45 | AR-FL |
| ZMYND11 | -0.45 | -0.77 | AR-V7 | ATP6V1E2 | -0.84 | -0.45 | AR-FL |
| PFAS | -0.47 | -0.77 | AR-V7 | RARA | -1.02 | -0.45 | AR-FL |
| FAM47E | -0.53 | -0.77 | AR-V7 | FTCDNL1 | -0.86 | -0.45 | AR-FL |
| RAB9A | -0.20 | -0.76 | AR-V7 | SHISA2 | -0.85 | -0.45 | AR-FL |
| RBPMS2 | -0.33 | -0.76 | AR-V7 | FAM228B | -1.57 | -0.45 | AR-FL |
| STAP2 | -0.23 | -0.76 | AR-V7 | GRIK5 | -1.01 | -0.45 | AR-FL |
| THBS4 | -0.52 | -0.76 | AR-V7 | ZNF214 | -1.43 | -0.45 | AR-FL |
| ARHGEF6 | -0.51 | -0.76 | AR-V7 | RHBDF2 | -1.15 | -0.45 | AR-FL |
| NMU | -0.13 | -0.75 | AR-V7 | SYDE1 | -1.12 | -0.45 | AR-FL |
| KCTD15 | -0.47 | -0.75 | AR-V7 | CYB5RL | -1.09 | -0.45 | AR-FL |
| PNP | -0.22 | -0.75 | AR-V7 | ENPP4 | -0.62 | -0.45 | AR-FL |

| | | | |
|---|---|---|---|
| FAM213B | -0.50 | -0.75 | AR-V7 |
| 2-Mar | -0.37 | -0.75 | AR-V7 |
| CTNNAL1 | -0.38 | -0.75 | AR-V7 |
| PRDX1 | -0.51 | -0.75 | AR-V7 |
| PHF12 | -0.37 | -0.75 | AR-V7 |
| TCF7 | -0.37 | -0.75 | AR-V7 |
| SCPEP1 | -0.30 | -0.75 | AR-V7 |
| CACYBP | -0.46 | -0.75 | AR-V7 |
| WASF3 | -0.40 | -0.75 | AR-V7 |
| PDCD2 | -0.17 | -0.75 | AR-V7 |
| ARHGAP32 | -0.14 | -0.74 | AR-V7 |
| NFU1 | -0.16 | -0.74 | AR-V7 |
| CCDC154 | -0.46 | -0.74 | AR-V7 |
| TTC28 | -0.28 | -0.74 | AR-V7 |
| CMTM7 | -0.53 | -0.74 | AR-V7 |
| MBNL3 | -0.41 | -0.74 | AR-V7 |
| CEP57L1 | -0.22 | -0.74 | AR-V7 |
| USP45 | -0.25 | -0.74 | AR-V7 |
| SAMD1 | -0.46 | -0.74 | AR-V7 |
| C1orf115 | -0.01 | -0.73 | AR-V7 |
| PCMTD1 | -0.25 | -0.73 | AR-V7 |
| LEO1 | -0.04 | -0.73 | AR-V7 |
| RNF6 | -0.25 | -0.73 | AR-V7 |
| MSRA | -0.03 | -0.73 | AR-V7 |
| G6PD | -0.19 | -0.73 | AR-V7 |
| TRAF3 | -0.26 | -0.73 | AR-V7 |
| NPW | -0.45 | -0.73 | AR-V7 |
| PIEZO1 | -0.04 | -0.72 | AR-V7 |
| UHRF1BP1L | -0.08 | -0.72 | AR-V7 |
| DCAKD | -0.38 | -0.72 | AR-V7 |
| CERS4 | -0.52 | -0.72 | AR-V7 |
| ERVMER34-1 | -0.34 | -0.72 | AR-V7 |
| HSP90AB1 | -0.25 | -0.72 | AR-V7 |
| RBL2 | -0.22 | -0.71 | AR-V7 |
| SMC2 | -0.50 | -0.71 | AR-V7 |
| MICU1 | -0.49 | -0.71 | AR-V7 |
| PCSK1N | -0.09 | -0.71 | AR-V7 |
| LRP11 | -0.41 | -0.70 | AR-V7 |
| CAT | -0.38 | -0.70 | AR-V7 |
| JAKMIP3 | -0.07 | -0.70 | AR-V7 |
| AKTIP | -0.38 | -0.70 | AR-V7 |
| ATP10D | -0.50 | -0.70 | AR-V7 |
| ELP4 | -0.55 | -0.70 | AR-V7 |
| SCFD2 | -0.12 | -0.69 | AR-V7 |
| A2ML1 | -0.46 | -0.69 | AR-V7 |
| TMSB15A | -0.09 | -0.69 | AR-V7 |
| SKOR1 | -0.89 | -0.45 | AR-FL |
| VIPAS39 | -0.80 | -0.45 | AR-FL |
| FUZ | -0.68 | -0.45 | AR-FL |
| STX2 | -0.85 | -0.45 | AR-FL |
| ZNF382 | -0.97 | -0.45 | AR-FL |
| POU4F1 | -1.45 | -0.45 | AR-FL |
| IKZF4 | -0.79 | -0.45 | AR-FL |
| LGALS8 | -0.93 | -0.45 | AR-FL |
| ATIC | -0.70 | -0.45 | AR-FL |
| GNB1L | -0.71 | -0.45 | AR-FL |
| SBF2 | -0.86 | -0.45 | AR-FL |
| ZNF496 | -0.91 | -0.45 | AR-FL |
| PLK3 | -1.12 | -0.45 | AR-FL |
| BIVM | -0.68 | -0.45 | AR-FL |
| ZNF234 | -0.81 | -0.45 | AR-FL |
| BCAS3 | -0.75 | -0.45 | AR-FL |
| NAP1L1 | -1.11 | -0.45 | AR-FL |
| RPSA | -0.66 | -0.45 | AR-FL |
| ACE | -1.53 | -0.44 | AR-FL |
| ADAM33 | -0.66 | -0.44 | AR-FL |
| GRIP2 | -1.61 | -0.44 | AR-FL |
| KIAA0895 | -0.93 | -0.44 | AR-FL |
| TPP2 | -0.69 | -0.44 | AR-FL |
| DNAJC27 | -0.97 | -0.44 | AR-FL |
| PAX8 | -2.04 | -0.44 | AR-FL |
| MKNK1 | -0.67 | -0.44 | AR-FL |
| METTL21B | -1.09 | -0.44 | AR-FL |
| EPHB4 | -1.06 | -0.44 | AR-FL |
| CHML | -1.25 | -0.44 | AR-FL |
| GRB7 | -0.94 | -0.44 | AR-FL |
| OSER1 | -0.71 | -0.44 | AR-FL |
| GREB1L | -1.42 | -0.44 | AR-FL |
| DECR1 | -1.19 | -0.44 | AR-FL |
| CEP95 | -1.26 | -0.44 | AR-FL |
| NR1H4 | -1.31 | -0.44 | AR-FL |
| PMS2 | -0.65 | -0.44 | AR-FL |
| ANAPC4 | -1.01 | -0.44 | AR-FL |
| MC1R | -1.38 | -0.44 | AR-FL |
| BEND3 | -0.98 | -0.44 | AR-FL |
| TMEM102 | -0.68 | -0.44 | AR-FL |
| PCDHB15 | -1.29 | -0.44 | AR-FL |
| NEFM | -1.44 | -0.44 | AR-FL |
| ZDHHC2 | -0.71 | -0.44 | AR-FL |
| TTC33 | -0.93 | -0.44 | AR-FL |
| YPEL3 | -1.35 | -0.44 | AR-FL |
| ZNF133 | -0.92 | -0.44 | AR-FL |

| | | | |
|---|---|---|---|
| ZFYVE9 | -0.28 | -0.69 | AR-V7 |
| ETFDH | -0.37 | -0.69 | AR-V7 |
| PKHD1L1 | -0.54 | -0.69 | AR-V7 |
| PRSS27 | -0.48 | -0.69 | AR-V7 |
| CCDC110 | -0.10 | -0.68 | AR-V7 |
| GATA6 | -0.47 | -0.68 | AR-V7 |
| CFD | -0.57 | -0.68 | AR-V7 |
| TSPEAR | -0.48 | -0.68 | AR-V7 |
| ZP3 | -0.30 | -0.68 | AR-V7 |
| NUAK1 | -0.57 | -0.68 | AR-V7 |
| TMEM123 | -0.57 | -0.68 | AR-V7 |
| ARSB | -0.28 | -0.68 | AR-V7 |
| SCML2 | -0.53 | -0.68 | AR-V7 |
| C2CD2 | -0.21 | -0.68 | AR-V7 |
| CCND1 | -0.27 | -0.68 | AR-V7 |
| STK24 | -0.17 | -0.68 | AR-V7 |
| PTMA | -0.20 | -0.68 | AR-V7 |
| TYRO3 | -0.24 | -0.67 | AR-V7 |
| POLD2 | -0.23 | -0.67 | AR-V7 |
| PRR19 | -0.15 | -0.67 | AR-V7 |
| NQO2 | -0.17 | -0.67 | AR-V7 |
| OSBP2 | -0.51 | -0.67 | AR-V7 |
| DBNDD1 | -0.19 | -0.67 | AR-V7 |
| MRPL14 | -0.16 | -0.67 | AR-V7 |
| C19orf66 | -0.13 | -0.67 | AR-V7 |
| R3HDM1 | -0.39 | -0.67 | AR-V7 |
| ADAT2 | -0.25 | -0.67 | AR-V7 |
| VWA1 | -0.37 | -0.66 | AR-V7 |
| IGSF3 | -0.34 | -0.66 | AR-V7 |
| CHTF18 | -0.54 | -0.66 | AR-V7 |
| PKN1 | -0.28 | -0.66 | AR-V7 |
| XPOT | -0.12 | -0.66 | AR-V7 |
| CPS1 | -0.43 | -0.66 | AR-V7 |
| CSPG5 | -0.33 | -0.66 | AR-V7 |
| SGMS1 | -0.50 | -0.66 | AR-V7 |
| POU2F1 | -0.42 | -0.66 | AR-V7 |
| TBC1D5 | -0.47 | -0.66 | AR-V7 |
| UHRF1BP1 | -0.10 | -0.66 | AR-V7 |
| RTN4RL2 | -0.42 | -0.66 | AR-V7 |
| SLC7A1 | -0.15 | -0.66 | AR-V7 |
| PAK2 | -0.51 | -0.65 | AR-V7 |
| ZFYVE28 | -0.53 | -0.65 | AR-V7 |
| EEFSEC | -0.52 | -0.65 | AR-V7 |
| PPP1R3E | -0.17 | -0.65 | AR-V7 |
| APOBEC3F | -0.27 | -0.65 | AR-V7 |
| SPHK1 | -0.52 | -0.65 | AR-V7 |

| | | | |
|---|---|---|---|
| C12orf76 | -0.99 | -0.44 | AR-FL |
| C12orf29 | -0.84 | -0.44 | AR-FL |
| SUSD5 | -1.64 | -0.44 | AR-FL |
| ATP1A3 | -1.95 | -0.43 | AR-FL |
| FBLL1 | -1.05 | -0.43 | AR-FL |
| YAE1D1 | -0.68 | -0.43 | AR-FL |
| HIBADH | -0.70 | -0.43 | AR-FL |
| PDK1 | -0.73 | -0.43 | AR-FL |
| NREP | -0.72 | -0.43 | AR-FL |
| RHOV | -0.75 | -0.43 | AR-FL |
| IKZF3 | -1.07 | -0.43 | AR-FL |
| BHLHB9 | -1.01 | -0.43 | AR-FL |
| EGR1 | -0.78 | -0.43 | AR-FL |
| TTR | -2.46 | -0.43 | AR-FL |
| ARHGAP5 | -0.68 | -0.43 | AR-FL |
| KIAA1958 | -0.90 | -0.43 | AR-FL |
| KLB | -0.98 | -0.43 | AR-FL |
| EML5 | -0.69 | -0.43 | AR-FL |
| SGCE | -0.98 | -0.43 | AR-FL |
| RAMP1 | -0.98 | -0.43 | AR-FL |
| SLC2A11 | -0.93 | -0.43 | AR-FL |
| KALRN | -0.80 | -0.43 | AR-FL |
| SETDB1 | -0.77 | -0.43 | AR-FL |
| SLC35D2 | -0.70 | -0.43 | AR-FL |
| SLC25A27 | -1.76 | -0.43 | AR-FL |
| TOMM20 | -0.76 | -0.43 | AR-FL |
| DOC2A | -2.38 | -0.43 | AR-FL |
| FAM171A1 | -1.10 | -0.43 | AR-FL |
| PGAP1 | -1.69 | -0.43 | AR-FL |
| KCNAB2 | -2.01 | -0.43 | AR-FL |
| ORC3 | -0.81 | -0.43 | AR-FL |
| HNRNPA1L2 | -0.80 | -0.42 | AR-FL |
| DTX3 | -0.86 | -0.42 | AR-FL |
| GTF2IRD2 | -0.79 | -0.42 | AR-FL |
| KCNJ12 | -1.80 | -0.42 | AR-FL |
| SGCB | -1.16 | -0.42 | AR-FL |
| KIAA1683 | -1.67 | -0.42 | AR-FL |
| FOXL2NB | -1.09 | -0.42 | AR-FL |
| ANKRD36C | -1.99 | -0.42 | AR-FL |
| AKAP5 | -0.89 | -0.42 | AR-FL |
| CYP7B1 | -0.99 | -0.42 | AR-FL |
| STRADB | -0.78 | -0.42 | AR-FL |
| KIAA1324 | -0.84 | -0.42 | AR-FL |
| HNRNPA1 | -0.93 | -0.42 | AR-FL |
| TFDP2 | -1.13 | -0.42 | AR-FL |
| GLMN | -0.63 | -0.42 | AR-FL |

| | | | |
|---|---|---|---|
| FAM216A | -0.58 | -0.65 | AR-V7 |
| YARS | -0.09 | -0.65 | AR-V7 |
| LRP8 | -0.13 | -0.65 | AR-V7 |
| MPP7 | -0.27 | -0.65 | AR-V7 |
| FKBP4 | -0.28 | -0.64 | AR-V7 |
| IGSF1 | -0.21 | -0.64 | AR-V7 |
| TUBA1A | -0.39 | -0.64 | AR-V7 |
| RRP1B | -0.15 | -0.64 | AR-V7 |
| DYNLT1 | -0.34 | -0.64 | AR-V7 |
| TFAP2E | -0.19 | -0.64 | AR-V7 |
| FKBP10 | -0.19 | -0.64 | AR-V7 |
| C3orf33 | -0.18 | -0.63 | AR-V7 |
| TKT | -0.24 | -0.63 | AR-V7 |
| FAM124A | -0.13 | -0.63 | AR-V7 |
| RRP1 | -0.14 | -0.63 | AR-V7 |
| ZNF704 | -0.56 | -0.63 | AR-V7 |
| TLE2 | -0.39 | -0.63 | AR-V7 |
| TSNARE1 | -0.34 | -0.63 | AR-V7 |
| TMEM97 | -0.38 | -0.63 | AR-V7 |
| GRASP | -0.05 | -0.62 | AR-V7 |
| SH3GLB2 | -0.42 | -0.62 | AR-V7 |
| SPTY2D1-AS1 | -0.21 | -0.62 | AR-V7 |
| NR1D2 | -0.34 | -0.62 | AR-V7 |
| PSMG4 | -0.29 | -0.62 | AR-V7 |
| TPK1 | -0.17 | -0.62 | AR-V7 |
| SPIN4 | -0.30 | -0.62 | AR-V7 |
| ARHGEF38 | -0.50 | -0.62 | AR-V7 |
| CC2D2A | -0.24 | -0.62 | AR-V7 |
| RAP2A | -0.27 | -0.62 | AR-V7 |
| FZD6 | -0.26 | -0.62 | AR-V7 |
| DIMT1 | -0.25 | -0.62 | AR-V7 |
| SNED1 | -0.36 | -0.61 | AR-V7 |
| LRRC8D | -0.56 | -0.61 | AR-V7 |
| NBEAL2 | -0.38 | -0.61 | AR-V7 |
| IPO4 | -0.21 | -0.61 | AR-V7 |
| SYNJ2 | -0.37 | -0.61 | AR-V7 |
| PSME1 | -0.37 | -0.61 | AR-V7 |
| OXCT1 | -0.19 | -0.61 | AR-V7 |
| TMEM243 | -0.37 | -0.61 | AR-V7 |
| HPRT1 | -0.24 | -0.61 | AR-V7 |
| SERTAD2 | -0.38 | -0.61 | AR-V7 |
| PKD2 | -0.28 | -0.61 | AR-V7 |
| SEMA3F | -0.24 | -0.61 | AR-V7 |
| ZCCHC24 | -0.14 | -0.61 | AR-V7 |
| CENPJ | -0.34 | -0.61 | AR-V7 |
| PSME2 | -0.04 | -0.61 | AR-V7 |
| ZCCHC11 | -0.72 | -0.42 | AR-FL |
| TMEM42 | -0.61 | -0.42 | AR-FL |
| EIF2S3 | -0.67 | -0.42 | AR-FL |
| DCAF4 | -1.38 | -0.42 | AR-FL |
| PNCK | -2.09 | -0.42 | AR-FL |
| RNF144A | -0.61 | -0.42 | AR-FL |
| C11orf57 | -0.68 | -0.42 | AR-FL |
| FJX1 | -1.35 | -0.42 | AR-FL |
| IGFALS | -1.10 | -0.42 | AR-FL |
| OTUD7A | -1.50 | -0.42 | AR-FL |
| CDHR1 | -3.83 | -0.42 | AR-FL |
| TMEM178A | -1.52 | -0.42 | AR-FL |
| CALD1 | -0.60 | -0.42 | AR-FL |
| SLC44A1 | -0.78 | -0.42 | AR-FL |
| CLIP2 | -0.58 | -0.41 | AR-FL |
| MAST1 | -1.44 | -0.41 | AR-FL |
| ZNF302 | -0.67 | -0.41 | AR-FL |
| EFEMP2 | -0.65 | -0.41 | AR-FL |
| FAM78B | -1.23 | -0.41 | AR-FL |
| LRIG2 | -0.78 | -0.41 | AR-FL |
| NECAB2 | -1.36 | -0.41 | AR-FL |
| ZNF501 | -0.75 | -0.41 | AR-FL |
| ZNF558 | -0.91 | -0.41 | AR-FL |
| TTC28 | -0.95 | -0.41 | AR-FL |
| TTC34 | -1.74 | -0.41 | AR-FL |
| COL6A1 | -1.71 | -0.41 | AR-FL |
| C11orf53 | -1.73 | -0.41 | AR-FL |
| AP1AR | -0.78 | -0.41 | AR-FL |
| NDRG2 | -0.65 | -0.41 | AR-FL |
| ZBTB44 | -0.93 | -0.41 | AR-FL |
| TBPL1 | -0.95 | -0.41 | AR-FL |
| RFX2 | -0.59 | -0.41 | AR-FL |
| TYMP | -0.99 | -0.41 | AR-FL |
| MCF2L2 | -0.88 | -0.41 | AR-FL |
| RIOK2 | -0.60 | -0.41 | AR-FL |
| SLC16A14 | -0.72 | -0.40 | AR-FL |
| COG8 | -0.59 | -0.40 | AR-FL |
| SLC24A1 | -0.79 | -0.40 | AR-FL |
| ZNF662 | -0.77 | -0.40 | AR-FL |
| WDSUB1 | -0.83 | -0.40 | AR-FL |
| CCDC115 | -0.89 | -0.40 | AR-FL |
| CGREF1 | -0.85 | -0.40 | AR-FL |
| NEK9 | -0.73 | -0.40 | AR-FL |
| AJUBA | -1.17 | -0.40 | AR-FL |
| YJEFN3 | -1.47 | -0.40 | AR-FL |
| SLC16A9 | -2.64 | -0.40 | AR-FL |

| | | | | | | | |
|---|---|---|---|---|---|---|---|
| ZRANB3 | -0.48 | -0.61 | AR-V7 | PALD1 | -1.35 | -0.40 | AR-FL |
| GTF2H2C | -0.28 | -0.61 | AR-V7 | ZBTB32 | -0.59 | -0.40 | AR-FL |
| RNF34 | -0.48 | -0.61 | AR-V7 | APC2 | -0.85 | -0.40 | AR-FL |
| PEX7 | -0.16 | -0.61 | AR-V7 | ST13 | -0.66 | -0.40 | AR-FL |
| NOD1 | -0.21 | -0.61 | AR-V7 | PUS10 | -0.79 | -0.40 | AR-FL |
| USP25 | -0.18 | -0.61 | AR-V7 | NOL8 | -0.97 | -0.40 | AR-FL |
| RCN1 | -0.02 | -0.61 | AR-V7 | HACE1 | -0.66 | -0.40 | AR-FL |
| FAM156B | -0.03 | -0.60 | AR-V7 | ABCA4 | -0.97 | -0.39 | AR-FL |
| RFX5 | -0.10 | -0.60 | AR-V7 | CHKB-CPT1B | -1.00 | -0.39 | AR-FL |
| CPNE7 | -0.37 | -0.60 | AR-V7 | RMDN2 | -0.82 | -0.39 | AR-FL |
| PCDHB14 | -0.57 | -0.60 | AR-V7 | STC2 | -1.02 | -0.39 | AR-FL |
| SMYD3 | -0.47 | -0.60 | AR-V7 | TRIM66 | -1.41 | -0.39 | AR-FL |
| MYH10 | -0.35 | -0.60 | AR-V7 | ZNF112 | -0.61 | -0.39 | AR-FL |
| KLF10 | -0.58 | -0.60 | AR-V7 | KLHL3 | -1.64 | -0.39 | AR-FL |
| HNF1B | -0.43 | -0.60 | AR-V7 | CLYBL | -1.53 | -0.39 | AR-FL |
| LRRC58 | -0.19 | -0.60 | AR-V7 | IRF6 | -1.01 | -0.39 | AR-FL |
| RGS3 | -0.50 | -0.60 | AR-V7 | ZNF404 | -0.90 | -0.39 | AR-FL |
| NOS1AP | -0.23 | -0.60 | AR-V7 | TOX | -2.26 | -0.39 | AR-FL |
| OVOL1 | -0.53 | -0.60 | AR-V7 | TIGD1 | -0.64 | -0.39 | AR-FL |
| MBOAT7 | -0.29 | -0.60 | AR-V7 | ZCCHC14 | -0.82 | -0.39 | AR-FL |
| IPO5 | -0.17 | -0.60 | AR-V7 | RANBP17 | -0.87 | -0.39 | AR-FL |
| BCAT2 | -0.51 | -0.60 | AR-V7 | NTAN1 | -0.93 | -0.39 | AR-FL |
| VPS54 | -0.34 | -0.60 | AR-V7 | FTO | -0.80 | -0.39 | AR-FL |
| ANGEL1 | -0.29 | -0.59 | AR-V7 | L3HYPDH | -0.94 | -0.39 | AR-FL |
| ZSCAN5A | -0.39 | -0.59 | AR-V7 | ZNF34 | -1.19 | -0.39 | AR-FL |
| MLLT1 | -0.21 | -0.59 | AR-V7 | DTWD1 | -1.03 | -0.39 | AR-FL |
| BCL2L11 | -0.17 | -0.59 | AR-V7 | SLC9A7 | -0.60 | -0.39 | AR-FL |
| AK6 | -0.01 | -0.59 | AR-V7 | ACBD4 | -0.71 | -0.39 | AR-FL |
| CMSS1 | -0.56 | -0.59 | AR-V7 | GPT | -0.79 | -0.39 | AR-FL |
| PAICS | -0.23 | -0.59 | AR-V7 | ACAT1 | -0.59 | -0.38 | AR-FL |
| TAF4B | -0.46 | -0.59 | AR-V7 | VPS13C | -0.67 | -0.38 | AR-FL |
| TMEM159 | -0.52 | -0.58 | AR-V7 | PANK1 | -0.80 | -0.38 | AR-FL |
| PRMT3 | -0.49 | -0.58 | AR-V7 | ELAVL2 | -0.90 | -0.38 | AR-FL |
| PCNT | -0.17 | -0.58 | AR-V7 | SPEF2 | -1.03 | -0.38 | AR-FL |
| CLSTN1 | -0.55 | -0.58 | AR-V7 | SLC19A2 | -1.22 | -0.38 | AR-FL |
| TIMELESS | -0.48 | -0.58 | AR-V7 | ZNF844 | -0.97 | -0.38 | AR-FL |
| TEX30 | -0.42 | -0.58 | AR-V7 | LRRC8C | -1.21 | -0.38 | AR-FL |
| SYNE3 | -0.27 | -0.58 | AR-V7 | EBF1 | -1.20 | -0.38 | AR-FL |
| EIF4EBP3 | -0.24 | -0.58 | AR-V7 | NAB1 | -1.23 | -0.38 | AR-FL |
| DNAJC13 | -0.19 | -0.58 | AR-V7 | HEXDC | -0.75 | -0.38 | AR-FL |
| DNAJC17 | 0.24 | 0.58 | AR-V7 | PNMAL1 | -1.78 | -0.38 | AR-FL |
| WDR13 | 0.34 | 0.58 | AR-V7 | PBLD | -0.92 | -0.38 | AR-FL |
| ZMYND8 | 0.36 | 0.58 | AR-V7 | IFIT5 | -0.58 | -0.38 | AR-FL |
| GPS2 | 0.31 | 0.58 | AR-V7 | ANGEL1 | -0.60 | -0.38 | AR-FL |
| ENDOG | 0.13 | 0.58 | AR-V7 | TTC27 | -0.80 | -0.38 | AR-FL |
| HIC2 | 0.41 | 0.59 | AR-V7 | 1-Sep | -1.00 | -0.38 | AR-FL |

| | | | | | | | |
|---|---|---|---|---|---|---|---|
| CELSR3 | 0.09 | 0.59 | AR-V7 | CCDC85B | -0.77 | -0.38 | AR-FL |
| EIF2B2 | 0.36 | 0.59 | AR-V7 | FSD1L | -0.93 | -0.38 | AR-FL |
| CIITA | 0.06 | 0.59 | AR-V7 | OXCT1 | -0.63 | -0.38 | AR-FL |
| SMN2 | 0.27 | 0.59 | AR-V7 | AHSA2 | -1.06 | -0.38 | AR-FL |
| ANKRD24 | 0.05 | 0.59 | AR-V7 | APAF1 | -0.60 | -0.38 | AR-FL |
| LAMB3 | 0.31 | 0.59 | AR-V7 | ZNF449 | -0.71 | -0.38 | AR-FL |
| HILPDA | 0.29 | 0.59 | AR-V7 | HPS4 | -0.70 | -0.38 | AR-FL |
| HACD4 | 0.10 | 0.59 | AR-V7 | PTCD2 | -1.24 | -0.38 | AR-FL |
| ANKIB1 | 0.18 | 0.59 | AR-V7 | PAK6 | -0.76 | -0.38 | AR-FL |
| WDSUB1 | 0.02 | 0.59 | AR-V7 | KISS1R | -1.14 | -0.38 | AR-FL |
| RP11-468E2.4 | 0.29 | 0.59 | AR-V7 | PLXDC1 | -2.09 | -0.38 | AR-FL |
| FAM63B | 0.31 | 0.59 | AR-V7 | CLUL1 | -0.67 | -0.37 | AR-FL |
| EHF | 0.43 | 0.60 | AR-V7 | SENP8 | -0.67 | -0.37 | AR-FL |
| TRIO | 0.49 | 0.60 | AR-V7 | TMEM69 | -0.62 | -0.37 | AR-FL |
| TUT1 | 0.43 | 0.60 | AR-V7 | WDR17 | -1.53 | -0.37 | AR-FL |
| NUTM2D | 0.05 | 0.60 | AR-V7 | PLXND1 | -1.16 | -0.37 | AR-FL |
| WDR5B | 0.44 | 0.60 | AR-V7 | CLTCL1 | -0.95 | -0.37 | AR-FL |
| SLC36A4 | 0.53 | 0.60 | AR-V7 | TXNIP | -0.86 | -0.37 | AR-FL |
| KIF16B | 0.03 | 0.60 | AR-V7 | EFNA2 | -0.70 | -0.37 | AR-FL |
| VAPA | 0.31 | 0.60 | AR-V7 | FAM161B | -1.27 | -0.37 | AR-FL |
| MED9 | 0.29 | 0.60 | AR-V7 | WWOX | -0.70 | -0.37 | AR-FL |
| ZNF622 | 0.47 | 0.60 | AR-V7 | PLCL2 | -1.09 | -0.37 | AR-FL |
| AHR | 0.13 | 0.60 | AR-V7 | MGEA5 | -0.90 | -0.37 | AR-FL |
| LARP1B | 0.56 | 0.60 | AR-V7 | SULT1A1 | -1.34 | -0.37 | AR-FL |
| LRFN4 | 0.25 | 0.60 | AR-V7 | SLC27A1 | -1.02 | -0.37 | AR-FL |
| MAGED1 | 0.37 | 0.61 | AR-V7 | PYROXD2 | -0.83 | -0.37 | AR-FL |
| NARF | 0.10 | 0.61 | AR-V7 | SMARCD3 | -1.35 | -0.37 | AR-FL |
| MLEC | 0.55 | 0.61 | AR-V7 | DENND6B | -0.95 | -0.37 | AR-FL |
| KCTD7 | 0.26 | 0.61 | AR-V7 | GLTSCR2 | -1.04 | -0.37 | AR-FL |
| HYPK | 0.18 | 0.61 | AR-V7 | LHX4 | -1.00 | -0.37 | AR-FL |
| NAGK | 0.14 | 0.61 | AR-V7 | ERMP1 | -1.17 | -0.37 | AR-FL |
| SLF1 | 0.04 | 0.61 | AR-V7 | QARS | -0.66 | -0.37 | AR-FL |
| DIXDC1 | 0.33 | 0.61 | AR-V7 | CBX7 | -1.10 | -0.37 | AR-FL |
| PNCK | 0.20 | 0.61 | AR-V7 | SUMF1 | -0.59 | -0.37 | AR-FL |
| HIVEP2 | 0.12 | 0.61 | AR-V7 | ETAA1 | -0.73 | -0.37 | AR-FL |
| ARRDC3 | 0.25 | 0.61 | AR-V7 | GEMIN5 | -0.77 | -0.36 | AR-FL |
| NME5 | 0.02 | 0.62 | AR-V7 | ARFGEF1 | -0.65 | -0.36 | AR-FL |
| SLC39A1 | 0.49 | 0.62 | AR-V7 | RNF165 | -0.71 | -0.36 | AR-FL |
| MTMR7 | 0.06 | 0.62 | AR-V7 | MXRA8 | -0.74 | -0.36 | AR-FL |
| DYNC2LI1 | 0.12 | 0.62 | AR-V7 | NCKAP5L | -1.24 | -0.36 | AR-FL |
| CBLB | 0.04 | 0.62 | AR-V7 | SEMA3B | -0.75 | -0.36 | AR-FL |
| DPH3 | 0.43 | 0.62 | AR-V7 | NLGN4Y | -0.66 | -0.36 | AR-FL |
| BRICD5 | 0.15 | 0.62 | AR-V7 | EIF3M | -0.63 | -0.36 | AR-FL |
| CREBRF | 0.23 | 0.62 | AR-V7 | RP5-862P8.2 | -0.71 | -0.36 | AR-FL |
| ZNF701 | 0.37 | 0.62 | AR-V7 | DARS | -0.58 | -0.36 | AR-FL |
| SLC27A3 | 0.06 | 0.62 | AR-V7 | EPHA10 | -1.01 | -0.36 | AR-FL |

| | | | |
|---|---|---|---|
| ENAH | 0.37 | 0.62 | AR-V7 |
| MIIP | 0.36 | 0.62 | AR-V7 |
| FERMT3 | 0.26 | 0.62 | AR-V7 |
| VPS37B | 0.31 | 0.62 | AR-V7 |
| TSPAN6 | 0.24 | 0.62 | AR-V7 |
| MGST3 | 0.41 | 0.63 | AR-V7 |
| KIFAP3 | 0.41 | 0.63 | AR-V7 |
| IL6R | 0.50 | 0.63 | AR-V7 |
| SMTN | 0.32 | 0.63 | AR-V7 |
| ARL1 | 0.52 | 0.63 | AR-V7 |
| IQCB1 | 0.43 | 0.63 | AR-V7 |
| ANKFY1 | 0.58 | 0.63 | AR-V7 |
| C4orf33 | 0.11 | 0.63 | AR-V7 |
| TBC1D20 | 0.23 | 0.63 | AR-V7 |
| ZNF781 | 0.10 | 0.63 | AR-V7 |
| ACBD7 | 0.29 | 0.63 | AR-V7 |
| ASCC1 | 0.22 | 0.64 | AR-V7 |
| C4orf48 | 0.04 | 0.64 | AR-V7 |
| TBC1D10A | 0.50 | 0.64 | AR-V7 |
| PKP2 | 0.51 | 0.64 | AR-V7 |
| SAFB2 | 0.20 | 0.64 | AR-V7 |
| AP4E1 | 0.53 | 0.64 | AR-V7 |
| NDUFV2 | 0.25 | 0.64 | AR-V7 |
| DDTL | 0.51 | 0.64 | AR-V7 |
| CPEB4 | 0.25 | 0.64 | AR-V7 |
| SLC26A6 | 0.21 | 0.64 | AR-V7 |
| MED10 | 0.28 | 0.65 | AR-V7 |
| PSENEN | 0.57 | 0.65 | AR-V7 |
| PRSS8 | 0.05 | 0.65 | AR-V7 |
| SNX1 | 0.17 | 0.65 | AR-V7 |
| SLC52A3 | 0.12 | 0.65 | AR-V7 |
| P4HA1 | 0.07 | 0.65 | AR-V7 |
| ZNF586 | 0.36 | 0.65 | AR-V7 |
| TAX1BP3 | 0.18 | 0.65 | AR-V7 |
| SORT1 | 0.34 | 0.65 | AR-V7 |
| TGFBR3 | 0.11 | 0.65 | AR-V7 |
| C15orf62 | 0.02 | 0.65 | AR-V7 |
| LDLRAD4 | 0.35 | 0.65 | AR-V7 |
| JUN | 0.49 | 0.65 | AR-V7 |
| MAP3K12 | 0.20 | 0.65 | AR-V7 |
| DOK4 | 0.01 | 0.65 | AR-V7 |
| MAP1LC3B | 0.30 | 0.66 | AR-V7 |
| PLXNA3 | 0.28 | 0.66 | AR-V7 |
| HPS3 | 0.11 | 0.66 | AR-V7 |
| DDN | 0.20 | 0.66 | AR-V7 |
| KIAA0040 | 0.33 | 0.66 | AR-V7 |
| MTA3 | -0.60 | -0.36 | AR-FL |
| KCNMB3 | -1.15 | -0.36 | AR-FL |
| SMO | -0.81 | -0.36 | AR-FL |
| GDAP1L1 | -1.61 | -0.36 | AR-FL |
| ZMIZ1 | -0.87 | -0.36 | AR-FL |
| SPIN3 | -0.75 | -0.36 | AR-FL |
| ZBED8 | -0.60 | -0.36 | AR-FL |
| ZNF835 | -1.20 | -0.36 | AR-FL |
| GSTO1 | -0.67 | -0.36 | AR-FL |
| ACADSB | -0.89 | -0.36 | AR-FL |
| ZBTB47 | -0.87 | -0.36 | AR-FL |
| EMX1 | -1.32 | -0.36 | AR-FL |
| ZNF286B | -1.38 | -0.36 | AR-FL |
| DOK1 | -0.63 | -0.36 | AR-FL |
| KBTBD3 | -1.32 | -0.35 | AR-FL |
| C5orf42 | -0.65 | -0.35 | AR-FL |
| ELP4 | -0.62 | -0.35 | AR-FL |
| TNKS | -0.68 | -0.35 | AR-FL |
| CASP8 | -0.69 | -0.35 | AR-FL |
| TGFA | -1.07 | -0.35 | AR-FL |
| SYTL3 | -1.28 | -0.35 | AR-FL |
| MAOA | -1.50 | -0.35 | AR-FL |
| KIAA1107 | -1.49 | -0.35 | AR-FL |
| CERS1 | -1.75 | -0.35 | AR-FL |
| AHNAK2 | -1.16 | -0.35 | AR-FL |
| MRPS27 | -0.67 | -0.35 | AR-FL |
| KLRG2 | -1.30 | -0.35 | AR-FL |
| ULK2 | -0.65 | -0.35 | AR-FL |
| IFFO2 | -0.70 | -0.35 | AR-FL |
| EXT1 | -0.91 | -0.35 | AR-FL |
| ENPP3 | -0.90 | -0.35 | AR-FL |
| C19orf44 | -0.98 | -0.35 | AR-FL |
| HEXIM2 | -0.82 | -0.35 | AR-FL |
| DDX50 | -0.63 | -0.35 | AR-FL |
| CEP85L | -0.87 | -0.35 | AR-FL |
| ALOXE3 | -1.17 | -0.35 | AR-FL |
| INSR | -0.62 | -0.35 | AR-FL |
| CRYZL1 | -0.75 | -0.35 | AR-FL |
| MRRF | -0.59 | -0.35 | AR-FL |
| ZNF846 | -0.82 | -0.35 | AR-FL |
| ARHGAP44 | -0.99 | -0.35 | AR-FL |
| SP4 | -0.79 | -0.34 | AR-FL |
| GNG12 | -0.59 | -0.34 | AR-FL |
| GGT7 | -1.24 | -0.34 | AR-FL |
| DYNC2H1 | -1.75 | -0.34 | AR-FL |
| FAM149A | -1.00 | -0.34 | AR-FL |

| | | | | | | | |
|---|---|---|---|---|---|---|---|
| UBE2V1 | 0.19 | 0.66 | AR-V7 | NAT6 | -0.67 | -0.34 | AR-FL |
| RAMP1 | 0.20 | 0.66 | AR-V7 | BACE1 | -0.95 | -0.34 | AR-FL |
| COMMD6 | 0.32 | 0.66 | AR-V7 | FCHSD2 | -0.76 | -0.34 | AR-FL |
| FAM104A | 0.37 | 0.66 | AR-V7 | TAPT1 | -0.71 | -0.34 | AR-FL |
| DSTN | 0.33 | 0.66 | AR-V7 | ZNF605 | -0.66 | -0.34 | AR-FL |
| HIST1H1B | 0.43 | 0.67 | AR-V7 | THBS2 | -1.17 | -0.34 | AR-FL |
| SYS1 | 0.12 | 0.67 | AR-V7 | RPRD2 | -0.62 | -0.34 | AR-FL |
| DHCR7 | 0.43 | 0.67 | AR-V7 | IL17D | -1.31 | -0.34 | AR-FL |
| HBP1 | 0.55 | 0.67 | AR-V7 | LRBA | -0.59 | -0.34 | AR-FL |
| POFUT2 | 0.18 | 0.67 | AR-V7 | NAPEPLD | -0.92 | -0.34 | AR-FL |
| CHKB | 0.12 | 0.67 | AR-V7 | SLC36A4 | -1.34 | -0.34 | AR-FL |
| CRADD | 0.37 | 0.67 | AR-V7 | MROH6 | -0.74 | -0.34 | AR-FL |
| SPRED3 | 0.25 | 0.68 | AR-V7 | KANSL1 | -0.63 | -0.34 | AR-FL |
| MSN | 0.29 | 0.68 | AR-V7 | WDR74 | -0.77 | -0.34 | AR-FL |
| GRIN2D | 0.04 | 0.68 | AR-V7 | FOXK1 | -0.65 | -0.34 | AR-FL |
| CYP51A1 | 0.40 | 0.68 | AR-V7 | ZNF790 | -0.70 | -0.34 | AR-FL |
| MIER1 | 0.34 | 0.68 | AR-V7 | APBB1 | -1.22 | -0.34 | AR-FL |
| NR6A1 | 0.20 | 0.68 | AR-V7 | BRWD1 | -0.65 | -0.33 | AR-FL |
| OR2A7 | 0.19 | 0.68 | AR-V7 | HEMK1 | -1.06 | -0.33 | AR-FL |
| KCNAB3 | 0.52 | 0.68 | AR-V7 | MTA1 | -0.90 | -0.33 | AR-FL |
| TSSK6 | 0.21 | 0.68 | AR-V7 | GPX7 | -0.95 | -0.33 | AR-FL |
| TMEM234 | 0.18 | 0.68 | AR-V7 | ZNRF3 | -0.72 | -0.33 | AR-FL |
| ADAM10 | 0.22 | 0.68 | AR-V7 | PPIL6 | -1.14 | -0.33 | AR-FL |
| ECM1 | 0.04 | 0.68 | AR-V7 | BCHE | -2.05 | -0.33 | AR-FL |
| SH3BGRL3 | 0.04 | 0.68 | AR-V7 | XDH | -0.64 | -0.33 | AR-FL |
| TUBB2A | 0.05 | 0.68 | AR-V7 | NBPF3 | -0.59 | -0.33 | AR-FL |
| SAXO2 | 0.14 | 0.69 | AR-V7 | KLHL21 | -0.78 | -0.33 | AR-FL |
| NDUFC2 | 0.27 | 0.69 | AR-V7 | ZNF502 | -0.75 | -0.33 | AR-FL |
| HSH2D | 0.48 | 0.69 | AR-V7 | ZNF740 | -0.65 | -0.33 | AR-FL |
| S100A2 | 0.31 | 0.69 | AR-V7 | FBXO46 | -0.73 | -0.33 | AR-FL |
| C10orf10 | 0.20 | 0.69 | AR-V7 | CLUAP1 | -0.95 | -0.33 | AR-FL |
| MSL3 | 0.12 | 0.69 | AR-V7 | ZNF233 | -0.77 | -0.33 | AR-FL |
| CES2 | 0.24 | 0.69 | AR-V7 | PAK3 | -2.03 | -0.33 | AR-FL |
| APAF1 | 0.18 | 0.69 | AR-V7 | PREX2 | -1.02 | -0.33 | AR-FL |
| ARL6IP4 | 0.08 | 0.69 | AR-V7 | KIAA1586 | -0.59 | -0.33 | AR-FL |
| EPOR | 0.29 | 0.69 | AR-V7 | FES | -0.88 | -0.33 | AR-FL |
| GPCPD1 | 0.19 | 0.69 | AR-V7 | RTN1 | -0.59 | -0.33 | AR-FL |
| ESF1 | 0.07 | 0.70 | AR-V7 | SPG20 | -1.40 | -0.33 | AR-FL |
| OSCP1 | 0.29 | 0.70 | AR-V7 | IFT74 | -0.61 | -0.33 | AR-FL |
| ORAI2 | 0.02 | 0.70 | AR-V7 | TET1 | -0.83 | -0.33 | AR-FL |
| YAF2 | 0.16 | 0.70 | AR-V7 | ZNF788 | -0.84 | -0.33 | AR-FL |
| WDR60 | 0.10 | 0.70 | AR-V7 | TSPAN10 | -0.72 | -0.33 | AR-FL |
| CCDC115 | 0.25 | 0.70 | AR-V7 | ZNF45 | -0.59 | -0.33 | AR-FL |
| CBX4 | 0.34 | 0.70 | AR-V7 | P2RY11 | -0.63 | -0.33 | AR-FL |
| TBCEL | 0.45 | 0.70 | AR-V7 | PIGZ | -1.30 | -0.33 | AR-FL |
| SAMD15 | 0.04 | 0.70 | AR-V7 | ZNF554 | -0.94 | -0.32 | AR-FL |

| | | | | | | | |
|---|---|---|---|---|---|---|---|
| ZNF625 | 0.01 | 0.70 | AR-V7 | ENTPD1 | -0.99 | -0.32 | AR-FL |
| SIRT1 | 0.54 | 0.70 | AR-V7 | P3H3 | -1.90 | -0.32 | AR-FL |
| NSUN3 | 0.24 | 0.70 | AR-V7 | XYLB | -0.90 | -0.32 | AR-FL |
| SMPDL3A | 0.44 | 0.70 | AR-V7 | RENBP | -0.61 | -0.32 | AR-FL |
| CFLAR | 0.36 | 0.70 | AR-V7 | ABCD4 | -0.70 | -0.32 | AR-FL |
| HIST1H2BJ | 0.32 | 0.70 | AR-V7 | ANO6 | -0.63 | -0.32 | AR-FL |
| WBP5 | 0.55 | 0.70 | AR-V7 | MSANTD4 | -0.62 | -0.32 | AR-FL |
| UNC13B | 0.25 | 0.70 | AR-V7 | TMEM231 | -1.09 | -0.32 | AR-FL |
| TSPAN31 | 0.07 | 0.71 | AR-V7 | MED13L | -0.65 | -0.32 | AR-FL |
| ARPC1B | 0.52 | 0.71 | AR-V7 | C17orf75 | -0.64 | -0.32 | AR-FL |
| CIB1 | 0.37 | 0.71 | AR-V7 | MAPT | -0.76 | -0.32 | AR-FL |
| ETAA1 | 0.05 | 0.71 | AR-V7 | SFRP4 | -2.16 | -0.32 | AR-FL |
| STX19 | 0.52 | 0.71 | AR-V7 | RPS29 | -0.70 | -0.32 | AR-FL |
| FLVCR2 | 0.02 | 0.71 | AR-V7 | ASTL | -0.79 | -0.32 | AR-FL |
| ZNF281 | 0.16 | 0.71 | AR-V7 | TRAPPC6B | -0.61 | -0.32 | AR-FL |
| COX17 | 0.50 | 0.71 | AR-V7 | SRCIN1 | -1.73 | -0.32 | AR-FL |
| THBS3 | 0.24 | 0.71 | AR-V7 | MTX3 | -0.70 | -0.32 | AR-FL |
| MAFF | 0.57 | 0.71 | AR-V7 | CLDN15 | -1.54 | -0.32 | AR-FL |
| DNAJC12 | 0.43 | 0.72 | AR-V7 | ARHGEF17 | -1.09 | -0.32 | AR-FL |
| MAT1A | 0.39 | 0.72 | AR-V7 | C10orf2 | -0.82 | -0.32 | AR-FL |
| WBP1L | 0.47 | 0.72 | AR-V7 | ERLIN2 | -0.63 | -0.32 | AR-FL |
| NOTCH2 | 0.18 | 0.72 | AR-V7 | MMP24 | -1.51 | -0.32 | AR-FL |
| RP11-505K9.4 | 0.36 | 0.72 | AR-V7 | CPS1 | -1.96 | -0.32 | AR-FL |
| MZT2A | 0.15 | 0.72 | AR-V7 | LRRC3 | -1.14 | -0.32 | AR-FL |
| FRS3 | 0.04 | 0.72 | AR-V7 | GPALPP1 | -0.83 | -0.32 | AR-FL |
| TTC13 | 0.39 | 0.72 | AR-V7 | FBXW8 | -0.78 | -0.32 | AR-FL |
| HOXA1 | 0.26 | 0.72 | AR-V7 | C9orf43 | -0.91 | -0.31 | AR-FL |
| VPS36 | 0.52 | 0.73 | AR-V7 | RSPO4 | -0.84 | -0.31 | AR-FL |
| PARD3 | 0.32 | 0.73 | AR-V7 | RIN3 | -1.40 | -0.31 | AR-FL |
| BTBD8 | 0.18 | 0.73 | AR-V7 | QRICH2 | -1.32 | -0.31 | AR-FL |
| HIST2H4A | 0.11 | 0.73 | AR-V7 | LINGO1 | -0.81 | -0.31 | AR-FL |
| CRY1 | 0.51 | 0.73 | AR-V7 | MACROD1 | -0.84 | -0.31 | AR-FL |
| CHST12 | 0.25 | 0.73 | AR-V7 | OTOGL | -2.35 | -0.31 | AR-FL |
| ADAMTS15 | 0.30 | 0.73 | AR-V7 | RECQL5 | -0.78 | -0.31 | AR-FL |
| ZNF274 | 0.54 | 0.73 | AR-V7 | ENDOV | -0.96 | -0.31 | AR-FL |
| CCDC107 | 0.23 | 0.73 | AR-V7 | SYNE3 | -1.04 | -0.31 | AR-FL |
| PYGO1 | 0.31 | 0.73 | AR-V7 | ANP32B | -0.72 | -0.31 | AR-FL |
| SLC29A2 | 0.56 | 0.74 | AR-V7 | ANKRD24 | -1.40 | -0.31 | AR-FL |
| HIST1H2BC | 0.39 | 0.74 | AR-V7 | NACA | -0.62 | -0.31 | AR-FL |
| KBTBD3 | 0.28 | 0.74 | AR-V7 | CCSAP | -0.81 | -0.31 | AR-FL |
| HDAC8 | 0.42 | 0.74 | AR-V7 | PIK3C3 | -0.59 | -0.31 | AR-FL |
| PPP1R3B | 0.54 | 0.74 | AR-V7 | NLRC3 | -1.06 | -0.31 | AR-FL |
| MEMO1 | 0.42 | 0.74 | AR-V7 | SIRT5 | -0.86 | -0.31 | AR-FL |
| OTUD6B | 0.54 | 0.74 | AR-V7 | MLLT10 | -0.60 | -0.31 | AR-FL |
| RIT1 | 0.57 | 0.74 | AR-V7 | NSA2 | -0.61 | -0.31 | AR-FL |
| DERL3 | 0.57 | 0.74 | AR-V7 | PCBD2 | -0.69 | -0.31 | AR-FL |

| | | | | | | | |
|---|---|---|---|---|---|---|---|
| AOC2 | 0.28 | 0.75 | AR-V7 | USP13 | -0.81 | -0.31 | AR-FL |
| UNC13A | 0.42 | 0.75 | AR-V7 | TUB | -0.62 | -0.31 | AR-FL |
| TRAPPC2P1 | 0.58 | 0.75 | AR-V7 | PAN2 | -0.98 | -0.31 | AR-FL |
| AC004381.6 | 0.51 | 0.75 | AR-V7 | BTBD2 | -0.66 | -0.31 | AR-FL |
| EOGT | 0.15 | 0.75 | AR-V7 | TRIM25 | -0.73 | -0.30 | AR-FL |
| CYB5R2 | 0.20 | 0.75 | AR-V7 | TRPV1 | -1.03 | -0.30 | AR-FL |
| HS1BP3 | 0.47 | 0.75 | AR-V7 | ST6GALNAC2 | -1.26 | -0.30 | AR-FL |
| WDFY1 | 0.47 | 0.75 | AR-V7 | ADGRF5 | -3.32 | -0.30 | AR-FL |
| SH3GLB1 | 0.52 | 0.75 | AR-V7 | PHGDH | -0.67 | -0.30 | AR-FL |
| NXPE3 | 0.46 | 0.75 | AR-V7 | MBNL2 | -0.85 | -0.30 | AR-FL |
| CCDC87 | 0.19 | 0.76 | AR-V7 | ZNF573 | -0.85 | -0.30 | AR-FL |
| ZNF682 | 0.08 | 0.76 | AR-V7 | CRTAP | -0.60 | -0.30 | AR-FL |
| SLC7A8 | 0.57 | 0.76 | AR-V7 | DNPH1 | -0.59 | -0.30 | AR-FL |
| AGFG1 | 0.42 | 0.76 | AR-V7 | C19orf68 | -0.98 | -0.30 | AR-FL |
| WDR78 | 0.08 | 0.76 | AR-V7 | VASH1 | -1.01 | -0.30 | AR-FL |
| CCDC186 | 0.54 | 0.76 | AR-V7 | RNPC3 | -0.76 | -0.30 | AR-FL |
| ZBTB25 | 0.35 | 0.77 | AR-V7 | QTRT1 | -0.61 | -0.30 | AR-FL |
| LRP12 | 0.20 | 0.77 | AR-V7 | NKX2-4 | -1.45 | -0.30 | AR-FL |
| STARD4 | 0.49 | 0.77 | AR-V7 | TFB1M | -0.87 | -0.30 | AR-FL |
| UGDH | 0.52 | 0.77 | AR-V7 | RPGRIP1L | -0.97 | -0.30 | AR-FL |
| LMTK3 | 0.21 | 0.77 | AR-V7 | RPL5 | -0.83 | -0.30 | AR-FL |
| CCDC24 | 0.32 | 0.77 | AR-V7 | ASB13 | -0.75 | -0.30 | AR-FL |
| STOML1 | 0.34 | 0.77 | AR-V7 | USP36 | -0.73 | -0.30 | AR-FL |
| HIST1H2BF | 0.33 | 0.77 | AR-V7 | TMEM182 | -1.15 | -0.30 | AR-FL |
| APLP2 | 0.52 | 0.78 | AR-V7 | NOC3L | -0.75 | -0.30 | AR-FL |
| CCBL1 | 0.47 | 0.78 | AR-V7 | GSTM4 | -1.05 | -0.30 | AR-FL |
| SPSB3 | 0.23 | 0.78 | AR-V7 | CNTFR | -2.07 | -0.30 | AR-FL |
| NICN1 | 0.27 | 0.78 | AR-V7 | GPC4 | -0.72 | -0.30 | AR-FL |
| CHKB-CPT1B | 0.11 | 0.78 | AR-V7 | KSR2 | -0.96 | -0.30 | AR-FL |
| GYG1 | 0.39 | 0.78 | AR-V7 | ZCCHC8 | -0.82 | -0.29 | AR-FL |
| FAS | 0.15 | 0.79 | AR-V7 | MGME1 | -1.25 | -0.29 | AR-FL |
| CUTC | 0.28 | 0.79 | AR-V7 | BBS9 | -0.92 | -0.29 | AR-FL |
| POLI | 0.27 | 0.79 | AR-V7 | ANKS6 | -0.78 | -0.29 | AR-FL |
| GCC2 | 0.42 | 0.79 | AR-V7 | SMC1B | -1.13 | -0.29 | AR-FL |
| ACSS2 | 0.35 | 0.79 | AR-V7 | BRD8 | -0.66 | -0.29 | AR-FL |
| TGS1 | 0.03 | 0.79 | AR-V7 | HNRNPDL | -0.62 | -0.29 | AR-FL |
| MVK | 0.53 | 0.79 | AR-V7 | RASL10B | -3.19 | -0.29 | AR-FL |
| HEPH | 0.44 | 0.80 | AR-V7 | LBHD1 | -0.68 | -0.29 | AR-FL |
| LDHD | 0.40 | 0.81 | AR-V7 | AP4S1 | -1.37 | -0.29 | AR-FL |
| ARG2 | 0.52 | 0.81 | AR-V7 | GSDMB | -1.29 | -0.29 | AR-FL |
| FOSL2 | 0.48 | 0.81 | AR-V7 | SLX4IP | -0.86 | -0.29 | AR-FL |
| HPD | 0.56 | 0.81 | AR-V7 | FOSL2 | -1.34 | -0.29 | AR-FL |
| THOC5 | 0.30 | 0.81 | AR-V7 | SNRPA | -0.62 | -0.29 | AR-FL |
| PTPN18 | 0.43 | 0.81 | AR-V7 | ZFHX2 | -0.89 | -0.29 | AR-FL |
| MYL6 | 0.22 | 0.82 | AR-V7 | TMEM39B | -0.75 | -0.29 | AR-FL |
| CASP10 | 0.28 | 0.82 | AR-V7 | KHK | -1.06 | -0.29 | AR-FL |

| | | | |
|---|---|---|---|
| ZNF165 | 0.41 | 0.82 | AR-V7 |
| PEBP4 | 0.20 | 0.83 | AR-V7 |
| ZNF527 | 0.19 | 0.83 | AR-V7 |
| FOS | 0.38 | 0.84 | AR-V7 |
| SOX15 | 0.06 | 0.85 | AR-V7 |
| C1orf54 | 0.12 | 0.85 | AR-V7 |
| SSX2IP | 0.36 | 0.85 | AR-V7 |
| SLC16A8 | 0.31 | 0.85 | AR-V7 |
| MMP13 | 0.24 | 0.85 | AR-V7 |
| NLGN3 | 0.18 | 0.85 | AR-V7 |
| STAT3 | 0.52 | 0.86 | AR-V7 |
| SRPK2 | 0.22 | 0.86 | AR-V7 |
| BIRC2 | 0.18 | 0.86 | AR-V7 |
| NARFL | 0.34 | 0.86 | AR-V7 |
| SYCE3 | 0.15 | 0.86 | AR-V7 |
| RNF170 | 0.27 | 0.86 | AR-V7 |
| FBLL1 | 0.42 | 0.86 | AR-V7 |
| ASCC3 | 0.30 | 0.86 | AR-V7 |
| RASSF1 | 0.28 | 0.87 | AR-V7 |
| PPP1R15A | 0.37 | 0.87 | AR-V7 |
| EIF2AK4 | 0.22 | 0.87 | AR-V7 |
| MVD | 0.48 | 0.87 | AR-V7 |
| GPR179 | 0.03 | 0.87 | AR-V7 |
| PRRG4 | 0.46 | 0.87 | AR-V7 |
| PCLO | 0.41 | 0.88 | AR-V7 |
| ARAF | 0.25 | 0.88 | AR-V7 |
| HIST1H2AE | 0.39 | 0.88 | AR-V7 |
| MSANTD4 | 0.26 | 0.88 | AR-V7 |
| C1S | 0.08 | 0.88 | AR-V7 |
| TMEM262 | 0.43 | 0.88 | AR-V7 |
| RELB | 0.38 | 0.88 | AR-V7 |
| CAPN9 | 0.24 | 0.88 | AR-V7 |
| KANK1 | 0.27 | 0.89 | AR-V7 |
| RP1-34B20.21 | 0.37 | 0.89 | AR-V7 |
| HS3ST1 | 0.26 | 0.89 | AR-V7 |
| NUDT18 | 0.31 | 0.89 | AR-V7 |
| RHBDL1 | 0.25 | 0.89 | AR-V7 |
| AK8 | 0.06 | 0.90 | AR-V7 |
| SPATA7 | 0.45 | 0.90 | AR-V7 |
| SLC22A31 | 0.44 | 0.90 | AR-V7 |
| C9orf3 | 0.18 | 0.90 | AR-V7 |
| PIGA | 0.53 | 0.90 | AR-V7 |
| GDF15 | 0.04 | 0.90 | AR-V7 |
| MAP6D1 | 0.28 | 0.90 | AR-V7 |
| GALNT2 | 0.43 | 0.90 | AR-V7 |
| TSC2 | 0.41 | 0.91 | AR-V7 |
| METTL12 | -0.58 | -0.29 | AR-FL |
| NPW | -1.03 | -0.29 | AR-FL |
| WDR83 | -0.59 | -0.29 | AR-FL |
| DLL3 | -0.65 | -0.28 | AR-FL |
| DENND3 | -0.70 | -0.28 | AR-FL |
| DOK3 | -0.76 | -0.28 | AR-FL |
| ESD | -0.73 | -0.28 | AR-FL |
| TRMT10B | -0.66 | -0.28 | AR-FL |
| PRKAR1B | -0.88 | -0.28 | AR-FL |
| LOXL2 | -0.94 | -0.28 | AR-FL |
| PROCA1 | -1.15 | -0.28 | AR-FL |
| IFT46 | -0.64 | -0.28 | AR-FL |
| CEP57 | -1.10 | -0.28 | AR-FL |
| RP11-511P7.5 | -0.60 | -0.28 | AR-FL |
| IPO5 | -0.72 | -0.28 | AR-FL |
| C19orf66 | -0.78 | -0.28 | AR-FL |
| PLIN4 | -0.83 | -0.28 | AR-FL |
| PLCD3 | -0.89 | -0.28 | AR-FL |
| THBS3 | -1.22 | -0.28 | AR-FL |
| ZDHHC11B | -1.75 | -0.28 | AR-FL |
| SULT4A1 | -1.39 | -0.28 | AR-FL |
| DLL1 | -1.26 | -0.28 | AR-FL |
| ARSJ | -1.06 | -0.28 | AR-FL |
| BOD1 | -0.66 | -0.28 | AR-FL |
| PRNP | -0.78 | -0.28 | AR-FL |
| CRYM | -1.88 | -0.28 | AR-FL |
| PARS2 | -0.77 | -0.28 | AR-FL |
| LTB4R | -1.38 | -0.28 | AR-FL |
| TCN2 | -0.82 | -0.28 | AR-FL |
| ZNF225 | -0.61 | -0.27 | AR-FL |
| RABGAP1L | -0.80 | -0.27 | AR-FL |
| ENOSF1 | -1.08 | -0.27 | AR-FL |
| FKBP10 | -0.79 | -0.27 | AR-FL |
| FBXO41 | -0.67 | -0.27 | AR-FL |
| AMER1 | -0.78 | -0.27 | AR-FL |
| NBPF11 | -1.22 | -0.27 | AR-FL |
| RIMS4 | -1.38 | -0.27 | AR-FL |
| ZNF221 | -1.46 | -0.27 | AR-FL |
| PITPNM2 | -1.43 | -0.27 | AR-FL |
| DAB1 | -4.09 | -0.27 | AR-FL |
| NEU3 | -1.01 | -0.27 | AR-FL |
| FAM135B | -2.80 | -0.27 | AR-FL |
| DUSP19 | -0.89 | -0.27 | AR-FL |
| EIF3L | -0.85 | -0.27 | AR-FL |
| DLX4 | -1.44 | -0.27 | AR-FL |
| EEF2 | -0.77 | -0.27 | AR-FL |

| | | | |
|---|---|---|---|
| TXNL4B | 0.41 | 0.91 | AR-V7 |
| PEAK1 | 0.20 | 0.91 | AR-V7 |
| ASIC3 | 0.04 | 0.92 | AR-V7 |
| CKAP2 | 0.15 | 0.92 | AR-V7 |
| FGF22 | 0.10 | 0.92 | AR-V7 |
| ASPG | 0.27 | 0.92 | AR-V7 |
| BZW1 | 0.58 | 0.93 | AR-V7 |
| FAM166B | 0.48 | 0.93 | AR-V7 |
| KIAA1161 | 0.11 | 0.93 | AR-V7 |
| AMT | 0.03 | 0.93 | AR-V7 |
| PLEKHG3 | 0.31 | 0.94 | AR-V7 |
| CCDC82 | 0.31 | 0.94 | AR-V7 |
| GPLD1 | 0.19 | 0.94 | AR-V7 |
| EPHX2 | 0.36 | 0.94 | AR-V7 |
| VWA8 | 0.12 | 0.94 | AR-V7 |
| GDF9 | 0.39 | 0.94 | AR-V7 |
| SPATA17 | 0.20 | 0.94 | AR-V7 |
| AOC3 | 0.10 | 0.95 | AR-V7 |
| THSD4 | 0.30 | 0.95 | AR-V7 |
| DCAF4L1 | 0.27 | 0.96 | AR-V7 |
| SLC22A18 | 0.36 | 0.96 | AR-V7 |
| PIP5KL1 | 0.22 | 0.96 | AR-V7 |
| MAP4K5 | 0.47 | 0.96 | AR-V7 |
| COL7A1 | 0.12 | 0.96 | AR-V7 |
| PQLC1 | 0.07 | 0.96 | AR-V7 |
| XKR9 | 0.44 | 0.96 | AR-V7 |
| BDH2 | 0.06 | 0.97 | AR-V7 |
| GALR3 | 0.49 | 0.98 | AR-V7 |
| CRIM1 | 0.19 | 0.98 | AR-V7 |
| LIMS1 | 0.54 | 0.98 | AR-V7 |
| IFI27 | 0.10 | 0.98 | AR-V7 |
| TRIP10 | 0.33 | 0.99 | AR-V7 |
| C22orf15 | 0.37 | 0.99 | AR-V7 |
| DUSP15 | 0.11 | 0.99 | AR-V7 |
| ADAMTSL3 | 0.35 | 0.99 | AR-V7 |
| PEG10 | 0.37 | 0.99 | AR-V7 |
| ZNF783 | 0.08 | 0.99 | AR-V7 |
| SLC22A4 | 0.44 | 1.00 | AR-V7 |
| PEX2 | 0.51 | 1.00 | AR-V7 |
| RWDD3 | 0.07 | 1.00 | AR-V7 |
| DNAJB5 | 0.55 | 1.00 | AR-V7 |
| EVA1C | 0.36 | 1.00 | AR-V7 |
| FAM129B | 0.30 | 1.00 | AR-V7 |
| TP53TG5 | 0.23 | 1.00 | AR-V7 |
| CETN2 | 0.33 | 1.01 | AR-V7 |
| LZTFL1 | 0.38 | 1.01 | AR-V7 |
| LRRC49 | -0.87 | -0.27 | AR-FL |
| RELN | -1.54 | -0.27 | AR-FL |
| WNT10B | -2.01 | -0.27 | AR-FL |
| ALPK1 | -0.69 | -0.27 | AR-FL |
| RNF8 | -0.60 | -0.27 | AR-FL |
| C5orf38 | -0.83 | -0.26 | AR-FL |
| TMEM25 | -0.68 | -0.26 | AR-FL |
| CEND1 | -0.66 | -0.26 | AR-FL |
| FBXO44 | -0.77 | -0.26 | AR-FL |
| ARHGEF40 | -1.61 | -0.26 | AR-FL |
| TCAP | -0.62 | -0.26 | AR-FL |
| STXBP6 | -0.96 | -0.26 | AR-FL |
| CMSS1 | -0.75 | -0.26 | AR-FL |
| DHRS12 | -0.62 | -0.26 | AR-FL |
| RPL15 | -0.73 | -0.26 | AR-FL |
| PERM1 | -0.70 | -0.26 | AR-FL |
| GNB2L1 | -0.77 | -0.26 | AR-FL |
| STAC3 | -1.21 | -0.26 | AR-FL |
| TPBGL | -2.39 | -0.26 | AR-FL |
| MPP6 | -0.72 | -0.26 | AR-FL |
| MTAP | -0.84 | -0.26 | AR-FL |
| BTF3 | -0.61 | -0.26 | AR-FL |
| ARID4A | -0.62 | -0.26 | AR-FL |
| FAN1 | -0.62 | -0.26 | AR-FL |
| TRAPPC6A | -0.74 | -0.26 | AR-FL |
| KLF2 | -0.93 | -0.26 | AR-FL |
| AATK | -1.27 | -0.26 | AR-FL |
| TMEM18 | -0.70 | -0.26 | AR-FL |
| LYSMD1 | -0.64 | -0.26 | AR-FL |
| CD274 | -1.50 | -0.26 | AR-FL |
| TXLNG | -0.58 | -0.26 | AR-FL |
| GNG7 | -1.24 | -0.26 | AR-FL |
| TPCN1 | -0.60 | -0.26 | AR-FL |
| PPIL3 | -0.69 | -0.26 | AR-FL |
| CDC25C | -0.63 | -0.26 | AR-FL |
| BCORL1 | -1.20 | -0.26 | AR-FL |
| EIF3F | -0.63 | -0.25 | AR-FL |
| ESYT1 | -0.73 | -0.25 | AR-FL |
| APOC1 | -0.93 | -0.25 | AR-FL |
| HOXB7 | -0.80 | -0.25 | AR-FL |
| C5orf30 | -0.74 | -0.25 | AR-FL |
| TRNAU1AP | -0.73 | -0.25 | AR-FL |
| WDR31 | -1.25 | -0.25 | AR-FL |
| MLLT6 | -0.74 | -0.25 | AR-FL |
| ASIC1 | -0.64 | -0.25 | AR-FL |
| TTYH2 | -1.65 | -0.25 | AR-FL |

| | | | | | | | |
|---|---|---|---|---|---|---|---|
| KIAA1109 | 0.24 | 1.01 | AR-V7 | DHFRL1 | -0.81 | -0.25 | AR-FL |
| SUCO | 0.54 | 1.01 | AR-V7 | RORA | -0.86 | -0.25 | AR-FL |
| TEX14 | 0.05 | 1.01 | AR-V7 | ITSN1 | -0.70 | -0.25 | AR-FL |
| ZNF454 | 0.09 | 1.01 | AR-V7 | PDK2 | -0.87 | -0.25 | AR-FL |
| SPRY4 | 0.55 | 1.02 | AR-V7 | ACAD10 | -1.02 | -0.25 | AR-FL |
| DUSP1 | 0.35 | 1.02 | AR-V7 | GRAMD4 | -1.01 | -0.25 | AR-FL |
| P4HA2 | 0.20 | 1.02 | AR-V7 | PALB2 | -0.62 | -0.25 | AR-FL |
| MYH9 | 0.50 | 1.02 | AR-V7 | NUFIP1 | -0.75 | -0.25 | AR-FL |
| CD24 | 0.11 | 1.02 | AR-V7 | CLK4 | -0.90 | -0.25 | AR-FL |
| SELM | 0.28 | 1.02 | AR-V7 | FAM184B | -0.85 | -0.25 | AR-FL |
| RAB20 | 0.44 | 1.03 | AR-V7 | USP11 | -0.84 | -0.25 | AR-FL |
| ITGA10 | 0.44 | 1.04 | AR-V7 | B4GALNT4 | -0.80 | -0.24 | AR-FL |
| ZNF501 | 0.29 | 1.04 | AR-V7 | CNNM1 | -0.92 | -0.24 | AR-FL |
| FAXDC2 | 0.09 | 1.05 | AR-V7 | IGSF22 | -0.71 | -0.24 | AR-FL |
| SMCHD1 | 0.34 | 1.06 | AR-V7 | AARSD1 | -0.72 | -0.24 | AR-FL |
| IRF7 | 0.28 | 1.06 | AR-V7 | NFKBIZ | -1.63 | -0.24 | AR-FL |
| ITGA7 | 0.15 | 1.07 | AR-V7 | DNMBP | -0.62 | -0.24 | AR-FL |
| SECTM1 | 0.00 | 1.07 | AR-V7 | WIPF3 | -1.16 | -0.24 | AR-FL |
| TLCD2 | 0.49 | 1.07 | AR-V7 | PGLS | -0.59 | -0.24 | AR-FL |
| WFDC3 | 0.27 | 1.08 | AR-V7 | AL589743.1 | -0.96 | -0.24 | AR-FL |
| LRGUK | 0.14 | 1.08 | AR-V7 | VPS13A | -0.72 | -0.24 | AR-FL |
| STX18 | 0.54 | 1.08 | AR-V7 | COL18A1 | -1.03 | -0.24 | AR-FL |
| CSRNP1 | 0.50 | 1.08 | AR-V7 | FHL1 | -2.73 | -0.24 | AR-FL |
| SMPD3 | 0.41 | 1.09 | AR-V7 | EIF3H | -0.64 | -0.24 | AR-FL |
| RAB32 | 0.43 | 1.09 | AR-V7 | RNASET2 | -1.23 | -0.24 | AR-FL |
| PPP2R3A | 0.30 | 1.09 | AR-V7 | TSGA10 | -1.09 | -0.24 | AR-FL |
| ZNF547 | 0.36 | 1.09 | AR-V7 | DENND5B | -1.16 | -0.24 | AR-FL |
| MFSD7 | 0.11 | 1.10 | AR-V7 | PPA1 | -0.76 | -0.24 | AR-FL |
| MFAP4 | 0.39 | 1.10 | AR-V7 | TRMT1 | -0.73 | -0.24 | AR-FL |
| MFSD6 | 0.15 | 1.10 | AR-V7 | TTLL3 | -1.11 | -0.24 | AR-FL |
| SMIM1 | 0.47 | 1.12 | AR-V7 | WBSCR27 | -1.02 | -0.24 | AR-FL |
| ZNF135 | 0.06 | 1.12 | AR-V7 | CCDC74B | -0.76 | -0.23 | AR-FL |
| AMOTL2 | 0.43 | 1.12 | AR-V7 | TSPAN3 | -0.64 | -0.23 | AR-FL |
| NCOR1 | 0.40 | 1.12 | AR-V7 | C1orf53 | -0.81 | -0.23 | AR-FL |
| ENG | 0.22 | 1.12 | AR-V7 | GAA | -0.67 | -0.23 | AR-FL |
| LMO7 | 0.56 | 1.13 | AR-V7 | CAMKV | -1.75 | -0.23 | AR-FL |
| RGPD2 | 0.31 | 1.14 | AR-V7 | WDR19 | -1.48 | -0.23 | AR-FL |
| PIH1D2 | 0.24 | 1.14 | AR-V7 | PIM2 | -0.89 | -0.23 | AR-FL |
| TYMP | 0.45 | 1.15 | AR-V7 | KLHL15 | -0.99 | -0.23 | AR-FL |
| HS6ST1 | 0.07 | 1.15 | AR-V7 | MDN1 | -0.60 | -0.23 | AR-FL |
| UBE2E3 | 0.14 | 1.15 | AR-V7 | SCNN1D | -1.24 | -0.23 | AR-FL |
| GGN | 0.37 | 1.16 | AR-V7 | SMYD2 | -0.63 | -0.23 | AR-FL |
| PRR15L | 0.19 | 1.16 | AR-V7 | GP1BA | -0.82 | -0.23 | AR-FL |
| GHDC | 0.52 | 1.17 | AR-V7 | TARBP1 | -0.87 | -0.23 | AR-FL |
| USP43 | 0.32 | 1.17 | AR-V7 | RPL4 | -0.76 | -0.23 | AR-FL |
| CLEC16A | 0.24 | 1.17 | AR-V7 | TMEM64 | -0.67 | -0.23 | AR-FL |

| | | | | | | | |
|---|---|---|---|---|---|---|---|
| SNX33 | 0.34 | 1.17 | AR-V7 | THEM4 | -0.64 | -0.22 | AR-FL |
| DDX53 | 0.10 | 1.17 | AR-V7 | WDR91 | -1.08 | -0.22 | AR-FL |
| ASB3 | 0.34 | 1.18 | AR-V7 | HEATR1 | -0.62 | -0.22 | AR-FL |
| DNAJB4 | 0.28 | 1.20 | AR-V7 | MATK | -1.44 | -0.22 | AR-FL |
| DRC3 | 0.33 | 1.20 | AR-V7 | OSBPL1A | -0.70 | -0.22 | AR-FL |
| CBWD5 | 0.43 | 1.21 | AR-V7 | C2CD4C | -0.99 | -0.22 | AR-FL |
| RP11-2C24.9 | 0.55 | 1.22 | AR-V7 | WDR27 | -1.06 | -0.22 | AR-FL |
| HELZ2 | 0.22 | 1.22 | AR-V7 | NUDT3 | -0.59 | -0.22 | AR-FL |
| COX7B2 | 0.18 | 1.22 | AR-V7 | MEST | -0.73 | -0.22 | AR-FL |
| ADAMTS20 | 0.00 | 1.22 | AR-V7 | FSCN2 | -0.73 | -0.22 | AR-FL |
| HIST1H1C | 0.46 | 1.22 | AR-V7 | RPS6KA5 | -1.00 | -0.22 | AR-FL |
| SLC47A2 | 0.36 | 1.23 | AR-V7 | PFAS | -0.88 | -0.22 | AR-FL |
| ZBED3 | 0.28 | 1.24 | AR-V7 | RASGRF1 | -0.85 | -0.22 | AR-FL |
| MFSD2A | 0.30 | 1.24 | AR-V7 | RPS4Y1 | -0.73 | -0.22 | AR-FL |
| MAP3K14 | 0.20 | 1.24 | AR-V7 | EMILIN3 | -2.11 | -0.22 | AR-FL |
| POF1B | 0.46 | 1.25 | AR-V7 | PTPRS | -0.63 | -0.22 | AR-FL |
| H2AFZ | 0.19 | 1.25 | AR-V7 | PER3 | -0.90 | -0.22 | AR-FL |
| HYDIN | 0.12 | 1.26 | AR-V7 | LRRC37A3 | -1.28 | -0.21 | AR-FL |
| C16orf86 | 0.48 | 1.26 | AR-V7 | LTB4R2 | -1.10 | -0.21 | AR-FL |
| BCL6 | 0.52 | 1.26 | AR-V7 | HADH | -0.62 | -0.21 | AR-FL |
| C4orf47 | 0.38 | 1.27 | AR-V7 | S100A13 | -0.84 | -0.21 | AR-FL |
| DNAH6 | 0.04 | 1.27 | AR-V7 | NOD1 | -1.22 | -0.21 | AR-FL |
| ERAP2 | 0.17 | 1.27 | AR-V7 | SFMBT2 | -1.23 | -0.21 | AR-FL |
| TFEB | 0.56 | 1.28 | AR-V7 | ZNF141 | -0.66 | -0.21 | AR-FL |
| PTK2B | 0.34 | 1.28 | AR-V7 | C20orf27 | -1.01 | -0.21 | AR-FL |
| AP3S1 | 0.18 | 1.32 | AR-V7 | ZNF585B | -0.59 | -0.21 | AR-FL |
| FAM188B | 0.23 | 1.34 | AR-V7 | EPHX4 | -1.87 | -0.21 | AR-FL |
| ZMYND10 | 0.09 | 1.34 | AR-V7 | RAD52 | -0.69 | -0.21 | AR-FL |
| TUBB3 | 0.12 | 1.34 | AR-V7 | KCTD15 | -1.01 | -0.21 | AR-FL |
| TBC1D8B | 0.45 | 1.34 | AR-V7 | RPL12 | -0.89 | -0.20 | AR-FL |
| TMEM47 | 0.51 | 1.35 | AR-V7 | MYL9 | -1.10 | -0.20 | AR-FL |
| MARS2 | 0.50 | 1.36 | AR-V7 | ATP9A | -0.76 | -0.20 | AR-FL |
| PGAP2 | 0.36 | 1.36 | AR-V7 | SFI1 | -1.16 | -0.20 | AR-FL |
| DUSP5 | 0.02 | 1.38 | AR-V7 | CSPP1 | -0.67 | -0.20 | AR-FL |
| DHRS3 | 0.33 | 1.38 | AR-V7 | LYRM7 | -0.65 | -0.20 | AR-FL |
| CMTM8 | 0.52 | 1.39 | AR-V7 | HOXA2 | -1.09 | -0.20 | AR-FL |
| VIM | 0.47 | 1.40 | AR-V7 | TBKBP1 | -0.68 | -0.20 | AR-FL |
| TACC2 | 0.35 | 1.40 | AR-V7 | TPR | -0.62 | -0.20 | AR-FL |
| GCLC | 0.27 | 1.40 | AR-V7 | ZNF780B | -0.59 | -0.20 | AR-FL |
| SLC25A18 | 0.11 | 1.46 | AR-V7 | RPS25 | -0.62 | -0.20 | AR-FL |
| PSTPIP2 | 0.19 | 1.49 | AR-V7 | RPS3 | -0.73 | -0.20 | AR-FL |
| NREP | 0.19 | 1.49 | AR-V7 | PLPP3 | -0.97 | -0.20 | AR-FL |
| ARSA | 0.36 | 1.51 | AR-V7 | IPO4 | -0.73 | -0.20 | AR-FL |
| IZUMO4 | 0.50 | 1.51 | AR-V7 | ECHDC2 | -0.60 | -0.20 | AR-FL |
| BBOF1 | 0.36 | 1.51 | AR-V7 | ICA1L | -0.68 | -0.20 | AR-FL |
| HSPB1 | 0.00 | 1.52 | AR-V7 | TDRD6 | -1.90 | -0.20 | AR-FL |

| | | | |
|---|---|---|---|
| GAB2 | 0.41 | 1.53 | AR-V7 |
| FLNA | 0.45 | 1.54 | AR-V7 |
| ZNF728 | 0.09 | 1.54 | AR-V7 |
| ALDH1A2 | 0.44 | 1.54 | AR-V7 |
| NFIL3 | 0.07 | 1.54 | AR-V7 |
| NRBP2 | 0.02 | 1.55 | AR-V7 |
| IGF2BP3 | 0.33 | 1.56 | AR-V7 |
| ZNF189 | 0.25 | 1.58 | AR-V7 |
| CERS6 | 0.47 | 1.60 | AR-V7 |
| TMEM60 | 0.18 | 1.61 | AR-V7 |
| CLU | 0.17 | 1.61 | AR-V7 |
| TNS2 | 0.01 | 1.62 | AR-V7 |
| CEP120 | 0.21 | 1.62 | AR-V7 |
| CHPT1 | 0.19 | 1.65 | AR-V7 |
| SNX22 | 0.29 | 1.65 | AR-V7 |
| IL17B | 0.10 | 1.65 | AR-V7 |
| SMOX | 0.17 | 1.65 | AR-V7 |
| KLHL31 | 0.45 | 1.66 | AR-V7 |
| CHRM3 | 0.22 | 1.68 | AR-V7 |
| PLAGL1 | 0.01 | 1.68 | AR-V7 |
| KCNH2 | 0.34 | 1.69 | AR-V7 |
| UPP1 | 0.24 | 1.70 | AR-V7 |
| S100P | 0.45 | 1.70 | AR-V7 |
| CNTD1 | 0.34 | 1.72 | AR-V7 |
| RAG1 | 0.06 | 1.73 | AR-V7 |
| TASP1 | 0.41 | 1.77 | AR-V7 |
| CRYM | 0.40 | 1.78 | AR-V7 |
| GBE1 | 0.17 | 1.78 | AR-V7 |
| VPS33B | 0.25 | 1.87 | AR-V7 |
| SPEF1 | 0.39 | 1.88 | AR-V7 |
| SLC22A1 | 0.44 | 1.88 | AR-V7 |
| SPATA4 | 0.39 | 1.89 | AR-V7 |
| CCDC108 | 0.55 | 1.92 | AR-V7 |
| MSX1 | 0.25 | 1.95 | AR-V7 |
| HIST1H2BG | 0.19 | 1.96 | AR-V7 |
| RAET1L | 0.43 | 1.96 | AR-V7 |
| FAM43A | 0.32 | 1.97 | AR-V7 |
| TMSB10 | 0.17 | 1.99 | AR-V7 |
| KCND1 | 0.37 | 2.03 | AR-V7 |
| XDH | 0.41 | 2.04 | AR-V7 |
| MAP1LC3A | 0.27 | 2.06 | AR-V7 |
| FAM151A | 0.25 | 2.07 | AR-V7 |
| TMEM56 | 0.56 | 2.08 | AR-V7 |
| GRIN2C | 0.14 | 2.09 | AR-V7 |
| ARHGEF28 | 0.20 | 2.09 | AR-V7 |
| IFI44 | 0.42 | 2.12 | AR-V7 |

| | | | |
|---|---|---|---|
| TMC4 | -0.69 | -0.20 | AR-FL |
| TMEM121 | -1.03 | -0.20 | AR-FL |
| ANKS1B | -1.42 | -0.20 | AR-FL |
| CCDC150 | -0.94 | -0.20 | AR-FL |
| TDRD10 | -0.81 | -0.20 | AR-FL |
| LRP5L | -0.58 | -0.19 | AR-FL |
| FBXO48 | -0.69 | -0.19 | AR-FL |
| ENPP1 | -0.97 | -0.19 | AR-FL |
| ENGASE | -0.80 | -0.19 | AR-FL |
| NUDT13 | -0.67 | -0.19 | AR-FL |
| FAHD2B | -1.03 | -0.19 | AR-FL |
| GEMIN2 | -0.83 | -0.19 | AR-FL |
| CYFIP2 | -0.83 | -0.19 | AR-FL |
| BMP8B | -0.90 | -0.19 | AR-FL |
| ZNF619 | -0.59 | -0.19 | AR-FL |
| FAM185A | -0.63 | -0.19 | AR-FL |
| CACNB1 | -1.55 | -0.19 | AR-FL |
| C2orf74 | -0.81 | -0.19 | AR-FL |
| TMCC2 | -0.69 | -0.19 | AR-FL |
| MYL5 | -1.22 | -0.19 | AR-FL |
| ABCC5 | -0.74 | -0.19 | AR-FL |
| CLMN | -1.22 | -0.19 | AR-FL |
| GEMIN6 | -0.58 | -0.18 | AR-FL |
| ANKRD61 | -0.74 | -0.18 | AR-FL |
| TCEA2 | -1.00 | -0.18 | AR-FL |
| RPL27A | -0.61 | -0.18 | AR-FL |
| TNFRSF21 | -0.81 | -0.18 | AR-FL |
| C1GALT1 | -0.86 | -0.18 | AR-FL |
| MMRN2 | -1.07 | -0.18 | AR-FL |
| RPL34 | -0.61 | -0.18 | AR-FL |
| SYNGR3 | -1.16 | -0.18 | AR-FL |
| L3MBTL1 | -1.47 | -0.18 | AR-FL |
| ANKRD18B | -0.90 | -0.18 | AR-FL |
| OSGEPL1 | -0.73 | -0.18 | AR-FL |
| CFAP70 | -0.79 | -0.17 | AR-FL |
| TBC1D10C | -1.23 | -0.17 | AR-FL |
| TMEM43 | -0.62 | -0.17 | AR-FL |
| INPP5B | -0.98 | -0.17 | AR-FL |
| ERCC6 | -0.65 | -0.17 | AR-FL |
| RPS15A | -0.72 | -0.17 | AR-FL |
| CYP2D6 | -0.70 | -0.17 | AR-FL |
| CEP250 | -0.58 | -0.17 | AR-FL |
| HNF1A | -0.65 | -0.17 | AR-FL |
| MKRN2OS | -0.61 | -0.17 | AR-FL |
| HSF4 | -0.60 | -0.17 | AR-FL |
| SLC44A3 | -0.59 | -0.17 | AR-FL |

| | | | |
|---|---|---|---|
| SRD5A1 | 0.23 | 2.13 | AR-V7 |
| EPHA7 | 0.52 | 2.19 | AR-V7 |
| FCHSD1 | 0.35 | 2.29 | AR-V7 |
| VILL | 0.21 | 2.29 | AR-V7 |
| PPM1B | 0.54 | 2.33 | AR-V7 |
| ALOX15B | 0.09 | 2.36 | AR-V7 |
| NAT2 | 0.25 | 2.39 | AR-V7 |
| DUOX1 | 0.52 | 2.42 | AR-V7 |
| SCIN | 0.30 | 2.46 | AR-V7 |
| ANXA1 | 0.03 | 2.47 | AR-V7 |
| STMN4 | 0.46 | 2.53 | AR-V7 |
| SLC45A1 | 0.19 | 2.57 | AR-V7 |
| ZNF560 | 0.15 | 2.57 | AR-V7 |
| TBX10 | 0.51 | 2.62 | AR-V7 |
| GLIPR1 | 0.43 | 2.67 | AR-V7 |
| CYP26B1 | 0.10 | 2.73 | AR-V7 |
| EPHA6 | 0.08 | 2.76 | AR-V7 |
| GCH1 | 0.18 | 2.90 | AR-V7 |
| GPR153 | 0.55 | 3.00 | AR-V7 |
| MYO1G | 0.04 | 3.07 | AR-V7 |
| MATN3 | 0.33 | 3.10 | AR-V7 |
| JPH4 | 0.31 | 3.19 | AR-V7 |
| TMEM169 | 0.51 | 3.28 | AR-V7 |
| MXRA7 | 0.28 | 3.35 | AR-V7 |
| ABLIM3 | 0.23 | 3.45 | AR-V7 |
| HRCT1 | 0.56 | 3.57 | AR-V7 |
| ETNK2 | 0.35 | 3.61 | AR-V7 |
| DDX4 | 0.48 | 3.75 | AR-V7 |
| TDRD9 | 0.54 | 3.90 | AR-V7 |
| WISP2 | 0.32 | 4.07 | AR-V7 |
| ITIH3 | 0.23 | 4.54 | AR-V7 |
| GNAI1 | 0.39 | 4.75 | AR-V7 |
| LAPTM5 | 0.36 | 4.75 | AR-V7 |
| TRPC3 | 0.43 | 4.77 | AR-V7 |
| MAGEA4 | 0.04 | 4.81 | AR-V7 |
| SKIV2L | 0.36 | 4.82 | AR-V7 |
| RALYL | 0.36 | 5.62 | AR-V7 |
| ABHD12 | 0.61 | 0.00 | AR-FL |
| B3GAT1 | 1.16 | 0.00 | AR-FL |
| CHPF | 0.89 | 0.00 | AR-FL |
| TBC1D30 | 0.95 | 0.01 | AR-FL |
| LYPLA2 | 0.64 | 0.01 | AR-FL |
| HIVEP1 | 0.72 | 0.01 | AR-FL |
| ERBB2IP | 0.59 | 0.01 | AR-FL |
| PLCH2 | 0.80 | 0.01 | AR-FL |
| PPP3CC | 0.59 | 0.01 | AR-FL |
| RPL23 | -0.65 | -0.17 | AR-FL |
| CPQ | -0.96 | -0.17 | AR-FL |
| RPS24 | -0.64 | -0.17 | AR-FL |
| GNB5 | -0.80 | -0.17 | AR-FL |
| PSMG4 | -0.71 | -0.16 | AR-FL |
| TIAF1 | -0.65 | -0.16 | AR-FL |
| PMP22 | -1.59 | -0.16 | AR-FL |
| AZIN2 | -1.03 | -0.16 | AR-FL |
| GPM6B | -0.58 | -0.16 | AR-FL |
| BMP4 | -1.15 | -0.16 | AR-FL |
| TUBB4A | -0.72 | -0.16 | AR-FL |
| SFR1 | -0.78 | -0.16 | AR-FL |
| COL6A2 | -0.92 | -0.16 | AR-FL |
| RPL14 | -0.72 | -0.16 | AR-FL |
| HMP19 | -1.02 | -0.16 | AR-FL |
| DDX47 | -0.73 | -0.16 | AR-FL |
| ADGRB1 | -1.07 | -0.16 | AR-FL |
| RPLP0 | -0.65 | -0.16 | AR-FL |
| RPL13A | -0.72 | -0.16 | AR-FL |
| SH3BGRL | -0.72 | -0.16 | AR-FL |
| VSTM2L | -0.93 | -0.16 | AR-FL |
| JUN | -1.00 | -0.16 | AR-FL |
| FAM132B | -1.36 | -0.16 | AR-FL |
| MAP3K1 | -0.86 | -0.15 | AR-FL |
| MICALCL | -0.59 | -0.15 | AR-FL |
| SP5 | -1.06 | -0.15 | AR-FL |
| GPLD1 | -1.28 | -0.15 | AR-FL |
| RPL3 | -0.81 | -0.15 | AR-FL |
| INO80B | -0.72 | -0.15 | AR-FL |
| NRP2 | -1.02 | -0.15 | AR-FL |
| INVS | -0.68 | -0.15 | AR-FL |
| ATHL1 | -0.61 | -0.15 | AR-FL |
| RPL10A | -0.67 | -0.15 | AR-FL |
| CD47 | -0.70 | -0.15 | AR-FL |
| DCAF16 | -0.63 | -0.15 | AR-FL |
| PODXL | -0.86 | -0.15 | AR-FL |
| SIPA1L2 | -0.97 | -0.15 | AR-FL |
| ZFYVE28 | -1.06 | -0.15 | AR-FL |
| RPS2 | -0.66 | -0.15 | AR-FL |
| ANKRD13A | -0.75 | -0.15 | AR-FL |
| FBL | -0.67 | -0.15 | AR-FL |
| ACTR3B | -0.63 | -0.14 | AR-FL |
| ZNF10 | -0.66 | -0.14 | AR-FL |
| AMT | -1.63 | -0.14 | AR-FL |
| ZNF551 | -0.71 | -0.14 | AR-FL |
| MBNL3 | -0.72 | -0.14 | AR-FL |

| | | | | | | | |
|---|---|---|---|---|---|---|---|
| ADPGK | 0.73 | 0.01 | AR-FL | RPL7A | -0.64 | -0.14 | AR-FL |
| UCHL1 | 1.13 | 0.01 | AR-FL | NME7 | -0.61 | -0.14 | AR-FL |
| G3BP2 | 0.77 | 0.02 | AR-FL | FUT10 | -0.63 | -0.14 | AR-FL |
| TAOK3 | 1.62 | 0.02 | AR-FL | NOXA1 | -1.01 | -0.14 | AR-FL |
| ASPHD2 | 0.96 | 0.02 | AR-FL | CUL4B | -0.73 | -0.14 | AR-FL |
| SEC14L2 | 2.43 | 0.02 | AR-FL | FAM78A | -0.90 | -0.14 | AR-FL |
| TFPI | 0.59 | 0.03 | AR-FL | TENM2 | -1.83 | -0.14 | AR-FL |
| TMEM79 | 2.36 | 0.03 | AR-FL | KCNAB3 | -1.50 | -0.14 | AR-FL |
| ZBTB24 | 1.95 | 0.03 | AR-FL | RCL1 | -0.62 | -0.14 | AR-FL |
| LRP2BP | 1.21 | 0.03 | AR-FL | H6PD | -0.58 | -0.14 | AR-FL |
| TMCC1 | 0.58 | 0.03 | AR-FL | RPL22 | -0.58 | -0.14 | AR-FL |
| REPS2 | 1.04 | 0.03 | AR-FL | PAPLN | -2.17 | -0.14 | AR-FL |
| KDM4B | 1.04 | 0.04 | AR-FL | SPAG16 | -0.68 | -0.14 | AR-FL |
| TMEM198 | 0.83 | 0.04 | AR-FL | SGSM1 | -1.62 | -0.14 | AR-FL |
| RNF185 | 0.58 | 0.05 | AR-FL | RTTN | -0.72 | -0.14 | AR-FL |
| SRP68 | 0.59 | 0.05 | AR-FL | ADSSL1 | -0.61 | -0.13 | AR-FL |
| MESDC1 | 0.79 | 0.05 | AR-FL | ACRC | -0.92 | -0.13 | AR-FL |
| TMEM50B | 0.60 | 0.05 | AR-FL | APBB3 | -0.77 | -0.13 | AR-FL |
| CAPS | 0.75 | 0.05 | AR-FL | RASGRF2 | -0.90 | -0.13 | AR-FL |
| ADRB1 | 1.73 | 0.06 | AR-FL | F5 | -0.94 | -0.13 | AR-FL |
| ACACA | 1.04 | 0.06 | AR-FL | LRRC61 | -0.88 | -0.13 | AR-FL |
| MTMR3 | 0.68 | 0.06 | AR-FL | RBFA | -0.65 | -0.13 | AR-FL |
| OSBP | 0.75 | 0.06 | AR-FL | KIF27 | -1.06 | -0.13 | AR-FL |
| CANT1 | 0.83 | 0.06 | AR-FL | PLSCR1 | -1.66 | -0.13 | AR-FL |
| CSGALNACT2 | 0.69 | 0.06 | AR-FL | RPL7 | -0.63 | -0.13 | AR-FL |
| TRAPPC11 | 0.63 | 0.07 | AR-FL | KATNAL2 | -1.91 | -0.13 | AR-FL |
| SLC17A5 | 0.76 | 0.07 | AR-FL | TYSND1 | -0.59 | -0.13 | AR-FL |
| TBK1 | 0.59 | 0.07 | AR-FL | C3orf67 | -1.33 | -0.13 | AR-FL |
| GDF11 | 0.62 | 0.07 | AR-FL | APCDD1 | -0.90 | -0.13 | AR-FL |
| DOCK11 | 2.31 | 0.07 | AR-FL | C2orf44 | -0.59 | -0.13 | AR-FL |
| EIF2S1 | 0.65 | 0.07 | AR-FL | AVIL | -0.67 | -0.13 | AR-FL |
| MXD1 | 0.70 | 0.07 | AR-FL | ISCU | -0.62 | -0.13 | AR-FL |
| MAPK6 | 0.92 | 0.07 | AR-FL | SPEG | -1.34 | -0.13 | AR-FL |
| MKLN1 | 1.12 | 0.08 | AR-FL | ULBP3 | -1.54 | -0.12 | AR-FL |
| CREB3L4 | 1.66 | 0.08 | AR-FL | NOS1 | -3.16 | -0.12 | AR-FL |
| FBXL8 | 0.82 | 0.08 | AR-FL | ARRB2 | -0.71 | -0.12 | AR-FL |
| NUDT9 | 1.26 | 0.08 | AR-FL | FAM53A | -0.96 | -0.12 | AR-FL |
| GBP2 | 0.96 | 0.08 | AR-FL | PRPF4 | -0.59 | -0.12 | AR-FL |
| RIPK4 | 0.64 | 0.08 | AR-FL | LAMC3 | -2.94 | -0.12 | AR-FL |
| ATMIN | 1.08 | 0.08 | AR-FL | PDIA2 | -2.36 | -0.12 | AR-FL |
| PLPP3 | 0.74 | 0.08 | AR-FL | PRRT4 | -1.21 | -0.12 | AR-FL |
| FAM214A | 0.72 | 0.08 | AR-FL | C1orf74 | -0.65 | -0.12 | AR-FL |
| BTAF1 | 0.94 | 0.08 | AR-FL | POLR2I | -0.61 | -0.12 | AR-FL |
| TRIM5 | 0.83 | 0.09 | AR-FL | EEF1A1 | -0.69 | -0.12 | AR-FL |
| PAK1IP1 | 2.70 | 0.09 | AR-FL | SNX7 | -0.69 | -0.12 | AR-FL |
| DLG2 | 2.47 | 0.09 | AR-FL | ZNF438 | -0.82 | -0.12 | AR-FL |

| | | | | | | | |
|---|---|---|---|---|---|---|---|
| HM13 | 0.78 | 0.09 | AR-FL | PHLDA1 | -0.66 | -0.12 | AR-FL |
| EPN2 | 0.87 | 0.09 | AR-FL | ARL4D | -0.94 | -0.12 | AR-FL |
| CCDC136 | 0.78 | 0.09 | AR-FL | TCEANC2 | -0.69 | -0.12 | AR-FL |
| UBE2J1 | 1.44 | 0.09 | AR-FL | JAG2 | -0.62 | -0.12 | AR-FL |
| RGPD6 | 1.26 | 0.09 | AR-FL | CAMKK1 | -0.87 | -0.12 | AR-FL |
| IL10RB | 1.10 | 0.09 | AR-FL | ALAD | -0.58 | -0.12 | AR-FL |
| PLEKHB2 | 0.66 | 0.09 | AR-FL | IDNK | -0.67 | -0.12 | AR-FL |
| PCTP | 1.78 | 0.10 | AR-FL | NFXL1 | -0.59 | -0.11 | AR-FL |
| COG5 | 0.66 | 0.10 | AR-FL | NPIPA1 | -1.01 | -0.11 | AR-FL |
| WSB2 | 0.65 | 0.10 | AR-FL | HSPA12A | -1.42 | -0.11 | AR-FL |
| COL4A5 | 0.84 | 0.10 | AR-FL | GRAMD1B | -1.37 | -0.11 | AR-FL |
| FECH | 0.89 | 0.10 | AR-FL | MAP6 | -0.84 | -0.11 | AR-FL |
| DYNLL2 | 0.59 | 0.10 | AR-FL | PSIP1 | -0.64 | -0.11 | AR-FL |
| TAF1A | 0.74 | 0.10 | AR-FL | ZNF165 | -0.66 | -0.11 | AR-FL |
| DNAH5 | 0.83 | 0.10 | AR-FL | RPS3A | -0.64 | -0.11 | AR-FL |
| MFN1 | 0.74 | 0.10 | AR-FL | ZNF497 | -0.74 | -0.11 | AR-FL |
| LARP4 | 0.65 | 0.11 | AR-FL | RPL18 | -0.62 | -0.11 | AR-FL |
| RALBP1 | 0.71 | 0.11 | AR-FL | TMEM240 | -0.70 | -0.11 | AR-FL |
| CCDC71L | 0.69 | 0.11 | AR-FL | NNT | -1.57 | -0.11 | AR-FL |
| GFM1 | 1.66 | 0.11 | AR-FL | NLRP11 | -0.61 | -0.11 | AR-FL |
| ERGIC2 | 1.12 | 0.11 | AR-FL | SYK | -0.77 | -0.11 | AR-FL |
| ACOX3 | 1.17 | 0.11 | AR-FL | ARRB1 | -0.76 | -0.11 | AR-FL |
| PEX10 | 2.23 | 0.11 | AR-FL | RPS14 | -0.62 | -0.10 | AR-FL |
| DHRS12 | 0.66 | 0.11 | AR-FL | CCDC82 | -0.81 | -0.10 | AR-FL |
| UCHL3 | 0.72 | 0.11 | AR-FL | PPP1R1C | -1.32 | -0.10 | AR-FL |
| PRKCH | 1.91 | 0.12 | AR-FL | PRKAB2 | -1.05 | -0.10 | AR-FL |
| SAMD9 | 1.23 | 0.12 | AR-FL | MAPK10 | -0.58 | -0.10 | AR-FL |
| PRDX6 | 0.65 | 0.12 | AR-FL | KBTBD11 | -0.70 | -0.10 | AR-FL |
| SEC16A | 0.61 | 0.12 | AR-FL | KCNQ4 | -1.00 | -0.10 | AR-FL |
| EFHC1 | 0.61 | 0.12 | AR-FL | CENPV | -0.66 | -0.10 | AR-FL |
| PPARA | 0.74 | 0.13 | AR-FL | NTN4 | -1.98 | -0.10 | AR-FL |
| TFPT | 0.90 | 0.13 | AR-FL | KCP | -1.53 | -0.10 | AR-FL |
| THYN1 | 0.59 | 0.13 | AR-FL | HIST1H2BC | -0.75 | -0.10 | AR-FL |
| ZNF761 | 0.64 | 0.13 | AR-FL | NEIL2 | -0.72 | -0.10 | AR-FL |
| APLN | 0.59 | 0.14 | AR-FL | EFCAB6 | -1.03 | -0.09 | AR-FL |
| GBA2 | 0.63 | 0.14 | AR-FL | GATM | -0.74 | -0.09 | AR-FL |
| TUSC3 | 0.79 | 0.14 | AR-FL | BCKDHB | -0.85 | -0.09 | AR-FL |
| SPCS2 | 0.64 | 0.14 | AR-FL | GPR143 | -0.68 | -0.09 | AR-FL |
| ACAA1 | 1.22 | 0.14 | AR-FL | NTN3 | -0.72 | -0.09 | AR-FL |
| HK2 | 0.75 | 0.14 | AR-FL | TFR2 | -1.02 | -0.09 | AR-FL |
| PSMD8 | 0.79 | 0.14 | AR-FL | CEP72 | -0.59 | -0.09 | AR-FL |
| OSTM1 | 0.61 | 0.14 | AR-FL | KIAA1161 | -0.63 | -0.09 | AR-FL |
| RGAG4 | 0.73 | 0.14 | AR-FL | MYOT | -0.63 | -0.09 | AR-FL |
| PERP | 0.62 | 0.15 | AR-FL | ZHX3 | -0.71 | -0.08 | AR-FL |
| CDK6 | 2.02 | 0.15 | AR-FL | PAIP2B | -0.71 | -0.08 | AR-FL |
| TPD52 | 1.73 | 0.15 | AR-FL | RCOR2 | -0.99 | -0.08 | AR-FL |

| | | | | | | | |
|---|---|---|---|---|---|---|---|
| GTF2E2 | 1.01 | 0.15 | AR-FL | ZNF529 | -0.92 | -0.08 | AR-FL |
| USP33 | 0.99 | 0.15 | AR-FL | SEC31B | -1.03 | -0.08 | AR-FL |
| SLC9A3R2 | 1.23 | 0.15 | AR-FL | ZNF483 | -0.75 | -0.08 | AR-FL |
| SMPD2 | 1.64 | 0.15 | AR-FL | NOTCH1 | -0.86 | -0.08 | AR-FL |
| SND1 | 0.59 | 0.16 | AR-FL | SPRED3 | -0.96 | -0.08 | AR-FL |
| SH3D19 | 0.72 | 0.16 | AR-FL | MECR | -0.80 | -0.08 | AR-FL |
| CDC42EP3 | 2.56 | 0.16 | AR-FL | AARD | -2.87 | -0.08 | AR-FL |
| ZFHX3 | 0.61 | 0.16 | AR-FL | FCGRT | -0.85 | -0.08 | AR-FL |
| RAB5A | 0.75 | 0.16 | AR-FL | NIPAL3 | -0.67 | -0.07 | AR-FL |
| SLC35C1 | 0.70 | 0.16 | AR-FL | OGT | -1.01 | -0.07 | AR-FL |
| SNAPC2 | 0.92 | 0.16 | AR-FL | ASB9 | -1.50 | -0.07 | AR-FL |
| CD164 | 0.74 | 0.17 | AR-FL | PRRG1 | -0.63 | -0.07 | AR-FL |
| TES | 0.92 | 0.17 | AR-FL | ZFHX4 | -1.04 | -0.07 | AR-FL |
| YIPF1 | 1.30 | 0.17 | AR-FL | PABPC1L | -1.14 | -0.07 | AR-FL |
| KIAA1324L | 0.73 | 0.17 | AR-FL | RWDD3 | -0.87 | -0.07 | AR-FL |
| PDE9A | 1.31 | 0.17 | AR-FL | ADAMTS17 | -1.48 | -0.07 | AR-FL |
| MTMR2 | 1.03 | 0.17 | AR-FL | PCSK4 | -1.12 | -0.07 | AR-FL |
| TRIM52 | 1.26 | 0.18 | AR-FL | NANOS1 | -0.62 | -0.06 | AR-FL |
| TBC1D15 | 0.74 | 0.18 | AR-FL | SKP2 | -0.72 | -0.06 | AR-FL |
| GLRX2 | 1.35 | 0.18 | AR-FL | JAK1 | -0.59 | -0.06 | AR-FL |
| MTRNR2L8 | 0.86 | 0.18 | AR-FL | EHHADH | -0.98 | -0.06 | AR-FL |
| TMEM125 | 0.92 | 0.18 | AR-FL | C8orf44 | -0.79 | -0.06 | AR-FL |
| GGPS1 | 0.72 | 0.18 | AR-FL | MST1 | -1.03 | -0.05 | AR-FL |
| PROSER2 | 1.07 | 0.18 | AR-FL | DSCAM | -2.21 | -0.05 | AR-FL |
| SRPRB | 0.68 | 0.18 | AR-FL | TRAF1 | -1.20 | -0.05 | AR-FL |
| GALNT7 | 1.06 | 0.18 | AR-FL | FAM161A | -0.82 | -0.05 | AR-FL |
| DCAF6 | 0.93 | 0.19 | AR-FL | ZNF665 | -0.73 | -0.05 | AR-FL |
| FBXO8 | 0.66 | 0.19 | AR-FL | BEST1 | -2.01 | -0.05 | AR-FL |
| ATP6V0A2 | 1.09 | 0.19 | AR-FL | EID2B | -0.92 | -0.05 | AR-FL |
| DGKA | 1.24 | 0.19 | AR-FL | ZSCAN18 | -0.69 | -0.04 | AR-FL |
| EZR | 0.87 | 0.19 | AR-FL | GOLGA8B | -1.64 | -0.04 | AR-FL |
| YES1 | 0.66 | 0.19 | AR-FL | DCHS1 | -0.79 | -0.04 | AR-FL |
| S100A11 | 0.86 | 0.20 | AR-FL | RILPL1 | -0.74 | -0.04 | AR-FL |
| MEGF9 | 0.86 | 0.20 | AR-FL | CAD | -0.98 | -0.04 | AR-FL |
| CBL | 0.61 | 0.20 | AR-FL | DDX51 | -0.63 | -0.04 | AR-FL |
| BAIAP2 | 0.95 | 0.20 | AR-FL | RCAN2 | -1.04 | -0.04 | AR-FL |
| SCYL1 | 0.59 | 0.20 | AR-FL | RCOR3 | -0.60 | -0.04 | AR-FL |
| CYTH1 | 1.61 | 0.20 | AR-FL | CA14 | -0.71 | -0.04 | AR-FL |
| SH3TC1 | 0.71 | 0.20 | AR-FL | LTBP3 | -0.70 | -0.03 | AR-FL |
| PREB | 0.66 | 0.20 | AR-FL | OPN3 | -0.89 | -0.03 | AR-FL |
| TMEM14C | 0.62 | 0.20 | AR-FL | HBQ1 | -1.43 | -0.03 | AR-FL |
| ZNF155 | 0.65 | 0.20 | AR-FL | CCDC188 | -0.70 | -0.03 | AR-FL |
| SLC22A23 | 0.97 | 0.21 | AR-FL | LDHD | -0.92 | -0.03 | AR-FL |
| SPATA2 | 0.62 | 0.21 | AR-FL | UBE2V1 | -0.59 | -0.03 | AR-FL |
| CTGLF11P | 0.84 | 0.21 | AR-FL | ZNF280C | -0.95 | -0.03 | AR-FL |
| JPH1 | 0.70 | 0.21 | AR-FL | ADAM23 | -2.10 | -0.03 | AR-FL |

| | | | |
|---|---|---|---|
| ZBTB37 | 0.64 | 0.21 | AR-FL |
| SLC25A20 | 1.17 | 0.21 | AR-FL |
| SLC35F2 | 2.07 | 0.21 | AR-FL |
| IGBP1 | 0.66 | 0.21 | AR-FL |
| ALG2 | 0.72 | 0.22 | AR-FL |
| SLC45A2 | 1.37 | 0.22 | AR-FL |
| WFS1 | 0.74 | 0.22 | AR-FL |
| TNIP2 | 0.64 | 0.22 | AR-FL |
| CRNKL1 | 0.71 | 0.22 | AR-FL |
| SH3BGRL | 1.43 | 0.22 | AR-FL |
| TMEM2 | 1.01 | 0.22 | AR-FL |
| PRODH | 0.87 | 0.22 | AR-FL |
| VMAC | 1.00 | 0.22 | AR-FL |
| SPRED2 | 0.77 | 0.22 | AR-FL |
| JKAMP | 0.61 | 0.23 | AR-FL |
| ABHD3 | 1.17 | 0.23 | AR-FL |
| COG3 | 0.61 | 0.23 | AR-FL |
| MMAA | 1.06 | 0.23 | AR-FL |
| PMM2 | 0.66 | 0.23 | AR-FL |
| PPM1D | 0.89 | 0.23 | AR-FL |
| GOLPH3 | 0.97 | 0.23 | AR-FL |
| CLCN6 | 0.61 | 0.23 | AR-FL |
| PSEN1 | 0.61 | 0.23 | AR-FL |
| HN1L | 0.62 | 0.23 | AR-FL |
| SEC61A1 | 0.66 | 0.24 | AR-FL |
| TMEM50A | 0.89 | 0.24 | AR-FL |
| SIAH2 | 0.76 | 0.24 | AR-FL |
| ANKRD27 | 0.60 | 0.24 | AR-FL |
| KIAA0513 | 0.59 | 0.24 | AR-FL |
| CHM | 0.66 | 0.24 | AR-FL |
| IMMP1L | 0.62 | 0.24 | AR-FL |
| SLC25A13 | 0.65 | 0.24 | AR-FL |
| PSMA6 | 0.98 | 0.24 | AR-FL |
| ACLY | 0.71 | 0.24 | AR-FL |
| MYO1C | 0.68 | 0.24 | AR-FL |
| PRDM4 | 0.65 | 0.24 | AR-FL |
| SMURF2 | 0.68 | 0.24 | AR-FL |
| SACS | 0.88 | 0.24 | AR-FL |
| ARCN1 | 0.69 | 0.24 | AR-FL |
| MON1B | 0.79 | 0.25 | AR-FL |
| ATP6V1D | 0.75 | 0.25 | AR-FL |
| TMEM87B | 1.49 | 0.25 | AR-FL |
| PGM3 | 1.51 | 0.25 | AR-FL |
| GMNC | 1.27 | 0.25 | AR-FL |
| BCL2L1 | 0.81 | 0.25 | AR-FL |
| ZG16B | 1.02 | 0.26 | AR-FL |
| C11orf96 | -1.70 | -0.02 | AR-FL |
| TCFL5 | -0.61 | -0.02 | AR-FL |
| DUT | -0.64 | -0.02 | AR-FL |
| RIPPLY3 | -0.96 | -0.02 | AR-FL |
| COL7A1 | -1.62 | -0.02 | AR-FL |
| OBSL1 | -0.67 | -0.02 | AR-FL |
| UBAP1L | -0.98 | -0.02 | AR-FL |
| GOLT1A | -0.74 | -0.02 | AR-FL |
| DYRK4 | -0.64 | -0.01 | AR-FL |
| AC092835.2 | -1.15 | -0.01 | AR-FL |
| SOD2 | -0.68 | -0.01 | AR-FL |
| MPP3 | -1.20 | -0.01 | AR-FL |
| ZBTB48 | -0.59 | -0.01 | AR-FL |
| ZNF550 | -0.78 | -0.01 | AR-FL |
| FAM229A | -0.59 | -0.01 | AR-FL |
| TCTN1 | -0.73 | -0.01 | AR-FL |
| FAM96A | -0.64 | -0.01 | AR-FL |
| ZFP36L1 | -1.48 | 0.00 | AR-FL |
| ACACB | -1.42 | 0.00 | AR-FL |
| MYH7B | -0.87 | 0.00 | AR-FL |
| MND1 | -0.97 | 0.00 | AR-FL |
| DNAJC19 | -0.60 | 0.00 | AR-FL |
| SLC22A10 | -0.02 | -2.25 | AR-V7 |
| MAB21L3 | -0.03 | -1.93 | AR-V7 |
| GPRC5A | -0.52 | -1.70 | AR-V7 |
| RIMS1 | -0.21 | -1.68 | AR-V7 |
| TRDN | -0.45 | -1.65 | AR-V7 |
| GBP3 | -0.23 | -1.58 | AR-V7 |
| ADAMTSL1 | -0.27 | -1.58 | AR-V7 |
| MYO18B | -0.38 | -1.53 | AR-V7 |
| PTPRK | -0.28 | -1.48 | AR-V7 |
| RPL21 | -0.56 | -1.44 | AR-V7 |
| BCAS1 | -0.23 | -1.40 | AR-V7 |
| PTPRE | -0.20 | -1.38 | AR-V7 |
| WNT5A | -0.53 | -1.35 | AR-V7 |
| IGFL2 | -0.23 | -1.34 | AR-V7 |
| LUZP2 | -0.15 | -1.32 | AR-V7 |
| RASSF6 | -0.50 | -1.31 | AR-V7 |
| ACPP | -0.03 | -1.29 | AR-V7 |
| SPRY4 | -0.57 | -1.24 | AR-V7 |
| SSTR5 | -0.48 | -1.24 | AR-V7 |
| GDPD1 | -0.12 | -1.23 | AR-V7 |
| SERTAD4 | -0.38 | -1.22 | AR-V7 |
| C10orf10 | -0.41 | -1.22 | AR-V7 |
| C16orf70 | -0.25 | -1.21 | AR-V7 |
| TCERG1L | -0.53 | -1.18 | AR-V7 |

| | | | |
|---|---|---|---|
| TTC12 | 0.60 | 0.26 | AR-FL |
| FBXL3 | 0.64 | 0.26 | AR-FL |
| TM2D2 | 0.58 | 0.26 | AR-FL |
| KCNRG | 0.63 | 0.26 | AR-FL |
| COPB1 | 0.63 | 0.26 | AR-FL |
| IMPDH1 | 1.01 | 0.27 | AR-FL |
| TESK1 | 0.63 | 0.27 | AR-FL |
| VAMP7 | 3.00 | 0.27 | AR-FL |
| CASD1 | 0.63 | 0.27 | AR-FL |
| MUC20 | 1.15 | 0.27 | AR-FL |
| CCDC53 | 1.25 | 0.27 | AR-FL |
| C20orf24 | 0.73 | 0.27 | AR-FL |
| FZD8 | 1.58 | 0.27 | AR-FL |
| SLC25A25 | 0.76 | 0.27 | AR-FL |
| TMEM41B | 0.75 | 0.27 | AR-FL |
| SRPR | 0.60 | 0.28 | AR-FL |
| C15orf61 | 0.97 | 0.28 | AR-FL |
| STAMBP | 0.59 | 0.28 | AR-FL |
| SLC33A1 | 0.87 | 0.28 | AR-FL |
| TMED10 | 0.71 | 0.28 | AR-FL |
| SERP1 | 1.11 | 0.28 | AR-FL |
| LRRC59 | 1.02 | 0.28 | AR-FL |
| MEGF6 | 1.35 | 0.28 | AR-FL |
| ZHX1 | 0.62 | 0.28 | AR-FL |
| ZNF697 | 1.13 | 0.29 | AR-FL |
| ARHGAP26 | 1.20 | 0.29 | AR-FL |
| PICALM | 0.75 | 0.29 | AR-FL |
| FMN1 | 1.63 | 0.29 | AR-FL |
| LA16c-380H5.3 | 0.98 | 0.29 | AR-FL |
| MTM1 | 0.90 | 0.29 | AR-FL |
| GORASP2 | 0.72 | 0.29 | AR-FL |
| FAM221A | 0.71 | 0.30 | AR-FL |
| ZNF540 | 0.98 | 0.30 | AR-FL |
| GNPNAT1 | 1.05 | 0.30 | AR-FL |
| EGFR | 1.04 | 0.30 | AR-FL |
| DNAJC3 | 1.23 | 0.30 | AR-FL |
| THAP5 | 0.62 | 0.30 | AR-FL |
| GOLGA5 | 0.83 | 0.30 | AR-FL |
| RBMXL1 | 0.78 | 0.30 | AR-FL |
| FHDC1 | 1.08 | 0.31 | AR-FL |
| NUMB | 0.59 | 0.31 | AR-FL |
| SEC23B | 0.83 | 0.31 | AR-FL |
| CORO1B | 1.26 | 0.31 | AR-FL |
| PTPN1 | 0.72 | 0.31 | AR-FL |
| MIPEP | 1.17 | 0.31 | AR-FL |
| CD2AP | 0.65 | 0.31 | AR-FL |
| DAB2 | -0.54 | -1.17 | AR-V7 |
| RELL1 | -0.06 | -1.17 | AR-V7 |
| TMOD1 | -0.56 | -1.16 | AR-V7 |
| MITF | -0.06 | -1.16 | AR-V7 |
| DACT2 | -0.31 | -1.15 | AR-V7 |
| TP53I3 | -0.30 | -1.15 | AR-V7 |
| ADRB2 | -0.44 | -1.15 | AR-V7 |
| CCDC146 | -0.38 | -1.13 | AR-V7 |
| ELK3 | -0.47 | -1.10 | AR-V7 |
| PELI1 | -0.50 | -1.09 | AR-V7 |
| ANKRD33B | -0.26 | -1.08 | AR-V7 |
| SMAD9 | -0.49 | -1.06 | AR-V7 |
| CTD-3088G3.8 | -0.28 | -1.06 | AR-V7 |
| TESK2 | -0.32 | -1.05 | AR-V7 |
| PVRL3 | -0.01 | -1.05 | AR-V7 |
| EYA2 | 0.00 | -1.04 | AR-V7 |
| TRIM2 | -0.35 | -1.03 | AR-V7 |
| SLC2A13 | -0.07 | -1.01 | AR-V7 |
| CNTNAP2 | -0.25 | -1.01 | AR-V7 |
| DRAM1 | -0.40 | -1.00 | AR-V7 |
| SLC22A15 | -0.36 | -0.99 | AR-V7 |
| CACNA1D | -0.37 | -0.98 | AR-V7 |
| NTF3 | -0.22 | -0.97 | AR-V7 |
| SRSF10 | -0.24 | -0.96 | AR-V7 |
| RGPD8 | -0.03 | -0.96 | AR-V7 |
| TRIM16L | -0.58 | -0.96 | AR-V7 |
| UNC80 | -0.50 | -0.96 | AR-V7 |
| MGLL | -0.22 | -0.95 | AR-V7 |
| NCEH1 | -0.50 | -0.95 | AR-V7 |
| PSD3 | -0.22 | -0.95 | AR-V7 |
| PAX5 | -0.29 | -0.95 | AR-V7 |
| TOX3 | -0.05 | -0.94 | AR-V7 |
| TTC9 | -0.29 | -0.94 | AR-V7 |
| NKX2-2 | -0.49 | -0.94 | AR-V7 |
| PDE11A | -0.27 | -0.93 | AR-V7 |
| CABP4 | -0.30 | -0.93 | AR-V7 |
| C1orf115 | -0.36 | -0.93 | AR-V7 |
| TSHZ2 | -0.34 | -0.93 | AR-V7 |
| ADPRM | -0.53 | -0.93 | AR-V7 |
| L3MBTL3 | -0.33 | -0.92 | AR-V7 |
| RWDD2A | -0.17 | -0.91 | AR-V7 |
| KRBA2 | -0.57 | -0.91 | AR-V7 |
| KSR1 | -0.05 | -0.91 | AR-V7 |
| APBB2 | -0.54 | -0.90 | AR-V7 |
| APOL6 | -0.43 | -0.90 | AR-V7 |
| ARHGAP42 | -0.05 | -0.89 | AR-V7 |

| | | | |
|---|---|---|---|
| SEC22C | 0.66 | 0.31 | AR-FL |
| KLHDC2 | 0.60 | 0.31 | AR-FL |
| SLC4A7 | 0.59 | 0.31 | AR-FL |
| LRIG1 | 1.95 | 0.31 | AR-FL |
| YOD1 | 0.75 | 0.31 | AR-FL |
| BCAR3 | 1.30 | 0.31 | AR-FL |
| ZNF350 | 2.27 | 0.31 | AR-FL |
| MSMB | 0.91 | 0.31 | AR-FL |
| C9orf91 | 0.98 | 0.31 | AR-FL |
| PDIA6 | 0.77 | 0.31 | AR-FL |
| HCFC2 | 0.75 | 0.31 | AR-FL |
| STX5 | 0.66 | 0.32 | AR-FL |
| TXNDC11 | 0.86 | 0.32 | AR-FL |
| MUC4 | 0.80 | 0.32 | AR-FL |
| SLC19A2 | 1.36 | 0.32 | AR-FL |
| SQLE | 0.62 | 0.32 | AR-FL |
| TIGD6 | 0.80 | 0.32 | AR-FL |
| SARAF | 0.80 | 0.32 | AR-FL |
| TRIM3 | 1.48 | 0.32 | AR-FL |
| SMURF1 | 0.74 | 0.32 | AR-FL |
| HYOU1 | 0.83 | 0.33 | AR-FL |
| C6orf52 | 0.69 | 0.33 | AR-FL |
| C11orf54 | 0.65 | 0.33 | AR-FL |
| GUCY1A3 | 1.23 | 0.33 | AR-FL |
| MTOR | 1.55 | 0.33 | AR-FL |
| STRN | 0.61 | 0.33 | AR-FL |
| TM4SF1 | 2.10 | 0.33 | AR-FL |
| TMEM8A | 0.62 | 0.33 | AR-FL |
| SLC35A2 | 0.65 | 0.33 | AR-FL |
| SEC22B | 0.64 | 0.33 | AR-FL |
| MAK | 2.58 | 0.34 | AR-FL |
| ARFGAP3 | 0.97 | 0.34 | AR-FL |
| CRISP3 | 1.91 | 0.34 | AR-FL |
| NANS | 1.09 | 0.34 | AR-FL |
| NOMO2 | 0.62 | 0.34 | AR-FL |
| HSPA13 | 0.98 | 0.34 | AR-FL |
| ACAD8 | 1.50 | 0.34 | AR-FL |
| BICD2 | 1.66 | 0.34 | AR-FL |
| TIFA | 0.76 | 0.34 | AR-FL |
| FKBP2 | 0.70 | 0.35 | AR-FL |
| COL4A6 | 1.38 | 0.35 | AR-FL |
| TIMM23B | 0.59 | 0.35 | AR-FL |
| ZNF613 | 1.26 | 0.35 | AR-FL |
| MPZL1 | 1.07 | 0.35 | AR-FL |
| MESDC2 | 0.62 | 0.35 | AR-FL |
| SSR1 | 1.04 | 0.35 | AR-FL |
| NRIP1 | -0.44 | -0.89 | AR-V7 |
| SH3RF1 | -0.54 | -0.89 | AR-V7 |
| LFNG | -0.26 | -0.89 | AR-V7 |
| LIN7A | -0.57 | -0.88 | AR-V7 |
| C21orf91 | -0.33 | -0.88 | AR-V7 |
| GALNT3 | -0.56 | -0.87 | AR-V7 |
| PCDHGA8 | -0.13 | -0.87 | AR-V7 |
| RAB25 | -0.51 | -0.86 | AR-V7 |
| ZNF860 | -0.41 | -0.85 | AR-V7 |
| UGT1A1 | -0.44 | -0.85 | AR-V7 |
| RALGAPA2 | -0.15 | -0.85 | AR-V7 |
| PPP2R3B | -0.56 | -0.84 | AR-V7 |
| DDX58 | -0.48 | -0.84 | AR-V7 |
| ARHGAP6 | -0.38 | -0.84 | AR-V7 |
| FAM171A2 | -0.48 | -0.84 | AR-V7 |
| SLC25A21 | -0.46 | -0.84 | AR-V7 |
| SLCO5A1 | -0.53 | -0.84 | AR-V7 |
| ZC3H12B | -0.51 | -0.84 | AR-V7 |
| HSD17B14 | -0.56 | -0.83 | AR-V7 |
| MYO6 | -0.15 | -0.83 | AR-V7 |
| POLN | -0.49 | -0.82 | AR-V7 |
| RNF186 | -0.36 | -0.82 | AR-V7 |
| NAALADL2 | -0.53 | -0.81 | AR-V7 |
| TSPAN6 | -0.30 | -0.81 | AR-V7 |
| TMEM44 | -0.44 | -0.81 | AR-V7 |
| HOXC11 | -0.45 | -0.81 | AR-V7 |
| TTC39C | -0.01 | -0.81 | AR-V7 |
| RNF128 | -0.23 | -0.80 | AR-V7 |
| ADAMTS1 | -0.47 | -0.80 | AR-V7 |
| MEX3B | -0.55 | -0.79 | AR-V7 |
| MIPOL1 | -0.48 | -0.78 | AR-V7 |
| C3orf80 | -0.04 | -0.78 | AR-V7 |
| CDR2L | -0.16 | -0.78 | AR-V7 |
| ANKMY2 | -0.02 | -0.77 | AR-V7 |
| CLSTN2 | -0.52 | -0.77 | AR-V7 |
| MIOS | -0.56 | -0.76 | AR-V7 |
| NBPF10 | -0.54 | -0.76 | AR-V7 |
| TP53INP1 | -0.52 | -0.76 | AR-V7 |
| AGMAT | -0.04 | -0.76 | AR-V7 |
| ANKRD29 | -0.56 | -0.76 | AR-V7 |
| PRICKLE2 | -0.41 | -0.75 | AR-V7 |
| TBL1Y | -0.39 | -0.75 | AR-V7 |
| SORBS2 | -0.22 | -0.74 | AR-V7 |
| ICK | -0.47 | -0.74 | AR-V7 |
| BPIFB1 | -0.48 | -0.73 | AR-V7 |
| SFRP1 | -0.28 | -0.73 | AR-V7 |

| | | | |
|---|---|---|---|
| SEC24C | 0.78 | 0.35 | AR-FL |
| MYCL | 0.99 | 0.35 | AR-FL |
| PPP1R13B | 0.61 | 0.35 | AR-FL |
| C1orf210 | 0.75 | 0.35 | AR-FL |
| PRRC1 | 0.81 | 0.35 | AR-FL |
| TMED2 | 0.68 | 0.35 | AR-FL |
| HES4 | 0.64 | 0.36 | AR-FL |
| SHROOM2 | 0.96 | 0.36 | AR-FL |
| ABAT | 0.59 | 0.36 | AR-FL |
| MRPS35 | 0.67 | 0.36 | AR-FL |
| RSPH9 | 0.64 | 0.36 | AR-FL |
| SPCS3 | 1.23 | 0.36 | AR-FL |
| KLB | 1.07 | 0.36 | AR-FL |
| SPTBN4 | 0.97 | 0.36 | AR-FL |
| SEPP1 | 2.25 | 0.36 | AR-FL |
| UBE2G1 | 1.20 | 0.37 | AR-FL |
| MEAF6 | 1.22 | 0.37 | AR-FL |
| BCAP29 | 1.07 | 0.37 | AR-FL |
| UBE2E1 | 0.59 | 0.37 | AR-FL |
| RAB1A | 0.66 | 0.37 | AR-FL |
| ZNF649 | 0.66 | 0.37 | AR-FL |
| MAP7D1 | 1.13 | 0.37 | AR-FL |
| C17orf62 | 0.68 | 0.37 | AR-FL |
| STT3A | 0.79 | 0.37 | AR-FL |
| ZNF518B | 0.96 | 0.37 | AR-FL |
| PEA15 | 0.92 | 0.37 | AR-FL |
| SNAP23 | 0.89 | 0.37 | AR-FL |
| SHOX2 | 0.71 | 0.37 | AR-FL |
| RREB1 | 0.75 | 0.37 | AR-FL |
| NOMO3 | 0.70 | 0.37 | AR-FL |
| PTPRJ | 0.93 | 0.37 | AR-FL |
| CMAS | 0.81 | 0.38 | AR-FL |
| VEGFA | 1.03 | 0.38 | AR-FL |
| C18orf8 | 0.71 | 0.38 | AR-FL |
| 5-Mar | 0.93 | 0.38 | AR-FL |
| ELOVL7 | 3.20 | 0.38 | AR-FL |
| GMPPB | 1.00 | 0.38 | AR-FL |
| ATG13 | 0.59 | 0.38 | AR-FL |
| SLC25A37 | 0.89 | 0.38 | AR-FL |
| ABCC11 | 1.13 | 0.38 | AR-FL |
| TMEM214 | 0.73 | 0.39 | AR-FL |
| MCPH1 | 0.73 | 0.39 | AR-FL |
| LATS2 | 0.99 | 0.39 | AR-FL |
| GLMP | 1.63 | 0.39 | AR-FL |
| ARF4 | 1.03 | 0.39 | AR-FL |
| KDELR2 | 1.08 | 0.39 | AR-FL |
| TMEM139 | -0.28 | -0.73 | AR-V7 |
| FAM122C | -0.49 | -0.73 | AR-V7 |
| FBXO25 | -0.57 | -0.72 | AR-V7 |
| DOCK4 | -0.32 | -0.72 | AR-V7 |
| ZNF280D | -0.58 | -0.72 | AR-V7 |
| ITGA5 | -0.12 | -0.72 | AR-V7 |
| SGMS2 | -0.49 | -0.72 | AR-V7 |
| SMYD3 | -0.24 | -0.72 | AR-V7 |
| MAP4K1 | -0.58 | -0.72 | AR-V7 |
| RBPMS | -0.26 | -0.71 | AR-V7 |
| GRAMD1A | -0.49 | -0.71 | AR-V7 |
| CYP4X1 | -0.56 | -0.71 | AR-V7 |
| KLF13 | -0.19 | -0.71 | AR-V7 |
| SETBP1 | -0.57 | -0.71 | AR-V7 |
| FBXL7 | -0.49 | -0.71 | AR-V7 |
| DIO1 | -0.12 | -0.71 | AR-V7 |
| MB | -0.05 | -0.70 | AR-V7 |
| ADCY1 | -0.04 | -0.70 | AR-V7 |
| ZNF117 | -0.31 | -0.70 | AR-V7 |
| UST | -0.38 | -0.70 | AR-V7 |
| CTIF | -0.40 | -0.70 | AR-V7 |
| CXCR6 | -0.53 | -0.70 | AR-V7 |
| IGF2 | -0.46 | -0.70 | AR-V7 |
| CNTN3 | -0.42 | -0.70 | AR-V7 |
| PAM | -0.13 | -0.70 | AR-V7 |
| SNX16 | -0.25 | -0.70 | AR-V7 |
| ATP12A | -0.40 | -0.69 | AR-V7 |
| AC013461.1 | -0.50 | -0.69 | AR-V7 |
| COL4A6 | -0.55 | -0.69 | AR-V7 |
| COL9A2 | -0.36 | -0.69 | AR-V7 |
| GDAP1 | -0.44 | -0.68 | AR-V7 |
| PCDHB16 | -0.30 | -0.68 | AR-V7 |
| ARHGEF6 | -0.41 | -0.68 | AR-V7 |
| TMCO4 | -0.30 | -0.68 | AR-V7 |
| CNTN4 | -0.40 | -0.68 | AR-V7 |
| ZNF532 | -0.29 | -0.68 | AR-V7 |
| NFKB2 | -0.09 | -0.67 | AR-V7 |
| ADSS | -0.43 | -0.67 | AR-V7 |
| SERPINB6 | -0.23 | -0.67 | AR-V7 |
| HELB | -0.52 | -0.67 | AR-V7 |
| RAP1GAP2 | -0.49 | -0.67 | AR-V7 |
| TRPV6 | -0.15 | -0.67 | AR-V7 |
| HEY1 | -0.31 | -0.67 | AR-V7 |
| ZNF680 | -0.37 | -0.67 | AR-V7 |
| MCU | -0.21 | -0.67 | AR-V7 |
| ISPD | -0.16 | -0.67 | AR-V7 |

| | | | | | | | |
|---|---|---|---|---|---|---|---|
| DNAJC10 | 1.47 | 0.39 | AR-FL | TLE2 | -0.49 | -0.67 | AR-V7 |
| POP1 | 0.62 | 0.39 | AR-FL | ZSWIM4 | -0.28 | -0.66 | AR-V7 |
| OTULIN | 1.11 | 0.39 | AR-FL | CTNNAL1 | -0.55 | -0.66 | AR-V7 |
| C16orf46 | 0.78 | 0.40 | AR-FL | PTPN13 | -0.31 | -0.66 | AR-V7 |
| APP | 1.45 | 0.40 | AR-FL | LPAR6 | -0.45 | -0.66 | AR-V7 |
| DERL2 | 0.85 | 0.40 | AR-FL | 6-Sep | -0.55 | -0.66 | AR-V7 |
| THUMPD1 | 0.61 | 0.41 | AR-FL | LRP4 | -0.29 | -0.66 | AR-V7 |
| MAP2K4 | 0.99 | 0.41 | AR-FL | CAPN5 | -0.56 | -0.65 | AR-V7 |
| GOLGA3 | 0.67 | 0.41 | AR-FL | NCKAP5 | -0.47 | -0.65 | AR-V7 |
| COPG1 | 0.99 | 0.41 | AR-FL | NR1H3 | -0.41 | -0.65 | AR-V7 |
| CAP1 | 0.73 | 0.42 | AR-FL | AAMDC | -0.48 | -0.65 | AR-V7 |
| GREB1 | 2.93 | 0.42 | AR-FL | GRHL2 | -0.16 | -0.65 | AR-V7 |
| SPRYD7 | 1.23 | 0.42 | AR-FL | SCAMP1 | -0.08 | -0.65 | AR-V7 |
| SNX32 | 0.77 | 0.42 | AR-FL | PLS1 | -0.32 | -0.65 | AR-V7 |
| TMED5 | 0.91 | 0.42 | AR-FL | KIF16B | -0.43 | -0.64 | AR-V7 |
| FOXO3 | 0.75 | 0.42 | AR-FL | CEP170 | -0.23 | -0.64 | AR-V7 |
| SYVN1 | 0.86 | 0.43 | AR-FL | RB1CC1 | -0.55 | -0.64 | AR-V7 |
| ATAD2 | 0.87 | 0.43 | AR-FL | ACSS1 | -0.50 | -0.64 | AR-V7 |
| SLC25A42 | 0.68 | 0.43 | AR-FL | SLC27A2 | -0.51 | -0.64 | AR-V7 |
| SLC35B1 | 0.65 | 0.43 | AR-FL | FGF12 | -0.35 | -0.64 | AR-V7 |
| PDIA3 | 0.75 | 0.43 | AR-FL | NEDD4L | -0.33 | -0.64 | AR-V7 |
| TMEM87A | 0.91 | 0.43 | AR-FL | ZNF717 | -0.53 | -0.64 | AR-V7 |
| SELK | 0.77 | 0.43 | AR-FL | HOXC4 | -0.55 | -0.64 | AR-V7 |
| CYB561D2 | 0.73 | 0.43 | AR-FL | LIPH | -0.01 | -0.64 | AR-V7 |
| CDK2AP2 | 0.68 | 0.43 | AR-FL | FOXP1 | -0.35 | -0.64 | AR-V7 |
| PNKD | 0.96 | 0.43 | AR-FL | STX16-NPEPL1 | -0.51 | -0.64 | AR-V7 |
| BET1 | 0.68 | 0.43 | AR-FL | SCCPDH | -0.46 | -0.64 | AR-V7 |
| NDFIP2 | 1.72 | 0.43 | AR-FL | IRF1 | -0.55 | -0.63 | AR-V7 |
| AGAP4 | 1.63 | 0.43 | AR-FL | B3GNT4 | -0.53 | -0.63 | AR-V7 |
| MARCKS | 0.70 | 0.43 | AR-FL | PCDHA10 | -0.17 | -0.63 | AR-V7 |
| TRADD | 0.99 | 0.44 | AR-FL | KLLN | -0.57 | -0.63 | AR-V7 |
| GANC | 0.66 | 0.44 | AR-FL | BICC1 | -0.44 | -0.62 | AR-V7 |
| PHF8 | 0.85 | 0.44 | AR-FL | PKIA | -0.53 | -0.62 | AR-V7 |
| PARG | 0.92 | 0.44 | AR-FL | SSC4D | -0.55 | -0.62 | AR-V7 |
| CORO2A | 1.25 | 0.44 | AR-FL | RAB27B | -0.42 | -0.62 | AR-V7 |
| PTRH2 | 0.72 | 0.44 | AR-FL | MIA2 | -0.27 | -0.62 | AR-V7 |
| ZSWIM8 | 0.59 | 0.44 | AR-FL | GLIS2 | -0.52 | -0.62 | AR-V7 |
| CCPG1 | 0.79 | 0.44 | AR-FL | SIAH1 | -0.21 | -0.61 | AR-V7 |
| SLC39A6 | 0.72 | 0.44 | AR-FL | PLEKHF2 | -0.14 | -0.61 | AR-V7 |
| XPNPEP1 | 0.65 | 0.44 | AR-FL | LRTM2 | -0.52 | -0.61 | AR-V7 |
| RCAN1 | 0.98 | 0.45 | AR-FL | DOCK9 | -0.17 | -0.61 | AR-V7 |
| CNPY3 | 0.95 | 0.45 | AR-FL | UTRN | -0.46 | -0.61 | AR-V7 |
| ZNF532 | 0.74 | 0.45 | AR-FL | ELF3 | -0.06 | -0.61 | AR-V7 |
| YWHAZ | 0.63 | 0.45 | AR-FL | KLHL7 | -0.10 | -0.61 | AR-V7 |
| GFPT1 | 0.97 | 0.45 | AR-FL | MAP4K4 | -0.51 | -0.60 | AR-V7 |
| FNBP1L | 1.11 | 0.45 | AR-FL | HDAC11 | -0.55 | -0.60 | AR-V7 |

| | | | |
|---|---|---|---|
| CHP1 | 0.65 | 0.45 | AR-FL |
| AFF4 | 0.75 | 0.45 | AR-FL |
| OCRL | 0.59 | 0.46 | AR-FL |
| ARSG | 1.56 | 0.46 | AR-FL |
| CAMSAP2 | 0.88 | 0.46 | AR-FL |
| RPS6KC1 | 0.74 | 0.46 | AR-FL |
| NRG4 | 1.01 | 0.46 | AR-FL |
| RNF103 | 0.70 | 0.46 | AR-FL |
| PJA2 | 0.78 | 0.46 | AR-FL |
| CYP46A1 | 0.68 | 0.46 | AR-FL |
| MRPS18A | 1.31 | 0.46 | AR-FL |
| TRIM24 | 0.60 | 0.46 | AR-FL |
| SEC61B | 0.81 | 0.46 | AR-FL |
| CCDC40 | 0.65 | 0.47 | AR-FL |
| ITPKC | 0.81 | 0.47 | AR-FL |
| MCEE | 0.90 | 0.47 | AR-FL |
| DYNLRB2 | 0.88 | 0.47 | AR-FL |
| PPM1K | 1.18 | 0.47 | AR-FL |
| ARID5B | 1.32 | 0.47 | AR-FL |
| ELMO2 | 1.22 | 0.47 | AR-FL |
| SPDEF | 2.30 | 0.47 | AR-FL |
| B3GNT2 | 0.74 | 0.47 | AR-FL |
| RAB43 | 0.70 | 0.47 | AR-FL |
| ZDHHC9 | 1.05 | 0.48 | AR-FL |
| ABCD3 | 0.65 | 0.48 | AR-FL |
| FBXO42 | 0.63 | 0.48 | AR-FL |
| ARHGAP1 | 0.80 | 0.48 | AR-FL |
| CREB3 | 0.73 | 0.48 | AR-FL |
| MTMR11 | 1.51 | 0.48 | AR-FL |
| USO1 | 0.79 | 0.48 | AR-FL |
| RAB27A | 1.98 | 0.48 | AR-FL |
| SEC24B | 1.32 | 0.48 | AR-FL |
| RDH11 | 0.60 | 0.49 | AR-FL |
| TXNDC5 | 0.75 | 0.49 | AR-FL |
| 2-Mar | 0.68 | 0.49 | AR-FL |
| CRACR2A | 2.24 | 0.49 | AR-FL |
| MORC4 | 0.78 | 0.49 | AR-FL |
| CRISPLD2 | 1.46 | 0.50 | AR-FL |
| COPB2 | 0.76 | 0.50 | AR-FL |
| CAPZB | 1.27 | 0.50 | AR-FL |
| SURF4 | 0.85 | 0.50 | AR-FL |
| SEC24A | 1.01 | 0.50 | AR-FL |
| CCDC90B | 0.60 | 0.51 | AR-FL |
| MPC2 | 1.70 | 0.51 | AR-FL |
| UCN | 0.68 | 0.51 | AR-FL |
| TRAM1 | 1.02 | 0.51 | AR-FL |

| | | | |
|---|---|---|---|
| EEFSEC | -0.54 | -0.60 | AR-V7 |
| QPCT | -0.26 | -0.60 | AR-V7 |
| NRN1 | -0.51 | -0.60 | AR-V7 |
| TMC5 | -0.34 | -0.60 | AR-V7 |
| LBH | -0.16 | -0.60 | AR-V7 |
| CD2AP | -0.41 | -0.60 | AR-V7 |
| RNF130 | -0.51 | -0.60 | AR-V7 |
| PERP | -0.52 | -0.60 | AR-V7 |
| FYCO1 | -0.31 | -0.60 | AR-V7 |
| AMOTL2 | -0.55 | -0.60 | AR-V7 |
| RAB30 | -0.33 | -0.59 | AR-V7 |
| POTEJ | -0.51 | -0.59 | AR-V7 |
| SYF2 | -0.51 | -0.59 | AR-V7 |
| DDI2 | -0.43 | -0.59 | AR-V7 |
| KCNT1 | -0.42 | -0.59 | AR-V7 |
| NYAP1 | -0.01 | -0.59 | AR-V7 |
| PCDHB13 | -0.33 | -0.59 | AR-V7 |
| PBXIP1 | -0.15 | -0.59 | AR-V7 |
| SIX4 | -0.56 | -0.59 | AR-V7 |
| GAREM | -0.49 | -0.59 | AR-V7 |
| DENND4C | -0.22 | -0.58 | AR-V7 |
| KRAS | -0.40 | -0.58 | AR-V7 |
| PCDHB9 | -0.31 | -0.58 | AR-V7 |
| COQ10A | -0.57 | -0.58 | AR-V7 |
| FRAT2 | -0.15 | -0.58 | AR-V7 |
| PIGQ | 0.44 | 0.58 | AR-V7 |
| DHTKD1 | 0.44 | 0.58 | AR-V7 |
| PYGB | 0.55 | 0.58 | AR-V7 |
| FBXL6 | 0.49 | 0.58 | AR-V7 |
| ACO2 | 0.56 | 0.58 | AR-V7 |
| CDCA3 | 0.13 | 0.58 | AR-V7 |
| NFE2L1 | 0.45 | 0.58 | AR-V7 |
| MMS22L | 0.01 | 0.58 | AR-V7 |
| HIF1A | 0.52 | 0.58 | AR-V7 |
| QPCTL | 0.46 | 0.59 | AR-V7 |
| LMBR1 | 0.10 | 0.59 | AR-V7 |
| U2AF1L4 | 0.12 | 0.59 | AR-V7 |
| MYH14 | 0.11 | 0.59 | AR-V7 |
| CDK11B | 0.50 | 0.59 | AR-V7 |
| BRI3BP | 0.38 | 0.59 | AR-V7 |
| SH2D5 | 0.04 | 0.59 | AR-V7 |
| TST | 0.18 | 0.59 | AR-V7 |
| PIP5K1C | 0.49 | 0.59 | AR-V7 |
| TTC3 | 0.24 | 0.59 | AR-V7 |
| ABCD3 | 0.04 | 0.59 | AR-V7 |
| FAH | 0.17 | 0.59 | AR-V7 |

| | | | |
|---|---|---|---|
| RALY | 0.62 | 0.51 | AR-FL |
| MYDGF | 0.92 | 0.51 | AR-FL |
| FRYL | 1.35 | 0.51 | AR-FL |
| CSNK1G3 | 0.61 | 0.51 | AR-FL |
| ARMCX3 | 0.77 | 0.51 | AR-FL |
| TTN | 1.19 | 0.51 | AR-FL |
| AADAT | 1.26 | 0.52 | AR-FL |
| P4HB | 0.65 | 0.52 | AR-FL |
| MYL9 | 2.24 | 0.52 | AR-FL |
| GBF1 | 0.69 | 0.52 | AR-FL |
| C9orf72 | 1.17 | 0.52 | AR-FL |
| PPP1R15B | 0.72 | 0.52 | AR-FL |
| APIP | 0.68 | 0.52 | AR-FL |
| FBXW11 | 0.67 | 0.52 | AR-FL |
| PAQR6 | 1.18 | 0.52 | AR-FL |
| MAP3K6 | 0.92 | 0.52 | AR-FL |
| EDEM1 | 1.28 | 0.52 | AR-FL |
| AMACR | 1.73 | 0.53 | AR-FL |
| APPBP2 | 1.95 | 0.53 | AR-FL |
| NKX3-2 | 1.29 | 0.53 | AR-FL |
| C15orf65 | 1.06 | 0.53 | AR-FL |
| SRP19 | 1.00 | 0.53 | AR-FL |
| RBM24 | 0.87 | 0.53 | AR-FL |
| SLC30A7 | 0.91 | 0.53 | AR-FL |
| TMCO3 | 0.61 | 0.53 | AR-FL |
| IDH1 | 1.04 | 0.54 | AR-FL |
| MAGT1 | 0.61 | 0.54 | AR-FL |
| ANXA5 | 0.67 | 0.54 | AR-FL |
| PDIA4 | 0.84 | 0.54 | AR-FL |
| MAP3K5 | 0.66 | 0.54 | AR-FL |
| PRKCA | 2.58 | 0.54 | AR-FL |
| STARD3NL | 0.85 | 0.54 | AR-FL |
| ERP44 | 0.91 | 0.54 | AR-FL |
| GOLGB1 | 1.04 | 0.54 | AR-FL |
| LMAN1 | 1.18 | 0.55 | AR-FL |
| SDF2L1 | 0.89 | 0.55 | AR-FL |
| MCFD2 | 0.84 | 0.55 | AR-FL |
| ERN1 | 2.01 | 0.55 | AR-FL |
| DHTKD1 | 0.66 | 0.55 | AR-FL |
| ZNF235 | 0.84 | 0.55 | AR-FL |
| OSTC | 0.98 | 0.55 | AR-FL |
| CSGALNACT1 | 5.08 | 0.55 | AR-FL |
| MAGED2 | 0.59 | 0.55 | AR-FL |
| CAMKK2 | 1.62 | 0.55 | AR-FL |
| UFM1 | 0.60 | 0.55 | AR-FL |
| SLC50A1 | 0.68 | 0.55 | AR-FL |
| PLK4 | 0.09 | 0.60 | AR-V7 |
| ARAF | 0.21 | 0.60 | AR-V7 |
| GLUD2 | 0.37 | 0.60 | AR-V7 |
| WDFY1 | 0.44 | 0.60 | AR-V7 |
| MDM1 | 0.18 | 0.60 | AR-V7 |
| ZNF778 | 0.56 | 0.60 | AR-V7 |
| MED30 | 0.19 | 0.60 | AR-V7 |
| VTN | 0.18 | 0.60 | AR-V7 |
| TADA2B | 0.34 | 0.60 | AR-V7 |
| KIF18B | 0.07 | 0.60 | AR-V7 |
| ACAP3 | 0.43 | 0.60 | AR-V7 |
| TNFRSF10B | 0.40 | 0.61 | AR-V7 |
| TMEM175 | 0.39 | 0.61 | AR-V7 |
| TM2D2 | 0.52 | 0.61 | AR-V7 |
| NCKIPSD | 0.10 | 0.61 | AR-V7 |
| ALKBH6 | 0.33 | 0.61 | AR-V7 |
| SLC2A4RG | 0.40 | 0.61 | AR-V7 |
| C9orf142 | 0.50 | 0.61 | AR-V7 |
| VWA5B2 | 0.47 | 0.61 | AR-V7 |
| ANKFY1 | 0.46 | 0.62 | AR-V7 |
| BTG3 | 0.26 | 0.62 | AR-V7 |
| NEMP2 | 0.01 | 0.62 | AR-V7 |
| ZNF184 | 0.40 | 0.62 | AR-V7 |
| ARL3 | 0.19 | 0.62 | AR-V7 |
| RBM47 | 0.54 | 0.62 | AR-V7 |
| DCDC2 | 0.17 | 0.62 | AR-V7 |
| PAGR1 | 0.31 | 0.62 | AR-V7 |
| MROH1 | 0.48 | 0.62 | AR-V7 |
| PRAME | 0.16 | 0.62 | AR-V7 |
| CCDC86 | 0.40 | 0.62 | AR-V7 |
| TRAF4 | 0.52 | 0.62 | AR-V7 |
| KLHL36 | 0.22 | 0.62 | AR-V7 |
| GYLTL1B | 0.07 | 0.63 | AR-V7 |
| RHOC | 0.33 | 0.63 | AR-V7 |
| ANKRD40 | 0.14 | 0.63 | AR-V7 |
| TMEM164 | 0.15 | 0.63 | AR-V7 |
| FBXO9 | 0.22 | 0.63 | AR-V7 |
| TMEM234 | 0.34 | 0.63 | AR-V7 |
| WDR13 | 0.51 | 0.63 | AR-V7 |
| PPP4R3A | 0.43 | 0.63 | AR-V7 |
| IQCK | 0.50 | 0.64 | AR-V7 |
| CCDC157 | 0.47 | 0.64 | AR-V7 |
| ALDH3B2 | 0.02 | 0.64 | AR-V7 |
| GRIPAP1 | 0.34 | 0.64 | AR-V7 |
| TK2 | 0.48 | 0.64 | AR-V7 |
| FIGNL1 | 0.44 | 0.64 | AR-V7 |

| | | | | | | | |
|---|---|---|---|---|---|---|---|
| FAR2 | 0.79 | 0.56 | AR-FL | TP73 | 0.11 | 0.64 | AR-V7 |
| HSPA5 | 0.69 | 0.56 | AR-FL | ZSCAN1 | 0.09 | 0.64 | AR-V7 |
| MGLL | 0.99 | 0.56 | AR-FL | SLC25A29 | 0.26 | 0.64 | AR-V7 |
| RPGRIP1 | 1.03 | 0.56 | AR-FL | RFC2 | 0.55 | 0.64 | AR-V7 |
| CTTN | 0.83 | 0.56 | AR-FL | EZH2 | 0.44 | 0.64 | AR-V7 |
| SSR2 | 0.97 | 0.56 | AR-FL | CENPP | 0.07 | 0.64 | AR-V7 |
| DFNB31 | 0.69 | 0.56 | AR-FL | NAIP | 0.13 | 0.64 | AR-V7 |
| RNF225 | 0.85 | 0.56 | AR-FL | NOS1AP | 0.49 | 0.65 | AR-V7 |
| HERPUD1 | 1.05 | 0.56 | AR-FL | PIGBOS1 | 0.41 | 0.65 | AR-V7 |
| EFNA5 | 1.31 | 0.57 | AR-FL | DYNLL1 | 0.57 | 0.65 | AR-V7 |
| FAM118B | 0.73 | 0.57 | AR-FL | PPP5C | 0.47 | 0.65 | AR-V7 |
| CCNYL1 | 0.74 | 0.57 | AR-FL | VRK1 | 0.52 | 0.65 | AR-V7 |
| CSNK1A1 | 0.72 | 0.57 | AR-FL | C4orf33 | 0.30 | 0.65 | AR-V7 |
| SEC61G | 0.90 | 0.57 | AR-FL | GALK1 | 0.12 | 0.65 | AR-V7 |
| FBXW2 | 1.16 | 0.57 | AR-FL | SRPK2 | 0.11 | 0.66 | AR-V7 |
| FOXO1 | 1.16 | 0.57 | AR-FL | AMOTL1 | 0.39 | 0.66 | AR-V7 |
| SEC11C | 1.30 | 0.57 | AR-FL | LMNB1 | 0.48 | 0.66 | AR-V7 |
| TMEM120A | 0.62 | 0.57 | AR-FL | STRADA | 0.43 | 0.66 | AR-V7 |
| BTN3A1 | 1.04 | 0.58 | AR-FL | ZNHIT1 | 0.53 | 0.66 | AR-V7 |
| YIPF5 | 0.66 | 0.58 | AR-FL | GOT2 | 0.41 | 0.66 | AR-V7 |
| WDYHV1 | 1.22 | 0.58 | AR-FL | ULK4 | 0.16 | 0.66 | AR-V7 |
| TMED7 | 0.88 | 0.58 | Both Isoforms | PRKX | 0.47 | 0.66 | AR-V7 |
| AKAP1 | 0.64 | 0.58 | Both Isoforms | TMSB10 | 0.42 | 0.66 | AR-V7 |
| ADH1C | 3.95 | 0.58 | Both Isoforms | AADAT | 0.52 | 0.66 | AR-V7 |
| CKMT1B | 0.81 | 0.59 | Both Isoforms | C12orf4 | 0.20 | 0.66 | AR-V7 |
| SEL1L | 0.60 | 0.59 | Both Isoforms | CHPT1 | 0.53 | 0.66 | AR-V7 |
| MGAT2 | 0.69 | 0.59 | Both Isoforms | SKA1 | 0.27 | 0.66 | AR-V7 |
| BET1L | 0.64 | 0.59 | Both Isoforms | TMEM39A | 0.53 | 0.67 | AR-V7 |
| COQ10B | 0.70 | 0.59 | Both Isoforms | CDKN1C | 0.39 | 0.67 | AR-V7 |
| RIMBP3 | 0.96 | 0.59 | Both Isoforms | IQCG | 0.39 | 0.67 | AR-V7 |
| MAP9 | 1.23 | 0.59 | Both Isoforms | TMEM144 | 0.47 | 0.67 | AR-V7 |
| GMPPA | 1.10 | 0.59 | Both Isoforms | KLHL26 | 0.27 | 0.67 | AR-V7 |
| ZNF432 | 0.78 | 0.60 | Both Isoforms | GLUD1 | 0.56 | 0.67 | AR-V7 |
| OPN1SW | 1.33 | 0.60 | Both Isoforms | NCAPG2 | 0.40 | 0.67 | AR-V7 |

| | | | | | | | |
|---|---|---|---|---|---|---|---|
| GOSR2 | 0.63 | 0.60 | Both Isoforms | C19orf71 | 0.15 | 0.67 | AR-V7 |
| OAT | 0.89 | 0.60 | Both Isoforms | LPIN1 | 0.27 | 0.67 | AR-V7 |
| KIF13B | 1.23 | 0.60 | Both Isoforms | PQLC1 | 0.27 | 0.68 | AR-V7 |
| SLC10A7 | 1.68 | 0.60 | Both Isoforms | OPLAH | 0.35 | 0.68 | AR-V7 |
| PIGH | 0.66 | 0.60 | Both Isoforms | SPA17 | 0.50 | 0.69 | AR-V7 |
| ARFIP2 | 0.97 | 0.60 | Both Isoforms | SLC43A3 | 0.30 | 0.69 | AR-V7 |
| SDCBP | 0.94 | 0.60 | Both Isoforms | DUSP3 | 0.49 | 0.69 | AR-V7 |
| PTPRM | 1.99 | 0.61 | Both Isoforms | SUV39H1 | 0.45 | 0.70 | AR-V7 |
| KLF4 | 1.14 | 0.61 | Both Isoforms | RBM34 | 0.06 | 0.70 | AR-V7 |
| TCAF1 | 1.56 | 0.61 | Both Isoforms | TMSB4Y | 0.30 | 0.70 | AR-V7 |
| MCCC2 | 0.99 | 0.61 | Both Isoforms | MYCBP | 0.56 | 0.70 | AR-V7 |
| HIST1H2BE | 0.60 | 0.61 | Both Isoforms | PXYLP1 | 0.00 | 0.70 | AR-V7 |
| ZNF587 | 0.75 | 0.61 | Both Isoforms | TRMT2B | 0.52 | 0.70 | AR-V7 |
| CALU | 1.19 | 0.61 | Both Isoforms | THOC2 | 0.55 | 0.71 | AR-V7 |
| ANXA4 | 0.59 | 0.61 | Both Isoforms | CDC14B | 0.29 | 0.71 | AR-V7 |
| CHPF2 | 0.61 | 0.61 | Both Isoforms | TPM1 | 0.46 | 0.71 | AR-V7 |
| HES6 | 1.98 | 0.61 | Both Isoforms | PPP1R16A | 0.45 | 0.71 | AR-V7 |
| MAFK | 0.71 | 0.61 | Both Isoforms | MOK | 0.18 | 0.72 | AR-V7 |
| TVP23C | 0.84 | 0.61 | Both Isoforms | CBS | 0.20 | 0.72 | AR-V7 |
| LRG1 | 0.84 | 0.62 | Both Isoforms | FAM217B | 0.08 | 0.72 | AR-V7 |
| FKBP11 | 0.96 | 0.62 | Both Isoforms | FLNC | 0.26 | 0.72 | AR-V7 |
| WDR44 | 0.91 | 0.62 | Both Isoforms | NCAPH2 | 0.38 | 0.72 | AR-V7 |
| SLC25A16 | 0.78 | 0.62 | Both Isoforms | CAPN2 | 0.08 | 0.72 | AR-V7 |
| TRIM13 | 1.18 | 0.62 | Both Isoforms | ZGRF1 | 0.11 | 0.72 | AR-V7 |
| GPR75 | 0.61 | 0.62 | Both Isoforms | SRSF2 | 0.28 | 0.73 | AR-V7 |

| | | | | | | | |
|---|---|---|---|---|---|---|---|
| AKAP13 | 1.05 | 0.62 | Both Isoforms | TRAIP | 0.17 | 0.73 | AR-V7 |
| AP1S2 | 0.73 | 0.62 | Both Isoforms | NASP | 0.05 | 0.73 | AR-V7 |
| PDXDC1 | 0.64 | 0.62 | Both Isoforms | TXNDC16 | 0.17 | 0.73 | AR-V7 |
| RRBP1 | 1.08 | 0.63 | Both Isoforms | EIF4EBP2 | 0.29 | 0.73 | AR-V7 |
| RTN2 | 0.70 | 0.63 | Both Isoforms | SPATA33 | 0.54 | 0.73 | AR-V7 |
| FABP3 | 0.64 | 0.63 | Both Isoforms | C1orf226 | 0.57 | 0.73 | AR-V7 |
| FAM19A4 | 1.64 | 0.63 | Both Isoforms | HIST1H1E | 0.34 | 0.73 | AR-V7 |
| GOLGA2 | 0.88 | 0.63 | Both Isoforms | HHEX | 0.13 | 0.73 | AR-V7 |
| SEC13 | 0.69 | 0.63 | Both Isoforms | NRGN | 0.07 | 0.73 | AR-V7 |
| ELOVL1 | 0.70 | 0.64 | Both Isoforms | FBXO36 | 0.48 | 0.74 | AR-V7 |
| GOLGA4 | 1.01 | 0.64 | Both Isoforms | TRIP13 | 0.29 | 0.74 | AR-V7 |
| EXOC8 | 0.82 | 0.64 | Both Isoforms | SERF1B | 0.10 | 0.74 | AR-V7 |
| KLF5 | 2.88 | 0.64 | Both Isoforms | STARD10 | 0.52 | 0.74 | AR-V7 |
| FAM46C | 0.65 | 0.64 | Both Isoforms | ENTPD3 | 0.32 | 0.74 | AR-V7 |
| DQX1 | 0.78 | 0.64 | Both Isoforms | TMPPE | 0.54 | 0.75 | AR-V7 |
| ID2 | 0.80 | 0.65 | Both Isoforms | TBCEL | 0.22 | 0.75 | AR-V7 |
| FEM1C | 0.66 | 0.65 | Both Isoforms | SIVA1 | 0.45 | 0.75 | AR-V7 |
| HARBI1 | 0.70 | 0.65 | Both Isoforms | SPRYD7 | 0.35 | 0.75 | AR-V7 |
| ADPRM | 1.33 | 0.65 | Both Isoforms | RGMA | 0.04 | 0.75 | AR-V7 |
| USP38 | 0.69 | 0.65 | Both Isoforms | KREMEN2 | 0.40 | 0.75 | AR-V7 |
| PDS5B | 0.84 | 0.65 | Both Isoforms | WDHD1 | 0.53 | 0.76 | AR-V7 |
| INHBC | 1.37 | 0.65 | Both Isoforms | TICRR | 0.04 | 0.76 | AR-V7 |
| LIG4 | 0.61 | 0.66 | Both Isoforms | NRSN2 | 0.30 | 0.76 | AR-V7 |
| ADAM9 | 1.00 | 0.66 | Both Isoforms | INPP1 | 0.12 | 0.76 | AR-V7 |
| GOLT1B | 0.76 | 0.66 | Both Isoforms | NARFL | 0.47 | 0.77 | AR-V7 |

| | | | | | | | |
|---|---|---|---|---|---|---|---|
| USP35 | 0.59 | 0.66 | Both Isoforms | NUDT1 | 0.57 | 0.77 | AR-V7 |
| MON2 | 0.64 | 0.66 | Both Isoforms | ZMAT3 | 0.46 | 0.77 | AR-V7 |
| TCEAL3 | 0.70 | 0.66 | Both Isoforms | KIF24 | 0.17 | 0.77 | AR-V7 |
| TRAFD1 | 0.82 | 0.66 | Both Isoforms | AKAP1 | 0.49 | 0.77 | AR-V7 |
| KLF7 | 0.59 | 0.67 | Both Isoforms | MBD2 | 0.52 | 0.77 | AR-V7 |
| PICK1 | 0.66 | 0.67 | Both Isoforms | PXMP2 | 0.53 | 0.77 | AR-V7 |
| LMLN | 0.60 | 0.67 | Both Isoforms | ADGRB2 | 0.11 | 0.78 | AR-V7 |
| PPIB | 0.78 | 0.67 | Both Isoforms | RNASEH2A | 0.37 | 0.78 | AR-V7 |
| C17orf107 | 0.62 | 0.67 | Both Isoforms | TMEM201 | 0.26 | 0.78 | AR-V7 |
| TIMM10B | 0.85 | 0.67 | Both Isoforms | TBC1D20 | 0.48 | 0.78 | AR-V7 |
| C19orf48 | 0.94 | 0.67 | Both Isoforms | C15orf59 | 0.11 | 0.78 | AR-V7 |
| TMED9 | 1.03 | 0.68 | Both Isoforms | ZNF527 | 0.15 | 0.78 | AR-V7 |
| ENTPD7 | 1.08 | 0.69 | Both Isoforms | CNN2 | 0.57 | 0.79 | AR-V7 |
| ORMDL2 | 0.73 | 0.69 | Both Isoforms | RAB15 | 0.20 | 0.80 | AR-V7 |
| TRMT6 | 0.61 | 0.69 | Both Isoforms | GPR137C | 0.44 | 0.80 | AR-V7 |
| RHOBTB2 | 1.00 | 0.70 | Both Isoforms | ASB16 | 0.47 | 0.80 | AR-V7 |
| HUS1 | 0.90 | 0.70 | Both Isoforms | PMEPA1 | 0.48 | 0.80 | AR-V7 |
| CUL4B | 0.84 | 0.70 | Both Isoforms | DPCD | 0.34 | 0.80 | AR-V7 |
| ENTPD4 | 0.61 | 0.70 | Both Isoforms | NPHP1 | 0.15 | 0.80 | AR-V7 |
| FRY | 1.56 | 0.70 | Both Isoforms | POLQ | 0.18 | 0.80 | AR-V7 |
| MPHOSPH9 | 1.66 | 0.70 | Both Isoforms | SERPINF1 | 0.35 | 0.80 | AR-V7 |
| LYPLAL1 | 1.04 | 0.71 | Both Isoforms | RFC4 | 0.27 | 0.80 | AR-V7 |
| FXYD3 | 1.10 | 0.71 | Both Isoforms | PC | 0.45 | 0.80 | AR-V7 |
| SYP | 0.60 | 0.71 | Both Isoforms | CDCA4 | 0.03 | 0.81 | AR-V7 |
| CMC2 | 1.26 | 0.71 | Both Isoforms | DSN1 | 0.51 | 0.81 | AR-V7 |

| | | | | | | | |
|---|---|---|---|---|---|---|---|
| LIMCH1 | 1.46 | 0.71 | Both Isoforms | RBL1 | 0.30 | 0.81 | AR-V7 |
| RPS6KA3 | 0.92 | 0.71 | Both Isoforms | MCM2 | 0.42 | 0.81 | AR-V7 |
| FBXO16 | 0.65 | 0.71 | Both Isoforms | RFC3 | 0.27 | 0.81 | AR-V7 |
| TMEM39A | 0.85 | 0.71 | Both Isoforms | CDCA5 | 0.18 | 0.81 | AR-V7 |
| PLPP5 | 1.39 | 0.71 | Both Isoforms | CUX2 | 0.42 | 0.82 | AR-V7 |
| NPC1 | 1.51 | 0.71 | Both Isoforms | ERVK3-1 | 0.07 | 0.82 | AR-V7 |
| FAM69A | 1.47 | 0.71 | Both Isoforms | FKBP1C | 0.21 | 0.82 | AR-V7 |
| MANF | 1.00 | 0.71 | Both Isoforms | NOMO3 | 0.12 | 0.82 | AR-V7 |
| ZNF449 | 0.94 | 0.72 | Both Isoforms | PITPNM3 | 0.14 | 0.82 | AR-V7 |
| PLEKHM3 | 0.87 | 0.72 | Both Isoforms | DDTL | 0.30 | 0.82 | AR-V7 |
| AFAP1L2 | 1.21 | 0.72 | Both Isoforms | 8-Sep | 0.26 | 0.82 | AR-V7 |
| C2orf76 | 1.40 | 0.72 | Both Isoforms | DNAH9 | 0.04 | 0.82 | AR-V7 |
| FZD5 | 2.48 | 0.72 | Both Isoforms | POLD3 | 0.49 | 0.83 | AR-V7 |
| PFKFB2 | 1.08 | 0.73 | Both Isoforms | TRABD | 0.12 | 0.83 | AR-V7 |
| CPEB3 | 1.76 | 0.73 | Both Isoforms | CENPU | 0.40 | 0.83 | AR-V7 |
| FAM13A | 1.31 | 0.73 | Both Isoforms | HRH3 | 0.10 | 0.83 | AR-V7 |
| HSD17B4 | 0.65 | 0.73 | Both Isoforms | UBXN11 | 0.17 | 0.83 | AR-V7 |
| SERF1B | 0.93 | 0.74 | Both Isoforms | EPAS1 | 0.07 | 0.84 | AR-V7 |
| WDFY2 | 0.83 | 0.74 | Both Isoforms | ARHGAP31 | 0.34 | 0.84 | AR-V7 |
| SNAPC1 | 0.80 | 0.74 | Both Isoforms | CCDC69 | 0.40 | 0.84 | AR-V7 |
| ANKRD37 | 2.13 | 0.74 | Both Isoforms | KANK2 | 0.27 | 0.85 | AR-V7 |
| TG | 4.73 | 0.74 | Both Isoforms | TOE1 | 0.33 | 0.86 | AR-V7 |
| RAB29 | 0.59 | 0.74 | Both Isoforms | NCOR1 | 0.21 | 0.86 | AR-V7 |
| KCTD21 | 0.85 | 0.74 | Both Isoforms | SPC25 | 0.10 | 0.86 | AR-V7 |
| ALDH6A1 | 0.76 | 0.75 | Both Isoforms | FANCI | 0.47 | 0.87 | AR-V7 |

| | | | | | | | |
|---|---|---|---|---|---|---|---|
| ZBTB1 | 0.77 | 0.75 | Both Isoforms | ZNF652 | 0.54 | 0.87 | AR-V7 |
| SGK3 | 1.25 | 0.75 | Both Isoforms | RIT1 | 0.43 | 0.87 | AR-V7 |
| CREB3L2 | 1.08 | 0.75 | Both Isoforms | RFC5 | 0.55 | 0.87 | AR-V7 |
| TBC1D8 | 2.05 | 0.75 | Both Isoforms | STXBP5 | 0.42 | 0.87 | AR-V7 |
| ELF2 | 0.87 | 0.75 | Both Isoforms | ATAD5 | 0.39 | 0.88 | AR-V7 |
| WWTR1 | 3.69 | 0.75 | Both Isoforms | RGPD6 | 0.02 | 0.88 | AR-V7 |
| PPP2CB | 1.08 | 0.75 | Both Isoforms | CCND1 | 0.58 | 0.88 | AR-V7 |
| YY2 | 0.78 | 0.75 | Both Isoforms | SHISA3 | 0.15 | 0.88 | AR-V7 |
| RASSF3 | 1.29 | 0.75 | Both Isoforms | CENPK | 0.39 | 0.89 | AR-V7 |
| PRR34 | 0.98 | 0.76 | Both Isoforms | FANCD2 | 0.52 | 0.89 | AR-V7 |
| ZNF587B | 0.91 | 0.76 | Both Isoforms | BAIAP3 | 0.15 | 0.89 | AR-V7 |
| ALOX12B | 2.09 | 0.76 | Both Isoforms | TMEM60 | 0.08 | 0.89 | AR-V7 |
| POLD4 | 0.70 | 0.76 | Both Isoforms | COMMD8 | 0.51 | 0.89 | AR-V7 |
| MTMR9 | 2.23 | 0.77 | Both Isoforms | GTPBP2 | 0.49 | 0.90 | AR-V7 |
| DISP2 | 0.69 | 0.77 | Both Isoforms | CTNNBIP1 | 0.25 | 0.90 | AR-V7 |
| HES2 | 2.84 | 0.77 | Both Isoforms | RPS6KL1 | 0.39 | 0.91 | AR-V7 |
| WDR1 | 0.74 | 0.77 | Both Isoforms | C17orf53 | 0.46 | 0.92 | AR-V7 |
| HGD | 1.77 | 0.77 | Both Isoforms | DNA2 | 0.23 | 0.93 | AR-V7 |
| TRO | 0.58 | 0.77 | Both Isoforms | ACADM | 0.29 | 0.93 | AR-V7 |
| VIMP | 1.25 | 0.77 | Both Isoforms | CKAP2 | 0.39 | 0.93 | AR-V7 |
| CLDN12 | 0.69 | 0.78 | Both Isoforms | BRIP1 | 0.27 | 0.93 | AR-V7 |
| GLYATL1 | 1.77 | 0.78 | Both Isoforms | CENPI | 0.42 | 0.93 | AR-V7 |
| HIST1H4H | 0.72 | 0.78 | Both Isoforms | CCBL1 | 0.46 | 0.94 | AR-V7 |
| PACSIN2 | 0.94 | 0.78 | Both Isoforms | TONSL | 0.51 | 0.94 | AR-V7 |
| GLYATL1P3 | 1.58 | 0.78 | Both Isoforms | FAM46C | 0.31 | 0.94 | AR-V7 |

| | | | | | | | |
|---|---|---|---|---|---|---|---|
| FRK | 1.88 | 0.79 | Both Isoforms | KRTCAP2 | 0.09 | 0.95 | AR-V7 |
| DNAJC1 | 0.66 | 0.79 | Both Isoforms | GAMT | 0.43 | 0.95 | AR-V7 |
| HERC4 | 0.73 | 0.79 | Both Isoforms | VCL | 0.42 | 0.95 | AR-V7 |
| FAM177A1 | 1.13 | 0.79 | Both Isoforms | NPHP4 | 0.24 | 0.96 | AR-V7 |
| CDR2 | 0.60 | 0.80 | Both Isoforms | MSX1 | 0.42 | 0.97 | AR-V7 |
| GCNT1 | 1.31 | 0.80 | Both Isoforms | POLD1 | 0.29 | 0.97 | AR-V7 |
| GNRH1 | 0.85 | 0.80 | Both Isoforms | XRCC2 | 0.33 | 0.97 | AR-V7 |
| IL17RA | 0.60 | 0.81 | Both Isoforms | KNTC1 | 0.07 | 0.98 | AR-V7 |
| ZNF589 | 0.97 | 0.81 | Both Isoforms | MAGI2 | 0.11 | 0.98 | AR-V7 |
| KLHL36 | 0.84 | 0.81 | Both Isoforms | ROR1 | 0.45 | 0.99 | AR-V7 |
| SCD | 0.62 | 0.81 | Both Isoforms | EYA1 | 0.18 | 0.99 | AR-V7 |
| HIST1H2BD | 0.63 | 0.81 | Both Isoforms | KCNC4 | 0.06 | 1.00 | AR-V7 |
| HMGCR | 1.55 | 0.82 | Both Isoforms | KIAA0101 | 0.42 | 1.00 | AR-V7 |
| CEP162 | 0.83 | 0.82 | Both Isoforms | APOD | 0.14 | 1.00 | AR-V7 |
| ZFP36 | 0.74 | 0.82 | Both Isoforms | STARD13 | 0.31 | 1.00 | AR-V7 |
| HIBCH | 0.90 | 0.82 | Both Isoforms | CBX4 | 0.58 | 1.00 | AR-V7 |
| COX18 | 0.78 | 0.82 | Both Isoforms | SPTB | 0.40 | 1.01 | AR-V7 |
| WDR93 | 0.67 | 0.82 | Both Isoforms | PANX2 | 0.39 | 1.01 | AR-V7 |
| BMPR1A | 1.74 | 0.82 | Both Isoforms | ACYP1 | 0.23 | 1.01 | AR-V7 |
| DUSP4 | 1.34 | 0.82 | Both Isoforms | DHFR | 0.43 | 1.02 | AR-V7 |
| MOB4 | 0.66 | 0.82 | Both Isoforms | HMGA2 | 0.40 | 1.02 | AR-V7 |
| THRB | 1.27 | 0.83 | Both Isoforms | H2AFX | 0.56 | 1.02 | AR-V7 |
| ECI2 | 1.54 | 0.83 | Both Isoforms | SORT1 | 0.55 | 1.02 | AR-V7 |
| FAM174B | 1.79 | 0.83 | Both Isoforms | BRICD5 | 0.48 | 1.02 | AR-V7 |
| WTIP | 0.67 | 0.84 | Both Isoforms | IRF7 | 0.50 | 1.03 | AR-V7 |

| | | | | | | | |
|---|---|---|---|---|---|---|---|
| NEK3 | 0.96 | 0.84 | Both Isoforms | PTK2B | 0.03 | 1.04 | AR-V7 |
| SC5D | 1.30 | 0.84 | Both Isoforms | CHKB | 0.35 | 1.05 | AR-V7 |
| RABAC1 | 0.81 | 0.84 | Both Isoforms | SAMD15 | 0.22 | 1.06 | AR-V7 |
| DHCR24 | 2.18 | 0.84 | Both Isoforms | NT5DC3 | 0.43 | 1.06 | AR-V7 |
| ILK | 0.59 | 0.85 | Both Isoforms | MLEC | 0.57 | 1.06 | AR-V7 |
| LAMC1 | 2.29 | 0.85 | Both Isoforms | EME1 | 0.27 | 1.06 | AR-V7 |
| ACOT9 | 0.65 | 0.85 | Both Isoforms | DAGLA | 0.10 | 1.06 | AR-V7 |
| EDEM2 | 0.95 | 0.85 | Both Isoforms | MAT2A | 0.26 | 1.07 | AR-V7 |
| KCNAB1 | 1.42 | 0.85 | Both Isoforms | TRIM36 | 0.13 | 1.07 | AR-V7 |
| SSR3 | 1.35 | 0.85 | Both Isoforms | RAD51AP1 | 0.30 | 1.07 | AR-V7 |
| LRRC63 | 1.36 | 0.85 | Both Isoforms | SCD5 | 0.55 | 1.08 | AR-V7 |
| CRELD2 | 0.88 | 0.85 | Both Isoforms | SPC24 | 0.32 | 1.08 | AR-V7 |
| PLEKHG2 | 1.55 | 0.86 | Both Isoforms | HSD17B4 | 0.34 | 1.10 | AR-V7 |
| CD4 | 0.88 | 0.86 | Both Isoforms | AKAP7 | 0.42 | 1.10 | AR-V7 |
| ABCA5 | 1.04 | 0.86 | Both Isoforms | VANGL1 | 0.47 | 1.10 | AR-V7 |
| NOX5 | 0.98 | 0.86 | Both Isoforms | RECQL4 | 0.53 | 1.10 | AR-V7 |
| EPPK1 | 1.20 | 0.86 | Both Isoforms | GALNT2 | 0.35 | 1.11 | AR-V7 |
| WWC1 | 0.90 | 0.86 | Both Isoforms | TMPO | 0.18 | 1.11 | AR-V7 |
| ACADM | 0.84 | 0.86 | Both Isoforms | SMARCA1 | 0.26 | 1.12 | AR-V7 |
| NAAA | 0.67 | 0.86 | Both Isoforms | FSIP1 | 0.15 | 1.12 | AR-V7 |
| C14orf28 | 1.01 | 0.87 | Both Isoforms | B3GALNT1 | 0.50 | 1.15 | AR-V7 |
| PDIA5 | 1.57 | 0.87 | Both Isoforms | FTH1 | 0.43 | 1.15 | AR-V7 |
| SYNJ1 | 1.72 | 0.87 | Both Isoforms | CSF1 | 0.54 | 1.16 | AR-V7 |
| MINPP1 | 0.78 | 0.88 | Both Isoforms | TTC21A | 0.50 | 1.17 | AR-V7 |
| VGF | 1.50 | 0.88 | Both Isoforms | DUSP2 | 0.32 | 1.18 | AR-V7 |

| | | | | | | | |
|---|---|---|---|---|---|---|---|
| RAB4A | 1.69 | 0.88 | Both Isoforms | MARS2 | 0.02 | 1.18 | AR-V7 |
| FBN1 | 0.86 | 0.89 | Both Isoforms | PLOD2 | 0.52 | 1.19 | AR-V7 |
| ABCC1 | 1.91 | 0.89 | Both Isoforms | E2F7 | 0.12 | 1.19 | AR-V7 |
| TNFAIP3 | 2.66 | 0.89 | Both Isoforms | TASP1 | 0.28 | 1.19 | AR-V7 |
| DNAJB14 | 0.69 | 0.89 | Both Isoforms | EP400NL | 0.44 | 1.19 | AR-V7 |
| VMP1 | 1.17 | 0.90 | Both Isoforms | LIMS1 | 0.38 | 1.26 | AR-V7 |
| DLEC1 | 1.16 | 0.90 | Both Isoforms | THOC5 | 0.21 | 1.26 | AR-V7 |
| MKRN2OS | 1.09 | 0.90 | Both Isoforms | VLDLR | 0.29 | 1.26 | AR-V7 |
| JMJD6 | 0.68 | 0.90 | Both Isoforms | IFI6 | 0.36 | 1.27 | AR-V7 |
| KCTD9 | 1.21 | 0.90 | Both Isoforms | AMOT | 0.45 | 1.28 | AR-V7 |
| HIF1A | 0.58 | 0.90 | Both Isoforms | CBLL1 | 0.38 | 1.28 | AR-V7 |
| PPM1A | 0.66 | 0.91 | Both Isoforms | DMRT2 | 0.30 | 1.29 | AR-V7 |
| DNM1L | 1.44 | 0.91 | Both Isoforms | SLC44A5 | 0.37 | 1.30 | AR-V7 |
| ZNF841 | 1.11 | 0.91 | Both Isoforms | C1orf21 | 0.29 | 1.31 | AR-V7 |
| KLK15 | 1.51 | 0.91 | Both Isoforms | CYSTM1 | 0.03 | 1.31 | AR-V7 |
| ZCCHC6 | 1.92 | 0.91 | Both Isoforms | OSGIN2 | 0.19 | 1.32 | AR-V7 |
| ASAH1 | 0.90 | 0.91 | Both Isoforms | GPER1 | 0.37 | 1.32 | AR-V7 |
| PCYT2 | 1.01 | 0.91 | Both Isoforms | LRRFIP2 | 0.55 | 1.33 | AR-V7 |
| PSME4 | 0.65 | 0.91 | Both Isoforms | MAP1A | 0.54 | 1.33 | AR-V7 |
| HIPK2 | 1.06 | 0.91 | Both Isoforms | NRBP2 | 0.08 | 1.35 | AR-V7 |
| C16orf71 | 0.73 | 0.92 | Both Isoforms | HELLS | 0.55 | 1.35 | AR-V7 |
| HMGXB3 | 1.59 | 0.92 | Both Isoforms | TYMS | 0.47 | 1.36 | AR-V7 |
| TBC1D4 | 1.15 | 0.92 | Both Isoforms | JAKMIP1 | 0.05 | 1.37 | AR-V7 |
| KIFC3 | 0.72 | 0.92 | Both Isoforms | MT1E | 0.06 | 1.38 | AR-V7 |
| ZC3H12A | 0.60 | 0.92 | Both Isoforms | RASL10A | 0.16 | 1.40 | AR-V7 |

| | | | | | | | |
|---|---|---|---|---|---|---|---|
| PFKFB1 | 0.85 | 0.92 | Both Isoforms | USP51 | 0.21 | 1.43 | AR-V7 |
| ACE2 | 0.96 | 0.93 | Both Isoforms | IZUMO4 | 0.07 | 1.44 | AR-V7 |
| B3GNT9 | 0.81 | 0.93 | Both Isoforms | SSX2IP | 0.57 | 1.44 | AR-V7 |
| FNDC3B | 0.80 | 0.93 | Both Isoforms | BOC | 0.46 | 1.45 | AR-V7 |
| SLCO2A1 | 1.06 | 0.93 | Both Isoforms | PTP4A1 | 0.39 | 1.45 | AR-V7 |
| PPP1CB | 1.16 | 0.95 | Both Isoforms | VGF | 0.06 | 1.46 | AR-V7 |
| PALLD | 0.63 | 0.95 | Both Isoforms | ESCO2 | 0.45 | 1.46 | AR-V7 |
| ATG101 | 0.99 | 0.95 | Both Isoforms | CDH23 | 0.00 | 1.47 | AR-V7 |
| AMHR2 | 0.68 | 0.95 | Both Isoforms | SMIM1 | 0.46 | 1.47 | AR-V7 |
| ZYX | 0.84 | 0.95 | Both Isoforms | TRIM35 | 0.38 | 1.48 | AR-V7 |
| CKAP4 | 0.66 | 0.95 | Both Isoforms | TMEM56 | 0.10 | 1.48 | AR-V7 |
| TSPAN13 | 1.00 | 0.95 | Both Isoforms | AKAP17A | 0.06 | 1.50 | AR-V7 |
| MLPH | 1.83 | 0.95 | Both Isoforms | ATOH8 | 0.16 | 1.51 | AR-V7 |
| SMIM14 | 1.09 | 0.96 | Both Isoforms | CEBPB | 0.09 | 1.54 | AR-V7 |
| ZCCHC9 | 0.90 | 0.96 | Both Isoforms | SBSPON | 0.00 | 1.55 | AR-V7 |
| CAP2 | 1.37 | 0.96 | Both Isoforms | MT1X | 0.58 | 1.55 | AR-V7 |
| GOT2 | 0.66 | 0.96 | Both Isoforms | HBEGF | 0.10 | 1.56 | AR-V7 |
| FAM151B | 1.14 | 0.96 | Both Isoforms | CCNE1 | 0.58 | 1.66 | AR-V7 |
| ANKH | 2.26 | 0.97 | Both Isoforms | ZBTB8B | 0.58 | 1.66 | AR-V7 |
| MBD2 | 0.59 | 0.97 | Both Isoforms | PIM3 | 0.25 | 1.67 | AR-V7 |
| BMPR1B | 1.64 | 0.97 | Both Isoforms | ZNF589 | 0.53 | 1.68 | AR-V7 |
| SLC39A10 | 1.59 | 0.97 | Both Isoforms | VEPH1 | 0.24 | 1.68 | AR-V7 |
| DNAJB9 | 2.01 | 0.98 | Both Isoforms | TIGD3 | 0.29 | 1.69 | AR-V7 |
| SLC25A33 | 0.82 | 0.98 | Both Isoforms | FOXD4 | 0.23 | 1.76 | AR-V7 |
| HK1 | 1.11 | 0.98 | Both Isoforms | LRCH2 | 0.47 | 1.85 | AR-V7 |

| | | | | | | | |
|---|---|---|---|---|---|---|---|
| AGR2 | 0.63 | 0.99 | Both Isoforms | GRAMD2 | 0.16 | 1.86 | AR-V7 |
| CYB5A | 0.70 | 0.99 | Both Isoforms | AC138969.4 | 0.51 | 1.87 | AR-V7 |
| ZNF652 | 0.93 | 0.99 | Both Isoforms | CITED4 | 0.56 | 1.92 | AR-V7 |
| ZNF737 | 0.60 | 0.99 | Both Isoforms | CDK6 | 0.12 | 1.93 | AR-V7 |
| SEC24D | 1.87 | 0.99 | Both Isoforms | SOWAHA | 0.21 | 1.95 | AR-V7 |
| DAPK3 | 0.95 | 0.99 | Both Isoforms | GALNT9 | 0.43 | 1.98 | AR-V7 |
| FICD | 1.84 | 1.00 | Both Isoforms | ZNF18 | 0.44 | 2.06 | AR-V7 |
| TMEM263 | 0.80 | 1.00 | Both Isoforms | NPY | 0.00 | 2.07 | AR-V7 |
| EXTL2 | 0.74 | 1.00 | Both Isoforms | LRRC75A | 0.52 | 2.08 | AR-V7 |
| SPAG17 | 2.70 | 1.00 | Both Isoforms | AASS | 0.37 | 2.10 | AR-V7 |
| EPDR1 | 1.42 | 1.00 | Both Isoforms | UVRAG | 0.14 | 2.13 | AR-V7 |
| SLC25A30 | 0.72 | 1.00 | Both Isoforms | MT2A | 0.56 | 2.16 | AR-V7 |
| LDLR | 1.39 | 1.00 | Both Isoforms | TCAF2 | 0.49 | 2.28 | AR-V7 |
| ZNF233 | 1.07 | 1.01 | Both Isoforms | PPM1B | 0.43 | 2.46 | AR-V7 |
| NMD3 | 0.64 | 1.01 | Both Isoforms | RAB9B | 0.13 | 2.54 | AR-V7 |
| MAPK8IP2 | 0.75 | 1.01 | Both Isoforms | ZBED3 | 0.28 | 2.70 | AR-V7 |
| INPP4B | 3.05 | 1.01 | Both Isoforms | HPCAL4 | 0.38 | 2.83 | AR-V7 |
| ZNF552 | 1.11 | 1.01 | Both Isoforms | SDC1 | 0.27 | 2.96 | AR-V7 |
| ERO1B | 0.60 | 1.01 | Both Isoforms | SERPINA7 | 0.30 | 3.16 | AR-V7 |
| HSP90B1 | 0.86 | 1.01 | Both Isoforms | GTPBP6 | 0.44 | 4.00 | AR-V7 |
| FAM214B | 0.88 | 1.01 | Both Isoforms | PCDH7 | 0.55 | 4.35 | AR-V7 |
| CAPN13 | 0.76 | 1.02 | Both Isoforms | NPTX2 | 0.00 | 4.40 | AR-V7 |
| FAM103A1 | 1.00 | 1.02 | Both Isoforms | SMOC1 | 0.44 | 4.44 | AR-V7 |
| MERTK | 2.43 | 1.02 | Both Isoforms | NEXN | 0.37 | 4.61 | AR-V7 |
| NFKBIA | 1.47 | 1.02 | Both Isoforms | ADAMTS8 | 0.43 | 6.34 | AR-V7 |

| | | | | | | | |
|---|---|---|---|---|---|---|---|
| VCL | 1.30 | 1.02 | Both Isoforms | HTATIP2 | 0.70 | 0.00 | AR-FL |
| CCDC153 | 1.01 | 1.03 | Both Isoforms | EXOC6 | 0.67 | 0.00 | AR-FL |
| SOX9 | 2.88 | 1.03 | Both Isoforms | ACOT9 | 0.85 | 0.01 | AR-FL |
| TRPM4 | 1.11 | 1.03 | Both Isoforms | MYADML2 | 1.04 | 0.01 | AR-FL |
| SLA2 | 2.63 | 1.03 | Both Isoforms | ITGA10 | 1.10 | 0.01 | AR-FL |
| PLA1A | 2.13 | 1.04 | Both Isoforms | CPT1A | 0.59 | 0.01 | AR-FL |
| RGPD3 | 1.35 | 1.04 | Both Isoforms | FRAT1 | 1.02 | 0.01 | AR-FL |
| NFIB | 1.44 | 1.04 | Both Isoforms | FXYD3 | 0.61 | 0.01 | AR-FL |
| ORC5 | 1.21 | 1.04 | Both Isoforms | DLGAP4 | 0.62 | 0.01 | AR-FL |
| CRLS1 | 1.20 | 1.04 | Both Isoforms | CCDC126 | 0.94 | 0.02 | AR-FL |
| EDEM3 | 1.49 | 1.04 | Both Isoforms | TNFSF13 | 0.69 | 0.02 | AR-FL |
| TMTC2 | 1.79 | 1.04 | Both Isoforms | SPG21 | 0.69 | 0.02 | AR-FL |
| BBX | 1.01 | 1.04 | Both Isoforms | TLN2 | 0.88 | 0.02 | AR-FL |
| PPP1R13L | 0.82 | 1.04 | Both Isoforms | GLMP | 1.20 | 0.02 | AR-FL |
| CHRNB1 | 0.73 | 1.05 | Both Isoforms | RHPN2 | 1.49 | 0.02 | AR-FL |
| SMS | 2.52 | 1.05 | Both Isoforms | FLYWCH2 | 0.72 | 0.02 | AR-FL |
| PHACTR2 | 1.01 | 1.05 | Both Isoforms | BCL3 | 1.75 | 0.02 | AR-FL |
| C1orf122 | 0.91 | 1.05 | Both Isoforms | HDAC7 | 0.59 | 0.02 | AR-FL |
| MYADM | 1.08 | 1.06 | Both Isoforms | SHB | 0.95 | 0.03 | AR-FL |
| ELK4 | 1.19 | 1.06 | Both Isoforms | ACTL10 | 0.88 | 0.03 | AR-FL |
| LSS | 0.89 | 1.06 | Both Isoforms | NOL3 | 0.92 | 0.03 | AR-FL |
| LAMA3 | 1.13 | 1.07 | Both Isoforms | STK10 | 0.80 | 0.03 | AR-FL |
| PDZRN3 | 1.48 | 1.08 | Both Isoforms | CD82 | 1.08 | 0.03 | AR-FL |
| MYL12A | 1.02 | 1.08 | Both Isoforms | MYCL | 1.05 | 0.03 | AR-FL |
| RFTN2 | 0.65 | 1.08 | Both Isoforms | C8orf33 | 0.63 | 0.03 | AR-FL |

| | | | | | | | |
|---|---|---|---|---|---|---|---|
| AC021106.1 | 0.76 | 1.09 | Both Isoforms | ST3GAL6 | 0.93 | 0.03 | AR-FL |
| NAMPT | 1.57 | 1.10 | Both Isoforms | CYTH1 | 0.69 | 0.03 | AR-FL |
| GPR37 | 1.35 | 1.10 | Both Isoforms | ZNF614 | 0.72 | 0.03 | AR-FL |
| TBKBP1 | 0.59 | 1.10 | Both Isoforms | STS | 0.75 | 0.04 | AR-FL |
| ODC1 | 2.22 | 1.10 | Both Isoforms | CALM2 | 0.64 | 0.04 | AR-FL |
| EVA1B | 1.03 | 1.11 | Both Isoforms | ACBD3 | 0.60 | 0.04 | AR-FL |
| NOSTRIN | 1.48 | 1.11 | Both Isoforms | SGPP1 | 0.83 | 0.04 | AR-FL |
| LGMN | 0.76 | 1.11 | Both Isoforms | GLIPR2 | 0.96 | 0.04 | AR-FL |
| TLX2 | 1.00 | 1.11 | Both Isoforms | FAM53C | 0.63 | 0.04 | AR-FL |
| CFAP99 | 0.66 | 1.11 | Both Isoforms | ANAPC15 | 0.69 | 0.04 | AR-FL |
| CYR61 | 1.13 | 1.12 | Both Isoforms | GNAI2 | 0.64 | 0.04 | AR-FL |
| PHOSPHO1 | 0.88 | 1.12 | Both Isoforms | ZNF217 | 1.04 | 0.04 | AR-FL |
| CXCR4 | 4.12 | 1.12 | Both Isoforms | PLOD1 | 0.66 | 0.04 | AR-FL |
| OCEL1 | 0.59 | 1.12 | Both Isoforms | EML1 | 0.76 | 0.04 | AR-FL |
| SGK223 | 3.46 | 1.12 | Both Isoforms | ADAT3 | 0.82 | 0.05 | AR-FL |
| KDM7A | 0.69 | 1.13 | Both Isoforms | RABEP1 | 0.83 | 0.05 | AR-FL |
| LRRC8A | 0.78 | 1.13 | Both Isoforms | RAP1GAP | 1.22 | 0.05 | AR-FL |
| NXF2 | 2.47 | 1.13 | Both Isoforms | CDC42EP3 | 1.93 | 0.05 | AR-FL |
| H1F0 | 0.88 | 1.13 | Both Isoforms | STRN | 0.82 | 0.05 | AR-FL |
| ERO1A | 1.03 | 1.14 | Both Isoforms | ALOX5 | 1.21 | 0.05 | AR-FL |
| HSD17B11 | 2.05 | 1.14 | Both Isoforms | RGS9BP | 1.06 | 0.05 | AR-FL |
| TPM4 | 0.82 | 1.14 | Both Isoforms | CCNJ | 0.65 | 0.06 | AR-FL |
| STARD13 | 0.92 | 1.14 | Both Isoforms | MYL12B | 0.65 | 0.06 | AR-FL |
| RANBP2 | 0.71 | 1.14 | Both Isoforms | FAM127C | 0.72 | 0.06 | AR-FL |
| ACTB | 0.63 | 1.15 | Both Isoforms | TTYH3 | 0.94 | 0.06 | AR-FL |

| | | | | | | | |
|---|---|---|---|---|---|---|---|
| IDI1 | 1.13 | 1.16 | Both Isoforms | PRDM10 | 0.71 | 0.06 | AR-FL |
| RHBDF1 | 0.86 | 1.16 | Both Isoforms | ZDHHC21 | 0.71 | 0.06 | AR-FL |
| VLDLR | 1.50 | 1.16 | Both Isoforms | GALNT7 | 0.97 | 0.06 | AR-FL |
| PRKCD | 0.92 | 1.16 | Both Isoforms | INTS8 | 0.59 | 0.06 | AR-FL |
| CFAP221 | 2.28 | 1.16 | Both Isoforms | PCTP | 0.66 | 0.06 | AR-FL |
| TSPYL2 | 0.62 | 1.16 | Both Isoforms | EPHB3 | 1.28 | 0.06 | AR-FL |
| CDYL2 | 2.48 | 1.17 | Both Isoforms | BAG3 | 0.68 | 0.07 | AR-FL |
| HIST4H4 | 0.98 | 1.17 | Both Isoforms | NQO2 | 0.71 | 0.07 | AR-FL |
| RAB3B | 2.04 | 1.18 | Both Isoforms | NPTXR | 1.15 | 0.07 | AR-FL |
| NEDD4L | 1.23 | 1.19 | Both Isoforms | P4HA1 | 0.84 | 0.07 | AR-FL |
| MALT1 | 1.76 | 1.19 | Both Isoforms | DYNC1LI1 | 0.62 | 0.07 | AR-FL |
| WDR41 | 0.65 | 1.19 | Both Isoforms | ITGA3 | 1.41 | 0.07 | AR-FL |
| EEF2K | 1.08 | 1.20 | Both Isoforms | ZNF443 | 0.96 | 0.07 | AR-FL |
| MAPK11 | 0.93 | 1.20 | Both Isoforms | TMEM63A | 0.91 | 0.07 | AR-FL |
| SMN1 | 0.77 | 1.21 | Both Isoforms | CYB5R4 | 0.67 | 0.07 | AR-FL |
| ITGAV | 1.70 | 1.21 | Both Isoforms | ZBTB7C | 0.78 | 0.07 | AR-FL |
| PLPP6 | 1.60 | 1.21 | Both Isoforms | LRTOMT | 0.58 | 0.07 | AR-FL |
| KRT18 | 1.89 | 1.21 | Both Isoforms | SNAP91 | 1.40 | 0.08 | AR-FL |
| EIF2AK3 | 1.13 | 1.21 | Both Isoforms | GPR180 | 0.63 | 0.08 | AR-FL |
| TNFRSF10B | 1.49 | 1.21 | Both Isoforms | CLEC11A | 0.62 | 0.08 | AR-FL |
| RRAS | 1.04 | 1.22 | Both Isoforms | WWTR1 | 1.15 | 0.08 | AR-FL |
| PTGES3L | 0.95 | 1.22 | Both Isoforms | ALDH1A3 | 2.82 | 0.08 | AR-FL |
| FMO4 | 1.09 | 1.22 | Both Isoforms | MMEL1 | 0.68 | 0.08 | AR-FL |
| SORD | 2.21 | 1.22 | Both Isoforms | SLC37A1 | 0.60 | 0.08 | AR-FL |
| TCTN2 | 0.65 | 1.23 | Both Isoforms | COL4A3BP | 0.64 | 0.08 | AR-FL |

| | | | | | | | |
|---|---|---|---|---|---|---|---|
| HERC3 | 2.75 | 1.23 | Both Isoforms | LHFPL2 | 0.81 | 0.08 | AR-FL |
| ARRDC1 | 1.29 | 1.23 | Both Isoforms | KDM6B | 0.95 | 0.08 | AR-FL |
| LRRC46 | 0.74 | 1.23 | Both Isoforms | FAM174B | 0.68 | 0.08 | AR-FL |
| ERLEC1 | 1.18 | 1.23 | Both Isoforms | RNF24 | 0.83 | 0.08 | AR-FL |
| PPFIBP2 | 2.02 | 1.24 | Both Isoforms | CNDP2 | 0.75 | 0.08 | AR-FL |
| GPR137C | 0.73 | 1.24 | Both Isoforms | ME1 | 0.99 | 0.08 | AR-FL |
| TMEM139 | 0.77 | 1.24 | Both Isoforms | CDS1 | 0.79 | 0.09 | AR-FL |
| PREX2 | 1.08 | 1.24 | Both Isoforms | TBK1 | 0.60 | 0.09 | AR-FL |
| CDC14B | 1.56 | 1.24 | Both Isoforms | ZG16B | 0.67 | 0.09 | AR-FL |
| SLC16A6 | 3.54 | 1.24 | Both Isoforms | ATP8B1 | 0.84 | 0.09 | AR-FL |
| PGM2 | 1.30 | 1.25 | Both Isoforms | TMEM206 | 0.89 | 0.09 | AR-FL |
| SAR1B | 1.02 | 1.25 | Both Isoforms | PM20D2 | 0.76 | 0.09 | AR-FL |
| PACS1 | 2.13 | 1.26 | Both Isoforms | BVES | 1.09 | 0.09 | AR-FL |
| TP53INP2 | 0.62 | 1.26 | Both Isoforms | S100P | 2.51 | 0.09 | AR-FL |
| C6orf201 | 1.65 | 1.26 | Both Isoforms | ACAA1 | 0.79 | 0.09 | AR-FL |
| NT5DC3 | 1.34 | 1.26 | Both Isoforms | TRIM68 | 0.84 | 0.09 | AR-FL |
| KDELR3 | 1.83 | 1.27 | Both Isoforms | SCUBE1 | 1.51 | 0.09 | AR-FL |
| RPN1 | 0.77 | 1.27 | Both Isoforms | RPGR | 0.58 | 0.10 | AR-FL |
| SHROOM3 | 1.52 | 1.28 | Both Isoforms | NFKBIE | 1.09 | 0.10 | AR-FL |
| TNFRSF19 | 2.24 | 1.28 | Both Isoforms | MTHFR | 0.61 | 0.10 | AR-FL |
| DNAL1 | 0.60 | 1.29 | Both Isoforms | SPTBN2 | 0.69 | 0.10 | AR-FL |
| B4GALT1 | 0.76 | 1.29 | Both Isoforms | ARMC5 | 0.67 | 0.10 | AR-FL |
| DBI | 1.71 | 1.30 | Both Isoforms | ABCC4 | 1.60 | 0.10 | AR-FL |
| LDB3 | 0.79 | 1.30 | Both Isoforms | HSD11B2 | 0.64 | 0.10 | AR-FL |
| SLC26A2 | 1.36 | 1.30 | Both Isoforms | LARP4B | 0.58 | 0.10 | AR-FL |

| | | | | | | | |
|---|---|---|---|---|---|---|---|
| EFCAB12 | 2.48 | 1.30 | Both Isoforms | TMED3 | 0.81 | 0.10 | AR-FL |
| CEBPD | 0.91 | 1.30 | Both Isoforms | NEK5 | 0.99 | 0.10 | AR-FL |
| KIF5C | 0.93 | 1.30 | Both Isoforms | TVP23B | 1.37 | 0.10 | AR-FL |
| MSMO1 | 0.90 | 1.31 | Both Isoforms | NCOR2 | 0.61 | 0.10 | AR-FL |
| CNKSR2 | 1.13 | 1.31 | Both Isoforms | FSTL1 | 1.10 | 0.10 | AR-FL |
| C1orf116 | 2.35 | 1.31 | Both Isoforms | GFPT1 | 0.66 | 0.10 | AR-FL |
| BHLHE40 | 1.14 | 1.31 | Both Isoforms | TINF2 | 0.61 | 0.10 | AR-FL |
| CFAP57 | 0.76 | 1.31 | Both Isoforms | SNTB2 | 0.76 | 0.10 | AR-FL |
| GRIN3A | 1.82 | 1.31 | Both Isoforms | ZNF765 | 0.73 | 0.11 | AR-FL |
| IGF1 | 2.60 | 1.31 | Both Isoforms | COPB1 | 0.67 | 0.11 | AR-FL |
| AASS | 1.16 | 1.32 | Both Isoforms | CBLC | 0.59 | 0.11 | AR-FL |
| HIST1H4B | 0.71 | 1.33 | Both Isoforms | MLKL | 1.36 | 0.11 | AR-FL |
| FBXO38 | 1.43 | 1.33 | Both Isoforms | KAT2B | 1.24 | 0.11 | AR-FL |
| PIAS1 | 1.38 | 1.33 | Both Isoforms | UBE2G1 | 0.71 | 0.11 | AR-FL |
| STK17B | 2.44 | 1.33 | Both Isoforms | FAM21A | 0.86 | 0.11 | AR-FL |
| ZNF18 | 1.39 | 1.34 | Both Isoforms | CAMKK2 | 0.64 | 0.11 | AR-FL |
| RARB | 1.13 | 1.34 | Both Isoforms | ZNF850 | 0.65 | 0.12 | AR-FL |
| GATSL3 | 0.63 | 1.34 | Both Isoforms | ZNF799 | 0.81 | 0.12 | AR-FL |
| KLF15 | 1.57 | 1.34 | Both Isoforms | NDFIP2 | 1.26 | 0.12 | AR-FL |
| AGAP2 | 2.50 | 1.35 | Both Isoforms | PCDHGB4 | 0.80 | 0.12 | AR-FL |
| TRANK1 | 0.90 | 1.35 | Both Isoforms | CCDC40 | 0.68 | 0.12 | AR-FL |
| TNFAIP8 | 1.87 | 1.35 | Both Isoforms | CXADR | 0.88 | 0.12 | AR-FL |
| PYGB | 0.63 | 1.35 | Both Isoforms | LRP3 | 1.13 | 0.12 | AR-FL |
| DHRS2 | 0.82 | 1.36 | Both Isoforms | PAQR5 | 0.89 | 0.12 | AR-FL |
| IFT57 | 0.76 | 1.37 | Both Isoforms | PARM1 | 2.22 | 0.12 | AR-FL |

| | | | | | | | |
|---|---|---|---|---|---|---|---|
| TMEM100 | 8.24 | 1.37 | Both Isoforms | EIF2B4 | 0.79 | 0.12 | AR-FL |
| CYP3A43 | 1.45 | 1.37 | Both Isoforms | PAK1IP1 | 0.60 | 0.12 | AR-FL |
| FAM114A1 | 0.60 | 1.38 | Both Isoforms | FAM120AOS | 0.65 | 0.12 | AR-FL |
| ZBTB10 | 2.27 | 1.38 | Both Isoforms | TMEM79 | 1.94 | 0.13 | AR-FL |
| POPDC2 | 0.64 | 1.38 | Both Isoforms | PRR19 | 0.73 | 0.13 | AR-FL |
| CHKA | 0.99 | 1.38 | Both Isoforms | RDH11 | 0.65 | 0.13 | AR-FL |
| CNTN3 | 1.31 | 1.39 | Both Isoforms | PRLR | 0.74 | 0.13 | AR-FL |
| ID4 | 1.37 | 1.39 | Both Isoforms | DHDH | 1.23 | 0.13 | AR-FL |
| PNPLA8 | 1.17 | 1.40 | Both Isoforms | RIMBP3 | 1.59 | 0.13 | AR-FL |
| CDH15 | 1.05 | 1.40 | Both Isoforms | LRRC8B | 0.61 | 0.13 | AR-FL |
| DISP1 | 1.71 | 1.42 | Both Isoforms | SLC9A2 | 0.71 | 0.13 | AR-FL |
| ERVK3-1 | 0.98 | 1.43 | Both Isoforms | MAPK6 | 1.11 | 0.13 | AR-FL |
| TBRG1 | 1.91 | 1.43 | Both Isoforms | ZDHHC9 | 0.85 | 0.13 | AR-FL |
| KLK4 | 1.53 | 1.45 | Both Isoforms | MESP1 | 1.39 | 0.13 | AR-FL |
| SCAP | 1.33 | 1.45 | Both Isoforms | PPP2R2D | 0.63 | 0.13 | AR-FL |
| ANGPTL4 | 0.58 | 1.48 | Both Isoforms | GOLGA5 | 0.80 | 0.13 | AR-FL |
| CCDC6 | 0.66 | 1.48 | Both Isoforms | PSEN2 | 0.77 | 0.13 | AR-FL |
| SLC38A2 | 0.99 | 1.48 | Both Isoforms | SLC16A3 | 1.21 | 0.13 | AR-FL |
| LIN7B | 1.69 | 1.49 | Both Isoforms | SLAIN2 | 0.88 | 0.13 | AR-FL |
| PLPP1 | 2.27 | 1.49 | Both Isoforms | PDCD10 | 0.67 | 0.13 | AR-FL |
| COL23A1 | 0.84 | 1.50 | Both Isoforms | MYBPC1 | 0.80 | 0.13 | AR-FL |
| FERMT2 | 1.21 | 1.51 | Both Isoforms | CAP1 | 0.70 | 0.14 | AR-FL |
| LDLRAD3 | 1.58 | 1.52 | Both Isoforms | C20orf196 | 1.09 | 0.14 | AR-FL |
| SLC2A12 | 1.57 | 1.53 | Both Isoforms | RANBP3L | 1.53 | 0.14 | AR-FL |
| ARAP2 | 1.44 | 1.55 | Both Isoforms | USP6NL | 0.74 | 0.14 | AR-FL |

| | | | | | | | |
|---|---|---|---|---|---|---|---|
| PDLIM5 | 1.75 | 1.55 | Both Isoforms | ZBTB42 | 0.69 | 0.14 | AR-FL |
| GBP1 | 3.67 | 1.55 | Both Isoforms | LRIG1 | 1.62 | 0.14 | AR-FL |
| ATP1A1 | 1.24 | 1.55 | Both Isoforms | CTC-534A2.2 | 0.63 | 0.14 | AR-FL |
| UNC80 | 2.25 | 1.55 | Both Isoforms | STRN3 | 0.60 | 0.14 | AR-FL |
| CBLL1 | 1.60 | 1.56 | Both Isoforms | TAX1BP3 | 0.89 | 0.14 | AR-FL |
| FDFT1 | 1.14 | 1.56 | Both Isoforms | ZNF143 | 0.60 | 0.14 | AR-FL |
| RHOBTB3 | 0.86 | 1.58 | Both Isoforms | TM9SF3 | 0.71 | 0.14 | AR-FL |
| RALGPS2 | 1.06 | 1.59 | Both Isoforms | CSK | 0.79 | 0.14 | AR-FL |
| KCNH6 | 2.41 | 1.60 | Both Isoforms | RABEP2 | 1.47 | 0.14 | AR-FL |
| CASC10 | 2.07 | 1.60 | Both Isoforms | PGM3 | 0.82 | 0.14 | AR-FL |
| ATP8B2 | 1.28 | 1.61 | Both Isoforms | RAB43 | 1.39 | 0.14 | AR-FL |
| PDE8B | 0.62 | 1.61 | Both Isoforms | ZMPSTE24 | 0.82 | 0.15 | AR-FL |
| HIST2H2BE | 0.72 | 1.62 | Both Isoforms | PPM1E | 1.01 | 0.15 | AR-FL |
| TLR6 | 0.63 | 1.62 | Both Isoforms | FZD9 | 2.01 | 0.15 | AR-FL |
| CADPS2 | 2.87 | 1.63 | Both Isoforms | CCDC149 | 0.64 | 0.15 | AR-FL |
| C17orf58 | 0.59 | 1.64 | Both Isoforms | TNFAIP3 | 1.20 | 0.15 | AR-FL |
| SH3D21 | 2.59 | 1.64 | Both Isoforms | DHX15 | 0.66 | 0.15 | AR-FL |
| GLYATL2 | 2.40 | 1.64 | Both Isoforms | EGF | 1.69 | 0.15 | AR-FL |
| ANG | 1.97 | 1.64 | Both Isoforms | CDCP1 | 0.63 | 0.15 | AR-FL |
| UNC13D | 0.79 | 1.66 | Both Isoforms | ADARB2 | 0.96 | 0.15 | AR-FL |
| WIPI1 | 2.80 | 1.67 | Both Isoforms | PRKCH | 1.58 | 0.15 | AR-FL |
| ATF3 | 0.85 | 1.67 | Both Isoforms | KMT2C | 0.70 | 0.16 | AR-FL |
| C1orf21 | 2.18 | 1.67 | Both Isoforms | GGT1 | 0.91 | 0.16 | AR-FL |
| PCSK5 | 2.62 | 1.68 | Both Isoforms | MBLAC1 | 0.70 | 0.16 | AR-FL |
| ATP8B3 | 1.52 | 1.68 | Both Isoforms | RAB2A | 0.61 | 0.16 | AR-FL |

| | | | | | | | |
|---|---|---|---|---|---|---|---|
| TSC22D1 | 1.99 | 1.68 | Both Isoforms | GLP2R | 0.99 | 0.16 | AR-FL |
| JAG1 | 1.39 | 1.68 | Both Isoforms | BORCS6 | 0.89 | 0.16 | AR-FL |
| OSBPL8 | 1.27 | 1.70 | Both Isoforms | FZD5 | 2.03 | 0.16 | AR-FL |
| KANSL1L | 0.82 | 1.71 | Both Isoforms | PRICKLE3 | 0.76 | 0.16 | AR-FL |
| MYO1E | 1.53 | 1.72 | Both Isoforms | MAP9 | 0.58 | 0.16 | AR-FL |
| SWT1 | 2.34 | 1.72 | Both Isoforms | TMEM165 | 0.60 | 0.16 | AR-FL |
| PLA2G5 | 2.21 | 1.73 | Both Isoforms | VPS37B | 0.61 | 0.16 | AR-FL |
| CENPN | 3.63 | 1.74 | Both Isoforms | VANGL2 | 1.60 | 0.16 | AR-FL |
| ZNF578 | 0.60 | 1.74 | Both Isoforms | PROSER1 | 0.61 | 0.16 | AR-FL |
| LPAR3 | 1.93 | 1.74 | Both Isoforms | CDK2AP1 | 0.82 | 0.16 | AR-FL |
| LRRFIP2 | 1.58 | 1.74 | Both Isoforms | GALNT18 | 0.71 | 0.16 | AR-FL |
| LAT2 | 3.01 | 1.74 | Both Isoforms | MTMR9 | 1.32 | 0.16 | AR-FL |
| HPX | 0.97 | 1.76 | Both Isoforms | REEP3 | 0.79 | 0.17 | AR-FL |
| LPXN | 1.35 | 1.76 | Both Isoforms | OTULIN | 0.69 | 0.17 | AR-FL |
| CITED2 | 0.79 | 1.77 | Both Isoforms | USP33 | 0.64 | 0.17 | AR-FL |
| EPSTI1 | 1.98 | 1.78 | Both Isoforms | MDFI | 1.28 | 0.17 | AR-FL |
| DLX2 | 1.46 | 1.78 | Both Isoforms | SLC51A | 0.65 | 0.17 | AR-FL |
| HMGCS1 | 1.70 | 1.80 | Both Isoforms | ESPN | 0.69 | 0.17 | AR-FL |
| NXF2B | 3.42 | 1.80 | Both Isoforms | WDR37 | 0.94 | 0.17 | AR-FL |
| GJC3 | 0.85 | 1.81 | Both Isoforms | TPD52 | 1.01 | 0.17 | AR-FL |
| ITGA1 | 1.33 | 1.83 | Both Isoforms | ACACA | 0.66 | 0.17 | AR-FL |
| GLUL | 1.26 | 1.84 | Both Isoforms | DHRS3 | 0.61 | 0.17 | AR-FL |
| ROPN1L | 0.71 | 1.84 | Both Isoforms | AK6 | 0.58 | 0.17 | AR-FL |
| ARHGEF37 | 0.68 | 1.85 | Both Isoforms | ECH1 | 0.64 | 0.17 | AR-FL |
| PSCA | 0.71 | 1.86 | Both Isoforms | SRMS | 0.73 | 0.18 | AR-FL |

| | | | | | | | |
|---|---|---|---|---|---|---|---|
| GSN | 1.02 | 1.86 | Both Isoforms | SEPHS2 | 0.67 | 0.18 | AR-FL |
| HACD2 | 0.71 | 1.87 | Both Isoforms | GALNT12 | 1.00 | 0.18 | AR-FL |
| KCND2 | 1.86 | 1.87 | Both Isoforms | ARHGEF16 | 0.61 | 0.18 | AR-FL |
| CADM2 | 3.27 | 1.87 | Both Isoforms | MANSC1 | 0.93 | 0.18 | AR-FL |
| TBC1D1 | 1.89 | 1.87 | Both Isoforms | ZNF579 | 0.65 | 0.18 | AR-FL |
| NR4A1 | 2.17 | 1.88 | Both Isoforms | LENG1 | 0.59 | 0.18 | AR-FL |
| TMEM164 | 1.34 | 1.88 | Both Isoforms | TRIB3 | 0.60 | 0.19 | AR-FL |
| SPATA6L | 1.47 | 1.88 | Both Isoforms | GPR108 | 0.69 | 0.19 | AR-FL |
| SDCBP2 | 1.41 | 1.88 | Both Isoforms | ATP2A3 | 1.40 | 0.19 | AR-FL |
| ANKRD29 | 1.74 | 1.89 | Both Isoforms | GCLM | 0.72 | 0.19 | AR-FL |
| ABCC4 | 3.67 | 1.90 | Both Isoforms | PSMD7 | 0.65 | 0.19 | AR-FL |
| SLC41A1 | 2.44 | 1.90 | Both Isoforms | CLDN7 | 0.95 | 0.19 | AR-FL |
| MAOA | 1.11 | 1.91 | Both Isoforms | YIPF3 | 0.79 | 0.19 | AR-FL |
| FAM189A2 | 1.88 | 1.92 | Both Isoforms | IGF2R | 0.62 | 0.19 | AR-FL |
| NAT1 | 1.94 | 1.92 | Both Isoforms | JPH1 | 1.10 | 0.19 | AR-FL |
| DUSP10 | 0.77 | 1.93 | Both Isoforms | PIN4 | 0.69 | 0.19 | AR-FL |
| CLGN | 1.41 | 1.96 | Both Isoforms | HOXC6 | 0.96 | 0.20 | AR-FL |
| FASN | 1.27 | 1.96 | Both Isoforms | PHACTR4 | 0.61 | 0.20 | AR-FL |
| FGD4 | 2.47 | 1.98 | Both Isoforms | NLK | 0.64 | 0.20 | AR-FL |
| PPIC | 0.89 | 1.99 | Both Isoforms | PDLIM4 | 2.85 | 0.20 | AR-FL |
| FADS2 | 1.03 | 1.99 | Both Isoforms | C9orf91 | 0.90 | 0.20 | AR-FL |
| NKX3-1 | 3.25 | 1.99 | Both Isoforms | RALGAPB | 0.58 | 0.20 | AR-FL |
| KLF6 | 1.62 | 1.99 | Both Isoforms | SSPO | 1.21 | 0.20 | AR-FL |
| PPP2R3B | 2.12 | 1.99 | Both Isoforms | CCDC47 | 0.61 | 0.20 | AR-FL |
| TRIM35 | 0.70 | 1.99 | Both Isoforms | RELL2 | 2.11 | 0.20 | AR-FL |

| | | | | | | | |
|---|---|---|---|---|---|---|---|
| RNF148 | 3.50 | 1.99 | Both Isoforms | PSMA6 | 0.65 | 0.20 | AR-FL |
| CYP2U1 | 2.83 | 2.00 | Both Isoforms | RTN4R | 0.60 | 0.20 | AR-FL |
| TPM1 | 1.56 | 2.00 | Both Isoforms | PPDPF | 0.58 | 0.21 | AR-FL |
| PVRL3 | 1.47 | 2.00 | Both Isoforms | CRISP2 | 1.34 | 0.21 | AR-FL |
| GLUD2 | 1.05 | 2.00 | Both Isoforms | CRNKL1 | 0.87 | 0.21 | AR-FL |
| ETS2 | 0.65 | 2.01 | Both Isoforms | MAGEA12 | 0.69 | 0.21 | AR-FL |
| ABHD2 | 2.51 | 2.03 | Both Isoforms | TDG | 0.60 | 0.21 | AR-FL |
| DNAH7 | 1.50 | 2.04 | Both Isoforms | CCDC22 | 0.61 | 0.21 | AR-FL |
| PGLYRP2 | 2.89 | 2.04 | Both Isoforms | TUBB6 | 0.84 | 0.21 | AR-FL |
| PPFIBP1 | 1.99 | 2.04 | Both Isoforms | ARFGAP3 | 0.69 | 0.21 | AR-FL |
| CACNG4 | 4.95 | 2.05 | Both Isoforms | BCCIP | 0.71 | 0.21 | AR-FL |
| NSDHL | 0.63 | 2.05 | Both Isoforms | SLC35F2 | 0.79 | 0.21 | AR-FL |
| LPIN1 | 1.04 | 2.06 | Both Isoforms | KIF21A | 1.21 | 0.21 | AR-FL |
| CNN2 | 1.50 | 2.06 | Both Isoforms | MPHOSPH9 | 0.90 | 0.21 | AR-FL |
| FAM110B | 1.73 | 2.06 | Both Isoforms | NAGA | 0.58 | 0.22 | AR-FL |
| IRS1 | 0.88 | 2.07 | Both Isoforms | IQCD | 0.86 | 0.22 | AR-FL |
| INSIG1 | 1.45 | 2.08 | Both Isoforms | PACS1 | 0.92 | 0.22 | AR-FL |
| CHRNE | 0.86 | 2.08 | Both Isoforms | TMEM41B | 0.81 | 0.22 | AR-FL |
| C3orf58 | 2.80 | 2.08 | Both Isoforms | PLPP5 | 0.69 | 0.22 | AR-FL |
| TSKU | 1.59 | 2.09 | Both Isoforms | LRFN3 | 0.73 | 0.22 | AR-FL |
| NPR2 | 2.28 | 2.09 | Both Isoforms | AK4 | 0.72 | 0.22 | AR-FL |
| MICAL1 | 4.37 | 2.10 | Both Isoforms | HMOX2 | 0.88 | 0.22 | AR-FL |
| TCAF2 | 2.95 | 2.11 | Both Isoforms | SNAP23 | 0.59 | 0.22 | AR-FL |
| AHNAK | 1.41 | 2.11 | Both Isoforms | MRPS24 | 0.75 | 0.22 | AR-FL |
| CROT | 1.08 | 2.11 | Both Isoforms | UBE2J2 | 0.63 | 0.22 | AR-FL |

| | | | |
|---|---|---|---|
| FOXD4L1 | 0.84 | 2.12 | Both Isoforms |
| CDKN1C | 1.23 | 2.13 | Both Isoforms |
| PLOD2 | 1.13 | 2.13 | Both Isoforms |
| TMEFF2 | 0.63 | 2.14 | Both Isoforms |
| PARVA | 1.27 | 2.14 | Both Isoforms |
| SP6 | 0.65 | 2.15 | Both Isoforms |
| C1orf168 | 1.13 | 2.16 | Both Isoforms |
| HOMER2 | 2.21 | 2.17 | Both Isoforms |
| ZBTB16 | 2.02 | 2.17 | Both Isoforms |
| TXNDC16 | 1.13 | 2.17 | Both Isoforms |
| STYK1 | 1.32 | 2.18 | Both Isoforms |
| GLRX | 0.61 | 2.19 | Both Isoforms |
| MAF | 5.07 | 2.20 | Both Isoforms |
| PRICKLE2 | 1.34 | 2.22 | Both Isoforms |
| DNAH3 | 0.78 | 2.22 | Both Isoforms |
| ROR1 | 1.24 | 2.23 | Both Isoforms |
| LONRF1 | 2.33 | 2.23 | Both Isoforms |
| FHL2 | 1.47 | 2.24 | Both Isoforms |
| HSPA2 | 1.63 | 2.24 | Both Isoforms |
| HERC5 | 1.94 | 2.25 | Both Isoforms |
| ASRGL1 | 2.20 | 2.26 | Both Isoforms |
| KRT8 | 2.35 | 2.27 | Both Isoforms |
| FADS1 | 0.93 | 2.29 | Both Isoforms |
| PTCRA | 4.70 | 2.29 | Both Isoforms |
| CDKL1 | 1.48 | 2.30 | Both Isoforms |

| | | | |
|---|---|---|---|
| BMPR1A | 0.62 | 0.22 | AR-FL |
| ATP1B3 | 0.80 | 0.22 | AR-FL |
| EPHX1 | 0.77 | 0.23 | AR-FL |
| ACOT1 | 1.16 | 0.23 | AR-FL |
| SLC26A11 | 0.60 | 0.23 | AR-FL |
| ARMC7 | 0.84 | 0.23 | AR-FL |
| PDZD11 | 0.60 | 0.23 | AR-FL |
| GSE1 | 0.66 | 0.23 | AR-FL |
| RNF185 | 0.74 | 0.23 | AR-FL |
| MYO5C | 1.02 | 0.23 | AR-FL |
| TNFRSF10A | 0.92 | 0.23 | AR-FL |
| ZBTB7A | 0.61 | 0.23 | AR-FL |
| DCAF6 | 0.94 | 0.23 | AR-FL |
| FAM114A1 | 0.62 | 0.23 | AR-FL |
| NQO1 | 1.13 | 0.23 | AR-FL |
| GALK2 | 0.64 | 0.23 | AR-FL |
| ARAP3 | 1.11 | 0.23 | AR-FL |
| SLC17A5 | 1.67 | 0.23 | AR-FL |
| BLOC1S3 | 0.69 | 0.23 | AR-FL |
| FBXL8 | 0.63 | 0.24 | AR-FL |
| NSUN5 | 0.58 | 0.24 | AR-FL |
| SLC2A9 | 1.22 | 0.24 | AR-FL |
| SMPD2 | 1.19 | 0.24 | AR-FL |
| SELT | 0.78 | 0.24 | AR-FL |
| SPRYD3 | 0.60 | 0.24 | AR-FL |

| | | | | | | | |
|---|---|---|---|---|---|---|---|
| SLC31A2 | 0.62 | 2.30 | Both Isoforms | XRCC4 | 0.77 | 0.24 | AR-FL |
| CD163L1 | 0.93 | 2.32 | Both Isoforms | HN1L | 0.89 | 0.24 | AR-FL |
| ACSM1 | 2.56 | 2.33 | Both Isoforms | NEBL | 0.80 | 0.24 | AR-FL |
| SH3TC2 | 1.02 | 2.34 | Both Isoforms | NDEL1 | 0.74 | 0.24 | AR-FL |
| ANXA2 | 2.97 | 2.34 | Both Isoforms | NACAD | 1.61 | 0.24 | AR-FL |
| TRIM36 | 1.57 | 2.38 | Both Isoforms | ALG3 | 0.60 | 0.24 | AR-FL |
| MMRN2 | 3.37 | 2.39 | Both Isoforms | SLC1A5 | 1.19 | 0.24 | AR-FL |
| MYH15 | 1.51 | 2.44 | Both Isoforms | WBP2 | 0.59 | 0.24 | AR-FL |
| RHOU | 3.32 | 2.47 | Both Isoforms | GRHL3 | 3.07 | 0.24 | AR-FL |
| NEK10 | 1.29 | 2.48 | Both Isoforms | TARDBP | 0.58 | 0.24 | AR-FL |
| RHOB | 0.73 | 2.50 | Both Isoforms | KRT14 | 1.90 | 0.24 | AR-FL |
| EGR1 | 1.02 | 2.50 | Both Isoforms | ARTN | 1.24 | 0.24 | AR-FL |
| IQUB | 2.04 | 2.51 | Both Isoforms | HERC3 | 1.72 | 0.24 | AR-FL |
| SDK1 | 4.17 | 2.51 | Both Isoforms | HPCAL1 | 0.74 | 0.24 | AR-FL |
| NTN4 | 0.98 | 2.52 | Both Isoforms | ZFAND4 | 1.06 | 0.24 | AR-FL |
| AKAP17A | 2.47 | 2.56 | Both Isoforms | NOP10 | 0.76 | 0.24 | AR-FL |
| FAM83E | 1.57 | 2.57 | Both Isoforms | LMO7 | 0.58 | 0.25 | AR-FL |
| KLK3 | 3.56 | 2.57 | Both Isoforms | MOB4 | 0.74 | 0.25 | AR-FL |
| PER1 | 1.77 | 2.59 | Both Isoforms | PELO | 0.64 | 0.25 | AR-FL |
| SERPIND1 | 2.16 | 2.59 | Both Isoforms | PCDHB2 | 0.66 | 0.25 | AR-FL |
| ANK1 | 2.78 | 2.60 | Both Isoforms | EFNA4 | 1.81 | 0.25 | AR-FL |
| C16orf54 | 5.68 | 2.60 | Both Isoforms | SEC31A | 0.64 | 0.25 | AR-FL |
| ARMC12 | 2.64 | 2.62 | Both Isoforms | KDM5C | 0.61 | 0.25 | AR-FL |
| CCDC160 | 2.71 | 2.64 | Both Isoforms | TPST2 | 1.43 | 0.25 | AR-FL |
| MBOAT2 | 2.84 | 2.65 | Both Isoforms | TEP1 | 1.08 | 0.25 | AR-FL |

| | | | | | | | |
|---|---|---|---|---|---|---|---|
| SAT1 | 2.33 | 2.65 | Both Isoforms | MARK2 | 0.66 | 0.25 | AR-FL |
| CDKN2B | 1.49 | 2.69 | Both Isoforms | ORAI1 | 0.94 | 0.25 | AR-FL |
| KRT6C | 2.59 | 2.69 | Both Isoforms | PAK4 | 0.66 | 0.25 | AR-FL |
| EAF2 | 3.89 | 2.70 | Both Isoforms | EFR3A | 1.18 | 0.26 | AR-FL |
| ITGA11 | 2.29 | 2.71 | Both Isoforms | TMEM223 | 0.74 | 0.26 | AR-FL |
| ASIP | 1.78 | 2.71 | Both Isoforms | MRPL27 | 0.69 | 0.26 | AR-FL |
| SPSB1 | 0.96 | 2.72 | Both Isoforms | 5-Mar | 0.81 | 0.26 | AR-FL |
| GLUD1 | 1.74 | 2.72 | Both Isoforms | AKT1S1 | 0.62 | 0.26 | AR-FL |
| SH2D3A | 0.92 | 2.72 | Both Isoforms | LIMK2 | 0.59 | 0.26 | AR-FL |
| PAPLN | 0.77 | 2.73 | Both Isoforms | DEXI | 0.63 | 0.26 | AR-FL |
| MYO1B | 2.20 | 2.73 | Both Isoforms | CACFD1 | 0.80 | 0.26 | AR-FL |
| SLITRK6 | 2.29 | 2.75 | Both Isoforms | S1PR3 | 1.90 | 0.26 | AR-FL |
| PTPN21 | 2.53 | 2.76 | Both Isoforms | MRPS18A | 0.72 | 0.26 | AR-FL |
| IGF1R | 3.24 | 2.76 | Both Isoforms | BCL2L12 | 1.02 | 0.26 | AR-FL |
| PHLDB2 | 4.28 | 2.76 | Both Isoforms | PRKCA | 0.99 | 0.26 | AR-FL |
| ALPK2 | 3.85 | 2.77 | Both Isoforms | YRDC | 0.64 | 0.26 | AR-FL |
| PNMA1 | 2.04 | 2.77 | Both Isoforms | CRELD1 | 0.86 | 0.26 | AR-FL |
| BTG1 | 0.72 | 2.79 | Both Isoforms | PSMD13 | 0.75 | 0.26 | AR-FL |
| ELOVL5 | 2.34 | 2.79 | Both Isoforms | FBXW10 | 1.68 | 0.26 | AR-FL |
| GADD45B | 1.05 | 2.82 | Both Isoforms | SCAND1 | 1.09 | 0.26 | AR-FL |
| SPATA13 | 1.16 | 2.84 | Both Isoforms | ERH | 0.59 | 0.26 | AR-FL |
| CBWD1 | 2.40 | 2.85 | Both Isoforms | WSB2 | 0.97 | 0.27 | AR-FL |
| LOX | 4.24 | 2.86 | Both Isoforms | SIPA1L3 | 0.63 | 0.27 | AR-FL |
| PMEPA1 | 3.39 | 2.86 | Both Isoforms | RAB11A | 0.94 | 0.27 | AR-FL |
| CRYAB | 0.78 | 2.88 | Both Isoforms | DNAJA4 | 1.03 | 0.27 | AR-FL |

| | | | | | | | |
|---|---|---|---|---|---|---|---|
| RASD1 | 2.66 | 2.89 | Both Isoforms | APOF | 0.71 | 0.27 | AR-FL |
| SLC1A2 | 0.92 | 2.89 | Both Isoforms | BPGM | 0.84 | 0.27 | AR-FL |
| FAM105A | 4.44 | 2.92 | Both Isoforms | MAP7D1 | 1.44 | 0.27 | AR-FL |
| TMPRSS2 | 4.58 | 2.96 | Both Isoforms | MTMR6 | 0.67 | 0.27 | AR-FL |
| NPNT | 1.57 | 2.97 | Both Isoforms | NFIB | 0.58 | 0.27 | AR-FL |
| B2M | 1.53 | 2.99 | Both Isoforms | FAM173B | 0.60 | 0.27 | AR-FL |
| MAP2 | 1.92 | 2.99 | Both Isoforms | HIPK1 | 0.61 | 0.27 | AR-FL |
| ANGPT2 | 5.37 | 3.01 | Both Isoforms | PNKD | 1.28 | 0.27 | AR-FL |
| POTEF | 3.39 | 3.07 | Both Isoforms | ERGIC2 | 0.98 | 0.27 | AR-FL |
| LAMA1 | 3.65 | 3.09 | Both Isoforms | MED8 | 0.60 | 0.27 | AR-FL |
| CAPN2 | 0.95 | 3.09 | Both Isoforms | SNRPA1 | 0.59 | 0.27 | AR-FL |
| GNMT | 3.59 | 3.20 | Both Isoforms | APMAP | 0.89 | 0.27 | AR-FL |
| SLC45A3 | 3.71 | 3.29 | Both Isoforms | BRD2 | 0.67 | 0.27 | AR-FL |
| AZGP1 | 2.43 | 3.32 | Both Isoforms | PIEZO1 | 0.60 | 0.27 | AR-FL |
| WDR46 | 0.64 | 3.34 | Both Isoforms | SLC7A6 | 0.68 | 0.27 | AR-FL |
| POTEJ | 2.26 | 3.35 | Both Isoforms | VTI1A | 0.62 | 0.27 | AR-FL |
| APOD | 1.27 | 3.35 | Both Isoforms | LPCAT3 | 0.89 | 0.27 | AR-FL |
| POTEI | 1.76 | 3.35 | Both Isoforms | BNIP3 | 0.90 | 0.28 | AR-FL |
| SCNN1G | 0.75 | 3.38 | Both Isoforms | PSMD3 | 0.63 | 0.28 | AR-FL |
| SNCG | 3.53 | 3.39 | Both Isoforms | COA6 | 0.60 | 0.28 | AR-FL |
| TMPRSS11E | 2.08 | 3.39 | Both Isoforms | SLC10A7 | 0.66 | 0.28 | AR-FL |
| NNMT | 3.49 | 3.40 | Both Isoforms | ZNF428 | 0.70 | 0.28 | AR-FL |
| POTEE | 3.34 | 3.40 | Both Isoforms | ZDHHC16 | 1.00 | 0.28 | AR-FL |
| PXDC1 | 2.49 | 3.45 | Both Isoforms | TXNDC9 | 0.80 | 0.28 | AR-FL |
| TIPARP | 2.08 | 3.46 | Both Isoforms | MAN1A2 | 0.67 | 0.28 | AR-FL |

| | | | | | | | |
|---|---|---|---|---|---|---|---|
| F2RL1 | 1.18 | 3.46 | Both Isoforms | RRAS | 1.40 | 0.28 | AR-FL |
| MEIS1 | 1.53 | 3.47 | Both Isoforms | USP4 | 0.58 | 0.28 | AR-FL |
| UGT2B11 | 3.31 | 3.49 | Both Isoforms | EXOC3L1 | 0.75 | 0.28 | AR-FL |
| SNAI2 | 5.73 | 3.49 | Both Isoforms | ZFPL1 | 0.78 | 0.29 | AR-FL |
| LIFR | 3.35 | 3.49 | Both Isoforms | MPG | 0.63 | 0.29 | AR-FL |
| KL | 4.93 | 3.52 | Both Isoforms | TTI2 | 0.73 | 0.29 | AR-FL |
| CUX2 | 4.17 | 3.52 | Both Isoforms | FAM89B | 0.76 | 0.29 | AR-FL |
| STK39 | 2.75 | 3.54 | Both Isoforms | MYADM | 0.64 | 0.29 | AR-FL |
| SOCS2 | 2.60 | 3.55 | Both Isoforms | GSR | 0.88 | 0.29 | AR-FL |
| TGFBR2 | 1.69 | 3.55 | Both Isoforms | LAMC1 | 1.59 | 0.29 | AR-FL |
| NDRG1 | 4.52 | 3.56 | Both Isoforms | ANXA7 | 0.62 | 0.29 | AR-FL |
| FOXG1 | 2.32 | 3.64 | Both Isoforms | STK17B | 0.93 | 0.29 | AR-FL |
| KLK2 | 4.55 | 3.68 | Both Isoforms | SRXN1 | 0.61 | 0.29 | AR-FL |
| C8orf4 | 2.90 | 3.70 | Both Isoforms | LPAR3 | 1.26 | 0.29 | AR-FL |
| IRS2 | 2.80 | 3.70 | Both Isoforms | ELOVL1 | 0.66 | 0.29 | AR-FL |
| XKR5 | 3.94 | 3.80 | Both Isoforms | WBP1L | 0.70 | 0.29 | AR-FL |
| NEXN | 1.57 | 3.81 | Both Isoforms | RNF26 | 0.66 | 0.29 | AR-FL |
| ACSL3 | 3.64 | 3.84 | Both Isoforms | SLC25A16 | 0.73 | 0.29 | AR-FL |
| CYP11A1 | 3.28 | 3.88 | Both Isoforms | KIAA0100 | 0.72 | 0.29 | AR-FL |
| FAM107A | 3.11 | 3.89 | Both Isoforms | PEPD | 0.74 | 0.29 | AR-FL |
| AFF3 | 4.07 | 3.89 | Both Isoforms | EIF4E | 0.66 | 0.29 | AR-FL |
| TRIM49D2 | 4.11 | 3.90 | Both Isoforms | KDELR1 | 0.88 | 0.30 | AR-FL |
| IL31RA | 2.30 | 3.92 | Both Isoforms | DTWD2 | 1.04 | 0.30 | AR-FL |
| PTPRN2 | 2.17 | 3.94 | Both Isoforms | DENND1C | 0.63 | 0.30 | AR-FL |
| KRT19 | 2.57 | 3.95 | Both Isoforms | GABARAP | 0.59 | 0.30 | AR-FL |

| | | | | | | | |
|---|---|---|---|---|---|---|---|
| CECR6 | 3.94 | 3.96 | Both Isoforms | CBR1 | 0.60 | 0.30 | AR-FL |
| HES1 | 0.90 | 4.03 | Both Isoforms | CCDC186 | 0.66 | 0.30 | AR-FL |
| NPPC | 5.58 | 4.04 | Both Isoforms | RABIF | 0.73 | 0.30 | AR-FL |
| ITGB8 | 1.30 | 4.05 | Both Isoforms | ASAH1 | 0.94 | 0.30 | AR-FL |
| ACTA2 | 3.04 | 4.06 | Both Isoforms | CCNYL1 | 1.57 | 0.30 | AR-FL |
| ATRNL1 | 2.45 | 4.06 | Both Isoforms | CKMT1A | 1.07 | 0.30 | AR-FL |
| KCNMA1 | 2.98 | 4.06 | Both Isoforms | SLC25A32 | 0.65 | 0.30 | AR-FL |
| HIST1H1E | 3.24 | 4.10 | Both Isoforms | PCMT1 | 0.85 | 0.30 | AR-FL |
| CTNNA2 | 5.53 | 4.11 | Both Isoforms | DCTPP1 | 0.75 | 0.30 | AR-FL |
| MAGEA11 | 2.48 | 4.16 | Both Isoforms | GPKOW | 0.60 | 0.30 | AR-FL |
| UGT2B28 | 5.82 | 4.20 | Both Isoforms | SPARC | 3.36 | 0.30 | AR-FL |
| TNS4 | 3.66 | 4.23 | Both Isoforms | PRAF2 | 0.58 | 0.30 | AR-FL |
| DNER | 1.44 | 4.23 | Both Isoforms | TMEM19 | 0.61 | 0.30 | AR-FL |
| MAOB | 1.54 | 4.39 | Both Isoforms | PCDHB3 | 1.02 | 0.30 | AR-FL |
| FOXD4 | 3.06 | 4.42 | Both Isoforms | AGTRAP | 0.91 | 0.30 | AR-FL |
| ACTG2 | 4.01 | 4.45 | Both Isoforms | CMAS | 0.61 | 0.30 | AR-FL |
| FKBP5 | 4.72 | 4.51 | Both Isoforms | TRMT12 | 0.80 | 0.31 | AR-FL |
| SIPA1L2 | 1.91 | 4.56 | Both Isoforms | CCDC159 | 0.90 | 0.31 | AR-FL |
| CA4 | 1.86 | 4.61 | Both Isoforms | KLHL25 | 0.79 | 0.31 | AR-FL |
| TBX15 | 6.07 | 4.62 | Both Isoforms | CREM | 0.66 | 0.31 | AR-FL |
| ASB9 | 1.67 | 4.64 | Both Isoforms | RAB27A | 1.51 | 0.31 | AR-FL |
| ADAMTS8 | 0.96 | 4.70 | Both Isoforms | GFM1 | 0.83 | 0.31 | AR-FL |
| ST6GALNAC1 | 4.83 | 4.77 | Both Isoforms | CHCHD1 | 0.62 | 0.31 | AR-FL |
| MYH7 | 1.77 | 4.85 | Both Isoforms | CRY1 | 1.18 | 0.31 | AR-FL |
| SULT1B1 | 2.16 | 4.92 | Both Isoforms | RHOU | 1.35 | 0.31 | AR-FL |

| | | | | | | | |
|---|---|---|---|---|---|---|---|
| EDN1 | 2.28 | 4.95 | Both Isoforms | MED27 | 0.83 | 0.31 | AR-FL |
| DLX1 | 2.77 | 5.01 | Both Isoforms | UQCC3 | 0.83 | 0.31 | AR-FL |
| ERRFI1 | 2.99 | 5.03 | Both Isoforms | INPPL1 | 0.62 | 0.31 | AR-FL |
| KRT73 | 6.95 | 5.03 | Both Isoforms | LARP4 | 0.62 | 0.31 | AR-FL |
| CHN2 | 2.47 | 5.04 | Both Isoforms | ASAP3 | 0.78 | 0.31 | AR-FL |
| MYH6 | 1.24 | 5.04 | Both Isoforms | UBE2J1 | 0.99 | 0.31 | AR-FL |
| ADAM7 | 3.87 | 5.06 | Both Isoforms | ZNF44 | 0.65 | 0.31 | AR-FL |
| PTGER4 | 3.90 | 5.07 | Both Isoforms | PTPN1 | 0.82 | 0.31 | AR-FL |
| ELL2 | 2.85 | 5.09 | Both Isoforms | NSF | 0.72 | 0.31 | AR-FL |
| RCSD1 | 4.44 | 5.20 | Both Isoforms | PROM2 | 0.62 | 0.31 | AR-FL |
| KY | 2.69 | 5.31 | Both Isoforms | PCDHGB1 | 0.91 | 0.31 | AR-FL |
| GP2 | 2.73 | 5.33 | Both Isoforms | PGK1 | 0.83 | 0.31 | AR-FL |
| SLC15A2 | 4.55 | 5.34 | Both Isoforms | ACP2 | 0.85 | 0.32 | AR-FL |
| TENM1 | 2.92 | 5.36 | Both Isoforms | TM9SF2 | 0.74 | 0.32 | AR-FL |
| TMCC3 | 5.06 | 5.45 | Both Isoforms | HAT1 | 0.81 | 0.32 | AR-FL |
| SPARC | 2.32 | 5.48 | Both Isoforms | ALDH16A1 | 0.63 | 0.32 | AR-FL |
| IVL | 2.11 | 5.56 | Both Isoforms | SLC2A7 | 0.72 | 0.32 | AR-FL |
| MYL7 | 2.99 | 5.69 | Both Isoforms | PSMA3 | 0.64 | 0.32 | AR-FL |
| TM4SF18 | 3.31 | 5.76 | Both Isoforms | SLC30A7 | 0.90 | 0.32 | AR-FL |
| F5 | 6.56 | 5.80 | Both Isoforms | THUMPD1 | 0.86 | 0.32 | AR-FL |
| PLA2G4E | 5.81 | 5.94 | Both Isoforms | CNOT3 | 0.64 | 0.32 | AR-FL |
| EMP1 | 4.57 | 6.06 | Both Isoforms | TIMM23 | 0.80 | 0.32 | AR-FL |
| PGC | 6.41 | 6.12 | Both Isoforms | RSPH9 | 0.62 | 0.32 | AR-FL |
| SLC6A3 | 2.12 | 6.16 | Both Isoforms | LAMA5 | 0.64 | 0.32 | AR-FL |
| TUBA3D | 4.22 | 6.19 | Both Isoforms | PSMA5 | 0.68 | 0.32 | AR-FL |

| | | | | | | | |
|---|---|---|---|---|---|---|---|
| PNLIPRP3 | 3.29 | 6.20 | Both Isoforms | PRKD2 | 1.11 | 0.32 | AR-FL |
| SERPINF2 | 2.62 | 6.21 | Both Isoforms | HCCS | 0.60 | 0.32 | AR-FL |
| STEAP4 | 10.11 | 6.36 | Both Isoforms | TMEM11 | 0.76 | 0.32 | AR-FL |
| NECAB1 | 0.70 | 6.44 | Both Isoforms | PYCR1 | 0.82 | 0.32 | AR-FL |
| MOGAT2 | 5.01 | 6.48 | Both Isoforms | ODF2 | 0.59 | 0.32 | AR-FL |
| TRIM49D1 | 5.86 | 6.58 | Both Isoforms | C1GALT1C1 | 0.75 | 0.32 | AR-FL |
| ZNF812 | 6.00 | 6.60 | Both Isoforms | HN1 | 0.82 | 0.33 | AR-FL |
| HOPX | 0.92 | 6.66 | Both Isoforms | PARK7 | 0.83 | 0.33 | AR-FL |
| MYBPC1 | 5.03 | 6.81 | Both Isoforms | ARL1 | 0.68 | 0.33 | AR-FL |
| DOCK8 | 5.74 | 6.85 | Both Isoforms | C21orf59 | 0.68 | 0.33 | AR-FL |
| PAEP | 3.31 | 6.95 | Both Isoforms | CREB3 | 1.07 | 0.33 | AR-FL |
| PDE6A | 2.09 | 6.97 | Both Isoforms | VAV2 | 0.80 | 0.33 | AR-FL |
| SLC26A3 | 5.99 | 7.00 | Both Isoforms | CLPB | 0.61 | 0.33 | AR-FL |
| SPOCK1 | 4.55 | 7.17 | Both Isoforms | MTM1 | 0.66 | 0.33 | AR-FL |
| NR5A2 | 1.28 | 7.27 | Both Isoforms | APLP1 | 0.73 | 0.33 | AR-FL |
| SLC2A3 | 10.76 | 7.46 | Both Isoforms | SETX | 0.60 | 0.33 | AR-FL |
| NPR3 | 1.04 | 7.57 | Both Isoforms | ROM1 | 0.90 | 0.33 | AR-FL |
| TUBA3E | 5.41 | 7.72 | Both Isoforms | PPP2CB | 0.69 | 0.33 | AR-FL |
| NNT | 4.40 | 7.72 | Both Isoforms | DYNLL2 | 0.75 | 0.33 | AR-FL |
| KRT6A | 2.68 | 7.98 | Both Isoforms | ACOX1 | 1.11 | 0.33 | AR-FL |
| ANKRD1 | 2.21 | 8.06 | Both Isoforms | ATP6V1E1 | 0.69 | 0.33 | AR-FL |
| TGM2 | 5.15 | 8.13 | Both Isoforms | CAMTA2 | 1.11 | 0.33 | AR-FL |
| CCDC141 | 9.88 | 8.15 | Both Isoforms | COMT | 0.70 | 0.33 | AR-FL |
| CHRNA2 | 6.63 | 8.36 | Both Isoforms | GOLPH3 | 0.88 | 0.33 | AR-FL |
| AKR1D1 | 6.08 | 8.36 | Both Isoforms | TRAPPC3 | 0.69 | 0.33 | AR-FL |

| | | | |
|---|---|---|---|
| GHRHR | 5.90 | 8.40 | Both Isoforms |
| SPINK13 | 1.90 | 8.40 | Both Isoforms |
| PDK4 | 3.44 | 8.54 | Both Isoforms |
| HPGD | 6.77 | 8.54 | Both Isoforms |
| IP6K3 | 3.75 | 8.56 | Both Isoforms |
| AKAP12 | 6.04 | 8.71 | Both Isoforms |
| OLAH | 3.74 | 8.74 | Both Isoforms |
| ACADL | 6.25 | 8.89 | Both Isoforms |
| FBLN5 | 1.51 | 9.70 | Both Isoforms |
| DCN | 3.29 | 9.77 | Both Isoforms |
| CXCL13 | 7.06 | 9.88 | Both Isoforms |
| SLC38A4 | 9.13 | 10.41 | Both Isoforms |
| CTSE | 4.78 | 10.46 | Both Isoforms |
| ORM2 | 6.39 | 10.89 | Both Isoforms |
| PNLIP | 9.72 | 12.00 | Both Isoforms |
| ORM1 | 6.65 | 12.38 | Both Isoforms |
| PCDH7 | 2.43 | 12.43 | Both Isoforms |

| | | | |
|---|---|---|---|
| H2AFY | 0.65 | 0.33 | AR-FL |
| NAPG | 0.72 | 0.33 | AR-FL |
| SMURF1 | 0.76 | 0.34 | AR-FL |
| SORD | 0.63 | 0.34 | AR-FL |
| ZNF672 | 0.91 | 0.34 | AR-FL |
| CLCC1 | 0.59 | 0.34 | AR-FL |
| PVR | 0.65 | 0.34 | AR-FL |
| OPA3 | 0.80 | 0.34 | AR-FL |
| SMIM4 | 0.72 | 0.34 | AR-FL |
| VWA1 | 0.72 | 0.34 | AR-FL |
| GRIN1 | 0.82 | 0.34 | AR-FL |
| GSPT1 | 0.74 | 0.34 | AR-FL |
| GSKIP | 1.00 | 0.34 | AR-FL |
| ARID1A | 0.64 | 0.34 | AR-FL |
| YWHAB | 0.59 | 0.34 | AR-FL |
| PSMC1 | 0.73 | 0.34 | AR-FL |
| DDX3X | 0.63 | 0.34 | AR-FL |
| NDUFB1 | 0.66 | 0.34 | AR-FL |
| ODC1 | 2.64 | 0.34 | AR-FL |
| TMEM50A | 1.08 | 0.34 | AR-FL |
| SAP30 | 0.77 | 0.34 | AR-FL |
| AUP1 | 0.78 | 0.34 | AR-FL |
| ZBTB17 | 0.66 | 0.35 | AR-FL |
| HIGD1A | 0.77 | 0.35 | AR-FL |
| PPCS | 0.74 | 0.35 | AR-FL |
| RREB1 | 0.59 | 0.35 | AR-FL |
| IGF2BP3 | 0.72 | 0.35 | AR-FL |
| SMDT1 | 0.77 | 0.35 | AR-FL |
| FGFR3 | 1.64 | 0.35 | AR-FL |
| ABCB6 | 0.60 | 0.35 | AR-FL |
| KMT2D | 0.80 | 0.35 | AR-FL |
| GTF2A2 | 0.65 | 0.35 | AR-FL |

| | | | |
|---|---|---|---|
| NOX5 | 0.90 | 0.35 | AR-FL |
| MAP3K9 | 0.67 | 0.35 | AR-FL |
| AP1G1 | 0.73 | 0.35 | AR-FL |
| RPL36AL | 0.63 | 0.35 | AR-FL |
| ARPC2 | 0.63 | 0.35 | AR-FL |
| XPNPEP1 | 0.65 | 0.35 | AR-FL |
| PLIN3 | 0.91 | 0.35 | AR-FL |
| LGALSL | 1.05 | 0.35 | AR-FL |
| TMEM87B | 1.33 | 0.35 | AR-FL |
| TMEM9B | 0.59 | 0.35 | AR-FL |
| ELP6 | 0.71 | 0.35 | AR-FL |
| PPFIA3 | 1.13 | 0.35 | AR-FL |
| CRB3 | 0.75 | 0.36 | AR-FL |
| CTNS | 0.66 | 0.36 | AR-FL |
| C6orf1 | 0.73 | 0.36 | AR-FL |
| ST3GAL4 | 1.26 | 0.36 | AR-FL |
| HIST1H3E | 0.64 | 0.36 | AR-FL |
| KIAA1549 | 1.16 | 0.36 | AR-FL |
| HMOX1 | 1.21 | 0.36 | AR-FL |
| RAB8A | 0.66 | 0.36 | AR-FL |
| OTUB1 | 0.61 | 0.36 | AR-FL |
| NDUFC1 | 0.75 | 0.36 | AR-FL |
| TTC13 | 0.69 | 0.36 | AR-FL |
| FAM98A | 0.69 | 0.36 | AR-FL |
| ZNF628 | 0.63 | 0.36 | AR-FL |
| ZNF440 | 0.63 | 0.36 | AR-FL |
| MICAL1 | 2.39 | 0.36 | AR-FL |
| COG3 | 0.76 | 0.36 | AR-FL |
| KIF5C | 0.59 | 0.36 | AR-FL |
| BTBD3 | 0.65 | 0.36 | AR-FL |
| SLC19A1 | 0.65 | 0.36 | AR-FL |
| ARCN1 | 0.79 | 0.36 | AR-FL |
| CTDP1 | 0.65 | 0.36 | AR-FL |
| PLPP6 | 1.00 | 0.36 | AR-FL |
| CD9 | 1.35 | 0.36 | AR-FL |
| RAB3D | 1.31 | 0.36 | AR-FL |
| BNIP1 | 0.60 | 0.36 | AR-FL |
| COMMD7 | 0.61 | 0.36 | AR-FL |
| DISP1 | 0.62 | 0.36 | AR-FL |
| SLC25A45 | 0.96 | 0.37 | AR-FL |
| ERCC4 | 0.62 | 0.37 | AR-FL |
| MYO1C | 1.01 | 0.37 | AR-FL |
| C19orf24 | 0.70 | 0.37 | AR-FL |
| MCOLN1 | 0.86 | 0.37 | AR-FL |
| C3orf70 | 0.75 | 0.37 | AR-FL |
| MTCH2 | 0.80 | 0.37 | AR-FL |

| | | | |
|---|---|---|---|
| MRPL20 | 0.58 | 0.37 | AR-FL |
| EPN3 | 0.82 | 0.37 | AR-FL |
| SLC45A1 | 0.76 | 0.37 | AR-FL |
| NARF | 1.08 | 0.37 | AR-FL |
| NDUFB3 | 0.65 | 0.37 | AR-FL |
| TRADD | 0.75 | 0.37 | AR-FL |
| FUT11 | 0.95 | 0.37 | AR-FL |
| PDXDC1 | 0.72 | 0.37 | AR-FL |
| SMAP1 | 0.85 | 0.37 | AR-FL |
| GOLGB1 | 0.79 | 0.37 | AR-FL |
| SIGIRR | 0.80 | 0.37 | AR-FL |
| ABHD13 | 0.62 | 0.37 | AR-FL |
| PMM2 | 0.67 | 0.37 | AR-FL |
| MKL1 | 0.63 | 0.37 | AR-FL |
| NDUFA8 | 0.64 | 0.37 | AR-FL |
| NAA16 | 0.58 | 0.37 | AR-FL |
| TBC1D7 | 0.74 | 0.37 | AR-FL |
| GK5 | 0.67 | 0.37 | AR-FL |
| TFPT | 1.28 | 0.38 | AR-FL |
| UQCRFS1 | 0.72 | 0.38 | AR-FL |
| MLPH | 0.65 | 0.38 | AR-FL |
| CHCHD3 | 0.63 | 0.38 | AR-FL |
| SLC52A3 | 0.65 | 0.38 | AR-FL |
| FAM177A1 | 0.84 | 0.38 | AR-FL |
| ECM1 | 1.15 | 0.38 | AR-FL |
| LAGE3 | 0.78 | 0.38 | AR-FL |
| OGDH | 0.94 | 0.38 | AR-FL |
| BAX | 0.73 | 0.38 | AR-FL |
| SPDEF | 2.16 | 0.38 | AR-FL |
| RPS19BP1 | 0.76 | 0.38 | AR-FL |
| ATG101 | 0.94 | 0.38 | AR-FL |
| TRPM4 | 0.99 | 0.38 | AR-FL |
| RUSC1 | 1.49 | 0.38 | AR-FL |
| TBXA2R | 0.94 | 0.38 | AR-FL |
| ALG11 | 0.66 | 0.38 | AR-FL |
| ARHGAP26 | 2.12 | 0.38 | AR-FL |
| PKM | 0.77 | 0.38 | AR-FL |
| POLD4 | 0.59 | 0.38 | AR-FL |
| HIST1H2AE | 0.79 | 0.38 | AR-FL |
| NAPRT | 0.90 | 0.38 | AR-FL |
| MYH9 | 0.82 | 0.39 | AR-FL |
| LRG1 | 1.46 | 0.39 | AR-FL |
| CENPBD1 | 0.68 | 0.39 | AR-FL |
| BEST3 | 0.77 | 0.39 | AR-FL |
| VEGFA | 0.70 | 0.39 | AR-FL |
| SCAMP2 | 0.80 | 0.39 | AR-FL |

| | | | |
|---|---|---|---|
| ACTR2 | 0.73 | 0.39 | AR-FL |
| ANO10 | 0.64 | 0.39 | AR-FL |
| SLC30A5 | 0.60 | 0.39 | AR-FL |
| IGHMBP2 | 0.81 | 0.39 | AR-FL |
| SEC63 | 0.69 | 0.39 | AR-FL |
| GJB1 | 1.07 | 0.39 | AR-FL |
| STX3 | 0.82 | 0.39 | AR-FL |
| COPB2 | 0.98 | 0.39 | AR-FL |
| MESDC1 | 1.26 | 0.39 | AR-FL |
| DAPK3 | 0.73 | 0.39 | AR-FL |
| SSNA1 | 0.75 | 0.39 | AR-FL |
| TOR2A | 0.86 | 0.39 | AR-FL |
| ZNF678 | 0.70 | 0.40 | AR-FL |
| CHUK | 0.77 | 0.40 | AR-FL |
| FXR2 | 0.86 | 0.40 | AR-FL |
| GOLT1B | 0.95 | 0.40 | AR-FL |
| TMEM222 | 0.60 | 0.40 | AR-FL |
| AES | 0.68 | 0.40 | AR-FL |
| DTX1 | 1.86 | 0.40 | AR-FL |
| CARTPT | 1.21 | 0.40 | AR-FL |
| EIF4G2 | 1.00 | 0.40 | AR-FL |
| TMC6 | 1.03 | 0.40 | AR-FL |
| PPP4C | 0.63 | 0.40 | AR-FL |
| RAC3 | 0.60 | 0.40 | AR-FL |
| ARPC4 | 0.94 | 0.40 | AR-FL |
| USP38 | 0.75 | 0.40 | AR-FL |
| IDH2 | 0.84 | 0.40 | AR-FL |
| BCL2L2 | 0.89 | 0.40 | AR-FL |
| GNA11 | 0.60 | 0.40 | AR-FL |
| HAGH | 0.70 | 0.40 | AR-FL |
| BAD | 0.59 | 0.40 | AR-FL |
| DPM3 | 1.12 | 0.40 | AR-FL |
| PITPNA | 1.04 | 0.40 | AR-FL |
| LAMTOR4 | 0.66 | 0.40 | AR-FL |
| FTSJ1 | 0.58 | 0.40 | AR-FL |
| AP3D1 | 0.63 | 0.41 | AR-FL |
| DUSP1 | 1.46 | 0.41 | AR-FL |
| SPATS2 | 0.68 | 0.41 | AR-FL |
| KCTD9 | 0.59 | 0.41 | AR-FL |
| RP11-111H13.1 | 1.17 | 0.41 | AR-FL |
| DOHH | 0.61 | 0.41 | AR-FL |
| NIPA2 | 0.86 | 0.41 | AR-FL |
| SBNO2 | 0.61 | 0.41 | AR-FL |
| G3BP2 | 1.39 | 0.41 | AR-FL |
| ZDHHC5 | 0.65 | 0.41 | AR-FL |
| BLOC1S1 | 0.63 | 0.41 | AR-FL |

| | | | |
|---|---|---|---|
| NEK3 | 1.35 | 0.41 | AR-FL |
| FKBP14 | 0.61 | 0.41 | AR-FL |
| TGFB2 | 1.41 | 0.41 | AR-FL |
| ELP5 | 0.68 | 0.41 | AR-FL |
| TNS3 | 0.70 | 0.41 | AR-FL |
| NAPA | 0.60 | 0.41 | AR-FL |
| LYPLA2 | 1.03 | 0.41 | AR-FL |
| ENO1 | 0.89 | 0.41 | AR-FL |
| SNRNP40 | 0.63 | 0.41 | AR-FL |
| CDC42SE1 | 1.01 | 0.41 | AR-FL |
| PPM1H | 1.48 | 0.41 | AR-FL |
| SCO2 | 0.64 | 0.41 | AR-FL |
| RANGRF | 1.46 | 0.41 | AR-FL |
| PPP6R1 | 0.66 | 0.41 | AR-FL |
| COPRS | 0.92 | 0.41 | AR-FL |
| POR | 0.66 | 0.41 | AR-FL |
| KLK15 | 2.04 | 0.41 | AR-FL |
| IRAK1 | 0.59 | 0.41 | AR-FL |
| BCAP31 | 0.79 | 0.41 | AR-FL |
| DYNLRB1 | 0.66 | 0.41 | AR-FL |
| COX6B1 | 0.80 | 0.41 | AR-FL |
| UNC13A | 0.90 | 0.41 | AR-FL |
| PLEKHF1 | 0.87 | 0.42 | AR-FL |
| HNRNPUL2-BSCL2 | 0.75 | 0.42 | AR-FL |
| ZNF410 | 1.12 | 0.42 | AR-FL |
| MPZL1 | 1.10 | 0.42 | AR-FL |
| MAPK7 | 0.84 | 0.42 | AR-FL |
| STT3B | 0.62 | 0.42 | AR-FL |
| SNRPG | 0.65 | 0.42 | AR-FL |
| ZNHIT2 | 0.65 | 0.42 | AR-FL |
| ARPC3 | 1.10 | 0.42 | AR-FL |
| SLC2A1 | 0.82 | 0.42 | AR-FL |
| AKT1 | 0.90 | 0.42 | AR-FL |
| TIMM10B | 0.72 | 0.42 | AR-FL |
| RNGTT | 0.65 | 0.42 | AR-FL |
| RPS6KA1 | 1.09 | 0.42 | AR-FL |
| CYP2U1 | 1.17 | 0.42 | AR-FL |
| KLK2 | 1.03 | 0.42 | AR-FL |
| SLC35A4 | 0.69 | 0.42 | AR-FL |
| TRAPPC9 | 0.62 | 0.42 | AR-FL |
| CLINT1 | 0.66 | 0.42 | AR-FL |
| PDIA5 | 1.17 | 0.42 | AR-FL |
| ADRBK1 | 1.40 | 0.42 | AR-FL |
| NR2F6 | 0.73 | 0.43 | AR-FL |
| SAMHD1 | 0.99 | 0.43 | AR-FL |

| | | | |
|---|---|---|---|
| ERG | 1.77 | 0.43 | AR-FL |
| CANX | 0.67 | 0.43 | AR-FL |
| HCFC1 | 0.77 | 0.43 | AR-FL |
| EIF4G1 | 0.72 | 0.43 | AR-FL |
| TMPRSS2 | 3.28 | 0.43 | AR-FL |
| PIAS3 | 1.00 | 0.43 | AR-FL |
| IL17RC | 0.59 | 0.43 | AR-FL |
| RHOG | 0.60 | 0.43 | AR-FL |
| ROGDI | 1.83 | 0.43 | AR-FL |
| JKAMP | 1.03 | 0.43 | AR-FL |
| GORASP2 | 0.86 | 0.43 | AR-FL |
| BCL7B | 0.74 | 0.43 | AR-FL |
| FRK | 1.32 | 0.43 | AR-FL |
| GNB2 | 0.98 | 0.43 | AR-FL |
| NDUFA1 | 0.69 | 0.43 | AR-FL |
| ALG5 | 0.69 | 0.43 | AR-FL |
| S100A11 | 0.84 | 0.43 | AR-FL |
| MIS12 | 0.68 | 0.43 | AR-FL |
| SERF2 | 0.74 | 0.43 | AR-FL |
| TSPAN17 | 0.69 | 0.43 | AR-FL |
| POLR3E | 1.07 | 0.43 | AR-FL |
| SMG5 | 0.83 | 0.43 | AR-FL |
| BCL2L1 | 1.05 | 0.44 | AR-FL |
| NFATC2IP | 0.81 | 0.44 | AR-FL |
| MLLT11 | 0.87 | 0.44 | AR-FL |
| WDR77 | 0.71 | 0.44 | AR-FL |
| SLC5A6 | 0.80 | 0.44 | AR-FL |
| EXOSC4 | 0.72 | 0.44 | AR-FL |
| CISD2 | 0.62 | 0.44 | AR-FL |
| PYCR2 | 0.63 | 0.44 | AR-FL |
| STEAP4 | 4.52 | 0.44 | AR-FL |
| RAB35 | 0.60 | 0.44 | AR-FL |
| ZSWIM8 | 0.61 | 0.44 | AR-FL |
| TLDC2 | 0.99 | 0.44 | AR-FL |
| FAM222A | 0.96 | 0.44 | AR-FL |
| GDPD5 | 0.79 | 0.44 | AR-FL |
| SELM | 1.20 | 0.44 | AR-FL |
| TRAM1 | 0.87 | 0.44 | AR-FL |
| TMCO1 | 0.67 | 0.44 | AR-FL |
| IL17RA | 0.69 | 0.44 | AR-FL |
| EIF4G3 | 0.75 | 0.44 | AR-FL |
| WWC2 | 0.88 | 0.44 | AR-FL |
| PROB1 | 0.72 | 0.44 | AR-FL |
| SLC22A5 | 1.40 | 0.45 | AR-FL |
| HIVEP3 | 0.77 | 0.45 | AR-FL |
| UCHL3 | 0.73 | 0.45 | AR-FL |

| | | | |
|---|---|---|---|
| SGSH | 0.99 | 0.45 | AR-FL |
| TMEM199 | 0.87 | 0.45 | AR-FL |
| BOLA3 | 0.65 | 0.45 | AR-FL |
| SPOCK3 | 0.85 | 0.45 | AR-FL |
| RAVER1 | 0.64 | 0.45 | AR-FL |
| RABGEF1 | 0.61 | 0.45 | AR-FL |
| PPP1R15B | 0.70 | 0.45 | AR-FL |
| C11orf84 | 0.61 | 0.45 | AR-FL |
| ATXN7L3 | 0.72 | 0.45 | AR-FL |
| TMED5 | 0.91 | 0.45 | AR-FL |
| SCAMP4 | 0.62 | 0.45 | AR-FL |
| GRID2IP | 0.64 | 0.45 | AR-FL |
| SETD3 | 0.81 | 0.45 | AR-FL |
| AIFM1 | 0.77 | 0.45 | AR-FL |
| PSME4 | 1.05 | 0.45 | AR-FL |
| RNF103 | 0.89 | 0.45 | AR-FL |
| ATP5G3 | 0.68 | 0.45 | AR-FL |
| HDLBP | 0.67 | 0.45 | AR-FL |
| COX18 | 0.79 | 0.45 | AR-FL |
| LDHA | 1.15 | 0.45 | AR-FL |
| FAM126B | 0.71 | 0.46 | AR-FL |
| SLC9A3R2 | 1.76 | 0.46 | AR-FL |
| FZD4 | 0.99 | 0.46 | AR-FL |
| CDC42EP4 | 1.18 | 0.46 | AR-FL |
| MFSD12 | 0.71 | 0.46 | AR-FL |
| SNAPC5 | 0.63 | 0.46 | AR-FL |
| SH3TC1 | 1.23 | 0.46 | AR-FL |
| PUF60 | 0.60 | 0.46 | AR-FL |
| MON1B | 0.80 | 0.46 | AR-FL |
| PACSIN1 | 1.10 | 0.46 | AR-FL |
| ALAS1 | 1.07 | 0.46 | AR-FL |
| GADD45B | 0.81 | 0.46 | AR-FL |
| DBNL | 0.63 | 0.46 | AR-FL |
| OLIG1 | 0.71 | 0.46 | AR-FL |
| RPS6KA4 | 1.22 | 0.46 | AR-FL |
| IRX4 | 1.19 | 0.46 | AR-FL |
| GOLGA2 | 1.00 | 0.46 | AR-FL |
| SERP1 | 0.85 | 0.46 | AR-FL |
| ANKRD52 | 0.82 | 0.46 | AR-FL |
| TMEM258 | 0.59 | 0.46 | AR-FL |
| BCL9L | 1.05 | 0.46 | AR-FL |
| DPM2 | 0.74 | 0.46 | AR-FL |
| CLN8 | 0.68 | 0.46 | AR-FL |
| SMIM10L1 | 0.66 | 0.46 | AR-FL |
| CALM1 | 0.73 | 0.46 | AR-FL |
| MARVELD2 | 0.68 | 0.46 | AR-FL |

| | | | |
|---|---|---|---|
| SNRPB | 0.65 | 0.46 | AR-FL |
| TECPR1 | 1.33 | 0.46 | AR-FL |
| PSME2 | 0.90 | 0.46 | AR-FL |
| HIST1H2BH | 0.76 | 0.46 | AR-FL |
| KPNA1 | 0.95 | 0.46 | AR-FL |
| ZWILCH | 0.58 | 0.46 | AR-FL |
| MRPL52 | 0.66 | 0.46 | AR-FL |
| DPP3 | 1.28 | 0.47 | AR-FL |
| TNKS1BP1 | 0.61 | 0.47 | AR-FL |
| TSSC4 | 0.69 | 0.47 | AR-FL |
| MAP3K11 | 0.98 | 0.47 | AR-FL |
| GPR137 | 1.12 | 0.47 | AR-FL |
| FAM91A1 | 0.76 | 0.47 | AR-FL |
| ATP6V1D | 0.83 | 0.47 | AR-FL |
| TUBB2A | 1.71 | 0.47 | AR-FL |
| DNAJA1 | 0.63 | 0.47 | AR-FL |
| NDUFA6 | 0.82 | 0.47 | AR-FL |
| PSME3 | 0.71 | 0.47 | AR-FL |
| FBXW5 | 0.86 | 0.47 | AR-FL |
| POMP | 0.64 | 0.47 | AR-FL |
| TMEM208 | 0.59 | 0.47 | AR-FL |
| NDUFB10 | 1.06 | 0.47 | AR-FL |
| ZNF511 | 0.89 | 0.47 | AR-FL |
| MEMO1 | 0.68 | 0.47 | AR-FL |
| CLTC | 0.72 | 0.47 | AR-FL |
| FAM57A | 0.89 | 0.47 | AR-FL |
| STK40 | 1.11 | 0.47 | AR-FL |
| CCDC124 | 0.72 | 0.47 | AR-FL |
| LSR | 0.90 | 0.47 | AR-FL |
| SEC22B | 0.91 | 0.47 | AR-FL |
| LSM1 | 0.71 | 0.47 | AR-FL |
| C2CD3 | 0.62 | 0.47 | AR-FL |
| PPL | 0.93 | 0.48 | AR-FL |
| ASPSCR1 | 0.62 | 0.48 | AR-FL |
| ZNF653 | 0.75 | 0.48 | AR-FL |
| TIMM17A | 0.72 | 0.48 | AR-FL |
| TVP23C | 0.96 | 0.48 | AR-FL |
| TOMM40L | 0.63 | 0.48 | AR-FL |
| PSMC2 | 0.92 | 0.48 | AR-FL |
| FASTKD5 | 0.69 | 0.48 | AR-FL |
| ZNF35 | 1.07 | 0.48 | AR-FL |
| TRIM5 | 1.12 | 0.48 | AR-FL |
| NDUFS5 | 0.62 | 0.48 | AR-FL |
| WDR24 | 0.78 | 0.48 | AR-FL |
| ATP6V0C | 1.01 | 0.48 | AR-FL |
| JMJD8 | 0.78 | 0.48 | AR-FL |

| | | | |
|---|---|---|---|
| COMMD5 | 1.09 | 0.48 | AR-FL |
| RAB3B | 1.16 | 0.48 | AR-FL |
| NABP2 | 0.73 | 0.48 | AR-FL |
| VPS13D | 0.88 | 0.48 | AR-FL |
| GBA | 1.04 | 0.48 | AR-FL |
| RBMS1 | 1.71 | 0.49 | AR-FL |
| NUDC | 0.59 | 0.49 | AR-FL |
| PPP1CA | 0.80 | 0.49 | AR-FL |
| TMEM63B | 1.12 | 0.49 | AR-FL |
| ENTPD4 | 0.76 | 0.49 | AR-FL |
| ATP6V0A2 | 1.37 | 0.49 | AR-FL |
| TPRKB | 0.73 | 0.49 | AR-FL |
| TMEM33 | 0.90 | 0.49 | AR-FL |
| YOD1 | 0.78 | 0.49 | AR-FL |
| GPRIN1 | 0.58 | 0.49 | AR-FL |
| SIL1 | 0.75 | 0.49 | AR-FL |
| ARF1 | 0.92 | 0.49 | AR-FL |
| IFT140 | 0.80 | 0.50 | AR-FL |
| UBLCP1 | 0.73 | 0.50 | AR-FL |
| GRINA | 0.73 | 0.50 | AR-FL |
| NXPH3 | 0.80 | 0.50 | AR-FL |
| RNF145 | 0.58 | 0.50 | AR-FL |
| LRWD1 | 0.72 | 0.50 | AR-FL |
| TMEM191B | 0.86 | 0.50 | AR-FL |
| ZNF611 | 0.92 | 0.50 | AR-FL |
| SDF4 | 0.77 | 0.50 | AR-FL |
| PDE8B | 0.77 | 0.50 | AR-FL |
| GET4 | 0.72 | 0.50 | AR-FL |
| SCYL1 | 0.97 | 0.50 | AR-FL |
| SMIM12 | 0.60 | 0.50 | AR-FL |
| ZNF66 | 0.98 | 0.50 | AR-FL |
| DNAJC25 | 0.67 | 0.50 | AR-FL |
| EMC7 | 0.67 | 0.50 | AR-FL |
| AAGAB | 0.76 | 0.50 | AR-FL |
| TMEM167A | 0.83 | 0.50 | AR-FL |
| SLC25A48 | 0.66 | 0.50 | AR-FL |
| HSPA13 | 0.68 | 0.50 | AR-FL |
| PRKAR2A | 0.84 | 0.51 | AR-FL |
| IL20RA | 2.02 | 0.51 | AR-FL |
| VPS53 | 1.02 | 0.51 | AR-FL |
| SWSAP1 | 0.90 | 0.51 | AR-FL |
| TMEM251 | 1.18 | 0.51 | AR-FL |
| ATP5E | 0.66 | 0.51 | AR-FL |
| CDC42EP2 | 2.55 | 0.51 | AR-FL |
| MRPL34 | 0.82 | 0.51 | AR-FL |
| ANKRD13D | 0.84 | 0.51 | AR-FL |

| | | | |
|---|---|---|---|
| SDHAF2 | 0.76 | 0.51 | AR-FL |
| COASY | 0.65 | 0.51 | AR-FL |
| POLR2K | 0.77 | 0.51 | AR-FL |
| PGAM5 | 0.69 | 0.51 | AR-FL |
| TAF10 | 0.70 | 0.51 | AR-FL |
| ASNA1 | 0.64 | 0.51 | AR-FL |
| CLPTM1L | 0.79 | 0.52 | AR-FL |
| GNG5 | 0.73 | 0.52 | AR-FL |
| ACOT7 | 0.65 | 0.52 | AR-FL |
| PTGES2 | 0.58 | 0.52 | AR-FL |
| FGD4 | 0.73 | 0.52 | AR-FL |
| SH3BP1 | 0.62 | 0.52 | AR-FL |
| ACTR1A | 1.13 | 0.52 | AR-FL |
| RER1 | 0.66 | 0.52 | AR-FL |
| CAPZB | 1.08 | 0.52 | AR-FL |
| ANGPT2 | 2.54 | 0.52 | AR-FL |
| HEXIM1 | 0.94 | 0.52 | AR-FL |
| EMC9 | 0.91 | 0.52 | AR-FL |
| GLB1L2 | 0.80 | 0.52 | AR-FL |
| VPS18 | 0.81 | 0.52 | AR-FL |
| CCPG1 | 0.65 | 0.52 | AR-FL |
| FAM219A | 1.09 | 0.52 | AR-FL |
| SOCS4 | 0.83 | 0.52 | AR-FL |
| C16orf71 | 0.79 | 0.52 | AR-FL |
| RHOT2 | 0.65 | 0.52 | AR-FL |
| BIRC5 | 0.65 | 0.52 | AR-FL |
| ATF5 | 1.00 | 0.52 | AR-FL |
| TPM3 | 0.85 | 0.52 | AR-FL |
| C10orf35 | 0.61 | 0.52 | AR-FL |
| FHOD1 | 1.02 | 0.52 | AR-FL |
| HCFC2 | 0.70 | 0.53 | AR-FL |
| PICALM | 0.68 | 0.53 | AR-FL |
| EFR3B | 1.20 | 0.53 | AR-FL |
| CAMK1 | 1.09 | 0.53 | AR-FL |
| OST4 | 0.66 | 0.53 | AR-FL |
| GOLGA4 | 0.83 | 0.53 | AR-FL |
| FRMD8 | 0.72 | 0.53 | AR-FL |
| SLC4A2 | 0.71 | 0.53 | AR-FL |
| MFAP3L | 0.63 | 0.53 | AR-FL |
| SOGA1 | 0.69 | 0.53 | AR-FL |
| TSPAN14 | 0.83 | 0.53 | AR-FL |
| CYB5B | 0.67 | 0.53 | AR-FL |
| UBL5 | 0.60 | 0.53 | AR-FL |
| C19orf47 | 0.66 | 0.53 | AR-FL |
| IGF1R | 0.69 | 0.53 | AR-FL |
| CTSD | 0.91 | 0.53 | AR-FL |

| | | | |
|---|---|---|---|
| PI4K2A | 0.69 | 0.53 | AR-FL |
| SLC35B2 | 1.09 | 0.53 | AR-FL |
| GPT2 | 0.82 | 0.53 | AR-FL |
| DRAP1 | 0.70 | 0.53 | AR-FL |
| TECR | 0.63 | 0.53 | AR-FL |
| PSMD8 | 1.04 | 0.53 | AR-FL |
| SHISA4 | 0.66 | 0.53 | AR-FL |
| COX8A | 0.83 | 0.54 | AR-FL |
| SYP | 0.82 | 0.54 | AR-FL |
| ZHX1 | 0.61 | 0.54 | AR-FL |
| TRIM11 | 1.18 | 0.54 | AR-FL |
| TXNDC17 | 1.14 | 0.54 | AR-FL |
| TMEM205 | 1.23 | 0.54 | AR-FL |
| FOS | 0.94 | 0.54 | AR-FL |
| CPSF2 | 0.88 | 0.54 | AR-FL |
| USMG5 | 0.82 | 0.54 | AR-FL |
| ZSCAN5A | 0.69 | 0.54 | AR-FL |
| HIST1H1C | 0.87 | 0.54 | AR-FL |
| DMWD | 0.83 | 0.54 | AR-FL |
| SLC9A1 | 0.81 | 0.54 | AR-FL |
| CHMP2A | 0.93 | 0.54 | AR-FL |
| MAPRE1 | 0.78 | 0.54 | AR-FL |
| MAPK11 | 0.82 | 0.54 | AR-FL |
| MRPS11 | 0.70 | 0.54 | AR-FL |
| CD63 | 1.17 | 0.54 | AR-FL |
| WHAMM | 0.63 | 0.54 | AR-FL |
| C17orf104 | 1.01 | 0.54 | AR-FL |
| ZNF513 | 0.77 | 0.54 | AR-FL |
| GPAT4 | 0.72 | 0.54 | AR-FL |
| PCCB | 0.73 | 0.55 | AR-FL |
| ATP5J2 | 0.87 | 0.55 | AR-FL |
| ABCA7 | 0.74 | 0.55 | AR-FL |
| KIAA1841 | 0.64 | 0.55 | AR-FL |
| RAB20 | 0.74 | 0.55 | AR-FL |
| ZNF697 | 1.79 | 0.55 | AR-FL |
| TAF6L | 0.63 | 0.55 | AR-FL |
| NTMT1 | 0.98 | 0.55 | AR-FL |
| DOT1L | 1.05 | 0.55 | AR-FL |
| RAB1B | 1.10 | 0.55 | AR-FL |
| KCNG3 | 1.37 | 0.55 | AR-FL |
| EPDR1 | 0.74 | 0.55 | AR-FL |
| PIP5K1A | 0.65 | 0.55 | AR-FL |
| SHFM1 | 0.75 | 0.55 | AR-FL |
| PTGS1 | 2.96 | 0.55 | AR-FL |
| GBA2 | 0.88 | 0.55 | AR-FL |
| ACSS2 | 0.60 | 0.55 | AR-FL |

| | | | |
|---|---|---|---|
| ECI2 | 0.77 | 0.55 | AR-FL |
| ALDH1A2 | 0.90 | 0.55 | AR-FL |
| ZCCHC24 | 1.16 | 0.55 | AR-FL |
| SNX11 | 0.89 | 0.55 | AR-FL |
| SC5D | 0.84 | 0.55 | AR-FL |
| RPS6KB2 | 0.94 | 0.55 | AR-FL |
| TMEM151A | 1.22 | 0.55 | AR-FL |
| TESK1 | 1.33 | 0.55 | AR-FL |
| NCLN | 0.73 | 0.55 | AR-FL |
| SLC25A53 | 0.72 | 0.55 | AR-FL |
| PPP6R2 | 1.20 | 0.55 | AR-FL |
| C9orf69 | 0.92 | 0.55 | AR-FL |
| ELMO3 | 0.68 | 0.55 | AR-FL |
| HYAL2 | 0.95 | 0.56 | AR-FL |
| HK1 | 1.18 | 0.56 | AR-FL |
| CASD1 | 0.92 | 0.56 | AR-FL |
| PCBP4 | 0.63 | 0.56 | AR-FL |
| MYEOV2 | 0.65 | 0.56 | AR-FL |
| SGOL1 | 0.66 | 0.56 | AR-FL |
| GHRHR | 1.13 | 0.56 | AR-FL |
| TEX2 | 0.82 | 0.56 | AR-FL |
| MTA2 | 1.01 | 0.56 | AR-FL |
| SLC25A1 | 0.73 | 0.56 | AR-FL |
| CANT1 | 1.55 | 0.56 | AR-FL |
| CYP4F12 | 1.82 | 0.56 | AR-FL |
| LSM4 | 0.65 | 0.56 | AR-FL |
| ATRNL1 | 0.80 | 0.56 | AR-FL |
| TALDO1 | 0.72 | 0.56 | AR-FL |
| SLC41A1 | 0.60 | 0.56 | AR-FL |
| IFT22 | 0.61 | 0.57 | AR-FL |
| IDH1 | 1.52 | 0.57 | AR-FL |
| LRRC16A | 0.78 | 0.57 | AR-FL |
| GTF3C6 | 0.97 | 0.57 | AR-FL |
| ISOC2 | 0.99 | 0.57 | AR-FL |
| ASL | 0.67 | 0.57 | AR-FL |
| RNF25 | 0.68 | 0.57 | AR-FL |
| EMC6 | 1.29 | 0.57 | AR-FL |
| CRADD | 0.65 | 0.57 | AR-FL |
| PSMB2 | 0.82 | 0.57 | AR-FL |
| LSM10 | 0.89 | 0.57 | AR-FL |
| PDXK | 0.88 | 0.57 | AR-FL |
| MESDC2 | 0.87 | 0.57 | AR-FL |
| ZNF668 | 0.97 | 0.57 | AR-FL |
| TSC22D3 | 0.64 | 0.57 | AR-FL |
| CABIN1 | 0.67 | 0.57 | AR-FL |
| CASKIN2 | 0.73 | 0.57 | AR-FL |

| | | | |
|---|---|---|---|
| RABAC1 | 1.14 | 0.57 | AR-FL |
| PRPF18 | 0.60 | 0.57 | AR-FL |
| UQCC2 | 0.61 | 0.57 | AR-FL |
| SLC30A2 | 1.90 | 0.57 | AR-FL |
| KIAA0513 | 0.85 | 0.57 | AR-FL |
| XKR8 | 0.68 | 0.57 | AR-FL |
| OSCP1 | 0.63 | 0.57 | AR-FL |
| AP2S1 | 1.10 | 0.57 | AR-FL |
| CTSV | 0.92 | 0.58 | AR-FL |
| ZNF341 | 0.83 | 0.58 | AR-FL |
| PSMB6 | 1.08 | 0.58 | AR-FL |
| CALML4 | 0.77 | 0.58 | AR-FL |
| CKAP4 | 0.74 | 0.58 | AR-FL |
| PHLDA2 | 1.27 | 0.58 | AR-FL |
| ATP6V0E2 | 1.36 | 0.58 | AR-FL |
| RNASEH1 | 0.63 | 0.58 | AR-FL |
| ATPIF1 | 0.70 | 0.58 | AR-FL |
| STIP1 | 0.91 | 0.58 | Both Isoforms |
| CISD3 | 0.65 | 0.58 | Both Isoforms |
| AGR3 | 2.93 | 0.58 | Both Isoforms |
| JUNB | 0.69 | 0.58 | Both Isoforms |
| ERLEC1 | 0.62 | 0.58 | Both Isoforms |
| CHRNA10 | 0.70 | 0.58 | Both Isoforms |
| RABL6 | 0.65 | 0.58 | Both Isoforms |
| TMED2 | 0.94 | 0.58 | Both Isoforms |
| ACTB | 0.69 | 0.58 | Both Isoforms |
| FBXO16 | 0.70 | 0.58 | Both Isoforms |
| KDELR2 | 1.38 | 0.58 | Both Isoforms |
| COX7B | 0.88 | 0.58 | Both Isoforms |
| ZC3HAV1 | 0.68 | 0.58 | Both Isoforms |
| C1orf210 | 0.70 | 0.58 | Both Isoforms |
| STX5 | 0.93 | 0.58 | Both Isoforms |
| PSMD14 | 1.04 | 0.59 | Both Isoforms |
| DPAGT1 | 1.19 | 0.59 | Both Isoforms |
| KIF1C | 1.30 | 0.59 | Both Isoforms |
| KLF4 | 1.78 | 0.59 | Both Isoforms |
| ZNF813 | 0.85 | 0.59 | Both Isoforms |
| KLHL18 | 0.76 | 0.59 | Both Isoforms |
| PSMC5 | 0.92 | 0.59 | Both Isoforms |
| PXN | 0.88 | 0.59 | Both Isoforms |
| RAP1A | 0.89 | 0.59 | Both Isoforms |
| TCEB2 | 0.76 | 0.59 | Both Isoforms |
| HMGCR | 1.07 | 0.59 | Both Isoforms |
| IKBKG | 0.90 | 0.59 | Both Isoforms |
| DIAPH1 | 0.63 | 0.59 | Both Isoforms |
| AC004381.6 | 0.64 | 0.59 | Both Isoforms |

| | | | | |
|---|---|---|---|---|
| | PQLC2 | 0.91 | 0.59 | Both Isoforms |
| | FLAD1 | 0.90 | 0.59 | Both Isoforms |
| | ALG8 | 1.16 | 0.59 | Both Isoforms |
| | NANS | 1.22 | 0.59 | Both Isoforms |
| | MICALL1 | 0.89 | 0.59 | Both Isoforms |
| | OSBPL7 | 0.96 | 0.59 | Both Isoforms |
| | MRPS26 | 0.66 | 0.59 | Both Isoforms |
| | PFN1 | 0.95 | 0.59 | Both Isoforms |
| | RALY | 0.80 | 0.59 | Both Isoforms |
| | LETM1 | 0.83 | 0.59 | Both Isoforms |
| | RNF181 | 0.66 | 0.59 | Both Isoforms |
| | KCNS3 | 0.65 | 0.59 | Both Isoforms |
| | UBA1 | 0.89 | 0.59 | Both Isoforms |
| | ABHD5 | 0.88 | 0.59 | Both Isoforms |
| | ANAPC11 | 0.79 | 0.59 | Both Isoforms |
| | PSMB3 | 0.94 | 0.59 | Both Isoforms |
| | MYL12A | 1.30 | 0.59 | Both Isoforms |
| | IFT27 | 0.67 | 0.59 | Both Isoforms |
| | NUP62CL | 0.83 | 0.59 | Both Isoforms |
| | WDR1 | 0.74 | 0.60 | Both Isoforms |
| | SWT1 | 1.22 | 0.60 | Both Isoforms |
| | MAP4K2 | 1.01 | 0.60 | Both Isoforms |
| | C17orf89 | 0.73 | 0.60 | Both Isoforms |
| | GALE | 0.69 | 0.60 | Both Isoforms |
| | ZNF467 | 2.12 | 0.60 | Both Isoforms |
| | ARHGDIA | 0.88 | 0.60 | Both Isoforms |
| | IDH3G | 1.06 | 0.60 | Both Isoforms |
| | SAPCD2 | 1.42 | 0.60 | Both Isoforms |
| | PLEKHB2 | 0.99 | 0.60 | Both Isoforms |
| | CHSY1 | 1.34 | 0.60 | Both Isoforms |
| | TUBA1C | 1.21 | 0.60 | Both Isoforms |
| | DNM2 | 0.68 | 0.60 | Both Isoforms |
| | BAHD1 | 0.59 | 0.60 | Both Isoforms |
| | WNT7B | 0.84 | 0.60 | Both Isoforms |
| | ALG14 | 0.87 | 0.60 | Both Isoforms |
| | CAMSAP3 | 0.89 | 0.60 | Both Isoforms |
| | NPLOC4 | 0.93 | 0.60 | Both Isoforms |
| | SLC25A10 | 0.73 | 0.60 | Both Isoforms |
| | FPGS | 0.62 | 0.60 | Both Isoforms |
| | KIF4B | 0.61 | 0.61 | Both Isoforms |
| | ABAT | 0.92 | 0.61 | Both Isoforms |
| | ZNF724P | 0.64 | 0.61 | Both Isoforms |
| | MTCH1 | 0.88 | 0.61 | Both Isoforms |
| | ZDHHC12 | 0.59 | 0.61 | Both Isoforms |
| | FUT1 | 0.70 | 0.61 | Both Isoforms |
| | FAAP24 | 0.79 | 0.61 | Both Isoforms |

| | | | | |
|---|---|---|---|---|
| | PPP2R3C | 0.86 | 0.61 | Both Isoforms |
| | KIF22 | 1.51 | 0.61 | Both Isoforms |
| | PCDHB5 | 1.00 | 0.61 | Both Isoforms |
| | ATP5G1 | 0.73 | 0.61 | Both Isoforms |
| | SLC25A13 | 1.02 | 0.61 | Both Isoforms |
| | UBE2M | 0.92 | 0.61 | Both Isoforms |
| | RAB1A | 1.06 | 0.61 | Both Isoforms |
| | ACADVL | 1.07 | 0.61 | Both Isoforms |
| | PSMC4 | 1.26 | 0.61 | Both Isoforms |
| | ADCY6 | 1.21 | 0.61 | Both Isoforms |
| | MPC2 | 1.73 | 0.61 | Both Isoforms |
| | TRIM3 | 1.07 | 0.61 | Both Isoforms |
| | COX14 | 0.68 | 0.61 | Both Isoforms |
| | HMG20B | 1.41 | 0.61 | Both Isoforms |
| | EDEM1 | 1.09 | 0.62 | Both Isoforms |
| | PSMD11 | 0.95 | 0.62 | Both Isoforms |
| | UGT2B28 | 2.97 | 0.62 | Both Isoforms |
| | ALDOA | 1.18 | 0.62 | Both Isoforms |
| | LMNA | 1.05 | 0.62 | Both Isoforms |
| | IWS1 | 0.81 | 0.62 | Both Isoforms |
| | CLPTM1 | 1.01 | 0.62 | Both Isoforms |
| | C17orf107 | 0.87 | 0.62 | Both Isoforms |
| | IMPDH1 | 1.22 | 0.62 | Both Isoforms |
| | TSEN15 | 0.66 | 0.62 | Both Isoforms |
| | EMC3 | 0.95 | 0.62 | Both Isoforms |
| | CIB1 | 1.12 | 0.62 | Both Isoforms |
| | IPPK | 1.66 | 0.62 | Both Isoforms |
| | MLX | 0.88 | 0.62 | Both Isoforms |
| | ALG12 | 0.90 | 0.62 | Both Isoforms |
| | TRAPPC10 | 0.75 | 0.62 | Both Isoforms |
| | TFRC | 1.09 | 0.63 | Both Isoforms |
| | APOO | 0.59 | 0.63 | Both Isoforms |
| | DOLPP1 | 1.03 | 0.63 | Both Isoforms |
| | ZFHX3 | 0.92 | 0.63 | Both Isoforms |
| | SEC61A2 | 0.74 | 0.63 | Both Isoforms |
| | TMEM141 | 1.22 | 0.63 | Both Isoforms |
| | PSMD2 | 1.02 | 0.63 | Both Isoforms |
| | EIF2B2 | 0.58 | 0.63 | Both Isoforms |
| | PNP | 0.59 | 0.63 | Both Isoforms |
| | ZNF726 | 0.60 | 0.63 | Both Isoforms |
| | RBX1 | 1.09 | 0.63 | Both Isoforms |
| | CAP2 | 1.75 | 0.63 | Both Isoforms |
| | DERL2 | 0.82 | 0.64 | Both Isoforms |
| | FZD8 | 1.86 | 0.64 | Both Isoforms |
| | SLC50A1 | 1.37 | 0.64 | Both Isoforms |
| | HNRNPAB | 0.90 | 0.64 | Both Isoforms |

| | | | |
|---|---|---|---|
| RAB4B | 1.07 | 0.64 | Both Isoforms |
| ZGPAT | 0.64 | 0.64 | Both Isoforms |
| C14orf1 | 0.86 | 0.64 | Both Isoforms |
| SAP130 | 0.82 | 0.64 | Both Isoforms |
| HID1 | 1.66 | 0.64 | Both Isoforms |
| GUK1 | 0.78 | 0.64 | Both Isoforms |
| PIGK | 0.93 | 0.64 | Both Isoforms |
| CPTP | 0.93 | 0.64 | Both Isoforms |
| CREB3L4 | 1.67 | 0.64 | Both Isoforms |
| SLC25A42 | 1.32 | 0.64 | Both Isoforms |
| LDLRAP1 | 1.04 | 0.65 | Both Isoforms |
| ZNF563 | 1.89 | 0.65 | Both Isoforms |
| ALDH4A1 | 1.09 | 0.65 | Both Isoforms |
| NDUFAF1 | 0.72 | 0.65 | Both Isoforms |
| MRPL36 | 1.01 | 0.65 | Both Isoforms |
| PCDHGC3 | 1.35 | 0.65 | Both Isoforms |
| MYH13 | 2.25 | 0.65 | Both Isoforms |
| PSMG3 | 0.60 | 0.65 | Both Isoforms |
| C16orf54 | 1.37 | 0.65 | Both Isoforms |
| SSR2 | 0.95 | 0.65 | Both Isoforms |
| FNDC3B | 1.09 | 0.65 | Both Isoforms |
| RXFP4 | 1.26 | 0.65 | Both Isoforms |
| UQCR10 | 0.81 | 0.65 | Both Isoforms |
| HIST1H2BJ | 0.81 | 0.65 | Both Isoforms |
| PDE12 | 0.95 | 0.65 | Both Isoforms |
| GPATCH4 | 0.70 | 0.65 | Both Isoforms |
| TMED7 | 1.14 | 0.65 | Both Isoforms |
| SEMA3G | 2.44 | 0.66 | Both Isoforms |
| SLC27A4 | 1.35 | 0.66 | Both Isoforms |
| CARD10 | 1.15 | 0.66 | Both Isoforms |
| PIK3C2A | 0.73 | 0.66 | Both Isoforms |
| NACC2 | 1.26 | 0.66 | Both Isoforms |
| LSS | 0.74 | 0.66 | Both Isoforms |
| TTC38 | 0.63 | 0.66 | Both Isoforms |
| HSD17B10 | 0.73 | 0.66 | Both Isoforms |
| SLC31A1 | 0.93 | 0.66 | Both Isoforms |
| CHKA | 0.81 | 0.66 | Both Isoforms |
| SNX19 | 0.92 | 0.66 | Both Isoforms |
| SLC33A1 | 0.75 | 0.66 | Both Isoforms |
| SRM | 0.67 | 0.66 | Both Isoforms |
| UBE2S | 1.14 | 0.66 | Both Isoforms |
| TMEM150C | 1.19 | 0.66 | Both Isoforms |
| VPS26B | 2.53 | 0.66 | Both Isoforms |
| PYM1 | 0.99 | 0.66 | Both Isoforms |
| SDSL | 1.09 | 0.66 | Both Isoforms |
| RAB5C | 0.99 | 0.66 | Both Isoforms |

| | | | |
|---|---|---|---|
| LGMN | 1.50 | 0.66 | Both Isoforms |
| IDH3A | 1.18 | 0.66 | Both Isoforms |
| GOLGA3 | 1.14 | 0.66 | Both Isoforms |
| ENG | 1.34 | 0.66 | Both Isoforms |
| GDI1 | 0.80 | 0.66 | Both Isoforms |
| TMCC3 | 0.63 | 0.66 | Both Isoforms |
| SHKBP1 | 1.15 | 0.66 | Both Isoforms |
| CABLES2 | 0.60 | 0.66 | Both Isoforms |
| BRMS1 | 0.97 | 0.66 | Both Isoforms |
| TOR3A | 0.68 | 0.66 | Both Isoforms |
| TMA7 | 0.64 | 0.66 | Both Isoforms |
| ZNF808 | 0.60 | 0.67 | Both Isoforms |
| IQCE | 0.79 | 0.67 | Both Isoforms |
| DLX2 | 1.48 | 0.67 | Both Isoforms |
| SCAMP3 | 1.02 | 0.67 | Both Isoforms |
| IFI27L1 | 0.78 | 0.67 | Both Isoforms |
| MPZL2 | 0.88 | 0.67 | Both Isoforms |
| MRPL41 | 0.92 | 0.67 | Both Isoforms |
| CNN3 | 0.63 | 0.67 | Both Isoforms |
| PRKACA | 0.73 | 0.67 | Both Isoforms |
| HERPUD1 | 0.95 | 0.67 | Both Isoforms |
| PSMC3 | 0.86 | 0.67 | Both Isoforms |
| TBRG1 | 0.59 | 0.67 | Both Isoforms |
| GREB1 | 1.03 | 0.67 | Both Isoforms |
| ZNF101 | 0.81 | 0.67 | Both Isoforms |
| SLC25A11 | 0.92 | 0.67 | Both Isoforms |
| SF3B5 | 0.90 | 0.67 | Both Isoforms |
| MEA1 | 0.76 | 0.68 | Both Isoforms |
| SEC24A | 1.41 | 0.68 | Both Isoforms |
| SLC35A2 | 0.79 | 0.68 | Both Isoforms |
| MFSD3 | 1.21 | 0.68 | Both Isoforms |
| RGMB | 0.69 | 0.68 | Both Isoforms |
| AP5B1 | 0.60 | 0.68 | Both Isoforms |
| WDYHV1 | 0.61 | 0.68 | Both Isoforms |
| PPFIA4 | 1.20 | 0.68 | Both Isoforms |
| DDA1 | 0.77 | 0.68 | Both Isoforms |
| ACAD8 | 1.38 | 0.68 | Both Isoforms |
| DPP9 | 0.67 | 0.68 | Both Isoforms |
| SQLE | 0.95 | 0.68 | Both Isoforms |
| DEF8 | 0.61 | 0.69 | Both Isoforms |
| TRPM2 | 1.25 | 0.69 | Both Isoforms |
| PCSK7 | 1.15 | 0.69 | Both Isoforms |
| FAM129B | 0.70 | 0.69 | Both Isoforms |
| SEC24C | 1.19 | 0.69 | Both Isoforms |
| PTS | 0.94 | 0.69 | Both Isoforms |
| DNAL1 | 0.65 | 0.69 | Both Isoforms |

| | | | |
|---|---|---|---|
| SCD | 1.82 | 0.69 | Both Isoforms |
| YKT6 | 1.03 | 0.69 | Both Isoforms |
| TMEM125 | 1.60 | 0.69 | Both Isoforms |
| RBM14 | 0.64 | 0.69 | Both Isoforms |
| ATP10A | 1.81 | 0.69 | Both Isoforms |
| SSR4 | 0.90 | 0.69 | Both Isoforms |
| SLC25A35 | 2.56 | 0.69 | Both Isoforms |
| DDX5 | 0.89 | 0.69 | Both Isoforms |
| PXMP4 | 0.78 | 0.69 | Both Isoforms |
| MGST2 | 1.11 | 0.69 | Both Isoforms |
| SPHK2 | 0.81 | 0.69 | Both Isoforms |
| CALU | 1.36 | 0.70 | Both Isoforms |
| PRDX4 | 0.81 | 0.70 | Both Isoforms |
| SERTAD3 | 1.22 | 0.70 | Both Isoforms |
| RYR3 | 0.77 | 0.70 | Both Isoforms |
| KLC2 | 1.34 | 0.70 | Both Isoforms |
| CTPS1 | 0.69 | 0.70 | Both Isoforms |
| OSTM1 | 1.48 | 0.70 | Both Isoforms |
| CCDC173 | 3.09 | 0.70 | Both Isoforms |
| TSPAN1 | 1.29 | 0.70 | Both Isoforms |
| APLN | 1.09 | 0.70 | Both Isoforms |
| CCDC64 | 1.20 | 0.70 | Both Isoforms |
| ICMT | 0.97 | 0.70 | Both Isoforms |
| VCP | 1.20 | 0.70 | Both Isoforms |
| MAGED1 | 1.48 | 0.70 | Both Isoforms |
| CYB561 | 1.05 | 0.70 | Both Isoforms |
| FAM213A | 0.62 | 0.70 | Both Isoforms |
| CACNG4 | 1.91 | 0.70 | Both Isoforms |
| CLPP | 0.66 | 0.70 | Both Isoforms |
| GNPNAT1 | 0.79 | 0.70 | Both Isoforms |
| PSMA7 | 1.09 | 0.70 | Both Isoforms |
| SLC9A3R1 | 1.54 | 0.70 | Both Isoforms |
| TIMM23B | 0.61 | 0.70 | Both Isoforms |
| TOB1 | 0.81 | 0.70 | Both Isoforms |
| CYB561D2 | 1.24 | 0.70 | Both Isoforms |
| GLA | 0.71 | 0.70 | Both Isoforms |
| SLX4 | 0.71 | 0.71 | Both Isoforms |
| SPCS2 | 1.03 | 0.71 | Both Isoforms |
| CCDC96 | 0.95 | 0.71 | Both Isoforms |
| ATP2A2 | 1.01 | 0.71 | Both Isoforms |
| TMEM50B | 1.14 | 0.71 | Both Isoforms |
| TMEM120A | 0.63 | 0.71 | Both Isoforms |
| MTOR | 1.44 | 0.71 | Both Isoforms |
| CD151 | 1.24 | 0.71 | Both Isoforms |
| NDOR1 | 0.60 | 0.71 | Both Isoforms |
| FAM64A | 0.64 | 0.71 | Both Isoforms |

| | | | | |
|---|---|---|---|---|
| | NDUFAF3 | 0.84 | 0.71 | Both Isoforms |
| | ACVR1 | 1.04 | 0.71 | Both Isoforms |
| | CFL1 | 1.25 | 0.71 | Both Isoforms |
| | NME4 | 1.21 | 0.72 | Both Isoforms |
| | ARFIP2 | 1.19 | 0.72 | Both Isoforms |
| | NDUFS7 | 0.84 | 0.72 | Both Isoforms |
| | ATP5I | 0.94 | 0.72 | Both Isoforms |
| | DAG1 | 1.02 | 0.72 | Both Isoforms |
| | AKIRIN1 | 1.16 | 0.72 | Both Isoforms |
| | NACC1 | 1.10 | 0.72 | Both Isoforms |
| | NXN | 0.92 | 0.72 | Both Isoforms |
| | TMED10 | 1.03 | 0.72 | Both Isoforms |
| | NDUFA3 | 0.82 | 0.72 | Both Isoforms |
| | BORCS8 | 0.98 | 0.72 | Both Isoforms |
| | THSD7A | 2.57 | 0.72 | Both Isoforms |
| | ADRM1 | 1.24 | 0.72 | Both Isoforms |
| | SYNGR2 | 1.61 | 0.73 | Both Isoforms |
| | LIG4 | 1.36 | 0.73 | Both Isoforms |
| | EMC8 | 0.83 | 0.73 | Both Isoforms |
| | PIR | 1.28 | 0.73 | Both Isoforms |
| | FAM110A | 1.10 | 0.73 | Both Isoforms |
| | UQCRQ | 1.24 | 0.73 | Both Isoforms |
| | LUZP1 | 0.93 | 0.73 | Both Isoforms |
| | MRPL33 | 0.69 | 0.73 | Both Isoforms |
| | ARPC5L | 1.08 | 0.73 | Both Isoforms |
| | SLC41A2 | 0.94 | 0.73 | Both Isoforms |
| | LINC00116 | 0.92 | 0.73 | Both Isoforms |
| | SEC23B | 1.06 | 0.73 | Both Isoforms |
| | AP2A1 | 0.66 | 0.73 | Both Isoforms |
| | MICALL2 | 1.19 | 0.73 | Both Isoforms |
| | NFKBIA | 0.64 | 0.74 | Both Isoforms |
| | TCF7 | 1.20 | 0.74 | Both Isoforms |
| | APIP | 0.80 | 0.74 | Both Isoforms |
| | TBC1D10A | 0.79 | 0.74 | Both Isoforms |
| | SLC36A1 | 1.07 | 0.74 | Both Isoforms |
| | ST14 | 0.76 | 0.74 | Both Isoforms |
| | PPP1CB | 1.06 | 0.74 | Both Isoforms |
| | TRIM37 | 1.35 | 0.74 | Both Isoforms |
| | ELMO2 | 1.01 | 0.74 | Both Isoforms |
| | PLPPR2 | 1.05 | 0.74 | Both Isoforms |
| | PRR4 | 0.61 | 0.74 | Both Isoforms |
| | ACLY | 1.49 | 0.75 | Both Isoforms |
| | GPRC5B | 2.67 | 0.75 | Both Isoforms |
| | PFKP | 0.71 | 0.75 | Both Isoforms |
| | DEFB132 | 1.76 | 0.75 | Both Isoforms |
| | PTGES3L | 1.28 | 0.75 | Both Isoforms |

| | | | |
|---|---|---|---|
| CHFR | 1.31 | 0.75 | Both Isoforms |
| SRA1 | 1.57 | 0.75 | Both Isoforms |
| C19orf48 | 0.68 | 0.75 | Both Isoforms |
| OCEL1 | 1.76 | 0.75 | Both Isoforms |
| ISG20 | 0.90 | 0.76 | Both Isoforms |
| KDELR3 | 2.43 | 0.76 | Both Isoforms |
| KL | 0.65 | 0.76 | Both Isoforms |
| MIEN1 | 1.40 | 0.76 | Both Isoforms |
| SIDT2 | 0.69 | 0.76 | Both Isoforms |
| ZNF701 | 0.75 | 0.76 | Both Isoforms |
| SLC35E4 | 0.78 | 0.76 | Both Isoforms |
| SYNJ1 | 0.68 | 0.76 | Both Isoforms |
| KLHDC7A | 1.48 | 0.76 | Both Isoforms |
| GPAA1 | 0.64 | 0.76 | Both Isoforms |
| MGAT2 | 0.64 | 0.77 | Both Isoforms |
| SHC1 | 0.91 | 0.77 | Both Isoforms |
| MYO9B | 0.92 | 0.77 | Both Isoforms |
| FAM81A | 1.12 | 0.77 | Both Isoforms |
| GALNT16 | 1.01 | 0.77 | Both Isoforms |
| PCNXL3 | 0.77 | 0.77 | Both Isoforms |
| VMP1 | 1.31 | 0.77 | Both Isoforms |
| ARF4 | 1.49 | 0.77 | Both Isoforms |
| ATP6V0D1 | 1.20 | 0.77 | Both Isoforms |
| MAP2K6 | 0.72 | 0.77 | Both Isoforms |
| C6orf226 | 1.41 | 0.77 | Both Isoforms |
| SWI5 | 1.04 | 0.78 | Both Isoforms |
| CCDC15 | 0.68 | 0.78 | Both Isoforms |
| LMAN2 | 0.95 | 0.78 | Both Isoforms |
| MPDU1 | 1.27 | 0.78 | Both Isoforms |
| SYVN1 | 1.10 | 0.78 | Both Isoforms |
| RAPH1 | 1.14 | 0.78 | Both Isoforms |
| MELK | 0.60 | 0.78 | Both Isoforms |
| ZNF718 | 0.71 | 0.78 | Both Isoforms |
| AP1S3 | 1.11 | 0.78 | Both Isoforms |
| ETV2 | 0.99 | 0.78 | Both Isoforms |
| CSRP1 | 1.39 | 0.78 | Both Isoforms |
| CRYAB | 1.99 | 0.78 | Both Isoforms |
| TANGO2 | 0.85 | 0.78 | Both Isoforms |
| LRRC26 | 2.42 | 0.79 | Both Isoforms |
| PLOD3 | 0.90 | 0.79 | Both Isoforms |
| ST3GAL1 | 0.79 | 0.79 | Both Isoforms |
| ALYREF | 0.88 | 0.79 | Both Isoforms |
| AGFG2 | 0.89 | 0.79 | Both Isoforms |
| OPN1SW | 1.51 | 0.79 | Both Isoforms |
| PPARGC1B | 0.70 | 0.79 | Both Isoforms |
| LTBR | 0.77 | 0.79 | Both Isoforms |

| | | | |
|---|---|---|---|
| C18orf8 | 1.33 | 0.79 | Both Isoforms |
| C4orf48 | 0.84 | 0.79 | Both Isoforms |
| CKS1B | 0.63 | 0.79 | Both Isoforms |
| YIF1A | 1.18 | 0.80 | Both Isoforms |
| INTS6 | 0.84 | 0.80 | Both Isoforms |
| PIP4K2A | 1.14 | 0.80 | Both Isoforms |
| PLXNB3 | 0.83 | 0.80 | Both Isoforms |
| MPPE1 | 0.67 | 0.80 | Both Isoforms |
| PLPP1 | 1.35 | 0.80 | Both Isoforms |
| SLC35C1 | 1.49 | 0.80 | Both Isoforms |
| CTTN | 0.89 | 0.80 | Both Isoforms |
| DPH3 | 0.98 | 0.80 | Both Isoforms |
| TRPT1 | 1.13 | 0.80 | Both Isoforms |
| HM13 | 1.16 | 0.80 | Both Isoforms |
| SAR1B | 1.16 | 0.80 | Both Isoforms |
| WWC1 | 0.96 | 0.80 | Both Isoforms |
| LMNB2 | 0.68 | 0.81 | Both Isoforms |
| SEC16A | 1.40 | 0.81 | Both Isoforms |
| PSMD1 | 1.11 | 0.81 | Both Isoforms |
| GPS1 | 0.82 | 0.81 | Both Isoforms |
| CLTB | 0.97 | 0.81 | Both Isoforms |
| CYCS | 1.10 | 0.81 | Both Isoforms |
| SMTN | 1.57 | 0.81 | Both Isoforms |
| PYCRL | 0.59 | 0.81 | Both Isoforms |
| TMEM55B | 0.86 | 0.81 | Both Isoforms |
| EPHB2 | 0.87 | 0.81 | Both Isoforms |
| ST20 | 1.10 | 0.81 | Both Isoforms |
| CYP4F11 | 0.81 | 0.82 | Both Isoforms |
| NDUFS8 | 1.02 | 0.82 | Both Isoforms |
| PDX1 | 1.44 | 0.82 | Both Isoforms |
| ZNF682 | 0.88 | 0.82 | Both Isoforms |
| ADAMTS7 | 2.79 | 0.82 | Both Isoforms |
| TSPAN13 | 1.91 | 0.82 | Both Isoforms |
| ENTPD2 | 1.90 | 0.82 | Both Isoforms |
| ANGPTL3 | 1.35 | 0.82 | Both Isoforms |
| PPP2R5B | 0.77 | 0.82 | Both Isoforms |
| TUBA1B | 0.80 | 0.82 | Both Isoforms |
| CDR2 | 1.01 | 0.83 | Both Isoforms |
| CBWD5 | 0.71 | 0.83 | Both Isoforms |
| YIF1B | 1.41 | 0.83 | Both Isoforms |
| ZC3H12A | 1.28 | 0.83 | Both Isoforms |
| C15orf65 | 1.13 | 0.83 | Both Isoforms |
| PFKFB4 | 1.36 | 0.83 | Both Isoforms |
| ZFP36 | 1.53 | 0.83 | Both Isoforms |
| GDF11 | 1.46 | 0.83 | Both Isoforms |
| DAD1 | 1.16 | 0.83 | Both Isoforms |

| | | | |
|---|---|---|---|
| ANO4 | 0.78 | 0.83 | Both Isoforms |
| C14orf142 | 1.10 | 0.84 | Both Isoforms |
| GK | 0.74 | 0.84 | Both Isoforms |
| PRKCD | 0.99 | 0.84 | Both Isoforms |
| FAM73B | 1.06 | 0.84 | Both Isoforms |
| KIAA1919 | 1.07 | 0.84 | Both Isoforms |
| BIN3 | 0.81 | 0.84 | Both Isoforms |
| C2CD2L | 1.29 | 0.84 | Both Isoforms |
| ITGAE | 0.88 | 0.84 | Both Isoforms |
| COPG1 | 1.36 | 0.85 | Both Isoforms |
| CAMK2N2 | 1.33 | 0.85 | Both Isoforms |
| MORF4L2 | 0.87 | 0.85 | Both Isoforms |
| LRCH4 | 1.06 | 0.85 | Both Isoforms |
| SLC25A22 | 0.63 | 0.85 | Both Isoforms |
| PCDHB7 | 1.04 | 0.85 | Both Isoforms |
| ZCCHC6 | 1.09 | 0.85 | Both Isoforms |
| SLC25A44 | 1.04 | 0.85 | Both Isoforms |
| B4GALNT1 | 1.72 | 0.85 | Both Isoforms |
| GCAT | 0.73 | 0.85 | Both Isoforms |
| RRP36 | 1.28 | 0.85 | Both Isoforms |
| TM9SF1 | 1.28 | 0.86 | Both Isoforms |
| MED31 | 0.85 | 0.86 | Both Isoforms |
| SPAG4 | 1.30 | 0.86 | Both Isoforms |
| SH3TC2 | 0.61 | 0.86 | Both Isoforms |
| GPR89B | 0.81 | 0.86 | Both Isoforms |
| MSTO1 | 0.82 | 0.86 | Both Isoforms |
| EDEM3 | 0.79 | 0.86 | Both Isoforms |
| MAPRE3 | 1.22 | 0.86 | Both Isoforms |
| RHBDF1 | 1.15 | 0.86 | Both Isoforms |
| PLCXD2 | 1.52 | 0.86 | Both Isoforms |
| PLEKHG2 | 1.89 | 0.87 | Both Isoforms |
| PIP4K2C | 1.13 | 0.87 | Both Isoforms |
| EPS8L2 | 0.77 | 0.87 | Both Isoforms |
| C7orf43 | 1.12 | 0.87 | Both Isoforms |
| PVRL1 | 1.68 | 0.88 | Both Isoforms |
| AEN | 1.08 | 0.88 | Both Isoforms |
| RHBDD3 | 1.09 | 0.88 | Both Isoforms |
| TTC12 | 1.48 | 0.88 | Both Isoforms |
| PSENEN | 0.98 | 0.88 | Both Isoforms |
| TCEAL3 | 1.85 | 0.88 | Both Isoforms |
| FAM214B | 1.50 | 0.88 | Both Isoforms |
| SLC38A10 | 0.79 | 0.88 | Both Isoforms |
| NOL12 | 0.69 | 0.89 | Both Isoforms |
| TXNDC11 | 1.39 | 0.89 | Both Isoforms |
| BBIP1 | 0.90 | 0.89 | Both Isoforms |
| CHEK1 | 0.61 | 0.89 | Both Isoforms |

| | | | | |
|---|---|---|---|---|
| | IDE | 1.48 | 0.89 | Both Isoforms |
| | EVI5L | 1.21 | 0.90 | Both Isoforms |
| | AP4E1 | 1.05 | 0.90 | Both Isoforms |
| | CYB5A | 0.68 | 0.90 | Both Isoforms |
| | C17orf62 | 1.93 | 0.90 | Both Isoforms |
| | SLC2A5 | 3.85 | 0.90 | Both Isoforms |
| | AUNIP | 0.61 | 0.90 | Both Isoforms |
| | ZNF761 | 1.47 | 0.90 | Both Isoforms |
| | SGK1 | 2.17 | 0.90 | Both Isoforms |
| | MTFR2 | 0.72 | 0.90 | Both Isoforms |
| | ABCC1 | 0.96 | 0.90 | Both Isoforms |
| | FAM43A | 0.91 | 0.90 | Both Isoforms |
| | SELK | 1.11 | 0.90 | Both Isoforms |
| | P4HA2 | 1.98 | 0.90 | Both Isoforms |
| | MFSD5 | 1.42 | 0.91 | Both Isoforms |
| | MAP2K4 | 0.77 | 0.91 | Both Isoforms |
| | POLR3K | 1.02 | 0.91 | Both Isoforms |
| | GPC2 | 2.48 | 0.91 | Both Isoforms |
| | FDPS | 1.42 | 0.91 | Both Isoforms |
| | C19orf73 | 0.87 | 0.91 | Both Isoforms |
| | PGAM1 | 1.17 | 0.91 | Both Isoforms |
| | GSG2 | 0.65 | 0.91 | Both Isoforms |
| | RTN4IP1 | 0.67 | 0.91 | Both Isoforms |
| | TAOK2 | 0.95 | 0.91 | Both Isoforms |
| | CHPF | 0.86 | 0.91 | Both Isoforms |
| | CCDC87 | 1.45 | 0.92 | Both Isoforms |
| | SEC13 | 1.20 | 0.92 | Both Isoforms |
| | VIMP | 1.26 | 0.92 | Both Isoforms |
| | RASSF7 | 1.16 | 0.92 | Both Isoforms |
| | MON2 | 0.61 | 0.92 | Both Isoforms |
| | B3GAT3 | 1.06 | 0.92 | Both Isoforms |
| | CDYL2 | 1.43 | 0.92 | Both Isoforms |
| | YIPF1 | 2.19 | 0.92 | Both Isoforms |
| | RCAN1 | 0.97 | 0.92 | Both Isoforms |
| | SRPRB | 0.80 | 0.92 | Both Isoforms |
| | DAP | 0.82 | 0.92 | Both Isoforms |
| | SLC43A1 | 1.01 | 0.92 | Both Isoforms |
| | SUOX | 1.19 | 0.93 | Both Isoforms |
| | RFX6 | 0.70 | 0.93 | Both Isoforms |
| | HSD17B7 | 1.03 | 0.93 | Both Isoforms |
| | PDXP | 1.06 | 0.93 | Both Isoforms |
| | SLC35F6 | 1.30 | 0.93 | Both Isoforms |
| | GBF1 | 1.16 | 0.93 | Both Isoforms |
| | EPS8L1 | 1.94 | 0.93 | Both Isoforms |
| | TAF13 | 1.24 | 0.93 | Both Isoforms |
| | OCRL | 0.76 | 0.93 | Both Isoforms |

| | | | |
|---|---|---|---|
| JMJD6 | 1.08 | 0.93 | Both Isoforms |
| SH3D21 | 1.16 | 0.93 | Both Isoforms |
| FDFT1 | 1.33 | 0.93 | Both Isoforms |
| CDK2 | 1.44 | 0.93 | Both Isoforms |
| ENTPD7 | 1.64 | 0.94 | Both Isoforms |
| CMTM3 | 1.19 | 0.94 | Both Isoforms |
| ATP6V0B | 1.52 | 0.94 | Both Isoforms |
| CCDC107 | 1.00 | 0.94 | Both Isoforms |
| TPI1 | 0.64 | 0.94 | Both Isoforms |
| GOT1 | 1.05 | 0.94 | Both Isoforms |
| FZD1 | 1.37 | 0.94 | Both Isoforms |
| HIST1H2AI | 0.82 | 0.95 | Both Isoforms |
| CENPO | 0.94 | 0.95 | Both Isoforms |
| TUBA4A | 1.22 | 0.95 | Both Isoforms |
| ARHGEF18 | 0.92 | 0.95 | Both Isoforms |
| MAGT1 | 1.16 | 0.95 | Both Isoforms |
| FKBP2 | 1.35 | 0.95 | Both Isoforms |
| KCNN3 | 1.26 | 0.95 | Both Isoforms |
| GORASP1 | 1.23 | 0.95 | Both Isoforms |
| CYP51A1 | 0.89 | 0.95 | Both Isoforms |
| FECH | 0.66 | 0.96 | Both Isoforms |
| NDUFS6 | 1.17 | 0.96 | Both Isoforms |
| CXCL8 | 1.69 | 0.96 | Both Isoforms |
| ORAI2 | 1.08 | 0.96 | Both Isoforms |
| KLK4 | 1.85 | 0.96 | Both Isoforms |
| MCM6 | 0.78 | 0.96 | Both Isoforms |
| ATP6V0E1 | 1.07 | 0.97 | Both Isoforms |
| LRRC45 | 1.62 | 0.97 | Both Isoforms |
| MSRA | 1.34 | 0.97 | Both Isoforms |
| STRA13 | 1.73 | 0.97 | Both Isoforms |
| SLC2A6 | 1.35 | 0.97 | Both Isoforms |
| CFAP53 | 0.67 | 0.97 | Both Isoforms |
| DDT | 1.53 | 0.97 | Both Isoforms |
| PRDM16 | 1.80 | 0.97 | Both Isoforms |
| MYDGF | 1.51 | 0.97 | Both Isoforms |
| C2orf76 | 1.37 | 0.97 | Both Isoforms |
| FITM2 | 1.22 | 0.97 | Both Isoforms |
| MCM8 | 0.89 | 0.98 | Both Isoforms |
| USP43 | 0.59 | 0.98 | Both Isoforms |
| USB1 | 1.59 | 0.98 | Both Isoforms |
| PHTF1 | 0.99 | 0.98 | Both Isoforms |
| CHP1 | 1.48 | 0.98 | Both Isoforms |
| TMED9 | 1.58 | 0.98 | Both Isoforms |
| FANCB | 0.91 | 0.98 | Both Isoforms |
| RPN2 | 1.31 | 0.98 | Both Isoforms |
| PLIN2 | 0.84 | 0.98 | Both Isoforms |

| | | | |
|---|---|---|---|
| MIDN | 1.15 | 0.98 | Both Isoforms |
| KRT10 | 1.22 | 0.98 | Both Isoforms |
| CENPM | 0.73 | 0.98 | Both Isoforms |
| SNCG | 1.70 | 0.98 | Both Isoforms |
| PDIA6 | 1.35 | 0.98 | Both Isoforms |
| STT3A | 1.36 | 0.99 | Both Isoforms |
| ARL13B | 1.29 | 0.99 | Both Isoforms |
| WDR34 | 0.71 | 0.99 | Both Isoforms |
| GNAO1 | 1.06 | 0.99 | Both Isoforms |
| SSR1 | 1.24 | 0.99 | Both Isoforms |
| PHKG2 | 1.00 | 0.99 | Both Isoforms |
| FAM151B | 1.00 | 0.99 | Both Isoforms |
| DUS1L | 0.80 | 0.99 | Both Isoforms |
| P2RY2 | 1.68 | 0.99 | Both Isoforms |
| LDLRAD3 | 0.92 | 0.99 | Both Isoforms |
| ABHD11 | 0.97 | 0.99 | Both Isoforms |
| ZFAND2B | 1.07 | 0.99 | Both Isoforms |
| SHROOM3 | 1.03 | 1.00 | Both Isoforms |
| DNAJB9 | 1.28 | 1.00 | Both Isoforms |
| TRAPPC2P1 | 1.15 | 1.00 | Both Isoforms |
| LIN52 | 0.84 | 1.00 | Both Isoforms |
| DNMT1 | 0.77 | 1.00 | Both Isoforms |
| PVRL2 | 1.89 | 1.00 | Both Isoforms |
| CST1 | 3.06 | 1.00 | Both Isoforms |
| ULBP2 | 1.06 | 1.00 | Both Isoforms |
| CNKSR2 | 1.32 | 1.00 | Both Isoforms |
| ERCC6L | 0.97 | 1.00 | Both Isoforms |
| PIAS1 | 0.81 | 1.00 | Both Isoforms |
| TMEM53 | 1.02 | 1.01 | Both Isoforms |
| PLEKHO2 | 2.07 | 1.01 | Both Isoforms |
| DHRS2 | 1.59 | 1.01 | Both Isoforms |
| UFD1L | 1.23 | 1.01 | Both Isoforms |
| IKZF1 | 2.77 | 1.01 | Both Isoforms |
| MCM3 | 0.71 | 1.01 | Both Isoforms |
| C20orf24 | 1.44 | 1.01 | Both Isoforms |
| RTEL1 | 0.74 | 1.01 | Both Isoforms |
| PUSL1 | 1.15 | 1.01 | Both Isoforms |
| TIMELESS | 0.68 | 1.01 | Both Isoforms |
| XBP1 | 0.98 | 1.02 | Both Isoforms |
| HOXA11 | 0.99 | 1.02 | Both Isoforms |
| METRNL | 1.39 | 1.02 | Both Isoforms |
| KRT18 | 1.52 | 1.02 | Both Isoforms |
| SAFB2 | 0.81 | 1.02 | Both Isoforms |
| COPE | 1.47 | 1.02 | Both Isoforms |
| UGGT1 | 1.15 | 1.02 | Both Isoforms |
| PDIA3 | 1.51 | 1.03 | Both Isoforms |

| | | | |
|---|---|---|---|
| GALNS | 0.97 | 1.03 | Both Isoforms |
| SLC25A20 | 1.90 | 1.03 | Both Isoforms |
| DOLK | 1.62 | 1.03 | Both Isoforms |
| EIF2AK3 | 1.10 | 1.04 | Both Isoforms |
| CTB-55O6.8 | 2.12 | 1.04 | Both Isoforms |
| TMEM214 | 1.48 | 1.04 | Both Isoforms |
| NUDT16L1 | 1.27 | 1.04 | Both Isoforms |
| PNPLA8 | 0.66 | 1.04 | Both Isoforms |
| SLC52A2 | 1.80 | 1.04 | Both Isoforms |
| KIAA0232 | 1.03 | 1.04 | Both Isoforms |
| GLRX | 1.11 | 1.04 | Both Isoforms |
| H2AFZ | 0.67 | 1.05 | Both Isoforms |
| UBE2T | 0.89 | 1.05 | Both Isoforms |
| CTXN1 | 1.41 | 1.05 | Both Isoforms |
| MBOAT2 | 1.36 | 1.05 | Both Isoforms |
| C19orf57 | 0.58 | 1.06 | Both Isoforms |
| LYZ | 1.79 | 1.06 | Both Isoforms |
| CHAF1B | 0.66 | 1.06 | Both Isoforms |
| AK1 | 1.57 | 1.06 | Both Isoforms |
| PRSS36 | 0.70 | 1.06 | Both Isoforms |
| CCDC141 | 3.00 | 1.07 | Both Isoforms |
| PACSIN2 | 1.08 | 1.07 | Both Isoforms |
| ORMDL2 | 1.60 | 1.07 | Both Isoforms |
| A2M | 1.08 | 1.07 | Both Isoforms |
| AQP3 | 0.69 | 1.07 | Both Isoforms |
| ACSM1 | 2.63 | 1.07 | Both Isoforms |
| XRCC3 | 0.64 | 1.07 | Both Isoforms |
| POLR2L | 1.34 | 1.07 | Both Isoforms |
| DNAJC10 | 2.10 | 1.07 | Both Isoforms |
| CHPF2 | 1.35 | 1.07 | Both Isoforms |
| JAG1 | 1.15 | 1.07 | Both Isoforms |
| ORC1 | 0.76 | 1.07 | Both Isoforms |
| CHRNA5 | 0.80 | 1.07 | Both Isoforms |
| DLX1 | 1.26 | 1.08 | Both Isoforms |
| MORN2 | 0.92 | 1.08 | Both Isoforms |
| CALR | 1.41 | 1.08 | Both Isoforms |
| EHD4 | 1.56 | 1.08 | Both Isoforms |
| SERTAD1 | 1.91 | 1.08 | Both Isoforms |
| CDKN2D | 0.94 | 1.09 | Both Isoforms |
| MERTK | 1.22 | 1.09 | Both Isoforms |
| BCAR1 | 1.55 | 1.09 | Both Isoforms |
| ROMO1 | 1.26 | 1.09 | Both Isoforms |
| POP5 | 0.80 | 1.10 | Both Isoforms |
| RHAG | 5.72 | 1.10 | Both Isoforms |
| GPS2 | 1.13 | 1.10 | Both Isoforms |
| GSTT2B | 1.25 | 1.10 | Both Isoforms |

| | | | |
|---|---|---|---|
| SRP19 | 1.09 | 1.10 | Both Isoforms |
| SURF4 | 1.64 | 1.10 | Both Isoforms |
| B3GNT2 | 2.23 | 1.11 | Both Isoforms |
| UNC13B | 0.79 | 1.11 | Both Isoforms |
| RHBDD2 | 1.31 | 1.11 | Both Isoforms |
| SRPR | 1.39 | 1.11 | Both Isoforms |
| BEAN1 | 1.73 | 1.11 | Both Isoforms |
| MASTL | 0.78 | 1.11 | Both Isoforms |
| NUDT22 | 1.46 | 1.11 | Both Isoforms |
| ENDOG | 0.88 | 1.11 | Both Isoforms |
| GINS1 | 0.65 | 1.11 | Both Isoforms |
| PPIB | 1.37 | 1.11 | Both Isoforms |
| ANKRD54 | 0.62 | 1.11 | Both Isoforms |
| GNRH2 | 1.45 | 1.13 | Both Isoforms |
| SLC39A3 | 1.35 | 1.13 | Both Isoforms |
| TNFAIP8 | 1.43 | 1.13 | Both Isoforms |
| PRSS8 | 0.86 | 1.13 | Both Isoforms |
| RFPL2 | 3.69 | 1.14 | Both Isoforms |
| DNAJB5 | 2.17 | 1.14 | Both Isoforms |
| CCNE2 | 1.19 | 1.14 | Both Isoforms |
| SEC14L5 | 1.95 | 1.14 | Both Isoforms |
| CORO1B | 2.71 | 1.14 | Both Isoforms |
| CKLF-CMTM1 | 1.74 | 1.14 | Both Isoforms |
| IL1R1 | 0.88 | 1.15 | Both Isoforms |
| SLCO3A1 | 1.85 | 1.15 | Both Isoforms |
| HSP90B1 | 1.34 | 1.15 | Both Isoforms |
| GPR75 | 1.14 | 1.15 | Both Isoforms |
| ATP1B1 | 1.25 | 1.15 | Both Isoforms |
| HES4 | 2.41 | 1.15 | Both Isoforms |
| SREBF1 | 1.63 | 1.15 | Both Isoforms |
| MVK | 1.34 | 1.15 | Both Isoforms |
| SEMA3C | 0.97 | 1.15 | Both Isoforms |
| FAM129A | 1.55 | 1.16 | Both Isoforms |
| PTPRM | 1.51 | 1.16 | Both Isoforms |
| TLCD2 | 1.15 | 1.16 | Both Isoforms |
| ELOVL6 | 1.25 | 1.16 | Both Isoforms |
| B3GAT1 | 1.26 | 1.17 | Both Isoforms |
| TMEM104 | 1.36 | 1.17 | Both Isoforms |
| SGK2 | 0.73 | 1.17 | Both Isoforms |
| SAMD12 | 1.61 | 1.17 | Both Isoforms |
| SH3BGRL3 | 1.49 | 1.17 | Both Isoforms |
| NR4A1 | 1.42 | 1.17 | Both Isoforms |
| MSMO1 | 1.51 | 1.17 | Both Isoforms |
| FKBP11 | 1.41 | 1.17 | Both Isoforms |
| SLC35B1 | 1.54 | 1.17 | Both Isoforms |
| PCYT2 | 1.18 | 1.17 | Both Isoforms |

| | | | |
|---|---|---|---|
| GIPC1 | 1.17 | 1.18 | Both Isoforms |
| LRRC46 | 0.83 | 1.18 | Both Isoforms |
| LDLR | 2.05 | 1.18 | Both Isoforms |
| CCDC167 | 1.40 | 1.18 | Both Isoforms |
| KBTBD8 | 1.25 | 1.18 | Both Isoforms |
| GCH1 | 1.35 | 1.18 | Both Isoforms |
| CRACR2B | 1.13 | 1.18 | Both Isoforms |
| GADD45G | 2.91 | 1.19 | Both Isoforms |
| ARRDC1 | 1.27 | 1.19 | Both Isoforms |
| KCNJ11 | 1.12 | 1.19 | Both Isoforms |
| SFXN5 | 2.00 | 1.19 | Both Isoforms |
| PSMC3IP | 1.07 | 1.19 | Both Isoforms |
| C16orf59 | 0.93 | 1.19 | Both Isoforms |
| LRRC59 | 1.87 | 1.20 | Both Isoforms |
| GMPPA | 1.51 | 1.20 | Both Isoforms |
| DHCR24 | 2.13 | 1.20 | Both Isoforms |
| SLC1A4 | 0.97 | 1.20 | Both Isoforms |
| SEC61G | 1.56 | 1.20 | Both Isoforms |
| PCNA | 0.98 | 1.20 | Both Isoforms |
| PREB | 1.52 | 1.20 | Both Isoforms |
| EGFR | 0.96 | 1.20 | Both Isoforms |
| CCDC57 | 0.84 | 1.20 | Both Isoforms |
| SERINC2 | 3.32 | 1.21 | Both Isoforms |
| ZWINT | 0.69 | 1.21 | Both Isoforms |
| KLHL2 | 1.34 | 1.21 | Both Isoforms |
| TMEM110 | 1.43 | 1.21 | Both Isoforms |
| SMPD1 | 1.44 | 1.21 | Both Isoforms |
| DTX2 | 1.09 | 1.21 | Both Isoforms |
| ISG15 | 1.92 | 1.22 | Both Isoforms |
| P4HB | 1.70 | 1.22 | Both Isoforms |
| GUCA2A | 2.10 | 1.23 | Both Isoforms |
| CDK2AP2 | 1.48 | 1.23 | Both Isoforms |
| BMPR1B | 1.51 | 1.23 | Both Isoforms |
| UBTD1 | 1.47 | 1.23 | Both Isoforms |
| LACTB | 1.11 | 1.23 | Both Isoforms |
| PEA15 | 1.28 | 1.23 | Both Isoforms |
| ATG4A | 1.62 | 1.23 | Both Isoforms |
| COQ10B | 1.81 | 1.24 | Both Isoforms |
| UNC5C | 1.00 | 1.24 | Both Isoforms |
| MCM5 | 1.07 | 1.24 | Both Isoforms |
| SLC25A33 | 0.98 | 1.24 | Both Isoforms |
| LRRC8A | 0.77 | 1.24 | Both Isoforms |
| TUBB4B | 1.50 | 1.24 | Both Isoforms |
| CLEC16A | 0.72 | 1.24 | Both Isoforms |
| PICK1 | 1.70 | 1.25 | Both Isoforms |
| AHCYL2 | 1.18 | 1.25 | Both Isoforms |

| | | | |
|---|---|---|---|
| CETN2 | 0.72 | 1.25 | Both Isoforms |
| ABHD3 | 1.67 | 1.25 | Both Isoforms |
| FAM58A | 0.76 | 1.26 | Both Isoforms |
| KLF15 | 0.65 | 1.26 | Both Isoforms |
| FKBP1B | 2.25 | 1.26 | Both Isoforms |
| PPARGC1A | 0.67 | 1.26 | Both Isoforms |
| DONSON | 0.96 | 1.26 | Both Isoforms |
| ALDOC | 0.94 | 1.27 | Both Isoforms |
| HSPA5 | 1.87 | 1.27 | Both Isoforms |
| POLA2 | 0.98 | 1.27 | Both Isoforms |
| SLC45A3 | 2.28 | 1.27 | Both Isoforms |
| AGR2 | 3.34 | 1.27 | Both Isoforms |
| PEBP4 | 2.18 | 1.27 | Both Isoforms |
| CREB3L2 | 1.49 | 1.27 | Both Isoforms |
| SPCS3 | 1.60 | 1.28 | Both Isoforms |
| DNAJB11 | 1.76 | 1.28 | Both Isoforms |
| SSR3 | 1.48 | 1.28 | Both Isoforms |
| USP5 | 0.60 | 1.28 | Both Isoforms |
| MVD | 1.47 | 1.28 | Both Isoforms |
| TBX15 | 4.18 | 1.28 | Both Isoforms |
| SEC61B | 1.67 | 1.28 | Both Isoforms |
| ABRACL | 1.07 | 1.29 | Both Isoforms |
| DDIAS | 1.74 | 1.29 | Both Isoforms |
| HIPK2 | 1.93 | 1.29 | Both Isoforms |
| RAB26 | 1.41 | 1.29 | Both Isoforms |
| PLAUR | 1.72 | 1.29 | Both Isoforms |
| SDK1 | 1.24 | 1.30 | Both Isoforms |
| RPN1 | 1.32 | 1.30 | Both Isoforms |
| 2-Mar | 2.22 | 1.30 | Both Isoforms |
| SLC5A4 | 3.24 | 1.30 | Both Isoforms |
| SEC61A1 | 1.47 | 1.31 | Both Isoforms |
| ST6GALNAC1 | 1.22 | 1.31 | Both Isoforms |
| ARPC1B | 2.16 | 1.31 | Both Isoforms |
| IVD | 0.61 | 1.31 | Both Isoforms |
| ADAM9 | 0.89 | 1.31 | Both Isoforms |
| HMGXB3 | 1.67 | 1.31 | Both Isoforms |
| POP1 | 0.92 | 1.31 | Both Isoforms |
| MAN2B2 | 1.28 | 1.31 | Both Isoforms |
| SLC22A18 | 1.66 | 1.32 | Both Isoforms |
| NNAT | 0.76 | 1.32 | Both Isoforms |
| CRIP2 | 1.51 | 1.32 | Both Isoforms |
| ABCB8 | 1.07 | 1.32 | Both Isoforms |
| SLCO4A1 | 0.77 | 1.33 | Both Isoforms |
| DNAJC3 | 2.01 | 1.33 | Both Isoforms |
| MSRB1 | 0.98 | 1.33 | Both Isoforms |
| ASRGL1 | 2.30 | 1.33 | Both Isoforms |

| | | | |
|---|---|---|---|
| HIST2H2BE | 1.16 | 1.34 | Both Isoforms |
| FANCA | 0.93 | 1.34 | Both Isoforms |
| RAPGEF3 | 2.40 | 1.34 | Both Isoforms |
| DGKA | 1.53 | 1.34 | Both Isoforms |
| CPE | 0.70 | 1.34 | Both Isoforms |
| HOMER2 | 1.55 | 1.34 | Both Isoforms |
| PDIA4 | 1.98 | 1.35 | Both Isoforms |
| GCHFR | 1.61 | 1.35 | Both Isoforms |
| GAL | 0.66 | 1.35 | Both Isoforms |
| DNAJB1 | 1.02 | 1.35 | Both Isoforms |
| WDFY2 | 0.87 | 1.35 | Both Isoforms |
| SLFN13 | 1.00 | 1.35 | Both Isoforms |
| ACADL | 0.97 | 1.36 | Both Isoforms |
| BRCA1 | 0.61 | 1.36 | Both Isoforms |
| SNAI2 | 1.82 | 1.36 | Both Isoforms |
| ACAT2 | 1.40 | 1.36 | Both Isoforms |
| NPIPA5 | 2.55 | 1.37 | Both Isoforms |
| GCC2 | 1.27 | 1.37 | Both Isoforms |
| KNSTRN | 0.95 | 1.37 | Both Isoforms |
| SLC22A3 | 1.00 | 1.37 | Both Isoforms |
| SHCBP1 | 1.37 | 1.37 | Both Isoforms |
| KCNAB1 | 0.92 | 1.37 | Both Isoforms |
| ZBTB7B | 1.68 | 1.37 | Both Isoforms |
| C15orf52 | 1.35 | 1.38 | Both Isoforms |
| ARSA | 1.97 | 1.38 | Both Isoforms |
| LAT2 | 2.29 | 1.38 | Both Isoforms |
| SLC22A31 | 0.93 | 1.38 | Both Isoforms |
| CST3 | 1.22 | 1.38 | Both Isoforms |
| NCAPD3 | 4.14 | 1.38 | Both Isoforms |
| KITLG | 2.04 | 1.39 | Both Isoforms |
| TMEM38A | 0.63 | 1.40 | Both Isoforms |
| GINS3 | 1.10 | 1.40 | Both Isoforms |
| CROT | 0.68 | 1.40 | Both Isoforms |
| SOCS2 | 1.93 | 1.41 | Both Isoforms |
| E2F8 | 0.78 | 1.41 | Both Isoforms |
| CST2 | 3.10 | 1.41 | Both Isoforms |
| CCDC102B | 0.64 | 1.41 | Both Isoforms |
| DRD2 | 1.47 | 1.41 | Both Isoforms |
| GINS4 | 1.01 | 1.41 | Both Isoforms |
| ORC6 | 1.24 | 1.42 | Both Isoforms |
| TRIM62 | 1.82 | 1.42 | Both Isoforms |
| SEC11C | 2.49 | 1.42 | Both Isoforms |
| SOX9 | 1.47 | 1.42 | Both Isoforms |
| AFMID | 0.73 | 1.42 | Both Isoforms |
| SLC4A7 | 1.17 | 1.42 | Both Isoforms |
| FAM13C | 2.29 | 1.43 | Both Isoforms |

| | | | |
|---|---|---|---|
| TFEB | 0.97 | 1.43 | Both Isoforms |
| RBM24 | 3.72 | 1.43 | Both Isoforms |
| LONRF1 | 1.07 | 1.43 | Both Isoforms |
| SLC22A17 | 1.12 | 1.43 | Both Isoforms |
| AP1B1 | 1.32 | 1.44 | Both Isoforms |
| PKMYT1 | 1.01 | 1.44 | Both Isoforms |
| SYTL4 | 1.01 | 1.45 | Both Isoforms |
| CCDC163P | 1.10 | 1.45 | Both Isoforms |
| SLC26A6 | 0.88 | 1.45 | Both Isoforms |
| CDC7 | 1.10 | 1.46 | Both Isoforms |
| WDR66 | 2.04 | 1.46 | Both Isoforms |
| GAS2L1 | 1.29 | 1.46 | Both Isoforms |
| ARHGEF37 | 0.73 | 1.46 | Both Isoforms |
| STK39 | 1.54 | 1.47 | Both Isoforms |
| IDI1 | 2.22 | 1.47 | Both Isoforms |
| HGH1 | 1.05 | 1.47 | Both Isoforms |
| WFS1 | 1.47 | 1.47 | Both Isoforms |
| MCM4 | 1.00 | 1.47 | Both Isoforms |
| TUBG1 | 2.40 | 1.47 | Both Isoforms |
| DHCR7 | 1.17 | 1.48 | Both Isoforms |
| EID3 | 0.90 | 1.48 | Both Isoforms |
| CEBPD | 0.68 | 1.48 | Both Isoforms |
| BLM | 0.73 | 1.49 | Both Isoforms |
| MPV17L2 | 1.59 | 1.49 | Both Isoforms |
| PGP | 1.91 | 1.49 | Both Isoforms |
| TSC22D1 | 1.36 | 1.49 | Both Isoforms |
| SPSB2 | 1.40 | 1.50 | Both Isoforms |
| TMEM169 | 1.14 | 1.50 | Both Isoforms |
| CITED2 | 0.59 | 1.50 | Both Isoforms |
| EXO1 | 0.87 | 1.51 | Both Isoforms |
| ASF1B | 0.82 | 1.52 | Both Isoforms |
| C15orf62 | 0.64 | 1.52 | Both Isoforms |
| CDO1 | 0.93 | 1.52 | Both Isoforms |
| HAUS8 | 0.99 | 1.53 | Both Isoforms |
| MAGEH1 | 1.53 | 1.53 | Both Isoforms |
| EMP1 | 2.79 | 1.53 | Both Isoforms |
| CRLF1 | 0.99 | 1.53 | Both Isoforms |
| CBWD1 | 0.79 | 1.53 | Both Isoforms |
| ADGRG2 | 2.16 | 1.54 | Both Isoforms |
| CMTM1 | 1.37 | 1.54 | Both Isoforms |
| PAQR4 | 2.33 | 1.54 | Both Isoforms |
| PIK3R3 | 1.28 | 1.55 | Both Isoforms |
| NDRG1 | 2.30 | 1.55 | Both Isoforms |
| POLE2 | 1.54 | 1.55 | Both Isoforms |
| PLEKHG3 | 0.92 | 1.55 | Both Isoforms |
| B4GALT1 | 1.33 | 1.55 | Both Isoforms |

| | | | |
|---|---|---|---|
| SYNC | 1.83 | 1.55 | Both Isoforms |
| MANF | 2.28 | 1.55 | Both Isoforms |
| PLA2G4F | 2.65 | 1.55 | Both Isoforms |
| TBXAS1 | 1.90 | 1.56 | Both Isoforms |
| EFCAB12 | 3.08 | 1.56 | Both Isoforms |
| SUCO | 0.86 | 1.57 | Both Isoforms |
| PCDH1 | 2.31 | 1.57 | Both Isoforms |
| EAF2 | 2.85 | 1.57 | Both Isoforms |
| SMS | 2.11 | 1.57 | Both Isoforms |
| HIST1H3C | 1.84 | 1.58 | Both Isoforms |
| NNMT | 1.40 | 1.58 | Both Isoforms |
| PRADC1 | 1.79 | 1.58 | Both Isoforms |
| SLF1 | 0.80 | 1.59 | Both Isoforms |
| C1orf122 | 1.48 | 1.60 | Both Isoforms |
| CMC2 | 1.98 | 1.60 | Both Isoforms |
| MAF | 1.70 | 1.60 | Both Isoforms |
| MTFP1 | 1.83 | 1.61 | Both Isoforms |
| USP18 | 1.12 | 1.61 | Both Isoforms |
| SLC6A17 | 0.68 | 1.61 | Both Isoforms |
| HYOU1 | 1.95 | 1.61 | Both Isoforms |
| RAD51 | 1.25 | 1.62 | Both Isoforms |
| RRBP1 | 1.95 | 1.62 | Both Isoforms |
| PRRG4 | 1.27 | 1.62 | Both Isoforms |
| ACSL3 | 3.05 | 1.63 | Both Isoforms |
| HOXA1 | 2.68 | 1.63 | Both Isoforms |
| B3GLCT | 0.92 | 1.65 | Both Isoforms |
| SLC15A2 | 2.75 | 1.65 | Both Isoforms |
| WBSCR17 | 1.71 | 1.67 | Both Isoforms |
| EBP | 2.03 | 1.67 | Both Isoforms |
| WDR76 | 1.50 | 1.68 | Both Isoforms |
| UHRF1 | 1.48 | 1.68 | Both Isoforms |
| MICAL2 | 0.63 | 1.68 | Both Isoforms |
| ROBO1 | 0.77 | 1.70 | Both Isoforms |
| IGFLR1 | 1.34 | 1.70 | Both Isoforms |
| CCDC189 | 0.82 | 1.70 | Both Isoforms |
| INSIG1 | 2.31 | 1.72 | Both Isoforms |
| DCXR | 1.60 | 1.72 | Both Isoforms |
| RELT | 1.65 | 1.73 | Both Isoforms |
| ZBTB16 | 0.83 | 1.74 | Both Isoforms |
| LIPG | 1.70 | 1.74 | Both Isoforms |
| LIFR | 1.63 | 1.75 | Both Isoforms |
| MYO1E | 1.98 | 1.75 | Both Isoforms |
| HMGCS1 | 1.91 | 1.75 | Both Isoforms |
| TBC1D8B | 0.79 | 1.76 | Both Isoforms |
| HARBI1 | 1.61 | 1.76 | Both Isoforms |
| ATP2B4 | 3.35 | 1.76 | Both Isoforms |

| | | | |
|---|---|---|---|
| TUBA3C | 2.26 | 1.78 | Both Isoforms |
| TK1 | 1.18 | 1.78 | Both Isoforms |
| RDH10 | 1.28 | 1.78 | Both Isoforms |
| CNIH2 | 1.29 | 1.78 | Both Isoforms |
| OLFM1 | 1.94 | 1.79 | Both Isoforms |
| KRT8 | 2.21 | 1.80 | Both Isoforms |
| MYBL2 | 1.36 | 1.80 | Both Isoforms |
| MUM1L1 | 1.34 | 1.80 | Both Isoforms |
| RAET1E | 3.13 | 1.80 | Both Isoforms |
| RAD54L | 1.22 | 1.81 | Both Isoforms |
| SMOC2 | 1.89 | 1.81 | Both Isoforms |
| ANK1 | 2.43 | 1.82 | Both Isoforms |
| CDC25A | 1.40 | 1.82 | Both Isoforms |
| KCNH6 | 1.66 | 1.82 | Both Isoforms |
| CENPN | 2.16 | 1.83 | Both Isoforms |
| COX17 | 1.87 | 1.83 | Both Isoforms |
| SLC20A1 | 1.30 | 1.83 | Both Isoforms |
| PDCD4 | 2.05 | 1.84 | Both Isoforms |
| NUDT11 | 0.79 | 1.84 | Both Isoforms |
| DTL | 1.56 | 1.84 | Both Isoforms |
| FAM189A2 | 2.69 | 1.85 | Both Isoforms |
| DERL3 | 1.49 | 1.85 | Both Isoforms |
| STYK1 | 1.42 | 1.87 | Both Isoforms |
| KRT15 | 2.94 | 1.87 | Both Isoforms |
| ITPR3 | 1.59 | 1.88 | Both Isoforms |
| VDR | 0.68 | 1.88 | Both Isoforms |
| GAS7 | 1.56 | 1.88 | Both Isoforms |
| STRIP2 | 1.74 | 1.88 | Both Isoforms |
| DOCK11 | 2.40 | 1.89 | Both Isoforms |
| ATP1A1 | 2.34 | 1.90 | Both Isoforms |
| TRIB2 | 1.72 | 1.91 | Both Isoforms |
| CHRNA2 | 1.57 | 1.91 | Both Isoforms |
| RRM2 | 1.07 | 1.91 | Both Isoforms |
| HS3ST4 | 1.28 | 1.91 | Both Isoforms |
| HPGD | 1.38 | 1.92 | Both Isoforms |
| CYP1B1 | 2.20 | 1.93 | Both Isoforms |
| TMEM204 | 2.17 | 1.94 | Both Isoforms |
| LOX | 1.18 | 1.94 | Both Isoforms |
| NUCB2 | 2.58 | 1.95 | Both Isoforms |
| CDC45 | 1.46 | 1.96 | Both Isoforms |
| C1orf87 | 3.20 | 1.97 | Both Isoforms |
| HACD2 | 1.13 | 1.97 | Both Isoforms |
| AFF3 | 1.26 | 1.97 | Both Isoforms |
| PPP1R3G | 1.34 | 1.98 | Both Isoforms |
| ELOVL5 | 1.62 | 1.99 | Both Isoforms |
| PLEKHD1 | 2.57 | 2.00 | Both Isoforms |

| | | | | |
|---|---|---|---|---|
| | CSRNP1 | 1.25 | 2.00 | Both Isoforms |
| | ANXA13 | 2.83 | 2.01 | Both Isoforms |
| | EPHA5 | 1.72 | 2.01 | Both Isoforms |
| | CPM | 1.62 | 2.01 | Both Isoforms |
| | SMOX | 0.86 | 2.01 | Both Isoforms |
| | FAM111B | 1.39 | 2.02 | Both Isoforms |
| | E2F2 | 1.92 | 2.03 | Both Isoforms |
| | OSGIN1 | 2.74 | 2.04 | Both Isoforms |
| | GMPPB | 2.52 | 2.04 | Both Isoforms |
| | SLCO2A1 | 2.17 | 2.05 | Both Isoforms |
| | JADE1 | 1.05 | 2.05 | Both Isoforms |
| | FEN1 | 1.61 | 2.05 | Both Isoforms |
| | CDC6 | 2.00 | 2.06 | Both Isoforms |
| | B2M | 0.83 | 2.07 | Both Isoforms |
| | MYO1G | 2.82 | 2.08 | Both Isoforms |
| | TESC | 3.25 | 2.08 | Both Isoforms |
| | LAMA1 | 3.46 | 2.09 | Both Isoforms |
| | ZNF367 | 1.71 | 2.09 | Both Isoforms |
| | KRT19 | 1.58 | 2.09 | Both Isoforms |
| | ASMTL | 1.93 | 2.10 | Both Isoforms |
| | CKLF | 2.31 | 2.11 | Both Isoforms |
| | SLC27A3 | 1.45 | 2.11 | Both Isoforms |
| | CDKN1A | 1.73 | 2.12 | Both Isoforms |
| | ACPT | 1.97 | 2.14 | Both Isoforms |
| | GINS2 | 1.12 | 2.14 | Both Isoforms |
| | RIBC2 | 1.19 | 2.15 | Both Isoforms |
| | SLC37A2 | 2.78 | 2.15 | Both Isoforms |
| | PER1 | 1.03 | 2.15 | Both Isoforms |
| | EVA1C | 1.39 | 2.15 | Both Isoforms |
| | ANGPT4 | 2.81 | 2.16 | Both Isoforms |
| | AMACR | 3.35 | 2.16 | Both Isoforms |
| | PFKFB2 | 2.12 | 2.16 | Both Isoforms |
| | C22orf42 | 7.11 | 2.16 | Both Isoforms |
| | SPOCK2 | 1.87 | 2.17 | Both Isoforms |
| | PRR15L | 1.98 | 2.17 | Both Isoforms |
| | WLS | 0.80 | 2.18 | Both Isoforms |
| | CLSPN | 1.84 | 2.18 | Both Isoforms |
| | KLK3 | 3.54 | 2.19 | Both Isoforms |
| | SEC24D | 2.61 | 2.19 | Both Isoforms |
| | ELL2 | 1.11 | 2.20 | Both Isoforms |
| | AZGP1 | 1.84 | 2.20 | Both Isoforms |
| | FAM196B | 1.36 | 2.20 | Both Isoforms |
| | ATAD2 | 1.18 | 2.20 | Both Isoforms |
| | IL18R1 | 1.18 | 2.24 | Both Isoforms |
| | TMEM8A | 3.15 | 2.24 | Both Isoforms |
| | E2F1 | 2.15 | 2.24 | Both Isoforms |

| | | | |
|---|---|---|---|
| MYO16 | 3.87 | 2.26 | Both Isoforms |
| CECR6 | 2.41 | 2.26 | Both Isoforms |
| STARD4 | 3.00 | 2.27 | Both Isoforms |
| FICD | 2.35 | 2.30 | Both Isoforms |
| MCM10 | 1.85 | 2.31 | Both Isoforms |
| TUBA3E | 2.20 | 2.32 | Both Isoforms |
| C9orf66 | 1.45 | 2.33 | Both Isoforms |
| SDF2L1 | 2.99 | 2.33 | Both Isoforms |
| KCNH2 | 0.98 | 2.33 | Both Isoforms |
| PATE2 | 1.41 | 2.37 | Both Isoforms |
| FSTL4 | 0.66 | 2.37 | Both Isoforms |
| LGI3 | 2.11 | 2.42 | Both Isoforms |
| CPEB2 | 1.52 | 2.43 | Both Isoforms |
| CRELD2 | 2.31 | 2.43 | Both Isoforms |
| FADS3 | 1.26 | 2.46 | Both Isoforms |
| CA2 | 1.37 | 2.46 | Both Isoforms |
| GPR153 | 0.64 | 2.46 | Both Isoforms |
| PRPS2 | 0.61 | 2.47 | Both Isoforms |
| RPRM | 1.18 | 2.53 | Both Isoforms |
| IRAK3 | 1.85 | 2.56 | Both Isoforms |
| SPOCK1 | 1.95 | 2.56 | Both Isoforms |
| PHOSPHO1 | 5.08 | 2.58 | Both Isoforms |
| WIPI1 | 3.28 | 2.59 | Both Isoforms |
| TGM2 | 1.71 | 2.60 | Both Isoforms |
| CTNNA2 | 2.36 | 2.60 | Both Isoforms |
| FGFR1 | 1.36 | 2.61 | Both Isoforms |
| DDIT4 | 2.05 | 2.61 | Both Isoforms |
| GNMT | 1.94 | 2.62 | Both Isoforms |
| TMEM47 | 1.75 | 2.63 | Both Isoforms |
| ZNF812 | 3.37 | 2.64 | Both Isoforms |
| NDRG4 | 1.11 | 2.65 | Both Isoforms |
| CKB | 2.29 | 2.65 | Both Isoforms |
| ARMC12 | 1.47 | 2.66 | Both Isoforms |
| FAM107A | 1.88 | 2.67 | Both Isoforms |
| CAPN14 | 1.47 | 2.67 | Both Isoforms |
| TUBA3D | 2.94 | 2.70 | Both Isoforms |
| FA2H | 2.17 | 2.70 | Both Isoforms |
| BHLHA15 | 3.87 | 2.70 | Both Isoforms |
| CYP4F8 | 1.83 | 2.71 | Both Isoforms |
| HSPH1 | 0.71 | 2.72 | Both Isoforms |
| DBI | 3.19 | 2.74 | Both Isoforms |
| THSD1 | 2.99 | 2.77 | Both Isoforms |
| MAP1B | 1.80 | 2.79 | Both Isoforms |
| GLIS3 | 1.24 | 2.81 | Both Isoforms |
| KCNMA1 | 3.05 | 2.81 | Both Isoforms |
| DYNC1I1 | 2.18 | 2.81 | Both Isoforms |

| | | | |
|---|---|---|---|
| CST4 | 4.37 | 2.84 | Both Isoforms |
| HSPA2 | 0.74 | 2.86 | Both Isoforms |
| AKAP12 | 0.93 | 2.91 | Both Isoforms |
| MAP3K6 | 2.43 | 2.93 | Both Isoforms |
| SLC26A2 | 2.61 | 2.95 | Both Isoforms |
| FAM166B | 0.90 | 2.96 | Both Isoforms |
| CLGN | 1.30 | 2.97 | Both Isoforms |
| NSDHL | 1.45 | 3.01 | Both Isoforms |
| SMIM24 | 2.50 | 3.03 | Both Isoforms |
| FADS2 | 2.76 | 3.06 | Both Isoforms |
| ELMO1 | 2.97 | 3.08 | Both Isoforms |
| FASN | 2.27 | 3.15 | Both Isoforms |
| IL31RA | 1.17 | 3.16 | Both Isoforms |
| MFSD2A | 2.26 | 3.21 | Both Isoforms |
| TMEM61 | 1.42 | 3.22 | Both Isoforms |
| RBM20 | 3.35 | 3.25 | Both Isoforms |
| TNFRSF1A | 0.60 | 3.27 | Both Isoforms |
| PDE6A | 1.12 | 3.28 | Both Isoforms |
| TDRD9 | 0.92 | 3.31 | Both Isoforms |
| RHOB | 1.67 | 3.37 | Both Isoforms |
| TIPARP | 1.27 | 3.38 | Both Isoforms |
| CHST2 | 1.59 | 3.46 | Both Isoforms |
| SP6 | 2.48 | 3.51 | Both Isoforms |
| ABCC8 | 0.64 | 3.52 | Both Isoforms |
| NEK10 | 2.12 | 3.56 | Both Isoforms |
| SLC2A3 | 5.06 | 3.59 | Both Isoforms |
| FAM196A | 0.62 | 3.59 | Both Isoforms |
| FADS1 | 2.33 | 3.60 | Both Isoforms |
| HES1 | 2.62 | 3.62 | Both Isoforms |
| HS6ST1 | 1.78 | 3.63 | Both Isoforms |
| RASD1 | 2.17 | 3.64 | Both Isoforms |
| DNER | 0.61 | 3.64 | Both Isoforms |
| SERPINF2 | 1.60 | 3.70 | Both Isoforms |
| NPR3 | 0.83 | 3.71 | Both Isoforms |
| ERRFI1 | 2.80 | 3.73 | Both Isoforms |
| ABHD6 | 1.43 | 3.82 | Both Isoforms |
| CDKN2B | 1.95 | 3.82 | Both Isoforms |
| ITGA11 | 3.81 | 3.85 | Both Isoforms |
| SLC31A2 | 1.46 | 3.85 | Both Isoforms |
| PPP1R16B | 0.78 | 3.86 | Both Isoforms |
| GP2 | 1.70 | 3.90 | Both Isoforms |
| TIMP3 | 1.67 | 3.91 | Both Isoforms |
| TSPYL2 | 1.62 | 3.96 | Both Isoforms |
| MOGAT2 | 2.47 | 3.97 | Both Isoforms |
| ADAMTS15 | 1.57 | 4.03 | Both Isoforms |
| SERPINA4 | 1.17 | 4.03 | Both Isoforms |

| | | | |
|---|---|---|---|
| SNTG2 | 1.45 | 4.16 | Both Isoforms |
| ABCG2 | 2.29 | 4.37 | Both Isoforms |
| NUDT10 | 2.41 | 4.54 | Both Isoforms |
| PLA2G4E | 4.66 | 4.56 | Both Isoforms |
| TMEM37 | 1.42 | 4.57 | Both Isoforms |
| NPR1 | 1.89 | 4.57 | Both Isoforms |
| CYP4F22 | 1.53 | 4.58 | Both Isoforms |
| ANGPT1 | 1.74 | 4.62 | Both Isoforms |
| EFEMP1 | 1.09 | 4.74 | Both Isoforms |
| EDN2 | 0.79 | 4.74 | Both Isoforms |
| TSPAN18 | 1.34 | 4.86 | Both Isoforms |
| SULT1C4 | 2.15 | 4.91 | Both Isoforms |
| FKBP5 | 4.29 | 4.99 | Both Isoforms |
| DISP2 | 3.48 | 5.09 | Both Isoforms |
| PGF | 0.69 | 5.09 | Both Isoforms |
| PGLYRP2 | 3.55 | 5.17 | Both Isoforms |
| FIBCD1 | 0.83 | 5.27 | Both Isoforms |
| PHGR1 | 3.74 | 5.31 | Both Isoforms |
| CRISP1 | 7.03 | 5.43 | Both Isoforms |
| DOCK8 | 3.55 | 5.49 | Both Isoforms |
| CA4 | 4.10 | 5.62 | Both Isoforms |
| IVL | 3.95 | 5.68 | Both Isoforms |
| NEFH | 2.41 | 5.89 | Both Isoforms |
| CD99 | 1.23 | 5.97 | Both Isoforms |
| THBD | 0.86 | 6.09 | Both Isoforms |
| SPINK2 | 2.29 | 6.33 | Both Isoforms |
| SLC38A4 | 4.14 | 6.42 | Both Isoforms |
| CPVL | 2.23 | 7.05 | Both Isoforms |
| SLC1A3 | 3.05 | 7.08 | Both Isoforms |
| ORM2 | 4.89 | 7.27 | Both Isoforms |
| TLR7 | 4.52 | 7.32 | Both Isoforms |
| ANGPTL5 | 3.58 | 7.35 | Both Isoforms |
| PGC | 8.31 | 7.42 | Both Isoforms |
| ATP1A2 | 6.12 | 7.46 | Both Isoforms |
| DPEP1 | 3.06 | 7.52 | Both Isoforms |
| SLC26A3 | 4.96 | 7.55 | Both Isoforms |
| SPINK13 | 2.22 | 7.72 | Both Isoforms |
| SERPINA3 | 2.23 | 7.74 | Both Isoforms |
| MORC1 | 4.52 | 7.76 | Both Isoforms |
| KRTAP9-6 | 4.52 | 7.86 | Both Isoforms |
| PNLIP | 5.27 | 8.05 | Both Isoforms |
| EBI3 | 2.22 | 8.06 | Both Isoforms |
| GPR6 | 3.06 | 8.53 | Both Isoforms |
| ORM1 | 4.74 | 9.00 | Both Isoforms |
| SLC2A14 | 3.05 | 9.09 | Both Isoforms |
| C7orf57 | 2.22 | 10.14 | Both Isoforms |

| | | | | |
|---|---|---|---|---|
| | PLP1 | 7.12 | 10.41 | Both Isoforms |
